# Evolution of vascular plants through redeployment of ancient developmental regulators

**DOI:** 10.1101/650358

**Authors:** Nicole van ’t Wout Hofland, Kuan-Ju Lu, Eliana Mor, Sumanth Mutte, Paul Abrahams, Hirotaka Kato, Klaas Vandepoele, Dolf Weijers, Bert De Rybel

**Author notes:** equal contribution (Nicole van ’t Wout Hofland, Kuan-Ju Lu).

## Abstract

Vascular plants provide most of the biomass, food and feed on earth; yet the molecular innovations that led to the evolution of their conductive tissues are unknown. Here, we reveal the evolutionary trajectory for the heterodimeric TMO5/LHW transcription factor complex, which is rate-limiting for vascular cell proliferation in *Arabidopsis thaliana*. Both regulators have origins predating vascular tissue emergence, and even terrestrialization. We further show that TMO5 evolved its modern function, including dimerization with LHW, at the origin of land plants. A second innovation in LHW, coinciding with vascular plant emergence, conditioned obligate heterodimerization and generated the critical function in vascular development. In summary, our results suggest that division potential of vascular cells may have been a major driver in the evolution of vascular plants.

## Introduction

Recent estimates suggest that plants contribute about 80% of all biomass on earth^1^. Most is formed by vascular plants (Tracheophytes), that can live for centuries and grow to large sizes. While non-vascular plants mostly rely on cell-to-cell symplastic transport, limiting movement of water and solutes to just a few cells^2^, Tracheophytes developed specialized vascular tissues with xylem and phloem cell types to facilitate transport over long distances^3^. The acquisition of conducting tissues can therefore be regarded as a major evolutionary innovation. Although vascular plants are defined by the presence of vascular tissue with characteristic secondary cell wall thickenings ensuring both structural support and transport over larger distances^4–7^, earlier diverging land plants in the Bryophyte group (mosses, liverworts and hornworts) also contain water-conducting cells (WCC) and food-conducting cells (FCC), analogous to xylem and phloem cells^6, 8^. Recent work in the moss *Physcomitrella patens* revealed that the differentiation of WCC and FCC hinges on NAC transcription factors orthologous to those that trigger xylem differentiation in the Tracheophyte *Arabidopsis thaliana*^9^. Thus, the transcriptional networks leading to a functional conducting tissue with secondary cell wall depositions are largely conserved. Given that the auxin hormone response pathway, that has been shown to trigger vascular tissue formation in Tracheophytes^10, 11^, is also conserved in Bryophytes^12–16^, a key question is what genetic innovation facilitated the establishment of elaborate vascular tissues in Tracheophytes. A fundamental difference in the conducting tissues between Bryophytes and Tracheophytes is its dimensions. The core genetic network driving vascular cell division leading to increase in girth in Arabidopsis involves a dimer of the TARGET OF MONOPTEROS 5 (TMO5) and LONESOME HIGHWAY (LHW) basic Helix-Loop-Helix (bHLH) transcription factors (TFs)^17–23^; representing a rate-limiting component in vascular cell proliferation. In this study, we show that proliferation of vascular cells may have been key to evolve elaborate vascular tissues.

## Results and Discussion

### TMO5/LHW acts as an obligate heterodimer in Arabidopsis vascular development

In Arabidopsis, both TMO5 and LHW have several paralogues, and higher-order mutants in either subclade strongly reduce the number of vascular cell files, leading to nearly indistinguishable phenotypes^17, 19–23^. Simultaneous misexpression of TMO5 and LHW triggers periclinal and radial cell divisions (PRD) in all root meristem cells^19, 20, 22–24^. Thus, TMO5 and LHW are functionally interdependent. Within the root meristem, overlapping *TMO5* and *LHW* expression patterns limits activity of the TMO5/LHW dimer: *TMO5* and homologs are expressed in young xylem cells, while *LHW* and homologs are more broadly expressed (**Fig. 1a**; Refs. 19, 22). Given that all LHW-clade genes are broader in expression domain than TMO5-clade genes, it is conceivable that LHW-clade genes also act independently from TMO5. However, the strong *lhw lhw-like1* (*lhw ll1*) double mutant phenotype was almost fully rescued in overall growth (**Fig. 1b**) and root vascular development (**Fig. 1c,d**) by expressing *LHW* only in *TMO5*-expressing cells (p*TMO5*::LHW in *lhw ll1*). Since p*TMO5*::LHW-GFP fluorescence was retained within the xylem cells (**Fig. 1e,f**), LHW protein does not move, which therefore suggests that LHW has no vascular function outside of the TMO5 domain, and establishes that TMO5/LHW acts as an obligate heterodimer in Arabidopsis. Given that TMO5 and LHW are phylogenetically distantly related bHLH proteins, this result urges the question of when and how the strict dependence on such an obligate heterodimer could have evolved.

**Fig. 1:**
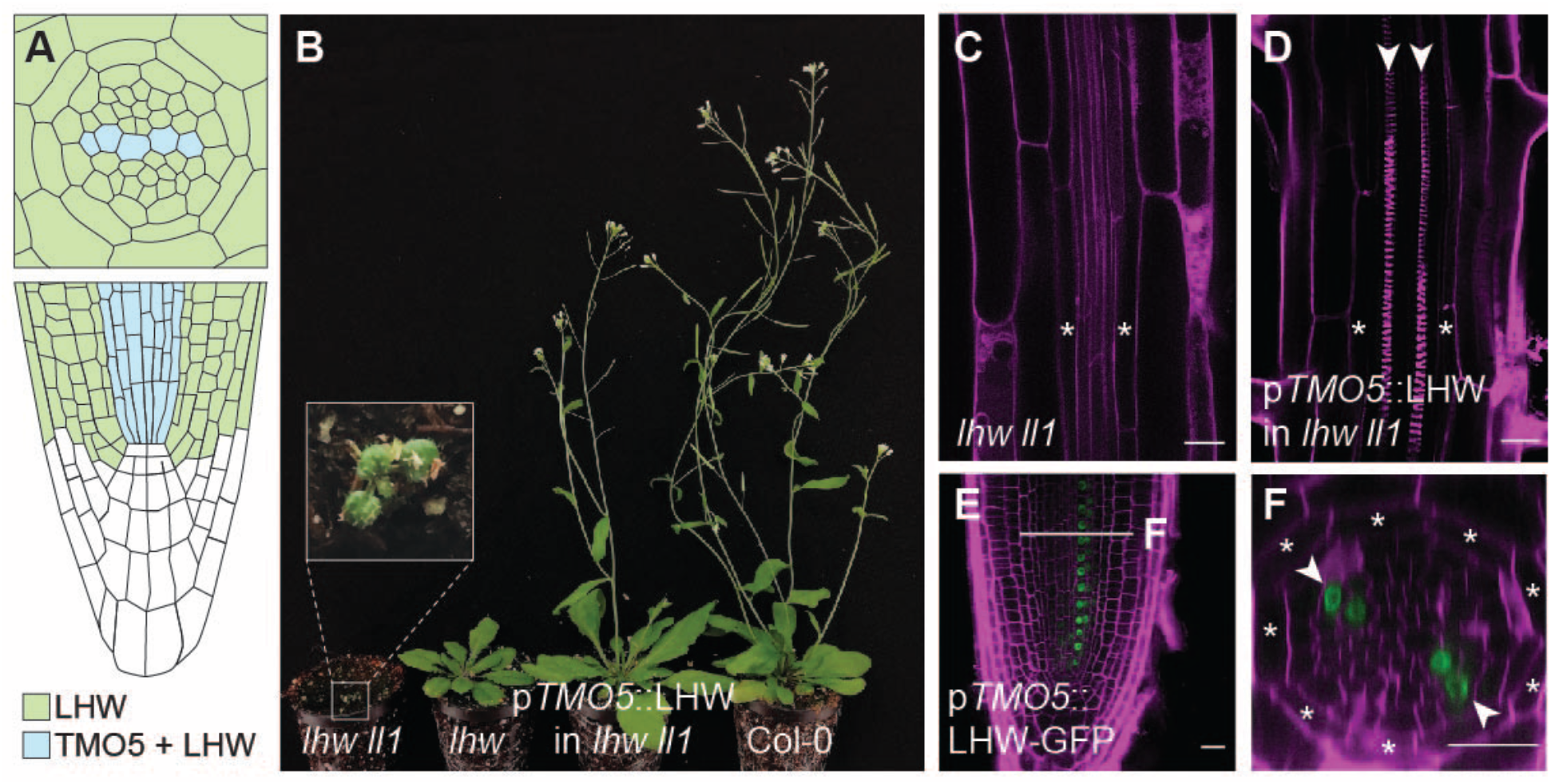
TMO5 and LHW act as an obligate heterodimer in Arabidopsis. (A) Schematic representation of the *TMO5* and *LHW* expression domains in the Arabidopsis primary root (transverse cross-section in upper and longitudinal section in lower panel). Within the LHW-domain (green), a subset of cells expresses both LHW and TMO5 (blue). (B-D) Complementation of the *lhw ll1* double mutant phenotype by a p*TMO5*::LHW transgene, as shown by flowering plant stature (B) and increased vascular tissue size and diarch pattern (D; surrounding endodermis layer marked by asterisks; xylem poles marked by arrowheads), compared to the *lhw ll1* mutant (B, C). (E, F) Accumulation of LHW-GFP protein in the xylem domain of p*TMO5*::LHW-GFP root in longitudinal (E) and transverse (F) cross-section. Asterisks mark surrounding endodermis and arrowheads mark xylem axis. Scale bars represent 20 µm.

### Deep evolutionary origin of TMO5 and LHW genes

To explore whether the cell division function triggered by TMO5/LHW may have contributed to the evolution of vascular tissues, we applied a phylogenomic strategy for identifying orthologs in the green lineage. We used full-length TMO5 and LHW Arabidopsis protein sequences as well as their homologs, TMO5-LIKE1 to 4 (T5L1 to T5L4) and LHW-LIKE1 to 3 (LL1 to LL3), to search against transcriptome assemblies and annotated genome sequences of a representative group of 58 plants ranging from Chlorophytes to Angiosperms (see details in Supplementary Material; **Table S3**). TMO5 and LHW orthologs were found to be present in all major plant lineages even including the Charophytes, algal sisters to land plants (**Fig. 2a,b**; for individual gene identifiers and bootstrap values, see **Fig. S1, S2**). We further observed that most Charophytes and Bryophytes contain only a single copy of each gene, whereas numbers increased in the Tracheophyte lineage (**Fig. 2k**). As bHLH family expansion occurred after Charophytes split from the Embryophytes^25^, this suggests that TMO5 and LHW orthologs were among the first bHLH proteins established in the plant lineage. Thus, *TMO5* and *LHW* genes did not appear with the establishment of vascular tissues during land plant evolution, but have a more ancient origin.

**Fig. 2:**
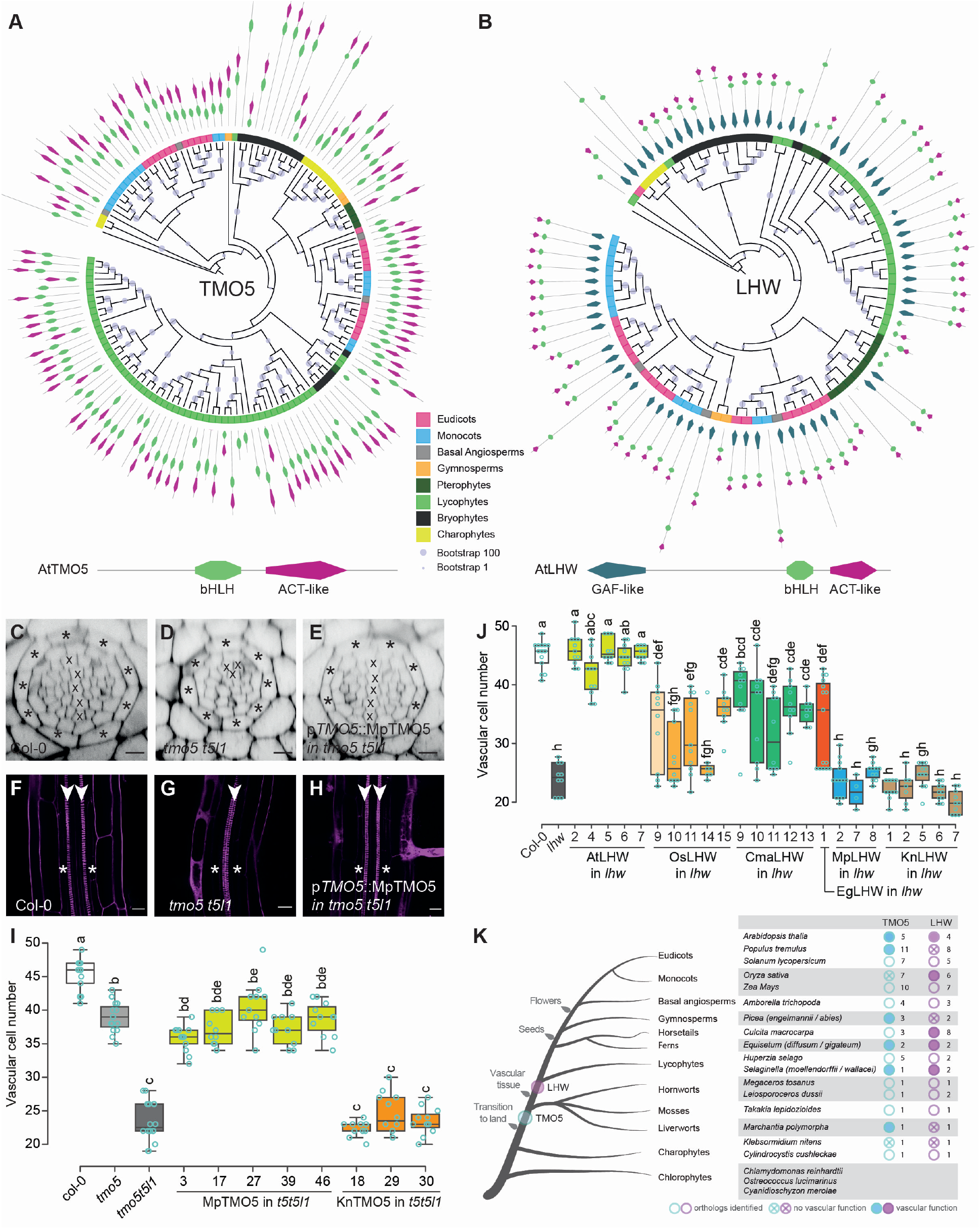
Acquisition of a modern function by ancient TMO5 and LHW proteins. (A and B) Schematic BIONJ phylogenetic trees and domain analyses of TMO5 orthologs (A) and LHW orthologs (B). Major plant clades are represented by different colors. Note that both TMO5 and LHW are present in Charophytes as well as in all land plant groups. In addition, all TMO5 orthologs contain both bHLH and ACT-like domains (lower panel); LHW orthologs have three conserved domains, with orthologs in charophytes, liverworts and mosses which lack the ACT-like domain. Blue circles in the trees indicate bootstrap values. (C-J) Inter-species complementation analysis of TMO5 (C-I) and LHW (J) orthologs. (C-H) Representative cross-section (C-E) and longitudinal sections (F-H) of a wild-type (C, F; Col-0), *tmo5 t5l1* mutant (D, G), and a line carrying the p*TMO5*::MpTMO5 transgene in the *tmo5 t5l1* background (E, H). Cell file numbers in the vascular tissue (encircled by the endodermis layer; asterisks) were quantified in cross-sections and represented in box plots for TMO5 (I) and LHW (J) orthologs. Small case letters in (I-J) indicate significantly different groups as determined using a one-way ANOVA with post-hoc Tukey HSD testing. Blue circles represent all data-points. Presence of a diarch vascular architecture was used as another sign of complementation. (K) Summary of the presence and numbers of TMO5 and LHW orthologs in the plant kingdom. Orthologs tested in complementation assays are marked by solid circle if the gene complemented, and by a crossed circle if no complementation was observed. Crosses in (C-E) mark xylem cell files; arrowheads (F-H) indicate xylem poles. Scale bars in (C-H) represent 20 µm.

### TMO5 function evolved early and is extremely conserved

Although TMO5 and LHW genes emerged early during plant evolution, it is plausible that the capacity to regulate vascular development is a derived property that evolved later. We therefore used a cross-species assay to investigate which TMO5 orthologs have the capacity to complement the Arabidopsis *tmo5 t5l1* double mutant (henceforth referred to as the ‘vascular function’). Due to reduced PRD, this mutant shows a strongly reduced vascular bundle with only one protoxylem pole (monarch vascular pattern) (**Fig. 2d,g**) instead of two protoxylem poles (diarch vascular pattern) in wild-type (**Fig. 2c,f**). We expressed the TMO5 orthologs (**Table S1** and **Table S3** for nomenclature) from the native At*TMO5* promoter (**Fig. 2e,h-j; Fig. S3, S4**). With the exception of OsTMO5, all Tracheophytic TMO5 orthologs (AtTMO5, PtTMO5, PaTMO5, EdTMO5 and SmTMO5) complemented the mutant phenotype (**Fig. S3**), revealing that the “vascular function” of this protein is conserved among vascular plants. Surprisingly however, the Bryophytic MpTMO5 was likewise able to restore vascular development (**Fig. 2e,h-j; Fig. S3, S4; Table S1**), suggesting a deep functional conservation of TMO5 extending beyond vascular plants to the Bryophytes. In contrast, the Charophytic KnTMO5 could not complement (**Fig. 2i; Fig. S3, S4; Table S1**). These data indicate that all land plant TMO5 orthologs tested are capable of performing the vascular function (**Fig. 2k**), and suggest that the protein function has remained extremely conserved despite hundreds of millions of years of evolution.

### Establishment of LHW vascular function correlates with emergence of vascular plants

The extreme conservation in TMO5 function suggests that changes in TMO5 protein function are not connected to the appearance of vascular tissues. We next dissected the emergence of a vascular function for LHW orthologs in a similar complementation study using the *lhw* mutant and a broad range of LHW orthologs (**Fig. 2j; Fig. S3, S4; Table S1**). Although only the AtLHW positive control showed a complete rescue of vascular bundle size (**Fig. 2j; Fig. S3, S4; Table S1**); the vascular orthologs OsLHW, CmaLHW and EgLHW could partly restore vascular cell number. Furthermore, restoration of the diarch pattern was observed in 9% of the SwLHW lines (**Table S1**). In contrast, non-vascular orthologs MpLHW and KnLHW did not show any complementation (**Fig. 2j; Fig. S3, S4; Table S1**), showing that LHW ortholog function is less conserved than TMO5 function. Strikingly, the ability to mediate vascular cell proliferation is limited to Lycophyte and Tracheophyte lineages (**Fig. 2k**), and thus coincides with the emergence of vascular plants. This raises the intriguing possibility that the establishment of a functional TMO5/LHW heterodimer contributed to the emergence of elaborate vascular tissues.

### TMO5/LHW heterodimerization is an ancient property

We next asked if the vascular function of TMO5 and LHW orthologues correlates with their ability to form heterodimers. We first examined the interaction of AtLHW with TMO5 orthologs with a vascular function (AtTMO5 and MpTMO5) as well as KnTMO5, without vascular function. Surprisingly, all TMO5 orthologs interacted with AtLHW (**Fig. 3a,c**), suggesting that the functional differences between KnTMO5 and MpTMO5 are not due to differential capacity to interact with AtLHW. To investigate if heterodimerization capacity is conserved among the natural TMO5/LHW pairs, we next tested interactions within the same species. Similar to the AtTMO5 and AtLHW, Bryophytic MpTMO5 and MpLHW can heterodimerize, but this was not the case for the Charophytic KnTMO5 and KnLHW (**Fig. 3b,c**). Given that bHLH proteins require homotypic interactions with another bHLH protein (either homo- or heterodimer) to bind DNA (*26, 27*), it is possible that KnTMO5/KnLHW proteins form homodimers instead. We therefore tested homo-dimerization potential of all TMO5 and LHW orthologues and found that indeed, all TMO5 orthologs formed homodimers (**Fig. 3b,c**). In contrast, none of the LHW orthologs homodimerized (**Fig. 3b**), suggesting that at least in Klebsormidium, LHW may have another bHLH interaction partner. In summary, these interaction studies indicate that the vascular function is not correlated to the capacity of forming TMO5/LHW heterodimers, as this ability already exists in pre-vascular Bryophytes. As such, it is possible that in Marchantia, MpTMO5 and MpLHW regulate another developmental process as heterodimers, and were later recruited for vascular development.

**Fig. 3:**
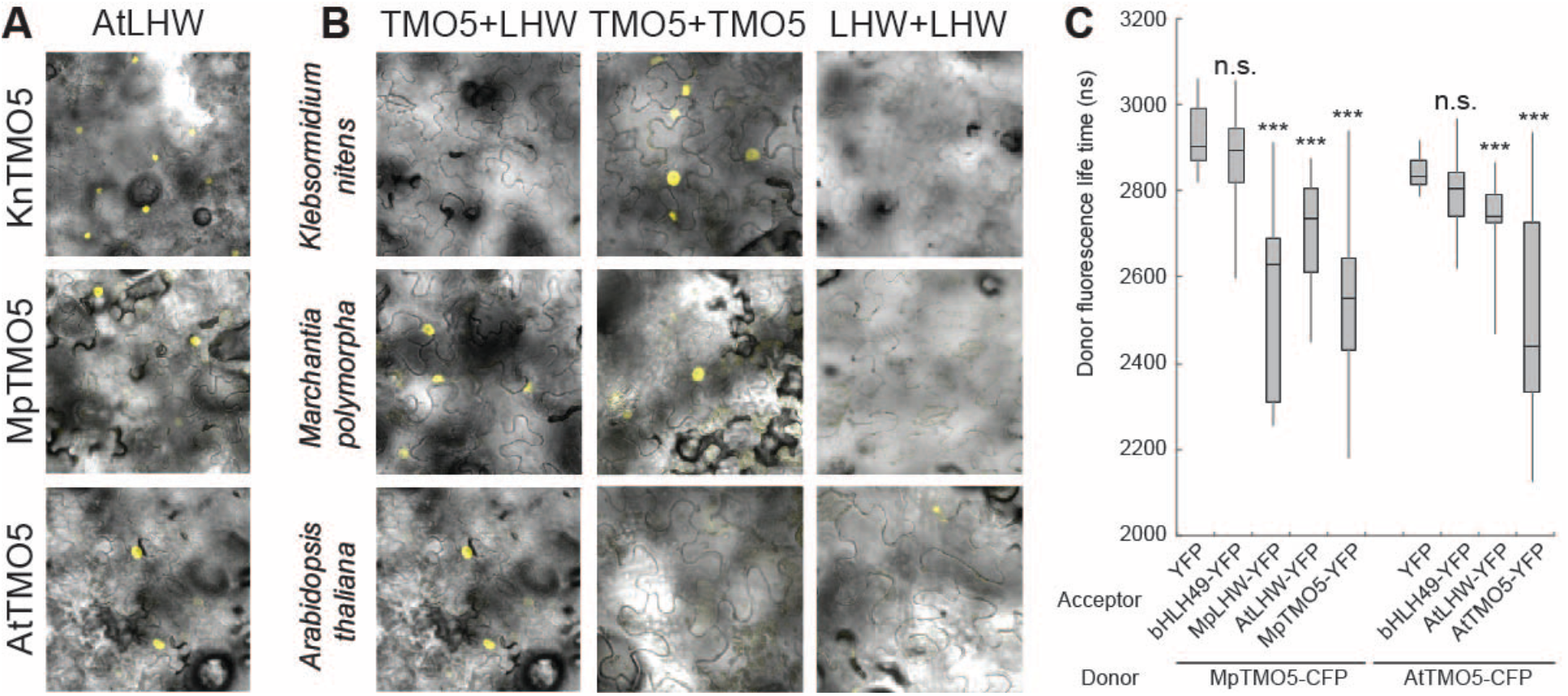
Homo- and heterodimerization among TMO5 and LHW orthologues. (A) Bimolecular Fluorescence Complementation (BiFC) analyses of heterodimerization of Klebsormidium (KnTMO5), Marchantia (MpTMO5) and Arabidopsis (AtTMO5) TMO5 orthologues with Arabidopsis LHW (AtLHW), tested in tobacco leaves. Yellow signal in nuclei indicate protein interaction. (B) Within-species homo- and heterodimerization of TMO5 and LHW orthologues. In each row, protein pairs from each species indicated on the left were tested in BiFC in all possible combination indicated above each row. (C) Förster Resonance Energy Transfer - Fluorescence Lifetime IMaging (FRET-FLIM) analysis of protein interactions among Marchantia (Mp) and Arabidopsis (At) TMO5 and LHW orthologues. Data are represented as mean ± SD (n≥15). YFP and the unrelated bHLH49 were used as negative controls. P-values were calculated by two-tail student t-tests; n.s.: not significant; ***: p-value < 0.01).

### A facultative interaction predated obligate heterodimerization

To understand the relation between TMO5 and LHW in a non-vascular plant, we turned to genetic analysis in *Marchantia polymorpha*, a liverwort with single TMO5 and LHW orthologs and an exquisite set of genetic and genomic tools^28–30^. We first analysed p*MpTMO5*::nCitrine and p*MpLHW*::nCitrine expression domains and found both to be ubiquitously expressed in gemmalings; with p*MpLHW*::nCitrine showing higher expression levels and stronger fluorescence in meristematic notches (**Fig. 4a,b**). Despite minor differences, MpTMO5 and MpLHW are thus co-expressed in most of their expression domains, consistent with heterodimerization potential. To explore *MpTMO5* and *MpLHW* function, we generated three independent CRISPR/Cas9 mutants for each gene (**Fig. S5**). Although general growth and horizontal expansion of thalli were reduced in both the *Mptmo5* and *Mplhw* mutants (**Fig. 4c-e; Fig. S5**), at the cellular level, no obvious defects were observed in young gemmalings (**Fig. 4f-h**). In summary, the mutant phenotypes show that both MpTMO5 and MpLHW are involved, but not critically required for normal development. This is in sharp contrast to Arabidopsis, where loss-of-function mutants are lethal^19^. Furthermore, while there is some phenotypic resemblance, it is unclear if MpTMO5 and MpLHW functions are interdependent.

**Fig. 4:**
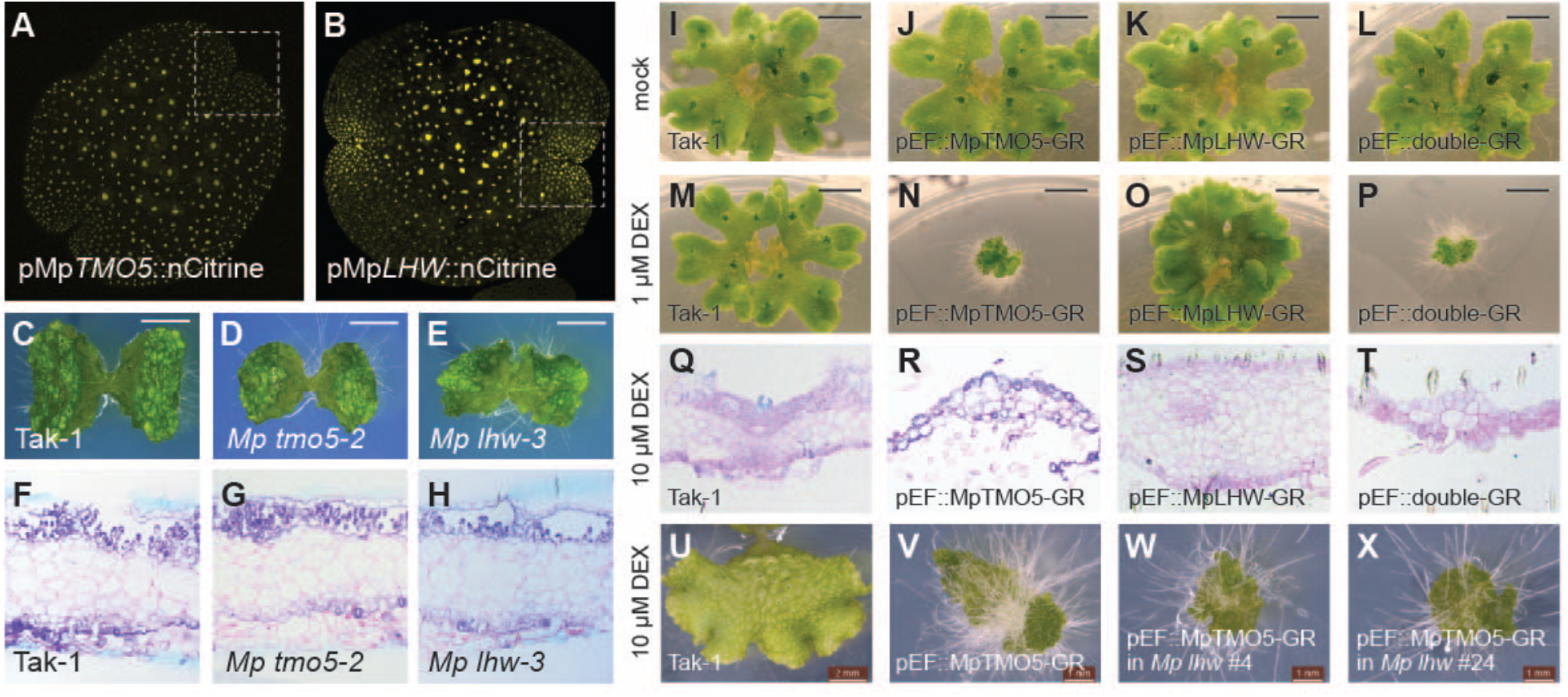
Marchantia TMO5 and LHW independently control gemmae and thallus development. (A, B) Expression of p*MpTMO5*::nCitrine (A) and p*MpLHW*::nCitrine (B) in gemmalings. Note that p*MpLHW* expression is ubiquitous in apical notches (marked by dashed box in B), while p*MpTMO5* expression is weak from the apical notch (dashed box in A). (C-H) Overall appearance (C-E) and histochemical cross-sections (F-H) of Tak-1 wild-type (C, F) *Mp tmo5-2* mutant (D, G) and *Mp lhw-3* mutant (E, H) gemmalings. (I-T) Overall appearance (I-P) and histochemical cross-sections (Q-T) of Tak-1 wild-type (I, M, Q), p*EF*::MpTMO5-GR (J, N, R), p*EF*::MpLHW-GR (K, O, S) and p*EF*::double-GR (L, P, T) thallus that were grown on mock medium (I-L), on 1 μM DEX (M-P) or on 10 μM DEX (Q-T). Cross-sections (Q-T) were made near the meristematic notches. (U-X) Overall appearance of Tak-1 wild-type (U), p*EF*::TMO5-GR in Tak-1 background (V) and two independent *Mp lhw* mutants in the same p*EF*::TMO5-GR background (W, X), all grown on 10 μM DEX for 14 days. Scale bars in (C-E, U) represent 2 mm and in (V-X) 1 mm.

Only simultaneous overexpression of AtTMO5 and AtLHW results in massive cell proliferation in Arabidopsis, while single misexpression of each gene induces very mild proliferation^20, 22–24^. Thus, to test functional interdependence, we next generated dexamethasone (DEX)-inducible single and double over-expression lines by driving MpTMO5-GR and MpLHW-GR from the strong p*EF* promoter^31^. In both single overexpression lines, we observed strong morphological changes upon DEX treatment, resulting in distinctive phenotypes that differed between MpTMO5-GR and MpLHW-GR lines (**Fig. 4i-k, m-o**). While induction of p*EF*::MpTMO5-GR resulted in dwarfed plants with increased rhizoid formation, p*EF*::MpLHW-GR induction caused a more compact thallus structure and absence of gemmae cups (**Fig. 4i-k, m-o**). Interestingly, combined overexpression did not result in additional phenotypes compared to the single misexpression lines (**Fig. 4l,p**). Closer examination of tissue anatomy in the misexpression thalli revealed a strongly reduced number of cells in the p*EF*::MpTMO5-GR; while p*EF*::MpLHW-GR induction caused a thickened thallus and cell proliferation (**Fig. 4q-t**). Hence MpTMO5 and MpLHW seem to induce distinct cellular phenotypes, consistent with a model in which they control development independently. To test this hypothesis directly, we generated CRISPR/Cas9 *Mplhw* loss-of-function mutations in the p*EF*::MpTMO5-GR background and asked if TMO5 misexpression phenotypes required LHW function. We observed the same phenotypic changes upon DEX treatment when the p*EF*::MpTMO5-GR construct was in a control Tak-1 or in a *Mp lhw* mutant background (**Fig. 4u-x** compared to **Fig. 4m-p; Fig. S6**), confirming that MpTMO5 does not need MpLHW to exert its function. These results hence support a model in which MpTMO5 and MpLHW function independently in *Marchantia polymorpha*, in contrast to the obligate heterodimer complex found in *Arabidopsis thaliana*.

To further support this hypothesis, we next determined if MpTMO5 and MpLHW regulate different sets of target genes by analysing genome-wide transcriptional changes upon single or double overexpression in *Marchantia*. First, we generated a comparable dataset in Arabidopsis roots by similarly inducing either only AtTMO5, only AtLHW, or both together, followed by RNA-seq analysis. As expected based on earlier transcriptome analysis^22, 23, 32^, AtTMO5 and AtLHW were clearly found to work cooperatively, as individual induction of AtTMO5 or AtLHW had a very limited transcriptional response (**Fig. 5** and **Table S4**), while simultaneous induction of both proteins caused a large transcriptional response with over 800 genes differentially induced, including the known target genes *LOG4* and *AT4G38650* (blue bars in **Fig. 5** and **Table S4**). We next performed the identical treatments on *Marchantia* gemmalings, followed by RNA-seq analysis. Confirming phenotypic data, and in stark contrast to the Arabidopsis data, single MpTMO5 or MpLHW induction resulted in respectively 108 and 248 differentially upregulated genes, of which most are only induced by one of the transcription factors (blue bars in **Fig. 5** and **Table S4**). Combined overexpression of MpTMO5 and MpLHW induced transcript levels of almost 300 additional genes (blue bars in **Fig. 5** and **Table S4**). When analysing both up- and downregulated genes together, a similar trend was observed (**Fig. S7** and **Table S4**). Thus, while each protein can act as a transcription factor on its own, regulating different sets of genes, there is cooperative regulation of a set of additional genes. Importantly, we did not recover the orthologs of the known Arabidopsis TMO5/LHW target genes in *Marchantia* and there was only a small overlap in induced genes from both species (**Table S4**). This suggests that MpTMO5 and MpLHW do not share transcriptional targets with Arabidopsis TMO5/LHW, indicating a switch in target specificity during vascular plant evolution. Our data thus suggests that the TMO5/LHW function evolved from two proteins with distinct functions.

**Fig. 5:**
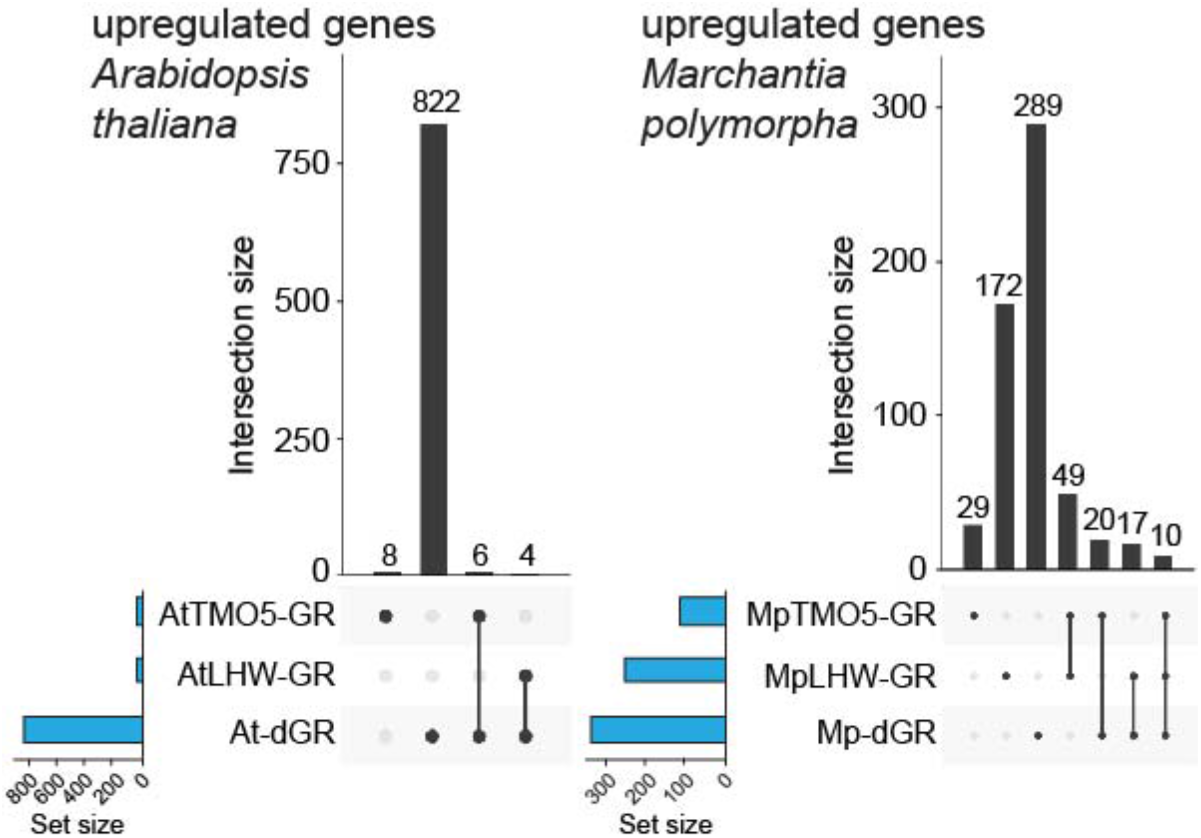
Independent transcriptional regulation by MpTMO5 and MpLHW. Comparison of genome-wide transcriptional changes, as measured by RNAseq, upon induction (upregulated genes only) of TMO5-GR, LHW-GR, or both in Arabidopsis (left graph) and Marchantia (right graph). UpSet plots show numbers of upregulated expressed genes in each DEX-treated transgenic line compared to DEX-treated wild-type (Col-0 in Arabidopsis, Tak-1 in Marchantia). Total numbers of genes in each line are shown in blue bars (set size), while gene numbers that are either unique to each line, or found in intersections (separated or connected black dots) are shown as black bars (intersection size).

### Two subsequent innovations led to a vascular function for TMO5/LHW

Clearly, modern TMO5/LHW function in vascular plants is not provided by gene absence or presence. Rather, properties of TMO5 and LHW proteins must underlie the acquisition of vascular activity. To identify the protein determinants conferring such activity, we analysed protein domains in detail. Besides the family-defining bHLH domain, an ACT-like domain was found in the C-terminal region of both AtTMO5 and AtLHW (**Fig. 2a,b; Fig. S1, S2**). ACT-like domains have been identified in other bHLH transcription factors, where these were shown to mediate protein-protein interactions^26, 33^. In addition, AtLHW contains a GAF-like domain at the N-terminus (**Fig. 2b; Fig. S2**), which is often associated with small-molecule binding and sensory functions such as oxygen and light sensing^34–36^. Interestingly, domain architectures of all TMO5 orthologs were identical across the plant kingdom (**Fig. 2a; Fig. S1**). While the ACT-like domain in LHW is lacking in Charophytes, it is present within Bryophytes in hornworts, but not in liverworts and mosses (**Fig. 2b; Fig. S2**). Given that all TMO5 orthologs tested can interact with AtLHW, but not *vice versa* (**Fig. 3; Fig. S8**), the domain architecture might be important for LHW function.

To first determine the importance of each domain for LHW function, we constructed domain deletions of AtLHW driven by its native promoter, and analysed the capacity to complement the *lhw* mutant (**Fig. 6a,b**). LHW proteins with a deletion of the ACT-like domain (LHW^ΔACT^) or the bHLH domain (LHW^ΔbHLH^) failed to rescue the mutant. Deletion of the N-terminal GAF-like domain (LHW^ΔGAF^) or the ∼300 amino acid spacer between the GAF-like and bHLH domain (LHW^Δspacer^) could partially rescue the reduced vascular bundle size, resulting a diarch phenotype in 29% and 40% of T1 roots, respectively (**Fig. 6a,b; Table S2**). Interestingly, having only the bHLH and the ACT-like domain (LHW^ΔGAFspacer^) was sufficient for partial rescue. Finally, removal of the 28 amino acid C-terminal tail of LHW (LHW^ΔC-terminus^) did not affect the vascular function and fully rescued the mutant phenotype (**Fig. 6a,b; Table S2**). We thus conclude that the bHLH and ACT-like domains are absolutely required for the vascular function of LHW, while the additional GAF domain and the spacer region are not. Given the importance of the ACT-like domain for a vascular function and considering its absence in non-vascular plants, we next tested if gain of the ACT-like domain was the sole causal evolutionary step required for LHW to adopt a vascular function. A chimeric *MpLHW* protein containing the ACT-like domain of *AtLHW* (MpLHW^AtACT^) could not complement the Arabidopsis *lhw* mutant (**Fig. 6c**), illustrating that other, yet unknown, modifications contribute to acquiring vascular function. The TMO5 domain architecture is conserved from Charophytes to Angiosperms, but KnTMO5 was not able to complement the Arabidopsis *tmo5 t5l1* double mutant (**Fig. 2i; Fig. S3i, S4d**). We therefore swapped domains of the complementing MpTMO5 into KnTMO5. Functionality of chimeric proteins was tested using complementation of the Arabidopsis *t5t5l1* double mutant (**Fig. 6d**). Although exchange of the N-terminal fragment (TMO5-C1) was not sufficient to complement the mutant phenotype, swapping both the bHLH and ACT-like domain (TMO5-C2) did provide complementation (**Fig. 6d**); suggesting that modifications to the bHLH or the ACT-like domain might be key to acquiring the vascular function. Single domain swaps of the ACT-like domain (TMO5-C3) did not result complementation. However, a swap of the bHLH domain and N-terminus (TMO5-C4) was able to rescue the mutant phenotype. Given that the HLH region of the bHLH domain is highly conserved between AtTMO5, MpTMO5 and KnTMO5 (**Fig. 6e**), we investigated the differences in the basic region, which is required for DNA binding specificity. Although AtTMO5 and MpTMO5 have similar basic regions, the first part is very different in KnTMO5 (**Fig. 6e**). Indeed, swapping only sixteen amino acids upstream of the HLH domain, including the basic domain (TMO5-C5) was sufficient to create a vascular function for KnTMO5 (**Fig. 6d, Fig. S9** and **Table S2**). Hence, we conclude that TMO5 is functionally conserved down to Bryophytes and with a small modification to the basic region even up to Charophytes. Given that TMO5 acts in a dedicated pathway associated with vascular plants, this is highly unexpected and establishes TMO5 as an ultra-conserved developmental regulator or a ‘molecular fossil’.

**Fig. 6:**
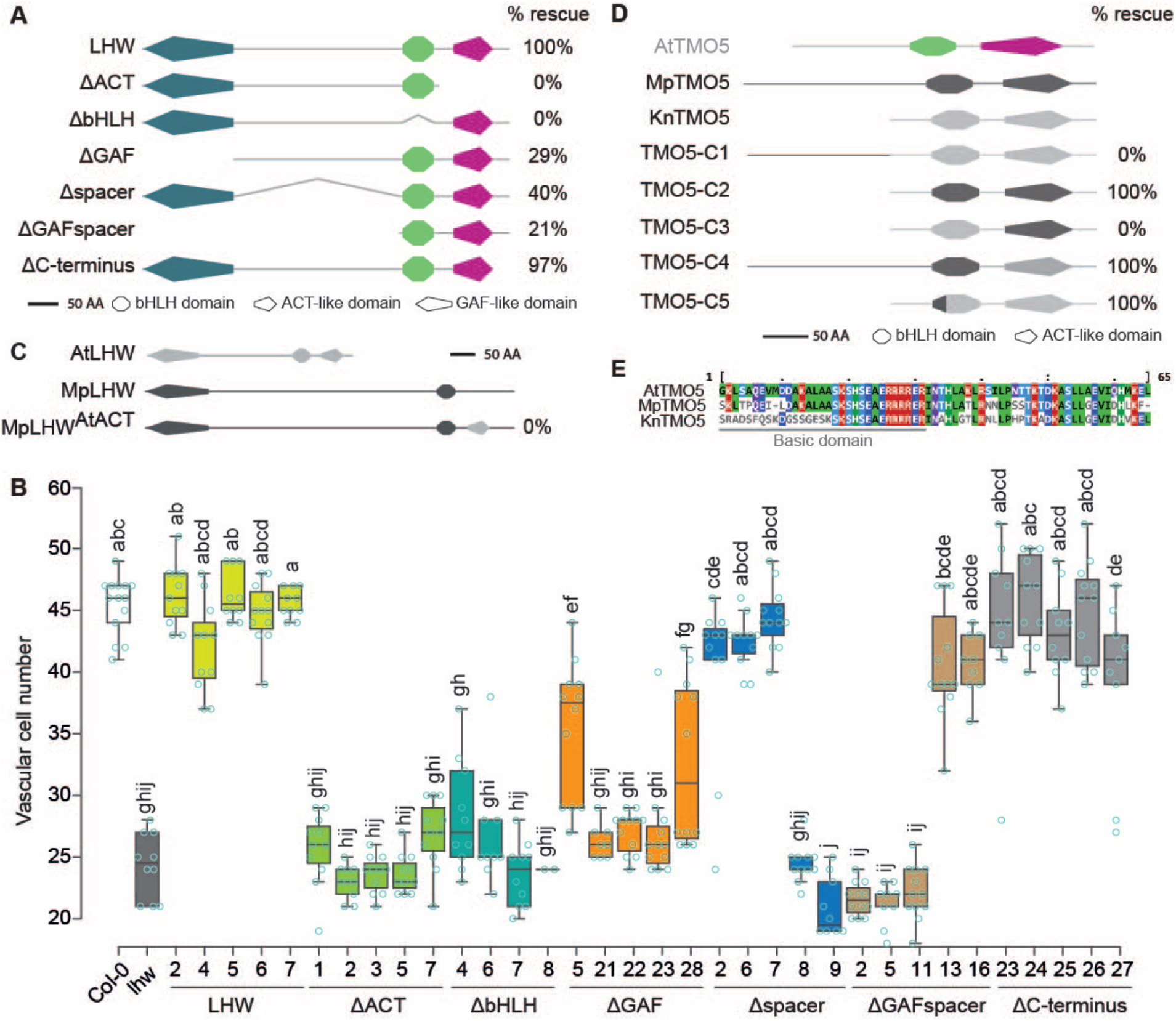
Contributions of LHW and TMO5 domains to modern vascular function. (A) Summary of domain deletions in Arabidopsis LHW, and their ability to complement the Arabidopsis *lhw* mutant. % Rescue indicates the percentage of transgenic lines with restoration of diarch vascular pattern in roots. (B) Quantification of vascular cell file number in radial sections of 5-day-old roots from lines carrying LHW domain deletions in the *lhw* mutant background. Data are plotted separately for multiple independent transgenic lines. Small case letters indicate significantly different groups as determined using a one-way ANOVA with post-hoc Tukey HSD testing. Blue circles represent all data-points. (C) Schematic representation of AtLHW, MpLHW and the chimeric MpLHW^AtACT^. Protein domains derived from AtLHW are in light grey and MpLHW domains in dark grey. No complementation (0 %) of the Arabidopsis *lhw* phenotype was observed with MpLHW^AtACT^. (D) Overview of domain swaps between MpTMO5 (dark grey) and KnTMO5 (light grey), and their ability to complement (% rescue) the diarch vascular pattern phenotype in the Arabidopsis *tmo5 t5l*1 mutant. Topology of Arabidopsis TMO5 is shown as reference. (E) Amino acid sequence alignment of *Arabidopsis*, *Marchantia*, and *Klebsormidium* TMO5 orthologues. The basic region is underlined. All proteins in (A, C, D) are drawn to scale (bar = 50 amino acids).

### Evolution of the TMO5/LHW dimer and the emergence of vascular plants

Here, we propose a two-step model for the evolution of the obligate TMO5/LHW heterodimer in vascular plants. In a first step with the emergence of land plants, point mutations in the basic region of the bHLH domain conferred the modern function to TMO5, which has been maintained in all subsequent lineages (**Fig. S10**). In a second step, gain of an ACT-like domain and other, yet unknown, modifications to LHW led to a preference for TMO5/LHW heterodimerization rather than homodimerization or interactions with other partners. This modification extended the range of target gene recognition and generated the function that is conserved among vascular plants (**Fig. S10**). Intriguingly, the extreme functional conservation of TMO5 suggests that modifications specifically to LHW were required to develop the vascular function of the TMO5/LHW complex. Importantly, the origin of these modifications to LHW that lead to an obligate TMO5/LHW heterodimer complex with vascular function are strongly correlated with the appearance of Tracheophytes. Although it is currently impossible to prove causality of this innovation to vascular tissue elaboration, it is worth highlighting that there is no such correlation for other pathways involved in vascular development, such as auxin signalling^16^, CK signalling^37^ or secondary cell wall formation through VND transcriptional regulators^9^. As such, we provide the first example of a developmental regulator for which the innovations leading to a vascular function are clearly correlated to the emergence of vascular plants. This leads to the tantalising hypothesis that the acquisition of cell division capacity leading to a larger bundle of cells might have been a crucial innovation leading to vascular plant evolution. This is further supported by genetic analysis in Arabidopsis that revealed a large degree of self-organisation in vascular tissue development, constrained by the number of cell files. Indeed, different doses of TMO5 and/or LHW lead to a range of vascular bundle cell file numbers. Thus, while cell division alone is likely not the only driver, it conceivably is an important evolutionary innovation enabling the emergence of vascular plants.

## Acknowledgments

The authors thank Veronique Storme and Helena Arents for help with statistical analysis. This work was supported by grants from the Netherlands Organization for Scientific Research (NWO; VIDI-864.13.001 to B.D.R; VICI-865-14-001 to D.W.), the European Research Council (ERC; StG TORPEDO; 714055 to B.D.R.) and the European Molecular Biology Organization (EMBO; ALTF 415-2016 to H.K.).

## Author contributions

B.D.R. and D.W. conceived the project; N.v.W.H., K-J.L., B.D.R. and D.W. designed experiments; N.v.W.H. and K-J.L. performed most of the experimental work, with assistance of P.A. and H.K.; S.M. assisted in the analysis of RNA-seq data; K.V. assisted in sequence alignments, phylogenetic tree construction and RNA-seq gene conservation analysis; E.M. performed the histological analyses; C.A. assisted in BiFC experiment; B.D.R. and D.W. supervised the project; N.v.W.H., K-J.L., D.W. and B.D.R. wrote the manuscript with input from all authors.

## Competing interests

Authors declare no competing interests.

## Data and materials availability

All RNA-sequencing data is available from NCBI as BioProject number PRJNA528622 (http://www.ncbi.nlm.nih.gov/bioproject/528622). All other data is available in the main text or the supplementary materials.

## Supplementary Materials

**Other Supplementary Materials for this manuscript include the following**

**Table S1:** Overview of the complementation experiment showing the percentage of rescue in independent T2 lines of *tmo5 tmo5-like1* and *lhw* mutants in Arabidopsis by introduction of TMO5 and LHW orthologs, respectively.

**Table S2:** Overview of the complementation experiment showing the percentage of rescue in independent T2 lines of lhw mutants in Arabidopsis by introduction of LHW domain deletion constructs (left) and overview of the domain swaps between MpTMO5 and KnTMO5, and their ability to complement (% rescue) the diarch vascular pattern phenotype in the Arabidopsis *tmo5 t5l1* mutant (right).

**Table S3:** Overview of the species used in the phylogenetic analysis, their abbreviation, clade, order and from which database they were retrieved.

**Table S4.** Overview of all transcriptomic analyses.

**Table S5.** DNA alignment of synthetic MpLWH CDS and that obtained from phytozome.

**Table S6.** Primers used in this research. Different colours indicate the adapter sequences for different cloning methods. Blue, TOPO cloning; Orange, ligation; Green, SLiCE

**Data S1.** FASTA file including all sequences used for the TMO5-related alignments

**Data S2.** FASTA file including all sequences used for the LHW-related alignments

### Materials and Methods

#### Sequence data

Genomic sequences of plants species *Cyanidioschyzon merolae*, *Ostreococcus lucimarinus*, *Chlamydomonas reinhardtii, Physcomitrella patens*, *Selaginella moellendorffii*, *Amborella trichopoda, Oryza sativa*, *Zea mays, Arabidopsis thaliana, Solanum lycopersicum and Populus tremulus* were retrieved from the comparative genomics database PLAZA 3.0, PLAZA 2.5 and PicoPLAZA (Ref. 38; http://bioinformatics.psb.ugent.be/plaza). Genomic data of *Marchantia polymorpha* was accessed via Phytozome v12 (Ref. 39, https://phytozome.jgi.doe.gov/pz/portal.html). *Klebsormidium nitens* genomic sequences were collected from the *Klebsormidium nitens* NIES-2285 genome project (Ref. 40; http://www.plantmorphogenesis.bio.titech.ac.jp/~algae_genome_project/klebsormidium/). Transcriptomic data of *Culcita macrocarpa, Equisetum diffusum, Picea engelmanni, Marchantia polymorpha* and a wide range of lycophytes, bryophytes and algae were derived from the transcriptome database OneKP, to which access was kindly provided by the developers of OneKP^41^ (**Table S3**). For extensive LHW and TMO5 analyses full length protein sequences of TMO5, TMO5-LIKE1 (T5L1), TMO5-LIKE2 (T5L2), TMO5-LIKE3 (T5L3), TMO5-LIKE4 (T5L4), LHW, LHW-LIKE1 (LL1) and LHW-LIKE2 (LL2) were used. PPR regions of LHW-LIKE3 (LL3) were excluded from the search query. Transcriptome assembly outputs resulting from tBLASTn searches were translated into protein sequences prior to alignments. Overlap between differentially expression gene sets from different species were performed using the Workbench in PLAZA Dicots 4.0.

#### Sequence alignments and phylogenetic tree construction

Full-length plant proteins were aligned using MAFFT version 7 (Ref. 42; http://mafft.cbrc.jp/alignment/server/), using default parameters. Gaps in the aligned sequences were removed in BioEdit v7.2.5 (Ref. 43). MSA in the Figures were visualized by Jalview v2.10.2 using clustal based colour coding. BIONJ phylogenetic trees were estimated using the program PhyML version 3.0 (*44;* http://www.atgc-montpellier.fr/phyml/), applying the LG amino-acid replacement matrix using 20 substitution rate categories and an estimated gamma distribution parameter. A Nearest Neighbor Interchange (NNI) topology improvement was applied. For TMO5 and LHW phylogenetic reconstruction, a Neighbour Joining (BIONJ) tree was calculated with an Approximate Likelihood-Ratio Test (aLRT). For confined TMO5 and LHW trees 100 bootstrap replicas were calculated (see **Fig. S1** and **Fig. S2**). The files containing all sequences used in FASTA format are present as **Data S1** for TMO5 and **Data S2** for LHW.

#### Plant material and growth conditions

All seeds were surface sterilized and grown on ½ MS plates. After a two-day stratification at 4°C, seedlings were grown under long day conditions (16 hours light, 8 hours dark) at 22°C. Arabidopsis ecotype Columbia-0 (Col-0) was used as wild-type. *lhw, lhw ll1, tmo5 and tmo5 t5l1* mutants were generated by and obtained from De Rybel et al., 2013 and De Rybel et al., 2014. *Marchantia polymorpha* strain Takaragaike-1 (Tak-1) was used as wild type and in all transformations and physiological analyses in this study. The growth condition for *Marchantia* is as follow: 50-60 μmol photos m^-2^s^-1^ white LED light at 22°C with continuous light.

#### Marchantia transformation

For gain-of-function and loss-of-function mutant generation, the agrobacterium dependent thallus transformation was performed as previously described^45^: each 14-day-old Tak-1 thallus was dissected into 8 pieces after removing the apical meristem niches and transferred onto half-strength Gamborg B5 (1/2 B5) medium with 1% sucrose and 1% Daishin agar plates for 3 days. The thallus pieces were further transferred and incubated with agrobacterium containing indicated plasmids in 50 ml 0M51C medium with 200μM Acetosyringone (4’-Hydroxy-3’,5’-dimethoxyacetophenone) in 200-ml flasks with agitation at 130 rpm for another 3 days. The thalli were further filtered and wished with water and transferred onto 1/2 B5 plates containing respective antibiotics for selection. Independent T1 lines were isolated and single G1 lines from independent T1 line were generated by subcultivating single gemmalings which emerged asexually from single initial cells^46^. The next generation of G1, called G2 generation, was used for analyses.

#### Cloning

Complementation vectors expressing p*TMO5* or p*LHW* were constructed through conventional cloning of pPLV28 (Ref. 47). 3 kb fragments upstream of the *TMO5* or *LHW* ATG were amplified from genomic DNA using Phusion Flash PCR Master Mix (Thermo Scientific). All further cloning procedures were performed using Seamless Ligation Cloning Extract (SLiCE), with 15 homologous bases^48^. Using primers with flanking LIC sites, a YFP was inserted in the LIC site of p*LHW*::LIC, creating p*LHW*::YFP. cDNA of *Populus trichocarpa, Oryza sativa, Picea abies and Selaginella moellendorffii* was used to amplify orthologs, while *MpTMO5*, *MpLHW*, *KnLHW*, *EgLHW*, *EdTMO5*, *SwLHW*, *CmaLHW* and *Cma2008862* were gene synthesized by GeneArt or GenScript. KnTMO5 CDS was amplified from genomic DNA (see below) by stitching PCR. The cDNAs of TMO5 orthologs were amplified and introduced in pGIIB-p*TMO5::*LIC-NOSt. The cDNAs of LHW orthologs, excluding the stop codon, were amplified and cloned into pGIIB-p*LHW::*LIC*-*YFP-NOSt. BiFC plasmids were constructed by cloning the orthologs into a modified pPLV22 or pPLV27 vector containing a p*35S*::LIC-nYFP or p*35S*::LIC-cYFP respectively. All constructs were verified by sequencing and transformed into Arabidopsis Col-0 plants by simplified floral dipping^49^. All *MpTMO5* related constructs and the *MpLHW* promoter construct used for generating Marchatnia transformants were cloned by PCR amplification from the gnomic DNA with primers listed in **Table S6**. For MpLHW overexpression and GR lines, the MpLHW CDS was synthesized (**Table S5**) and sub-cloned by PCR amplification with primers listed in **Table S6**. The amplified fragments were cloned into pENTR-D-TOPO vector (Invitrogen) according to the user manual and further subcloned into pMpGWB vectors as previously described^50^. For FRET-FLIM analysis, MpTMO5 genomic fragment and MpLHW coding sequence were amplified by PCR with primers listed in **Table S6** and cloned into pMON999 vectors with sCFP3A and sYFP2 to create MpTMO5-sCFP3A and MpLHW-sYFP2 constructs by SLiCE cloning method^51^. The Arabidopsis *TMO5* and *LHW* constructs were adapted as previously described^52^. For genomic DNA analysis, small pieces of thalli were collected by forceps and homogenized with liquid nitrogen in the Retsch MM400 machine (Retsch GmbH) and DNA was extracted with 2x CTAB buffer (2% CTAB, 1.4M NaCl, 100 mM Tris-HCl pH 8.0, and 20 mM EDTA). The desired fragments were amplified by primers listed in **Table S6** and were further analyzed by sequencing (sequencing primers are indicated in **Table S6**). Sequence alignment was performed on Benchling (benchling.com) by using genomic sequence obtained from phytozome (Ref. 39; https://phytozome.jgi.doe.gov/) (Marchantia polymorpha v3.1).

#### Complementation assays

For complementation assays, plasmids carrying the LHW orthologs were transformed into *lhw* plants by simplified floral dipping^49^. Rescue was analyzed in T1 as well as T2 plants. Approximately 10 individual seedlings were screened per T2 line. Plasmids carrying the TMO5 orthologs were transformed to *tmo5 t5l1* plants, homozygous for *tmo5*, heterozygous for *t5l1* (obtained by crossing *tmo5* mutant plants with *tmo5 t5l1* double mutant plants). In T1 lines, rescue was considered when 80% or more of the independent lines showed complementation. Homozygous *tmo5 t5l1* double mutant backgrounds were selected in MpTMO5 and KnTMO5 T2 lines by screening vascular pattern in seedlings that lost the transgene by segregation.

#### DNA extraction Klebsormidium

Klebsormidium, freshly grown on BCD agar medium, was inoculated in 200ul undiluted Edwards solution (200 mM Tris-HCl pH 7.5, 250 mM NaCl, 25 mM EDTA, and 0.5% SDS) for 1 hour at 90°C. After centrifugation, the supernatant was diluted 20 times to obtain a working stock Klebsormidium DNA.

#### Bimolecular Fluorescence Complementation (BiFC)

*Agrobacterium tumefaciens* containing BiFC plasmids were cultured overnight at 28°C/250 rpm in 5ml LB medium containing 50 µg/ml kanamycin, 25 µg/ml rifampicin, 2 µg/ml tetracyclin and 200 µM acetosyringone. The bacteria were collected by centrifugation (4000rpm, 10 minutes) and resuspended in MMA infiltration medium (20 g/l sucrose, 5 g/l MS-salts, 2 g/l MES, pH 5.6) containing 200 µM acetosyringone to an optical density (OD_600_) of 0.3. BiFC samples were mixed in a 1:1 ratio to a total OD_600_ of 0.6. Samples were incubated for 1-2 hours at room temperature (RT) under continuous shaking. The abaxial side of the two youngest, fully expanded leaves of 5 – 6 weeks old *Nicotiana benthamiana* plants were infiltrated with a 1 ml needless syringe. Infiltrated leaves were harvested after 2 – 3 days and imaged with a confocal microscope.

#### FRET-FLIM

The FRET-FLIM analysis was performed as previously described^53^. FRET-FLIM measurements were performed on a Leica SP8 system with an HyD detector. A diode-pulse laser is used to generate the single-photon excitation with 40 Mhz. FRET between sCFP3A and sYFP2 was detected by monitoring donor emission. Images with a frame size of 128 × 128 pixels were acquired, and the average count rate was around 10^4^ photons per second for an acquisition time of ±60s. Donor FLIMs (sCFP3A) were analyzed with SPCImage 3.10 software (Becker & Hickl) using a two-component decay model. Several cells (n > 15) were analyzed, and average FLIMs of different combinations were exported for generating a box plot. Statistical significance of differences between samples was determined using a two-tailed Student’s t test.

#### Plant imaging and Quantification of vascular cell numbers

For confocal imaging of monarch or diarch root architecture, 5 – 7 day-old-seedlings were stained with 10 µg/ml Propidium Iodide. Confocal imaging was done one a Leica SP5 confocal microscope, with a HyD detector. Modified Pseudo Schiff – Propidium Iodine (mPS-PI) staining was performed as described previously^54^. Radial cross sections and quantification of vascular cell numbers were obtained using a SP5 confocal microscope on 5 day-old-roots that were stained using mPS-PI. Sections were taken at the middle of the root meristem: half way between the QC and the first elongating cortex cell. The data was visualized using BoxPlotR.

#### Histological analysis and sectioning

All histological analysis, sectioning and counterstaining using toluidine blue and ruthenium red was performed as described previously^52, 55^.

#### RNA-sequencing

For *Arabidopsis thaliana*, seeds of Col-0, p*RPS5A*::TMO5-GR, p*RPS5A*::LHW-GR and p*RPS5A*::TMO5-GR x p*RPS5A*::LHW-GR (dGR; Ref. 52) were surface sterilized and stratified for 2 days in the cold room then were plated on MS plates with nylon meshes (pore size 100 μm). All plates were incubated in a climate chamber at 22 °C with 16h/8h day/night cycle for 7 days and then transferred to MS plates containing 10 μM dexamethasone for 1hr. All seedling roots (about 1.5 cm) of each line were harvested and subjected to RNA isolation. For *Marchantia polymorpha*, gemmalings of Tak-1, pEF::MpTMO5-GR, pEF::MpLHW-GR and pEF::dGR were transferred to 1/2 B5 medium plates with nylon mesh (pore size 100 μm) and incubated in an incubator at at 22°C with 16h/8h day/night cycle for 10 days then transferred to 1/2 B5 plates containing 10 μM dexamethasone for 1hr. All gemmalings of each line were harvested and subjected to RNA isolation. Total RNA (at least 500 ng) of each sample with was delivered to BGI (HongKong, China) for f RNA-sequencing. Raw reads were mapped to the respective genomes using HISAT2 (Ref. 56) with additional parameters “--trim5 10 --dta”. Read counts were calculated using feature Counts implemented in the subread package (v1.6.0). The obtained raw read counts were normalized and differentially expressed genes (Padj <0.01) were identified using DEseq2 (Ref. 57) implemented in R Bioconductor package. To determine the overlap of induced genes by combined overexpression of TMO5 and LHW between Arabidopsis and Marchantia (**Table S4**), we used the PLAZA Dicots 4.0 Workbench^58^. First, starting from the Mp double-GR genes, Arabidopsis orthologs were identified using the integrative orthology method requiring at least one evidence, considering Tree-based orthologs, Best Hit Family and Orthologous gene family information, yielding a set of 1322 Arabidopsis genes. Next, this set was compared with the At double-GR genes, yielding an overlap of 145 genes (9%, 145/1647). All RNA-sequencing data is available from NCBI as BioProject number PRJNA528622 (http://www.ncbi.nlm.nih.gov/bioproject/528622).

---

**Fig. S1.**
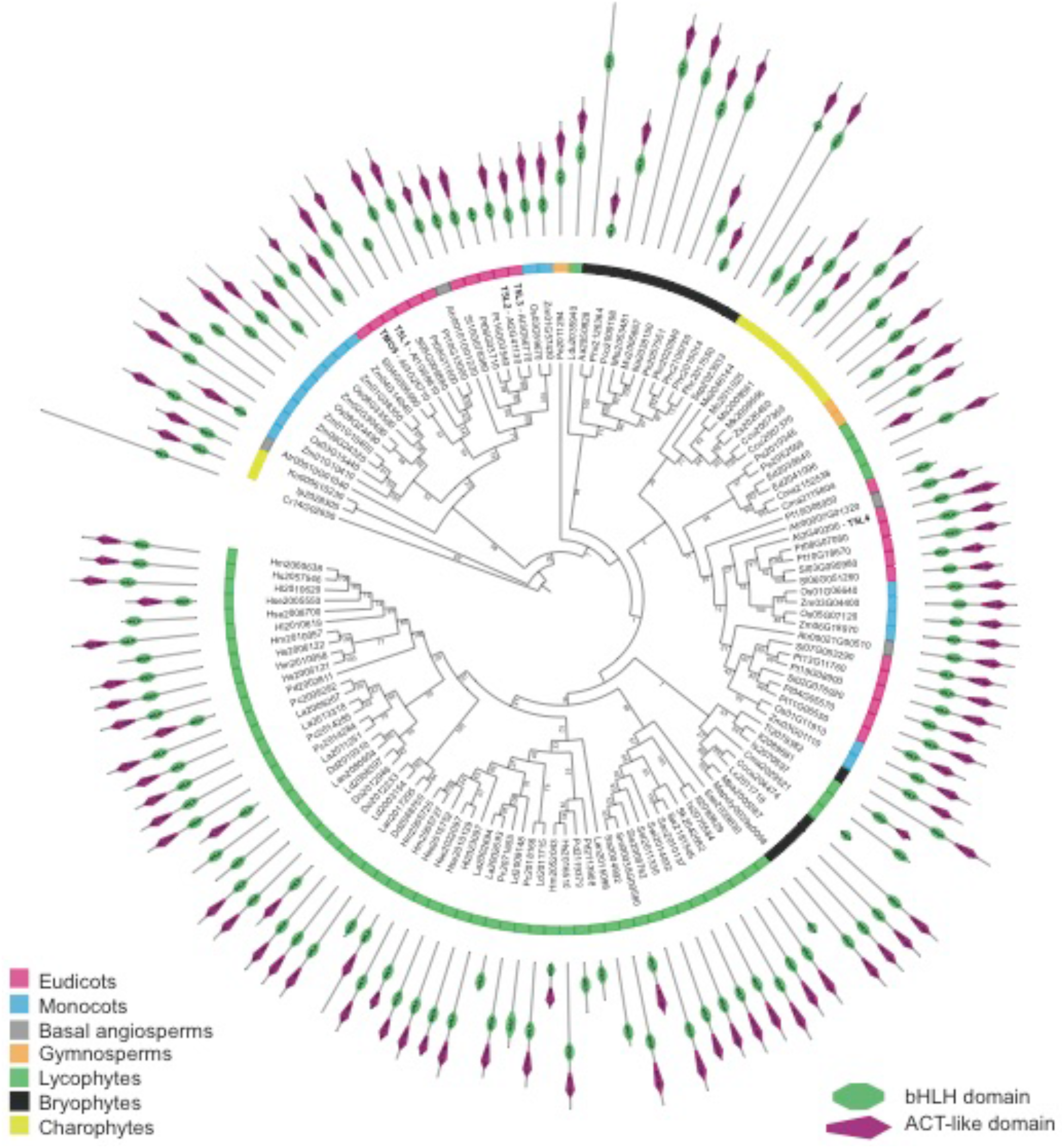
The Neighbour Joining phylogenetic tree of TMO5 orthologs in the plant kingdom. The phylogenetic tree was rooted with an algal protein and 100 bootstrap replicas were calculated. The coloured ring indicates different clades of species which is referred to the legend. Additionally, the domain architecture is shown for each protein in which the green shape represents the bHLH domain and the magenta shape the ACT-like domain.

**Fig. S2.**
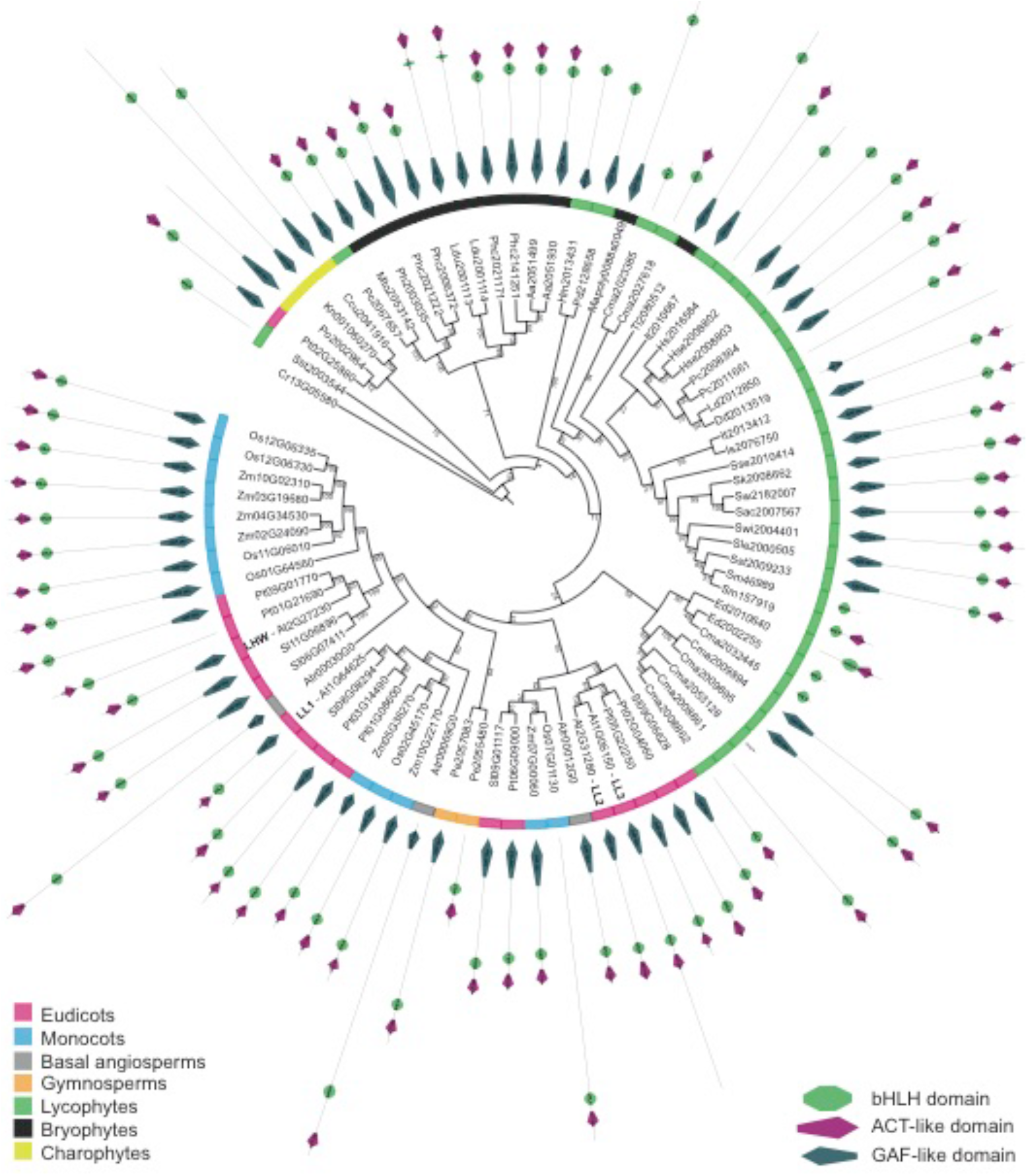
The Neighbour Joining phylogenetic tree of LHW orthologs in the plant kingdom. The phylogenetic tree was rooted with an algal protein and 100 bootstrap replicas were calculated. The coloured ring indicates different clades of species which is referred to the legend. Additionally, the domain architecture is shown for each protein in which the green shape represents the bHLH domain, the magenta shape the ACT-like domain and the dark green shape the GAF-like domain.

**Fig. S3.**
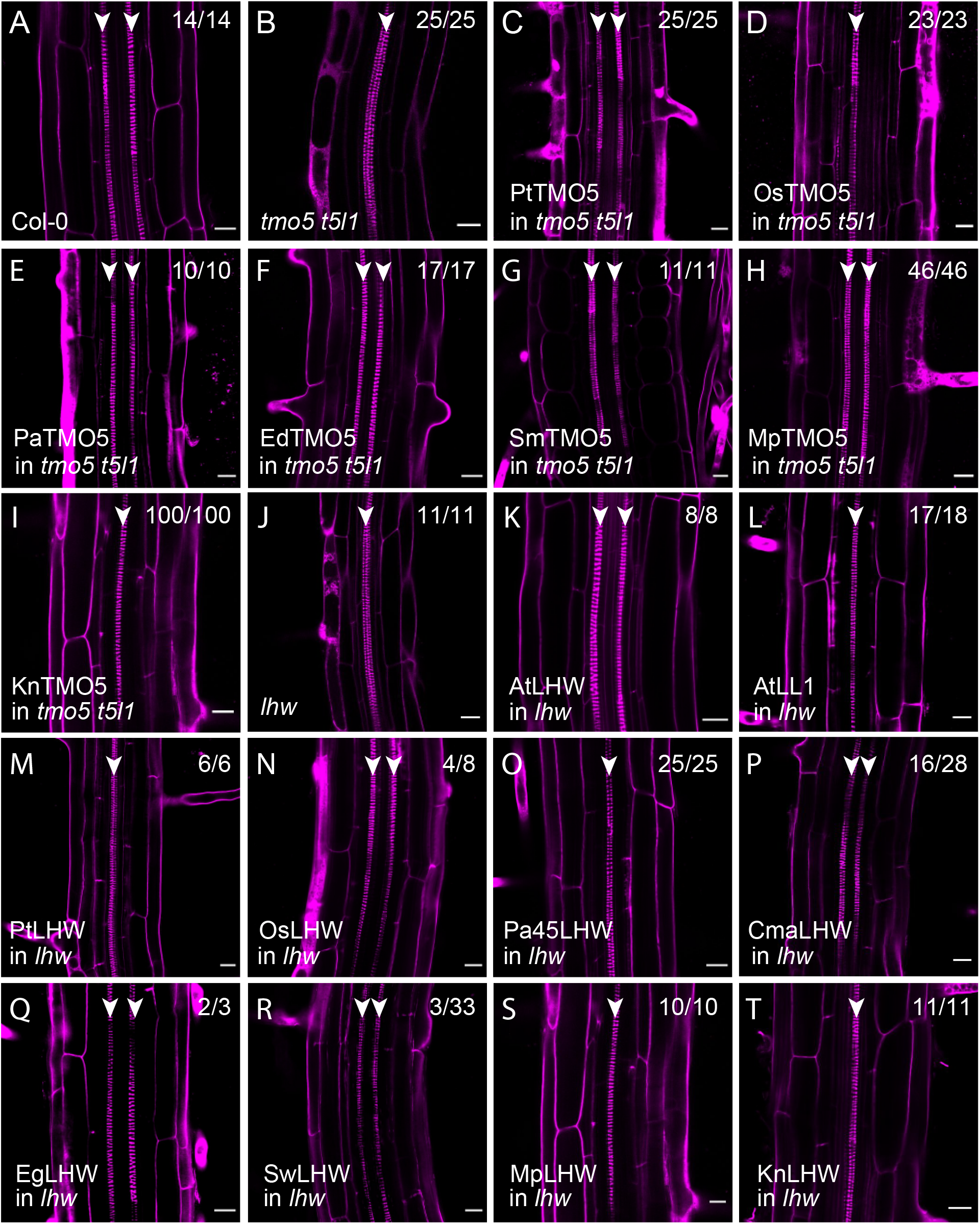
Complementation of the *t5t5l1* double and *lhw* single mutant vascular phenotypes by their respective orthologs. **A.** Wild type Arabidopsis root with two xylem poles. **B.** Arabidopsis *t5t5l1* double mutant root showing only one protoxylem pole. **C-I.** Complementation of the *t5t5l1* vascular phenotype by introduction of the TMO5 orthologs from *Populus trichocarpa* (*Pt -* Pt08G11600), *Oryza sativa* (*Os -* Os03G15440), *Picea abies* (*Pa -* Pa65818g0010), *Equisetum diffusum* (*Ed -* Ed2039640), *Selaginella moellendorffii* (*Sm -* Sm00038G00580), *Marchantia polymorpha* (*Mp -* Mapoly0039s0068) and *Klebsormidium nitens* (*Kn -* Kn000610230). **J.** Arabidopsis *lhw* mutant root showing only one protoxylem pole. **K-T.** Complementation of the *lhw* vascular phenotype by introduction of the LHW homologs from *Arabidopsis thaliana* (AtLHW - At2G27230 and AtLL1) and the orthologs from *Populus trichocarpa* (*Pt*), *Oryza sativa* (*Os -* Os01G64560), *Picea abies* (*Pa*), *Cultita macrocarpa* (*Cma -* Cma2009894), *Equisetum giganteum* (*Eg -* Eg1939), *Selaginella wallace* (*Sw -* Sw2182007), *Marchantia polymorpha* (*Mp -* Mapoly0088s0049) and *Klebsormidium nitens* (*Kn -* Kn001060270). Scale bars represent 20 mm; arrowheads indicate protoxylem poles. In all panels, the number indicated in the right top corner how many show the phenotype out of the total number of individual T1 lines analysed.

**Fig. S4.**
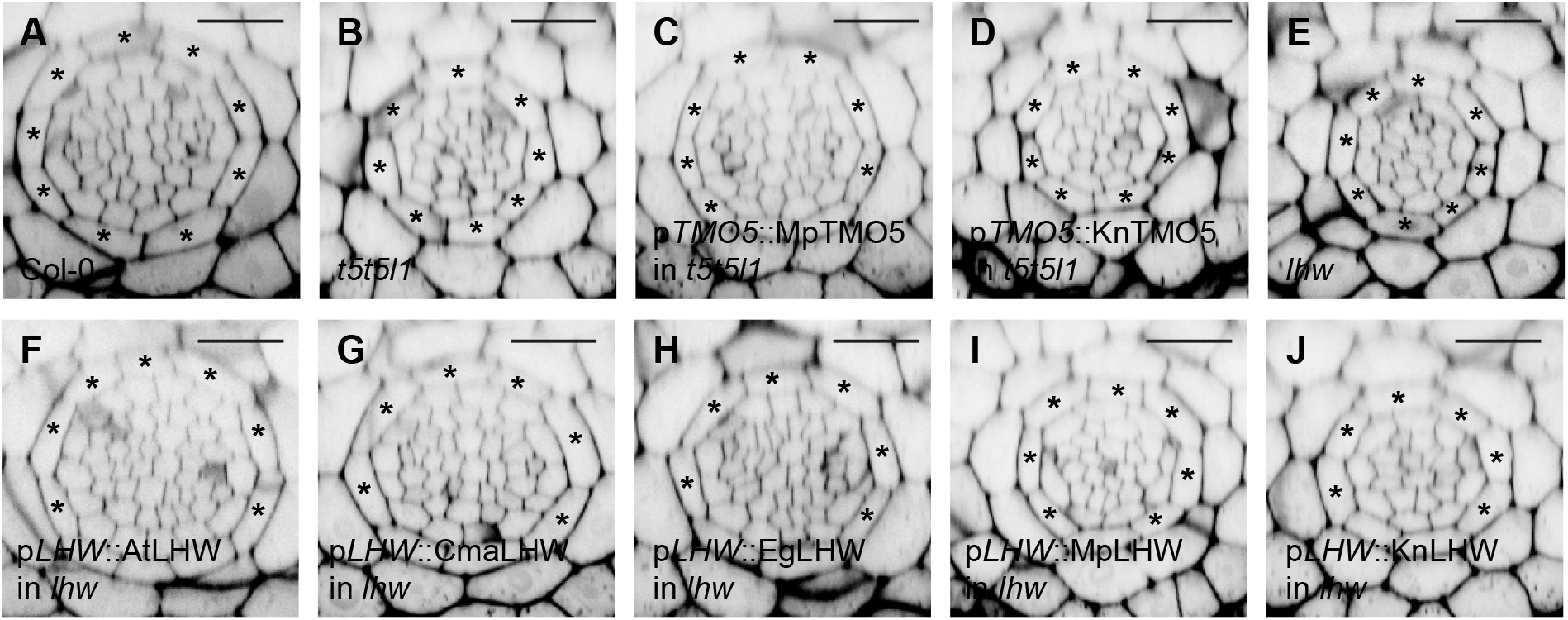
Representative confocal radial cross sections of the complementation study. **A.** Col-0. **B.** The *tmo5 t5l1* double mutant. **C-D.** Complementation of the *tmo5 t5l1* double mutant with MpTMO5 (Mapoly0039s0068) (C) and KnTMO5 (Kn000610230) (D). **E.** The *lhw* single mutant. **F-J.** Complementation of the *lhw* mutant with AtLHW (At2G27230) (F), CmaLHW (Cma2009894) (G), EgLHW (Eg1939) (H), MpLHW (Mapoly0088s0049) (I) and KnLHW (Kn001060270) (J). Scale bars represent 20 mm. Asterisks indicate endodermis cells.

**Fig. S5.**
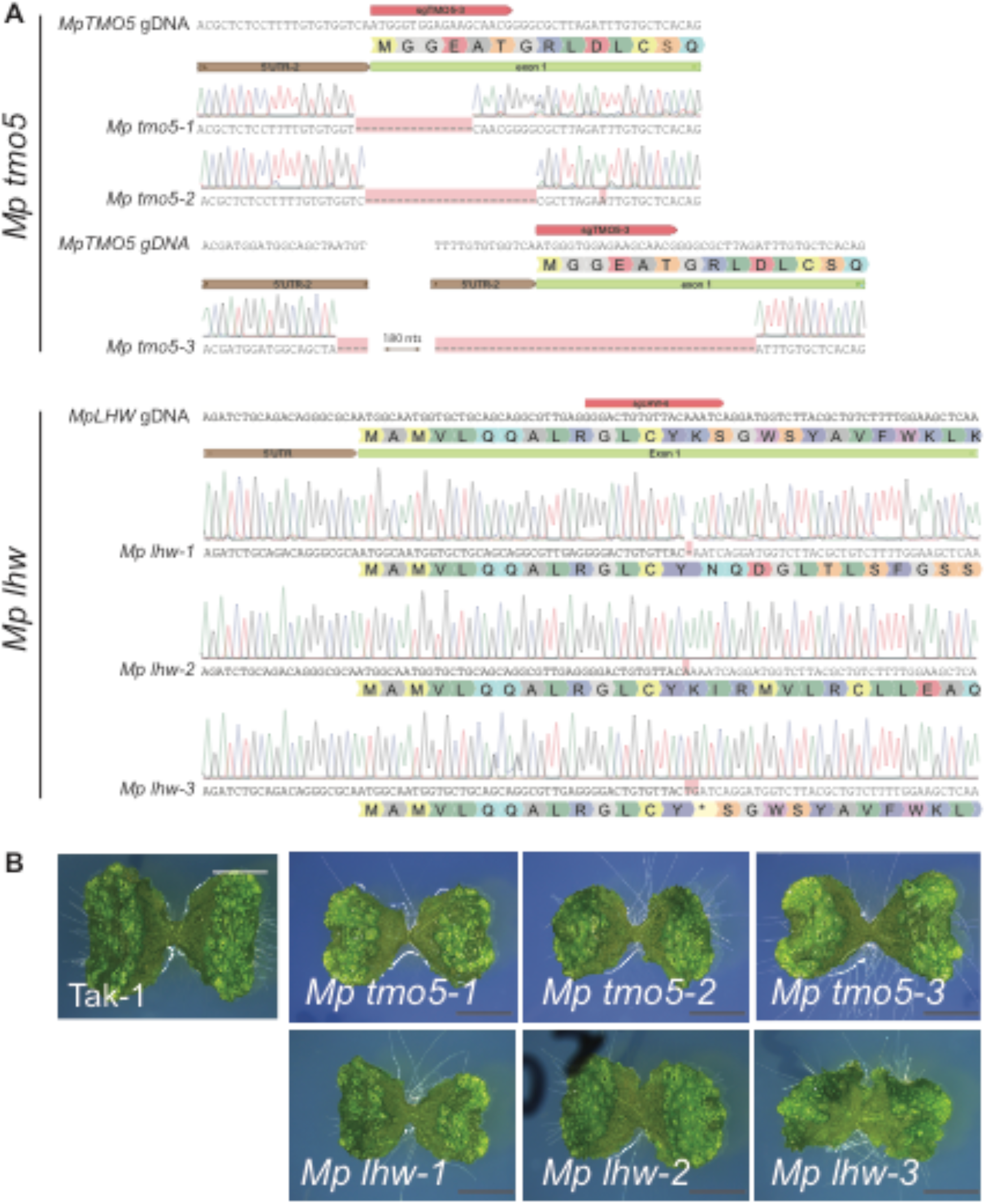
Molecular and phenotypical characterisation of multiple independent *Mp tmo5* and *Mp lhw* CRISPR/Cas9 mutant lines. **A.** Location of the *Mp tmo5* and *Mp lhw* CRISRP/Cas9 mutations compared to the control genomic DNA sequence. Red tags above the genomic DNA indicate the target region of sgRNAs and mutations are labeled by red colors. The translated amino acid sequences are indicated for each *lhw* mutant. Note that the *Mp tmo5* mutants all lack the original start codon. **B.** Resulting young gemmaling phenotypes of the CRISPR/Cas9 mutants compared to the control plants (Tak-1).

**Fig. S6.**
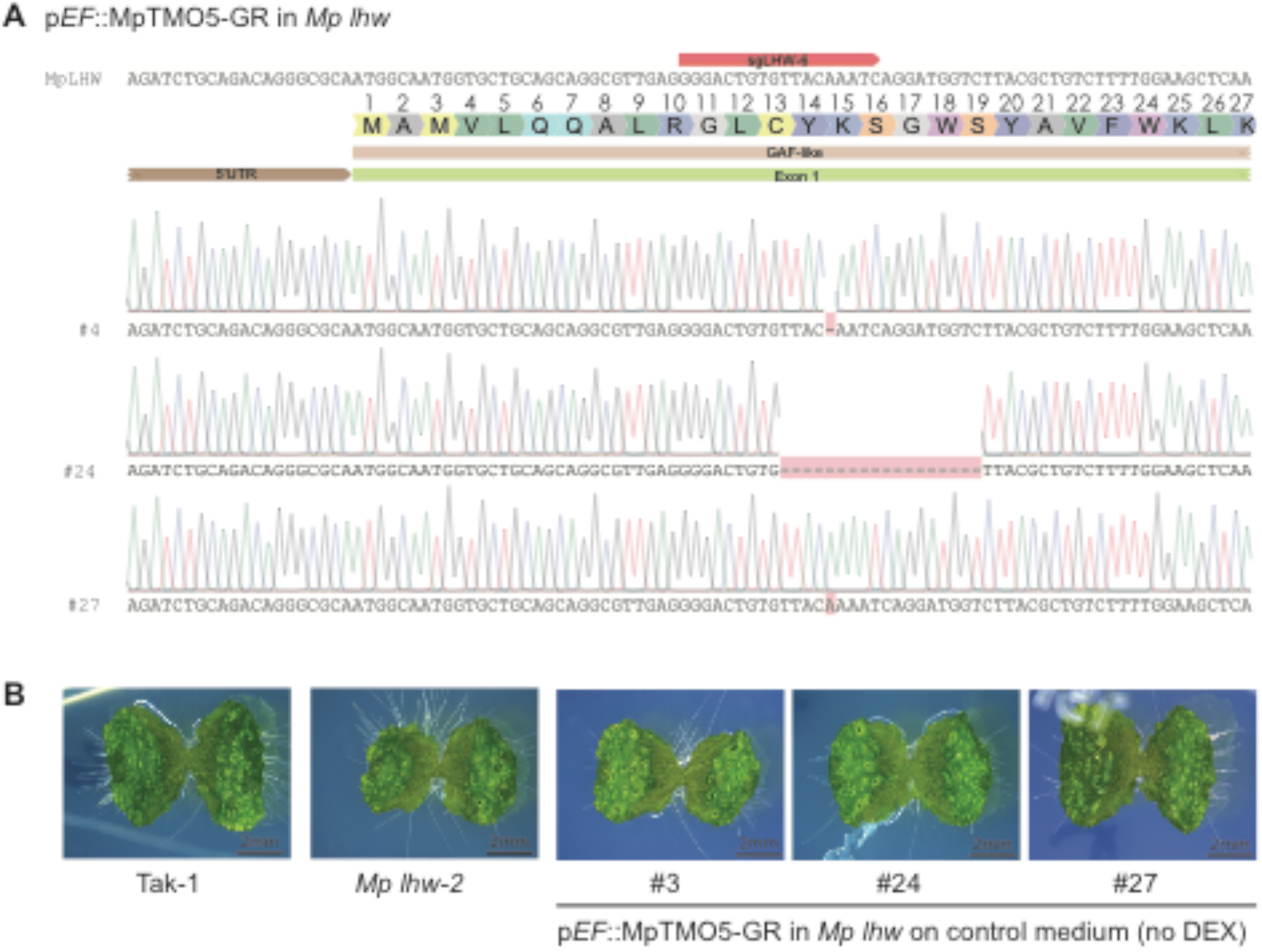
pEF::MpTMO5-GR in *Mp lhw* mutants show similar phenotype to *Mp lhw*. **A**. Location of the CRISRP/Cas9 mutations in MpLHW compared to the control genomic DNA sequence. Red tags above the genomic DNA indicate the target region of sgRNAs and mutations are labeled by red colors. **B.** Young gemmaling phenotypes of the control Tak-1, the CRISPR/Cas9 *Mp lhw-2* mutant, and 3 independent lines of the *Mp lhw* mutant carrying a pEF::MpTMO5-GR construct. Scale bars represent 2 mm.

**Fig. S7.**
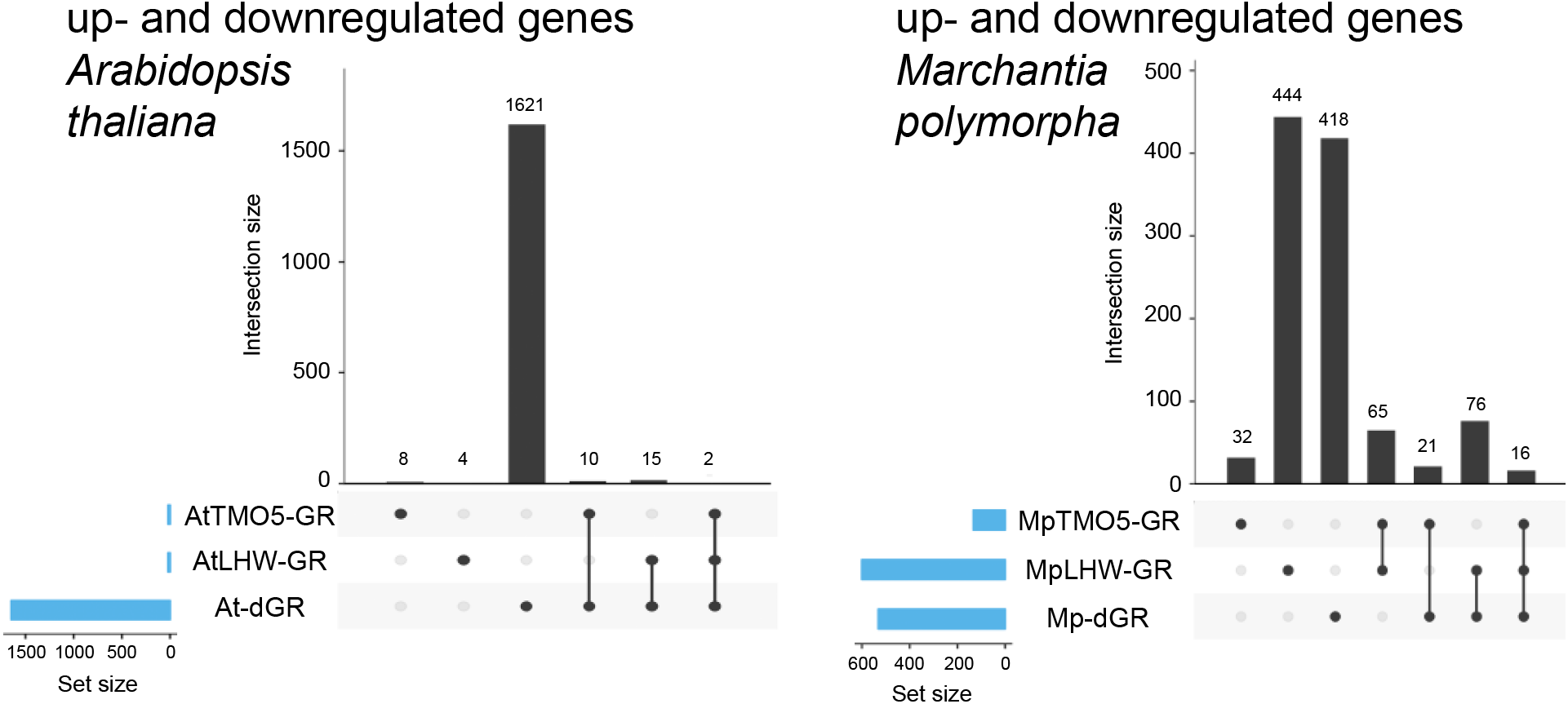
Independent transcriptional regulation by MpTMO5 and MpLHW. Comparison of genome-wide transcriptional changes (up- and down-regulated genes combined), as measured by RNAseq, upon induction of TMO5-GR, LHW-GR, or both in Arabidopsis (left graph) and Marchantia (right graph). UpSet plots show numbers of differentially expressed genes (DEG) in each DEX-treated transgenic line compared to DEX-treated wild-type (Col-0 in Arabidopsis, Tak-1 in Marchantia). Total DEG numbers in each line are shown in blue bars (set size), while DEG numbers that are either unique to each line, or found in intersections (separated or connected black dots) are shown as black bars (intersection size).

**Fig. S8.**
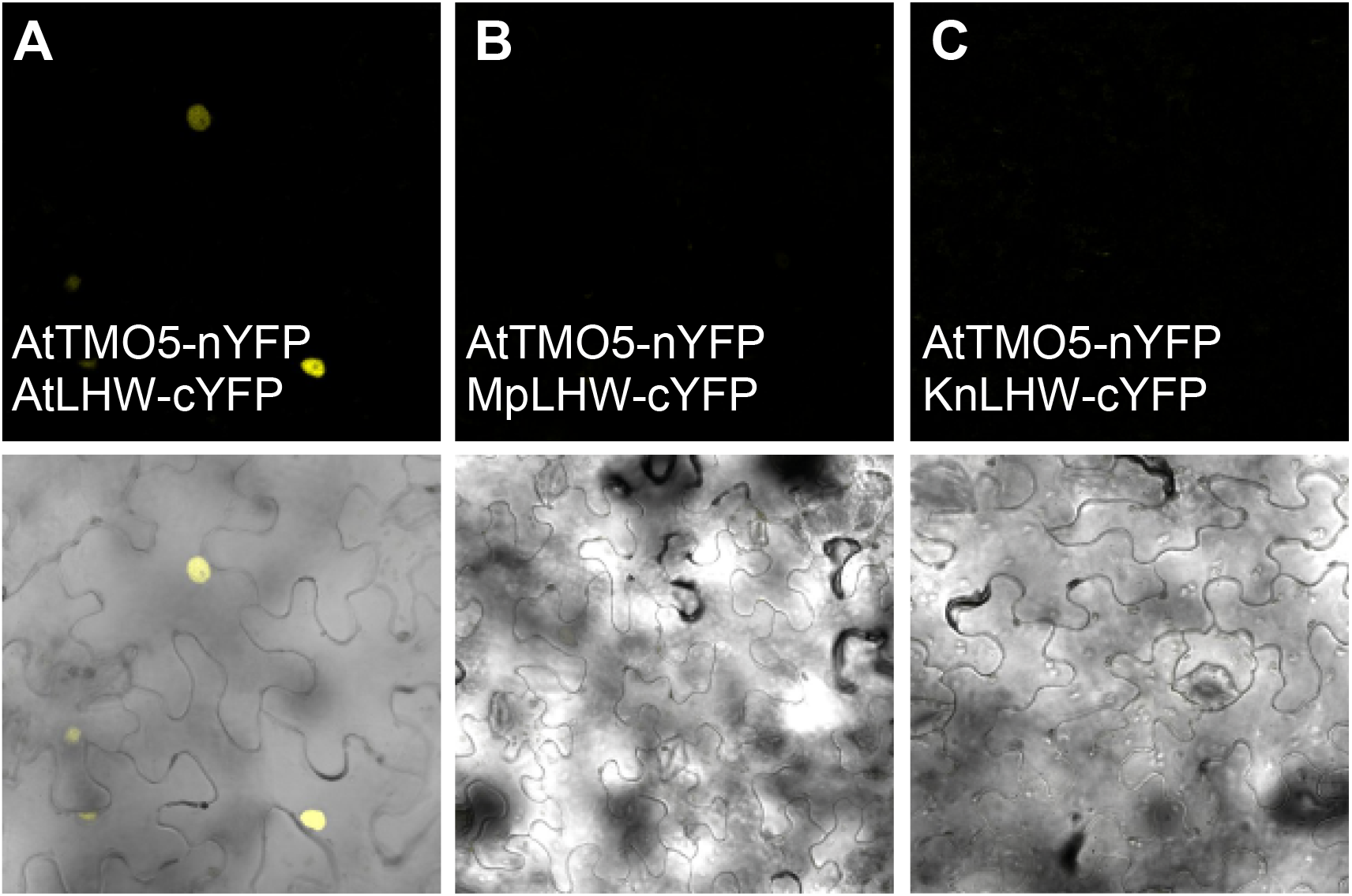
LHW orthologs do not have conserved dimerization capacities. BiFC results of tested interactions between AtTMO5 and AtLHW (**A**), MpLWH (**B**) and KnLHW (**C**).

**Fig. S9.**
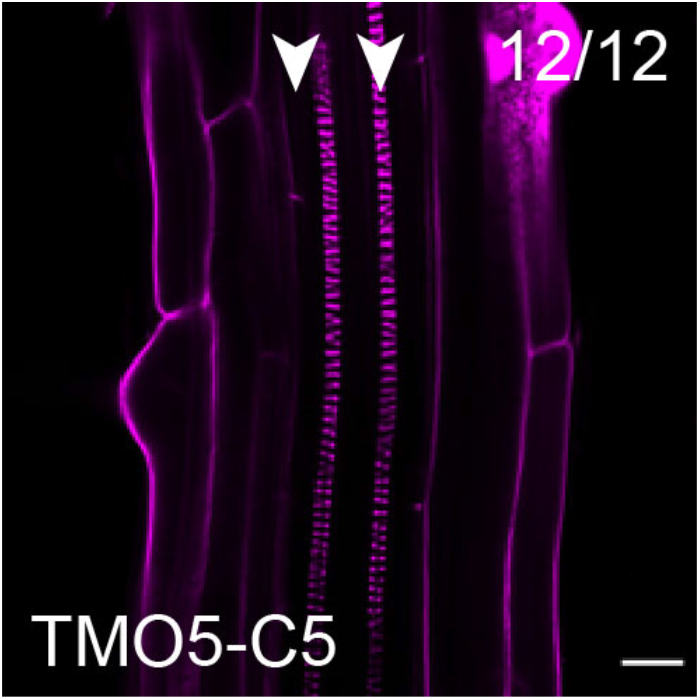
Domain swaps between MpTMO5 and KnTMO5, and complementation to a diarch vascular pattern phenotype in the Arabidopsis *tmo5 t5l1* mutant using the TMO5-C5 construct. Scale bar represents 20 µm; arrowheads indicate protoxylem poles. The number in the right top corner indicated how many show the phenotype out of the total number of individual T1 lines analyzed.

**Fig. S10.**
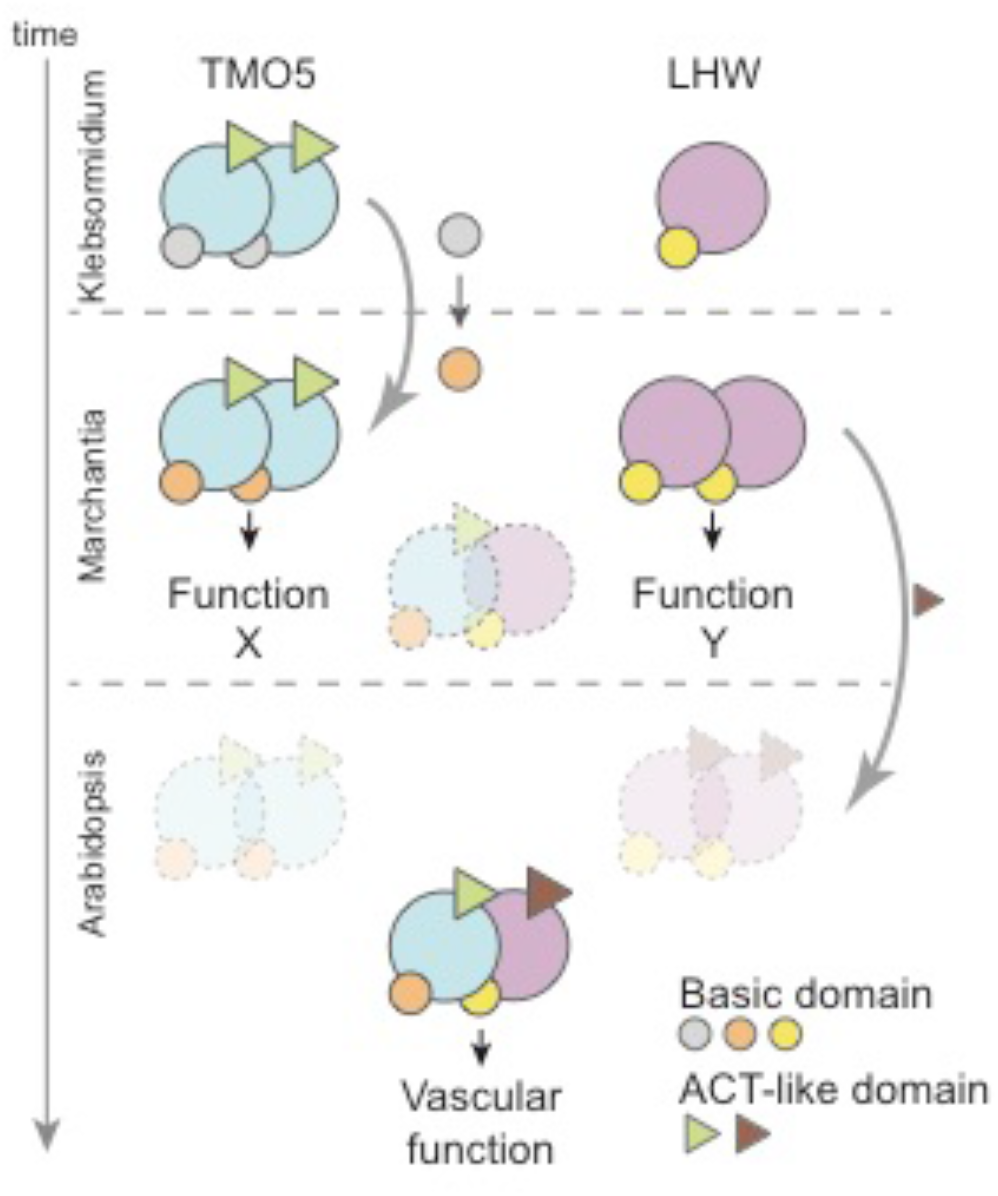
A two-step model for TMO5/LHW evolution. Evolutionary scenario depicting the innovations in TMO5 and LHW proteins towards the modern vascular function. The presumed ancestral TMO5 and LHW proteins (represented by orthologs in extant charophytes, including Klebsormidium) can form homodimers, but do not heterodimerize. At the origin of land plants, TMO5 (represented by the ortholog in the extant Bryophyte *Marchantia polymorpha*) gained its modern vascular function potential through mutations in its basic domain (orange circles). At the same time, TMO5 and LHW acquired the capacity to form heterodimers, yet control development independent from one another. A critical innovation to LHW in the ancestor of vascular plants (represented by the extant flowering plant Arabidopsis) favored obligate heterodimerization with TMO5 and completed the modern vascular function of this dimer. These changes included mutations in the ACT-like domain (brown triangle) together with another, yet unknown modification. Along with this switch to obligate heterodimerization, functions of each homodimer were lost.

## Notes

http://www.ncbi.nlm.nih.gov/bioproject/528622

