## Supplemental Dataset 1 for "Evolution of vascular plants through redeployment of ancient developmental regulators"

>Hs2006121-Huperzia_squarrosa|m.290

MGTSRLEDRLHPNGWILGSESAATGRLLSNVSPSGDPFGLMQEINSNWITLKALPKQEHH

PDLQANPPMLPIINPLTDWHRAAADSLAPGKHSEAKTPRTHAPFPYGLNRTFPLLCYNYG

DLINMPSMTPKEIMDAKALAASKSHSEAERRRRERINTHLATLRSRLPGTIKTDKASLLA

EVVHHVKELKRQAAEIAQISSLPTDADELQVDTDTSLGEDRVLIRASLCCDDRPGLLSDI

IRAIEEIKLHTTKIEIVALGGRVKNVIFMTGGNHASLKQEEVSATCVQDALRAAIDKSAS

NELTTSNFFANKRQRMGYHGSSQL

>Hs2006122-Huperzia_squarrosa|m.292

METSRLEEKMHPNGWILAHQSTATGRLLSNASPSGDPFGLMQEINSNWITFNALTKQQQH

HTELQANPLMLPISLTDKHRGPAESLAFGKVSEPKTPRLHVPFTYGMNRTFPLLCYNHGD

LISMPTMTSKEIMDAKALAASKSHSEAERRRRERINTHLATLRSRLPGTIKTDKASLLAE

VVHHVKDLKRKAAELAQISSLPTAADELQVDTDSSLGDDRVLIRASLCCDDRPGLLSDLI

RAIEEMKLHTTKIEIVALGGRVKNVIFMTAGNCPSLKQEGLSVTGVQDALRAVIEKSASS

ELTAFNFFTNKRQRMGHYGSSQV

>Hs2016510-Huperzia_squarrosa|m.296

MTLDLLCIINFLKALIDLCGSFWMQRSHSGPAYQFTTVARSLSYFRIPSLSVHCVSFAMG

SGGPLELGYMLANQISSFDMSGASNIVPEHFASSPDAQQEPTWNLSSQRYPDSTCNVFPQ

IKFQQQTPYQRSILHPSFNAVPIFSTFNHALESRCNSGILQIGSCRLNGLPPFPASHSLE

DKMFLTKQAGSSFPTAINEASMFFSRKHGNIGKIAKMTPQEIMEAKALAASKSHSEAERR

RRCRINTHLATLRSMLPSTTKTDKAYLLAEVIQHLKELKRQAAEISIVDPVPSDADEVSV

EIDPSFIGDCTLVRASLCCDDRPGLLSDLIRTLRCLNLQIVKAEMSMLGGRVKISILLTN

PLGTSEKQDASFMSSVQEALQAVLDKSCSNGIIPSNFEGKRQKVGYGNNDLIPF

>Hs2057546-Huperzia_squarrosa|m.293

MGASIDASVEARFPIFATADPANGILDSVRDSFGLMQGISSNCITIKKPSKQEPTTSFPP

NSCLHHLPLPQDSPSPRFSSFRPWRGAAAAESLAPEAYPSIIHYPLAYGLSRAFPLCYHH

GDLINMPNLMTPKEIMDAKALAASKSHSEAERRRRKRINSHLATLRSLLPATIKTDKASL

LAEVVHRVKDLKRQAAELAQISSLPSDADELQVDTNASLGEDGVLIRASLCCEDRPALFS

DLSRAIKEMNLRVTKVEIVTLGGRVRNVIFMTRGDRVKQEGLSVTSVQDTLKAVIKNSAS

NELTTTNFFDTKRPRLGPMALHKYTKSISFNSFDA

>Hl2010619-Huperzia_lucidula|m.300

RCLLPERLAKTIHLLGVDLSFMGTSIDASLEAKLPPTAWILAPESANGTFFSTSGDPFGL

MQGINSNWITLKNPPKQEPSSPLLKNSASISCLHHLPDPQANSPRFPNFNPLTDWHRAAA

ESLAPGNYSEAKAPRIHVPFPYGLNRTFPLLCYNHGDLINMSNMTPKEIMDAKALAASKS

HSEAERRRRERINTHLATLRSRLPGTIKTDKASLLAEVVHHVKELKRKAAELAQISSLPT

DADELQVDTDSSLGEDRVLIRASLCCDDRPGLLSDLIRAIEEIKLHTTKIEIVALGGRVK

NVIFMTGGNRASLKQEGLSATGVQDALRAVIEKSTSSELTTSNFFANKRQRMGHHGSSQV

>Hl2010620-Huperzia_lucidula|m.301

RCLLPERLAKTIHLLGVDLSFMGTSIDASLEAKLPPTAWILAPESANGTFFSTSGDPFGL

MQGINSNWITLKNPPKQEPSSPLLKNSASISCLHHLPDPQANSPRFPNFNXWHRAAAESL

APGNYSEAKAPRIHVPFPYGLNRTFPLLCYNHGDLINMSNMTPKEIMDAKALAASKSHSE

AERRRRERINTHLATLRSLLPGTIKTDKASLLAEVVHRVKELKRQAAELAQISSLPSDAD

ELQVDADASLGEDRVLIRASLCCDDRPALFSDLSKAIKEIKLRATKIQIVALGGRVKNVI

SMTCGDRVSFKQEGLPVTSVQDTLRAVIEKSASNELTTTNFFGTKRQRLGHYGSSQV

>Hl2023097-Huperzia_lucidula|m.298

MAISHGGSVELAFPFLDPNLNSVFGVTTNVLPDFSSEVETRRAPWLAPFSSSVGSMHKIY

PELDSQLHSVKRDSTTCLLPMFAASDSSYTPINDCHINPLFMPFGNQGNRLRASPVPQSY

VETLLFRNREGKPSFPHALSGGAPLFISGKHGDIMAKMTPQEIMNAKALAASKSHSEAER

RRRERINTHLATLRSLLPNTTKTDKASLLAEVINHVKDLKRKAAEIAEWAPIPSDTDELR

VDTISSDGAEDGVLIKVSLCCDDHPTLLSDLTKTLRNLKLQTVKAEIASLGGRMKNVILL

TSQDTTSGKPDESLLENVHEALREMLERSGSNQSSPAVFGSGSISKRQKLGF

>Hse2005550-Huperzia_selago|m.303

MSNMTPKEIMDAKALAASKSHSEAERRRRERINTHLATLRSLLPGTIKTDKASLLAEVVH

RVKELKRQAAELAQISSLPTDADELQVDTDSSLGEDRVLIRASLCCDDRPGLLSDLIRAI

EEIKLHTTKIEIVALGGRVKNVIFMTGGNRASLKQEGLSATGVQDALRAVIEKSASSELT

TSNFFANKRQRMGHHGSSQV

>Hse2022097-Huperzia_selago|m.302

IYPELDSQLHSVKRDSTTCLLPMFAASDSSYTPINDCHINPLFMPFGNQGNRLRASPVPQ

SYVETLLFRNREGKPSFPHALSGGAPLFISGKHGDIMAKMTPQEIMNAKALAASKSHSEA

ERRRRERINTHLATLRSLLPNTTKTDKASLLAEVINHVKDLKRKAAEIAEWAPIPSDTDE

LRVDTILSDGAEDGVLIKVSLCCDDHPTLLSDLTKTLRNLKLQTVKAEIATLGGRMKNVI

LLTSQDTP

>Sw2181545-Selaginella_wallacei|m.305

MGGHSDVGFPFAEASLGISYAGFSKLKQELPPPLGPPSASGMASSRPPWALPPMMGPSLG

SSAATTEHHGLPLLGHGGRSFSELHSAPVSSSSSFGVPYEYKPSPLAAASSAYGGFKDRG

GAGGGGGARHQHYPLQYNGGSLVLDHDRGELINLSKMTPQEILDAKALAASKSHSEAERR

RRERINNHLNTLRSLLPSTTKTDKASLLAEVILHVKELKRQAAEIAEGGPVPSDVDEIKV

DADSSSTDGNLVMKASLCCDDRPDLLSDLTKVLKTLKLKTLKAEIATLGGRVKNVIVIGK

DATAVGEGADDEMAGSSSSSSGQQDRPSVNCVQEALRAVIERSSNDASASGYSSGGKRQK

MNHHHESSSSGY

>Sw2181894-Selaginella_wallacei|m.315

MDSSSCFSKAMAASKSHSEAERRRRRRINNHLETLRQLLPGTAAKVGLVVFVLLKPTTHQ

LVLCSCVCLSAACRQGISAGRSHWKDQGAQAESSRDLRARAGSQRSR

>Swi2008637-Selaginella_willdenowii|m.321

SRCWNPERVHAGRFVFLVLSWKLVLECNMNPWSLREKGCEDLPSSSYPKSAFQVTGLEPL

FASPSLANATSQDQLNMKNACSSSSSQLQDAKALAASKSHSEAERRRRERINNHLSTLRT

LLPNTAKVESRISS

>Swi2014032-Selaginella_willdenowii|m.317

MGGPSTNGLDVGFPFGESPTALGMSYTGFSKLKQELPAGLVTPSPSTVHASTSRPPWALP

PTLAGASVVPPKLASLSEFGSHPQHFMKQDHGTATTATTTNSSPRFHELPFSYDSQVNNL

YKSSLGSSASAFGSGVNYNKERSSGRPYPLQYNGGSLVLDHHRGELINLSKMTPQEILDA

KALAASKSHSEAERRRRERINNHLNTLRGLLPSTTKTDKASLLAEVIEHVKDLKRQAAEI

AEGGPVPTDVDELKVDTDPSSSDGNFVLKASLCCEDRPDLLTDLTKALKTLKLKTLKAEI

ATLGGRVKNVLLIGKDNSAVAEAGGEQERPTVSCVQDALRAVIERSADLSSPSYGSKRQK

MNSSMH

>Sse2039830-Selaginella_selaginoides|m.324

IDSSPYAGFGKLKQELPGTAAADIVGSSSRPPWMLPPSFAPTSKPFADGQYSGNASGRFP

DLNPAPFSLPFDNNQVLYKSSLMDYKDRGRHYPLQYNGGSLVLDQHRGELINLSRMTQQE

IQDAKQLAASKSHSEAERRRRERINRHLATLRSLLPS

>Sa2002423-Selaginella_apoda|m.334

MLRLSPAQALVDSEKALAASRSHSEAERRRRERINKHLSTLRTLLPGTAKTDKASLLAEV

IDRIKELKEQVAEISQLGPVPSDADELNVDVLDPPDEQGKVLIKASICCADRPSLIRDLV

RTLKSLHLRTLRAEMATMEGRTKNVFVMTIKEDAELLEPTLACVEEALKSVMEESAPEEN

PDDRESFPASC

>Sa2004992-Selaginella_apoda|m.325

MGGPSSGSNVDVGYGPFGGGGGGAEASPSGFGGGGGGISYNTGGGGGGGGNFSKLKQELV

TSRHPPWTIPPPLASSTVPKPSSSFPDLPAAHHHQFVANQQPDHRHHHHHHHQHHHQPHV

SLLGSGSGGGTSTTAATAGRFHDLHGSSSPFSSLPYDLYKSGALATSSSPAGYGVGLGNR

SYKDRGRQHYPLQYNGGSLVLDHNRGELINLSKMTPQEIIDAKALAASKSHSEAERRRRE

RINNHLNTLRGLLPSTTKTDKASLLAEVIEHVKDLKRQAAEIAEGGPVPTDVDELKVDTD

ASSSDGNFVLKASLCCEDRPDLLTDLTKALRTLKLKTLKAEIATLGGRVKNVILIGKDDD

NSGHHQGGGDGGGGESSAAGGGNSTGDRPSVNCVQEALRAVIERSGELSAPGYGSSGKRQ

KMSSAAASSPSSY

>Hse2010139-Huperzia_selago-A|m.347

MAISHGGSVELAFPFLDPNLNSVFGVTTNVLPDFSSEVETRRAPWLAPFSSSVGSMHKIY

PELDSQLHSVKRDSTTCLLPMFAASDSSYTPINDCHINPLFMPFGNQGNRLRASPVPQSY

VETLLFRNREGKPSFPHALSGGAPLFISGKHGDIMAKMTPQEIMNAKALAASKSHSEAER

RRRERINTHLATLRSLLPNTTKTDKASLLAEVINHVKDLKRKAAEIAEWAPIPSDTDELR

VDTILSDGAEDGVLIKVSLCCDDHPTLLSDLTKTLRNLKLQTVKAEIATLGGRMKNVILL

TSQDTPSGKPDESLLEXTP

>It2089681-Isoetes_tegetiformans|m.352

DSAFSCRMPRILGASNSAGGKELFNFSKMATSQEMLEAKALAASKSHSEAERRRRERINS

HLATLRTLLPSSTKTDKASLLAEVIDYVKELKRQAAEIAEGGPVPTDVDELNVDTDSSIE

EDKVLIRASLCCEDRPDLLSDLIRTLRSLKLQTVRAEMASHGGRIKNV

>It2099829-Isoetes_tegetiformans|m.349

MASSAGNGVEVNFPFADSNSLVHGGYGGSGSGGNSFKLLQDYNTPASSVQPPWMVPMPST

ASMSKALADLAANQQQMPKEQPLSSFLASSPMPSFGRFAPTSHVPPFPLSLEAQQSFRHT

SRGYGGQTRQFPLQYVGNGGSLVLDRARGELINLSKMTPQEILDAKAIAASKSHSEAERR

RRERINNHLATLRSLLPSTTKTDKASLLAEVIEHVKELKRQAAEIAEGGPVPTDVDELKV

DSEAGDGNVVLRASLCCDDRPNLLSDLTKAIKSLKLKTLKAEFATLGGRVKNVILLTNDP

SGICTSSTSSSSSDNSEADQSIPSASSVQEALRAVIERSAELTPANGYGGNKRQKLVNDF

PSTY

>Ld2003154-Lycopodium_deuterodensum|m.356

MAASEGASLEQGLASSGWNLAPEPAKSKFLSNSSPSCAMSFGGSSSGLMQGYYTGWSVEQ

YPPKQEPGSPLPNFSAHIPSTHQFPDVHANRPMLPILYHSNDWRRATADSLSFGSNSEGK

ATKKHDLFHFQMDRGLPVVWGNHGDIITMSKMIPKEILNAKALAASKSHSEAERRRRERI

NTHLGTLRSLIPSTTKSDKASLLAEVVHHVKRLKRQAAEIAQVCPVPTAIDELQVDTDFS

LGGDRVFIRASLCCDDRPDLLSDLIRASKDLELQTVKAEIFMLGGRVKNVILMTRGDGTS

CKQEGPLVTSVQEALREVMEKSGSNELSPSNFFSNKRQRMGYHSSSRV

>Ld2008397-Lycopodium_deuterodensum|m.355

MGTSGGASLEHSLPSNGWILTPDVANDKIFPNFSASCVMPFGGSFGLMQKPYSDWSSLQH

RPKQEPVSSLPKFSAPISSFHQLPDVHVNPPMLPFISQFNDWHRGGADSLTFGSNLEAKA

PGKRDLFPYVTNRTLPVLCCNHGEFMNMSKMSPKEIMDAKALAASKSHSEAERRRRERIN

SHLATLRSLLPSTTKTDKASLLAEVVHHVKELKRQAAEIAEISPVPTDVDELQVDIDSSL

GEGRVLIRATLCCDDRPGLLSDLTSALRDLKLQTVKAEIATLGGRVKNVILMTRGDRASC

KQEGPSLTSVQEALKVVMEKSSNELSPSNFFSNKRQRTGYHCSSQV

>Ld2009148-Lycopodium_deuterodensum|m.354

LLDPSLSSAFGGTANLLPELPSGGGIQRADLRSTPFSSSVGSVLKAFPGMESQQYFSKQS

SSTASLFPLSTAPVTSLSSIPDFHVNPLLLPFQNQANGLRMNRSPLIPQNYVENLLFRDR

GGQSSLPACAVSGASLLLHGKHGDSINMANFSPQEVMDAKALAASKSHSEAERRRRERIN

THLVTLRSLLPNTTKTDKASLLAEVIQHVKELKRKTAEIADRDPIPSDVDELRVDTDTSY

DEDRVLIKASLCCDDRPGLLSDLIKTLRNLKLQTVKAEISTIGGRVKNVILLTSEDAASV

EEDGRSLITNVQEALRAVMDRSGSSELSSLGFGSNKRQKLGS

>Ld2011715-Lycopodium_deuterodensum|m.353

MATSHGGAVDLGFPFLDPSSAFGGTTNLSSDLPSGVGSQRPPWLTPFNSSTGSTRKVFPG

LNPQDHFTKRDSPACLLPSFAAPISALSQIPDFHVNPLLLPFQNHANALRMTRASTLPQS

YGDNILFRDRVGQSSLPFTMSGASLLLSGKHGELINMSKLTPQEIMDAKALAASKSHSEA

ERRRRERINTHLATLRGLLPNTTKTDKASLLAEVIQHVKELKRQAAEIAEGGPVPSDVDE

LRVDTDSSYSKDRILIKASICCDDRPGLLSDIIRTLRNLKLQTVKAEIATLGGRVKNVIF

LTSEDTISGTKDGPLVTSVQEALRAVMERSGSNELSPSGFGSNKRQRLGSQDFTPF

>Is2070897-Isoetes_sp.|m.360

MMGSQGRIGWQMGSADPEASRNNKGGALEMPWNMAADVASSNFLLQKALMQQQNHHMQQN

NLQQDQLYNPTSRPGIPSYMPAGPLQFSFDHRAPKPWPNPASNFYGADDSAFSCRMPRIL

GASNSAGGKELFNFSKMATSQEMLEAKALAASKSHSEAERRRRERINSHLATLRTLLPSS

TKTDKASLLAEVIDYVKELKRQAAEIA

>Is2075584-Isoetes_sp.|m.357

MASSAGNGVEVNFPFADSNSLVHGGYGGSGSGGNGFKLLQDYNTPASSVQPPWMVPMPST

ASMSKALADLAANQQQMPKEQPLSSFLASSPMPSFGRFASTSHVPPFPLSLEVQQGFRHT

NRGYGGQTRQFPLQYVGNGGSLVLDRARGELINLSKMTPQEILDAKAIAASKSHSEAERR

RRERINNHLATLRSLLPSTTKTDKASLLAEVIEHVKELKRQAAEIAEGGPVPTDVDELKV

DSEAGEGNVVLRASLCCDDRPNLLSDLTKAIKSLKLKTLKAEFATLGGRVKNVILLTNDP

SGICTSSTSSSSSDNSEADQSIPSASSVQEALRAVIERSAELTPANGYGGNKRQKLVNDF

PSTY

>La2002693-Lycopodiella_apressa|m.366

AAFGGRALTYLSDSSPGVMGVERVDPWLSSFVSSSASLTQPEILPVLESQQHNFYTKRPS

KAPCDHFLPAFAVPISSSTPVHGSFNGMDHLQLPASFLKAHANNFHLPPSVSIHVPQNYR

DLNVVSSQDMKGEGQSSLPNYVVNGTSLLIKTGNHGDKRSMQEMAKLTSREIMEAKALAA

SKSHSEAERRRRERINTHLATLRRLLPNTAKTDKASLLGEVIQHVRELKRQTAEIAEGTL

LPSDTDELRVDTELIHDENRVLIKAFICCDDRPGLLSDIVKAVRNLKLQTVKAEIYTLEG

RVKSVILLRREDTGTVLVEENAGPLLTHVQEALRTVMAKAEYSGEFSSPFGLGSNKRQKV

GSYVKISR

>La2002694-Lycopodiella_apressa|m.364

AAFGGRALTYLSDSSPGVMGVERVDPWLSSFVSSSASLTQPEILPVLESQQHNFYTKRPS

KAPCDHFLPAFAVPISSSTPVHGSFNGMDHLQLPASFLKAHANNFHLPPSVSIHVPQNYR

DLNVVSSQDMKGEGQSSLPNYVVNGTSLLIKTGNHGDKRSMQEMAKLTSREIMEAKALAA

SKSHSEAERRRRERINTHLATLRRLLPNTAKTDKASLLGEVIQHVRELKRQTAEIAEGTL

LPSDTDELRVDTELIHDENRVLIKAFICCDDRPGLLSDIVKAVRNLKLQTVKAEMSTLGG

RVKNVIVLSSEDRTSTTLLEEX

>La2009267-Lycopodiella_apressa|m.367

SPAMPFGGSSGFMQKLPSDWGAFQQTPKQEPVSSLQKFSASISSFSHLPDVHANAAMPPF

FGQMNGWSRARTGPLISGSNSEANNNSGRRDLISLTSNRTLQVLCCNSEEFINASTMTQK

EITDAKALAASKSHSEAERRRRQRINTHMANLRSLLPSTTKTDKASLLAEVVHHLRELKR

QAADIAQVSQVPTDVDELQVDIGTSFGEEKMLIKASLCCDDRPGLLLDTKNALRDLKLQI

VKAEIATLGGRVKTVILMSRMGPAPCTQEGPSLISVQEALKAVMEKSASNELAPYSYFNN

KRQKTTYHCPQGLGQFC

>La2011251-Lycopodiella_apressa|m.365

EFMNISKMTTKEIVDAKASAASRSHSEAERRRRQRINTHMATLRTLLPSTTKTDKASLLA

EVVHHVKDLKRQAAEIAQISPVPTDIDELHVEFDSSFGEGRMLIKASFCCDDRPSLFSDL

TKGLKDLKLQTVKAEIATLGGRIKSVMLMTRVDQVSCSQEGPSLTGVQEALRAVMEKPPS

NELALSNFFSEKRQRTTYYCA

>La2013318-Lycopodiella_apressa|m.368

KPFSSFSPRAALAFGGSGLMQEPHTNWSSLQHTLKQEPVSSPNKLSAPISSRHQRPDAPA

NAPVTPLFGQIDDWSKTGTDPFVFGSNSDIEVSGTCDLSSYASRRTLPVICCNAGVFRNV

PKMTSKEIADAKALAASKSHSEAERRRRLRINTHMSTLRTLVPSTTKTDKASLLAEVVHH

LRELKQQAAENAQLSXAVDELLVDTDASVGEDSVLIKASFCCDDRPGLLPDITKALRDLK

LRIVKTEIVTLGGRVKIVMLVTRVDHVSCKEEGPSLTSVQEALKAVMEKSCSNEIVLPNF

FSYKRQRTTYHYS

>Pc2009292-Pseudolycopodiella_caroliniana|m.370

MGTFEVASLEHTNPPNCWILAPDMAKGKAFSSFSPSPAMPFGGSSGLMQKPHLGWGALQQ

TLKQEAFSSLSEFSPPISSFPLFPEVHASAAMLPLFGQISSWSSGGTDPFCAGNNSEPKT

SGKRDLPRLASNGNLPVPCCSSGEFINVSRMTPKEIMDAKALAASKSHSEAERRRRQRIN

THMAKLRSLLPSTTKTDKASLLAEVVHHVRELKRQAAEIAKISPIPTDFDELNVDIGSSS

CEDKALVKASLCCDDRSGLLSDITKSLKDLELQIVKCEIATLSGRVKIAMLICRMDHAPC

KQEGSSLNSVREALKAVMEKSASNELAPYSFFNNKGQRISYHCS

>Pc2014284-Pseudolycopodiella_caroliniana|m.372

MGAFGGASLEHSYPRNGWMLAPDKPFSNFSPSAMLFGGSSGLIQKPHSDCISLQQALKQE

PVSSLHQYSAPISSXLHQLPNIHESAPILPFFGPVNDWSRAGNDPIVFGRNSGTKVPGKN

DLLSYSSSRTLPVLYCDRGEFMDMSKMTPKEIMDAKALAASKSHSEAERRRRLRINTHMT

TLRSLLPSTIKTDKASLLAEVVHHVKELKRHAAEIAQISPVPSDVDELDVDMDSSLGEDS

VIMKASLCCDDRPGLLLDIAKALKDLKLQPVKAEIATLGGRVKHVMLIRRLDCGSCKQEG

PSLTNVQEALRAVMEKPPSNEIAPPNFFSDKRQRTTYYCA

>Pc2014285-Pseudolycopodiella_caroliniana|m.369

MAGLLLPMFRMISLFLAFLPXVLPVGGSRLIQKNHSDWSSRKHPLKQEPVSAPYKLGAPI

SIHQPPDVRANVPTVPVFGQINDWRRAGTDPFAFGSNSETKVSGKFDLSSYASNRSLPVI

FCNSMGSMDIPKMTPKEIMDAKALAASKSHSEAERRRRQRINTHMATLRTLVPSTTKTDK

ASLLAEVVHHLRELKQQATKIAQISPVPTEIDELHVDFDSASSEDKLLLKASLCCDDRDG

LLSDIAKALRDLKLQIVKAEIVTVGGRVKNVMLMTRADHLSCKQEGPSLTSVQEALKTAM

EKSPSDELSPSNFLSYKRHRTTYHCF

>Pc2018166-Pseudolycopodiella_caroliniana|m.374

DANILPASSLVGTQRPPCSTPFNSPAASMRVAFPGLDPEMHFAKRDSIACNFLPPFASSI

SSLSRIPNFHVNPLPLPFQNQANNVRLTGTSTCPAYAENFPFRCNGVSASMPNSAMAGPS

VFLNGKHGEIIDMTKLTPQEIMDAKALAASKSHSEAERRRRERINGHLTTLRGLLPNTTK

TDKASLLAEVITHVKDLKRQVAEIEDEGPVPSDVDELRVDKELSFEKDRVLIKASLCCDD

RPSLISDI

>Pc2071655-Pseudolycopodiella_caroliniana|m.376

LKDKGGGQSSLPNYLVNGASFLLKAGINLGDRPMQEIAKLTSQEMMEAKALAASKSHSEA

ERRRRERINTHLASLRRLLPTTTPKTDKASLLGEVVEHVRELK

>Dd2010310-Diphasiastrum_digitatum|m.378

MGTSGGASLEHSFQSNGWILAPDVANDKLFPNFSPSCVMPFGGSFGLMQKPYFDWSSQQH

IPKQEPVSSLPKFSTPISSFHQPSDVHVNPPMLPFLGQMNDWHRAGADSLGFGNNSEAKA

SGKRDLFPYATNRTLPVLCCNHGEFMNMSKMTPKEIMDAKALAASKSHSEAERRRRERIN

SHLATLRSLLPSTTKTDKASLLAEVVHHVKELKRQAAEIAEMSPVPTDVDELQVDIDSSL

GEDRVLIRASLCCDDRPGLLSDLTRXEI

>Dd2068750-Diphasiastrum_digitatum|m.377

MTTSGGGSLELGFPSSGWNLAPEPVEDKLFSNSSPSCVMPYGGSSFGLMQGCYSGWSVLQ

HPPKQEPASLLPNFSALISSLHQLPDLHANPPMLPILCRSNDWRRAAADSLPLKSNSEEK

APRKHGLFHYQMGRTLPAVWGNYGELIGMSKMIPKEILNAKALAASKSHSEAERRRRERI

NTHLGTLRSLIPSTAKTDKASLLAEVVHCIKKLKRQAAEIAQVCPIPTDIDELQVDTDSS

LGGDRVLIKASLCCDDRPDLLSDLIRASRDLKLQTVKAEIFMLGGRVKNVILLTRGDGTS

CKQEGPSVSSVQEALREVMEKSGLNELSPSNFLSNKRQRMGYHSSSKV

>Do2012233-Dendrolycopodium_obscurum|m.380

SKSEKTAPRKHDLFHYRMDRTLPVVWGNELITMSKMIPKDILKAKALAASKSHSEAERRR

RERINTHLGTLRSLIPSTTKTDKASLLAEVVHHVKKLKQQAAEIAQVCSVPTDIDELQVD

TDSSLGGDRVLIRASLCCDDRPDLLSDLIRASKDLKLQTVKAEIFMLGGRVKNVILMTHG

DGTSCKQEGPSVTSVQEALREVMEKSGSNELSSSNIFSNKRQRTGYHSSSKV

>Do2012946-Dendrolycopodium_obscurum|m.379

MGTSGGASLEHSFPSNGWILAPDVANDKVFSNFSASCVMPFGGSFGFMQRPYSDWSSLQH

PPKQEPVSSLNKFSAPISSFHQRPDVHVNPPMLPFLSQINEWHRAGADSLTFGNNTEAKA

PGKRDLFPYATNGTLPVLCCNHGEFMNMSKMTPKEIMDAKALAASKSHSEAERRRRERIN

SHLATLRSLLPSTTKTDKASLLAEVVHHVKELKRQAAEIAEISPVPTDVDELQVDIDSSL

GEDRVLIRASLCCDDRPGLLSDLTRALRDLKLQTVKAEIATLGGRVKNVILMTRGDHASC

KQEGPSLTSVQEALKTVMEKSSNELSPSNFFSNKRQRTGYHCSSQV

>Hse2008700-YHZW-Huperzia_selago-2_samples_combined|m.383

MGTSIDASLEAKLPPTAWILAPESANGTFFSTSGDPFGLMQGINSNWITLKNPPKQEPSS

PLLKNSASISCLHHLPDPQANSPRFPNFNPLTDWHRAAAESLAPGNYSEAKAPRIHVPFP

YGLNRTFPLLCYNHGDLINMSNMTPKEIMDAKALAASKSHSEAERRRRERINTHLATLRS

LLPGTIKTDKASLLAEVVHRVKELKRQAAELAQI

>Hse2015752-YHZW-Huperzia_selago-2_samples_combined|m.381

MAISHGGSVELAFPFLDPNLNSVFGVTTNVLPDFSSEVETRRAPWLAPFSSSVGSMHKIY

PELDSQLHSVKRDSTTCLLPMFAASDSSYTPINDCHINPLFMPFGNQGNRLRASPVPQSY

VETLLFRNREGKPSFPHALSGGAPLFISGKHGDIMAKMTPQEIMNAKALAASKSHSEAER

RRRERINTHLATLRSLLPNTTKTDKASLLAEVINHVKDLKRKAAEIAEWAPIPSDTDELR

VDTILSDGAEDGVLIKVSLCCDDHPTLLSDLTKTLRNLKLQTVKAEIATLGGRMKNVIXX

XXXXXXXXSLLENVHEALREMLERSGSNQSSPAVFGSGSISKRQKLGF

>Sk2006839-Selaginella_kraussiana|m.395

CLPLAAAMYDVYLSHCWVAGTSVKIADIVANIISIEARHLLPRLDFHSGTSVSKLVSQVT

WNNKGIPTGRRREFLNLSPPDGICLLEKPXAASKSHREAERRRRGRINEHLATLNKLLPG

TIKADKASILAGVIQHIKRLKEQVAEISTFWPVPSAHEASQGVHLLQ

>Sk2007004-Selaginella_kraussiana|m.392

MGSPYFPTFPINTALDDDDSSELALRRACVDAKALAASKSHSEAERRRRGRINDHLATLK

ALLPNPTKVDKASILAEVIERIKELKQQVAEISEFCPVPSEVDELNVHIDRPAKLLKASI

CCNDRPGLFGDLVKTLKTLGLETVRAEMATLDGRTKNVFLMTTREGQPLLDEETLACVQE

ALKVVMEQPEETSRLKGGKLIPSKSSK

>Sk2042682-Selaginella_kraussiana|m.386

MGNNGVEVGFSAFGEASAANLGMTYSGFSKLKQELPSSSASSSRPPWAIPPLGPSTSGSS

SSGATVAAVAPKSFADLTPSHSHYQNVVKHHQQQQQQQQDHHHHHPLLGGSPSARSFAEL

HAAAAAAASPFALPYETQSRSSGGGLYKTSSSPSAAYNGGSRSRHYPLQFTGGSLVLDHH

RGELINMSKMTQQEILDAKALAASKSHSEAERRRRERINNHLNTLRSLLPSTTKTDKASL

LAEVIEHVKELKRQAAEIAEGGPVPSDVDELKVDTDSSSTDGNLVMKASLCCDDRPDLLT

DLTKVLKTLKLKTLKAEIATLGGRVKNVIVIGKDAAGAGGGATAGAGPCGDNEPGSSQDS

QRPSVNCVQEALRAVIERSNELSASGYSGSKRQKMNHHHEGASSY

>Sac2010137-Selaginella_acanthonota|m.397

MGGHSDVGFPFGEASLGISYAGFSKLKQELPPPLGPPSASVATGMASSRPPWALAPMMGP

ASLGSSAATTEHHGLPLLGHGGRSFSELHSTPVASSSSSFGVPYDASSMYKPSPLAAASS

AYGSLKERGGGGGARHQHYPLQYNGGSLVLDHHRGELINLSKMTPQEILDAKALAASKSH

SEAERRRRERINNHLNTLRSLLPSTTKTDKASLLAEVIQHVKELKRQAAEIAEGGPVPSD

VDEIKVDADSSSTDGNVVMKASLCCDDRPDLLSDLTKVLKTLKLKTLKAEIATLGGRVKN

VIVIGKDAAAVGEGADEMAGAGSGSSSSSGQQDRPSVNCVQEALRAVIERSSNDASASGY

SSGGKRQKMSHHHESSPSGY

>Pd2002811-Phylloglossum_drummondii|m.403

MNRSLPLCYGHGDIFSIPSMTQKEIMDAKALAASKSHSEAERRRRERINTHLTTLRSLLP

ATIKTDKASLLSEVVHQVKELKRQADALPQISSLQPTDTDELQVDVESSLEDTVCMRALL

CCDDRPGLLADLIRAIKEIKLNVTKIEFVSLGGRVKHVILMTRREHASSKQEGPSVAGVQ

DALRAVIEKSAPCELAPNFFSNKRQRMGRHGNFLQV

>Pd2113908-Phylloglossum_drummondii|m.405

KMSKMTPHEIIEAKALAASKSHSEAERRRRCRINTHLETLRRMLPSSTKTDKASLLGEVI

QHLMELKSYATEISDVDPVPGDINEVIVEIDPAMTGESLTLRVSLCCDDRPG

>Pd2119379-Phylloglossum_drummondii|m.404

NQEGSPSFPTVTKNGASVWINGKHRNFSKTSQEIMEAKALAASKSHSEAERRRRCRINAH

LATLRSMLPSTTKTDKAYLLAEVIQNLKELKRRATEISIVDFVPSDEDEVSVETVPGRLV

GDCELLRASLSCDDRPGILSDLNR

>Sst2011330-Selaginella_stauntoniana|m.406

MGGPSTNNVDVGFPFVDSSGLGMTYAGFSKLKQELPVSLVTAAGHSSSRPPWTIPPLGPG

CTAAQATSTAAFPELAAASSQHGQFVTKQDHVSSLLHASGGGGGSSAPGRFHDLHGSSSP

FSTLPYDLYKSGALASSSSSGYGVGLSYKDRGRHHYPLQYNGGSLVLDHNRSELINLSKM

TPQEIIDAKALAASKSHSEAERRRRERINNHLNTLRGLLPSTTKTDKASLLAEVIEHVKD

LKRQAAEIAEGGPVPTDVDELKVDTDASSSDGNFVLKASLCCEDRPDLLSDLTKALRTLK

LRTLKAEIATLGGRVKNVILIGKDHSDEQGGAAMESSSDGXRPRGTEGSDRAI

>Sle2008762-Selaginella_lepidophylla|m.268

MGGPSSSSVDVGYGPYGGGGEASSSGFGGGGGGGITYSAGGGNFSKLKQELIPAGLVTSR

HPPWTIPPPLASSTVPAPKTTIMTSPAATSFPDLAAAHQHHHHHQFVAKPPDHHHLLGGS

GGSSTTAATAGRFHDLHGSSSPFSTLPYDLYKSGALATAASSSPSGYGVGLGSRSSYKDR

GRQHYPLQYNGGSLVLDHNRGELINLSKMTPQEIIDAKALAASKSHSEAERRRRERINNH

LNTLRGLLPSTTKTDKASLLAEVIEHVKDLKRQAAEIAEGGPVPTDVDELKVDTDASSSD

GNFVLKASLCCEDRPDLLTDLTKALRTLKLKTLKAEIATLGGRVKNVILIGKDNSENHQG

GAAAVESSGDAGGGGGGGDRPSVSCVQEALRAVIERSGELSAPGYGSGKRQKMSSAFSSS

SY

>Hm2005726-Huperzia_myrisinites|m.276

MRRSSNREAKPLFSHAPSWGGPLVISGRHGEIMAKMTPKEIFNAKALAASKSHSEAERRR

RERINTHLATLRSLLPSTTRNDKASLLAEVIHHVKELKRKAADIAEWTPIPSDTDELRVD

TIFSEGAEDGVLMKVSLCCDDHPTLLSDLTKTLRNLKLQTVKSEIVTLGGRMKIVLLLSR

QDKTSREPDESLLENVREALREMLERSVSNQFSPAVHGSISKRQKLCFKD

>Hm2005727-Huperzia_myrisinites|m.282

MRRSSNREAKPLFSHAPSWGGPLVISGRHGEIMAKMTPKEIFNAKALAASKSHSEAERRR

RERINTHLATLRSLLPSTTRNDKASLLAEAIHHVKELKRKAAEIA

>Hm2010056-Huperzia_myrisinites|m.274

MGTSRLEDRLHPNGWILGPESAATGRLLSNVSPSGDPFGLMQEINSNWITLKALPKQEHH

PDLQANPPMLPIINPLTDWHRAAADSLAPGKHSEAKTPRTHVPFSYGLNRTFPLLCYNYG

DLINMPRMTPKEIMDAKALAASKSHSEAERRRRERINTHLATLRSRLPGTIKTDKASLLA

EVVHHVKELKRQAAEIAQISSLPTDADELQVDTDTSLGEDRVLIRASLCCDDRPGLLSDI

IRAIEEIKLHTTKIEIVALGGRVKNVIFMTGGNHASLKQEEVSATCVQDALRAAIEKSAS

NELTTSNFFANKRQRMGYHGSSQL

>Hm2010057-Huperzia_myrisinites|m.284

METSRLEEKMHPNGWILAHQSAATGRLLSNASPSGDPFGLMQEINSNWISFNALPKQQQH

HTDIQANPLMLPISLTDKHRAPAESLAFGKVSEPKTPRLHVPFAYGINRTFPLLCYNHGD

LISMPTMTSKEIMDAKAQAASKSHSEAERRRRERINTHLATLRSRLPGTIKTDKASLLAE

VVHHVKDLKRKAAELAQISSLPTAADELQVDTDSSLGDDRVLIRASLCCDDRPG

>Hm2010058-Huperzia_myrisinites|m.275

MGTSRLEDRLHPNGWILGPESAATGRLLSNVSPSGDPFGLMQEINSNWITLKALPKQEHH

PDLQANPPMLPIINPLTDWHRAAADSLAPGKHSEAKTPRTHVPFSYGLNRTFPLLCYNYG

DLINMPRMTPKEIMDAKALAASKSHSEAERRRRERINTHLATLRSRLPGTIKTDKASLLA

EVVHHVKELKRQAAEIAQISSLPTDADELQVDTDTSLGEDRVLIRASLCCDDRPGLLSDI

IRAIEEIKLHTTKIEIVALGGRVKNVIFMTGGNHASLKQEEVSATCVQDALRAAIEKSAS

NELTTSNFFANKRQRMGYHGSSQL

>Hm2052043-Huperzia_myrisinites|m.283

RRCRINTHLATLRSMLPSTTKTDKAYLLAEVIQHLKELKRLAAEISIIDPVPSDTDEVSV

ENDPSFIGDCTLVRASLCCDDRPGIISDLIRTLRCLNLQIIKAEMSMLGGRVKISIQLTN

ALGTSAKQD

>Hm2060638-Huperzia_myrisinites|m.278

MGASIDASVEARYPIFAAADRTNGSLDSVRDSSGLMQGISSNCITIKKPSKQEPTTSFPP

PIPCFHHLPLPQASPSPRFSSFRPWRGAAAAESLPPEAYPSIIHSPFAYGLSRAFPLCYH

HGDLIKIPNLMTPKEIMDAKALAASKSHSEAERRRRKRINSHLATLRSFLPATIKTDKAS

LLAEVVHRVKDLKRQAAELAQISSLPSDADELQVDTNASLGEDGVLIRASLCCEDRPALF

SDLSRAIKEMNLHVTKVEIVTLGGRVRNVIYMTGGDQVKQEGLSVTSVQDTLKAVIKNST

SNELTTTNFFDTKRPRLGSMALHKYTKSISFNSFDA

>Lan2015098-Lycopodium_annotinum|m.289

IDLLAKYLNLFCRQGISMGEASVSNTVPLDWLLPNPNSKAELVQSQKLVEEPLSLVQNIQ

TFRWPIPFNENTPSFIPTTLSDCQIPRHLFGHPESISSDISTQIPFIHFPTFTLDPRFVP

YENSSFTAASESAQCHGSTFDLRARXXXGAHYVPIGFSGRSVFVNHGELLNIPKINSQDL

LDAKALAASKSHSEAERRRRERINTHLATLRSLLPTTTKTDKASLLAEVINQVKELKRKA

ATIAEGSPVPS

>Lan2017295-Lycopodium_annotinum|m.285

MATSGGASLEQGFPSSSWNLAPEPPDDKLFSNSSPSCVMPFGGSSFGLMQGCYSGWSVLQ

HPPKQEPASPLPNFSAVISSIHQLPDLHVNPPMLPILCQSNDWRRAAADSLPFGSNSEEK

ALRKHGLFQYQMDRTLPVAWGNYGELITMSKMIPKEILKAKALAASKSHSEAERRRRERI

NTHLGTLRSLIPGTTKTDKASLLAEVVRYVKKLKRQAAEIAQVCPVPTDIDELQVDTDSS

LGGDRVLIRASLCCDDRPDLLSDLIRASRDLKLQTVKAEIFMLGGRVKNVILMTRGDGTS

CKQEGPSVTSVQEALREVMEKSGSNELSPSNFLSNKRQRMGYHSSSKI

>Lan2080604-Lycopodium_annotinum|m.287

CVMPFGGSFGLMQKPCSDWSSLQHLPKQEPVSSLPKFSAPISSFHQLADVHVNPPMLPFL

HQMNDWHRAGADSLTFGNSSEAKGPGKRDLFPYATNRTLPVLCCNHGEFMNMPKMTPKEI

MDAKALAASKSHSEAERRRRERINSHLATLRSLLPSTTKTDKASLLAEVVHHVKELKRQA

AEIAEISPVPTDVDELQVDIDSSLGEDRELIRASLCCDDRPGLLLDLTRA

>Mpa2005587-Marchantia_paleacea-non_mycorrizal|m.255

MGGEATGRLDLCSQWQITPDSGFTRPFNKPVLEEPNFAMPALPNSGVWPVAHSLATSTSS

GPSALLSVSTAAASRGLLDHHTASGSMQFNHLLPKQEVFISPNFPSMLGHSTSSLGYDHL

AQPHLFGSPLLPNAEHQLLGRNSLNSFSYDDGLVHGSSSSGGARERGKQQLPPPPGPGQA

NGGSLVLDVSKGELINLSKLTPQEILDAKALAASKSHSEAERRRRERINTHLATLRSNLP

SSTKTDKASLLGEVIDHLKFLKRQAADIAEGGPVPSDVDELTVDVDPSSSETGDGRTYYR

ASLCCDDRPDLYPDLMRTLHTLRLQTVKAEIATLGGRIKNVLLMTRSDEGSDEDDKEAPS

VSSVMEALRAVMERSGLGDQSPGSSSKRQRLASLDSSSPSM

>Mpa2002731-LFVP-Marchantia_paleacea-2_samples_combined|m.264

MGGEATGRLDLCSQWQITPDSGFTRPFNKPVLEEPNFAMPALPNSGVWPVAHSLATSTSS

GPSALLSVSTAAASRGLLDHHTASGSMQFNHLLPKQEVFISPNFPSMLGHSTSSLGYDHL

AQPHLFGSPLLPNAEHQLLGRNSLNSFSYDDGLVHGSSSSGGARERGKQQLPPPPGPGQA

NGGSLVLDVSKGELINLSKLTPQEILDAKALAASKSHSEAERRRRERINTHLATLRSNLP

SSTKTDKASLLGEVIDHLKFLKRQAADIAEGGPVPSDVDELTVDVDPSSSETGDGRTYYR

ASLCCDDRPDLYPDLMRTLHTLRLQTVKAEIATLGGRIKNVLLMTRSDEGSDEDDKEAPS

VSSVMEALRAVMERSGLGDQSPGSSSKRQRLASLDSSSPSM

>Mpa2002732-LFVP-Marchantia_paleacea-2_samples_combined|m.262

MGGEATGRLDLCSQWQITPDSGFTRPFNKPVLEEPNFAMPALPNSGVWPVAHSLATSTSS

GPSALLSVSTAAASRGLLDHHTASGSMQFNHLLPKQEVFISPNFPSMLGHSTSSLGYDHL

AQPHLFGSPLLPNAEHQLLGRNSLNSFSYDDGLVHGSSSSGGARERGKQQLPPPPGPGQA

NGGSLVLDVSKGELINLSKLTPQEILDAKALAASKSHSEAERRRRERINTHLATLRSNLP

SSTKTDKASLLGEVIDHLKFLKRQAADIAEGGPVPSDVDELTVDVDPSSSETGDGRTYYR

ASLCCDDRPDLYPDLMRTLHTLRLQTVKAEIATLGGRIKNVLLMTRSDEGSDEDDKEAPS

VSSVMEALRAVMERSGLGDQSPGSSSKRQRLASLDSSSPSM

>Ef2004374-Entransia_fimbriat|m.104

MDSGIDEEVQWPPDNDFWAMDTPVDHLGPLCPTMPANQVTVGGGITTRCHAVSSGPAVSL

APATASALASIFVDLGPANAVPAPAPPQRPIPGPNPHPGTLSPQPALPAMAPPLPLSSAT

LCQQQVQPQQKHTLQLPPISSLLRGPPSSPSRSSLNMRLVSMFEETQRVLTSSGGAGLDM

ASLVRDLARTPPPAVAMAPSPSVSHPPSPSTPSTPTTPTTPTTPVTATSIPSFSSLSSIP

PSPSQSTASGLPPISTLTSLPRFAAAPVPNFDLLASHLRAHRSPICIPVPVPPASLMDRA

VEAAARGEAQKLAEAKKATAEVEENKAAQSRNHSEAERKRREKINQHLATLRKMLPQSAK

AXASLLGEVIDYVQQLKALPEEGSDCSALLHGYLKQQLPAENDELSVKADSLPNGDEEFV

SVVRTVVCCEDHEGLLGDMVSTITGLKLRIVRAEMTAHNGRVRNVIIAALSPSAREGGHK

TTVPPLSALCKQVKGAVSSLLRKRPAASGGTGSICPSEPQTLSGCNTSDLSTGCNKRRKV

ESSPSQ

>Ip2028305-Interfilum_paradoxum|m.108

MEPSNAGGSDAVTRSGELGTPPASSLPPHMQNTASISRADSYQSTKDGSSGESKSKSHSE

AEKRRRERINAHLGTLRNLLPHNTKADKASLLGEVIEHVKELQRQADEVAEDGTPAGGDE

LLVTLETQEEGAQVVKAIMSCEDKPELLAEVVKAVQSCKLRTLRAEITTLGGRVRIEALL

GHEEEGSFDKLPKDACTAVKEALAAVMDKSAQPLGTVWPESKRQKTANAQTVSAS

>Kn000610230 –Klebsormidium nitens

MESSKSGAGDPTLRNGDSGTPPAPALPPRLQAKDNSLSRADSFQSKDGSSGESKSKSHSE

AERRRRERINAHLGTLRNLLPHPTKADKASLLGEVIDHVKELQRQADEVAGEGVPSGGDK

VLVTLDEQEGAQVLKVAMSCEDRPELLTEVVKAIQRCKLRTLRAEMTTLGGRVRLDLLLG

SEEESKEQKLPKDVCKAVRDALTAVIERNASLSNPNWPESKRQKTGNGQTVTAS*

>Mc2011025-Mesotaenium_caldariorum|m.112

MSMEQEALEWSILNKPELLLATLAGEAPLPADMMASMAPMANMSQIRRYTNNMNLGGNRR

SAIMPVSGSYTAELAQLMQNELLGLNRNEEDIHNQAGFSSGNLSLQTLSQMHSRQLQGKT

NDYGLLSRETDMLKEKMYDGMTSEEIQEVKTWAQSRSHSEAERRRRERINKHLSTLRDML

PSTPKPDKATLLAEVVNRVKQLDQRLEDLSKLAPVPNDVEEVTAVKICSQGTEEQRNQRV

KVSLCCDDRPDLLTSVIKTLQVLNLHVFRLDTSTLGSRVKHVLVLGLLTSQEDSAVGAIN

TFPTTGVILEAFKKVVDPYRSRHLDRVPSSSGYLGKRMRFSKFESSSSQSED

>Ccu2007369-Cylindrocystis_cushleckae|m.118

MPSSTWQQSVGSMGLSDSAAGSAGQSLRGVVSGPLKDLPSHFSARHGGSFTDAHLPLSNL

ANGLTSARLRGFCSGNDLGGQGGANLGLDADMLAHAQSLHNQLQHQEQSQFQLQQTNRLL

AGLDAFAGEFQGSLLSPQDMLEAKSLAQSRSHSEAERRRRERINHHLGTLREMLPASGKV

SLGMGELENFRPMWLFNDSHVRAETT

>Ccu2007370-Cylindrocystis_cushleckae|m.113

MPSSTWQQSVGSMGLSDSAAGSAGQSLRGVVSGPLKDLPSHFSARHGGSFTDAHLPLSNL

ANGLTSARLRGFCSGNDLGGQGGANLGLDADMLAHAQSLHNQLQHQEQSQFQLQQTNRLL

AGLDAFAGEFQGSLLSPQDMLEAKSLAQSRSHSEAERRRRERINHHLGTLREMLPASGKA

DKASLLADVVDKIRELEKLMAELEEDGPMPSVTEDLSVVEERTEKKGGETSMQLRATICC

YDRPDMLTALVRTLQGIRLKVVRADMATYGSRVKHVLVLVAASSQRARERELPTVETVEE

ALRGVINPSRARHMAELASSSGSSGKRPRLSPIESSSSPSE

>Zs2026450-Zygnemopsis_sp.|m.121

MAWQQPLEDYGLLSTSGVQSLHLPCVTSASMLQGIRGSGALLSPSLVSRNSLVGPDFGRI

GGGFGDELGAIDVGVFGSIQGLAASEAQVQGQAYRMLPSGSWQPSAPAAGPGLLPIDHFH

RRSRGMPTDFMAQAQGPAGAAEFLLEDPAEDAGLPQDSAEARAQAQMRSHKEAEKQRRGR

INENLATLEKMVTVNGKVHKASVLGAVIDQVTSLGKRLRQYQSVGPVPTEAEELTVVREA

VQGDDAEGGEEQFRVTLCCDDKLDVYPILIRSLQALKLEVAHKKVTTLGRRVKLELEIKG

VDSLPKGTALPPTEAVEETLIGILDPSKGRAVADHPSSSGGSSGKRQRLSGNESSWSLSE

>Mk2009666-Mesotaenium_kramstei|m.124

MPSLQLSSVTSAGINRRLSQPNINLARPSSPGNDLAGMDSLEMEFGEFGEFFGDSLQATT

GHNHSGQFLGAEVAGGAENGLNSYGLPGLEANEAQLREALTSSWHHSAGGGGQLLSSIGL

QSCQQQRGLDSPRVRPPGVTMHGDSGEKPLLLSSSWAGNAGASQLQGFGSESGSAGPGRR

VSVEDAPQLLHSLTSQQQHQQLQLRQGNRAGAVGAVEALSGDPQQLFSSIQETSEARSLA

QLKSHSEAERRRRERINHHLATLREMLPSAGKDKASVLAEVIDKVKELAQQVSQMQEAGP

VPGNVTELTVVQEGAEEDGGEGSNRLRATLCCEDRPDLLMALIRSLQALKLQMVRAEMVS

LGTRVKHVLLLAQEAAEEEDNGGGRVELPSAETVEAALKAVVDPGRGRSMGDHPSSWGSS

GKRQRLSAIESNSSPSDS

>Ssp2023033-Spirotaenia_sp.|m.127

IGGNAGAHGGSSGAGGVGSMDTFDIPGAADMSPAELAEAKALAQSKSHSEAERRRRERIN

SHLGTLRSLLPSGNKQQSDKASLLGEVIEYVKDLKRQARDANAAASAGGPATIVPGDADE

LQVELEAVPADGSGGAGKRLICAAMCCEDRPDLLADLVXAQAADGACGDCDAGDACEERV

CNGAGAARRPSSRAATAAGGRGQAGGRRWPRWSAAGGHPGGAAGGDAAGRCRRQQRW

>Me2046144-Mesotaenium_endlicherianum|m.130

MEMPGSSPINSQLYLQQLQEQLDPERLMTLDELQEAKLLAQSRSHSEAERRRRERINLHL

GTLRNLLPNTNKTNKASLLAEVIEHVQSLKRSAAKMEAASGPLPSDADDLQVEVLKSSAG

EGVGPRVRATVNCSDRPDLLPALLRAVQALKLKLVRADMAVHSGRVKHVLLLTSDATSGT

TRSIETPGFSLEESVQNAIQSVLDCRHRGGSEQPQGSSESTGKLPRMSPSSSIPSSQSL

>Ms2009561-Mougeotia_sp.|m.132

MDGDLPWEAFSNFYGQVPVSSRWLSDLGYARETLHEGTTLEELQGDKALVQSKSHSEAER

RRRERINNQLNSLRETLPNSKKSDKATLLAEVIERVNFLDRQINEQGKNSLMPVDFDDVK

VVSLLPPPIEQDASGPCFTVSLSCENRPDILPGLSRVVQGLGLQICQSDLCSLGTRVTYS

LILGKGPRREAGREQELPSVAVVLEAMKRVLDPNRAQHLGMAQASPKNAKKRLKSSSSSH

SD

>Tl2078382-Takakia_lepidozioides|m.414

METLEIGEPSTSGPVQWSSAEGHFPAHYADLVTSSNFGTRGIPLRASWEVPSDEASLSFS

AADMPARLVASTQQSPPRDETLSRMHPLRGGSGPALPLSSFPQLPNVPTPSSSLSSFPQF

PIVHTSSPSSHYDRHAHQVDGGQGPAPVGAFSLTQSLDPKASDYPYVPISSMGGSLVLDG

SRGELINIPKMTPQEMLDAKALAASKSHSEAERRRRERINTHLTTLRSLLPSSTKTDKAS

LLAEVIMHLKDLKRQAAEIGEEASVPTDVDELQVETDPSFGDQLVIRACICCNDRPDLIK

DLIKVVNSLQLHTVKAEISTLGGRIKIVLAVTNKTEGAMDGGTHRNLSLETLTNSLRVVV

DKEGSWEVSQGSSGSIKRPRLVPVQPPTSLNLRTSDGKA

>Mv2005867-Megaceros_vincentianus|m.170

MGGGPMSSVDVHWPHFIDPGVSSVASGGLAYGNPIDDYSLAMAAPAALWGNSSSRSQMTS

AAGLGAGVAGNMYSHYGTLNMSSHLPLKPPPQLSVGFGGGGGGGGGGGGVAGLGALINNS

PTHQQLYARPSPLGPVSAGALGNGIFVDVAQLASGLEAGMLGGGGGMSPLLLHQPPQMAT

GLFGTRLAAGGGSFTDMSIPLSHLQQQQQQQQQRPVLYDGLGMGGSAHLFDTATGGFPLQ

PCRASVGDGRVVGGDVLLGMSAPLPPQPSPLRPAPAIHVAREGVAAASGDLLNMSTSLPL

EPRVAAAAGNGSSRFMDMSSQFHLTSIPDLPDLSDLADFPPIPAMGSQQAYTLKEEPAQN

LSCLLTGDVPFCDRTGIGSDNIATPPLRFADLNVASVPYDTRFRQAFTDHLPPPLHPPPL

LTRRGELRDVGMPVFPLPHTPGGLPGGSLILDGARSELINISKMTPQEILDAKALAASKN

HSEAERRRRERINTHLGTLRTLLPSSTKTDKASLLGEVIEHVKELKRQAAEIEGPLPSEV

DDLTVEEGEPYAEQPGKSVFKVIVSCDDRPCLLTDLGATLDRCNLENFHMELATVSGRIN

VVMLAYSRDDDVPEAGKHEAFMSSVQEAIRSVVDKSGGTDNSSGSSGGGLKRQRTMSNSP

P

>Aa2050828-Anthoceros_agrestis-A|m.177

EGEGLRVEVECYSSMGGGSGVEVHWPNLIDSNLCAISGGYGSGPQMQMGGSFVDDYSLAM

AAPAALWSGGMAPSAAPVSASVCAPAGAQGTGLSAGLVNFTSSQLHFGSGFGFGLGSGGS

LNMPGMNMSMSMAQSGNMGLVSPEPTLFEMVNAVPQSLRALGPGFGAGGMLDMAGAMQQQ

QQQQPLDSPGLGLGSGCSAAAGAFALQPELEAAELVGVAGNGDLHALPPMPPLLGGGLGD

DAAASLLNMAVNVALPMPMPLQQGPGLGEGSNDGLFGLPAGMSMPMEAGVSGSGAGGSFM

DLAARFQLNSFSDLADLSAITSQPLNYILQQDPGAGMQPMVSAVGPRIGGVPDNIPAPAP

PPPPPPLRFADLNVTSIPYDSRYRQPYGASVPPGLLFLKPESKLDHGFPGSSPLHAGGGI

GRGVGVGGLGLGLGGGVLPGGSLVLDGARSELINISKMTPQEILDAKALAASKSHSEAER

RRRERINTHLGTLRTMLPSSTKTDKASLLGEVIEHVKELKRQAADIDGPVPTEVDELGVE

MSEASEKHPGKLIIKLTVCCDDRPGLLTAIGRAVEACKLDNFSVELSTLAGRLKVVMLAA

CTESVQEAKQDAVMSSLQEAVRAALDKVGPATAAATSTRTNSSGSSGGLKRQRTGMEASS

PSN

>Mto2053481-Megaceros_tosanus|m.183

MGGGPISSVDVHWPHFIDSALSSIATAGLGYGNPVDDYSLAMAAPAALWGNSTTNTRAQI

TPAAGLGAVVPGNMHTHYGALLNMSTHLAAARPPPQLSAGFGGGAAAAATGMAAALINNC

PTHQQQQQQQFYARPSPLGPASAGGALLGNGTLVDVPQLASGLEAGMLGGGGGMSPLLVH

QHHQPPQQMASGIFGNRLAPGGASFTDMSIALSDLQQQQRPMLYDRARLFDAPTTTGFPL

QQQQPSSRGSAGDPSIVGGDVLMMGMSRPLPSQPLSSRVSAAAVRAGREGVGADGADLLN

MSTSLPLLQPGVAGAQGTGSRFMDMSSQFQLTSIPDLPDLSDLADFPAIPAMGSHQAYSL

KQEPAQNLTCRLTVDVPFFDRTGIGSDTIATPPLRFADLNVASVPYDTRFRQVFADHLAP

PLPPPLLLARRGELRDVGMPVFPIPQTAGGGLPGGSLILDGARSELINISKMTPQEILDA

KALAASKNHSEAERRRRERINTHLGTLRTLLPCSTKTDKASLLWEVIDHVKELKRQAAEI

EGPIPSEVDDLTVEEGESCAEQPGKAVFKVTVSCDDRPCLLTDLGATLDRCNLENFHMEL

ATVSGRMNVVMLAYSRDEDVPEPGKHEAFMSSVQEALRSVVDKSGVTDNSSGSSGGGLKR

QRTMSNSPP

>Phc2017550-Phaeoceros_carolinianus-gametophyte|m.186

MGGGATSSVDVHWPRFIDSSPSSIGSGFGYGNQVIDDYSLAMAAPPSVWGNSSSSMSQIS

PGAAGNVLNSGYGQLNYGNGLFNMHVASSFKPPQVSVGFGPGAASGLGGINLGSLMNNSS

VLQAGSRPMPSALGTGTLMDVTQVAPGLGQTGGLAGNWNLVTGMSRAPLLQQQQPTAGAS

PGFGSGGGLAGTGPFMDMSTVVPGLHLQQQQQRVGSGFGSLFDMTSFGFPSALDNSGLPS

ALDNGGLLGMPSPLPQQQAPELAGNVVDLLSMPVSLPLQQPRVLPGSTNGSFVDLSSQFQ

LTSIPDLPDLVGLSDLADLPSIPAAVGTQAYKQQPSQNYMTLSAADAPVTFLGSSTGLAD

PIVAPPLRFADLNVASIPYDNRLGLRPAFGTQLPPPLLSSGRGDLRDAGMSSLFPSPPTL

GVLGLPGGSLVLDGSRSELINISKMTPQEILDAKALAASKNHSEAERRRRERINTHLGTL

RTLLPSSTKTDKASLLGEVIEHVRGLKRQAAEVEGPVPSETEELTVERGEPYAEQPGRPV

IKVTVSCDDRPGVLTDLARTLDGCKLENFHMELATISGRIKVVMQAYSTDETLAERGVSE

AFLANVQDALRSVVDRAGATDNSSGSSGGGLKRQRTMSNSPP

>Phc2015014-Phaeoceros_carolinianus-gametophyte_1575|m.191

MGGGATSSVDVHWPRFIDSSPSSIGSGFGYGNQVIDDYSLAMAAPPSVWGNSSSSMSQIS

PGAAGNVLNSGYGQLNYGNGLFNMHVASQLSFKPPQVSVGFGPGAASGLGGISLGSLMNN

SSIIQAGSRPMPTALGTGTLMDVTQVAPGLGQTGGLVGNRNLVTGMSRAPLLQQQQPTAG

ASSGFGIGGGLAGTSPFMDMSTVVPGLHLQQQQQRVGSGFGSLFEMTSFGFPSALDNGGL

LGMPSPLPQQQAPELAGNVVDLLSIPVSLPLQQPRVMPGSTNGSFVDLPSQFQLTSIPDL

PDLVGLSDLADLPSIPAAVGTQAYKQEPAQNYMTLSAADAPVTFLGSSTGLSDPIVAPPL

RFADLNVASIPYDNRLGLRPAFGTQLPPPLLSSGRGDLRDAGMSSLFPSPPALGVLGLPG

GSLVLDGSRSELINISKMTPQEILDAKALAASKNHSEAERRRRERINTHLGTLRTLLPSS

TKTDKASLLGEVIEHVRGLKRQAAEVEGPVPSETEELTVERGEPYAEQPGRPVIKVTVSC

DDRPGVLTDLARTLDGCKLENFHMELATISGRIKVVMQAYSTDETLAERGVSEAFLANVQ

DALRSVVDKAGATDNSSGSSGGGLKRQRTMSNSPP

>Phc2106750-Phaeoceros_carolinianus-sporophyte_1575|m.197

ERINTHLGTLRTLLPSSTKTDKASLLGEVIEHVRGLKRLAAEVEGPVPSETEELTVERGE

PYAEQPGRPVIKVTVSCDDRPGVLTDLARTLDGCKLENFHMELATISGRIKVVMQAYSTD

ETLAERGVSEAFLANVQDALRSVVDKAGATDNSRSS

>Pc02009198-Phaeomegaceros_coriaceus|m.134

MEGVAAASGDLLNMSTSLPLQPGVAGNGSRFMDMSSQFHLTSIPDLPDLSDLADFPAIPA

TGSQQAYALKQEPVQNLTCRLTGDVPFFDRTGIGSDTIATPPLRFADLNVASVPYDNRFR

QAFANQLPPPLPPPPLLSRRGELRDVGMPLFPLPQPAGGLPGGSLVLDGARSELINISKM

TPQEILDAKALAASKNHSEAERRRRERINTHLGTLRTLLPSSTKTDKASLLGEVIEHVKE

LKRQAAEIEGPVPSEVDDLTVEEGEPCAEQPGKTVFKVTVSCDDRPCLLTDLGATLDRCN

LENFHMELATVSGRINIVMLAYSRDGNVPEAGKHEAFMSSVQEALRSVVDKAGGTDNSSG

SSGGGLKRQRTMSNSPP

>Ldu2035940-Leiosporoceros_dussii-B|m.139

MAGVPVDLHWPSVVDSSLCARDDYSLAVAASPIPAPSSWPAATALQDAGGGGGGGHTGVV

NLMNEVELAASRSSGCWDVYTRSSSLPQPVQPCWLGAGNSLLDLPRIGGDVVKMDPGSHS

TFFLNSSPPACHPQLFPAALVSNLQEEGPLRFADLNVSSIPYDSRYRQPGPQIGVGGSLV

LDGARGELINISKMTPQEILDAKALAASKSHSEAERRRRERINTHLGTLRSMLPSSTKTD

KASLLGEVIDHVKDLKRQAAQTEGPVPSEADELVVEPGEARDADPGKAVFKVTVCCDDRP

GLLTELSRVLEDFNLDNFNAELATLAGRMKAVLLVLSSQPEACNRNAFAGTVQEALQAVL

DKAGGGAVSSGSSGPLKRLRTSSTSPSATKDG

>Na2038150-Nothoceros_aenigmaticus|m.151

HYGTLNMSSHLPPKPPPQLSVGFGGGFGGGEAVAGLGALINNSPTHQQLYARPSPLGPVS

AGALGNGIFVDVAQLASGLEAGMLGGGGGMSPLTLHQPPQMATGLFGTRLAAGGGSFTDM

SIPLSHLQQQQQQQQQQQRPVLYDVLGMGGSAHLFDTATGGFPLQPCRASVGDGRVVGGD

VLMGMSAPLPLQPSPLRPAAPLHVAREGVAAASGDLLNMSTSLPLETRVAAAAGNGSSRF

MDMSSQFHLTSIPDLPDLSDLADFPPIPAMGSQQAYTLKEEPAQNLSCLLTGDVPFCDRT

GIGSDNIATTPMRFADLNVASVPYDTRFRQAFTDHLPPPLHPPPLLNRRGELRDVGMPVF

PLPHTPGGLPGGSLILDGARSELINISKMTPQEILDAKALAASKNHSEAERRRRERINTH

LGTLRTLLPSSTKTDKASLLGEVIEHVKELKRQAAEIEGPLPSEVDDLTVEEGEPYAEQP

GKSVFKVIVSCDDRPCLLTDLGATLDRCNLENFHMELATVSGRINVVMLAYSRDEDVPEA

GKHEAFMSGVQEAIRSVVDKSGGTDNSSGSSGGGLKRQRTMSNSPP

>Ph2057551-Paraphymatoceros_hallii|m.155

MGGGPMSSVDVHWPHFIDSSLSSIGSGLGYGNQVVDDYSLAMAAPASVWGNSGSSMSQIS

PGAAGNVNNGYGQLYGSGLFNMHMSSQLSFKPPQVSMGFGPGAAATGGGINFGSLINNSS

ISQVGTRPLPAAASAAMGTGTLLDVTHLDSGLGQAGMSGNRNLVGMSPLLQQPAAAASPG

FGIGGGMAGTSFMNMSTVVPGLNLQQQQRGMGSGIGSLFDMTSFGFPSAVRDGGLLGRPS

PLPQQQRQQPPELAGNVVDLLSMPISLPLQQPSRVLPGNSNGSFGDLSSQFHLTSIPDLP

DLVGLSDLADLPSIPAVASEAYKQEPAQNYTLSAAGEVPVPFFAGSTGLSDSIVAPPLRF

ADLNVASIPYDNRLGLRQAFGTQLPSPLPPPLLGSGRGDGISSLFPSPPTLGGLGLQGGS

LVLDGSRSELINISKMTPQEILDAKALAASKNHSEAERRRRERINTHLGTLRTLLPSSTK

TDKASLLGEVIEHVKGLKRQAAEVEGPVPSETDELNVERGEPYAEQPGRPVIKVTVSCDD

RPGVLTDLARTLDSCKLENFHMELATISGRIKVVMQAYSTDETLAEAGVTEAFLTNVQDA

LRSVVEKSGATDNSSGSSGGGLKRQRTTSNSPT

>Phc2020949-Phaeoceros_carolinianus-sporophyte|m.165

APPLRFADLNVASIPYDNRLGLRPAFGTQLPPPLLSSGRGDLRDAGMSSLFPSPPALGVL

GLPGGSLVLDGSRSELINISKMTPQEILDAKALAASKNHSEAERRRRERINTHLGTLRTL

LPSSTKTDKASLLGEVIEHVRGLKRQAAEVEGPVPSETEELTVERGEPYAEQPGRPVIKV

TVSCDDRPGVLTDLARTLDGCKLENFHMELATISGRIKVVMQAYSTDETLAERGVSEAFL

ANVQDALRSVVDKAGATDNSSGSSGGGLKRQRTMSNSPP

>Phc2126364-Phaeoceros_carolinianus-sporophyte|m.167

THLGTLRTMLPSSTKTDKASLLGEVIEHVKELKRQAADIDGPVPTEVDELGVEMSEASEK

HPGKLIIKLTVCCDDRPGLLTAIGRAVEACKLDNFSVELSTLAGRLKMVMLAACTESVQE

AKQDAVMSSLQEAVRAALDKVGPATAAATSTRTNSSGSSGGLKRQRTGMEASS

>Lc2011718-Lunularia_cruciata|m.266

MGGEATGRVDFCSQWQITSDPGLSRSSNKSVLEEPNFSIPVLPNSGVWTVGQSLAATASA

GCSALLSVSAVAASRGLLDNHAASGTMQCNHLLLKSEPFISSQFPLMVGHTVPPLTYDQL

GQQQHDFGNSPMHNSECNHLGRSPHINSLGCFNNNGVQGHSGGPVHRERGKQPQIASLPG

GQSTGGSLVLDASKGELINLSKLTPQEILDAKALAASKSHSEAERRRRERINTHLATLRS

NLPSSTKTDKASLLGEVIDHLKYLKRQAADTAEGGPVPSDVDELTVDLDPSTNEMGDGRV

YYRASICCDDRSDLYPDLMRTLQNLRLQTVKAEIATLGGRIKNVLLMTRSDGGNDEDDKE

SPSVSSVMEALRVVMERSGLGDQSSGSSSKRQRLASLDSSSPSM

>Mpa2010922-Marchantia_paleacea-mycorrizal|m.257

MGGEATGRLDLCSQWQITPDSGFTRPFNKPVLEEPNFAMPALPNSGVWPVAHSLATSTSS

GPSALLSVSTAAASRGLLDHHTASGSMQFNHLLPKQEVFISPNFPSMLGHSTSSLGYDHL

AQPHLFGSPLLPNAEHQLLGRNSLNSFSYDDGLVHGSSSSGGARERGKQQLPPPPGPGQA

NGGSLVLDVSKGELINLSKLTPQEILDAKALAASKSHSEAERRRRERINTHLATLRSNLP

SSTKTDKASLLGEVIDHLKFLKRQAADIAEGGPVPSDVDELTVDVDPSSSETGDGRTYYR

ASLCCDDRPDLYPDLMRTLHTLRLQTVKAEIATLGGRIKNVLLMTRSDEGSDEDDKEAPS

VSSVMEALRAVMERSGLGDQSPGSSSKRQRLASLDSSSPSM

>Coco2044748-Conocephalum_conicum|m.259

QLVHIAVWSMAGESAGRLDLRARWQITETDFMRPLNKTLLEESNFAMPPLPNSGVWPMGR

SVPTSAGSGPSALLSVSAVAASRGLLDRHTASEAMQFSNLLPKQEPLVSSHLSSILGHSA

SSMAYDHSAQQHLFGSTSMRNAEHHLLGRSQLNLFSYDDVNGHNATGTSAARERGKQSQQ

FLPPTAPGSSNGGSLVLDVSKGELINLSKLTPQEILDAKALAASKNHSEAERRRRERINT

HLGTLRNNLPSSTKTDKASLLGEVIEHLKYLKRQASDIAEGGPVPSDVDELTVDVDPSSN

EFGDGRIYYRASICCDDRPDLYPDLMKTLRNLRLQTIKVEIATLGGRIKNNLLMTRSDED

YDEADKEGPSRSSVMEALRAVMERSGVGDQSPRPTKRHRLASLDSSSSSL

>At1G01260

MNIGRLVWNEDDKAIVASLLGKRALDYLLSNSVSNANLLMTLGSDENLQNKLSDLVERPNASNFSWNYAIFWQISRSKAG

DLVLCWGDGYCREPKEGEKSEIVRILSMGREEETHQTMRKRVLQKLHDLFGGSEEENCALGLDRVTDTEMFLLSSMYFSF

PRGEGGPGKCFASAKPVWLSDVVNSGSDYCVRSFLAKSAGIQTVVLVPTDLGVVELGSTSCLPESEDSILSIRSLFTSSL

PPVRAVALPVTVAEKIDDNRTKIFGKDLHNSGFLQHHQHHQQQQQQPPQQQQHRQFREKLTVRKMDDRAPKRLDAYPNNG

NRFMFSNPGTNNNTLLSPTWVQPENYTRPINVKEVPSTDEFKFLPLQQSSQRLLPPAQMQIDFSAASSRASENNSDGEGG

GEWADAVGADESGNNRPRKRGRRPANGRAEALNHVEAERQRREKLNQRFYALRSVVPNISKMDKASLLGDAVSYINELHA

KLKVMEAERERLGYSSNPPISLDSDINVQTSGEDVTVRINCPLESHPASRIFHAFEESKVEVINSNLEVSQDTVLHTFVV

KSEELTKEKLISALSREQTNSVQSRTSSGR

>At1G02340

MSNNQAFMELGWRNDVGSLAVKDQGMMSERARSDEDRLINGLKWGYGYFDHDQTDNYLQIVPEIHKEVENAKEDLLVVVP

DEHSETDDHHHIKDFSERSDHRFYLRNKHENPKKRRIQVLSSDDESEEFTREVPSVTRKGSKRRRRDEKMSNKMRKLQQL

VPNCHKTDKVSVLDKTIEYMKNLQLQLQMMSTVGVNPYFLPATLGFGMHNHMLTAMASAHGLNPANHMMPSPLIPALNWP

LPPFTNISFPHSSSQSLFLTTSSPASSPQSLHGLVPYFPSFLDFSSHAMRRL

>At1G06170

MGGGGMFEEIGCFDPNAPAEMTAESSFSPSEPPPTITVIGSNSNSNCSLEDLSAFHLSPQDSSLPASASAYAHQLHINATPNCDHQFQSSMHQTLQDPSYAQQSNHWDNGYQDFVNLGPNHTTPDLLSLLQLPRSSLPPFANPSIQDIIMTTSSSVAAYD

PLFHLNFPLQPPNGSFMGVDQDQTETNQGVNLMYDEENNNLDDGLNRKGRGSKKRKIFPTERERRVHFKDRFGDLKNLIP

NPTKNDRASIVGEAIDYIKELLRTIDEFKLLVEKKRVKQRNREGDDVVDENFKAQSEVVEQCLINKKNNALRCSWLKRKS

KFTDVDVRIIDDEVTIKIVQKKKINCLLFVSKVVDQLELDLHHVAGAQIGEHHSFLFNAKISEGSSVYASAIADRVMEVL

KKQYMEALSANNGYHCYSSD

>At1G09530

MPLFELFRLTKAKLESAQDRNPSPPVDEVVELVWENGQISTQSQSSRSRNIPPPQANSSRAREIGNGSKTTMVDEIPMSV

PSLMTGLSQDDDFVPWLNHHPSLDGYCSDFLRDVSSPVTVNEQESDMAVNQTAFPLFQRRKDGNESAPAASSSQYNGFQS

HSLYGSDRARDLPSQQTNPDRFTQTQEPLITSNKPSLVNFSHFLRPATFAKTTNNNLHDTKEKSPQSPPNVFQTRVLGAK

DSEDKVLNESVASATPKDNQKACLISEDSCRKDQESEKAVVCSSVGSGNSLDGPSESPSLSLKRKHSNIQDIDCHSEDVE

EESGDGRKEAGPSRTGLGSKRSRSAEVHNLSERRRRDRINEKMRALQELIPNCNKVDKASMLDEAIEYLKSLQLQVQIMS

MASGYYLPPAVMFPPGMGHYPAAAAAMAMGMGMPYAMGLPDLSRGGSSVNHGPQFQVSGMQQQPVAMGIPRVSGGGIFAG

SSTIGNGSTRDLSGSKDQTTTNNNSNLKPIKRKQGSSDQFCGSS

>At1G10120

MGGESNEGGEMGFKHGDDESGGISRVGITSMPLYAKADPFFSSADWDPVVNAAAAGFSSSHYHPSMAMDNPGMSCFSHYQ

PGSVSGFAADMPASLLPFGDCGGGQIGHFLGSDKKGERLIRAGESSHEDHHQVSDDAVLGASPVGKRRLPEAESQWNKKA

VEEFQEDPQRGNDQSQKKHKNDQSKETVNKESSQSEEAPKENYIHMRARRGQATNSHSLAERVRREKISERMRLLQELVP

GCNKITGKAVMLDEIINYVQSLQQQVEFLSMKLATVNPEINIDIDRILAKDLLQSRDRNTPTLGLNPFAGFQGNIPNLSA

TTNPQYNPLPQTTLESELQNLYQMGFVSNPSTMSSFSPNGRLKPEL

>At1G10610

MMMMRGGERVKEFLRPFVDSRTWDLCVIWKLGDDPSRFIEWVGCCCSGCYIDKNIKLENSEEGGTGRKKKASFCRDDHNK

HRIRTLACEALSRFPLFMPLYPGIHGEVVMSKSPKWLVNSGSKMEMFSTRVLVPVSDGLVELFAFDMRPFDESMVHLIMS

RCTTFFEPFPEQRLQFRIIPRAEESMSSGVNLSVEGGGSSSVSNPSSETQNLFGNYPNASCVEILREEQTPCLIMNKEKD

VVVQNANDSKANKKLLPTENFKSKNLHSERKRRERINQAMYGLRAVVPKITKLNKIGIFSDAVDYINELLVEKQKLEDEL

KGINEMECKEIAAEEQSAIADPEAERVSSKSNKRVKKNEVKIEVHETGERDFLIRVVQEHKQDGFKRLIEAVDLCELEII

DVNFTRLDLTVMTVLNVKANKDGIACGILRDLLLKMMITSI

>At1G12860

MNSDGVWLDGSGESPEVNNGEAASWVRNPDEDWFNNPPPPQHTNQNDFRFNGGFPLNPSENLLLLLQQSIDSSSSSSPLL

HPFTLDAASQQQQQQQQQQEQSFLATKACIVSLLNVPTINNNTFDDFGFDSGFLGQQFHGNHQSPNSMNFTGLNHSVPDF

LPAPENSSGSCGLSPLFSNRAKVLKPLQVMASSGSQPTLFQKRAAMRQSSSSKMCNSESSSEMRKSSYEREIDDTSTGII

DISGLNYESDDHNTNNNKGKKKGMPAKNLMAERRRRKKLNDRLYMLRSVVPKISKMDRASILGDAIDYLKELLQRINDLH

TELESTPPSSSSLHPLTPTPQTLSYRVKEELCPSSSLPSPKGQQPRVEVRLREGKAVNIHMFCGRRPGLLLSTMRALDNL

GLDVQQAVISCFNGFALDVFRAEQCQEDHDVLPEQIKAVLLDTAGYAGLV

>At1G18400

MANFENLSSDFQTIAMDIYSSITQAADLNNNNSNLHFQTFHPSSTSLESLFLHHHQQQLLHFPGNSPDSSNNFSSTSSFL

HSDHNIVDETKKRKALLPTLSSSETSGVSDNTNVIATETGSLRRGKRLKKKKEEEDEKEREVVHVRARRGQATDSHSLAE

RVRRGKINERLRCLQDMVPGCYKAMGMATMLDEIINYVQSLQNQVEFLSMKLTAASSFYDFNSETDAVDSMQRAKARETV

EMGRQTRDGSPVFHLSTWSL

>At1G25330

MARFEPYNYNNGHDPFFAHINQNPELINLDLPASTPSSFMLFSNGALVDANHNNSHFFPNLLHGNTRRKGNKEESGSKRR

RKRSEEEEAMNGDETQKPKDVVHVRAKRGQATDSHSLAERVRREKINERLKCLQDLVPGCYKAMGMAVMLDVIIDYVRSL

QNQIEFLSMKLSAASACYDLNSLDIEPTDIFQGGNIHSAAEMERILRESVGTQPPNFSSTLPF

>At1G26260

MSDKDEFAAKKKDLVNTPVDLYPPENPMLGPSPMMDSFRETLWHDGGFNVHTDADTSFRGNNNIDIPLEMGWNMAQFPAD

SGFIERAAKFSFFGCGEMMMNQQQSSLGVPDSTGLFLQDTQIPSGSKLDNGPLTDASKLVKERSINNVSEDSQSSGGNGH

DDAKCGQTSSKGFSSKKRKRIGKDCEEEEDKKQKDEQSPTSNANKTNSEKQPSDSLKDGYIHMRARRGQATNSHSLAERV

RREKISERMKFLQDLVPGCDKVTGKAVMLDEIINYVQSLQCQIEFLSMKLSAVNPVLDFNLESLLAKDALQSSAPTFPHN

MSMLYPPVSYLSQTGFMQPNISSMSLLSGGLKRQETHGYESDHHNLVHMNHETGTAPDHEDTTADMKVEP

>At1G27740

MDVFVDGELESLLGMFNFDQCSSSKEERPRDELLGLSSLYNGHLHQHQHHNNVLSSDHHAFLLPDMFPFGAMPGGNLPAM

LDSWDQSHHLQETSSLKRKLLDVENLCKTNSNCDVTRQELAKSKKKQRVSSESNTVDESNTNWVDGQSLSNSSDDEKASV

TSVKGKTRATKGTATDPQSLYARKRREKINERLKTLQNLVPNGTKVDISTMLEEAVHYVKFLQLQIKLLSSDDLWMYAPL

AYNGLDMGFHHNLLSRLM

>At1G32640

MTDYRLQPTMNLWTTDDNASMMEAFMSSSDISTLWPPASTTTTTATTETTPTPAMEIPAQAGFNQETLQQRLQALIEGTH

EGWTYAIFWQPSYDFSGASVLGWGDGYYKGEEDKANPRRRSSSPPFSTPADQEYRKKVLRELNSLISGGVAPSDDAVDEE

VTDTEWFFLVSMTQSFACGAGLAGKAFATGNAVWVSGSDQLSGSGCERAKQGGVFGMHTIACIPSANGVVEVGSTEPIRQ

SSDLINKVRILFNFDGGAGDLSGLNWNLDPDQGENDPSMWINDPIGTPGSNEPGNGAPSSSSQLFSKSIQFENGSSSTIT

ENPNLDPTPSPVHSQTQNPKFNNTFSRELNFSTSSSTLVKPRSGEILNFGDEGKRSSGNPDPSSYSGQTQFENKRKRSMV

LNEDKVLSFGDKTAGESDHSDLEASVVKEVAVEKRPKKRGRKPANGREEPLNHVEAERQRREKLNQRFYALRAVVPNVSK

MDKASLLGDAIAYINELKSKVVKTESEKLQIKNQLEEVKLELAGRKASASGGDMSSSCSSIKPVGMEIEVKIIGWDAMIR

VESSKRNHPAARLMSALMDLELEVNHASMSVVNDLMIQQATVKMGFRIYTQEQLRASLISKIG

>At1G51140

MESEFQQHHFLLHDHQHQRPRNSGLIRYQSAPSSYFSSFGESIEEFLDRPTSPETERILSGFLQTTDTSDNVDSFLHHTF

NSDGTEKKPPEVKTEDEDAEIPVTATATAMEVVVSGDGEISVNPEVSIGYVASVSRNKRPREKDDRTPVNNLARHNSSPA

GLFSSIDVETAYAAVMKSMGGFGGSNVMSTSNTEASSLTPRSKLLPPTSRAMSPISEVDVKPGFSSRLPPRTLSGGFNRS

FGNEGSASSKLTALARTQSGGLDQYKTKDEDSASRRPPLAHHMSLPKSLSDIEQLLSDSIPCKIRAKRGCATHPRSIAER

VRRTKISERMRKLQDLVPNMDTQTNTADMLDLAVQYIKDLQEQVKALEESRARCRCSSA

>At1G59640

MDPSGMMNEGGPFNLAEIWQFPLNGVSTAGDSSRRSFVGPNQFGDADLTTAANGDPARMSHALSQAVIEGISGAWKRRED

ESKSAKIVSTIGASEGENKRQKIDEVCDGKAEAESLGTETEQKKQQMEPTKDYIHVRARRGQATDSHSLAERARREKISE

RMKILQDLVPGCNKVIGKALVLDEIINYIQSLQRQVEFLSMKLEAVNSRMNPGIEVFPPKEVMILMIINSIFSIFFTKQY

MFLSRYSRGRSLDVYAVRSFKHCNKRSDLCFCSCSPKTELKTTIFSQNMTCFCRYSRVGVAISSSKHCNEPVTLCFYSYC

LRKIYHFLLWNLKYKIQKSVLFS

>At1G63650

MATGENRTVPDNLKKQLAVSVRNIQWSYGIFWSVSASQPGVLEWGDGYYNGDIKTRKTIQAAEVKIDQLGLERSEQLREL

YESLSLAESSASGSSQVTRRASAAALSPEDLTDTEWYYLVCMSFVFNIGEGIPGGALSNGEPIWLCNAETADSKVFTRSL

LAKSASLQTVVCFPFLGGVLEIGTTEHIKEDMNVIQSVKTLFLEAPPYTTISTRSDYQEIFDPLSDDKYTPVFITEAFPT

TSTSGFEQEPEDHDSFINDGGASQVQSWQFVGEEISNCIHQSLNSSDCVSQTFVGTTGRLACDPRKSRIQRLGQIQEQSN

HVNMDDDVHYQGVISTIFKTTHQLILGPQFQNFDKRSSFTRWKRSSSVKTLGEKSQKMIKKILFEVPLMNKKEELLPDTP

EETGNHALSEKKRREKLNERFMTLRSIIPSISKIDKVSILDDTIEYLQDLQKRVQELESCRESADTETRITMMKRKKPDD

EEERASANCMNSKRKGSDVNVGEDEPADIGYAGLTDNLRISSLGNEVVIELRCAWREGILLEIMDVISDLNLDSHSVQSS

TGDGLLCLTVNCKHKGTKIATTGMIQEALQRVAWIC

>At1G66470

MALVNDHPNETNYLSKQNSSSSEDLSSPGLDQPDAAYAGGGGGGGSASSSSTMNSDHQQHQGFVFYPSGEDHHNSLMDFN

GSSFLNFDHHESFPPPAISCGGSSGGGGFSFLEGNNMSYGFTNWNHQHHMDIISPRSTETPQGQKDWLYSDSTVVTTGSR

NESLSPKSAGNKRSHTGESTQPSKKLSSGVTGKTKPKPTTSPKDPQSLAAKNRRERISERLKILQELVPNGTKVDLVTML

EKAISYVKFLQVQVKVLATDEFWPAQGGKAPDISQVKDAIDAILSSSQRDRNSNLITN

>At1G68810

MCAKKEEEEEEEEDSSEAMNNIQNYQNDLFFHQLISHHHHHHHDPSQSETLGASGNVGSGFTIFSQDSVSPIWSLPPPTS

IQPPFDQFPPPSSSPASFYGSFFNRSRAHHQGLQFGYEGFGGATSAAHHHHEQLRILSEALGPVVQAGSGPFGLQAELGK

MTAQEIMDAKALAASKSHSEAERRRRERINNHLAKLRSILPNTTKTDKASLLAEVIQHVKELKRETSVISETNLVPTESD

ELTVAFTEEEETGDGRFVIKASLCCEDRSDLLPDMIKTLKAMRLKTLKAEITTVGGRVKNVLFVTGEESSGEEVEEEYCI

GTIEEALKAVMEKSNVEESSSSGNAKRQRMSSHNTITIVEQQQQYNQR

>At1G68920

MDLSAKDEFSAEKRNPDNYDSVNNPSGDWRVDSYPSENLISAGPASCSPSQMMDSFGQTLWYDPTSVQAVGYAGFNGGNA

SSSSFRGSIDRSLEMGWNLPNLLPPKGNGLFLPNASSFLPPSMAQFPADSGFIERAARFSLFSGGNFSDMVNQPLGNSEA

IGLFLQGGGTMQGQCQSNELNVGEPHNDVSVAVKESTVRSSEQAKPNVPGSGNVSEDTQSSGGNGQKGRETSSNTKKRKR

NGQKNSEAAQSHRSQQSEEEPDNNGDEKRNDEQSPNSPGKKSNSGKQQGKQSSDPPKDGYIHVRARRGQATNSHSLAERV

RREKISERMKFLQDLVPGCNKVTGKAVMLDEIINYVQSLQRQVEFLSMKLATVNPQMDFNLEGLLAKDALQLRAGSSSTT

PFPPNMSMAYPPLPHGFMQQTLSSIGRTITSPLSPMNGGFKRQETNGWEGDLQNVIHINYGAGDVTPDPQAAATASLPAA

NMKVEP

>At1G69010

MRTGKGNQEEEDYGEEDFNSKREGPSSNTTVHSNRDSKENDKASAIRSKHSVTEQRRRSKINERFQILRELIPNSEQKRD

TASFLLEVIDYVQYLQEKVQKYEGSYPGWSQEPTKLTPWRNNHWRVQSLGNHPVAINNGSGPGIPFPGKFEDNTVTSTPA

IIAEPQIPIESDKARAITGISIESQPELDDKGLPPLQPILPMVQGEQANECPATSDGLGQSNDLVIEGGTISISSAYSHE

LLSSLTQALQNAGIDLSQAKLSVQIDLGKRANQGLTHEEPSSKNPLSYDTQGRDSSVEEESEHSHKRMKTL

>At1G73830

MANLSSDFQTFTMDDPIRQLAELSNTLHHFQTFPPPFSSSLDSLFFHNQFPDHFPGKSLENNFHQGIFFPSNIQNNEESS

SQFDTKKRKSLMEAVSTSENSVSDQTLSTSSAQVSINGNISTKNNSSRRGKRSKNREEEKEREVVHVRARRGQATDSHSI

AERVRRGKINERLKCLQDIVPGCYKTMGMATMLDEIINYVQSLQNQVEFLSMKLTAASSYYDFNSETDAVESMQKAKARE

AVEMGQGRDGSSVFHSSSWTL

>At2G14760

MEAMGEWSTGLGGIYTEEADFMNQLLASYEQPCGGSSSETTATLTAYHHQGSQWNGGFCFSQESSSYSGYCAAMPRQEED

NNGMEDATINTNLYLVGEETSECDATEYSGKSLLPLETVAENHDHSMLQPENSLTTTTDEKMFNQCESSKKRTRATTTDK

NKRANKARRSQKCVEMSGENENSGEEEYTEKAAGKRKTKPLKPQKTCCSDDESNGGDTFLSKEDGEDSKALNLNGKTRAS

RGAATDPQSLYARLKQLNKVHCMMVQKRRERINERLRILQHLVPNGTKVDISTMLEEAVQYVKFLQLQIKLLSSDDLWMY

APIAYNGMDIGLDLKLNALTR

>At2G16910

MESNMQNLLEKLRPLVGARAWDYCVLWRLNEDQRFVKWMGCCCGGTELIAENGTEEFSYGGCRDVMFHHPRTKSCEFLSH

LPASIPLDSGIYAETLLTNQTGWLSESSEPSFMQETICTRVLIPIPGGLVELFATRHVAEDQNVVDFVMGHCNMLMDDSV

TINMMVADEVESKPYGMLSGDIQQKGSKEEDMMNLPSSYDISADQIRLNFLPQMSDYETQHLKMKSDYHHQALGYLPENG

NKEMMGMNPFNTVEEDGIPVIGEPSLLVNEQQVVNDKDMNENGRVDSGSDCSDQIDDEDDPKYKKKSGKGSQAKNLMAER

RRRKKLNDRLYALRSLVPRITKLDRASILGDAINYVKELQNEAKELQDELEENSETEDGSNRPQGGMSLNGTVVTGFHPG

LSCNSNVPSVKQDVDLENSNDKGQEMEPQVDVAQLDGREFFVKVICEYKPGGFTRLMEALDSLGLEVTNANTTRYLSLVS

NVFKVEKNDNEMVQAEHVRNSLLEITRNTSRGWQDDQMATGSMQNEKNEVDYQHYDDHQHHNGHHHPFDHQMNQSAHHHH

HHQHINHYHNQ

>At2G18300

MLEGLVSQESLSLNSMDMSVLERLKWVQQQQQQLQQVVSHSSNNSPELLQILQFHGSNNDELLESSFSQFQMLGSGFGPN

YNMGFGPPHESISRTSSCHMEPVDTMEVLLKTGEETRAVALKNKRKPEVKTREEQKTEKKIKVEAETESSMKGKSNMGNT

EASSDTSKETSKGASENQKLDYIHVRARRGQATDRHSLAERARREKISKKMKYLQDIVPGCNKVTGKAGMLDEIINYVQC

LQRQVEFLSMKLAVLNPELELAVEDVSVKQFQAYFTNVVASKQSIMVDVPLFPLDQQGSLDLSAINPNQTTSIEAPSGSW

ETQSQSLYNTSSLENSCGNYNKISKILLSTKCTHQYVPIRRVWV

>At2G20180

MHHFVPDFDTDDDYVNNHNSSLNHLPRKSITTMGEDDDLMELLWQNGQVVVQNQRLHTKKPSSSPPKLLPSMDPQQQPSS

DQNLFIQEDEMTSWLHYPLRDDDFCSDLLFSAAPTATATATVSQVTAARPPVSSTNESRPPVRNFMNFSRLRGDFNNGRG

GESGPLLSKAVVRESTQVSPSATPSAAASESGLTRRTDGTDSSAVAGGGAYNRKGKAVAMTAPAIEITGTSSSVVSKSEI

EPEKTNVDDRKRKEREATTTDETESRSEETKQARVSTTSTKRSRAAEVHNLSERKRRDRINERMKALQELIPRCNKSDKA

SMLDEAIEYMKSLQLQIQMMSMGCGMMPMMYPGMQQYMPHMAMGMGMNQPIPPPSFMPFPNMLAAQRPLPTQTHMAGSGP

QYPVHASDPSRVFVPNQQYDPTSGQPQYPAGYTDPYQQFRGLHPTQPPQFQNQATSYPSSSRVSSSKESEDHGNHTTG

>At2G22750

MNSLVGDVPQSLSSLDDTTTCYNLDASCNKSLVEERPSKILKTTHISPNLHPFSSSNPPPPKHQPSSRILSFEKTGLHVM

NHNSPNLIFSPKDEEIGLPEHKKAELIIRGTKRAQSLTRSQSNAQDHILAERKRREKLTQRFVALSALIPGLKKMDKASV

LGDAIKHIKYLQESVKEYEEQKKEKTMESVVLVKKSSLVLDENHQPSSSSSSDGNRNSSSSNLPEIEVRVSGKDVLIKIL

CEKQKGNVIKIMGEIEKLGLSITNSNVLPFGPTFDISIIAQKNNNFDMKIEDVVKNLSFGLSKLT

>At2G22760

MDEDFFLPDFSLVDIDFDFNIYEENNLSPDESLSNSRRADQSSKFDHQMHFECLREKPKAAVKPMMKINNKQQLISFDFS

SNVISSPAAEEIIMDKLVGRGTKRKTCSHGTRSPVLAKEHVLAERKRREKLSEKFIALSALLPGLKKADKVTILDDAISR

MKQLQEQLRTLKEEKEATRQMESMILVKKSKVFFDEEPNLSCSPSVHIEFDQALPEIEAKISQNDILIRILCEKSKGCMI

NILNTIENFQLRIENSIVLPFGDSTLDITVLAQMDKDFSMSILKDLVRNLRLAMV

>At2G22770

MDDSSFMDLMIDTDEYLIDDWESDFPICGETNTNPGSESGSGTGFELLAERPTKQMKTNNNMNSTSSSPSSSSSSGSRTS

QVISFGSPDTKTNPVETSLNFSNQVSMDQKVGSKRKDCVNNGGRREPHLLKEHVLAERKRRQKLNERLIALSALLPGLKK

TDKATVLEDAIKHLKQLQERVKKLEEERVVTKKMDQSIILVKRSQVYLDDDSSSYSSTCSAASPLSSSSDEVSIFKQTMP

MIEARVSDRDLLIRVHCEKNKGCMIKILSSLEKFRLEVVNSFTLPFGNSTLVITILTKMDNKFSRPVEEVVKNIRVALAE

>At2G24260

MMNSSLLTPSSSSSSHIQTPSTTFDHEDFLDQIFSSAPWPSVVDDAHPLPSDGFHGHDVDSRNQPIMMMPLNDGSSVHAL

YNGFSVAGSLPNFQIPQGSGGGLMNQQGQTQTQTQPQASASTATGGTVAAPPQSRTKIRARRGQATDPHSIAERLRRERI

AERMKALQELVPNGNKTDKASMLDEIIDYVKFLQLQVKVLSMSRLGGAASVSSQISEAGGSHGNASSAMVGGSQTAGNSN

DSVTMTEHQVAKLMEEDMGSAMQYLQGKGLCLMPISLATAISTATCHSRNPLIPGAVADVGGPSPPNLSGMTIQSTSTKM

GSGNGKLNGNGVTERSSSIAVKEAVSVSKA

>At2G28160

MEGRVNALSNINDLELHNFLVDPNFDQFINLIRGDHQTIDENPVLDFDLGPLQNSPCFIDENQFIPTPVDDLFDELPDLD

SNVAESFRSFDGDSVRAGGEEDEEDYNDGDDSSATTTNNDGTRKTKTDRSRTLISERRRRGRMKDKLYALRSLVPNITKM

DKASIVGDAVLYVQELQSQAKKLKSDIAGLEASLNSTGGYQEHAPDAQKTQPFRGINPPASKKIIQMDVIQVEEKGFYVR

LVCNKGEGVAPSLYKSLESLTSFQVQNSNLSSPSPDTYLLTYTLDGTCFEQSLNLPNLKLWITGSLLNQGFEFIKSFT

>At2G31210

MYEESSCFDPNSMVDNNGGFCAAETTFTVSHQFQPPLGSTTNSFDDDLKLPTMDEFSVFPSVISLPNSETQNQNISNNNH

LINQMIQESNWGVSEDNSNFFMNTSHPNTTTTPIPDLLSLLHLPRCSMSLPSSDIMAGSCFTYDPLFHLNLPPQPPLIPS

NDYSGYLLGIDTNTTTQRDESNVGDENNNAQFDSGIIEFSKEIRRKGRGKRKNKPFTTERERRCHLNERYEALKLLIPSP

SKGDRASILQDGIDYINELRRRVSELKYLVERKRCGGRHKNNEVDDNNNNKNLDDHGNEDDDDDDENMEKKPESDVIDQC

SSNNSLRCSWLQRKSKVTEVDVRIVDDEVTIKVVQKKKINCLLLVSKVLDQLQLDLHHVAGGQIGEHYSFLFNTKIYEGS

TIYASAIANRVIEVVDKHYMASLPNSNY

>At2G31220

MEEERESLYEEMGCFDPNTPAEVTVESSFSQAEPPPPPPQVLVAGSTSNSNCSVEVEELSEFHLSPQDCPQASSTPLQFH

INPPPPPPPPCDQLHNNLIHQMASHQQQHSNWDNGYQDFVNLGPNSATTPDLLSLLHLPRCSLPPNHHPSSMLPTSFSDI

MSSSSAAAVMYDPLFHLNFPMQPRDQNQLRNGSCLLGVEDQIQMDANGGMNVLYFEGANNNNGGFENEILEFNNGVTRKG

RGSRKSRTSPTERERRVHFNDRFFDLKNLIPNPTKIDRASIVGEAIDYIKELLRTIEEFKMLVEKKRCGRFRSKKRARVG

EGGGGEDQEEEEDTVNYKPQSEVDQSCFNKNNNNSLRCSWLKRKSKVTEVDVRIIDDEVTIKLVQKKKINCLLFTTKVLD

QLQLDLHHVAGGQIGEHYSFLFNTKICEGSCVYASGIADTLMEVVEKQYMEAVPSNGY

>At2G40200

MENSYDSSKWSDSTTPYMVSWSLQSESSDSDWNRFNLGFSSSSFGGNFPADDCVGGIEKAESLSRSHRLAEKRRRDRINS

HLTALRKLVPNSDKLDKAALLATVIEQVKELKQKAAESPIFQDLPTEADEVTVQPETISDFESNTNTIIFKASFCCEDQP

EAISEIIRVLTKLQLETIQAEIISVGGRMRINFILKDSNCNETTNIAASAKALKQSLCSALNRITSSSTTTSSVCRIRSK

RQRWFLSSHYSHNE

>At2G41130

MQPETSDQMLYSFLAGNEVGGGGYCVSGDYMTTMQSLCGSSSSTSSYYPLAISGIGETMAQDRALAALRNHKEAERRRRE

RINSHLNKLRNVLSCNSKTDKATLLAKVVQRVRELKQQTLETSDSDQTLLPSETDEISVLHFGDYSNDGHIIFKASLCCE

DRSDLLPDLMEILKSLNMKTLRAEMVTIGGRTRSVLVVAADKEMHGVESVHFLQNALKSLLERSSKSLMERSSGGGGGER

SKRRRALDHIIMV

>At2G41240

MCALVPPLYPNFGWPCGDHSFYETDDVSNTFLDFPLPDLTVTHENVSSENNRTLLDNPVVMKKLNHNASERERRKKINTM

FSSLRSCLPPTNQTKKLSVSATVSQALKYIPELQEQVKKLMKKKEELSFQISGQRDLVYTDQNSKSEEGVTSYASTVSST

RLSETEVMVQISSLQTEKCSFGNVLSGVEEDGLVLVGASSSRSHGERLFYSMHLQIKNGQVNSEELGDRLLYLYEKCGHS

FT

>At2G42300

MDLTQGFRARSGVVGPVAGLESLNFSDEFRHLVTTMPPETTGGSFTALLEMPVTQAMELLHFPDSSSSQARTVTSGDISP

TTLHPFGALTFPSNSLLLDRAARFSVIATEQNGNFSGETANSLPSNPGANLDRVKAEPAETDSMVENQNQSYSSGKRKER

EKKVKSSTKKNKSSVESDKLPYVHVRARRGQATDNHSLAERARREKINARMKLLQELVPGCDKIQGTALVLDEIINHVQT

LQRQVEMLSMRLAAVNPRIDFNLDSILASENGSLMDGSFNAESYHQLQQWPFDGYHQPEWGREEDHHQANFSMGSATLHP

NQVKMEL

>At2G43010

MEHQGWSFEENYSLSTNRRSIRPQDELVELLWRDGQVVLQSQTHREQTQTQKQDHHEEALRSSTFLEDQETVSWIQYPPD

EDPFEPDDFSSHFFSTMDPLQRPTSETVKPKSSPEPPQVMVKPKACPDPPPQVMPPPKFRLTNSSSGIRETEMEQYSVTT

VGPSHCGSNPSQNDLDVSMSHDRSKNIEEKLNPNASSSSGGSSGCSFGKDIKEMASGRCITTDRKRKRINHTDESVSLSD

AIGNKSNQRSGSNRRSRAAEVHNLSERRRRDRINERMKALQELIPHCSKTDKASILDEAIDYLKSLQLQLQVMWMGSGMA

AAAASAPMMFPGVQPQQFIRQIQSPVQLPRFPVMDQSAIQNNPGLVCQNPVQNQIISDRFARYIGGFPHMQAATQMQPME

MLRFSSPAGQQSQQPSSVPTKTTDGSRLDH

>At2G46510

MNMSDLGWDDEDKSVVSAVLGHLASDFLRANSNSNQNLFLVMGTDDTLNKKLSSLVDWPNSENFSWNYAIFWQQTMSRSG

QQVLGWGDGCCREPNEEEESKVVRSYNFNNMGAEEETWQDMRKRVLQKLHRLFGGSDEDNYALSLEKVTATEIFFLASMY

FFFNHGEGGPGRCYSSGKHVWLSDAVNSESDYCFRSFMAKSAGIRTIVMVPTDAGVLELGSVWSLPENIGLVKSVQALFM

RRVTQPVMVTSNTNMTGGIHKLFGQDLSGAHAYPKKLEVRRNLDERFTPQSWEGYNNNKGPTFGYTPQRDDVKVLENVNM

VVDNNNYKTQIEFAGSSVAASSNPSTNTQQEKSESCTEKRPVSLLAGAGIVSVVDEKRPRKRGRKPANGREEPLNHVEAE

RQRREKLNQRFYALRSVVPNISKMDKASLLGDAISYIKELQEKVKIMEDERVGTDKSLSESNTITVEESPEVDIQAMNEE

VVVRVISPLDSHPASRIIQAMRNSNVSLMEAKLSLAEDTMFHTFVIKSNNGSDPLTKEKLIAAFYPETSSTQPPLPSSSS

QVSGDI

>At2G46810

MFVLRVSNQSFKLHQQVQCKDEIFCLDQKVNVRRSLQVQETVEDHQSFALEEEEQQLSTPSLLQDTTIPFLQMLQQSEDP

SPFLSFKDPSFLALLSLQTLEKPWELENYLPHEVPEFHSPIHSETNHYYHNPSLEGVNEAISNQELPFNPLENARSRRKR

KNNNLASLMTREKRKRRRTKPTKNIEEIESQRMTHIAVERNRRRQMNVHLNSLRSIIPSSYIQRGDQASIVGGAIDFVKI

LEQQLQSLEAQKRSQQSDDNKEQIPEDNSLRNISSNKLRASNKEEQSSKLKIEATVIESHVNLKIQCTRKQGQLLRSIIL

LEKLRFTVLHLNITSPTNTSVSYSFNLKMEDECNLGSADEITAAIRQIFDS

>At2G46970

MEAKPLASSSSEPNMISPSSNIKPKLKDEDYMELVCENGQILAKIRRPKNNGSFQKQRRQSLLDLYETEYSEGFKKNIKI

LGDTQVVPVSQSKPQQDKETNEQMNNNKKKLKSSKIEFERNVSKSNKCVESSTLIDVSAKGPKNVEVTTAPPDEQSAAVG

RSTELYFASSSKFSRGTSRDLSCCSLKRKYGDIEEEESTYLSNNSDDESDDAKTQVHARTRKPVTKRKRSTEVHKLYERK

RRDEFNKKMRALQDLLPNCYKDDKASLLDEAIKYMRTLQLQVQMMSMGNGLIRPPTMLPMGHYSPMGLGMHMGAAATPTS

IPQFLPMNVQATGFPGMNNAPPQMLSFLNHPSGLIPNTPIFSPLENCSQPFVVPSCVSQTQATSFTQFPKSASASNLEDA

MQYRGSNGFSYYRSPN

>At3G06120

MSHIAVERNRRRQMNEHLKSLRSLTPCFYIKRGDQASIIGGVIEFIKELQQLVQVLESKKRRKTLNRPSFPYDHQTIEPS

SLGAATTRVPFSRIENVMTTSTFKEVGACCNSPHANVEAKISGSNVVLRVVSRRIVGQLVKIISVLEKLSFQVLHLNISS

MEETVLYFFVVKIGLECHLSLEELTLEVQKSFVSDEVIVSTN

>At3G07340

MENELFMNAGVSHPPVMTSPSSSSAMLKWVSMETQPVDPSLSRNLFWEKSTEQSIFDSALSSLVSSPTPSNSNFSVGGVG

GENVIMRELIGKLGNIGDIYGITASNGNSCYATPMSSPPPGSMMETKTTTPMAELSGDPGFAERAARFSCFGSRSFNSRT

NSPFPINNEPPITTNEKMPRVSSSPVFKPLASHVPAGESSGELSRKRKTKSKQNSPSAVSSSKEIEEKEDSDPKRCKKSE

ENGDKTKSIDPYKDYIHVRARRGQATDSHSLAERVRREKISERMKLLQDLVPGCNKVTGKALMLDEIINYVQSLQRQVEF

LSMKLSSVNTRLDFNMDALLSKDIFPSSNNLMHHQQVLQLDSSAETLLGDHHNKNLQLNPDISSNNVINPLETSETRSFI

SHLPTLAHFTDSISQYSTFSEDDLHSIIHMGFAQNRLQELNQGSSNQVPSHMKAEL

>At3G19860

MGIRENGIMLVSRERERARRLENRESIFAEPPCLLLAHRISPSPSILPAEEEVMDVSARKSQKAGREKLRREKLNEHFVE

LGNVLDPERPKNDKATILTDTVQLLKELTSEVNKLKSEYTALTDESRELTQEKNDLREEKTSLKSDIENLNLQYQQRLRS

MSPWGAAMDHTVMMAPPPSFPYPMPIAMPPGSIPMHPSMPSYTYFGNQNPSMIPAPCPTYMPYMPPNTVVEQQSVHIPQN

PGNRSREPRAKVSRESRSEKAEDSNEVATQLELKTPGSTSDKDTLQRPEKTKRCKRNNNNNSIEESSHSSKCSSSPSVRD

HSSSSSVAGGQKPDDAK

>At3G23690

MNMDKETEQTLNYLPLGQSDPFGNGNEGTIGDFLGRYCNNPQEISPLTLQSFSLNSQISENFPISGGIRFPPYPGQFGSD

REFGSQPTTQESNKSSLLDPDSVSDRVHTTKSNSRKRKSIPSGNGKESPASSSLTASNSKVSGENGGSKGGKRSKQDVAG

SSKNGVEKCDSKGDNKDDAKPPEAPKDYIHVRARRGQATDSHSLAERARREKISERMTLLQDLVPGCNRITGKAVMLDEI

INYVQSLQRQVEFLSMKLATVNPRMEFNANASLSTEMIQPGESLTQSLYAMACSEQRLPSAYYSLGKNMPRFSDTQFPSN

DGFVHTETPGFWENNDLQSIVQMGFGDILQQQSNNNNNNCSEPTLQMKLEP

>At3G24140

MDKDYSAPNFLGESSGGNDDNSSGMIDYMFNRNLQQQQKQSMPQQQQHQLSPSGFGATPFDKMNFSDVMQFADFGSKLAL

NQTRNQDDQETGIDPVYFLKFPVLNDKIEDHNQTQHLMPSHQTSQEGGECGGNIGNVFLEEKEDQDDDNDNNSVQLRFIG

GEEEDRENKNVTKKEVKSKRKRARTSKTSEEVESQRMTHIAVERNRRKQMNEHLRVLRSLMPGSYVQRGDQASIIGGAIE

FVRELEQLLQCLESQKRRRILGETGRDMTTTTTSSSSPITTVANQAQPLIITGNVTELEGGGGLREETAENKSCLADVEV

KLLGFDAMIKILSRRRPGQLIKTIAALEDLHLSILHTNITTMEQTVLYSFNVKITSETRFTAEDIASSIQQIFSFIHANT

NISGSSNLGNIVFT

>At3G25710

MYAMKEEDCLQTFHNLQDYQDQFHLHHHPQILPWSSTSLPSFDPLHFPSNPTRYSDPVHYFNRRASSSSSSFDYNDGFVS

PPPSMDHPQNHLRILSEALGPIMRRGSSFGFDGEIMGKLSAQEVMDAKALAASKSHSEAERRRRERINTHLAKLRSILPN

TTKTDKASLLAEVIQHMKELKRQTSQITDTYQVPTECDDLTVDSSYNDEEGNLVIRASFCCQDRTDLMHDVINALKSLRL

RTLKAEIATVGGRVKNILFLSREYDDEEDHDSYRRNFDGDDVEDYDEERMMNNRVSSIEEALKAVIEKCVHNNDESNDNN

NLEKSSSGGIKRQRTSKMVNRCYN

>At3G26744

MGLDGNNGGGVWLNGGGGEREENEEGSWGRNQEDGSSQFKPMLEGDWFSSNQPHPQDLQMLQNQPDFRYFGGFPFNPNDN

LLLQHSIDSSSSCSPSQAFSLDPSQQNQFLSTNNNKGCLLNVPSSANPFDNAFEFGSESGFLNQIHAPISMGFGSLTQLG

NRDLSSVPDFLSARSLLAPESNNNNTMLCGGFTAPLELEGFGSPANGGFVGNRAKVLKPLEVLASSGAQPTLFQKRAAMR

QSSGSKMGNSESSGMRRFSDDGDMDETGIEVSGLNYESDEINESGKAAESVQIGGGGKGKKKGMPAKNLMAERRRRKKLN

DRLYMLRSVVPKISKMDRASILGDAIDYLKELLQRINDLHNELESTPPGSLPPTSSSFHPLTPTPQTLSCRVKEELCPSS

LPSPKGQQARVEVRLREGRAVNIHMFCGRRPGLLLATMKALDNLGLDVQQAVISCFNGFALDVFRAEQCQEGQEILPDQI

KAVLFDTAGYAGMI

>At3G56770

MQPEVSDQIFYAFLTGGLCASSTSTTVTSSSDPFATVYEDKALASLRNHKEAERKRRARINSHLNKLRKLLSCNSKTDKS

TLLAKVVQRVKELKQQTLEITDETIPSETDEISVLNIEDCSRGDDRRIIFKVSFCCEDRPELLKDLMETLKSLQMETLFA

DMTTVGGRTRNVLVVAADKEHHGVQSVNFLQNALKSLLERSSKSVMVGHGGGGGEERLKRRRALDHIIMV

>At3G56970

MCALVPSFFTNFGWPSTNQYESYYGAGDNLNNGTFLELTVPQTYEVTHHQNSLGVSVSSEGNEIDNNPVVVKKLNHNASE

RDRRKKINTLFSSLRSCLPASDQSKKLSIPETVSKSLKYIPELQQQVKRLIQKKEEILVRVSGQRDFELYDKQQPKAVAS

YLSTVSATRLGDNEVMVQVSSSKIHNFSISNVLGGIEEDGFVLVDVSSSRSQGERLFYTLHLQVENMDDYKINCEELSER

MLYLYEKCENSFN

>At3G56980

MCALVPPLFPNFGWPSTGEYDSYYLAGDILNNGGFLDFPVPEETYGAVTAVTQHQNSFGVSVSSEGNEIDNNPVVVKKLN

HNASERDRRRKINSLFSSLRSCLPASGQSKKLSIPATVSRSLKYIPELQEQVKKLIKKKEELLVQISGQRNTECYVKQPP

KAVANYISTVSATRLGDNEVMVQISSSKIHNFSISNVLSGLEEDRFVLVDMSSSRSQGERLFYTLHLQVEKIENYKLNCE

ELSQRMLYLYEECGNSYI

>At3G61950

MERFQGHINPCFFDRKPDVRSLEVQGFAEAQSFAFKEKEEESLQDTVPFLQMLQSEDPSSFFSIKEPNFLTLLSLQTLKE

PWELERYLSLEDSQFHSPVQSETNRFMEGANQAVSSQEIPFSQANMTLPSSTSSPLSAHSRRKRKINHLLPQEMTREKRK

RRKTKPSKNNEEIENQRINHIAVERNRRRQMNEHINSLRALLPPSYIQRGDQASIVGGAINYVKVLEQIIQSLESQKRTQ

QQSNSEVVENALNHLSGISSNDLWTTLEDQTCIPKIEATVIQNHVSLKVQCEKKQGQLLKGIISLEKLKLTVLHLNITTS

SHSSVSYSFNLKMEDECDLESADEITAAVHRIFDIPTI

>At4G00050

MSQCVPNCHIDDTPAAATTTVRSTTAADIPILDYEVAELTWENGQLGLHGLGPPRVTASSTKYSTGAGGTLESIVDQATR

LPNPKPTDELVPWFHHRSSRAAMAMDALVPCSNLVHEQQSKPGGVGSTRVGSCSDGRTMGGGKRARVAPEWSGGGSQRLT

MDTYDVGFTSTSMGSHDNTIDDHDSVCHSRPQMEDEEEKKAGGKSSVSTKRSRAAAIHNQSERKRRDKINQRMKTLQKLV

PNSSKTDKASMLDEVIEYLKQLQAQVSMMSRMNMPSMMLPMAMQQQQQLQMSLMSNPMGLGMGMGMPGLGLLDLNSMNRA

AASAPNIHANMMPNPFLPMNCPSWDASSNDSRFQSPLIPDPMSAFLACSTQPTTMEAYSRMATLYQQMQQQLPPPSNPK

>At4G00120

MENGMYKKKGVCDSCVSSKSRSNHSPKRSMMEPQPHHLLMDWNKANDLLTQEHAAFLNDPHHLMLDPPPETLIHLDEDEE

YDEDMDAMKEMQYMIAVMQPVDIDPATVPKPNRRNVRISDDPQTVVARRRRERISEKIRILKRIVPGGAKMDTASMLDEA

IRYTKFLKRQVRILQPHSQIGAPMANPSYLCYYHNSQP

>At4G00870

MYNLTFSPSLSSSLLSFTQQTPAAIVSSSPPDLVLQQKLRFVVETSPDRWAYVIFWQKMFDDQSDRSYLVWVDGHFCGNK

NNNSQENYTTNSIECELMMDGGDDLELFYAASFYGEDRSPRKEVSDESLVWLTGPDELRFSNYERAKEAGFHGVHTLVSI

PINNGIIELGSSESIIQNRNFINRVKSIFGSGKTTKHTNQTGSYPKPAVSDHSKSGNQQFGSERKRRRKLETTRVAAATK

EKHHPAVLSHVEAEKQRREKLNHRFYALRAIVPKVSRMDKASLLSDAVSYIESLKSKIDDLETEIKKMKMTETDKLDNSS

SNTSPSSVEYQVNQKPSKSNRGSDLEVQVKIVGEEAIIRVQTENVNHPTSALMSALMEMDCRVQHANASRLSQVMVQDVV

VLVPEGLRSEDRLRTTLVRTLSL

>At4G01460

MSGLMSFGELEDQFGQISDTTMEEKIPFLQMLQCIEHPFTTTEPNQFLQSLLQIQTLESKSCLTLETNIKRDPGQTDDPE

KDPRTENGAVTVKEKRKRKRTRAPKNKDEVENQRMTHIAVERNRRRQMNEHLNSLRSLMPPSFLQRGDQASIVGGAIDFI

KELEQLLQSLEAEKRKDGTDETPKTASCSSSSSLACTNSSISSVSTTSENGFTARFGGGDTTEVEATVIQNHVSLKVRCK

RGKRQILKAIVSIEELKLAILHLTISSSFDFVIYSFNLKMEDGCKLGSADEIATAVHQIFEQINGEVMWSNLSRT

>At4G02590

MASNNPHDNLSDQTPSDDFFEQILGLPNFSASSAAGLSGVDGGLGGGAPPMMLQLGSGEEGSHMGGLGGSGPTGFHNQMF

PLGLSLDQGKGPGFLRPEGGHGSGKRFSDDVVDNRCSSMKPVFHGQPMQQPPPSAPHQPTSIRPRVRARRGQATDPHSIA

ERLRRERIAERIRALQELVPTVNKTDRAAMIDEIVDYVKFLRLQVKVLSMSRLGGAGAVAPLVTDMPLSSSVEDETGEGG

RTPQPAWEKWSNDGTERQVAKLMEENVGAAMQLLQSKALCMMPISLAMAIYHSQPPDTSSVVKPENNPPQ

>At4G09180

MQPTSVGSSGGGDDGGGRGGGGGLSRSGLSRIRSAPATWLEALLEEDEEESLKPNLGLTDLLTGNSNDLPTSRGSFEFPI

PVEQGLYQQGGFHRQNSTPADFLSGSDGFIQSFGIQANYDYLSGNIDVSPGSKRSREMEALFSSPEFTSQMKGEQSSGQV

PTGVSSMSDMNMENLMEDSVAFRVRAKRGCATHPRSIAERVRRTRISDRIRKLQELVPNMDKQTNTADMLEEAVEYVKVL

QRQIQELTEEQKRCTCIPKEEQ

>At4G09820

MDESSIIPAEKVAGAEKKELQGLLKTAVQSVDWTYSVFWQFCPQQRVLVWGNGYYNGAIKTRKTTQPAEVTAEEAALERS

QQLRELYETLLAGESTSEARACTALSPEDLTETEWFYLMCVSFSFPPPSGMPGKAYARRKHVWLSGANEVDSKTFSRAIL

AKSAKIQTVVCIPMLDGVVELGTTKKVREDVEFVELTKSFFYDHCKTNPKPALSEHSTYEVHEEAEDEEEVEEEMTMSEE

MRLGSPDDEDVSNQNLHSDLHIESTHTLDTHMDMMNLMEEGGNYSQTVTTLLMSHPTSLLSDSVSTSSYIQSSFATWRVE

NGKEHQQVKTAPSSQWVLKQMIFRVPFLHDNTKDKRLPREDLSHVVAERRRREKLNEKFITLRSMVPFVTKMDKVSILGD

TIAYVNHLRKRVHELENTHHEQQHKRTRTCKRKTSEEVEVSIIENDVLLEMRCEYRDGLLLDILQVLHELGIETTAVHTS

VNDHDFEAEIRAKVRGKKASIAEVKRAIHQVIIHDTNL

>At4G16430

MGQKFWENQEDRAMVESTIGSEACDFFISTASASNTALSKLVSPPSDSNLQQGLRHVVEGSDWDYALFWLASNVNSSDGC

VLIWGDGHCRVKKGASGEDYSQQDEIKRRVLRKLHLSFVGSDEDHRLVKSGALTDLDMFYLASLYFSFRCDTNKYGPAGT

YVSGKPLWAADLPSCLSYYRVRSFLARSAGFQTVLSVPVNSGVVELGSLRHIPEDKSVIEMVKSVFGGSDFVQAKEAPKI

FGRQLSLGGAKPRSMSINFSPKTEDDTGFSLESYEVQAIGGSNQVYGYEQGKDETLYLTDEQKPRKRGRKPANGREEALN

HVEAERQRREKLNQRFYALRAVVPNISKMDKASLLADAITYITDMQKKIRVYETEKQIMKRRESNQITPAEVDYQQRHDD

AVVRLSCPLETHPVSKVIQTLRENEVMPHDSNVAITEEGVVHTFTLRPQGGCTAEQLKDKLLASLSQ

>At4G17880

MSPTNVQVTDYHLNQSKTDTTNLWSTDDDASVMEAFIGGGSDHSSLFPPLPPPPLPQVNEDNLQQRLQALIEGANENWTY

AVFWQSSHGFAGEDNNNNNTVLLGWGDGYYKGEEEKSRKKKSNPASAAEQEHRKRVIRELNSLISGGVGGGDEAGDEEVT

DTEWFFLVSMTQSFVKGTGLPGQAFSNSDTIWLSGSNALAGSSCERARQGQIYGLQTMVCVATENGVVELGSSEIIHQSS

DLVDKVDTFFNFNNGGGEFGSWAFNLNPDQGENDPGLWISEPNGVDSGLVAAPVMNNGGNDSTSNSDSQPISKLCNGSSV

ENPNPKVLKSCEMVNFKNGIENGQEEDSSNKKRSPVSNNEEGMLSFTSVLPCDSNHSDLEASVAKEAESNRVVVEPEKKP

RKRGRKPANGREEPLNHVEAERQRREKLNQRFYSLRAVVPNVSKMDKASLLGDAISYISELKSKLQKAESDKEELQKQID

VMNKEAGNAKSSVKDRKCLNQESSVLIEMEVDVKIIGWDAMIRIQCSKRNHPGAKFMEALKELDLEVNHASLSVVNDLMI

QQATVKMGNQFFTQDQLKVALTEKVGECP

>At4G21330

MGGGSRFQEPVRMSRRKQVTKEKEEDENFKSPNLEAERRRREKLHCRLMALRSHVPIVTNMTKASIVEDAITYIGELQNN

VKNLLETFHEMEEAPPEIDEEQTDPMIKPEVETSDLNEEMKKLGIEENVQLCKIGERKFWLKIITEKRDGIFTKFMEVMR

FLGFEIIDISLTTSNGAILISASVQTQELCDVEQTKDFLLEVMRSNP

>At4G28790

MTWKPKMLILSHDLISPEKYIMGEDDIVELLGKSSQVVTSSQTQTPSCDPPLILRGSGSGDGEGNGPLPQPPPPLYHQQS

LFIQEDEMASWLHQPNRQDYLYSQLLYSGVASTHPQSLASLEPPPPPRAQYILAADRPTGHILAERRAENFMNISRQRGN

IFLGGVEAVPSNSTLLSSATESIPATHGTESRATVTGGVSRTFAVPGLGPRGKAVAIETAGTQSWGLCKAETEPVQRQPA

TETDITDERKRKTREETNVENQGTEEARDSTSSKRSRAAIMHKLSERRRRQKINEMMKALQELLPRCTKTDRSSMLDDVI

EYVKSLQSQIQMFSMGHVMIPPMMYAGNIQQQYMPHMAMGMNRPPAFIPFPRQAHMAEGVGPVDLFRENEETEQETMSLL

LREDKRTKQKMFS

>At4G28800

MQFDESDARVWIWFEIRREDDIVELLWQSGQVVGTNQTHRQSYDPPPILRGSGSGRGEENAPLSQPPPHLHQQNLFIQEG

EMYSWLHHSYRQNYFCSELLNSTPATHPQSSISLAPRQTIATRRAENFMNFSWLRGNIFTGGRVDEAGPSFSVVRESMQV

GSNTTPPSSSATESCVIPATEGTASRVSGTLAAHDLGRKGKAVAVEAAGTPSSGVCKAETEPVQIQPATESKLKAREETH

GTEEARGSTSRKRSRTAEMHNLAERRRREKINEKMKTLQQLIPRCNKSTKVSTLDDAIEYVKSLQSQIQGMMSPMMNAGN

TQQFMPHMAMDMNRPPPFIPFPGTSFPMPAQMAGVGPSYPAPRYPFPNIQTFDPSRVRLPSPQPNPVSNQPQFPAYMNPY

SQFAGPHQLQQPPPPPFQGQTTSQLSSGQASSSKEPEDQENQPTA

>At4G28815

MMIISSQILLLFGFKLFFETRGEDDIVELLCKIGQTQIPSSDPLPILRGSGSGGREENTPLPPPLPHQNLFIQEDEMSSW

PHHPLRQDYLCSELYASTPAPHPQSSVSLAPPPPKPPSSAPYGQIIAPRSAPRIQGTEEARGSTSRKRSRAAEMHNLAER

RRREKINERMKTLQQLIPRCNKSTKVSMLEDVIEYVKSLEMQINQFMPHMAMGMNQPPAYIPFPSQAHMAGVGPSYPPPR

YPFPNIQTFDPSRVWLQSPQPNPVSNQPQMNPYGQFVGHHQMQQSLPPPLQVILSQYPLCLFLCSNK

>At4G29930

MEDLDHEYKNYWETTMFFQNQELEFDSWPMEEAFSGSGESSSPDGAATSPASSKNVVSERNRRQKLNQRLFALRSVVPNI

SKLDKASVIKDSIDYMQELIDQEKTLEAEIRELESRSTLLENPVRDYDCNFAETHLQDFSDNNDMRSKKFKQMDYSTRVQ

HYPIEVLEMKVTWMGEKTVVVCITCSKKRETMVQLCKVLESLNLNILTTNFSSFTSRLSTTLFLQVTLSLSPSLISLFGN

VITSTNYKILNASREYCTCLVLV

>At4G30980

MNSSSLLTPSSSPSPHLQSPATFDHDDFLHHIFSSTPWPSSVLDDTPPPTSDCAPVTGFHHHDADSRNQITMIPLSHNHP

NDALFNGFSTGSLPFHLPQGSGGQTQTQSQATASATTGGATAQPQTKPKVRARRGQATDPHSIAERLRRERIAERMKSLQ

ELVPNGNKTDKASMLDEIIDYVKFLQLQVKVLSMSRLGGAASASSQISEDAGGSHENTSSSGEAKMTEHQVAKLMEEDMG

SAMQYLQGKGLCLMPISLATTISTATCPSRSPFVKDTGVPLSPNLSTTIVANGNGSSLVTVKDAPSVSKP

>At4G33880

MEAMGEWSNNLGGMYTYATEEADFMNQLLASYDHPGTGSSSGAAASGDHQGLYWNLGSHHNHLSLVSEAGSFCFSQESSS

YSAGNSGYYTVVPPTVEENQNETMDFGMEDVTINTNSYLVGEETSECDVEKYSSGKTLMPLETVVENHDDEESLLQSEIS

VTTTKSLTGSKKRSRATSTDKNKRARVNKRAQKNVEMSGDNNEGEEEEGETKLKKRKNGAMMSRQNSSTTFCTEEESNCA

DQDGGGEDSSSKEDDPSKALNLNGKTRASRGAATDPQSLYARKRRERINERLRILQNLVPNGTKVDISTMLEEAVHYVKF

LQLQIKLLSSDDLWMYAPIAFNGMDIGLSSPR

>At4G34530

MNGAIGGDLLLNFPDMSVLERQRAHLKYLNPTFDSPLAGFFADSSMITGGEMDSYLSTAGLNLPMMYGETTVEGDSRLSI

SPETTLGTGNFKKRKFDTETKDCNEKKKKMTMNRDDLVEEGEEEKSKITEQNNGSTKSIKKMKHKAKKEENNFSNDSSKV

TKELEKTDYIHVRARRGQATDSHSIAERVRREKISERMKFLQDLVPGCDKITGKAGMLDEIINYVQSLQRQIEFLSMKLA

IVNPRPDFDMDDIFAKEVASTPMTVVPSPEMVLSGYSHEMVHSGYSSEMVNSGYLHVNPMQQVNTSSDPLSCFNNGEAPS

MWDSHVQNLYGNLGV

>At4G36540

MDLSVLDRLKWLQQQQMVSPEFLQILGSDGREELKRVESYLGNNNDELQSFRHFPEFGPDYDTTDGCISRTSSFHMEPVK

NNGHSRAITLQNKRKPEGKTEKREKKKIKAEDETEPSMKGKSNMSNTETSSEIQKPDYIHVRARRGEATDRHSLAERARR

EKISKKMKCLQDIVPGCNKVTGKAGMLDEIINYVQSLQQQVEFLSMKLSVINPELECHIDDLSAKQFQAYFTGPPEGDSK

QSIMADFRSFPLHQQGSLDYSVINSDHTTSLGAKDHTSSSWETHSQCLYNSLRTDSVSNFFSLK

>At4G36930

MISQREEREEKKQRVMGDKKLISSSSSSSVYDTRINHHLHHPPSSSDEISQFLRHIFDRSSPLPSYYSPATTTTTASLIG

VHGSGDPHADNSRSLVSHHPPSDSVLMSKRVGDFSEVLIGGGSGSAAACFGFSGGGNNNNVQGNSSGTRVSSSSVGASGN

ETDEYDCESEEGGEAVVDEAPSSKSGPSSRSSSKRCRAAEVHNLSEKRRRSRINEKMKALQSLIPNSNKTDKASMLDEAI

EYLKQLQLQVQMLTMRNGINLHPLCLPGTTLHPLQLSQIRPPEATNDPLLNHTNQFASTSNAPEMINTVASSYALEPSIR

SHFGPFPLLTSPVEMSREGGLTHPRLNIGHSNANITGEQALFDGQPDLKDRIT

>At4G37850

MSILSTRWFSEQEIEENSIIQQFHMNSIVGEVQEAQYIFPHSFTTNNDPSYDDLIEMKPPKILETTYISPSSHLPPNSKP

HHIHRHSSSRILSFEDYGSNDMEHEYSPTYLNSIFSPKLEAQVQPHQKSDEFNRKGTKRAQPFSRNQSNAQDHIIAERKR

REKLTQRFVALSALVPGLKKMDKASVLGDALKHIKYLQERVGELEEQKKERRLESMVLVKKSKLILDDNNQSFSSSCEDG

FSDLDLPEIEVRFSDEDVLIKILCEKQKGHLAKIMAEIEKLHILITNSSVLNFGPTLDITIIAKKESDFDMTLMDVVKSL

RSALSNFI

>At4G38070

MEKVYEELDEVKAVNEKLRIDYRNKTELLENLKKVQNEQLIEIREARLVNEKHGFEIEEKSREIAELKRANEELQRCLRE

KDSVVKRVNDVNDKLRANGEDKYREFEEEKRNMMSGLDEASEKNIDLEQKNNVYRAEIEGLKGLLAVAETKRIEAEKTVK

GMKEMRGRDDVVVKMEEEKSQVEEKLKWKKEQFKHLEEAYEKLKNLFKDSKKEWEEEKSKLLDEIYSLQTKLDSVTRISE

DLQKKLQMCNGALTQEETRRKHLEIQVSEFKAKYEDAFAECQDARTQLDDLAGKRDWEVAELRQTLSMKDAYFKEMKYEN

GKLEQENRELLGSLKELQEATIQGSGNSALSKLKNKFRNLENIHKNCSANLRSKEAEWSSQVEKMVEEINDYKLQLQSKE

AALKEVELELENCRSSTAKMRLQYEEISIMFLVLSRTVSEAQSRLANAKDKQIKDEKREGNCYSLLMEQLDQKNAALAKA

QMEIKEERESVACLLKRIEMLDLFENQNIQMQKEVERFKEMVEESSRFQTQMQEKMKEAENDYEEKLLQVCDALDNTNID

LVAEREKVVSLTRQIESLGTVKEKNLVMEKETQEYKEMLEESEKCRVLLEEQISQLESDSNENIRELCSKVDIAYAKLAE

EVEKTASLVRKSESIDLNEEHRQRELDHYKEMLEESTKTQLLLQEKVVDVENDSKRKLADVSEALEIANSELSDKTSEVF

QIEFQLWVWKSIAKRLKAELEQNQNLRKRVEASLLEQVGVGEAIKQEKNELVHKLKVISHARSSDSEKKESLMRDKDEML

ESLQREVELLEQDSLRRELEDVVLAHMIGERELQNEREICALQQKDQDLCEVKHELEGSLKSVSLLLQQKQNEVNMLRKT

WEKLTARQILTAVETESKKMMIIELEGEISSLSQKLETSNESVSCFRQEATKSRAELETKQTELKEVTTQMQEKLRTSEA

EKTELVKEVASLSTEKRNLLSFISEMEDGMLKLYDGDTKLMKTLERVTQCCDGFGKENNNGETIGSPRLAMKHEEDVVTE

DREGASQNRTAPDLKPIETCRKLLKSREFLSVRYIIQVNQLLKSKRHDDDDQFEQVEAICDLLIAAENRLPNGLKDYYGM

IRANRFGPVLLEDFEKLTKLLDKKHNYFPFSQEAADKASLAWSLLSKPPIKAHYDLAISAFFGECSKVKRKIFIPKKQIE

VVVISDDDDEEEFSCLKISLNRYYYYDYLLFVFIYIYIHNKFNIPSNVLLSPSLCVTMVLLHHVSLSHYQNSSSLFSSSS

ESILCLFLVLCVMQLEQGMRPISRCYNPTAYSTTMGRSFFAGAATSSKLFSRGFSVTKPKSKTESKEVAAKKHSDAERRR

RLRINSQFATLRTILPNLVKQDKASVLGETVRYFNELKKMVQDIPTTPSLEDNLRLDHCNNNRDLARVVFSCSDREGLMS

EVAESMKAVKAKAVRAEIMTVGGRTKCALFVQGVNGNEGLVKLKKSLKLVVNGKSSSEAKNNNNGGSLLIQQQ

>At5G08130

MELPQPRPFKTQEFRTGRKPTHDFLSLCSHSTVHPDPKPTPPPSSQGSHLKTHDFLQPLECVGAKEDVSRINSTTTASEK

PPPPAPPPPLQHVLPGGIGTYTISPIPYFHHHHQRIPKPELSPPMMFNANERNVLDENSNSNCSSYAAASSGFTLWDESA

SGKKGQTRKENSVGERVNMRADVAATVGQWPVAERRSQSLTNNHMSGFSSLSSSQGSVLKSQSFMDMIRSAKGSSQEDDL

DDEEDFIMKKESSSTSQSHRVDLRVKADVRGSPNDQKLNTPRSKHSATEQRRRSKINDRFQMLRQLIPNSDQKRDKASFL

LEVIEYIQFLQEKADKYVTSYQGWNHEPAKLLNWSNNNQQLVPEGVAFAPKLEEEKNNIPVSVLATAQGVVIDHPTTATT

SPFPLSIQSNSFFSPVIAGNPVPQFHARVASSEAVEPSPSSRSQKEEEDEEVLEGNIRISSVYSQGLVKTLREALENSGV

DLTKASISVEIELAKQSSSSSFKDHEVREPVSRTRNDNVKQTRKPKRLKTGQ

>At5G10570

METELTQLRKQESNNLNGVNGGFMAIDQFVPNDWNFDYLCFNNLLQEDDNIDHPSSSSLMNLISQPPPLLHQPPQPSSPL

YDSPPLSSAFDYPFLEDIIHSSYSPPPLILPASQENTNNYSPLMEESKSFISIGETNKKRSNKKLEGQPSKNLMAERRRR

KRLNDRLSLLRSIVPKITKMDRTSILGDAIDYMKELLDKINKLQEDEQELGSNSHLSTLITNESMVRNSLKFEVDQREVN

THIDICCPTKPGLVVSTVSTLETLGLEIEQCVISCFSDFSLQASCFEVGEQRYMVTSEATKQALIRNAGYGGRCL

>At5G37800

MSLINEHCNERNYISTPNSSEDLSSPQNCGLDEGASASSSSTINSDHQNNQGFVFYPSGETIEDHNSLMDFNASSFFTFD

NHRSLISPVTNGGAFPVVDGNMSYSYDGWSHHQVDSISPRVIKTPNSFETTSSFGLTSNSMSKPATNHGNGDWLYSGSTI

VNIGSRHESTSPKLAGNKRPFTGENTQLSKKPSSGTNGKIKPKATTSPKDPQSLAAKNRRERISERLKVLQELVPNGTKV

DLVTMLEKAIGYVKFLQVQVKVLAADEFWPAQGGKAPDISQVKEAIDAILSSSQRDSNSTRETSIAE

>At5G38860

MNSHDIDDQLEADVYSNLPSRNDSSTGRRNRNSCRSKHSETEQRRRSKINERFQSLMDIIPQNQNDQKRDKASFLLEVIE

YIHFLQEKVHMYEDSHQMWYQSPTKLIPWRNSHGSVAEENDHPQIVKSFSSNDKVAASSGFLLDTYNSVNPDIDSAVSTK

IPEHSPVSAVSSYLRTEPSLQFVQHDFWQPKTSCGTINCFTNELLTSDEKTSASLSTVCSQRVLNTLTEALKSSGVNMSE

TMISVQLSLRKREDREYSVAAFASEDNGNSIADEEGDSPTETRSFCNDIDHSQKRIRR

>At5G41315

MATGQNRTTVPENLKKHLAVSVRNIQWSYGIFWSVSASQSGVLEWGDGYYNGDIKTRKTIQASEIKADQLGLRRSEQLSE

LYESLSVAESSSSGVAAGSQVTRRASAAALSPEDLADTEWYYLVCMSFVFNIGEGMPGRTFANGEPIWLCNAHTADSKVF

SRSLLAKSAAVKTVVCFPFLGGVVEIGTTEHITEDMNVIQCVKTSFLEAPDPYATILPARSDYHIDNVLDPQQILGDEIY

APMFSTEPFPTASPSRTTNGFDQEHEQVADDHDSFMTERITGGASQVQSWQLMDDELSNCVHQSLNSSDCVSQTFVEGAA

GRVAYGARKSRVQRLGQIQEQQRNVKTLSFDPRNDDVHYQSVISTIFKTNHQLILGPQFRNCDKQSSFTRWKKSSSSSSG

TATVTAPSQGMLKKIIFDVPRVHQKEKLMLDSPEARDETGNHAVLEKKRREKLNERFMTLRKIIPSINKIDKVSILDDTI

EYLQELERRVQELESCRESTDTETRGTMTMKRKKPCDAGERTSANCANNETGNGKKVSVNNVGEAEPADTGFTGLTDNLR

IGSFGNEVVIELRCAWREGVLLEIMDVISDLHLDSHSVQSSTGDGLLCLTVNCKHKGSKIATPGMIKEALQRVAWIC

>At5G43175

MENEAFVDGELESLLGMFNFDQCSSNESSFCNAPNETDVFSSDDFFPFGTILQSNYAAVLDGSNHQTNRNVDSRQDLLKP

RKKQKLSSESNLVTEPKTAWRDGQSLSSYNSSDDEKALGLVSNTSKSLKRKAKANRGIASDPQSLYARKRRERINDRLKT

LQSLVPNGTKVDISTMLEDAVHYVKFLQLQIKLLSSEDLWMYAPLAHNGLNMGLHHNLLSRLI

>At5G46690

MTLEALSSNGLLNFLLSETLSPTPFKSLVDLEPLPENDVIISKNTISEISNQEPPPQRQPPATNRGKKRRRRKPRVCKNE

EEAENQRMTHIAVERNRRRQMNQHLSVLRSLMPQPFAHKGDQASIVGGAIDFIKELEHKLLSLEAQKHHNAKLNQSVTSS

TSQDSNGEQENPHQPSSLSLSQFFLHSYDPSQENRNGSTSSVKTPMEDLEVTLIETHANIRILSRRRGFRWSTLATTKPP

QLSKLVASLQSLSLSILHLSVTTLDNYAIYSISAKVEESCQLSSVDDIAGAVHHMLSIIEEEPFCCSSMSELPFDFSLNH

SNVTHSL

>At5G46760

MNGTTSSINFLTSDDDASAAAMEAFIGTNHHSSLFPPPPQQPPQPQFNEDTLQQRLQALIESAGENWTYAIFWQISHDFD

SSTGDNTVILGWGDGYYKGEEDKEKKKNNTNTAEQEHRKRVIRELNSLISGGIGVSDESNDEEVTDTEWFFLVSMTQSFV

NGVGLPGESFLNSRVIWLSGSGALTGSGCERAGQGQIYGLKTMVCIATQNGVVELGSSEVISQSSDLMHKVNNLFNFNNG

GGNNGVEASSWGFNLNPDQGENDPALWISEPTNTGIESPARVNNGNNSNSNSKSDSHQISKLEKNDISSVENQNRQSSCL

VEKDLTFQGGLLKSNETLSFCGNESSKKRTSVSKGSNNDEGMLSFSTVVRSAANDSDHSDLEASVVKEAIVVEPPEKKPR

KRGRKPANGREEPLNHVEAERQRREKLNQRFYSLRAVVPNVSKMDKASLLGDAISYINELKSKLQQAESDKEEIQKKLDG

MSKEGNNGKGCGSRAKERKSSNQDSTASSIEMEIDVKIIGWDVMIRVQCGKKDHPGARFMEALKELDLEVNHASLSVVND

LMIQQATVKMGSQFFNHDQLKVALMTKVGENY

>At5G46830

MINTDDNLLMIEALLTSDPSPPLLPANLSLETTLPKRLHAVLNGTHEPWSYAIFWKPSYDDFSGEAVLKWGDGVYTGGNE

EKTRGRLRRKKTILSSPEEKERRSNVIRELNLMISGEAFPVVEDDVSDDDDVEVTDMEWFFLVSMTWSFGNGSGLAGKAF

ASYNPVLVTGSDLIYGSGCDRAKQGGDVGLQTILCIPSHNGVLELASTEEIRPNSDLFNRIRFLFGGSKYFSGAPNSNSE

LFPFQLESSCSSTVTGNPNPSPVYLQNRYNLNFSTSSSTLARAPCGDVLSFGENVKQSFENRNPNTYSDQIQNVVPHATV

MLEKKKGKKRGRKPAHGRDKPLNHVEAERMRREKLNHRFYALRAVVPNVSKMDKTSLLEDAVCYINELKSKAENVELEKH

AIEIQFNELKEIAGQRNAIPSVCKYEEKASEMMKIEVKIMESDDAMVRVESRKDHHPGARLMNALMDLELEVNHASISVM

NDLMIQQANVKMGLRIYKQEELRDLLMSKIS

>At5G48560

MDNELFMNTEFPPPPEMATHFEHQQSSSSAMMLNWALMDPNPHQDSSFLWEKSTEQQQQQSIFDSALSSLVSSPTPSNSN

FSGGGGDGFLIRELIGKLGNIGNNNNNSGEIYGTPMSRSASCYATPMSSPPPPTNSNSQMMMNRTTPLTEFSADPGFAER

AARFSCFGSRSFNGRTNTNLPINNGNNMVNNSGKLTRVSSTPALKALVSPEVTPGGEFSRKRKSVPKGKSKENPISTASP

SPSFSKTAEKNGGKGGSKSSEEKGGKRRREEEDDEEEEGEGEGNKSNNTKPPEPPKDYIHVRARRGQATDSHSLAERVRR

EKIGERMKLLQDLVPGCNKVTGKALMLDEIINYVQSLQRQVEFLSMKLSSVNDTRLDFNVDALVSKDVMIPSSNNRLHEE

GLQSKSSSHHHQQQLNIYNNNSQLLPNISSNNMMLQSPMNSLETSTLARSFTHLPTLTQFTDSISQYQMFSEEDLQSIVG

MGVAENPNNESQHMKIEL

>At5G50915

MATFSYFQNYPHSLLDPLLFPTPHSSINLTSFIDQNHLYPLPNISTVEDISFLEYNVDKTENSGSEKLANTTKTATTGSS

SCDQLSHGPSAITNTGKTRGRKARNSNNSKEGVEGRKSKKQKRGSKEEPPTDYIHVRARRGQATDSHSLAERVRREKISE

RMRTLQNLVPGCDKVTGKALMLDEIINYVQTLQTQVEFLSMKLTSISPVVYDFGSDLDGLILQSEMGSPEVGTSFTNAMP

TTTPIFPSLLDNSVVPTHAQVQEEGEERENFVDRSGFNNNNFCSFP

>At5G53210

MQEIIPDFLEECEFVDTSLAGDDLFAILESLEGAGEISPTAASTPKDGTTSSKELVKDQDYENSSPKRKKQRLETRKEED

EEEEDGDGEAEEDNKQDGQQKMSHVTVERNRRKQMNEHLTVLRSLMPCFYVKRGDQASIIGGVVEYISELQQVLQSLEAK

KQRKTYAEVLSPRVVPSPRPSPPVLSPRKPPLSPRINHHQIHHHLLLPPISPRTPQPTSPYRAIPPQLPLIPQPPLRSYS

SLASCSSLGDPPPYSPASSSSSPSVSSNHESSVINELVANSKSALADVEVKFSGANVLLKTVSHKIPGQVMKIIAALEDL

ALEILQVNINTVDETMLNSFTIKIGIECQLSAEELAQQIQQTFC

>At5G56960

MMHLILSCSYLISMDGYYNEASEEPSSSSSSGSLARSLFHEYRQSVIPLQNGHVPSMAFMNNLPYVEIRPQESQRLAFND

TQRLFYQMKIEASLREWFPEDFNRKSSPANSDYLRPPHYPSSSSSSLSPNNISEYSSLLFPLIPKPSTTTEAVNVPVLPP

LAPINMIHPQHQEPLFRNRQREEEAMTQAILAVLTGPSSPPSTSSSPQRKGRATAFKRYYSMISDRGRAPLPSVRKQSMM

TRAMSFYNRLNINQRERFTRENATTHGEGSGGSGGGGRYTSGPSATQLQHMISERKRREKLNESFQALRSLLPPGTKKDK

ASVLSIAREQLSSLQGEISKLLERNREVEAKLAGEREIENDLRPEERFNVRIRHIPESTSRERTLDLRVVLRGDIIRVDD

LMIRLLEFLKQINNVSLVSIEARTLARAEGDTSIVLVISLRLKIEGEWDESAFQEAVRRVVADLAH

>At5G57150

MEDIVDQELSNYWEPSSFLQNEDFEYDRSWPLEEAISGSYDSSSPDGAASSPASKNIVSERNRRQKLNQRLFALRSVVPN

ITKMDKASIIKDAISYIEGLQYEEKKLEAEIRELESTPKSSLSFSKDFDRDLLVPVTSKKMKQLDSGSSTSLIEVLELKV

TFMGERTMVVSVTCNKRTDTMVKLCEVFESLNLKILTSNLTSFSGMIFHTVFIELRPNIYWVVWFLVFMSIFGPTIIVIW

SIWFIKKKIILSLWRMKKNKRCCG

>At5G58010

MENGNGEGKGEFINQNNDFFLDSMSMLSSLPPCWDPSLPPPPPPPQSLFHALAVDAPFPDQFHHPQESGGPTMGSQEGLQ

PQGTVSTTSAPVVRQKPRVRARRGQATDPHSIAERLRRERIAERMKSLQELVPNTNKTDKASMLDEIIEYVRFLQLQVKV

LSMSRLGGAGSVGPRLNGLSAEAGGRLNALTAPCNGLNGNGNATGSSNESLRSTEQRVAKLMEEDMGSAMQYLQGKGLCL

MPISLATAISSSTTHSRGSLFNPISSAVAAEDSNVTATAVAAPEASSTMDDVSASKA

>At5G61270

MSNYGVKELTWENGQLTVHGLGDEVEPTTSNNPIWTQSLNGCETLESVVHQAALQQPSKFQLQSPNGPNHNYESKDGSCS

RKRGYPQEMDRWFAVQEESHRVGHSVTASASGTNMSWASFESGRSLKTARTGDRDYFRSGSETQDTEGDEQETRGEAGRS

NGRRGRAAAIHNESERRRRDRINQRMRTLQKLLPTASKADKVSILDDVIEHLKQLQAQVQFMSLRANLPQQMMIPQLPPP

QSVLSIQHQQQQQQQQQQQQQQQQQFQMSLLATMARMGMGGGGNGYGGLVPPPPPPPMMVPPMGNRDCTNGSSATLSDPY

SAFFAQTMNMDLYNKMAAAIYRQQSDQTTKVNIGMPSSSSNHEKRD

>At5G62610

MDPPLVNDSSFSAANPSSYTLSEIWPFPVNDAVRSGLRLAVNSGRVFTRSEHSGNKDVSAAEESTVTDLTAGWGSRKTRD

LNSEDDSSKMVSSSSSGNELKESGDKKRKLCGSESGNGDGSMRPEGETSSGGGGSKATEQKNKPEPPKDYIHVRARRGQA

TDRHSLAERARREKISEKMTALQDIIPGCNKIIGKALVLDEIINYIQSLQRQVEFLSMKLEVVNSGASTGPTIGVFPSGD

LGTLPIDVHRTIYEQQEANETRVSQPEWLHMQVDGNFNRTT

>At5G65640

MELSTQMNVFEELLVPTKQETTDNNINNLSFNGGFDHHHHQFFPNGYNIDYLCFNNEEEDENTLLYPSSFMDLISQPPPL

LLHQPPPLQPLSPPLSSSATAGATFDYPFLEALQEIIDSSSSSPPLILQNGQEENFNNPMSYPSPLMESDQSKSFSVGYC

GGETNKKKSKKLEGQPSKNLMAERRRRKRLNDRLSMLRSIVPKISKMDRTSILGDAIDYMKELLDKINKLQDEEQELGNS

NNSHHSKLFGDLKDLNANEPLVRNSPKFEIDRRDEDTRVDICCSPKPGLLLSTVNTLETLGLEIEQCVISCFSDFSLQAS

CSEGAEQRDFITSEDIKQALFRNAGYGGSCL

>At5G67110

MGDSDVGDRLPPPSSSDELSSFLRQILSRTPTAQPSSPPKSTNVSSAETFFPSVSGGAVSSVGYGVSETGQDKYAFEHKR

SGAKQRNSLKRNIDAQFHNLSEKKRRSKINEKMKALQKLIPNSNKTDKASMLDEAIEYLKQLQLQVQTLAVMNGLGLNPM

RLPQVPPPTHTRINETLEQDLNLETLLAAPHSLEPAKTSQGMCFSTATLL

>Atr002Ga00080

MIPFVPPGFTLRDGYCYMNNTTTTTTTTTQCWDQGRYLQKLACKSSSSPCSVLTLLNDDNNNNSEKVNICWREKEAKAAA

ASKSHSDAERRRRERINTHLNTLRNLLPNTTKTDKASLLAEVVRHVKELKQHAKGTEGHGPIPGEADELTVVHPEPSSSL

IMATLCCEDRPGLLSELTNAFKALRLRTVRAEITTLGGRVKNVFFIDREHSETSLSCLEDTLKAIMVKPGPPPIVRHESK

RPRINYHNNDFMV

>Atr002Gb02470

MAPPQPAGLWSDDNNAMLEAFMAAADLPIPSTPSTWFDPSPSNATTTHTTTSSITTQGANATSPSQDSLQHRLQSLIETA

HEPWTYAIFWQPSLPSQALAWADGYYKGEPLKLARPPAPPSAEQINRKLVLRQLHSLISQTSSATNDESVDEEVTDTEWF

YLVSMMQSFAGPTALPSHSFFSGAPVWLAGGGRLATCPCERARQAVAFGLQSLVCIPTSSGVVELGSTEAVPHNPELVRG

AQALFTWPDPHAQHDDPSLWLTDPAPEPPEPAKTTTQTPPVTKPVPPPPPPPPVVENTSNSTVTQTAFSLNFSDFGFEGL

ATNNQRGALNTYEDSHRDGKVGNLQPCKPESGEILSFGGGNGGGNSMSRGGSSSGLMGRVDEKRGGRLGPTMGPTRQNGG

VDEGILSFSSGVVLQSELKSGADSDHSDLEASVREVESSRIVDTAAERRPRKRGRRPANGREEPLNHVEAERQRREKLNQ

RFYALRAVVPNVSKMDKASLLGDAIAYINELRGKLQGLESENDELQTQVEALKKKESQLFNGSLKSQALSGDSDLKAHST

HSSANSGKYPGLEIEVKILGWEAMIRIQSNKQNHPAARFMVALKDLDLEVHYASVSVVKDLMIQQATVKMTSRIYTQDQL

SAALYSKVADIGSVR

>Atr007G01320

MHQQADGSHSPAFISPSLQRGPVPSPRALGTKTPSDDAQAVASQKSHSEAERRRRERINGHLSTLRQLLPCTSKMDKASL

LREVVDSLKELKRKASEISKDLSCPTDTDELRVELCNSGSPNSAFPLIKASFCCEDRPQLVADLIRALQSLRLRTVKANI

STLGGRVRNDFIFMARERVSVTTIQEALRAVLIRGSNGESTLDACNKRQKRILSFVHGLF

>Atr010G01040

MEGSEAHGSKAFSDFNGGGYIFPEMNQQRPMPPPYMTTELGSYNNNSGGGLAIPGNSVVAWPMPPPPPPPPPPPPVVPSF

NPTVDPAVHFAVQAREHERFLACMGGGGLTNRPSIYSGLGSSSSSSPSTFQGFGRRGIQFGYERENNVIYGGVSDGVGIG

IGIGGGGDAGKMSAQEIMDAKALAASKSHSEAERRRRERINTHLHTLRSLLPNTTKTDKASLLAEVIQHVKELKRQTAEI

AETMPIPTETDELTVDPDDSYGEGKLVIKASICCEDRSDLFPDIIKALKALRLRTLKAEITTLGGRIKNVLLVTNEEEGE

GREQDSNAPSINSIQDALKAVMERPLSEESSSTNIKRQRTNAAMLEHRSM

>Atr021Ga00510

MGSSFFGSDFLCFYGNGDDECCFDYGNGAVGFGGVIGEEWGWGRKKALVLDGSRGELVEEVPRAVVTEAKNAAALRSHSD

AERRRRERINAHLNTLRALISGPNKMDKASLLAQVINHLRDSKRKADEISKGSLIPTAFDEVRVENEADGPNKEGVFLRV

SLCCDDRPDLLIDLKRTLQNLRLKIISAEISTLGGRVKNVFVLTTCKENGDANKHQAVNDAAKNLVTCSDNSHAKCADEG

ADLLHEDNGGRSTSRAQWRLPRHLMRP

>Atr021Gb02190

MKAEMVSARVWSGEDKATIEAILGPKAVEFLTTTKFSFEGLVMATTADCNLQNKLCDLVEGHHFSKFSWNYAIFWQISSS

KSGELVLGWGDGYCKEPKEEEVVSENGLDLCNEKETQQKMRKRVLELLHLFFGGSDEENYELGLDRVSDTEMFYLVSMYF

SFPNGFGIPGRVFRTRKPLWLCDGPNSLADYCYRTFIARLTGIQTLVCIPVANGVLELGSIESIPQDQEALHKIVSVFAH

NDGQPNLRSNAWAFSSPTPLKSERPPSPLIEKNPKIFGQDLKLVQSQVERHYSVFNPSQENIGLCDNGERISSIGNAKAL

QAFNWNRIHGGNMGCTNGGAEKNRVNQIQQQKQHKQISFNGDHSDAEATCRESKLPQSSERKPRKRGRKPANGREEPLNH

VEAERQRREKLNQRFYALRAVVPNISKMDKASLLGDAIAYINELQNKLKDIETENEKQASSDLMEMALETEDQNKGVSQI

DIEVQSSSNEVLVKIECGLNEHPVSKIINLLREAQVTVEEPKISLSGDTVLHTFVIKSLGIERVTKDKLVSALSKLSNGL

>Atr025G00260

MVPTPSSSSKPNWSMATAETPALSSAAAQLQRLLQLAVQSVQWTYSVFWQICPQQGVLIWGDGYYNGAIKTRKTVQPMEV

NAEEVCLQRSQQLRELYDSLSAGETNQPSKRPCAALSPEDLTESEWFYLMCISFTFPLGIGIPGRAASRRHYIWLTGANE

EDSKVFTRAILAKVVPVQTIVCIPVMDGVLELGTTERVQEDSSMVQQLKALFMDDQGQLQPQKPVHSEHSTSKPATSTDS

FPYHRDHHRPHRQHHHQQHQHHQQPHDQQQEQPPLPWQAVEIEEEGDGESESESEREMQTKDIQVGLSPEGCSDPNPINM

TAHTSLTNPPCQEQHLPMPTNNLLLDDTNPWPLLHDDISIGLPSSGATMHRDDMVATQEDGHYSKTVAAVLQRNNTNNNA

APPFLVTYSCQFPKSEDTVFSKWKWNNQAATTTSQGGGGQQWLLKYILFSVPFLHSKYRDENSPKARGDGESGSRLRRGV

TTPQDELSANHVLAERRRREKLNERFIILRSLVPFVTKMDKASILGDTIEYVKQLRKRIQDLESRNRQMEINLKTRASVS

SETQKLSSTKDRTNSNTSAVLTTAQTLNDRSRTMALDKRKRHILEGARTKMAAGGCITDVQVSIIESDALLELQCPYRNR

LLLEIMQTLNELHFETQSVQSSSDNGVLIAEFRAKVKENPNGEKVTLVEAKQAIHHILENC

>Atr045G01440

MTKEKWDFAQPKMTNCSTDLAFVPDHEFVELLWENGQIVLQGQTSRNRKDASSSGYSSHPPKTHLNHNSGDAVLPVASKG

GRYGTLESVLHDFSSGQASGCPLVQEMKWPDHEFVELLWENGQIVLQGQTSRNRKDASSSGYSSHPPKTHLNHNSGDAVL

PVASKGGRYGTLESVLHDFSSGQASGCPLVQEDEMVPWLNYPLDDSLERDYCSEFFPELSGVNLTPLSKPTNVVVSEKAC

NLDTTSHKNLSMEHSSQAPSRLRALQPFNSQQGQPSNPILRQFPSNTHKSSCGNSSSPGANPAIGIVNSKTEQPNHGSTR

PPQPTHTNLMNFSHFSRPAALVKANLQSMGMAGRLKNNNDKASVTDSNPVESSIVDSAVVSKSGKTKLDQQCQASLHPTK

AKDISSSPAREKQSVEEDVMYVEDASTSQNRSPDRTLCPSSSFAASTALGISESVKAVEPVVASSSVCSGNSGGRVSKEP

RHGLKRKVGEQEESGYQSEDVEGESVGTRKQATGRSATTKRSRAAEVHNLSERRRRDRINEKMRALQELIPNCNKVDKAS

MLDEAIEYLKTLQLQVQIMSMASGMCMPPMMMTAGMQHMPPPHMTHFSPMGVGMGMGMGIGMGMGMGMLDIGGSPGCPLI

PLPSIRGPQIPCSSISGPMGLPGMPGSNLQMYGMPMARAPFLPFSGFPQAKAVAGPDISAPTSSGAFSELNPPSSSKDQM

QNLISPNVQQKVTDSQQLEASNQVISEHSAQPNLVPGHNQSSQVIGSNRNGVAPNGTTSYD

>Atr046G00260

MPFSPSSSSAPSSFSSSFPSQNSDFALAMAAGRAAGFIPQANWEAFHHLKTFQPHPSPFLASSEFSGDNKTLHEQPLRPE

FPVLGDFARRNLPLVSSSVGDETAEGTERGVCGTSGLSEAGVSGRQSSIGDQSSPRRDSEPCKKKKAHNDNDLDDLDCES

EEGQEPSEEMSKPAPSRSSTKRSRAAEVHNLSEKRRRSRINEKMKALQNLIPNSNKTDKASMLDEAIEYLKQLQLQVQML

SMKSGINLAPMCMPGQLQSMQLPQICMGFTTENGTLPITMGMGLLPVNQQDSSSQPSFDLPNISPPSSSLQSLVMPNIPN

VIPNSGGCAETSSFGLDSSQGLVRPYQLSSPAEDIYRADILPQQQVDMIQSVRSTAENQTKTVASSLSMDGQVTVLERAN

YLEAFMCVGDKVRPGLPKDVNSQSMVQRLHCSKIERGISNTEPSK

>Atr064G01050

MDNVFFMDVDTRTLFIRALVRKLGWTYICLWACQLYPSKCLVFMDGWFNDEGGQPSSSSGSFPRTLFESYCQSLLSLDNG

VPGLAHSHGMEDGLWLGGEGLINLASTDNQRRFYQGAGIENAFFMSCQSGVIELGTSNPNVDLNMNPKNLITDDVIRQFQ

QREPPPWADPSRASSSSSSLPSLSVGSPEYSNPAAITPPHLTIPAPAAHSITLTRMLPGGIPTSYEAIITRAILDVVSSS

PSSLSRVGASRPSQRPGAFQRYNSSGGVKAERRVGVSQRLFKRSIAVLRRIREVRQRVEEGAAAGTRPASSTQLHHVISE

RRRREKLNELFQALRAMLPLDSKKDKASILSRTREYLGELKVQVKELNERNISLETRLSSRGVIIAETALVASQEIETVE

ITETADDREIELRVRVVLEHHGLEDLVVRILQRLKRMEGFRVLSVSADSQVQPQFDTAIFRFNVQGGRWDKSWFEEEIRK

ELIELQHT

>Atr065G01560

MQGVNSMTTCLTSLQDLQQQQHTTLHQNQSQNQNQQITQQIQPHFDQHHDDLFDQMLSTLPSSWAELSKSPWPELNPQTL

QNSKLFSISLNSNSGTKTENSSENISPYYDESSLLASRLRQHQISSQSSPGLKLNNAGIGAAQLISPRGVARSPGSSSCL

SDGGMMPPLPLSLGQGGQDSNEETQIMVDRARDEVEASFKAGNGVGDGLVPGLYNGFSAASQPRASQPNNQHFYSQGQLP

SQTFGGGNPAMNHAPPGSGPTGNAAPARPRVRARRGQATDPHSIAERLRRERIAERMKALQELVPNANKTDKASMLDEII

EYVKFLQLQVKVLSMSRLGGAAAVAPLVADISSEGGGDCIQANGIGRNASGGQTPTQDSLTVTEHQVAKLMEEDMGSAMQ

YLQGKGLCLMPISLASAISSATCHTRNPSSVLMGPTNADVPSSPSMSVLTVQSAMANGTDVGAVKDAASVSKP

>Atr101G01220

MWTKREVNQESESYGYYWGAHEELEPGPLMADPSSSLGVSANLDAFSSYYCYGGNGFHGVEVGGMREKGAVDAKAMAALR

SHSEAEKRRRERINGHFDKLRHLLPSNVKRDKASLLAEAIRRLKELKLQTSEISAMGPIPDESDEVTVECSGEFPSDSGK

SSGQPVVMASFSCDDRSDLLPELITTLKSLRLKTIRAEIQTLEGRMRHVLHVTCEDSGQNSLDCGDLTAGMLQEALKQVV

HRSPEDRKMFSGGLKRQRTNSGF

>Atr106G00960

MSQCVPSWDLKSKLGLHVDEDDFQNLYDPLLESDYQVAELTWENGHLTMHGLGPPKPPIKPVQRCTWEKPQSGGGGGGGS

GGSGGGGGGGSGGGGGGGGGGGGTLECVVNQAKRASHGTLVTVEEPTTLAPWSVDALVPCSDPKGPARACAIDHASATCA

NVPAVPGLTPSQRGPGTCVASCSYPNIVPSHFQTLLKADEGTSKRKRARSMHVCEPMYTRVSGQGIDDSKNRSMRMGTCE

RSFDLGLASTSMGSQDKTSSARRLATEDHDSASHSRPQREDDGEHDEKRKCQGERSSVSTKRSRAAAVHSQSERKRRDRI

NQKMKALQKLVPNSSKTDKASMLDEVIEYLKQLQAQVHMISRMNMSQMMMPITMQQLQMAMMAQMGMGMSMGMGMGMGMD

INSAMRPAAIPNGMPPIIHPAFLPFPMSWDNSGDRLPASGNVMEDPFSAFLACHPQSMTMDAYGKMAALYQHLHQQQTPP

NTSVSRD

>Atr121G00050

MEPPTMAASSSLQQPWLTQQQVPLQQSLHHLLNTRTEWWEYAIFWEASHDSLPQAPVLVWGEGFFRGPTDGCNPSQMTKS

EAQQMERKKVLRDLQAMMEVDPDGDGFSLDSDVSDLEWFYMVSLTKSFMGAEGLPAHVFVAGRPIWLTGSHSIQSYNCER

TKEAHLHGIQSMACIPIANGVLELGSMDLIPEDWSLLETFTSIFGSGQLQRWGSDNPPPPSVPATEEGAVAVSSGPGWSG

VDSEHSDAEDIAAAAAAAEAIAADRRPKKRGRKPGNGRVVPLNHVEAERQRREKLNRRFYALRAVVPHVSKMDKASLLAD

AVSYINQLKAKVQHLESALQKLGSDEKTTQKDSDGKLMPDTSGSEGAEWNGIAELEVKIMGQDAIIRVQSESTGHPAAKL

MRALEELGLPVHHASVSNVKGLMVQDVVVRVPEGSPCSDDELKAALLRTGQLSRGPNSS

>Atr129G00950

MMDDFLDQFLSSSSWSDLSSTTRSSWNGAAGQTNGLLADPVGLYVDGPKNPTTTVVNPSLIIENLGNQRHAHEGTTENPN

YGLNKGLFPVQNQPTREAQDLGSAQSEHNKEASDRTFPLQQALESNATSGSLGHQLNASGSVTPTCSSAQYLPGVGGSNG

VVGGLIEHGSGGTKGNETVGFQQSIGDSLPINSAGALLRPSYGGVPLAPIMGQGKLEGFGLRNDYMEGDANLLGKGYLGV

DKLPLLENIPATSEQDEPCNSHLPSFATGPQIISPRTGGLPTHQQHGGNQSISQLHPVAATGGGCNGGVRPRVRARRGQA

TDPHSIAERLRREKIAERMKNLQELVPNSNKTDKASMLDEIIDYVKFLQLQVKVLSMSRLGAAGAVVPLITETEGSSNLL

LSTSAGQAPDLSESQDNIAFEQEVMKLMECNMTSAMQYLQNKGLCLMPIALASAISNSGKQSPENKYSMAGIQPLSPGSD

NLIVENILERERKELPNGAVIKQQEKQKPLQRI

>Cm12G02390

MLNFTTSAVLVTRTWRRRCTGCQQAPSVAYRKSTCFGVAGVRLCAESSPPTPAGAASLERSAGAGTETADGGVSGSGQGF

TPSITADNESQYYATRLVVTCRDRKGLLSDLTDALKSIGLQIRRAVARTKDGIASDEFFVTRDGSQLSDTDLDAVEQALQ

PVMGTSGPTCPVPQNTERRLPAPQSPVRFVDHNRGVHVYVDNHASQHYTTITVNAPDRPNLLNEIIDVLHELELNITFAC

LSTYADENKYRHDIFHVTTMSGEQVDAVLRDEIMNTLYFLLTSRESDEQSF

>Cm17G04120

MFTSIGLGWGVGSLISRAKLSRLKAPGTGEVCSGRTRHPFWERSKRACVTLRAEEAVSAPHVNPVTVTRPHNVPTADEDS

GPDLSRSRNFGSSVKPVASSSTALASDDFGATAAVLSAHPEATATVERPRKPGRARRQQALAQQAREDAEAVIRDVRREI

RRRNWPTAERLLRALVDREPTNGRAWLLLAQLYAYRLRDLEKARATFAAALQVNCTNTRLAHAFAMFEARCMQNTDAARS

LLDKALKIDPSDGVIWQAYALLEERTGHSERAQELFEAGLARDPRNVFLLQAFGMFHLRNGASVEACAYFERAVESNPSH

VPSWQAYGIALSKQGDWEHAAAKFEQALRLDPVSVPTLQAYGIAEARQGHYERARNLFQRAAELWPSHVPVYHAWATMED

RLGNHEEARKVFERGILAAGGRVDDVRRVRSELEGAKRTDPMLKAWADMEQRLGHISPAPEWNVYRSNRSDPETRERRRE

TIGERLLMLRKLIDRRSEEDLRTVLTFIAERTRMDRQARAAITERSRDDLDRVRRWAERRNEEDVRAFQKWFEKRYEEDR

LVGQYVFGWNLGPPLRSTVVSSPSTENTGGGDSASAVPPEWLRAAPMPRPLSEYEKELYSGERELQLVPAMAWISSFAEN

LSSRAAISTLLLGLMMFLSVGFAHIYELGRYEIGNVESTLLQILSTRVPEGVDAALIDLNPSALDDIAAEHW

>Cm2007185

KEQLCNVLRATDDRRGVKTDFSMGLTHPERRLHQMMFADRDYEGSSLSSQAKHEHGAHIT

IENCNEKGYSVVSVRCRDRPKLLFDTVCTLTDMQYVVSHASIGSDGPYAVQEYYIRHMDG

CILDTQGEQQRVVKCLEAAIERRASQGIRLELCTSDRVGLLSDVTRIFRENGLSVTRADV

TTQGNKAVNIFYVTDASGGPVDMKTIETIRNEIGQQILHVKDVPKFGRAYATDGPGKPGF

SFGSLIKSQWERFSSNLGLVKSCS*

>Cm2146157

QELIPNANKPDKASLLDAVIEYVKLLQSQLQVMSMRTGIGIPSGTISNMSMQHLQMPALP

QVRLGMDMGMGVGLGMGMRMGMGLMDVNTPAVGTGCKIMPMPQFSGQPSIGHSPHAAGLQ

PHNHLQNPGIMNPYNLYLAPSQVQPVSMNPNMNQNVAMYNAYMQWQQQQLQVQLQQSHQL

QGNNGRKSSQ*

>Cma2001047

SNSIHGGGGRLGLHSVNKAMQDMSGSSAPSKTVNSQDHLMAERKRREKLSQRFIALSAIV

PGLKKMDKASVLGDAIKYVKNLQERLKALEDQVPKTVSVAIQKSGKGSADPSIVSEARKD

NPEGLQQPDIEVRVIEKNALIRVHCDKSKSVLVKVLAELEKLQLSVVNANIL

>Cma2001048

SNSIHGGGGRLGLHSVNKAMQDMSGSSAPSKTVNSQDHLMAERKRREKLSQRFIALSAIV

PGLKKMDKASVLGDAIKYVKNLEERVKTLEDQVPKTVSVAAQKSGKGSADPSIVSEARKD

NPEGLQQPDIEVRVIEKNALIRVHCDKSKSVLVKVLAELEKLQLSVVNANIL

>Cma2001431

MQSGHGISMSPYDSHTGLAQSQGAVGVVTGAPKPRVRARRGQATDPHSIAE

RLRRERIAERMAALQELVPNSNKTDKVSMLDEIIEYLKFLQLQVKVLSMSRLGGAAAVAP

LVADLPAEGKNSLAAAALGQNAGLPQDAMTAAEHEVASLMEEDMX

>Cma2001432

XTGAPKPRVRARRGQATDPH

SIAERLRRERIAERMAALQELVPNSNKTDKVSMLDEIIEYLKFLQLQVKVLSMSRLGGAA

AIAPLVADLSAEGKSSLAAAALHQNAGLPQDAMMAAEHDVASLMEEDMGTAMQFLQSKGL

CLMPISLATAX

>Cma2001433

RERIAERMKALQELVPNSNKTDKASMLDEIIEYVKFLQLQVKVLSMSRLGGAAAVAPLVA

DLPAEGKNSLAAAALGQNAGLPQDAMTAAEHEVASLMEEDMX

>Cma2006004

RSTSRRNSGRIVEGNVDVMQRNEMARQHMIAERKRRRRQNDSFKALKTVVPSITKKDKIT

ILEHTLNHLKQLQERVDELEQEKNLLIIQATVHHANSSSVT

>Cma2006434

XHGIPINKHWQQQQQHGLEVLDGNKRPAR

VPSEDDDDEDDESLDHGYNPKKEGRYQKCDTTQQLEGKSMDSKGITPRSKHSATEQRRRS

KINDRFQMLRQLLPNADQKRDKASFLLEVIEYIQMLHEKVKKYEAAEHGGQQQERTKTIS

WDRNSGRRDPAANALYVPVVSTVTYDTKEAPDSNGYMRSRSSPLRVTCENGITGSELMSS

MAQVSALQNRLLQGAQACLSNYESKSLVSKPPPLTYTQPPAISTLERASTSMFHTVPQSM

PEGRTMFGHLLHQEELAAIKAKCSLYSKVGGTALPQLQQVSNLLSPQTSNCVLDQDHEKF

RSPGRNSAISNHRRPDEELEERTRSSQVIDMYQTDVGSTSKESGKDDGRPAHALDGKDDC

RPVHGVIDGPKGECKGEVPANEQEPPAILGGVINISSVYSQGLLDTLTQALQSSGIDLAQ

ATX

>Cma2006435

MAQLESGIPSSQQQGPPPLPQQSFDLEVRDLQLPAASEGKEATIHDFLSLCGSRSTS

STVPVTKNGSQSSITTQDLLQPLQKSIKPGSSNLLPHGFVLSATRPNTSSGDYCAGQQTT

YALVNGTLELPSNLQSKYRSSDISYVTKAHVPHGIPINKHWQQQQQHGLEVLDGNKRPAR

VPSEDDDDEDDESLDHGYNPKKEGRYQKCDTTQQLEGKSMDSKGITPRSKHSATEQRRRS

KINDRFQMLRQLLPNADQKRDKASFLLEVIEYIQMLHEKVKKYEAAEHGGQQQERTKTIS

WDRNSGRRDPAANALYVPVVSTVTYDTKEAPDSNGYMRSRSSPLRVTCENGITGSELMSS

MAQVSALQSRLLQGAPACLSNYESKSLVTKPTPLTFTQPLAISTLERASTPMFHSVPQSM

PEERTMFGHLLHQEEQAAVKAKCSLYNKVEAIALPQLQQVSNLLAPQTSNYVLDQDHEKF

RSPGRNSAISNHRRPDEELEERTRSSQVIDMYQTDVGSTSKESGKDDGRPAHALDGKDDC

RPVHGVIDGPKGECKGEVPANEQEPPAILGGVINISSVYSQGLLDTLTQALQSSGIDLAQ

ATX

>Cma2006436

MDSKGITPRSKHSATEQRRRS

KINDRFQMLRQLLPNADQKRDKASFLLEVIEYIQMLHEKVKKYEAAEHGGQQQERTKTIS

WDRNSGRRDAAANASYLPLASTVTYDTKEAPDSNGYMRSRSNPLRVTCENGITGSELMSS

MAQVSALQSRLLQGAPACLSNYESKSLVTKPTPLTFTQPLAISTLERASTPMFHSVPQSM

PEERTMFGHLLHQEEQAAVKAKCSLYNKVEAIALPQLQQVSNLLAPQTSNYVLDQDHEKF

RSPGRNSAISNHRRPDEELEERTRSSQVIDMYQTDVGSTSKESGKDDGRPAHALDGKDDC

RPVHGVIDGPKGECKGEVPANEQEPPAILGGVINISSVYSQGLLDTLTQALQSSGIDLAQ

ATX

>Cma2006673

GSSAPPKPGNSQDHIMAERKRREKLSQRFIALSAIVPGLKKMDKASVLGDALKYVKHLQE

RLKTLEEQAPKTVSVAVQKSGRDANGLNAVPDVKKGNSEGFQQPDIEVRX

>Cma2006674

GSSAPPKPGNSQDHIMAERKRREKLSQRFIALSAIVPGLKKMDKASVLGDAIKYVKQLQE

RLKTMEEQAPKTVSVAVQKSGRDAAGLNAVSDVQKDNSESSQ

>Cma2010596

MQPSAQMFKA

TATFPLGTSPTFCGRXPQMYGNTAAKAAAGYGGAAVKAPEAQNSNSFVRAPALQSYGLSN

KLPEAHNLTRAFKVAEVKDHKYVPNALGMQGQALDYNYVSKGQAEMQSYGQGMKPSENSS

FSQPAKSFGDARMQGISDNQSREEKFKGSLQQEDILALPMNGAIRSSVESEHSDAEASFK

DAECSEAVQERKPRKRGRKPANGREEPLNHVEAERQRREKLNQRFYALRAVVPNVSKMDK

ASLLNDAALYIQELKCKLQDLEIENKSLIAQLKTSKKEAFSHADHGDPSTWTPDIIKSKG

PVPFLPAGGRVTVKVHFLGGREAMIKVDSPKESHPVARVMVVLQDLQLQVHHASVSTVQD

IIRQSILVKMRGQCFFTEDQLTAAIFGRGD*

>Cma2011585

MSFCVPDWDGNGFLGDFSSNTRGKWEAARPISMESPFVP

DDEFEELCWEDGQLMLVMPSQNNRPPHKQLSSWAASVSPLNKPVSRDQVDASLYMGDIAT

NHDETLDAVVDDGDGSRVAGFPPAISDDEMIAWLQYPLDDELRVDDSDPVLHSGHHAVSY

LPSLPFQKSDSVHRNIGAMQRERPSIGITKGKQAATTTNTDLAQAIARRGSQTESFSASF

GNSTSALTTDGATSDPKPGTQYLEQTIFSSKAASLLPPLVGSPYCSQQSDPQKILQSISR

PVTDMKPATGVPNWPASLKHVKHIDRGNVEVCMSTDSSIESTITGRSLQEPLGDRSGGLT

GPNVSHCMTATTSTLHQLLSVKGPDAAACIDDFAVCSETVVADIPSGSKLFCSADANQTV

EKGSGAFESTDTSCAAGSGNTTATNGGKVDSNGSYGKRKVKDLDDSESQSEDLGKQPPLS

QATSTKRGRAAEVHNNSERKRRERINEKLKALQELIPNANKTDKASMLGEAIEYLKALQS

HILVMSYRTGITIPPMLISHGLPHPHVPPHAGIGFGAGMGVKTGVMDMSAATSTCSMPKA

LALGCLIPSVTIPALSTPATNVATLPQNGLSRPVVGYSTNYFPFSTFGAQPFLVHTNQLQ

NPPATTSHKTPSLHRHTHQPSQLLSTDMFNPSMQEWQQQQQHMHKQ*

>Cma2011784

HGHGLDHSTYSTPTTTISAEDLDLLDSPTMLNPLDQDMFLLDTSSPGMEIDWNLPSVEIF

PVWKPHEFPSLTQNVNDDHPQHHHQQETHDHHERPHQQETHETISDKIQVGNQVRLRWYG

ERSKEDHTRVRPMASSSSPNETKGSSTISTGSPSLPKSCGLKRPKSAIDLMKQLEHSDAM

EDPNGQAPMSKNLVSERRRRKKLNERLYSLRAIVPKISKMDKSSIVGDAISYVQDLQKQV

EDVQNDILTLQTGKDAVHEGSSESRSTNEGGGGMDAKFPYQNLGEHKILELDVSSMEENT

YQLRIYCKKGPGVLVQLTRALEALNFEIVNANLTLVTDHILNTIVVKVPVWWTGATTCVW

MCMYMDAGTNGFMFMPHMCGCASCLWWMCIMFMLHAMFIVHVCGCWHKWSYVYATSVWAG

CA*

>Cma2011785

HGHGLDHSTYSTPTTTISAEDLDLLDSPTMLNPLDQDMFLLDTSSPGMEIDWNLPSXXXP

HEFPSLTQNVNDDHPQHHHQQETHDHHERPHQQETHETISDKIQVGNQVRLRWYGERSKE

DHTRVRPMASSSSPNETKGSSTISTGSPSLPKSCGLKRPKSAIDLMKQLEHSDAMEDPNG

QAPMSKNLVSERRRRKKLNERLYSLRAIVPKISKMDKSSIVGDAISYVQDLQKQVEDVQN

DILTLQTGKDAVHEGSSESRSTNEGGGGMDAKFPYQNLGEHKILELDVSSMEENTYQLRI

YCKKGPGVLVQLTRALEALNFEIVNANLTLVTDHILNTIVVKVPVWWTGATCVWTYGCWH

KWVYVYATYVWMCIMFMVDVHYVYATCYVYSACVWMLAQMELCLCHICVGWMCINGAVCL

DTCR*

>Cma2012864

MSSQGAEIREWSPEDLAMIKQLLG

PSAIAHLSWNLQGLTPAGGRRLNEAALQQRLQSLVEGSAVNWTYAIFWQLSCTPKGEEVL

GWGDGYFKGPKENISEPSQGMEKDEDQQLKRRVLRTLQAQFCATDDDLMDTADEDLVSDT

ELFYLISMFYSFQRGMGIPGASFESQNHVWMTGTNRDTSNVCTRGPLARMADIKTIVCVP

TLRGVAELGSTDLIYENRKLIHEIKVSFTDDVWEQQIENRSSPLPMAPAKHIPFPACASD

PLAAPMPALRDMMSCRTQETGRHRSSLVSEPFHGGASQDFRNGRMAPGLHPEHLQSLPAP

FQLNPFLAAQRAAAVKVFQAHNWHPKHNVEAAKGLNPCLKQTGLEMHHRNQIIKGSSLSM

DLGTQVSSVLGGEEKFKQGLGHSFSAMVRSCVESEHSDVEASCKEVDFKPAFEERRPRKR

GRKPANGREEPLNHVEAERQRREKLNQRFYALRSVVPNISKMDKASLLGDAIAYIQELQA

KLKEAQNEKDELQARSMANGADASALFETKEGLHSPSGMQNVELQVLNGEATVKASCPKE

NYPLTKVMEVLQELKLEMPSATVALLNESIVHTIKLNLRGSDMVTEEQLHAAITKSS*

>Cma2014229

KGGFRRMSSSSDLLKQVDSSDNSDGMEGMDEGGKNRGNGRSKGPMSKNLVSERRRRKKLN

ERLYSLRALVPKISKMDKASIVGDAINYVRDLQKQVEEMKVDVESLEASKEAVANGLIDP

KKDLKRVMRCHGKKVVLQEHHIMELDVTQMEEQTYHLRIHCKKSPGVLVQLTRALEALEF

EILNANLTSVNDHILNTLVIEVKNGHLMKSEEVRQMTLEVIPRFGLFL*

>Cma2014775

MESLEDTPHSDKLLESMGRLRPLSTTDLFQTMTGNNSAYSAPMESPPITISTEDLEI

LNSPTIMNSFDHEMLLSGLSTSSMMGINGDEFTPLLEHSEKWNLPSEPIFSTWNPSSEHA

RFHYRPDVVDGASSKSRSGSLEGGKGQRSWCQKTNKQEETRVNAGGRFTVISSSPEEWRG

SSVTSVSTGAQGFRRWDSANDLSRQFDSSDNSDAMEDEEGDKQSPNGKAPMSRNLASERR

RRKKLNERLYSLRALVPKISKMDKX

>Cma2015959

TDTEWFFLVSMMCSFPIGTGTPGQAFASSQYAWLNGADQLRGRDCMRAELAQRFGIRTIL

CVPTQNGVVELGSTEVIGEDLAFLNIIRHSFDPLLSNMDMPYFSSPMRSFTDVNPALVAE

FGYLNNSGNEVSAGDSGQTKMSSSGMMMQDLQLGATVEKAGQFLGNGTHGDPLHSLLPQD

SPYSDMMFPGNESERLLLPMIWQQSSFSQPAVAASEDGRALTTANQSHDPQLLQKAEFED

VTKRRGMSPDFSHYDHYVKQLKDDSQASGFKARECHSYKETTKVESSSEFMKPVAVQNYS

HSMVGEMQSQSQTLPQNYTPFVKTAHIDNFNQFGMSTVPQTYPHHSVKVTEMQSHSQMST

MAAGPQIYPNGIKTAEMQSHSQVGMTIVPQSFTKTSEVQSYSQPVRAPETESHFDGGKAP

DLFNHDHFEKETLGTLSHSQAAPASAIKTSDSARKSPDHIVPMRSHVQSYNQTSKLWGLD

VQTYSLKPADSTSSYQQDVKAVELLTFDKRSTELNKQSSDEKEGGLLQLDESVAAQVTGA

VRSSVESEHSDVEASFKEADCSQSVVEKKPRKRGRKPANGREEPLNHVEAERQRREKMNQ

RFYALRAVVPNVSRMDKASLLGDATSYIEELRAKVHGLEIEKKKLLLRLDGGKADRMMSH

GGSFKNSGQGDPPARTSPTDRKSMNSLACPHGKVAINVHFLQGREALIQVESSRENFPVA

KLMLALQELQLEVHHATAAVVQGMLFQKIIVIMKTPSYITEQQLTAALSTRAVDCNCC

>Cma2015962

TDTEWFFLVSMMCSFPIGTGTPGQAFASSQYAWLNGADQLRGRDCMRAELAQRFGIRTIL

CVPTQNGVVELGSTEVIGEDLAFLNIIRHSFDPLLSNMDMPYFSSPMRSFTDVNPALVAE

FGYLNNSGNEVSAGDSGQTKMSSSGMMMQDLQLGATVEKAGQFLGNGTHGDPLHSLLPQD

SPYSDMMFPGNESERLLLPMIWQQSSFSQPAVAASEDGRALTTANQSHDPQLLQKAEFED

VTKRRGMSPDFSHYDHYVKQLKDDSQASGFKARECHSYKETTKVESSSEFMKPVAVQNYS

HSMVGEMQSQSQTLPQNYTPFVKTAHIDNFNQFGMSTVPQTYPHHSVKVTEMQSHSQMST

MAAGPQIYPNGIKTAEMQSHSQVGMTIVPQSFTKTSEVQSYSQPVRAPETESHFDGGKAP

DLFNHDHFEKETLGTLSHSQAAPASAIKTSDSARKSPDHIVPMRSHVQSYNQTSKLWGLD

VQTYSLKPADSTSSYQQDVKAVELLTFDKRSTELNKQSSDEKEGGLLQLDESVAAQVTGA

VRSSVESEHSDVEASFKEADCSQSVVEKKPRKRGRKPANGREEPLNHVEAERQRREKMNQ

RFYALRAVVPNVSRMDKASLLGDATSYIEELRAKVHGLEIEKKKLLLRLDGGKADRMMSH

GGSFKNSGQGDPPARTSPTDRKSMNSLACPHGKVAINVHFLQGREALIQVESSRENFPVA

KLMLALQELQLEVHHATAAVVQGMLFQKIIVIMKTPSYITEQQLTAALSTRAVDCNCC*

>Cma2017379

EEDIIDYVSSSMVKSGAGDVHGFLGKGSAGLPSHHQVFGLNAPLMAGGVSLHRKLASQLA

TSELSAVTDTEENQRTSLPPVPKDLFSDELSDVPEKERGSFLKAEDSQLMEEAKSDQLCK

ANIETLSHNSEFEIDKEKEDNFDDMGDGNAMIHDGDDGLGFDEPIQVQASTVSAEAGKGK

KGLPAKNLHAERRRRKKLNERLYLLRSVVPKISKMDRASILGDAIDYLKDLLQQINDLQM

ELKSPPSADSTLLPCIPSIAMGTSPSTGACVNEEFSTLMEGPEGPPPKVEVDTREGRALN

IHMVCSKQPGLLLATVKKLDELGLDIQQAVVSCFNGFALDVFRAEQPSEKDVQPEDIKTA

LLQTVGCQQYML*

>Cma2018129

MHARNSAGMQDLQFGTSKQHTTLDL

EMPQLSLASVQEGASQSTTHDFLSLYKSAAPGGKTGPAALQGFVSSTATQDFLQPLEQCA

KSVSECTIPCGLIASTGFKNRVRHDGEAEASYGPYNETMEFSSIVSVCTNGGCKDEGGSE

CKENTSRSKHSATEQRRRSKINDRFQMLRQLVPYSDQKRDKVSFLLEVSEYIQVLQAKVQ

KYEATKLAWQHERSQKMSWDISQRNSGMQHYAGNSPVVFDPL

>Cma2018130

MHARNSAGMQDLQFGTSKQHTTLDL

EMPQLSLASVQEGASQSTTHDFLSLYKSAAPGGKTGPAALQGFVSSTATQDFLQPLEQCA

KSVSECTIPCGLIASTGFKNRVRHDGEAEASYGPYNETMEFSSIVSVCTNGGCKDEGGSE

CKENTSRSKHSATEQRRRSKINDRFQMLRQLVPCSDQKRDKASFLLEVIEYIQALQERVQ

KYEATELAWEHERSQKIXACNFC*

>Cma2021101

EKVSSSAEGRILIRVSLCCDDRPDLMADLRDALDTLRLRTLKAEISTLGGRIKNVFIVTL

KEDTTEPDQDISVKCVQEALKLVMDGPGGGELSPGSSNKRQRVSPFESPGCT*

>Cma2022893

MSAANNSSKSGSKSCSSPDDFYEDELVTWLQYPLEDSFEKSYCSDLF

GQWPSAAAGAALAKTAKTPPDMGASKPPAGTSMAMGRSVCTESGSLSRSDSARALVDTGN

CGAKEKAGGGPLLRSTSAEGALALGAERAAGLASQSGRDAFSKVRTSMQPPAGSQSSPLS

SMKATGKTGSNSMANPGSFASKLPLQPTQGGLHSFPNSPRPMAGLRSQVSNSLAVGVKGA

ISKSMSFTSDRSRKGSETHLQFSSAQQSGCSRITEEATIRRESGYTSSASSRGHLCTTQM

NGGMLARLTPSIVEGHGQKRTGVGGASEMALASCSGGSQTSADLSMREATCSGKRKFEDS

ECPSEDAEGESMEIKQPRKVSSSKRNRAAEVHNLSERKRRNRINEKLKALQDLIPNSSKT

DKASVLDEAIEYMKTLQMQLQMVSMRSGMNVSPMMIPTGMPHLQMAPLHPVGLGVPIGHA

GMGMGMGMGLGLSMGMGMMDLPQGVMARPFQSIPAAMTHFGMPFQGGSVVSSQDCSQGST

KMELTPSELFPVPHNSSQPSDQMHPRSHSMSDAHHLHQQHEQGQHLAHHSLDHETREHFN

EQTGKED*

>Cma2025364

RLTEFPASIPGVEMLDWLDYPLVEPVDGEATIVTSCVSPSNMPRALASSHSFQPFGRMSP

VDFSQTSARAEHVGTTCGPSALVTSKDFGKVDVEVGMPVVSSLHSTMTRKLIPEDKDALL

SCSAPHCMTSSPITQSLQNTLFTQRTGAAVFFDGSPAARNAKTSTASNCVNPYASVEKSP

ASDIESVFLEASEPTHISFSCGSGHDPDKGDRFGHACGKRKTEAMNDFDSESEDSYGDVV

VARKPRQERSASTKRTRAAETHNQSERKRRGRINEKLNALQGLIPNANKTDKASLLSEAI

DYLKKLRLQLQIMSYRSGMSISPLLISQGVQHLQMPHVGMGVGADATSTSFRSLPTAPII

GSTAPFVSAQICPGSTGPPIAQASLQDPFVSTSSHFPPLPTAVPRPSGSQDLSRSVSFHH

PFQV*

>Cma2025672

HVKEGLSVSTAQSNATAMTGMGSSRLHPYRVETARKPFIAHRKEGLSASVTKSTTTGVTS

MGSCKLQAGASGPTNVERARQPSMAHTKGGLSSCAAESTTIVGPGTGSKLQAGVNGSMSN

IETGRQPSLSQEGLSCSGAESAATVTTDRDPSVHSELARHGSTNHRRSLPPTPPYGKEPI

AGKELELSTRSVTAARALTPPEKLCETLETGDVTMASPSRGSGNSTEKSTKEAAASSTKR

KTPPAEDSESESKDFEEGKLASGRPSGAKRTRAAEIHNLSERKRRNKINVKMKALQDLIP

NANKTDKASILDEAIEYMKMLQSEVQLMSIRTGMVVPPVLLPPGMQSPPX

>Cma2027188

MASIDAFENSAQCDKLLESMRALRPLLGGDSNNHE

VFHGYMNSSPMGATSPTTISTEDLDLLDSPTMLSSLDQEMLLSSLGGDSSSASMDINCDE

WNLSSEGSFTPWKLHELQDQVNLPTQQHPCGPILTNTAANAAVAAHSNTGKSRLSWYQEN

ETNKGNKSASLPPSRPSEGKSSSLSSESHGFKGTKSAVDLLKQLDSSDNSDAMDDDDGDK

HSPNGKIPMSKNLVSERKRRKKLNDRLYSLRAIVPKISKMDKASIVGDAISYVQDLQQQV

EDMRSDILTLQTSRDGSVEGLPDCANDGMAPMLPSIKLPEHKILELDVSKMEELTYHLRI

HCKKGPGVLVQLTRALEALDFEIVNANLTSVSDHILNTIVMKVKNGEMLEREELREMILG

VVPKFGLSL*

>Cma2029521

ERRRRERINTHLATLRNLLPSSTKTDKASLLAEX

>Cma2084096

TDSHSLAERVRRKKIGERMKLLQDLVPGCDKISGKAMMLDEIIKYV

>Cma2104022

LNQHFFALRAVVPSALKMDKASLLADAASYIQELKGKVHGLESEKKVLLSEVETLSKKEI

MG

>Cma2104515

EKLVALRELMPNSSKTDKASILDEAIEYVKLLQSQLQAWSTRTGIGFPPALMPNMGMQHL

QMP

>Cma2115804

DKASLLAEVIDQVKTLKRQVSEIYEYGPMPTDLDELSVEKVSSSAESRILIRASLCCDDR

PDLMTDIRDALDTLX

>Cma2120370

ALRAVVPNVSKMDKASLLGDAVAYIEELRSKVKELESEQEELIALAKPNGEENLSSETMP

SPDQPSAIKEQSSCTSLDAKDX

>Cma2122492

TRERPLSSCESVGDGKIADPSLMQEMGNYEGHPTAPPKTAGHTSDHILAEHKRREKLSQR

FIALSAMLPGLKKMDKATILGDAI

>Cma2132468

VESEHSDVEASFKDADCSQSVVDKKPRKRGRKPANGREEPLNHVEAERQRREKMNQRFYA

LRAVVPNVSKMDKASLLADATSCIEELRAKVHELEIGNTNLLAQLDG

>Cma2143532

PNTNPNPNPSHNPSPSLSLSRNFNPRPNHNPNPRQSGFSRVDSSNELFKQFDSSDNSDVM

DDGDESGGKRKAPMSKNLMSERKRRKKLNERLYSLRAIVPNISKLDKASIVSDAITYVRD

LQKQVDEIQADITDLESQKEGAAVADSRAQDPRRIKAIVCYKKKVKQEHX

>Cma2145178

LMPNANIYMDMDPVVQAVVEGLGHCNSVKSSDKRFNFLQASASQSYNHSNIKTSEMHGHK

QGSNLSSTIVYRTCVSKAPDMQIHNHSQSVSPRIKLDFIHPSHSADDARMHDIEDGEEKL

NGISLTMKGAVCIDADSEHADMADDKKPKKRGRKPANGREEPLNHVEAERQRREKLNQHF

YALRAVVPN

>Cma2148027

SHGMKEAAKNALTRILEGSSGAVGLNESPSVWSNGSETSSPQQISCDTLSPPLVTPMGHP

PYFREVLSLSPGMEQTKPQNLFFQARAGLHERMLDTLDGAATQVNQVQDAIWHSHSQDAC

GAVAGTPRPRVRARRGKATDPHSIAERLRRERIAERMKSLQELVPNTNKTDRAVMLDEII

EYVKFLLLQVKVLSMSRLGGAGFFVPLVAGLPEGKSMHMPTAVPGEASGPSHDSIQGVEE

HVARLX

>Cma2149492

IGPETLETVDVMLTSPSGGSGNNTEKSAKDTVASSCKRKSHPAEESECESKDADEEVAAS

EKNPATGRPSPAKRTRASEIHNQSERKRRNKINDRMKALQELIPNANKTDKASMLDEAIE

YVKTLQEQLQLMSMRTGMVVPPLMMPPGMQYPSIQQRLGSSSHMNMGMGMSMGLGMGIGM

LDASAIGVGAGRSIVAMPPFLGQPVPGHAYHAAGYTPYGSSSIIHEAQGPVHINEGHNLL

STQQLQPLNLNLQMSADMYRAFMQQHQAQENLTSKEP*

>Cma2149870

GHLAPELEAPAEEDEDCEALFHMSDPDHHMNSMMDGVSAALETSADEQGASQLAATAGSA

SAATGHNGTNDTANRGGGGRGKVKGPPAKNLMAERRRRKKLNDRLYTLRSVVPKISKMDR

ASILGDAIEYLKELLQRINDLHNELEATAPDNRPGASLPPTPNSFMNQSSLTPTSTSNLP

SFVKLEECPSSSMQAMTAGPDQSQPPKIEVRTREGRALNIHMFCARRPGLLLSTMRALDG

LGLDVQQAVISCFNGFALDVFRAEVISHLRSHTKI*

>Cma2150653

QELYTRLPLLEDPNNHNSFSICKDEEATLPFTAQNVPNEPSLGSSLCGINYVSLAPQHPS

PSQYSDPSSTLISGENYFNEGQDEANCFAQQSNQASLDSNACSLPHANHSLISSSIKSIN

GLAASANHEFPCLMAFEQCGSRNNINEKKKNINMAVGDGVNVTTIHEKKKRKRTRSRKNG

EEMESQRMTHIAVERNRRRQMNEHLSALRSLMPSSYIQRGDQASIVGGAIRFIEELGQAL

QSLRLQKQLRGREDESSQVHPAKLGQLVDDDLPYFRPHFGLPYNSTLENITRSSYDLREE

SYACSKSSMANVEVKMVVSSAIFVKVLAQKRPHQLLHTLLALNNLHLKVMHLNITTIEHT

ILYSFNLEMEDECQLQTPIELA

>Cma2152538

MKGVELSYHQQRGVAFPFGFSGKEEYVNESPILRGGGGGGGGNPN

SPWLGSSNANTIRGLSFESCPSLQPSYSGFYSNNSVNPIPHTIQQMQGTLPLEGYNWSGS

SDTKDLVMHFSPYSALNAGSLVLDSSRGELVNALKMTPKEILEAKALAASKSHSEAERRR

RERINSHLATLRTILPSSIKTDKASLLAEVIDQVKRLKRQVSDVYQYGSMPTERDELSVE

RDPASRKGRTVIRASLCCDDRPDLMTDLKHTLDTLKLRTVKAEISTLGGRIKNVFVVTLK

DDTSEQDLDMSIKQVQEALRLVMERPERGELLQGRFSKRQRLASLEAQVDS*

>Cma2152588

MLSSRPPAAASLSSSSSSSSSPVSALLPSSITTISALNH

NSKDEFVCQVALDCPSDFLQCANPPIDGIGLLNNGTVSKSLLSPNAPKVRKSGVDCVQCV

EAFHTDTMNHCVPEWDAAGRDLDVIWDPTELFSPSKGKWDSGQTRRPDLAFVPDHDFEEL

CWEDGQLLVVMQGLNSKGNFKNRSHWQVSSGETAPSSSGLPGAGMDGTFDTLGTGEFMDP

SSVNLHTQEDELLSWLQHPSDELVDKLEIPSDTPTLNAPSTGCERSCPGDFIHHGIPQNT

MNAGDSSIGPASKNLRHDAFPPFQTTTLHKPGHDHGDVPQHIHVNTLTGSSITPTTIMSS

KEQFEHSSRMARLEMPPPQMHPAMAGVTCGPSVPSGSMSSMNFSNFSRPAAAFKANLHSI

GMASGPSGVARLKQLDRVAVDAFKSESIDSTTKHLRLIAPRTEGVVLLTVPNISQSIGAS

VVHPPPHSFSLGREAEQGVSANGAVNGSSGIVGKSNGWNSSVTSRAALGTDRSTEVLEPS

DTSSSAGSEESDGVDGKGAAGVGKRKGYDEDLEIQSKALESESTGTKKHATGGGNSMKRS

RASENHNQSERRRRDRISEKMKALQALVPNSHKTDKASMLEEAIAYVKALKAQIQMISYR

SGMYFSPMLIPQGMQHVQMAPWPHMGMGMGMGMSMGMGMGAGMGMLDMTMAAATTTARQF

PPMLSIPGAVMPTIHAAPAATLDSARPVHGVANKHVVLPPAHQMIVPARTGNPAAGTHDP

TPNVVSHKLFPASQQQHPVQCTSPQIVAARVHRNRTPHPHQLPQSTTFMTGMGPLQQQ*

>Cr01G01160

MLGYKLRGCVSRGARDQSQGKKLPSVCPGARGRRSSTRRAPGLGRSTVVTVRAGPVEIDEDDLIQGKDRYKWYEAKFCAE

DQPAWNIGSFVSNVQVSPLYRAVTMSVEVSRERIPLAAAYKAAGQRAVLRVNNGLERHCAAATPPFPEEINNEPLFKVRG

DLFAQEIKAVREPISVKAHLTVLVTRKEAPEVWGLGPDDVVEVGPF

>Cr02G07750

MDFPTAEPGAGDFSFAAEAAAGSRAPVSHSTVEKQRRDRINSLIDELRDLVPPTQQQQQQQQQIGVVTIGVSDNPEASSR

RPKHVVLADTINLLKALRQRVSFAAVTAELQQLPAGGSGGGGGGGGGALPLPLPVPGMYGAVAGAGMVPGMPGSGVQPVK

QEPQGSSSQDDDDMGHPGGPGVTVKKGPDCFYVQVTCPDRKGLLSDITDTLRNLSLEVRTAAVTTNGGSVRDVFEVVPPD

GAVALAPEAVQSMVQGALSQRVAEGQQEVTAGKRPRA

>Cr04G00860

MNEALDFGIGDSQYVFTDLELNELLGVIERKAAGEAEPDALDFLRATDGNGLALQFQPRSQKDNGSGCSLEQSAVAAAVK

LEDSALSSALASPVDTPALTGVADPASLYGSGAEISIMPMPHAAAASAPTSLHAYTLPGTAGHAALVGSSPALQAQHNAQ

LAAAAAGCLHVHAPLQLARFASVPAPPGKAMSMSMSMAEPKGQISHSTVEKQRRDRINSLIDELRELVPPQQRGGANGAA

AAAANDAGGLEARRPKHVVLADTIQLLKHLQLKIPCQMTQMSGVTVERGPDCYYVQVKCRDRKGLLSDIINALRQLPLEI

RTAAVTTTNGTVRDVFEVKLDDPGLSPEDVQNLVHDALFQSHLLAAQSESLAAAGKRPRA

>Cr14G02650

MQPTSLTGRQQQQLGLLGLAGRTQPDMLEAARVAALLNGQGQFNLQTSGLGSGALSGQSLNLSLGSLPTSSLQQTSQQAP

MQQSSQLGLPDQLALLSGFPAALFPQQYGSGDRDLQLGGLRNVGKTKSSDSRSSSAYASRHQAAEQRRRTRINERLELLR

KLVPHAERANTACFLEEVIKYIEALKARTLDLESQVEALTGKPVPKSLALPTGMPSVLAGGSTSADNTNASPRMVGAATS

SQGGPAGSLPSGQPGAGGAGAGSLASPSTTPPPTMTAQQASQQLSLMQSGGQAGGSQGLPSQLTLPSGGAGAGLLSAAQQ

SLLGFPQSGGLSLSGAGLSLGGSGLGHGTSGISLTQFAGNLQAAAAAAAAASHGAGSQSHSQSQSQHSGLSLGSHHVTAS

QLNELQAMQMMQSLQQHHNQHAAAAAVVAAAGGGGGSRPGSTFHPTNNKAFLHFNEDAYAFSGKPELSLPARSLLGAAAA

SAATPSTSLQLTTVQLPADSNTLLQVEMARKAASGSPVSSEESGVPLKKRKVLVL

>Ed2000242

AKSCLPVDEEEEGGQSFMALLLPEDASPCSKNIDGNRLSEEREREQRKDCNEEREMDEEA

SLLGMFKDGKDGNGDGMDESIDGSGVLHETEQVVGVVGVGGGGXXXXXXXXXXXXXXXXL

MAERRRRKKLNDRLFMLRSVVPKISKMDKASILGXXXXXLLQRINELTNELEATPDRVLS

QNPPLLHPLVLGSANIPNSIKEESSPNVVDTQPIKVDVQTREGRGLNIHMFCARKPGLLL

STMRALDSLGLDVQQAVISCFNGFALDVFRAEQLQEVPAEEIKKMLIMTASSPSC*

>Ed2001071

GGGGGEPPGNTMREASAGSPLPTAQPWFGSRKSSNRPAVEADSSKGKKPQRDNRTNSTVT

GSAVSATDGETEAFEAQEQTRTRTRTQTRSRDSTDESTEESGKEAASSLKRKKSQGDESE

YQSEDLEDAATLTVKPSTGRTKRRRAAEIHNESERRRRDRINEKMRALQALVPNANKTDK

ASMLDEAIEYLKLLQAQVQMMSMRTGMGMPPMMMTMGMPHLHHPQQVPPLTQMGVGMGMG

MMDASSGHAMVPTLANPVQPFRGPGYPVNIFRPAMPMNTNECQYYLQTPGAVEPYGAFVA

SQQFQSANANMHMYNAVLQQQHMLQLQNNHIQAQQSVGNNPGNASTKEGP*

>Ed2001072

GGGGGEPPGNTMREASAGSPLPTAQPWFGSRKSSNRPAVEADSSKGKKPQRDNRTNSTVT

GSAVSATDGETEAFEAQEQTRTRTRTQTRSRDSTDESTEESGKEAASSLKRKKSQGDESE

YQSEDLEDAATLTVKPSTGRTKRRRAAEIHNESERRRRDRINEKMRALQALVPNANXSIG

TQCQQGRSPKPVLSHLGLEMNQKVFRSKCLFAD*

>Ed2002292

KPAGIQTILCVPAGNGVVELGSTELIPEDPKFIQHIKHSFSHGIWEQAENPTHSNNNHHS

VVPSSLTTTPSPLNGHSVEHAAQKLRFVTGNRTVGSFMPPDMGNPGKTAPIVEENDRFRP

LVVAPFAPPNPFFTVTAKVFQGNSWHPRQDLEMRKGVDLSKTHQGIKITHQDAKXNLEPR

FASGLGGLHSIARIPPPPHLVERDSGKAPESEXXXXXXXXXXGGGGEEPLNHVEAERQRR

EKLNQRFYALRAVVPNISKMDKASLLGDAIAYIQELQENLKELEIEKEDLQARCAASQST

DVSSNCARIGESDMEVRVEDRDAIIEVCCPKETHPMAKVMLALQNLRVEVRNSSISLAKD

SVVHTLSVRIESSGSSSFSKNQLLEAMTGTQGLSSDPDGRIIKM*

>Ed2003762

MPTRRRLPRWRMCTPPTSSRCRCKCRYRCRCKCSSSSSKASRRSTSTPAPAP

STAALGTGLPTWPLWWATRALHTRLXTTTTTTTTTTTTAATTATCPNTSSAGGAPPCIDP

HRVDLGLNVGADLDVDRKLGRSSSRKRRAENEEGATPTPKMDDGESEDSSSNKRHRCSNK

STGGPKLGAGGKPNKCTKENNQSHSHSHSPPTSSRSTKDSSKQSKSGVVDPTQDYVHVRA

RRGQATDSHSLAERVRREKISERMKFLQDLVPGCNKTIGKAMMLDEIINYVQSLQHQIEF

LSMKLAAMNPKIEFDIESLLGGIGEMLPNVAMEAANNVEQLGSMCDMVFNPSTNMSFQML

LQTPLPPVDSTSFPKSHPGMLNVGNVQSTSIPQLPSAWDSRGMSLRRDEMLYYGSAPTEP

SAGGNNHPSMPKVEF*

>Ed2003879

SHSLAERVRREKISERMKFLQDLVPGCSKVTGKAVMLDEIINYVQSLQRQVEFLAMKLAA

LNPSLQNFNLDGYFHSEGINRGSTMVLSSNPEINTVT

>Ed2004142

SSIIGAANMIPSLDASNAMINLQKNYLAEGIGVDRLRSVEDAQIGDENSVVIPANSAAPY

CDSRKPGDAAETEEVVTVKXQEEEGSCYQREAVSTDTNSGKKRKCGSRDFTKPNKEFKTD

AETAEKCXXXXRTKSYQEAAADDKSGPKEELPASKALNPASEPRKQIQASNSGTSGESDS

KPTASDQPPTKPVYIHVRARRGQATDSHSLAERVRREKISERMKLLQDLVPGCNKVTGKA

MMLDEIINYVQSLQRQVEFLSMKLAAVNPRLDYYNIENLLLNKEVLPNRTPVSAFGAEVL

PPPPYPSLPSTCTTTTNTSQTSLLLQQQKKIVNNMPGGAELAATLNASIATAAVSSATHP

SIRSMMMMAPNSFNVADHNPSQVQQQVWGGDELHHVVQHMGGRGPFPLVSEPSSSSSVLV

GSIAAALKAEM*

>Ed2004236

EGEGEGEEEEEEKKAKRNSSTIARSGKNSSKQSLNNNSAAPPKASQGYVHLRARRGQATD

SHSLAERVRREKISERMKLLQDLVPGCSKTLGKAMMLDEIINYVQSLQHQVEFLSMKLAA

LSPQMDFQFDNLVIGEKWRSSGAMKTTENKNVEEVGNMLFNPGSNMSFQMLLNTPLTLPS

HSTDFPHFKGGGGYNEDALLNAVHAAIN*

>Ed2004237

DSHSLAERVRREKISERMKILQDLVPGCSKVTGKAMMLDEIINYVQSLQRQVEFLSMKLA

ALSPQMDFQFDNLVIGEKWRSSGAMKTTENKNVEEVGNMLFNPGSNMSFQMLLNTPLTLP

SHSTDFPHFKGGGGYNEDALLNAVHAAIN*

>Ed2004238

DSHSLAERVRREKISERMKILQDLVPGCSKVTGKAMMLDEIINYVQSLQRQVEFLSMKLA

AXEH*

>Ed2004707

MSTEADMMEKLTWEEDSNAFLEAFIGGGASPYDNANSSFWQDHSMASASTP

LLSLPLPLPLSLPPLPSPLPATPPLMNEDSLQQRLQMLVEGGHYAWTYAIFWQLTLSASG

EQVLGWGDGYFNPKECEQRQTQEGVSEADQQLRRRILRELQALIGQSNGGGSGGSGGGDS

PGHLDALEADVTDAEWFYLVSMMCSFQIGDGAPGRAYATNKHLWIRGASCDPDLQRCRRV

QLAQRFGIQTLLCVPTTAGVLELGSAESIADDHTFVRFSMSFFTDTQWDTQGIPTGPPLD

PFFSAGNDLSFMSSSESRISTRRSSSSMDGFGGATSSCSGTASTMGGSGAGQIPSLGRRD

GAGPSRMKSASSIFPDSNLLHSSGGNSGPAETDKLVSMSGWAATTTQFPKLGEINKGGGI

ATKIDSRLHSFDSSSGLIQGMDHQMGKMSASAAEMNTMQPPPDMQHLYHHNHHQTINGVD

PVHKYNDDNNRGGLTLNANEANSYKLGLNTAHVSNPYNMQVGMASQNMQQNQAHTGMKPA

ESLHGSQGSIFSQFAKPLTAASTEIKPPQVQLTRQKSEETFANYLKQEDAIPSSGTGFGA

IRSSLESELSDAEVSLKEMTDCPLTAERKPRKRGRKPANGREEPLNHVEAERQRRERLNQ

RFYALRAVVPNVSKMDKASLLADAVSYINELTKKLEQLEIEKKELAMKSGQTGGGAVGKD

ASLARNHGTKSYQSPEPSTKSGLKEQSSCTTVDVKSVSSGACPQHRLVVNVTYILEREAL

IRVDSVKSNNPVGRLTVALQELQLDIHHASVSTVQDITSQTVLVKFRNAYWNEDQLAAAI

SARASDCGCSV*

>Ed2004834

SISINNEEELMMSEELMMSVRAMCNYDSSASVGFFLPHEEEEEEDPLITATQSDTNPLSS

SSKEEQDEDNEVLEEAEHRREXQEEEEEERKEKEEEEEGRRRKRKRVKISKDPQSVAARN

RRHRISQKIRVLRKLVPGGMKMDTASMLDEAVQYLKCLKAQVRSMEALMNALENAEPFPF

PFSMPMPMPMEYYY*

>Ed2005089

MLTSRSKMDTAETTTPMTEIGTFRAEKGTSRAEKVAFRAETSTPRTEMGAFRAEMGTLRT

EMGTAEMDTSRTEMGAADMDSSRIEMGTPRTEMDTPRTERAADKVGITRTEIGASISKVP

RTSPTSGPDTALIRPLTCTKKSLKEVDGQMEDVEGESTSFKHHHRHHHHHPLPRISLKRS

RSTEIHNLSERRRRDRINEKMRALQELIPNSNKTDKASVLDEAIDYLKMLQAQLQMMTMR

SGMYVPPMMFNPAMQVPPPMAPLNPLHHHPGMGMVGPGTGMGIGIIGRPDFPPPPPPPPL

SSSSSSSSSSSTSSSSSSPSSSSSCCLAQTSNCTPGLSSCGTMPSSADSFVASGTPYYAP

LAFPYPQYQVSPQAFQPINMDAYNAFLLHQQYLQHHHQHHHHNQHHQPEQPHQQQQQQPQ

NGSPPNG*

>Ed2005658

MEGAPALEV

NEFEKFLRRMNPPRVSVDNESCETATLVKVDSVNRPGILLDVVQVLADLDLSIRKAYISS

DGGWFMEVFYVTDSEGNKLTDETRINCMVQTITVDNHFSRRVGVQTASEHTAIELTGPDR

PGLLSEIFALLTELKCNVVSSELWTHNNRVACLIHVADDCAHGPIQDDGRLYRIKEALYN

VLKGHTDTRAARTDFASSVTHVERRLHQMMFADRDYETLGSAKSIGGDVSLTQSPGNQEV

DKVTVDNCDEKGYSVVNIECRNRPKLLFDAVCTLTDMGYVVFHATIDCEGLRAYQEYYIR

HRDGCPLNSEAERQRLIQCLRAAIERRVPEGLRLELCTRDRVGLLSEVTRIFREYGLSVT

KAGVTTRGDRAVNVFYVTDAAGNPASSKVIERLQKEIGKPFLQVKDDELLREETSSTKVK

FSFGNLFGRPTQALLNLGLMKSFI*

>Ed2006157

MAITVASVSRILNSSTSSPCSIASPAGCVCLGRRSRHSPTFGSWVPTVTPDMTRVPRSPS

VPESRPSSASPPCAVVLSSLAPPSASTRTASSCTKSSSPSTTPSGSTAPIXXXXPLPPTL

RNKQIVPTPSLPPSFPPSLNPFLAAKVLHQQHHDPDTLRKAKNPTPTLHPHQGW*SEPNK

FTLGPLHSIARLDPKGGLSKEIEFSSIDDRKPRKRGRKPANGRDEPLNHVEAERQRREKL

NQRFYALRAVVPNISKMDKASLLGDAISYIQELQVKLKEMQVEKEELEARCSTSSAIVVV

GSSQLTATDASXXY*

>Ed2006218

MLTWAKGVLKYSLTKMMKTMKDLLNKVLIRKKIXLSEGDPSQTLDGRVHDVKGRTLRSKHSATEQ

RRRTRINERFQMLRQLVPNYDQKRDKXXXXXXXHDALPISPREG*

>Ed2006605

LNQQTDWLDSGVSKGGGLMRYNSAPSSILVDYARFEGILNVASEMDFPPLFPIDDDSSSS

SSSSSSSAALITHHQLHNNNRRRSLPPPTTMEDMNHSSALSAILEHLPSPHMHTLHYGHL

VASDSSDSPHPEGLIMVDNSYLLPPQQQQKRNKLIRHSSSPAGLLNHLASEVRMKNEPNN

TNTNSNTNNGHLSNLGYPDVIMEHASEGNISSHLQSASSRGVSHSYDDIFCPSSSSIIIS

PPSQSGIMRQSSLPVGIMSKLTLESICEDGDKMISSPGNSTDDANLGLLYGSFPMMNNDM

QDEAARRRRMDPHSDSLAGQMNHLGMAGXELNMDSASSSSVLCKARAKRGCATHPRSIAE

RVRRTKISEKMKKLQDLIPNMDKMQQANTADMLDEAVEYIKTLQHLVEELTISQSKCDRS

CKNRQKNSP*

>Ed2006642

MGVEDGSGDLSSDE

APKQQGKRSKVNNGKAITRSRQPEYATNHCMAERKRREKLNNQFQALKDLIRPYIRSKDD

KASLLGDTIEYIKELQGRVHEKENTPCTCQGQYPLKSKSMDPVQLDAVQSEDLRTEKAGG

YLSESLDNKKGAVIKGGDNGPTIEVRGMEDDLVMIKIVCTDRPGLFRDLMQAIHELKLQV

QYANVNTNAGIVHQVIHAKVDITKEQHANQLVADSFRCAVLKASGVTR*

>Ed2006649

DQPGDTRNTAATLSQIPVKTEDGVEVEECAFPSGGGVEPLGPCPNPITPTPSGSSRRRRR

CQAGEEPLLLHAATVEGSIPNMARVSPTPSIKMENTVGGGGMEDGSGDLSSDDIPNMLAG

SNKRSRISAAEKIPRDRREEYARNHCIAERKRREKLNNQFQALKELLRPHIRSKEDQASL

LGDTIEYVKELQRRLDEEEVVPLVCQCKNAVKGDSADLLTRLDSRKLEGQLSEPLENKIE

NKREASSSQGNCEPKIEVRGMKGELLVVKIVGSDRPGLFGDLMEAILDLKLEVPFVNLSI

NAGMAHQVIHVKVDTTNEQHTNQLIADRLRHALVK*

>Ed2007836

EEEECIHSPLTTTFPNPRSSPSSNPNPSSSPDPNSRPNCSPNHNPYPNPNLSPNPYPSPS

PNPRPSRSCNPRPSRSPKPNPSSSPNPNPNPRHRPSRSCIPRPISSPNPGSSTSPNPKPR

PSRNPNPRSSRSPNPISSSSPNQISSLNANPGASPNPSPNPNPSPSSLGDSGKAPVAVSA

EKMALAALPAKVKTGIPLEDGGGDLSSDGVERPGKRPKRNGSGSKALAAQHPDYAAVHCM

AERKRREKLNGHFLALKDLIRPHICSNKDDKASLLGDAIEYIKELQCSLQEKVSAPCTCQ

YNHLAQSDLMDLKTKSDSIQSEGLSNGKIVHVSQPIVKKNDVCIIEGDSEPTIEVREMED

DLIMIKIVCAEKPNLFEDLMLTMLGLKLQVQYANVNTNGGNLHQVIHAKVDNTKEQHTTQ

LIADSLRHAVLKNNNRHIVDSTLLIPIXPSI*

>Ed2008279

ATKKSRATEVHNLSERRRRDRINEKLKALQELVPNSRKTDKASLLEDAIQHMKMLKAQLQ

MMYARTGMDMPQMLISPGMQQLQMSPFPPPFGFTPLFPPPPPPFTMTMGTPRHHITPRHH

KPRHHPKTPP*

>Ed2008615

FFLYFFFISVCVPLSLSLSPFVCLSFCLFFSVCLCVFLLVRKVRAVQGVMSSMGRKEEHD

SERVLEDERKPQKPSQESETEQCVGDEPVGDKDNDLLSRSTPFELTDSVNDRLVMLQQLI

PHGQEIDKEEILQVAVEYVRALEQEIQTFDKSKQDGNIMEQGCLEKSPNANGNALQQRGL

CVVPISALDKVF*

>Ed2008741

RNNIASNPQWGXLNGVTSVDVSLTDGETEDLASLRVPPP

PWTRSRHSCNGTEDSGKEAAAAASSSKRKRNHGDGASHRQTVTTGRPKRSRAADIHNESE

RRRRDRIKEKMRELQELIPNANKMDKASMLDEAIEYLKFLQAQIQMMSMRTGMGVVSPMM

LSMGMQHPQMPPVSQMGLGIGMGMMDASTMVHSFKGPNSSYMPNVFHPQPTPFSTSETQY

GLQSIGAMDNAVLQQQYILHLQNSLFQPQLLQGNNHGNVAMKKF*

>Ed2009077

MKNDECCGRSCSRIEALDKSFGIIIGGNCDDGDFEGDLDELGDC

SAVLNEAAEAFDGKGDSGPLSANIVSTEAGKIKRNMPAKNLMAERRRRKKLNDRLSMLRA

VVPKISKMDRASILVDAISYLKELLQQITELQIELESQPGDQFNLVCSSGPHIITKVHNE

LPVIIRKEPPKLQNMEIPQPPKVEVKVEEEGRSLNIHMDCPGQPGLLFETVKALDTLGID

MEQAVISCFDRFALDIFRAQPLTEGSIRPEEIETVLMRMIWMQQSQARS*

>Ed2009741

PPPPPSPPSSSDHLSFQSLLQRCSRTTLPFPTEAILPSTGNPDPNPNHNQNPNLSPNPNL

YPNPNPNPNPDLDGFRDGEQKPALVFSGQMGPGNHKGESGTGAAEYEEKEIPMQFLGAHD

ASASDDSAVPYPRVKVEEGGAEDGGGGDVSSDDAPKQLSQGGGGGKRSKSNSGRAALRSQ

HPEYASNHCMAERKRREKLNNQFQALRDLIRPHIHSKDDKASLLGDTVEYIKELQRRLRE

KESALCVCQYKHPAQSDSADLSMSRSEEGQRKGKECPVSLPLENKKKDGDGPIIEVREMS

DERIMITVRCSDRPSLFEDLMLAVLDLKLQVKYADVNTNAGILHQVIHAKVDRTKEQHAT

QHITDSLRHAVLKSSLLHH*

>Ed2010936

DFCEADARNMVPDTWMTMLKSSFPPVPCVTEFRDHMGQQHYSSLHFSFDQDLQAKPAAQK

PTLSPTKVNGVREETLNSFYSTSEEGLVPCTPEYFNNGTSFGNNIIIDPPLTLGCSIEES

VGPPYGLWEATNACVENNMMPSCIDFNTLLPNSQHMLESAGSSIESEITSRMLPEYSINS

LPSPFLSAVLSDIASNPLLNDELENPISESSIGLRRSGWINNDNNVNASLLVDRYSASGS

SLQRQLVEESLENSKIGSFLCSYNGLAANLGHGYESLQPEFKCKVENGVKRKTSSDQEGG

YSLNNESCGLISDACRVDGDLDERRECSESSGKGKAKRNMPAKNLMAERRRRKKLNDRLC

TLRSVVPKISKMDRASILGDAIDYLKELLQQINELHMELGLQEGSDGVNLVSPVLPLIDP

PAIRKELPPKVEVKVEEGRGVNIHMACQGQPGLLFETVKALDTLGIDIEQAVVSCFDEFA

LDIFRAQLLTETNIRPEEIQNILMQTASCYQKVF*

>Ed2010995

KKGDVMQRFDGRGNDAKPNTPRSRHSATEQRRRSKINDRFQMLRQLVPNNDQKRDKASIL

LEAIEYIQSLQEKAKTFEDREQLKHNETSKPAVDTAQDSLTALDIKPTTAIVCPARSRLP

GELQECHSSEIMMQELDKHADVIRNSECNGDNKALTSGINISSSKSTGRLPFEPASGHRL

HPPSVLQVLINSPSDKQESVNLSQLQRDPTHFPPTISMHNQNLDIQATMNPLREHSMXFT

GKLDSRVQQSDVSSSMETQNEDTVVSSSDIKAISGDSKRPAYLSRDDQTADQELRIQGGS

ISISSSYSQGLLEKLTQALQIAGLDLSQACISVQIDLGGSGVLARSPHTQPMGSRLNADQ

TQESRGHGKNAHPLSELESQSKKPKLEKDC*

>Ed2011036

MASPGSSSLLNRALEQPQSFGMYSSSNSEQTACLGQQL

EEPEGAVSCHAFCQPLFLGDFHPVVPELQSCTDMQNFSILGSHGGACSNPTTKGSTAEER

LSARKVHKADREKLRRDKLNEHFAELARALDPDRPKNDKATVLGESVQVFKDLQAEVKRL

KTEHTSLLTESHDLTQEKNELREEKTALKHETEQLQSQLQQRMNAMSPWMAMDPSLVMGP

TPFQFPTAAMPX

>Ed2028142

ERNRRERIAERMKALQELVPDANKTDKASMLDEIITYIRFLQMQVKCLSACRLGG

>Ed2033055

REKGEKRERKEAEEEGKKRKRKRVKISKDPQSVAARNRRHRISQKMRVLRKLVPGGMKMD

TASMLDEAIQHLKRLKDQVRSMAALMNALEN

>Ed2034662

IAERLRRERIADRMRTLQDLVPSANKTDKALMLDEIIEYVKFLQLQVKVLSMSRLGATVP

GIGAEGYNNVAMGASNGGASGSSQDSMSATEKHVARLMEEDMGSAMQYLQX

>Ed2038307

MKANLHSIGIANGPSCTERLK

QMDKVVDKGCIFAHSSMESAPSGLNLALNGLNSASILQQPKTSAASVSGVPGGVESSCIG

FKPWMTQSSSVTSNVGLRSEKGQEAVEPEPTVSSSSGASGGESRGKAGKTVGSSSKRKAS

ELEDFECQSDDPDDELVDTKKPLQSRTTSTKRSRAAEIHNLSERRRRDRINEKMKLLQEL

IPNSX

>Ed2038559

QAEVFFDTSGSSNEKAYGHIAPEAGEFQLDFGFSQRCLGQESCLTDAYPGGISGTKDTVL

SPSESIHGGHKPRLFPNDMNLHRFDESTGYTPLPPAKPPGHTQDHIMAERKRREKLSQQF

IALSALVPGLKKMDKASVLGDTIKYVKLLQERLNQLEEQISRGSVAVVSCAKKKKKRQEE

SLSALDSEDSSGSSCDADESEKAEIEVRVLGKDLLIRVHCKKKQGVLGKCLLELEKLRLV

VSNASILSFTDTTMDLTLTAQME

>Ed2038678

PGKKRRHTIMTSPESLKWMAFLENNLLEDMVVGDQPDLLWCDTTLNDQIECSADNESLKK

NLDEQDKCCLRKRIRDESSTAIGSKACREKMRRDRLNERFMELSAILDPDKTPKVDKASI

LADAVQVLNQLKMEAAQLKKSNLELHESIKELKAEKNEIRDEKVKLKAEKERLEQQMKAI

VVPSGFIPHPAASFQAALAIQSQAANKGATVPGYPGRIGMWQWMPPSAVDTSQDHLLRPP

AA*

>Ed2039284

MGGCFGSKREVNPDLNSGVLLQRMDSCPAEFSSRGGGGGGLEVPTLNSTPASSESLVDRR

GFNRGGGGGGRSGFLGSFPVASWEDAVLANGNLLHGMHDGSSSINLKKRELSQLDQNMSA

RSLFQRDVYDSGLGGTNQFTLTSGNSLGRSSSSPEKLLSDCTPCQARAKRGCATHPRSIA

ERVRRTKITERMKKLQELVPNMDKQTNTAEMLDEVVDYVKSLQQEVKELRQKASQCDGSC

QIVKTDSSS*

>Ed2039330

RFPASLGLGAGFAPEPPFPAAFAYGQFPVKTEEQQQRTSLDDGSSGGDVTSDEVPDFAPT

PPSGSNPHLLHRKRPRSGGEGEREGEREREDVPPWRQQYASKHCMAERKRREKLNDRFQI

LKELIRPYLRSKDDKTSLLADTVEYIKDLQQRLDQKECVCQYSNAAAATTSAASPIDVGG

SNSSQCDLAASHSPAQTADVKSHIEVQSVKDDMVMIEIFCEEKSGLFGHLTLALLDLKLE

VQYLNATIHSGMIHQVIHAKVDASYKEEHMKQAIADCVQQALLKNLADR*LVAFR*PFFF

FFGYISMEVGFEFCHSSSCIIIQAWKRIKIRDTP

>Ed2039615

KKQGKRGKKNAVGRMNKMGSSQNITTNNNSKWMMSFFEDNLLEEEEEEVAQPAVNSFFWC

PQPTQDNPDCSMDNECCNMDGDEQGKSNSKKRMRDESCGRSGSKACREKKRRDRLNDRFQ

ELGALLDPGKPPKTDKIAILADALRALNQLKAEAQQLRESNQQLENGIKELKTEKNELRE

EKQRLKTEKERVEQQVKAASFPSPYLSHPAAAMHAASLAAFAAHSQLSSNKVAAPVPGYP

SPSGGMWQWMPPSSVDTSQDHLLRPPVA*

>Ed2039640

THSRFLLREFAQFAFVLTGISPLPTTPIMDFGDGLLRKILESGHMKPHLAAPGFPYPPSR

PPSNSFYGAHELPPHVTTPLQDTMGMGMGMGMGMGMGMGVSSLPHISTSSLVLDSARDEL

VSASKMTPKEILEAKALAASKSHSEAERRRRERINLHLSTLRTLLPTSTKTDKASLLADV

IGQLKLLKRQVSEVYELGVTPTDVDELKVEKDMASSAEVKCLIKASLCCEDRPDLMPDLI

RAFDSLGLRALKAEISTLGGRSRNIVYLTVKEDALEQDEGGPSVDSVQDVLRQVVERPRG

SDVSPGTSNKRPRLGPY*

>Ed2040222

FVGDGLNSSSLSLSSMQALLSSTTNLQSLDGNQQLYSNTNNSCQIIPPKVLRPSETYPSL

GSQPTLFQKRAAHRFTSSIPTLSPIIPPLTTTLLRRSDAFPSPKMKEQKKGGGGSLPEDM

ANGLDETDDGSGVLYETDEGGGCGAASLDKRKGKKGIPAKNLMAERRRRKKLNDRLFMLR

SVVPKISKMDRASILGDAIDYLRELLKRINDLTKELESTPAAADQRLPLLQQTGAGLIQA

LPPSIGLFPCQVKQEFPDPSSANASDTHPIKVEVRVGEGRALNIHMVCARKPAGLLLSTM

RALDSLGLDIQQAVISCFNGFALDVFQAEQMEEVAAEEIKRVLLLTAATPSPPSNLI*

>Ed2040338

SKSHSEAERRRRERINSHLATLRTLLPSSTKTDKASLLADVIGQLKLLKQQVSEVYEVGV

MPTDVDELKVEEEAPSAEGKLLIKASVCCEDRTDLTSDLIHAFDSLGLRVLKAEISTLGG

RSRNIVYLTTVDKVADALEDEEKGGPSVNNVQEALRQVMERARGGSSDMSPGSSNKRLRL

GSFELN*

>Ed2040719

MEEYIEQIFSTPTWVDLNGG

TRAPWDFNNAPGVETGALVGNGGSEESMPSLGQATSVASTWRQPYIAGIEPPVSLGVLGQ

PKAENLSPEEGASNGSHLMGKRSRDEEEDRCGPQENMFGTFMGQPQSARTGALQSMHQLQ

PMPGVSASAYGNHASLVQSQASGATIAAPARPRVRARRGQATDPHSIAERLRRERIADRM

KALQDLVPSANKTDKASMLDEIIEYVRFLQLQVKVLSMSRLGGAGASMPCITELFTEGYS

DATMGASNGVQTPSQDGIATTEKHVARLMEDNMGSAMQYLQSKGLCLMPVSLVNKMPTRP

PSASGVQDISSSPPSNAVLPTTLLPPLNTNCDGATNDLSKGVRNPPTEHQVHDHLTSPST

SAKSGGNNSKEEPVRRG*

>Ed2041096

MEPTNAYPSGF

ERVGFRSSNSSSGDFSVGGVGTKNHHPFSSFPPHISFHASSVVWTDSSKGGSGGGPDLGF

TRMTPQEILQAKALAASRNHSEAERRRRERINSHLDTLRTLVPSSSKTDKASLLASTIHQ

LKSLKQQASEICKLGPMPTDVDELKVEKDPASSIEGKLLIKASLCCEDRPELFSDLSRAI

NSLRLLPLKVEISTLGGRSRNVILLTVKGSTSDLEEEGGEPLVNNVHEALRQVMERPSVS

KKPRLGTFESA*

>Mp2000609

MWTIAGILASQVVAGGGGGXLGTEKHQSQGAEVQEAPE

PTITSSSGGSGNSADRGKEAASSGKRRTREGEESECQSEVSNLLQPQGSQVGAVAHRQDG

DDEMAQQKKPQQRGSTTKRSRAAEVHNLSERRRRDRINEKMKALQELIPNSNKTDKASML

DEAIEYLKMLQLQLQQMMSMRSGMSIPSMVMPAGMQHLQIPQLPQMGPMPPMGMGMGMGM

GMGPMGMGMSMMDMGGGAPGRPMIPHPSHSGPSHSHNVSATSPAVSMVDTHDHRLQNPSV

MESLNVFMAQQRQMQMAPQPMSMDAFSAFMIHQHQQQHQQQHQQHQLQHQHQHSLHHLPS

GMGGGAPQ*

>Mp2001754

LVLGSHAPECQTSIVAAAATFTSIRISARRRALARPRRXXXXXXXXXXXRQQPLGSTSAF

VDWRMTAGSAPKAPLPGNQQWMLKNALFHITHLYSAPRSDEATLVSADNRSRSGGPRKPA

GAQEDLNVSHVLAERRRREKLNDRFMSLRALVPFVTKMDKASILGDAIEYVKQLQKRIQE

LELRDKQAESSHLPRSHADDLSKSQLGDSTPKTSPSETFNSSHFEIVSNGNHMTGIETNV

NVSFDGASNSDPKVPSGDNEMKSSPSLASDREDPGFCNSDGESNATQAKRLDGVKANVKV

TADDDVSLELNCPWRKTLLVDVLRTLNDLQLDVFVVQASTSDGTLATVFKGKMRESTHSL

TATKTKLQTALEETAAGMGSR*

>Mp2001906

ERDEAKPPTPPPPPSHSLAERVRREKISERMKFL

QDLVPGCSKVTGKAVMLDEIINYVQSLQHQVQFLSMKLAAVNPRLDFNIENLLAKEVSRS

QGMIMGPETSYPQLHPPHQPQLQHGVPCGLEYPVQSLGNPSQVALRRSISAPIPAVIPRE

DGFADGVPTGVWDDGELQSIVMGFGQLRSLSSQALLHGHLPPGHMKVEL*

>Mp2002970

MKALQELVPNSNKTDKASMLDEIIEYVKFLQLQVKVLSMSRL

GGAGAVAPLVADLPSEGPSSYVSATIGRSNGAPGPSQDGLALTERQVARLMEEDMGSAMQ

YLQSKGLCLMPISLATAISSTNSRSQGGGAGAGAGAAGTQQQQQGSGDRHRSESGAAG

>Mp2004519

DYLFSEDFAFGDVSIDNEKELQSASTFTGSYRGKVSCQEDKISSVGIAKKSLPAGQAQNL

TRGMETGEILNPPELRQGRDILQVNDVASTRIKEEGTIKTSKHKLSSSLKSQHPEGIGTL

ASAYVRSSIESEQDIDSEAEVSYKEVECSFTVGQKPPRKRGRKPANDREEPLNHVQAERQ

RREKLNQRFYALRSVVPNVSKMDKASLLGDAIAHIQELHSKVQEMEFQIKGLEAQAKLLM

EKSSDPESSRQCFNNGSREYLHPSTVLTGTSASSASRISMDVKPTIAVHTLGHEAMIRLH

CMKESYALANLMFTLEELKLEVQHANLSILEDSMVHIILVKMKNDILPTEGQLISAIEKS

YSLTNFSSVRTHQDPYLAEGKAENNKCDIH*

>Mp2005672

DIAEGGPVPSDVDELTVDGDPSSSEIGDGRTYYRASLCCDDRPDLYPDLMRTLHTLRLQT

VKAEIATLGGGIKKGLLLLMTRSDEGSDEDDKEALSVSSVMEALRAVMERSGLGDQSPGS

SNKRQRLASLDSSSPSM*

>Mp2008486

MAASSTNSWLHTSLLSPSSQQQSAQ

HALYATDLPMHHPSSHLAQQHHVGLGQQDHHQEHLPCDTFCQPLHHSLNPDFQHSAESHQ

LNGFNGGVGGSCSGRVGVGGENGGHVSARKVHKADREKLRRDRLNEQFAELAGVLDPDRP

KNDKATILGDSVQVVKELRAEVKRLRTEHTSLLDESRDLAQEKTELREEKASLKNETEQL

QGQLQQRLRIMLPWMGMDPSLMMGAAPYAYPMAVPQPGPPSSEGTQTSTAPPVPQSVVAP

AAYIPMAPTAPGTFAVHPALQAYTMFGNRPGEGGAPYMPYPHYPPQVNSHSHVERPYAQY

PSPVQPLPGYLMQMQPHQGQPPISGAPPGPSMYRSYGPGMPLMPPQPQANQQSVQARASQ

HPVAPYAHYPASGMQPYSGPSLSSSPQASPGQSVQTTLQLQTPSQPQPQKPFEASPVNST

DGSSKAQSGGSGSASSGHKKEPKQAAASGVSDSIASSPISSAPAASVSSTGNGGDAGHLD

TGACPNRDTSQTHDTSFGRPLVK*

>Mp2008643

MAELNTTL

WDLLQTIQWSYAALWTLTPDRRMLKWQEGWFNLKAAGDTNGSDSDNKNAELFYFLYNQLT

FSPGVGFAGHALQHMGDRILWLAGDMALGSIPNPVEKQSFFIKAAGIQTTICIPLEDKVL

ELGTINLVPENDEIAHCIKKIMYSLLDRYLKKPEPCHFPDIQFGLEMNNLTPTSSGAGPS

MVIPLPNDFSTLSSKPMNLMAMNLPTVYDDLDASYIVPQVAMSAPELMQSRLSWTPEQLS

MPLGHNDLDIWSTASTLAALQSHGISTSSASSTLMVDHIEQALPGFTLPMPFIESDVTLL

APSPSPPIVEATTSLRDRSSSRSSLRSGAHLEDNEISSVAKVYADCQISEPKLLAEAGRE

SDSRVSSGRLRRSPSPKEMSPDSLSSLPKPRAPSHGLQQPLMFHPGSTSEPSHQRPRLER

QNSSTLKPADSTPVLVQLGNVLQLGNASGQVAEAPPQETDEAAVSHMLAERRRRIKQNEN

FSQLKALIPSATKMDKATVLAETIVYINSLKARLEALEQCNQSLKFLIGDRATDIAGPSN

APAIRLPTPPVPSSREPERVEVTPDGEGNLRIEIRSPCAQPENAIHILTSLKQMQLRVLS

SQMSIDDNRITATFVVKMKRKNSTGQNEVCANVTSELRKCLGL*

>Mp2008644

MAELNTTL

WDLLQTIQWSYAALWTLTPDRRMLKWQEGWFNLKAAGDTNGSDSDNKNAELFYFLYNQLT

FSPGVGFAGHALQHMGDRILWLAGDMALGSIPNPVEKQSFFIKAAGIQTTICIPLEDKVL

ELGTINLVPENDEIAHCIKKIMYSLLDRYLKKPEPCHFPDIQFGLEMNNLTPTSSGAGPS

MVIPLPNDFSTLSSKPMNLMAMNLPTVYDDLDASYIVPQVAMSAPELMQSRLSWTPEQLS

MPLGHNDLDIWSTASTLAALQSHGISTSSASSTLMVDHIEQALPGFTLPMPFIESDVTLL

APSPSPPIVEATTSLRDRSSSRSSLRSGAHLEDNEISSVAKVYADCQISEPKLLAEAGRE

SDSRVSSGRLRRSPSPKEMSPDSLSSLPKPRAPSHGLQQPLMFHPGSTSEPSHQRPRLER

QNSSTLKPADSTPVLVQLGNVLQLGNASGQVAEAPPQETDEAAVSHMLAERRRRIKQNEN

FSQLKALIPSATKMDKATVLAETIVYINSLKARLEALEQCNQSLKFLIGDRATDIAGPSN

APAIRLPTPPVPSSREPERVEVTPDGEGNLRIEIRSPCAQPENAIHILTSLKQMQLRVLS

SQMSIDDNRITATFVVKVRQTGYGMDIALICLMIHNAKIPIQWNDSLSFTIFTRLACVCV

CFW*

>Mp2008660

MGQGEQLRELVRSMGWSYAV

LWSIPPLMTELVWTDGWYEASNLKVNVERLFNSSYKTCSFAPGYGYVGKVFTQGQHIWVT

GDAVHRQSTTSGQATFFSSAGIQTLLCFPWFNGVVELGAEGLVPQNDDLLQQIRRFLSNT

PASKLQQQQQHLRGIPAGDDSCHSSRFSSLSPSQGPLTSQGLSSDDAEGGGLSDTTLNQP

VDSCPSLSMEDRMRQHFRLTAMSGIDGSCYWAQGPAAVDAEILLPAKSSSMSQVTGAAAG

GLHHQGAHSGITGMIGADWRLDFGSDIHQLTSSPTSTLQQAVFTPLPGTEHEEFESSSFL

VASSQLNQTAKSLYQQQQHLQQMAAKSPTSSGVEISSNQQQHLRGFLSTKKTGTPAFKPW

KGQKQVIGVKRQKSQSNQVLLQRSIKMVHQISLLNQKKDEKKRTEQMALAMRRAPTEARE

YHSCHDEAAINHMLAERKRREKQKENFSALRALIPFVSKTDRASILGDAIQYVKQLRSRV

QELESINRELEAQIPNSQRKPNSSV*

>Mp2008678

MHSIVGRGRRPAFREEQGWKEHSSPHTPLSPNFHADPLFGT

PSGSLQDILRSVHMVEGSAFVSVPKSSGSQAPPDIIAPQQQQQVSSTARNVSVDAPVIDD

GLGKLQRKQLQCHTSSSWDHDMRNEHVFEGYPHVGLADISNTQLQLDSFKPVGTWTDTPG

ADHEQQWTDLRRSSKTTMLSPGPNLDSGTCQPSVWRERWSDGSRVHASSHQERHRAAAPA

AAGATAAAAAHLEESTDGGSDPMDEKNSRRFSNTAGYCSSRNLFSERKRRKKMNDGLYSL

RSIVPNISKMDKASIIGDAIKYVMELERQVKELEGEMELDEQRKNKINSSSDTLGSLAAT

GVQHGGDAAAQNDKDVDWNKNSGGNVSKLDTKDQPNVTLELSKMKTGMYNLRMFYQQQPG

IFMHMTQAIESLDLNILNSHVSTCDGYVVHSMIAKMTSWDDTVVGDVEKHILGLVRQYQ*

>Mp2008820

GPRSSFHRRIYLAITTSTAPADCHSLPKPKVSFSSCCIKLLQESRNENVLHQIQRWLSSV

TTPLNIQPSPGAVGSENDSGGSREAMQMSPSIASSMFADGSNCNVHFGPSSDILLKRNNS

GQGSGAGGFSVSGHENASASSGAPGAGNRVQVQVQVRKKNGLGRYPPHSHDAISLLNRHS

QDVLSQRRSISGQEKTLEASSFVEEWGPAAAGTSSIFQQVVGHDSDPIISCSGMGIPGIS

SLMPPDTFSQQQQQQQQRSHPRGNASISLDQALLTSLPMTMATDISASLALPMDMLQDST

ANLYGLHSKNSREWPTLEPSKEQSLVQMLEEFTDDDHLQRLQSPDHALIQTAVASSSDTM

AEWASTGTASISHQQQHHRHYDHHTQQRLQQQQRQQQQQQQQRVVQRQFEFPNNPAVSSL

QNMTTPRPALANLISHQQHQHQPQQHRPQLVPHGGGQQPQLLPLGQQQQQQQQVNQPGPP

SWQQNISFTTPTAWVRWVSRKQASLLRYKARDPNGTSPAALRVNQMVLKRLLTEMVPKIA

KEQKAREQEAGQALDTETSSHGHRVRMISYPTTTSTKPSESGTLIQDPQARTHMLAERRR

RGRENEHYAALRKLIPPTSRMDKATVLAETIKLVRMLRRRVRQLEKGNPDNSGTPNLSLV

TSEDSQIGTTSPSSRGEPGSPIPARTGAGSSEPELQEGKPPGGRFLDSCLVPAPAPSQED

NKEIVEVDRRACIPTELRAKIRTSHRPDTVIHILTCLKDLGLEVLSTNISIEKDWVYADF

QLKMNEVHICRVSQFCEELQEAIRKAIFPAESSNADSTLDGRSHSQDS*

>Mp2009030

MTTTETTVSCSTRRRQLQNSHLRRRRIWECRTQACQTHQLEEVWX

XXXXXXPPYQPNFSNPGPLQQINSFGLDTISFAPSLLNGLMREQGPELNLQQAAAAAGSW

DLSLEQQLMQDQNMIPANLKHKLTGPTSALTKSGSCVASFEQALFHPHAQHIPQRRALPV

ESTGKLLSDFQAQLFNQSQLQQQSSLMQNSAETVVSDSPHHSPVPGQPLPDSLKHAHSPD

IHWRSVTHSKGIGSPRRLLTVESGKGLAEVSNIDDPSLKCHQLASPSEESHQHNFTDLIP

PSNHGHGQQTQYLAELTGEAHTGQSADHSWSSEAISPTISEQEQDPEDGYSLWEDTSETL

ENGRFLGDSFKHDRSKKARLCGLNMNGHHQSNGTANGMGSFLDHPHMDERTRQENEKFQV

LRAMLQPHATCKGDKATVLGDAIEHMRDLQRRVKESESTTTSSASDDGQNPRSLERSSSL

TTSLVQGVNHVNLRRDYGHRGVRQQQYGRSGGGDNAPLLKLSDVCLDTNEELDGVVLAVL

DDIPQIEVRMMENERVMIKLSSKDRPNLFTDIMFALQDLHLQVQHANIRTVAGMVHDVFA

AKVDVALLEKQSTHTIAEAIRRAIERGNSPNSCSPNPATPDANPNVNSPRKRSLQLT*

>Mp2009677

MAFSSSGT

PMSMKEHSLGVSIGPGASVGVSSAGYTTFAPASHDFQGPVEFSQQQREYLAQRSMMGQAA

EQQHSQGRGQGGGHSQGQHINMNEGAKPSTVSTHDFLSLYADNSNQYAEHHDNMRGLSSA

LTTKDFLQPLETRGNHAAFREGQPSRGEKHPRQPEGLPHGSTPAPPAVVERLLSTGMNVY

NGGHGVPNYGRVAKGAIDLPANTMVRMVGEGNVEMRGRGGFNPSAGSMVTREDHDARNDL

PKDQDAAPRGDNLAYWSNVRMKYAMLPPELNYHSQPNQPTSHVVMENVQQPPVPNMPKPW

PQGVQGVIDMTKRPTRVLSEDDEDDEEEYGDRPRETRSKEPLSKADPAQKVDGKAPENRG

GTPRSKHSATEQRRRSKINDRFQMLRELVPHSDQKRDKASFLLEVIEYIQVLQDKVRKFE

TAEQGRHQDRLKNMVWDLCKRGGSGPGDTSIEATCMDALAYAKEFVPSSAGLPIDVKQVT

EAAEVVRQRAVANADENGSTELSHNPQHTHKSGHHGASSSHDEKEGAAVLGKSPLSSQQH

QGQTTFSSRNLPHASQSKQLSAASDAQANVAGEGQVQNSSTIAPATGAYDQHDKANEVEP

SQQTRTQASTQQSQGEGDTSDQQHPDTRHRHGTTTAEDTGNGHAASFTDDQSNTKPHMKS

DRQHGYGSSPRSTHNSQATDNSQANWQGQQHGSSAENSQQKEESRGATTSAQGQDPLTIQ

GGVINVSSVYSQGLLDTLTRALQSSGLDLSQANISVQIDLGKSISSAALPPSTPKVSGQP

AESSGQRHQQQSESHARQQGGSSTDPEHPPLKRPKLEMEL*

>Mp2009885

MDSFTAADPNLHEILERG

AAAAVSAAPQSGTGPDFLTALGDGVGVGVGVLGPHSSSSDSFLRQAEVPKWPAAPRWMIQ

QHIGPPPSFPLAAAAGAAAPVPAAVVVPSAASGGSKPSKPASLAGPVPGPGPGSSDDRPA

QVWCDRNSKQALLKSEGENAPPSKLIAMAKCDSHNTNIINNNNSNITNGSKSSMVLDTLP

AAAAAAADSPRGNVIAKDGVDVDLKQIDSSDCSDHMEEQEDKATNRTVSKNLVSERKRRK

KLNEGLYTLRALVPRISKMDKASIIGDAIDYVRELQKEVDDLQTAISEVEASRKSGNLDT

EEMSQNAQIMENSMVTEDFEKASRNNTPGSNDDTTTEQKILKLDVAKMEEKSYHLRIFCT

KGPGVFVLLMRSLEALGMEVLSANLTSFEENILNTFIAEIKDLEAMKTEEVKAAILDVCA

RFGLQTT*

>Mp2010146

MNVNPSSAPFLMSSNVRSIYQMQQPVLAGLGAPADLTKVAGPSAKGGLMA

NMAPHHLLFGPNFPAIANLDPAAIENSRPKRRNVKISKDPQSVAARHRRERISDRIRVLQ

RLVPGGTKMDTASMLDEAISYVKYLKLQVQTLESCGNAGFDPRMPYPAYYMPVRSAQGDC

STMSQISPGGGSADLGIYGAGLGQSTMKFPQFCH*

>Mp2020606

MGGEATGRLDLCSQWQITPDSGFIRPFNKPVLEEPNFAMPALPNSGVWPGAQSLATCTSS

GPSALFSVSTAAASRGLLDHHTASRSMQFNHLLPKQEAFISSNFPSMLGQSTSSLGYDRL

PQTQLFGSPLLPNAEHQLVGRSSLNSFSYDDGLVHGSSSSGASRERGKQQLPPPSGPAQA

NGGSLVLDVSKGELINLSKLTPQEILDAKALAASKSHSEAERRRRERINTHLATLRNNLP

SSTKTDKASLLGEVIDHLKFLKRQAADIAEGGPVPSDVDELTVDVDPSSSEIGDGRTYYR

ASLCCDDRPDLYPDLMRTLHTLRLQTVKAEIATLGGRIKNVLLMTRSDEGSDEDDKEALS

VSSVMEALRAVMERSGLGDQSPGSSNKRQRLASLDSSSPSM

>Mp2038954

MDKKPLIIPGD

NRLDSSQNLHHSVLVGAAHEMFQSGHGLGTSSNMTPGSMCNLHPPLSSFGAESNLHELLA

HASPHAGSHAQSLNHSSNHPADFLATLGDGVLGESSDNHMFHRQMEFSKQKWTTPSKLTP

GPFWMLPHHLDPANYPLPGLAVSGTSSPKTNVGPSAGEDSSNPNRQVWCERGANQVLMKS

EDNVATLKLPMVVKPDRSLLCDGSGAPVDSPKGANGNLINRDTSDMDAKQLDSSDCSDHN

MDEQEEKDQATRNGAGRTVSKNLVSERKRRKKLNDGLYTLRALVPKISKMDKASIVGDAI

DYVRELQKQVEELQADISDIESTKPTTGPGGEASLSSVAEPSTAQGSTAPSSGRSWEEEG

HRTSEGKPRQEMSEAPFEDTTEQKILELDVAKMEEQIYHLRIFCTKGPGVFVQLMQSLEA

LGLEIRNANLTSFQENLLNTFIAEIKDWEMMKTEEVKKAILDVAARFGLQP*

>Mp2039223

MEVVSTPAG

NGAKQWLSELGLEEPSLVRHLDSLFLPTSNQYQSVKGYNSLNSYELGGSFLNVKQRDASP

VTMNNTFTSTGYSSHWDALLNNERPAKMPKTMSWDGSFSNQGMGNSLTHQPLLPLIFSHK

QQQQHHEPQIQHQQHHQHQHSHMNQFFSPQQQLCQGIDGFVASLPKLSDDLHDSVAPRTP

TMEIGGSQVLGGGAQDTCNSDVHSVEAMSMSSTCNRDSSPIADSFRGKRKPEFFPSPTDM

RVQRPPPMATTRAPSPPSKATGHTQDHIMAERKRREKLSQRFIALSAIVPGLKKMDKASV

LGDAIKYVKSLQDRVQKLEEQAPKKNVSSIAVVKNGKDSKGSGSDGETGADSEVVESGET

PDIEARVNGKNVLIKMHCEMKIGSVAKCISELEKLHLIVLNANILSFTNTSVDLTLSAQT

EDGHEVTADEIVAALQSFFKTL*

>Mp2039284

MEFAPSSTSAMQWLSEMGLEDPSLVRHLDSLFPPTSESLVNVVGHQ

NGHNLSFPSLMPGFKTGNDMKIPSAVPPHNFSSYSSPLDSLCNLERPMKMLRSNAWEGNI

SQQGQRSSTVPIYSGKPQHRMVQFLQPEKIGNHRTEGFTSLSNIPDETLESVASTTTSIP

CRNGLRIQENCNDVHSSIETESNSCGRESPNSLGESMPGKRKPDMDYVDVKPDRVNAVQA

PFTTVKGSGHTQDHIMAERKRREKLSQRFIALSAIVPGLKKMDKASVLGDAIKYVKQLQD

RVKTLEESTPKKGLRTVYTVKKQSRNASDEEDTSENTTVTTVEDEEQAAEIEARMIEKNV

LIKLHCEKKKGVLVKSLAELEKLRLVVVSANILSFSATALDLTFNAKVEEGSELTADDIV

NALQKLFKTFT*

>Mp2039334

MDVSECYPSSLTSLLPNVTGALPYPCSPGVNAGPINQDFRLTRSVSGSLIPSSQQR

LGMEKSIPLGSGWNSVDHGNLGLTVDPAKGGFALQGVGLTSSPIARFSCNPGFAERAARF

SAFGNGHFSQSYSHSDNGSVRSGSYENENKATKLSARGEISTGAIGSPEVSCRSQANALD

VGNDGEDTKSRLYRSTSAACSPAQTDEGCNAHDNERPAVGSPSESGATNGSRGQETSKID

ADQGRSTADQLSGKKRKASGDEAAKEIASPVTKEQKNSGAQSKRSKGAEDKEDQKSKAER

SDNSSSSSLAAAAKDSKHVDHPKTDYIHVRARRGQATDSHSLAERVRREKISERMKFLQD

LVPGCSKVTGKAVMLDEIINYVQSIQRQVELLSMKLAAVTPRPDFNIDSFLTKQLQMGQN

CGLPNMVEHTESTIQYPHLPSLHSQQSLHMQQVLSNVLDNSQSVESTLRRMNSAPPMSLP

RAPMDIFGDGLSQLTGLWEGDLQSVVQMGFTQGRPMTGLGAFQCDLPPGHMKIEI*

>Ol17G00340

GGASVGTEKPVPIVIIDNKSDAFATVVEVSFGNYLGELLDTVAALKNLGLDINKGDVQMSGDSTKTSKFYVIDRENGEKV

TKSERLEEIRQTILTNMMAFHPEAAEYIQAKAPTRAGGEGVLGKVKKKVQTGIKCAPERYHSKLEIETTDRPGLLVDVVR

TLKDLSLCVVSAEVDTIGDKASDIIYVTHKGGPLSPPMEQLVVNSLSYYLSLTEEESY

>Os01G06640

MAECQPLQLQEGKKLQELQPYDGCNPSVYRGPILLPRQANSAPPAVPPEMSSSSGSGRSATEARALKIHSEAERRRRERI

NAHLTTLRRMIPDTKQMDKATLLARVVDQVKDLKRKASEITQRTPLPPETNEVSIECFTGDAATAATTVAGNHKTLYIKA

SISCDDRPDLIAGITHAFHGLRLRTVRAEMTSLGGRVQHVFILCREEGIAGGVSLKSLKEAVRQALAKVASPELVYGSSH

FQSKRQRILESHCSIMSI

>Os01G11910

MGSAPFGDAVAGGGLYEYQGYHGGFAGGHGLGQPAGRAPALDDGETEGMDASAAAAVAAMEMAKRNCGGGREEKAAMALK

SHSEAERRRRERINAHLATLRTMVPCTDKMDKAALLAEVVGHVKKLKSAAARVGRRATVPSGADEVAVDEASATGGGGEG

PLLLRATLSCDDRADLFVDVKRALQPLGLEVVGSEVTTLGGRVRLAFLVSCGSRGGAAAAAMASVRHALQSVLDKASSGF

DFAPRAASLLGSKRRKVSTFESSSSSS

>Os01G13460

MVMKMEADEDGANGGTGGTWTDEDRALTASVLGTDAFAYLTKGGGAISEGLVAASLPVDLQNRLQELVESDRPGAGWNYA

IFWQLSRTKSGDLVLGWGDGSCREPHDGEMGPAASAGSDEAKQRMRKRVLQRLHSAFGGVDEEDYAPGIDQVTDTEMFFL

ASMYFAFPRRAGGPGQVFAAGVPLWIPNTERNVFPANYCYRGYLANAAGFRTIVLVPFETGVLELGSMQQVAESSDTLQT

IRSVFAGAIGNKAGVQRHEGSGPTDKSPGLAKIFGKDLNLGRPSAGPGTGVSEADERSWEQRTGGGSSLLPNVQRGLQNF

TWSQARGLNSHQQKFGNGILIVSNEATPRNNGGVDSSTATQFQLQKAPPLQKLPQLQKSHQLVKPQQLVSQQQLQPQAPR

QIDFSAGTSSKPGVLTKKPAGIDGDGAEVDGLCKDEGPPPALEDRRPRKRGRKPANGREEPLNHVEAERQRREKLNQRFY

ALRAVVPNISKMDKASLLGDAITYITDLQKKLKEMEVERERLIESGMIDPRDRTPRPEVDIQVVQDEVLVRVMSPMESHP

VRAIFQAFEEAEVHAGESKITSNNGTAVHSFIIKCPGAEQQTREKVIAAMSRVMNSG

>Os01G39330

MDEVWCSLDSMPADIAAGIHQPPPPDLHDPSFWPAFADCAASFIAGGGGDNACFDELMAGGSSGDTRMVAMDDGDDGSGF

LVGDAEAEHLMLSSSSPSSLSSGRSLSIDSAGSMSSFSLDAAAALAMSTLAVPHPYPPPVAHGMFASGDGGGGGGGAVDD

HEDAIMRAMMAVISSASASPSSSGGSASSPTPFSRDSGAHHQPAGQPAMAAPQQPRGGNGGHVVVKSSSSSGGLAVPMDQ

KPGGGGRGRQQEEAAAASATNSSQLYHMMSERKRREKLNDSFHTLRSLLPPCSKKDKTTVLINAAKYLKSLETEITELEG

TNTKLEKHIAGGGGAADAAMRARRAQQRAKVQISKAADSQSQQLVSLTVMVMVECDVVELVLHILECLRWMKEISVLSVY

ADTYSPQLLLKAIANIKLQIVGGDWNEASFHEAMTKAANDATISCAPLAITAAQ

>Os01G39480

MVPEEPNVVNRITTAFWEFQLLACSDEPISSGTPSSPSSPLTKETGDANTVLIDDLFLAHSDAILAGGDQEDHQLGNDLG

QQQAATAMEIDDDMIYSLIRNWDNDSSSSWIELLDHVVVSPASCFVPWKRTELDKQAVAGGGEAAQRLLKKAVGGGGAWM

NRAAGSSIKNHVMSERRRWEKLNEMFLTLKSLVPSIDKVDKASSLAETIAYLKELERRVQELESGKKVSRPAKRKPCSER

IIGGGDAGAVKEHHHWVLSESQEGTPSNVRVIVMDKDELHLEVHCRWKELMMTRLFDAIKSLRLDVLSVQASAPNGLLGL

KIRAKVVSLT

>Os01G39580

MSRVPTSWGFVYSRVNKVSEEPNVVNRITTAFWELQLPACSDEPISSGTPSSPSSPPTKETGDANTVVIDDLFLAHSDAI

LPAGGDQKEEHQLGDDLGQQQAATAMEIDDDVIYSLIRNWDNDSSSSWIELLDHAIVSPASCFVPWKRTELDKEAVAGGE

AAQRLLKKVVGGGGAWMNRAAGSCSIKNHVMSERRRREKLNEMFLILKSLVPSIDKVDKASILSETIAYLKELERRVQEL

ESGKKVSRPAKRKPCSETIIGGGGGGGGAGAVKEHHHWVLSESQEGTPSDVRVIVMDKDELHLEVQCRWKELMMTRVFDA

IKSLRLDVVSVQASAPDGLLGLKIRAKYASSAAVVPAMISETLRTAVAGY

>Os01G50940

MSWSETDAALFAAVLGHDAAHHLATTPPHLDAPEGSPSSAELQASLHDLVERQGGAWTYGIFWQESRGAGAASGRAARAV

LGWGDGHCRDGAGHGEVGAAERSVARKRVLLRLHALYGGGDEDGADYALRLDRVTGAEMYFLASMYFSFPEGSGGPGRAL

ASGRHAWADVDPHPSGSGSAPGWYVRSSLAQSAGLRTVVFLPCKGGVLELGSVVAIRETPEVLRAIQSAMRAVPAPPEDF

MRIFGKDLSPGRPSQPMGCDAPWTPRLVVQTTPVRPAKKEVVKAKPAEPPKSLDFSKANVQEQAGGQERRPRKRGRKPAN

GREEPLNHVEAERQRREKLNQRFYALRAVVPKISKMDKASLLSDAIAYIQELEARLRGDAPVPARADGPAVEVKAMQDEV

VLRVTTPLDEHPISRVFHAMRESQISVVASDVAVSDDAVTHTLMVRSAGPERLTAETVLAAMSRGVSVTTPSP

>Os01G70310

MDEAEAAAAAAKMDELAGGGGGGGGDWSYLAADALAAASFTAFPFHHHHHHHHRDVLSASTPSSLLLNMDAATAAAMFDF

QAAFPSSSVPPPPPTTTAALPPFHDFASSNPFDDAPPPFLAPPGQKLGFLGPPGGAFGGGMGWDDDDEIEQSVDASSMGV

SASLENAAPVAAGGGGGGGGGGGRGKKKGMPAKNLMAERRRRKKLNDRLYMLRSVVPKISKMDRASILGDAIEYLKELLQ

RINDLHNELESAPSSSLTGPSSASFHPSTPTLQTFPGRVKEELCPTSFPSPSGQQATVEVRMREGHAVNIHMFCARRPGI

LMSTLRALDSLGLGIEQAVISCFNGFAMDVFRAEQCRDGPGLGPEEIKTVLLHSAGLQNAM

>Os02G02820

MGRGDHLLMKNSNAAAAAAAVNGGGTSLDAALRPLVGSDGWDYCIYWRLSPDQRFLEMTGFCCSSELEAQVSALLDLPSS

IPLDSSSIGMHAQALLSNQPIWQSSSEEEEADGGGGAKTRLLVPVAGGLVELFASRYMAEEQQMAELVMAQCGGGGAGDD

GGGQAWPPPETPSFQWDGGADAQRLMYGGSSLNLFDAAAADDDPFLGGGGGDAVGDEAAAAGAWPYAGMAVSEPSVAVAQ

EQMQHAAGGGVAESGSEGRKLHGGDPEDDGDGEGRSGGAKRQQCKNLEAERKRRKKLNGHLYKLRSLVPNITKMDRASIL

GDAIDYIVGLQKQVKELQDELEDNHVHHKPPDVLIDHPPPASLVGLDNDDASPPNSHQQQPPLAVSGSSSRRSNKDPAMT

DDKVGGGGGGGHRMEPQLEVRQVQGNELFVQVLWEHKPGGFVRLMDAMNALGLEVINVNVTTYKTLVLNVFRVMVRDSEV

AVQADRVRDSLLEVTRETYPGVWPSPQEEDDAKFDGGDGGQAAAAAAAAGGEHYHDEVGGGYHQHLHYLAFD

>Os02G15760

MADGGGGGLCELFSDDHRDIRHVADADLFRILETWEECINGGAGGGGGGVSLAGVADQGAAAASTGAGGGARTTTTTTAA

NGRRREGRDEEKGGGGGGGPPAQKKQKGSSSSSSSPAALAAAVGDGDGAAKMSHITVERNRRKQMNEHLAVLRSLMPCFY

VKRGDQASIIGGVVDYIKELQQVLRSLEAKKNRKAYADQVLSPRPSPAAAALMVKPTPPISPRFAAAAAAGVPISPRTPT

PGSPYNKHAAAAATARPPHPAAATSSCSVAYSMSPAMTPTSSSSTTTTTTHELSPAPAFLPILDSLVTELAARGGASCRP

LVIPSSAAAIAGIVGVPDVRVEFAGPNLVLKTVSHRAPGQALKIIAALESLSLEILHVSICTVDDATVLSFTIKIGIECE

LSAEELVQEIQQTFL

>Os02G35660

MGGFAYPFTPSPAWSRDAVFAGSPWAAGGVSSLADALVSYGAVDDEEAAFLGKTAASSPSTARLHEQQQLLLEAELLRHG

DGLGFAAMDDDGGAAMLGALEPCAMPLTDSGGPPVICSSSSNDSSGSEHSAAMPAGGGFLVGEQQQHVPPAAYAAGGVLP

SMAAGEETPQSFGFGSLFNGDLLQEATVSKYHHHQQQQQLGVVPSSQPHHLNDDIDFNTGKLMSFASGQQHVTPSIDSLQ

IDQKEFSSGLHHLNLSSLISGPLASFNATQSHRQPAEACGGKNGGAAPFVNLSEVLPKGNGSGSAGNGAPKPRVRARRGQ

ATDPHSIAERLRREKISDRMKDLQELVPNSNKTNKASMLDEIIDYVKFLQLQVKVLSMSRLGAAEAVVPLLTETQTESPG

FLLSPRSSSGERQAGAGAVTGGLPGDQPELLDGGAMFEQEVVKLMEDNMTTAMQYLQSKGLCLMPVALASAISAQKGTSS

AAVRPEKKKNGDGDGGGDEEDVKGEFDAPRRPPVGRPKEMRSRV

>Os02G55250

MQPNARQMRGGGGGGGGGQDDFFDQMLSTLPAVWSELGSGKPAWDLTAGAVGGGGGASDDHSAAAFDDSALLASRLRQHQ

IDGGGDKPIMLQLSDLHRHHGLAAGDDSGGAAGFLPLSLFADRSQDDIDAAFKSPNGARGDHALYNGFGAAGMHGAAAMQ

PPPFGQGGSMPAQSFGGGAAASGGGGGGSASAAAAAGASSGGGAAAPPRQRQRARRGQATDPHSIAERVYHSPTTFPFSP

PFFIASMPCCLRRERIAERMKALQELVPNANKTDKASMLDEIIDYVKFLQLQVKVLSMSRLGGASAVAPLVANMSSESNG

NGNATSSSGNGEAANGSSNGDNNGGGTLRVTEQQVAKLMEEDMGSAMQYLQGKGLCLMPISLATAISSATSSSLLPRTGG

GAGGSLHEGGNGTSPPLVNGTATGCDDAGVFFSVKFVVELSFLLLNEDCRGKEESKLLVQKGP

>Os02G56140

MMDGRGSQEEEHLDLIMRHHASMGLDRCESEEALGSSESEQPTRPARPRGKRSRAAEVHNLSEKRRRSRINEKMKALQSL

IPNSSKTDKASMLDDAIEYLKQLQLQVQMLSMRNGLYLPPVNLSGAPEHLPIPQMSAALDQNSAKASDPSVVLQPVNQTS

GALLPFELASQHKPLFLPGVPNATALEPRFLVESSRSNLQSLRFTEPAEMIYPDEMMLKHRLTSASESTIVPGTDEKSVR

QNTYMMNADRFDRYALSKDQLQHIMPKNTESVLDMPHLQR*

>Os03G04310

MELDEESFLDELMSLRRDGSAPWQAPPYPGGGGGGGGGGMMMSDLLFYGGDGGSAEARGGMDASPFQELASMAAPPPQHPHEEFNFDCLSEVCNPYRSCGAQLVPSEAASQTQTQLTPLRDAMVAEEETSGDKALLHGGGGSSSPTFMFGGGAGESSEMMAGIRGVGGGVHPRSKLHGTPSKNLMAERRRRKRLNDRLSMLRSIVPKISKMDRTSILGDTIDYVKELTERIKTLEEEIGV

TPEELDLLNTMKDSSSGNNNEMLVRNSTKFDVENRGSGNTRIEICCPANPGVLLSTVSALEVLGLEIEQCVVSCFSDFGM

QASCLQEDGKRQVVSTDEIKQTLFRSAGYGGRCL

>Os03G12760

MSTLSNKVSLSSNLLNLQTGLAEEPEELTYMYHQEEHARMQEQFAGTPLVEQPVRFDQFYPASMAPNQFHPSHCSSFPAFGGSSALPSLAFGAVATTKKEQVQQPSPSSSNVLSFAGQVQGSTTTLDFSGRGWQQDDGVGVFQQPPERRSRPPANAQEHVIAERKRREKLQQQFVALATIVPGLKKTDKISLLGSTIDYVKQLEEKVKALEEGSRRTAEPTTAFESKCRITVDDDDGGSA

SSGTDDGSSSSSSPTVEASIHGNTVLLKICCKERRGLLVMILSELEKQGLSIINTSVVPFTDSCLNITITAKARLALPVY

YS

>Os03G12940

MKLRLRSMDQRGGAGGAAETHRVQLPDTATLSDVKAFLATKLSAAQPVPAESVRLTLNRSEELLTPDPSATLPALGLASG

DLLYFTLSPLPSPSPPPQPQPQAQPLPRNPNPDVPSIAGAADPTKSPVESGSSSSMPQALCTNPGLPVASDPHHPPPDVV

MAEAFAVIKSKSSLVVGDTKREMENVGGADGTVICRLVVALHAALLDAGFLYANPVGSCLQLPQNWASGSFVPVSMKYTLPELVEALPVVEEGMVAVLNYSLMGNFMMVYGHVPGATSGVRRLCLELPELAPLLYLDSDEVSTAEEREIHELWRVLKDEMCLPLMISLCQLNNLSLPPCLMALPGDVKAKVLEFVPGVDLARVQCTCKELRDLAADDNLWKKKCEMEFNTQDTCGCMMCKCIYSDQRKDIVLADKYTCGNYMQKPVTQPGRWLIILVYHSLLCQYITIGLSLLWYHLVDLVQDAPAAGIHFDCIIPLPINPYQLPPSAGACCSTTQASASAKDGGNMYSPPCSAAASSQGHCFAVGANQLASLDLAMDFDEPILFPVHNASLQEGIQFYNPTGDTQLSRNMSIDKCLKGSKRKGSGEGSSSLHSQEETGEMPQRELSMEHAGEKAGDADASREEYVHVRAKRGQATNSHSLAERFRREKINERMKLLQDLVPGCNKITGKAMMLDEIINYVQSLQRQVEFLSMKLSTISPELNSDLDLQDILCSQDARSAFLGCSPQLSNAHPNLYRAAQQCLSPPGLYGSVCVPNPADVHLARAGHLASFPQQRGLIWNEELRNIAPAGFASDAAGTSSLENSDSMKVE

>Os03G15440

MGAHGDHRHHHHHQEAGVLVDEEEEEVIEQACGGPTSGVVEQEVGGDGGGVCQDAAGMVFEATSSVGSVSATMGPPPIMCWPPPAQPVHGAIHHHHNLGGGGGQQSPFFPLLPPLPPQPPPPPPFFADFYARRALQYAYDHSGGASSSSDPLGLGGLYMGHHGSHVAGMMMPPPFAPSPFGDLGRMTAQEIMDAKALAASKSHSEAERRRRERINAHLARLRSLLPNTTKTDKASLLAEVIQHVKELKRQTSEITEEACPLPTESDELTVDASSDEDGRLVVRASLCCDDRTDLLPDLIRALKALRLRALKAEITTLGGRVKNVLVVTGDDSAAAAACAGTDGDGEQQEEAMQAPMSPQHTVASIQDALRAVMERTASATEESGGSGAGGGLKRQRTTSLSAILENRSI

>Os03G42100

MESGGVIAEAGWSSLDMSSQAEESEMMAQLLGTCFPSNGEDDHHQELPWSVDTPSAYYLHCNGGSSSAYSSTTSSNSASGSFTLIAPRSEYEGYYVSDSNEAALGISIQEQGAAQFMDAILNRNGDPGFDDLADSSVNLLDSIGASNKRKIQEQGRLDDQTKSRKSAKKAGSKRGKKAAQCEGEDGSIAVTNRQSLSCCTSENDSIGSQESPVAAKSNGKAQSGHRSATDPQSLYARKRRERINERLKILQNLVPNGTKVDISTMLEEAMHYVKFLQLQIKLLSSDEMWMYAPIAYNGMNIGIDLNLSQH

>Os03G43810

MNQFVPDWSNMGDASRTLGEDDNLIELLWCNGHVVMQSQNHHRKLPPRPPEKAAAAAVQEDEAGLWFPFALADSLEKDIF

SDLFYEAPVAATAEAAPAGPGAGADGEGKTCKGDAAMAEEERGGPGAASEAPRELMPPPKSTNASCSRQQTMSLADGGDN

AGDLSELVRARRSSGGAARRKAEAGGGGGGASSSMLSAIGSSICGSNQVQVQQRTASEPGRRGAPPSAVGSANAIPCGGR

DHGHGHEATTVASSSGRSNCCFGTTTTTEPTSTSNRSSKRKRLDTTEDSESPSEDAESESAALARKPPAKMTTARRSRAA

EVHNLSERRRRDRINEKMRALQELIPHCNKTDKASMLDEAIEYLKSLQLQLQMMWMGSGMAPPVMFPGVHQYLPRMGVGM

GAAAAAMPRMPFMAAPQPVVPTPPVNHLDLGVNHLQPPPTQGVGYYPLGAKAVQQQQNPPLHVPNGSIMPPPENAPNTGS

GMGSFYFYFYFSSD

>Os03G46790

MDDSSLFLHWAVSTLQHQHPAAVAVVADDDATFFSFQELCDTEEVVVVPVQEEVITEAHGGGASRIGLAVAVDEHGGWSR

SPNPGARPPSGGCGSNNLPLMSWDFSAASVAVQLEHVVAERKRREKINQRFMELSAVIPKLKKMDKATILSDAASYIREL

QEKLKALEEQAAARVTEAAMATPSPARAMNHLPVPPEIEVRCSPTNNVVMVRIHCENGEGVIVRILAEVEEIHLRIINAN

VMPFLDQGATMIITIAAKASSSLLY

>Os03G46860

MEDSSLFMEWAMETLQHLHPLPATPPPAGGGYAGDNATFPSLQALRESSVSQNGMAPPEPTAHEGHRASNSWSSGDTDSV

SGGGGGAVMEHDGWSTSPNSVRCAAGGGGGGGGGGLWPVSWNFSSAMTQPCNDQATPSNPPTTTRARYGGGGVRYLPAAV

SPSPSAQTRRASSKGNGGGGSGSSSAAPYAQEHIIAERKRREKINQRFIELSTVIPGLKKMDKATILSDAVRYVKEMQEK

LSELEQHQNGGVESAILLKKPCIATSSSDGGCPAASSAVAGSSSSGTARSSLPEIEAKISHGNVMVRIHGENNGKGSLVR

LLAAVEGLHLGITHTNVMPFSACTAIITIMAKVEDGVSVTAEDIVGKLNTVLQQNSRNSARETKS

>Os03G51580

MATQWFSNMVMDEPSFFHQWQSDGLLEQYTEQQIAVAFGQAGEADAAAAAAAMMVQQQQYAAAAAAEHRPRKAAKVNTSW

DSCITEQGSPADSSSPTILSFGGHADAAAAAAFASAGQAQSAPYYGGASAAALKPKQELDAAAAPFSQARPVKRSYDAMV

AADVAKAPAAAASRPASQNQEHILAERKRREKLSQRFIALSKIVPGLKKMDKASVLGDAIKYVKQLQDQVKGLEEEARRR

PVEAAVLVKKSQLSADDDDGSSCDENFDGGEATAGLPEIEARVSERTVLVKIHCENRKGALITALSEVETIGLTIMNTNV

LPFTSSSLDITIMATAGENFSLSVKDIVKKLNQAFKLSL

>Os03G56950

MDGNARSAANQTKQIVTDNELVELLWHNGGVVAQPQAAQARVVSSSGRGQSASVLTGDDTETAAWFPDTLDDALEKDLYT

QLWRSVTGDAFPAAAAAGPSSHHAPPPDLPPPAARPPMRSGIGSSWTGDICSAFCGSNHIPETAAQRCRDAGAALPPERP

RRSSTHDGAGTSSSGGSGSNFGASGLPSESASAHKRKGREDSDSRSEDAECEATEETKSSSRRYGSKRRTRAAEVHNLSE

RRRRDRINEKMRALQELIPHCNKTDKASILDEAIEYLKSLQMQVQIMWMTTGMAPMMFPGAHQFMPPMAVGMNSACMPAA

QGLSHMSRLPYMNHSMPNHIPLNSSPAMNPMNVANQMQNIQLREASNPFLHPDGWQTVPPQVSGPYASGPQVAQQNQIPK

ASASTVLPNSGAEQPPTSDGI

>Os03G58330

MAGQQPQQQGPPEDDFFDQFFSLTSSFPGAAPGGRAAGDQPFSLALSLDAAAAAEASGSGKRLGVGDDAEGGGSKADRET

VQLTGLFPPVFGGGGVQPPNLRPTPPTQVFHPQQSKQGGAAVGPQPPAPRPKVRARRGQATDPHSIAERLRRERIAERMR

ALQELVPNTNKTDRAAMLDEILDYVKFLRLQVKVLSMSRLGGAGAVAQLVADIPLSVKGEASDSGGNQQIWEKWSTDGTE

RQVAKLMEEDIGAAMQFLQSKALCMMPISLAMAIYDTQQTQDGQPVKHEPNTPS

>Os03G59670

MFPVEVAAAAAAAGRMQGEAVVPMMLPPFFMDSGIWPAAAGVVDVAASAEEEAAAAAAAAQDRALAASRNHREAEKRRRE

RIKSHLDRLRAVLACDPKIDKASLLAKAVERVRDLKQRMAGIGEAAPAHLFPTEHDEIVVLASGGGGVGGAGGAAAVFEA

SVCCDDRCDLLPELIETLRALRLRTLRAEMATLGGRVRNVLVLARDAGGAGEGGDGDDDRAGYSAVSNDGGDFLKEALRA

LVERPGAAAGDRPKRRRVVSDMNMQAAA

>Os04G47040

MASAPPVQEEPLQPGTNHFRSLLAAAVRSISWSYAIFWSISTSCPGVLTWNDGFYNGVVKTRKISNSADLTAGQLVVQRS

EQLRELYYSLLSGECDHRARRPIAALSPEDLADTEWYYVVCMTYSFQPGQGLPGKSYASNASVWLRNAQSADSKTFLRSL

LAKSASIQTIICIPFTSGVLELGTTDPVLEDPKLVNRIVAYFQELQFPICLEVLMSTSPSPNETEDADIVSEGLITHNAI

EEGQMVVSDECVSNANRDPITMEIDELYSIYEDLDLDMDLDLDTVRFLEDNGWPVNPSSFQLVPASSTEAVAAAAAANDV

DGVANSQVSCFMAWKSAKSNEMAVPVVTGIESQKLLKKVVDCGARMSTGRGSRAALTQESGIKNHVISERRRREKLNEMF

LILKSIVPSIHKVDKASILEETIAYLKVLEKRVKELESSSEPSHQRATETGQQRRCEITGKELVSEIGVSGGGDAGREHH

HVNVTVTDKVVLLEVQCRWKELVMTRVFDAIKSLCLDVLSVQASAPDGLLGLKIQAKFACSGSVAPGMISEALQKAIGG

>Os04G47059

MQAIDLFGCAMLSLQIAKPFLRALLAKSASIQTIVCIPFMSGVLELGTTDPVSEDPNLVNRIVAYLKELQFPICLEVPSS

TPSPDETEDADTVFDGLIEEDQMVILQGEDELGDVVVAECETNGANPETITMETDEFYSLCEELDLDLGSYQLVPTSARE

TVAAAAAAANDVDGVAYSHASCFVSWKRANPAEKVVAVPMTAGIESQKLLKKAVGGGTAWMSNIDDRGSVAITTTPGSNI

KSHVMSERRRREKLNEMFLILKSLLPSVRKVDKASILAETITYLKVLEKRVKELESSSREPSRWRPTEIGQGKAP

>Os04G47080

MEETPLPSGKNFRSQLAAAARSINWTYAIFWSISTSRPGVLTWKDGFYNGEIKTRKITNSMNLMADELVLQRSEQLRELY

DSLLSGECGHRARRPVAALLPEDLGDTEWYYVVCMTYAFGPRQGLPGKSFASNEFVWLTNAQSADRKLFHRALIAKSASI

KTIVCVPFIMHGVLELGTTDPISEDPALVDRIAASFWDTPPRAAFSSEAGDADIVVFEDLDHGNAAVEATTTTVPGEPHA

VAGGEVAECEPNSDNDLEQITMDDIGELYSLCEELDVVRPLDDDSSSWAVADPWSSFQLVPTSSPAPDQAPAAEATDVDD

VVVAALDSSSIDGSCRPSPSSFVAWKRTADSDEVQAVPLISGEPPQKLLKKAVAGAGAWMNNGDSSAAAMTTQGSSIKNH

VMSERRRREKLNEMFLILKSVVPSIHRVDKASILAETIAYLKELEKRVEELESSSQPSPCPLETRSRRKCREITGKKVSA

GAKRKAPAPEVASDDDTDGERRHCVSNVNVTIMDNKEVLLELQCQWKELLMTRVFDAIKGVSLDVLSVQASTSDGLLGLK

IQAKFASSAAVEPG

>Os04G52770

MEARRPTPTRRSRSAEFHNFSERRRRDRINEKLKALQELLPNCTKTDKVSMLDEAIDYLKSLQLQLQMLVMGKGMAPVVP

PELQQYMHYITADPSQIPPIRPSEPRPFQITHATQQRQSNVESDFLSQMQNLHPSEPPQNFLRPPKLQLYTPEQQRRGLA

SSSGHNSGWITERNSSYNFLE

>Os05G04740

MLRGNDTGSDLAELLWDNGAPAPLRPPPPPPFQPFTCSAAATTSPPAHDYLFIKNLMRGGGAANHHHHDDDDDDDDDVPW

LHYHPVVDDDDDADADTAPLPPDYCAALLSGLSDHLPPPAAAASRVDPDPCSSSHGAVVPSTSAAAAKQARTSGGGGGGV

MNFTFFSRPLQQRPSGGETASASASAAATSTVPVESTVVQAATNRLRSTPLFSDQRMAWLHPPKPSPRAAAPPPPPPLAP

TTRHRLDTAAATATVAQRLPPSEARAPDAPPPAATATATTSSVCSGNGDRRQLNWRDSHNNQSAEWSASQDELDLDDELA

GVHRRSAARSSKRSRTAEVHNLSERRRRDRINEKMRALQELIPNCNKIDKASMLEEAIEYLKTLQLQVQMMSMGTGMFVP

PMMLPAAAAAMQHHHMQMQQMAGPMAAAAHFPHLGAAAAMGLAGFGMPAAAQFPCPMFPAAPPMSMFAPPPPPPPFPHAA

ATAVEQTPSPPGAADAGNAPAVKQA

>Os05G07120

MAACQQQIWQEGKQQQHLHHGGYDDLSSVYRGTVVLPRRQGGLAPEPPPPRPSSSSGRSAAAQATAMTIHSEAERRRRER

INAHLATLRRILPDAKQMDKATLLASVVNQVKHLKTRATEATTPSTAATIPPEANEVTVQCYAGGEHTAAARTYVRATVS

CDDRPGLLADIAATFRRLRLRPLSADMSCLGGRTRHAFVLCREEEEEEDAAAEARPLKEAVRQALAKVALPETVYGGGGR

SKRQRLMMESRYSTAVVHTHVDPQYCWYNSR

>Os06G06900

MDEQRGRGGFDELVLLHQQQEQRRRREQQQEEEEEEEVRRQMFGAVVGGLAAFPAAAAALGQQQVDCGGELGGFCDSEAG

GSSEPEAAAGARPRGGSGSKRSRAAEVHNLSEKRRRSKINEKMKALQSLIPNSNKTDKASMLDEAIEYLKQLQLQVQMLS

MRNGVYLNPSYLSGALEPAQASQMFAALGGNNVTVVHPGTVMPPVNQSSGAHHLFDPLNSPPQNQPQSLILPSVPSTAIP

EPPFHLESSQSHLRQFQLPGSSEMVFHGEIMPKHHLSSHQESLPGNEMNSIRKESSMLNTNNFDGVSLSKEQS

>Os06G09370

MDYSAGSYMWPGNSGSENYNFVDGSSESYAEEGSLPPSGYFMGAGSDRSLKITENERNPTMLANGCLPYNTQAHPLSGQI

LPKGELPNNLLDLQQLQNSSNLRSNSIPPGVLQCNSTSGTFDAKLDTPGLAELPHALSSSIDSNGSDISAFLADVHAVSS

APTLCSAFQNVSSFMEPVNLDAFGFQGAQNVAMLNKTSLPNGNPSLFDNAAIASLHDSKEFLNGGSIPSFGTVLQALGAG

GLKAAQQEQNIRNIPLPTFTSGSHLAVTDAQGPPLPSKIPPLIHDHNSEYPINHSSDVEPQANSAPGNSANAKPRTRARR

GQATDPHSIAERLRREKISERMKNLQVLVPNSNKADKASMLDEIIDYVKFLQLQVKVLSMSRLGAPGAVLPLLRESQTEC

HSNPSLSASTISQGPPDMPDSEDSSAFEQEVVKLMETSIISAMQYLQNKGLCLMPIALASAISNQKGMAAAAAIPPEK

>Os07G05010

MDAGATARSSSSSAMMMNQKKPLLSDGELVELLWQDGGVVAHAQTRHRSSDVLARSGVTGEEETASAWFADGGGGGGGDD

DALGLGMGRDIYSQLWHSFANVDGHAAGALALATPTPTPRAAARSDDVSSRLDEADLSICGSNAVVAPALPADDDDDIDA

AAPREEEEEEEEGPGAARAAGASSSGGSGSGSGSYPLFKRGREELVDSLSEVADETRPSKRPAAKRRTRAAEVHNLSERR

RRDRINEKLRALQELVPHCNKTDKASILDEAIEYLKSLQMQVQIMWMTTGIVPMMFPGTHQLMPPMGMGLNTACMPGAQA

QGLNQMQRTTYYMNNSLPNQMPQIPSPAMNAPSVPDDMQNDNRIRGPRNPFLHCNDTLTATAQVPGLFTYGSQIAEQNEI

QELLSGAVIPSSSDGTIK

>Os07G39940

MAQFLGAHGDHCFTYEQMDESMEAMAAMFLPGLDTDSNSSSGCLNYDVPPQCWPQHGHSSSVTSFPDPAHSYGSFEFPVM

DPFPIADLDAHCAIPYLTEDLISPPHGNHPSARVEEATKVVTPVATKRKSSAAMTASKKSKKAGKKDPIGSDEGGNTYID

TQSSSSCTSEEGNLEGNAKPSSKKMGTRANRGAATDPQSLYARKRRERINERLRILQNLVPNGTKVDISTMLEEAVQYVK

FLQLQIKLLSSDDTWMYAPIAYNGVNISNIDLNISSLQK

>Os08G33590

MWEAVGGGDGTALLPWPGSAAAAPLYMPPAAAAAAPFAAGEQLPVEQPFYFDGGGGVAGHNHHPHHHQYGMEAPPPMTMM

QMGGGGSSSSRMVVSGLLGTLQAELGRMTAKEIMDAKALAASRSHSEAERRRRQRINGHLARLRSLLPNTTKTDKASLLA

EVIEHVKELKRQTSAMMEDGAAGGEAAAAPVVLLPTEDDELEVDAAADEGGRLVARASLCCEDRADLIPGIARALAALRL

RARRAEIATLGGRVRSVLLIAAVEEEDPDEAGNDDDGEHGYGVAASHRRHELVASIHEALRGVMNRKAASSDTSSSGAGG

GGGSIKRQRMISAHDQQGSFNSSGW

>Os08G37290

MVTDGECSAAAVRKGGSPAVRSHSEAERKRRQRINAHLATLRTLVPSASRMDKAALLGEVVRHVRELRCRADDATEGADV

VVPGEGDEVGVEDEDDDEGERDEGCYVVGGGDRRWRRRVRAWVCCADRPGLMSDLGRAVRSVSARPVRAEVATVGGRTRS

VLELDVVVASDAADNDRAVALSALRAALRTVLLNREELLAAAAAAATDGYKRPRFSPRCSSLT

>Os08G42080

MMACGSPSTEVVDEFEKLVIRMNPPRVTVDNTSDMTATLVKVDSANKYGTLLEVVQVLTELKLTIKRAYISSDGEWFMDV

FHVVDQDGNKLYDGQVIDRIELSLGAGSLSFRAPPERSVEVEAEAAAAQTAIELIGKDRPGLLSEVFAVLTDLKCNIVSS

EVWTHDARMAALVHVTDADTLGAIDDQDRLDTVKRLLRHLLRGGGAGARDRKATARAAIPAPRRDGAAAHAPRRLHQMMH

DDRAAAAPQPSSSSGDGGGRGRPVVEVVDCAERGYTLVNVRCRDRPKLLFDTVCTLTDMQYVVFHGTVIAEGSEAYQEYY

IRHLDDSPVTSGDERDRLGRCLEAAIQRRNTEGLRLELYCEDRVGLLSDVTRIFREHGLSVTHAEVATRGARAANVFYVV

AASGEPVEAHAVEAVRAEIGEQVLFVREDAGGGEPRSPPGRDRRSLGNMIRSRSEKFLYNLGLIRSCS

>Os08G43070

MDELVLSPSSFSATACFPTLDFEFCEVPDQWLLGLGHDELDKDAAASALAAAAASQSASNDDVPRNPPATTTTTKRRGRK

PGPRSGGGGAPPIGHVEAERQRREKLNRRFCELRAAVPTVSRMDKASLLADAVDYIAELRRRVERLEAEARRAPLAPSAA

AAAAWAAGLGAGAIGRDDLVVRMIGRDAAILRLTTAAAAARHAPARMMCAVRALNLAVQHASVARVGGATVQDVMVDDVP

AALQDEARLRAALLHTLQLADTT

>Os09G24490

MWEGGSHDAAAQLLPWFVGEPAAAAVGGYGGCVDVVGQGGVFGFGFEAAAAPVVTRQQRGGAAAAEGSSRGGGGKPAVVS

GLLGSLQAELGRVTAREIMDAKALAASRSHSEAERRRRQRINGHLARLRSLLPNTTKTDKASLLAEVIEHVKELKRQTTA

IAAAAAAGDYHGNDEDDDDAVVGRRSAAAQQLLPTEADELAVDAAVDAEGKLVVRASLCCEDRPDLIPDIARALAALRLR

ARRAEITTLGGRVRSVLLITADEQQQQHCDDVDDDEDGHRLLLRHGIDGAGAAADDDDECAASHRRHECIATVQEALRGV

MDRRAAASSGDTSSSGGAVVAGGGGGSIKRQRMNYGVHEQCSV

>Os09G28900

MVAARSGDDAELRLDVECLAAAPRWTRARRSHSEAERKRRERINAHLDTLRGLVPSASRMDKAALLGEVVRYVRKLRSEA

AGSAAVVPGEGDEVVVEEEEVEVEGCSCDAGERQAARRVKASVCCADRPGLMSELGDAERSVSARAVRAEIATVGGRTRS

DLELDVARTAAAGGGSNGASQLPALQAALRAVIMSQEELLAVESYKQRRFSADFA

>Os09G34330

MDELLSPCSSFSPPSPSSMFSTGAAAAAAHAVLEFTSCEVPDEWLMGDVVMAKNEEDVGGGELWPVFAGGSLSPDSELSE

LPRSFEAAAAQRPAKRRGRKPGPRPDGPTVSHVEAERQRREKLNRRFCDLRAAVPTVSRMDKASLLADAAAYIAELRARV

ARLESDARQAAAARFEPSSCGGGGNASYHGGGGGGGAAPGLDEAVEVRKMGRDAAAVRVTTTGARHAPARLMGALRSLEL

PVQHACVMRVHGATTVQEVLVDVPAALQDGDALRAALLQRLQDS

>Os10G01530

MQQLAESLANELFNQPQEQQEEQHGYHNPSLRVLPFVGDINKPEGHTPAAAAIRDSFFSLTNGSSSSLNFSALEQQQDSG

PMTKFCSPLSEMKRGGRRATSSMQEHVIAERKRREKMHQQFTTLASIVPEITKTDKVSVLGSTIEYVHHLRERVKILQDI

QSMGSTQPPISDARSRAGSGDDEDDDGNNNEVEIKVEANLQGTTVLLRVVCPEKKGVLIKLLTELEKLGLSTMNTNVVPF

ADSSLNITITAQIDNASCTTVELVKNLKSTLRNF

>Os10G40740

MNQCVPSWDLDDPVGGGGIGGGGGGGHRVVSGGGGGFMPVAVPTSDQYNEVAELTWEKGNISSHGLLLNRPAPPKFPPHQ

QLQAAMGGGGGGGVVGDRETLEAVVGEAAARSSSSSHLAARARPVPAPWLGSVGVVAAADALVPCDADAAEGRSKRPREV

VGEDGRRACASQGSAAPGRRGESTLLTLDACCGTAADDVCGFTTTTNNSTSLEDRTEDKGSPETENTSIAGGASDSRCFS

RRSQSQRGGMCDEDEHVVIRGEGAMRSSISTKRSRAAAIHNESERKRRDRINQKMKTLQKLVPNSSKTDKASMLDEVIDY

LKQLQAQVQVMSRMGSMMMPMGMAMPQLQMSVMAQMAQMAQIGLSMMNMGQAGGYAPMHMHTPPFLPVSWDAAASSSSAA

AADRPPQPTGAATSDAFSAFLASQAAQQNAQQPNGMEAYNRMMAMYQKLNHQQQQQQDQPSNSRQ

>Os10G42430

MWVLLSPLLTTKNPFHPIPIPTFPLLLFSSSLVGVLFQIKSNLEEEEIEIKSMNLWTDDNASMMEAFMASADLPAFPWGA

ASTPPPPPPPPHHHHQQQQQQVLPPPAAAPAAAAFNQDTLQQRLQSIIEGSRETWTYAIFWQSSIDVSTGASLLGWGDGY

YKGCDDDKRKQRSSTPAAAAEQEHRKRVLRELNSLIAGAGAAPDEAVEEEVTDTEWFFLVSMTQSFPNGLGLPGQALFAA

QPTWIATGLSSAPCDRARQAYTFGLRTMVCLPLATGVLELGSTDVIFQTGDSIPRIRALFNLSAAAASSWPPHPDAASAD

PSVLWLADAPPMDMKDSISAADISVSKPPPPPPHQIQHFENGSTSTLTENPSPSVHAPTPSQPAAPPQRQQQQQQSSQAQ

QGPFRRELNFSDFASNGGAAAPPFFKPETGEILNFGNDSSSGRRNPSPAPPAATASLTTAPGSLFSQHTPTLTAAANDAK

SNNQKRSMEATSRASNTNNHPAATANEGMLSFSSAPTTRPSTGTGAPAKSESDHSDLEASVREVESSRVVAPPPEAEKRP

RKRGRKPANGREEPLNHVEAERQRREKLNQRFYALRAVVPNVSKMDKASLLGDAISYINELRGKLTALETDKETLQSQME

SLKKERDARPPAPSGGGGDGGARCHAVEIEAKILGLEAMIRVQCHKRNHPAARLMTALRELDLDVYHASVSVVKDLMIQQ

VAVKMASRVYSQDQLNAALYTRIAEPGTAAR

>Os11G15210

MYDDNGAADLPTSQSASIKTIVCVPFIMHGVLELGTTDPVSEDPALVDRITASLWDTPPRAAFSSEAGVADIVVFEDLDH

GNTAVEATTTMVPGEPEPHAVAGGEVAECESNAHNDLEQITMDDIGELYSLCEELDVLDDDSSSWVADPWSSFQLVPTAE

ATDVDDAVVAALGAIDGSCRPSPSSFVAWKRTPDSDEVQAVPLISGEPPQKLLKKAVAGAGAWMNNADGSAATMTTDQGS

SIKNHVMSERRRREKLKEMFLILKSVVPSIHKVDKASILAETIAYLKELEKRVEELESSSQPSPRPMETTRRRCCKSTGK

KVSAGARAKRKAPAPEDTDGERRHCVSNVNVTIMDNKELLLELQCQWKELLMTRVFDAIKGVSLDVLSVQASTSDGLLGL

KIQAKVVVSAAKSSQQICSIVYLSIYQSLYLRLFGVLILAILLLHACSLPHLLPSNLG

>Os11G32100

MLPRFHGAMWMQDDGGGDQEHGQAAPPGQEQHHHDQHLMALAAAAAGGAGFGAAQAPAPLLDEDWYFDAAGGGGGGAHGS

MMLGLSSVHGGIGAGTSGGGHGQQFSLLNMGAAAAPFDVSGFDLGIACGGVGGGGDVVSFLGGGNASNTALLPVGNAGFL

GTFGGFGTAASQMPEFGGLAGFDMFDAGAVNTGGSSSSSSAAAAAASASAHVSNTAPFSGRGKAAVLRPLDIVPPVGAQP

TLFQKRALRRNAGEDDDDKKRKAAAGAGAGALSADGADMVLDDGDDDGLSIDASGGLNYDSEDARGGEDSGAKKESNANS

TVTGDGKGKKKGMPAKNLMAERRRRKKLNDRLYMLRSVVPKISKMDRASILGDAIEYLKELLQKINDLQNELESSPATSS

LPPTPTSFHPLTPTLPTLPSRIKEEICPSALPSPTGQQPRVEVRLREGRAVNIHMFCARRPGLLLSAMRAVEGLGLDVQQ

AVISCFNGFTLDIFKAEQCKDGPGLLPEEIKAVLMQSAGFHTMI

>Os11G34470

MGWWRGGDLEVISYRRGPKFPKFLRRSSYRRRRRTSWQWTQWCRVRASLSCEDRPDIARTFAALQLHARRAEITTLFGHA

WSVLLIIADEQQRNVRRRPGLVHAIFFAGCMIGSSIFILDGRCIETGASRFYYDSSRKRLRPARQRRREEEKERMKKMIN

ASSNHAEETSTKRPQLPPPELPTRAMHSCLRPPENAYSKVIEGSL

>Os12G41650

MNQFVPDWNTTSMGDGFAPLGEDDGLVELLWCNGHVVMQSQAPRKPPRPEKTTAAAAAAMAEDESASWFQYPVDDVLEKD

LFTELFGEMTAAGGGGGDVRRAACKEERGAVAAFQSRMMPPPWPARGKAEFGDVDDVCGVSEVVMAKMDGAAAAETVGES

SMLTIGSSICGSNHVQTPPVGNGKAGAGTAGAARRAHDTATVASSSMRSRSCTAKAEPRDVAAAGVGGKRKQRGGAAMES

GSPSEDVEFESAAATCSPAQKTTTAKRRRAAEVHNLSERRRRDRINEKMKALQELIPHCNKTDKASMLDEAIEYLKSLQL

QLQMMWMGGGMAPPAVMFPAAGVHQYMQRMGAVGMGPPHMASLPRMPPFMAPPPAAVQSSPVVSMADPYARCLAVDHLQP

PPPMHYLQGMSFYQLAAAKNLQQQQNTAEAPPPPPAGGNRAAADS

>Os12G43620

MDDSSFFMQWAMDTLHQLPSDSTAAAAYATDVAGDSGAFPSLQALRNASAAGGGGGFRDLTVQVDQVHRANSWSSSDSPG

GGAATAAAGWSPHVTGGGGRGHRPMSWNFSAASAQPTTEDSGGGGGGGVVPAPLQAMETTATARAAVKKGGGGGSSSSAA

APGYVQDHIIAERRRREKINQRFIELSTVIPGLKKMDKATILGDAVKYVKELQEKVKTLEEEDGGGRPAAMVVRKSSCSG

RQSAAGDGDGEGRVPEIEVRVWERSVLVRVQCGNSRGLLVRLLSEVEELRLGITHTSVMPFPASTVIITITAKASSLSNH

PALLCLYICIKLPCVISS

>Pe2005324

MDQITDGSHSTQHLL

YSLQQPQEQQSEVTLLPDNPNNMNAMDMCMLDRQRKGPNAQPQLNWPQQQQFLRTNYNSE

QMHGPFTSLLTSSEMFSDGIITSEWQQLAEQNGHLVTANPEYTMNIGRPMSSQGSALRQT

LSRSMSSPNGPLNLHTDWSNLSNNTESMCTVESEANGRSAIKAPACLGPFPSDPAFAQRA

AKFSSLNGKNWNGRLSTSQRASDGEAETGSLEFLSDVARNEPQVACSDDSESKKAFCRTT

SCPPTVEHAAAETEKSSVVVEESAVSEKSSARDAKTSNNNAVKKRKSGRTSTQNHRAADS

EESKEKRSKTVHPDTEEKELVMAKTEQSNSDNSVDSSPKPARANPKPLENQKQDYIHVRA

RRGQATDSHSLAERVRREKISARMKYLQDLVPGCNKVTGRAVMLDEIINYVQSLQRQVEF

LSMKLAAVNPSLEFNIDNFFAREMMMPVPPPGNFSSLEGMSPNLAPYIHFQQMLQQQASL

QAGACSGLDVPTLDTTPSLHRPGSAEMFGDAANFQGHAPSTWDSELQNVFSMGLVQDRQP

PFSSEGLPGHLEPNHMKMEM*

>Pe2006916

MLNSRAQAITTGLNIPSMSVSHSDANGIQMSFSQYSMDSEKMPQMQQQ

TNNVDFMDLDYKPSVCNVSQADGVMALTERGSSNNFQGPTGLMRYHSAPSSFFSSLGEEE

NNNIISEYFSGSSANALQSNTKPLQQNQSSILSFHLRENEPAKHLGEHNEYLKRNARQLP

SLKRSAGNVAREDLGKHRPLDAILENVPDVSQDSFGTSQMSLMSQVQVSEPSGQHIDSSY

QMNSVCCNTLDQSGGRMSNAYSSTSKNTLIRHSSSPAGLLSELVAEGRGTFESGMQNGTN

FTARKRAKELDIKMMQSLNNSDHQKVEAGIRGASALTNHPYSLPRSTSSELAMEEFLQDA

VPCKVRAKRGCATHPRSIAERVRRTRISERMRKLQELVPNSDKQTVNIADMLDEAVEYVK

SLQKQVQELTDNRDKCTCTHKPDCAYKT*

>Pe2006950

MNLYPAPSAEELLQEAAAIVLKKEFYLVSNFSEKYI*

>Pe2006951

MNLYPAPSAEELLQEAAAIVLKKEFYLVSNFSEKYI*

>Pe2007211

EKKMGSEAAAASRFWNEEDKATVNSLLGPSAFEHLMMSYLSSEGLVSG

INDCALQQKLQNLVESSSFNWTYAIFWQLSRSKNGDVVLGWGDGSFKGPREGQEADQARG

FDQRFAETDQQLKKKVLQKLQSFFGGGGEEDNNFVSGLDNVSDTEMFYLASMYYSFPRGI

GVPGQALASGKNIWLNEPSKLPTNMCSRAYLAKTGGIQTLVCLPMEHGVVEVGSVEMIRE

SKHAIDKIRSSFNENACDGNRGQPTVKGSLVAPFSPNPIRVNAVNAKAAPPLKPSHDWKI

FGQELSKSSESVVTKVEERDRHYHPVFRPPYSHAAPYVTNEQRISYTNANQNGLQSPNWS

HISNGEGGEIYNTRDLIKQSSRISPISVAGPSLSAVTARPPLMESEEHSDVEASERRPVV

VEERRPRKRGRKPANGREEPLNHVEAERQRREKLNQRFYALRAVVPNISKMDKASLLGDA

ISYIQELQNKVKDMETEKEKQQQPQLQQAKSNIQDGRIVDPISDIDVQMMSGEATVRVSC

PKESHPVGRVMLALQRLQLDVHHANISAANENILHTFVIKLGGAQVLTKDQLLEAISGWS

PRQKQ

>Pe2007372

ICLFTMELADERSILRYKKP

KLSKNVVSERRRRQKMNKLLYTLRALVPNISKMDKASILGDAIEYVDKLKKQVERAESDV

QSTNVSAQSADPRIEETDSFTVRLDAANRYGSDRKYHVQVEAHVVEESIVEVRIRCKQEH

GILFHLTRALESLPFRLKNVTFSTLNEYLLLLNAVLQADHCFPAMSDYGVRQSIEDSFGK

QGLILTSGNSSGPSWQELK

>Pe2008905

MESTDQPSMLRYKKPKL

SKNVVSERRRRQKMNKLLYTLRALVPNISKMDKASILGDAIEYVNELKQQVERAECDVQS

TNVSSQSAEAPVDETNSLGLRLDAANRYGSDRKYHVQVETRVLGESIAELRIRCKKECGI

LFHLTRALESLPFRLKNVSFTTLDDYLLLLNVVLQAEAECFPATGDCGVRQSIEDSFGKQ

GLILTSGNSNGHSWQELK

>Pe2009306

MDTLMASSAVDQRFSQETL

QQRLQTLVETASIVWTYAIFWQVSYESSGAIQLCWGDGYYKGSRNTEEDERLRMRSRLTV

SPADQELRKKVLRDLHSMISGSDEGNQQDNSSVSVDEEVTDAEWFYLISMMQSFLSGFGV

PGTAFSTGAPVWIVGAERLRVSTCERARQAHDLGIQTLVCVPIQGGVVEFGSTEDIVENW

LFLEQVNRSFKYNLNQTHDNLFQIQSLWPEETLSVKSSNTMQSAPCIEPVNNAEIQSLNS

ALARELPVTGKQKASVFAEQSSLVVKDDKSLLHPLTQQTEALEAPAIRIPETVNGTEPQT

RALGFKGSEKNVIKPSIKEDTIGLLSNPPGIAIGGLRSSIESELSDAEPSASIKDSTSAV

VERKPRKRGRKPANGREEPLNHVEAERQRREKLNQKFYELRAVVPNVSKMDKASLLGDAA

AYIKDLCSKQQDLESERVELQDQIESVKKELLMNSLKLAAKEATDLSSIDLKGFSQGKFP

GLNSEVRILGREAIIRIQCTKHNHPVARLMTALQELDLEVLHASISTVKDSLIIQTVIVK

MTRGLYTEEQLHALLCKKVADLN*

>Pe2009619

TNSQRARQHNSGVEVPGSNNMPASIVESSGGLCANSDESYPSSCRASTLSSKNQPPPLNM

VRVYSKPVPTCETNASDHDCITAKAVEDQNRGNNNSMSESATIAKERTVSENYEGVEVTL

TSTSGGSPNNSMTQKSGKEPCSTRGKRKFSGEEDSGCQSEDPDDNESAQAKKPQTGRRSG

SSNRSRAAAVHNLSERRRRDRINERMRALQKLIPNSSKTDKASMLDEAIEYLKHLQAQLQ

MMILRNGMNIPPMMMPLGMQQLQMSLLASLGPTGLGMGMTGVGLGMGMGMMDLNSVAAGR

VPMHQWPNLFLQ*

>Pe2010025

MLDFTDLKFDSCACPISSVLGVVDGSSSCNVFCSDGKRSSYGAASKNTISERRRRRRLNE

KLYSLRAIVPMITKMNKTSIIEDSIDYIQQLQNQVRVSEEDVVRLKENSTPTSEKKTGRE

FSHDDYKGESIKELLEFDVSEVKENVYSICLYCKCSPRVVLEVNRVIESMEMTVTYANFN

SFDGYILINIIVQRGENGKKLQVDALKSIIAKVIME*

>Pe2010345

TRSHGLKSEQIFSLPNDVYGLPRVPDFQGLKFPYSYSGGRINTNNSSSNSLGMSAMCERM

GLYMHLNGVGGAGGGGSRLVLDGVRGELVNAAKMMTPKELMEAKAVAASKSHSEAERRRR

ERINSHLATLRTLLPSTTKTDKASLLAEVIDHVKDLKRRASEIAKGNPVPTDVDELRVEE

ADGDGDNSKGRLVIKASLCCDDRPDLLADLNRTLHSLRLRAVKAEISTLGGRMKNVFIIT

STEGVPEKDQDSPSVNCVQEALRAVMERAASNELSSGNLASNKRQRILPLDATSSAI*

>Pe2010837

VYASSVGRGRKNGLPAKNLMAERRRRKKLNDRLFMLRSVVPKVSKMDRASILGDAVEYLK

ELLQRINDLHIELMAGSSNSKPLVPTMPDFPYRMNQESQASLLNPEVEPATVEVSTREGK

ALNIHMFCSKKPGLLLSTLRALDELGLDVKQAIISCLNGFALDVFRAEQSMGGDVTAEEI

KALLLHTADNEDGL*

>Pe2011133

MALSPRANATAGSPKEIVTLSVSQATVCPPSSLHP

GPVRSSDLSYSAALNSSFGFPFQDFRTQRIPPPNSHCMELPHPQLPFEGSKPPHDFLSLY

GDSSSNRVEHRSSQANSTHLKTQDFLQPLERGGKNSNIKGLEVSSTEASAATPASSVEHV

LPGGIGTYTISHISGYSQINGKSEIPVVHVNVAETKPEVTRAASSTNYGGGAFTLWEERI

RAVEGNRTPDGLAATDFSKEYLEKSGYWPAVRAKTDLSFPAKTAFSLSTVETSQQPLPSS

SKQRLQTGQGFVEMIKSIKAISEDDEDEEDEYVDRSRESRKECSSQKGDPPPKADGRNND

QKANTPRSKHSATEQRRRSKINDRFQMLRDLVPHSDQKRDKASFLLEVIEYIQVLQEKVR

KYETTEQSWNQERMKTMVWDLCKKSPGHGEVPMDGSRIVVGTLSDSKQNNEVGDPARSKV

TPNALHYENGTAGTDMHPTSQNLMQPNKSTPVIPGCAPSYKEKENIGAFNKSIPPTLSMQ

QCVYSPFGRSYIHSTHPKGQATSEIHSLQSIQGITPTCIPANAYNFDKEKDGDSAAPVQL

SHFQGNTVTVAKTTSGHESQRSELRSSQLSPIQQYSSQNGGILLKDEAADTKSASKSENQ

HSSAYFNSSPRSSQYVPQSEAELASRQPQPRSSSPDQHRASEPRLAASEQEELVIQSGVI

NVSSVYSQGLLDTLTQALQNSGLDLSQANISVQIDLGRRGGTSGATTNTKEHSHNHEESS

QNHQLQGQSRAADTSSECEQPQKRPKIERDV*

>Pe2011294

MQQDEXNARRADQDSKMLFPTWPTIAVPSHNVAQGHLEPFQTQ

VIPGRDQILAAMASEGGSSSSNSFSGFQDLINGGAPLQFSYDYRGGHMLQVPAGLANFSS

MDHKSVGDRLFQPVLMSRSALEMSKMTAQEIVEAKALAASKSHSEAERRRRERINNHLAT

LRSLLPNTTKTDKASLLAEVIDHVRELKRMVDDIGQASPVPSEADEVSLEYPDSSEEGRP

VIRASLCCEDRPDLLSDLMKTLRSLRLRAVKAEIATLGSRVKNVLVVTSGGSQDPNDVSE

TPSASSIQEALKAVMERQNSGDLSPATSSGGNKRQRQVG*

>Pe2011779

MMAQRLDI

SGGNDSSISELEQTNAHPDLTSKDIINLRKRKCLSNPKVKVADVHSIPPKTKETDETERN

GKHYKVGESTKDKDDLKDKLEESNSAETAESCPKQTVDNAKPSSVSVKQDYIHVRARRGQ

ATDSHSLAERVRREKISERMKLLQDLVPGCNKVTGKAVMLDEIINYVQALQCQVEFLSMK

LAAVNPQLDCNVEGGYLTRDVLQPHCSSISKMFAPDTTAAASQINQLQKTPLQHGLQCRA

DRQELAIRGMMDTQFTCMNGYADPTFQLQMSQGWDDEFQNAVDIGLDQNRSNPLKSHGFH

GVLPTGHMKVEL*

>Pe2011837

DNNVMIEAFMGNLDYSYSFPWNGIDANPSALPSPAHFASSAASVAIATPFNQDTLQQRLL

ALVEGATESWTYAIFWQLSSEASGSPVLGWGDGYYKGPRDMTEEERASKKAASVDATAAD

QELRKKVLRDLHALINPNATGDTDPAEFSGDDLTVDGEVTDAEWFYLVSMMQSFVNGCGV

PGQAFSSAMPVWIIGSERLQGYNCDRARQAQQFGIQTMVCIPTLNGVVELGSTDLIPQNW

DLIQKARDSFTFTLPDTALWEENHTQNDPDPALWLTEPPAEPKTETEKKQQQIPNAETEA

LHSFFPHELGFSDLGFLNGEGENASQKLVVEECGQKEDKSSQGFTGSQVSYQQNWQAQTT

CKTEVVDIPVFQPGKRTNTNGITLSFENPYGAQGLVSWGDEKMKRPVRNGNEDGGSGLCF

SSEVTAVSASTSATAAAAALPVNGAAGVRSSVESEHSDIEASFKEAECSQAIVERRPRKR

GRKPANGREEPLNHVEAERQRREKLNQRFYALRAVVPNVSKMDKASLLGDAISYINELRN

KVQDSDSHKKDLQAQLEALKKELVARESVASGFSGSNFGLLKNPSAADPSNLDVKGFGLK

NQCPNIELEVRILGREAMVRVQCPKQNHPVARLMVAFKELELEVHHASVSTVKELMIQTV

ILNMTGIVYTQEQLNAALLRKVADPGLR*

>Pe2012024

MDPDFGVERQAFEAVPDSKSLVSLHSSLSCSFNTMDTIFCDESRYSQKCTGEFGEIASQNA

FGGAPESIISNAGGAFFADPSFAAKYLQKSSFSPVMAKETSNADGSGSIGGPNVPILANS

VQFHTPSVKNSVNARRWRNLEASHLGSEVSKFGIGSVPSALVQFPSDPGFVEQAAKFSPF

NNIDRNSSENVVCGASNLPPIVGGVQQGGGENMSVFSEPTKCVDDGNGKKRRAKSLTSAE

NSKEPEEAKAKRCRLGESSEIDDDDNNDETSDSNSKKGKEKNSNVSQKDDNYIHVRARRG

QATDSHSLAERVRREKINQRMKFLQDLVPTCNKVTGKAVMLDEIINYVQSLQHQVEFLSM

KLATVNPKLDFNIDNFFAKEMSGSFSSKGMSPTYFHLDQLKQASLQTVPSPGSDIPSTMS

SVDSAEMFDGTNFQRQEGWDSELQNMYNMGLLQSRLQQHQHQQRPL*

>Pe2024173

PELLSDLIKTLGTLGLCTIKAEITTLGGRVKHELLIDSSNEX

>Pe2032305

TLRSLVPRITKMDKASIVGDAISYVQDLQKQVKDIQAEIEGLRSNLSGQNDST

>Pe2050038

HEEKSNTELDVVRRDPPELLSNVVTKETKAESPKDYIHVRARRGQATDRHSLAERVRRQK

ISDRMNILQNLVPGCNKVTNKALLLEEIINYVQSLQIQVEILSMKLASTSYLRAFDDQIT

GHPFKELLPAEATTGPMIASGIAPHSVDHTRTVAQHRAPGTFV*

>Pe2051696

DFTERENRSATTGLDGRQGPLLTNYARSQPGQQNTGLQVLQSSQGNSLQQQGNRSVISQD

QLAATAGAVLNGAPRTRVRARRGQATDPHSIAERERRERIAKNLKSLQELVPNANKTDKA

SMLDEIIDYVKFLQLQVKVLSMSRLGGAGAVAPLIADVPAEGSGSLASAALGQAGGSLSQ

DGLAFEQNVVKLMEKDMTAAMQYLQNKGLCLMPIALATAISSSTGKPLLGSGVTNSVDST

SDRQSSGNQPASLTVDSCAGALGMGSAGFGSDFLLKENTVEKRGRELVPTAESNGAEFGI

ISX

>Pe2052556

MAMFTGNYSLRKSLAKRGFHQQTFPSGYSDPTSLTHLPD

FQIGAFPFSSYGSHLQTTVSYLNTNGVRHCEQNQLDQNGGSSLVLDNARGELVNASKMTL

KQILEAKALAASKNHSEAERRRRQRINSHLATLRGLLPSGTKTDKASLLAEVIDHVKDLK

RRASEIAQVGPVPTDVDELKVEEDGAIEEGKILIKASLCCDDRPDLMPDLIQTFQTLGLI

TVRAEISTLAGRVKNVFLVTDGEKDFEESQTRLSINSIQEALRAVMERTAASGELSAGSS

ISNKRLLKIVLNVAVSNTYSSVFLGIS*

>Pe2052825

ASHLGLRTSGGYLGDERPAKKSKPNNLCVSPFHGQQRFSSEVSTSPSIYVPNSQQKSQHR

QIDKYFTNFFKLNGDETYENAALTTPSIQAEIVASGGHHQSCISDTICHSHPDAFNVSSG

QFFRTNHGPQVRISSSTIGNRHYNNLPNKTSGHSLDHIMAERKRREKLSQRFIALSAIVP

GLKKMDKASVLGDAIKYVKQLEDRIKALEEQAPKITVQSVVYVKKEELCTDDQEDSDKFS

SINSSSDNAVIEGVLTPEIEARLVDKNVLIHIHCEKKKHLLVEFLAELEKLQLTVLNASI

LSFSESSFDLTFNAKMEEKCDLTGKVIAKALQALFKKTTSIASPPN*

>Pe2052902

MPSSTQVGSLSPCLQFAGNDYNGLLVNGGVADSGLSGLLVRGGHDPRSSSLGSRSKAERG

QVTLFQKRATARHSSPTPLPDEESPNTRALRKSPKGDGIRVQSSNLEEEGEEPLGGEDNM

DDDSGGGGSGLLYETDDAKPEQVMSGQDGPAPGPGPPADKGKKKGLPAKNLMAERRRRKK

LNDRLYMLRSVVPKISKMDRASILGDAIEYLKELLQRINDLHTELEGTSERPLSATALPL

PLNLPVPSPFHYPLTPSTPPLIPCRIKEECPPISLPAHSEASDSQPPRVEVKTRDGRSLN

IHMFCARRPGLLLSTMRALDELGLDVQQAVISCFNGFALDVFRAEQSRGGGDIALEEIKA

VLLHTANSHQITM*

>Pe2054274

MAHKCLEPSFLVHNPVLWDDEDALN

RNYNYNYYTDLWGGLPLSSSVATTARTSAEDSRPEVVAVSEALESDYGRLNNNNNNNVSA

DTQLSCETAATSSSGGSMSSPSNGNKVIKSSKKKVLMGTKDIECQSQKAQEDSGENFKQC

STGTSSSKRSRAAEVHNLSEKRRRNRINEKMKALQNLIPNSNKTDKASMLDEAIEYLKKL

QLQVQMLSARSGIDISSMRWLAQMPHLQIQQMPKACMTTDQHAGVSISMPVGSGLMNTNQ

GSEKRPLPLHDLYISGALGNTAIPINLPSARIDDHQSCKMDRPQGHFTLPQLPTTTQEVH

SALSLQEK*

>Pe2054955

RTSNSISRRSFVARSGGIQTILCVPTDTGVVELGSVDSIKENKEVVQMVKSIFNDHQVQQ

LGHSRGSALSRSPARVPEIHPLSREKDVQESAVVPFKLYEYGAHQKVVAQDYRKVFGQDL

NSGRTRTLVDDKFISSKMENRLLQPVLRAPNGHQHEYYLNGDKIFYGSNKKGFQSITLNQ

VCSSEEGDINDSRHDYHQQKHGNGVLILNNDNNNLKWSKEENKIKERQRQQQQSQMEFSG

TSRSIVSLRHCVAESEHSDADAPCKDDKSPLVDERRPRKRGRKPANGREEPLNHVEAERQ

RREKLNQRFYALRAVVPKISKMDKASLLGDAIAYIQELQKKLKDMEKEKETQEKTSHAPA

VAPQGNIPAVVNNNSISDVDVQIVNGDAVIRVTCPRESHPVARVMLALQEMQLDVQQADV

STAKEMIFHTFVVKLNGAQELTKEKLVAAISG*

>Pe2055341

MLNNMLHF

QSQLTAGPKLEPLSTVNHPDDPLRSVLNISSSYQSPWMNPSCFSSPESVPFDEKALELSE

MDTLVHQMGDIESWGNLDQVHGCVKPLNAASMQACSIVSDPQLEILNVFLKKHNPAEDVH

VNLNPDSDFLPRVRSANVLQMDAVVQQDAFGSILSDEQEPMENKSKDHMLKVPAKRSSGT

RPAQLRPTKRGAAAAPQQDVSGISQREIHIQSERQRRKGMNHLFERMRSLLPNPTQKTDK

STTVSEIINYIQALHQNLEDLNKKRAKMLPSPRGGTVHVKTEPTSAESIQSECTGSEDSL

PVSSEEECTRQTPHVTLHFNGNDIYVTVSCLNKANLLAAIIFVVEDHNLQVISANFSATD

TVAFHCLHVKALETPDAVTREALKCALQKFVCSYTQICLK*

>Pe2055367

MEIIEEMIEIPR

LNDNLGLNIPWLHGGLFRKRLTSSPLHSEPAGKQGNTGPNLENYFTRLVQGFEPVESCVT

ALSELKPGGNIDTFNAVHSHSDTAEDGISDTTQEDQQNAISMAVPAADAGKGRKRKRLKV

SKNSEEVLSQRITHIEVERNRRKQMNAHLDVLRSLIPQSYIQRGDQASIIGGAIDLVKEL

EQILQSLQFQKIKRENENRTDPKNCLSILPPHQLSVPCNSKCVVDTSSSQLREAEDKSSI

ADIEVTLIETHGSLKILSKKKHGQLLKIITALEGFHMTILHLNITSMDEAVLFSFNVKVE

DACPLTSANDIARSLHQILRIVHAC*

>Pe2056043

MDDFLDQILSSSSSWMDMSGGQANGMSGDSMGPFQETLKHPTIPSDTLHRL

VESNTVAGGPTSLQLNTSVPVTRQGSTSQFLTAVGGPSRTAVSLAELASAGSSSSEASGF

QQALADSHPPAPVWSESYTVTSALPAAVGQGKIEGFTLKEETVQGDGHLLGKRSHSEDKM

LARENRSGDTEHDDLQGTLLTSYTGPQGGQTVLPRTPGSQSHQQNLSTQTIQSGEGTSLQ

QYGNHSAVSQLQSGAGGGNAANGAARPRVRARRGQATDPHSIAERLRRERIAERMKSLQE

LVPNSNKTDKASMLDEIIEYVKFLQLQVKVLSMSRLGGAGAVAPLIADVPSQGSGGVMST

ALGQASGPLALSQDGLAFEQEVARLMESNMTSAMQYLQNKGLCLMPIALATAISSSSGKP

ILATVPGTGVDSSEKQNSDMQSIVLPFSSSSTTMGLRASASGSDAPINENSMNKASIEKV

VAEKSNGISPGLSSDVPKVGSQNREELLNRAQ*

>Pe2056101

PIISISAQAASECKSFKFSKQAAGFESRDCEIIRRGRIPLRLDILASLVNFLVDMQPSTS

MLGTGPVQGVSSVTACLSSSPQVGLQEQMHSQQHQINNQQQHFSSQFDQTNQPHQNMDDF

LEQMLFMPQWSDVAGGKSPWEFNPNNAPQGNNASNNNNSPQSLQSNTQKLFTMSLMPPGI

GSQGLNSQENDGHAGEHMQYSYDQSPFLANRLRQHQLSGNSSINQTPNQVSQGINETNTS

PGRSMVLQLSTGNASASQLLASMGNSPRGTAVMTRSPSTGGSCNGSDGGMLPLPLSLGQA

GKSGDMNEPGSREEIEASFKSVNNARDSNLGGLFQPFAVSPRGVRPTGQNFHAQPGQVPL

QGYGGMPQPQHQNPPPTGVGAAPPVRPRVRARRGQATDPHSIAERLRRERIAERMKALQE

LVPNSNKTDKASMLDEIIDYVKFLQLQVKVLSMSRLGGAGAVAPLVADIPAEGANAPQGT

RTNGSQNSSPDGLALTERQVAKLMEEDMGTAMQYLQGKGLCLMPISLASAINNSGSRPQA

PSTPSLQGLLASNVNDRQVLEPSSNSALSALTSTSMTIQPSISSPGSGITEQGDNAHKAG

NGTRNPKDSREANSVTKSNGIGPALSKNAVKGEDEPQRRTGQ*

>Pe2056666

MLSRMNLNGGVWGLEDATDLHCNNDENLNLPTFK

AMLDAEEWFTSHSNTSNNDITTASINNANSHPDGPCFNIDMKDFGCYSSLVNPNPEPPNM

LLQHAMNSCSSSPASIFSLDPSQVQSFLAGGSNHHHHHHVSPFPDVAGSSPFQELACEGL

VSSNNASSNSSSNSYLYGGLPGGIFRQLSPSFAGSSPRTNTVTPTPNLSSPHSQMGGSST

PNLSPRMMPTSNLNLSDTGLFSSSNHTTTTNNNNLSNLLGLSSLMSPRIPAPLNFSSPKM

LRPLEICAPVGAQPTLFQKRAALRHTALSRCTTTPCIEVLDEESPNGNSNSNVKGKSKLM

LVGADNKEDEDVDESNDGSGVHYDSDDVGANNNSYKVDQIGEDGLASAGGTGNVAAVNNN

INGGGDKGKKKGLPAKNLMAERRRRKKLNDRLYMLRSVVPKISKMDRASILGDAIEYLKE

LLQKINDLHNELESTSQGPVLPGTSNFHPLTPTTPSLPCRVKEECPTSLPSPNAQPARVE

VRMREGHALNIHMFCARRPGLLLSTMRALDGLGLDVQQAVISCFNGFALDVFRAEQAKEG

EIAPEEIKAVLLHTASCHTAI*

>Pe2056881

MCLKAMAVMASANSSAIRKQLESVVQ

SIQWTYSIFWQLSNQQGVLEWSDGYYNGDIKTRKTVQPMELSNEELCLQRTLQLRELYES

LSAGESNQPARRPCAALSPEDLTDTEWYYLVCMSYTFAPGVGLPGRTLANGRLVWLCQAN

EADSKVFPRALLAKSASIQTVVCIPIGDGVLEFGTTELEREDPGLVQRTISFFMDYPKPI

CSEQSTSSPQCSDRDEKDQVGMMTLLSSDSIVCLGRNQIGASTVTDCGQYLPTSHEDLDL

PIQTFEQKDKISITEDPQQHGLNESMQVEICEDYKASGSPEDHCCNGDAGPHEFPLISAE

NDCLQNGHVNLNPAALEGWPYMEDNTSHGLQASGECVTQSIVDPSPQLCTYSQRDMNMNM

NVLLGLEQGSNALETMLESAAQTLDEDGHYSRTLSTILEQQQAGNLTESTGFISTKPGKD

WRSRHSRQGSGFIHWKSNGNCVVGIKAVASPQRILKKVLFNLARLHSKYKEDPNYSPKLG

EEEIGSKLVGRKIGQEDLSVSHVLAERRRREKLNEKFIVLRSLVPFVTKMDKASILGDAI

EYLKQLQRRVEELEASSKVMEAEMRKTQNRNLPKRSCSSTEDMRMARHGGNHVDSCLQSS

CLDGELGWTLTDTKQPPSKMPRLESKRKLNDLHKKGSCTLPAREDTEVSVSVIEDDAVLI

EIQCPCRHGVLLDIMQRLSSLHLDTCSVQSSTADKMFAAVLKAKVQEKFGGSKRPNIAEV

KEAVELVASKC*

>Pe2056926

MGSDGKEEFGFSMNYKASG

SPPSSNWQSAVANVAMNNVQESSLRVPTSSESLSSSFFSLNWEPLIEQSIPFQSSLSPMV

SPSPPATSLPSDSIAIRELIGRLGGNICSVSGPLGSTMDTVLNMGTAWNPLNSSPNLNLA

VDPSKGALSIQGAKHTPPHNLAQFSSDPGFAERAARFSCFGNRNYPELATPFNFPEGEPS

YRSAPVDNSKIPRVQSNQSLKAGLPINLPNMAETKESNASETPHEGSEADPRFTDRKISR

ISRSSTPVSTDDMKQRLATSGNESDEAEFSTGREESSCSDHIAGREPGSKTLNEVNGRKR

RVLSKAKAKDTPSAVASSGGRETKSLEADESPTKRYKGAEAGSNEKDDAKSKAEQSTILS

TGESSPKQTKDIVKTPEPPKDYIHVRARRGQATDSHSLAERVRREKISERMKFLQDLVPG

CSKVTGKAVMLDEIINYVQSLQRQVEFLSMKLATVNPRLDFNMDGLIAKDMLQSHGSSPR

MLFSTDPTAAFPQLHQPQQGPVQVGVTCGTEGHRMGHPVEGALRRTMNAQPPCIDGYGDS

IPQVANVWDEDLQSVVQMGFGQNRQSPFTSQGFHGSVPTSHMKIEL*

>Pe2056957

MALVDFTEVPRTRLRQQMQAAVQSIQWTYSVFWQFSHPEGLLVWMDGFYNGGIKTR

KTVQPMELSPEELCAQRSLQLRELFESLSAGETNPPTRRPCAALSPEDLTESEWFYLMCM

SFTFAPGVGLPGRALAKRHHVWLSQANEADSKLFSRAILAKSARIQTVACIPLADGVLEI

GSTELIREDIGLIHQVTSLLADHSKPVCSEQSTSNPHSENASRALPPDQLSMQPPDGIVF

EEQNVKATEYNSDNAHENAHIPMQVSGLQSEKTVTEFGQNGSEVMQLDMSPEDCSNDIGS

ELQVTGGNSCMQVKHSTNAWPDISHGLQSSGTQQYLDQGSDDESGHYSKTVSTILQQQRP

SQWTETTTLQLVRGNINPQDITQRCAFSCWTGNGGVSVQKSMNPQWVLKYILFSVPNLHS

RDRDESSPKLREGENGCRVRRSGQDDITVNHVLAERRRREKLNERFIILRTLVPFVTKMD

KASILGDAIEYVKQLRRRIQDLEVRSKQMEAELKKSAEPRKQLSTTSQEKIVRQKSGGTT

NNNNVDQELVSSCRNFSDHSDQQQYKISRFEKRKIRVLERSEPLTIIDDCSTDVQVSIIE

HEALVELQCPWREGLLLDIMQTLSNLLFETHSVQSSVVNNTFVAKIRAKVKAANIGEKPT

ITKVKNAVLECIPSRC*

>Pe2057023

MPWAGRNSTPVRPPAGNEEGGLCSRIADKSYKEPPGPNESLFCQQIDLKDSMNH

CVPNWDMENGIIQTSDFLPMAKVNWDYVQPQKRSMMLEHDIEELLWENGQVVNKVVKRSL

PTQLSKTLQNDDAVLPGKPSPVGKDNTLESVVHDMSAANASSIHEDVMDSWLHYPLDDSL

EKEYCSDFFVGMPSTNVHMLRDSLAGQMNNVPSEKVSNLHVQKESSLSSESNSVPWTGVF

AGMVNTTNAAGAIKTGKESPPKSISSDKAMALGAGRASGIIPQSGTDTFAKVRTTTQLPV

SKWHTDVQGSSSSKDDQPPCKKSVTQVSVSPLSMPPPKMQATDLASMKLSRSKLVNFSHF

SRPAAAMKANLQSIGSASGLSTIGIQNRLDKIRMDGNATAEPSIIESNSTGMTTIGSSSG

ANSRAQDNGSCQTCQNPPLGKNLDVATCSEDVTDSSAKASEQVVCQNSDAGRSLASCGTE

KCGDAGDGPEPTITSSSGGSGNSAGRAGKEATNTSKRKGRDLEDSECQSEDVEYESADTK

KQAPSRTTTSKRSRAAEVHNLSERRRRDRINEKMRALQELIPHCNKSDKASMLDEAIEYL

KTLQLQVQIMSMGGGMGMPPLVFPGGMQHFQVPQMAHLSPMGMGIGMGYSMGMLDMAATS

GRPVMSLPSMHVSALPGSAIHCQAALPLSGMPGPSIPMSRLPGPGLQVSGLPVSTIPVSG

LPGTTQPGLVSNSASGSTDLQDHMQNANVMGHYKHYMNHHQMQGPPQVINGNLYNASMAQ

PPPQNPVQSSGRGIHNAASASGKTGTTG*

>Pe2057055

KKVLRELHELIDGASDPAADEEVTDVEWFYLVSMTHSFTGAEGVPGHAFMSSAPVWLSGS

LKLESFGCQRARQASQFGIQTMVCIPTPNGVVELGSMDLVCENWGLLQQAKSSFTFSSSF

WEDNGNGNINHNNNHNNNYGTNNQSLWNPGSPFLTQESILGDLSFLNNEESQNRNSSAQK

SLSILEENRNPLPPFTVQKQVALEENRIPPSFPAQKSAVLDERTSLPLVQKTGISEGAHN

PLPFLSKPPPGGGFDEKSNTLPYSAVQRPAIIDENCNSLLSQKSSICSEILNSYPYVTGP

KSVDFSETCNSYPVPALQKTGIENSNQNLLQSAAVQKPNIADQIGNPNSNPNPLLFSVQK

TCTIDQSKGFIKDLLIEDDKPKPLLDFQSNNFQRHTLSFANGYSRNMVEEKMVKPISIDD

EKPKSLPTISSGAVFGGVRSSIESDHSDVEAASFKEANQAVIEKKPRKRGRKPANGREEP

LNHVEAERQRREKLNQRFYALRAVVPNVSKMDKASLLGDAVSYINELQSRVQDIESEKKE

LQAQIEATKKESSSSHSAFSGTNLGFIKDQSGSSQKPDVKRFGTKECSALDLEVRILGPD

AMIRIQSTKKNHPAARLMTSLQDLELEVHHASVSTVDELMLQNVIVKLPSSLYTEEQLNA

ILLKKLSDPKFK*

>Pp001G01120

MPSHSQIGQVGLQLQHPAQLLSYPMPPCTPQAHCRSDSLLPYSLESWFPASSTTFEATTRSCGSPPXWSNFQQPVETPEG

SVGDAIRISTGSAISQPADFQCDTWWSPITVQAAMPNDNLISQRGSEISCDLEAGASYPMIFTTDCELDSRQRAIGNIGG

CDQNEPRKYPTSALAVVNVAQIPASVSRSKGASRKEILQLAKEKLARTPTSPSEVRLGKEKQLGVQKSKGSGKRPLSQRE

NHIWSERQRRKGMNYLFSTLRSLLPQPNPKTDKSTVIGEIIKYIQSLQVKLEMLTKKRQQVMTAVLPRPGSSASHCTGLT

LLDHSNFDSSSMTAITALPPPGRESCLQSYLGTNVGLHVCGLNVFITTSSPRGRRGLLQQLLLTIQRYNLEVINATISTS

SASIFHCLHCQASQNAEVLNNDLHSALQTAITNFGLTQF

>Pp001G02200

MYRKLRHLSGSSIPPATIAQQLRNSLQAYCPTSLPADTPPSNEATMNHKPIARSDSHFPDPFWLLQDFSSDIPDQRDVNP

NPLVPNSSQTSTSMWGGVVGDREFSTQPVTAPASKHSQSSPIIGSPGDDVMEIPANSSDTAEEKPGGRKCSHSRCVASKN

LVSERKRRKKLNEGLFQLRAVVPKISKMDKASIIGDAIAYVRELQKELEEIESEIDDLEQKCTGSVGEETGSVEEAGTGA

NFSSPTYSNPASGVEIQGAEPGVDSVDVVSADATQVQLPARLAQKILEVDVARLEEQTYHFRIFCQRGPGVLVQLVQAVE

SLGVQVINAHHTAFQENILNCFVAEMDSKMETEDVKRTIFSAAAQYGLAQG

>Pp001G02570

MYRKLRHLSGSSIPPATIAQQLRNSLQAYCPTSLPADTPPSNEATMNHKPIARSDSHFPDPFWLLQDFSSDIPDQRDVNP

NPLVPNSSQTSTSMWGGVVGDREFSTQPVTAPASKHSQSSPIIGSPGDDVMEIPANSSDTAEEKPGGRKCSHSRCVASKN

LVSERKRRKKLNEGLFQLRAVVPKISKMDKASIIGDAIAYVRELQKELEEIESEIDDLEQKCTGSVGEETGSVEEAGTGA

NFSSPTYSNPASGVEIQGAEPGVDSVDVVSADATQVQLPARLAQKILEVDVARLEEQTYHFRIFCQRGPGVLVQLVQAVE

SLGVQVINAHHTAFQENILNCFVAESFMTSQQMDSKMETEDVKRTIFSAAAQYGLAQG

>Pp003G01120

MAGNDATPLLLRCNKSASIQTVICVPLKDGFREFGTTQNVPVDSKMVQHLLSFLEQPKHNRSQSSLVIGQRYEPSNMTCQ

KSEHEAHGNQSPSSGKTQRSRNVDIRRTHSFLHFALAQWIQVFLHNFDGILRHASARNPKRAKLSNPPRTKVEDKMSLNN

VNISTRLSRQGPRTCVHIFQRLQGSASRRNGEQFPEGSKRTSPASAFVIWKNSLTPPRKYRKAGNRQWILKEVLFHVTQL

FNGTLKEINVEVESSLVTTEKVGEDRRASELESHKPVPSIEETNASHVLAERRRREKLNERLLSLRALVPNVSKMDKASI

LGDAIEYVKELQSRLQAFQIRAEP

>Pp004G00390

MPEPCDRAAPGELVAETLKRVLKLFSFLKLGFRGIRPSDVPLYDDTGVQDLNSFTPAYQQVIPGSHGVPVPVSWSATDPT

LLLNEQSSGKGLLSWQAESCVQQSLQYSQAGFDTVGHVEMMAALNADPYKSLGNYELTQLVQLYSTYYQDPYPYSNLSFS

DSVTFQPDVSLPTSSDCLQVVSSSGSDSLSTTETWRSETSGGCAPSSYENVLTNLNESSSVTLVRDSHDSNFQPQRPHLD

MLKHDQAVSLPPVHEMWIQNQIQAGVGRNTLHTVLALTQDMPSYDVLPVKSSSDPPTFKRPRTWGEALQQGSAHCDSLRH

SLASKTSPTLDSTQLSSLSRSIPSRQFSFQGKLSPSHLSGAGGSTYSRLPGLPEEGRLCNGKPAATSVEPQSVAARHRRK

KISERIRVLEKLIPGGNKMDTATMLDEAIEYVKFLQLQVQILESDTLDNAPLTASNGQNLPLQGSRAKQGLKRKGDTSLG

GDSSPALSVRAATSPLILSEVLQQQLFKQKLCLVSIRQCPPRAASTQAAAAPSVLRKN

>Pp004G00440

MRCEIRVTYSIDAVVFVFCSGQICWLAKAHGCETRCSRQLPGGRLKTLELLCLRESDCVYWTGCMNRFHSYCPQLDSGLH

SSFGSHATACPLKASDQRESSWGTPHALLRPSKLSPNPTSHQAQDDNGAPHKMTKEHPQDDEPEDLRRDRTWFSIPVKRR

AEQKNVQRGNRSMEPPPKECLLASGAILYLLGKVRVLTIAFEGRERIQVILRCITWMSRSSTSELRNAFEFIALGLLMLS

QKVCHSTSMNGFELSLLCLRSARIFRQAEAGSRAITSHGDREVSNEHILNSMYKSCTFTPNFGSVGTAYAEGRHIWLNGA

AVHLSAGSTEQAQFLRHAGIQTAICIPWSDIVLELGTCENVAEDLKLMERIRIFITERILPALLGTSQSPPNSPPIVYNI

GSPFSSYSSMGHLSGDLNYQEFDPNSLNQQMRRLTKAPTSLNTVEMGSGSDTDLLLDPALWTIPTTSKPTCGILQSQPAD

QMPPQSIPCRGQHRSQSATTSTSGSSLEDRFNTRLSDNSPEQLHMSQCMTSSVRCNSSPNSRSLPRGGGSPHSPESMNEL

HDTNTDPITAFQSLPQHEGSQKAFSFMQNEEVELSTQEDLQHVYNPSLKSGNASERNLLRSFASILAEDHGEQPDSKSCP

RAHVLAPIFNTNFDGSSQKLVRKAVEIMKQIPNLQQGIQASGSAPPNIEQQQPPLASCSSPKASKDADEARDPFGQDAPW

SGRKRPCRGSRIPRTDQVHRAHGEAATNHMLAERRRRVKQKENFNALRKLVPIISKADKASILGDAIFYLKDLQKQLEEL

EAISTQTENQYKILRSSYNNLQRQNEELEAIARNDALCHTIPTRLNSC

>Pp004G00490

MIFACNMIILHRNLEWRGGHFNVGSRAITSHGDREVSNEHILNSMYKSCTFTPNFGSVGTAYAEGRHIWLNGAAVHLSAG

STEQAQFLRHAGIQTAICIPWSDIVLELGTCENVAEDLKLMERIRIFITERILPALLGTSQSPPNSPPIVYNIGSPFSSY

SSMGHLSGDLNYQEFDPNSLNQQMRRLTKAPTSLNTVEMGSGSDTDLLLDPALWTIPTTSKPTCGILQSQPADQMPPQSI

PCRGQHRSQSATTSTSGSSLEDRFNTRLSDNSPEQLHMSQCMTSSVRCNSSPNSRSLPRGGGSPHSPESMNELHDTNTDP

ITAFQSLPQHEGSQKAFSFMQNEEVELSTQEDLQHVYNPSLKSGNASERNLLRSFASILAEDHGEQPDSKSCPRAHVLAP

IFNTNFDGSSQKLVRKAVEIMKQIPNLQQGIQASGSAPPNIEQQQPPLASCSSPKASKDADEARDPFGQDAPWSGRKRPC

RGSRIPRTDQVHRAHGEAATNHMLAERRRRVKQKENFNALRKLVPIISKADKASILGDAIFYLKDLQKQLEELEAISTQT

ENQYKILRSSYNNLQRQNEELEAIARNDALCHTIPTRLNSC

>Pp005G00980

METGPPNLWDATDPLMVEAFIGGYGIPGYETQVDLGCTAGQDLEQNDSVLQRRLHTLVEESSENWIYGIFWQRSLSPSGE

SILGWGDGYYKGPNDSDEFDSRQTLTEEHQLQRKKVLRELQALVSCLDDDATEDVSNTEWFYLVSMCHSFALGVGPSRIY

YSSRKLDWVTLCFECSTPGQALALGQHIWLEEADKASNKICTRANLAKTILCVPTMNGVVELGSTDLIHRRWDVVEHIKM

VFQDSTWGLDDMQIMSHSQVANFDSTLMPYNSSMTLDPTSIAGTTSITSNTDPGLADHESVDFNARLFHTDKLLLHSGGL

GFNGFDHLWGQTNDFHCNDPLPDDVENSLGQSMYDILGKLPLQEEQLPFLASSSFPKNPESNSRHSVFQMNNVGKLPQLD

HRSASLPAMEKPHSKLQPTYTFPQYSGDFEVGEMYNPPGHIPTMRPKLPNQEERVQFPLMPAVEKASVIEKPMPLLKPIP

QSPSPPVSKPAGPSVSANGLKLTDHLGQNFVDPESVTIKVNVMEAPKLPRKRGRKPANDREEPLNHVQAERQRREKLNKR

FYALRAVVPNVSKMDKASLLGDAIAHINYLQEKLHDAEMRIKDLQRVCSAKRERGQEALVIGAPKDDTQLKPERNGTRPV

FGIFPGGKRFSIAVNVFGEEAMIRVNCVRDAYSVVNMMMALQELRLDIQHSNTSSTSDDILHIVVAKAQESLNRLSAGCT

MGLKGYPDEEVLERVEILRKALEAAGALEVDDMDSRSVSILWFQRAVYVCSGSCKSKRCEYFLEAVQDFAGSFASFFKQL

ANCTWFFCEYRMNSTQSHHIAAQKEKQLESMRDALGLKNVKEGEASDRELQEQKRQERIMAREEKEQENREKQAAIEEEE

RRQKKELRREEKLRHREQQEEERKQRKLERKVRREQDEQERYQKGKESRELMEPRESKEKRDQEYEERDHKKKESRERRK

NEDEEPHERIKDPREKRENREH

>Pp005G01070

METGPPNLWDATDPLMVEAFIGGYGIPGYETQVDLGCTAGQDLEQNDSVLQRRLHTLVEESSENWIYGIFWQRSLSPSGE

SILGWGDGYYKGPNDSDEFDSRQTLTEEHQLQRKKVLRELQALVSCLDDDATEDVSNTEWFYLVSMCHSFALGVGPSRIY

YSSRKLDWVTLCFECSTPGQALALGQHIWLEEADKASNKICTRANLAKMAGIQTILCVPTMNGVVELGSTDLIHRRWDVV

EHIKMVFQDSTWGLDDMQIMSHSQVANFDSTLMPYNSSMTLDPTSIAGTTSITSNTDPGLADHESVDFNARLFHTDKLLL

HSGGLGFNGFDHLWGQTNDFHCNDPLPDDVENSLGQSMYDILGKLPLQEEQLPFLASSSFPKNPESNSRHSVFQMNNVGK

LPQLDHRSASLPAMEKPHSKLQPTYTFPQYSGDFEVGEMYNPPGHIPTMRPKLPNQEERVQFPLMPAVEKASVIEKPMPL

LKPIPQSPSPPVSKPAGPSVSANGLKLTDHLGQNFVDPESVTIKVNVMEAPKLPRKRGRKPANDREEPLNHVQAERQRRE

KLNKRFYALRAVVPNVSKMDKASLLGDAIAHINYLQEKLHDAEMRIKDLQRVCSAKRERGQEALVIGAPKDDTQLKPERN

GTRPVFGIFPGGKRFSIAVNVFGEEAMIRVNCVRDAYSVVNMMMALQELRLDIQHSNTSSTSDDILHIVVAKMKPTERYT

QEQLAALLERSDQATGYLTKREGSDTALQKLGNPP

>Pp006G00650

MTDLISILESSGSSREEMCPVAVPSSVASSCERLIWEGWTAQPSPVEESTTSKLLPKLLPELETSSYSALTLQQPDALSS

ILSVLHPFSHYSSASLELARNPDWSLKSSNPLRESSSEAGIRTSSFEGLYSGQHTTKKIHLGVIPYHLSEDQRQCAVSPP

ENECRLLSANSSGSLHWWHSIGPESPSSTLAFHNIGIQHSTFEKCEPRGQSHSSWPAASGTSPTVQYFHAHSADNEGVEV

VKQDDSQISKALATYQPHGDHSLVLNSDRIASTTSHSEDPCGPKPGRRPAASYDTEMILSPSESFLTTPNMLSTLECVIS

GASNISDQYMNFVREPQEQRLSSISDLSLIPDSHADPHSIGFISGTFRTDSHGTGIRKNRIFLSDEESDFLPKKRSKYTV

RGDFQMDRFDAVWGNTGLRGSSCPGNSVSQMMAIYEFGPALNRNGRPRVQRGSATDPQSVHARARREKIAERLRKLQHLI

PNGGKVDIVTMLDEAVHYVQFLKRQVTQSDADGGPDFEICVANAASEIRRVLDVRHADLVPEQIRRLQSGSRREQLILQE

SWHVATVRQHGKSAVPPKSTMCQ

>Pp007G00140

MSPRSTLKSRSKSQVMHPKQHYIHVRARRGQATDSHSLAERVRREKISERMKFLQDLVPSCSKVTGKAVMLDEIINYVQS

LQRQIEFLSMKLAAVDPRLDINLLNLLNKEVHNLICEMA

>Pp011G01580

MNSHLVLYQGQMAGIQIRESWEVVQNVKMVFDEPMMWAAHEIQAVAHSLPLSSDATSMRPSSPSLMSIATISASISNASQ

IGRCSNPSQDHEAHFVGRKNPPSSRPPMRFYKPRVEELSSDTPETNLVDNKYIVGKSVSFHHLSKIGQRGMPGPPTTANR

LPCFAPSEVDCHESEADVSVKENVVESSTNLEPKPRKRGRKPANDREEPLNHVQAERQRREKLNQKFYALRSVVPNVSKM

DKASLLEDAITYINELQEKLQKAEAELKVFQRQVLASTGESKKPNPSRRDSTESSDEERFRLQESGQRSAPLVHTSENKP

VISVFVLGEEAMIRVYCTRHSNFIVHMMSALEKLRLEVIHSNTSSMKDMLLHVVIVKIDSTQTYTQEQLGVSETRNWCW

>Pp011G01800

MSQLLNAWEVADSAMIEAFMGTGYNCVEGFEVQDDPDGQLHLNESVLLRRLHSLVEESTVDWTYAIFWQLSALREGEMML

GWGDGYFRSAKENEINDARNMKGGSQEEDQQMRRKVLRELQALVNGSEDDVSDYVTDTEWFYLVSMSHSYAAGVGTPGRA

LASDRPVWLIGANKAPDNNCSRVQLAKVHSSMILQTILCIPSKSGVVELGSTDLAKSWEVVQNVKMVFDEPMMWAAHEIQ

AVAHSLPLSSDATSMRPSSPSLMSIATISASISNASQIGRCSNPSQDHEAHFVGRKNPPSSRPPMRFYKPRVEELSSDTP

ETNLVDNKYIVGKSVSFHHLSKIGQRGMPGPPTTANRLPCFAPSEVDCHESEADVSVKENVVESSTNLEPKPRKRGRKPA

NDREEPLNHVQAERQRREKLNQKFYALRSVVPNVSKMDKASLLEDAITYINELQEKLQKAEAELKVFQRQVLASTGESKK

PNPSRRDSTESSDEERFRLQESGQRSAPLVHTSENKPVISVFVLGEEAMIRVYCTRHSNFIVHMMSALEKLRLEVIHSNT

SSMKDMLLHVVIVKVR

>Pp025G01320

MSISIPYKGDWFISPTRNQRSRCATNKTIQSESSTFWNEKARYTKIAIISHLAIILHGSLRAVMSKQAGGGVQERLRSIV

GPKGWDYAVFWQLRDETRTLDWIGCCCSGAEAPGNDHLGASSSTRYLESSTGCPDVRGFHPATHICSLLAAMPSSVSLDS

GIQGRVFLGGQPKWVHGDSSMEGQELTVQTKVCIPVQSGLVELGVANHVTENAALVNYVRNRCGEQWQAGTVSKQGGSGN

TTLGTMGQQAVKMYYTRHFANNLDSSWPSSHPWEQEDPTLESQLMGSMDQDLMNQLPSSTTHFPHHGTLTDSPRGSGLSK

DDGEVKQEMRGESSDCSDPIDDDDEKGGTRSTRRHLSKNLVAERKRRKKLNERLYSLRALVPKITKMDRASILGDAIEYV

KELQQQVKELHEELVDNKDNDMTGTLGFDEEPVTADQEPKLGCGIVIGRSCPKVDSQSVVIEVVDRKGDHELTQPMQVEV

NKMDGRLFSLRIFCEKRPGVFVKLMQALDVLGLNVVHANITTFRGLVLNIFNAEVRDKELIGVEQMRDTLLEMATLQSGH

RPLPLPPSDGTSLQTSIMSGDQRLEKVTHLGPLP

>Pp025G01330

MRGESSDCSDPIDDDDEKGGTRSTRRHLSKNLVAERKRRKKLNERLYSLRALVPKITKMDRASILGDAIEYVKELQQQVK

ELHEELVDNKDNDMTGTLGFDEEPVTADQEPKLGCGINLNWVIQVEVNKMDGRLFSLRIFCEKRPGVFVKLMQALDVLGL

NVVHANITTFRGLVLNIFNAEVIASPS

>Pp028G02130

QDYIHVRARRGQATDSHSLAERVRREKISERMKFLQDLVPGCSKITGKAVMLEEIINYVQSLQRQIEFLSMKLAAVDPRL

DIN

>Pp030G01900

MSSNSARNLEDLTWKAPTSSQNTTTDVVKDTAEKSNFDKGVLEEEWYTPETSLMELSYSSSYEMQDAPSNFSLLEPSLNF

DSVNSHASFRLAPSINLGMSQLQGNGLLDNLACTGQGSSVSLLSNIPPGGQLGHISTMDSYSQGIIFPGSSVGNIASSSI

NARSVNLPPSFADAYSVSALNQNKPGEGGSGVSRNDFVLHHTSKQAKSCLPGQQTLFQKGAASRRSVAGGTVPSASKSPS

RVFTSASNDSSMDTRDKESPHTGNASRAEMLSGSDDPNDIRLDGDDYDPDESGDGSGGPYEVEEGAGNGAENHGNSKIKG

KRGLPAKNLMAERRRRKKLNDRLYMLRAMVPKITKMDRASILGDAIEYLKELLQRINDIHSELDAAKQEQSRSMPSSPTP

RSAHQGCPPKAKEECPMLPNPETHVVEPPRVEVRKREGQALNIHMFCARRPGLLLSTVRALDALGLDVQQAVISCFNGFA

LDLFRAEAKDADVEPDEIKAVLLLTARRGMHPLQ

>Pp030G01870

MTRKDSNFKAGGREAYDSSGRIVLLNKLQKFCDHSYTLKLAPEDLTWKAPTSSQNTTTDVVKDTAEKSNFDKGVLEEEWY

TPETSLMELSYSSSYEMQDAPSNFSLLEPSLNFDSVNSHASFRLAPSINLGMSQLQGNGLLDNLACTGQGSSVSLLSNIP

PGGQLGHISTMDSYSQGIIFPGSSVGNIASSSINARSVNLPPSFADAYSVSALNQNKPGEGGSGVSRNDFVLHHTSKQAK

SCLPGQQTLFQKGAASRRSVAGGTVPSASKSPSRVFTSASNDSSMDTRDKESPHTGNASRAEMLSGSDDPNDIRLDGDDY

DPDESGDGSGGPYEVEEGAGNGAENHGNSKIKGKRGLPAKNLMAERRRRKKLNDRLYMLRAMVPKITKMDRASILGDAIE

YLKELLQRINDIHSELDAAKQEQSRSMPSSPTPRSAHQGCPPKAKEECPMLPNPETHVVEPPRVEVRKREGQALNIHMFC

ARRPGLLLSTVRALDALGLDVQQAVISCFNGFALDLFRAEAKDADVEPDEIKAVLLLTARRGMHPLQ

>Pp031G00290

MAERRRRMKQKENFTALRRLVPTISKADKASTLIDAITYLKDLQNKIQEMKASKEDINQRCETLENKCRELEDRNQQLVA

MLSINHPSSSNQLDT

>Pp031G00600

MRLKLAVATHCLGWTYSIFWKLISEQQVLVWGEGFHNSLNPNFALRRSEQLRNFFIAMNATRDTAAQRVSATPPPLAPEE

ISATEWFYMGSMACSFAAGAGFPGRVLAERSFIWHCGPVGAGGSSRVFTREHLAQTIVCIPAPDGVIEFGTTALKEEVQI

QYSNEAETSAQLYSGMRSFRGGPYSSGGSSISAHARNERNRRQKLHDRFMTLRSLVPNITKPDKVSLLGDAVLYVQDLHR

RVTELEASKAPTPKTPTEPRVEVTIEKNTAYLKLSSPWQDGLIIHILERLHDFHLEVVDVSARVSHDAVLNAQLKAKVSD

NTRYST

>Pp031G00820

MAASSNSWLHQNLLPQHNAQQQGSPHPLYHHSELPHVGSSQSGPLQHMNLAHVGHGMDSHPPLHHGMHSALHMPREMQQQ

QDPHIPVHVGTGTSGRAVGPSENGDAASVRKVHKADREKLRRDRLNEQFGELAGVLDPDRPKNDKATILGDSVQVVNELR

SEVKRLKCEQTALLDESRDLQQEKSELREEKAALKSETENLQNQLQQRLRGMLPWISMDPTAALMGAPPYPYPHPMPVPQ

PPPSPPPSSTPPASESASGLTTSQPSVVGPSPFLSLAPPGVYMHPYAMYGARSPDGGHPYMPYPPYPGSVGNHSQVERPY

AQYPSPIPTMPGYLVQMQPPPPQGPANASAVPMYRPPPPPYAVSGISLVPPGHQPRVPPHSTNPPINGPQAASPEPHGVE

TSLQLQTPGGSSRSPSTSELPPSQRSGEPSKTHERADLELTPPGTKEASDLAGSKPDDINGGDSISTLTLVSKSLQSRAD

ATPSPSGGSNSEEGHLRGSTLEPSLRSAEQLPKL

>Pp031G01350

MQWVTELGMEDVRQFDPLFHQSNAEAYGGGLSSVSGSGNNYAACLESFKQQKERQSFADLDGLYHQERPQKMLKSNSSMP

TLNDFHYQASPNWSMSMGLSSGLGGISFGSQQPLAQNPHQQVHQMIPQHLDAYAMSVQHLQEDSAVHPRTPGSQVDCGSS

PVSVGTNGCKPDAVSVESLRGSSPLSSIEYVQQGKQKSSGFFASNVSGSRLTMPPQPPPPVKSTGHTQDHIMAERKRREK

LSQRFIALSAIVPGLKKMDKASVLGDAIKYVKTLEEKLKALEERLPKKRMRSLSVKNMPPVPPSSSNSQGCSKLAPAVKQ

QLGEEVVDEDDGSQPEIEARKIDKNVLIRMHCEKRKSLLVKSLAELEKMKLVILNANILSFSATTVDLTCCAHMTDGCDI

NTDEIVRTLQDLYYTLED

>Pp031G02090

MEVSRFGNAKARARAGARRSAFQRWDYARLTKPPHTKQGSHQQQMLKTSLLSIPRLFTVRRGIDEHDEQLSGNAMESAET

ADGHIQRGDGRHMMAERKRREKLNDRFVTLRSLVPYVSKQDKVSLLGDAIDFIKDLQRQVEELESRRKISENPSKPRVEI

TVENNRAVFEISSPWRQDLLIAILETFVGTHMQVEDVAAKVSKDTFKATLKAKVTSSGIDDDDDDEDKKSIKQIRETLLS

VVNGNNSPTFEKLC

>Pp031G02110

MESAETADGHIQRGDGRHMMAERKRREKLNDRFVTLRSLVPYVSKQDKVSLLGDAIDFIKDLQRQVEELESRRKISENPS

KPRVEITVENNRAVFEISSPWRQDLLIAILETFVGTHMQVEDVAAKVSKDTFKATLKAKVSDTVVVS

>Pp033G01440

MRDVHAPSFCPIIFLARRCLVAGWVVSHLEEAEDKGEDLVDWLRHLVLEDGQRTDRGFRFRRGNEVKSGSSDIAGFSFCE

SDSRFEEDRYRGWRKGPVSSRWGLGLAPLSTQRMALNMGMPMEYNDALDHCNYVDQEYWSDLTDTESNDDYTFTPRAFEH

IVPRSSKRTYGVFFASEAVLEQKKRSRINEQFELLRAAIPSSCTDVKASILTGAYEYIKKLERQVQDLQHELDAESCCED

DASYSDDDLSTCEVDLRQWFTEEKRPVGCNAASEAELTSCHGCPQPTNLHAAKYLSSHFEALRDQVEVVQTQGRLKIHVE

CEKRSGLLVDIMEVLESSGLNVEQASITCQEHLVFDGVGSEVLRIAHLAVILELLIITCLVSRQRDHISNVEEDDGAECR

RVSYVNAEHVESRLRCLIVKQKD

>Pp034G01160

MGYDDLLDLYSPCKTRARRGYATHPRSIAERNRRSRISERMKKLQDLVPNMDKQTNTADMLDEAVEYVKHLQTQVKDLSE

TIVRLKNSIRAQTAS

>Pp037G00600

MAFVATAGQKQLVPERAIRSSSSLGYSSFGELAPGDSVTQQQREFQVVHRYTMAQQHAERAQSRHFAHSEDTKPSTHDFL

SLYDKGSQQAENVSSAGLSSSLTTHDFLQPLERNGARPTSRDAPVLREKHSSGENGQSRHSTPGPGEQAHPTSSPNYAAI

KVPYTVPPNSMLRMVGEAEAVRQGRNYPIATLSSREDHSGSETFKGVEIAVGPEAANAQENLAHWPGVRFQFGMPPDLTF

TAGNPAAKSVVENGHQLLPTMTKHWAQAAQGMIESPKRPARPPSDEDEDDDEDSSERPKVNRKEPMLRKGDGAQRMDSKN

ADLRGVTPRSKHSATEQRRRSKINDRFQMLRDLVPHSDQKRDKASFLLEVIEYIQVLQDKVRKYETVEQGRHQERLKSMV

WDLCKRGAAGSGDETSQINALVFAKDLNSGVRSVDIKPGTEVPADRVPRSSEEAAPISEAHSTGGLNAQQASSRPACSIH

SCVSSLGGEKESATAVTAVGVPLQQAIPSPHAFPIVPSKGQREQDSNRGTPASNSNAPRKEKTSDMDLVSSRENGYAHDT

TPDHQFEVRPRISSGQADQNSDHQPKDHQMSATTIVGRMDGVQAGLLSSAPRQPQGGPQHKEDANKHTRHPQTNQNDHMA

TRDGFAVFPNIPQPFPRIQGGVINVSSVYSQGLLDTLTRALQSSGVDLSRANISVQIDLGKNATVAPGDTSAKVATTPES

SAQSHQSQGRTRPAPSVPESEPARKRPKLLSSAYTKHHCLLRSGCGKIEKDRFGFRLCYDQHDAGFWHNECRSFGLSFAK

ISLSGLEMAIWRTALLRFVISFEPTSAGKEGVHAVVACSLITIFH

>Pp042G00840

MQWLSEMGMEDHQLVRQFDPLFNQTSVEQYGGGLPSGSSNDYVAPLESLKQQKERQSFADLDGFYHQERPQKMLKPNCYE

PNLSNFQYQANPHWNTSMSFSSGLDFMTFATHQPSGQNPHQKQEHQMLPQQLDTYAMPVQHLHDDGALHPRTPGSRVDSG

SSAVSLGANGSKLDALSMEVLRGSSPLSSIEYVQQGKQKITGFFSPNGGGSRLSMPTQPPPPVKSTGHTQDHIMAERKRR

EKLSQRFIALSAIVPGLKKMDKASVLGDAIKYVKTLEEKLKTMEERLPKKRIRSLSNKKSSQPSTTPGPVSQGESKPAVV

VKQQLSDDVVDEDDCSQPEIEARKIDKNVLIRMHCEKRKSLLVKSLAELEKMKLVILNANILSFSAATVDLTCCAQMSEG

CEVNTDEIVRCLQELDSQTMPSAISGLLPFPQVKMDMAIASEFFNVVNPFTIFSTVQVLFLQISLVPVLVHISKECVYLE

LCPELYSEGSDSMTSLTTFSAASTRVMAVFAKAVADFAWFLFHSRVLGTSVVIHTRDDPLWG

>Pp048G01150

MELGRRHASMIARRELDTIFAEVTRRCNTQSVGGLGVFGTDSAISKLDIPIKLLSEYLYLRWALVELYSFFGRRRVSRAS

SLSLQMSLTFRADDCNELPSMENWLNAGADWNEHHGIVDVDATATAPLFYHTAEASTTSGEDSLFDALKKGQEAISTHPT

FLKLDTLELCKQNPDAVLVRTGSSSMVDSSSNEVDVSDHANSLPSTGMLVSNLLDRTGSLDESGLYMEADSPRGPLSPAA

LQLGKSPKGVAIKRSFEDASEASWMPAAEATHIRARKSAKQQLPKNVALSQEDLQASAAAVPYPPFVAMASSGEMAPLAI

RSGVAPAFVPASAANVPPLLFPTPLTLPNVPSMEEIAQSRPKRRNVRISKDPQSVAARHRRERISDRVRVLQHFVPGGTK

MDTASMLDEAIHYVKFLQQQLQTLEQIGNMSDPRFMAQPGGPMMMPQMGAMNASANAATRIWNHRPMDTFKSIGGCFEGE

VWRGPLVLRGDHPRLTAEIPHRSRIVKILRFVLMVLPNRPWPLHVERCYSCFSLSVCLWLRWLTTRVMCSGVNVPLRHAV

YNKANVLKNKQGILHYTFVAAVLREDLVGWADPFSLLKLLRNNFKPCLIRCSDKLYIVIATISLHLCCVIAEVSKGNPVN

QYLWTSFQNRRTAKSSKSDNWTARRVQHED

>Pp051G00900

MASQSNSDMWMGAASSEISLPSNPALELEEETSWKVSTSSQSTSNIEVKENVGTPSFSKGVLDEEWYTPETSLMELSYSL

PYGISDTRTGFGMLESSLNFDSSSNLMSSFRPAPSALSMGLESNRSLEDLVCTGQGSSNVGLLSSLSPGGQLGRSTVMES

FSSGLPTSFNQGIINAGGSITNITSSNINNVRSNFPLMASPSNFSDAYRARSVSEDKSGKVVGSGGPRNELVPYHRNKGA

ETRSHGQGQQTLFLKRAASRRCAGSSGTVSPVSKSPPRVVTSASNDSSVDTPDKDSPHPRNAHLQSASGRLNINSGSDDP

NDMGLDGDDYDAKDDDDLDESGDGSGGPYEVEEGAGNGADQSIGKGNGKGKRGLPAKNLMAERRRRKKLNDRLYTLRSVV

PKITKMDRASILGDAIEYLKELLQRINEIHNELEAAKLEQSRSMPSSPTPRSTQGYPATVKEECPVLPNPESQPPRVEVR

KREGQALNIHMFCARRPGLLLSTVKALDALGLDVQQAVISCFNGFALDLFRAEAKDVDVGPEEIKAVLLLTAGCDLHSLQ

>Pp051G00950

MSSFRPAPSALSMGLESNRSLEDLVCTGQGSSNVGLLSSLSPGGQLGRSTVMESFSSGLPTSFNQGIINAGGSITNITSS

NINNVRSNFPLMASPSNFSDAYRARSVSEDKSGKVVGSGGPRNELVPYHRNKGAETRSHGQGQQTLFLKRAASRRCAGSS

GTVSPVSKSPPRVVTSASNDSSVDTPDKDSPHPRNAHLQSASGRLNINSGSDDPNDMGLDGDDYDAKDDDDLDESGDGSG

GPYEVEEGAGNGADQSIGKGNGKGKRGLPAKNLMAERRRRKKLNDRLYTLRSVVPKITKMDRASILGDAIEYLKELLQRI

NEIHNELEAAKLEQSRSMPSSPTPRSTQGYPATVKEECPVLPNPESQPPRVEVRKREGQALNIHMFCARRPGLLLSTVKA

LDALGLDVQQAVISCFNGFALDLFRAEAKDVDVGPEEIKAVLLLTAGCDLHSLQ

>Pp055G00060

MRAGVRSCCREEHKAGRGQLESFHGDLQKILLSPDRPNSHRFVEVVITMCRESSVLRPPRSLLKSPFPSPNLFSEFFSVR

HALGDVEDIHIGDVLEYLLAISVTMTLSLRPDDRSALRTMEKWSNSGAGWSNHQHPMLESSISPLFNYTQSHNGLQSQSM

FLHPMGAAEQQALVQAFVQANVGCMMDQPSQSGCGNLDSLLKSSNSSSMVDSSASEMTSPDDDCTGAPENNLHMNCIPSL

GGGPVSPVLTSMLDRTASNGASLMAAHPQSKNDFFRMRSPGGPLSPGASACAESEDSLHGGEACAPAMTGLKRRSFEGDD

GDWMVDQMHLSQKVEKQHHQQQPEKALAFQESFGGYNVEVASTVGLTATYSDSLTIPSLMQPSPQPFLRSSGNCGAASPV

DLDEFASMRAILFRHASQPVPSLEEIASSRPKRRNVRISKDPQSVAARHRRERISDRIRVLQRLVPGGTKMDTASMLDEA

IHYVKFLKLQLQTLEQIGNNGCDPRSFLEQGGATEELANLVRPFDLNCTAAFQTWPPQTTTMVNHGDTSPQGHNSTEWAC

EANQDDCPISPIDVPDFEITDLGWKNLYKWYDLP

>Pp057G00920

MPNGALQAVTALVKRHAEQIGITGVPLEHQYQHIQRQLQNLLQQLRWSYVALWSLSHLAPPQTRQTSQNETAMIRGTYPQ

WVADTCRSVGMAHAEGRNFWMNGSSVHLTAGSMEQAQFLQHAGIETAMCIPWSDSVLELGTTERVAEDPSLMERIRGFMT

EIIPPALPDTSQAQQNRFGGTSITEYSPFSCLSAAGDLPDNLNYQAFDSNSLNQQLRSVSPSARLATSEMVLVKSDDSQV

NYLKSHHCLDTGLLPDPAMLRFPRASRSASDILQMQHQVLPPASSNQFRGRCRSHSSTTSPSASNLEGRFNTRLSNSPEN

INTTFSRSRNSRSTTSSSSRRGTPSQGPEFKLENTNTYTGLVPQLPRDDDEIQKVLTFMQDGGESLGTHREDWQDLYKSS

LETENVMRETYLPSSSGVRVEDGETELGSVSWQRTRAVGPILDPRNDRRSPVLERKAARHCKEIPSLQQGTQNSGAAGQQ

QPVLTFSGAETSTNTCRGQDAFYLGPLTDQRRVRRVSRIASLGPVNGAHEDAAVNHMMAERRRRVKQKENFTALRKLVPI

ISKADKASTLGDAIIYLKELQMKIEELKASTTKTENRYKILELSYYNLKKRNEELESITGDGDFSYTHPLRKNSYQ

>Pp057G01290

MGHKEWTNNDGGFDKSIDLINIGVDLTFMFIYCTTTMERHELKILVAKSERQLPMPNGALQAVTALVKRHAEQVLSLFHL

HAFSDEEPGVGWSRTDHRRSCGNVWDSRLTQHKSRSGFLQQQTCHLSRLQSCTASCHCRALVSTPSHLTDWNYRGPLGTS

IPAHTEAAAEFVATAEVVLCCALVTEPSGSAADKKLGMSISLNPLTSSAHSLPALGKAPTRPSSHRMRHFVITIFNRSAC

EVLEECVRNFVLNKTSQNETAMIRGTYPQWVADTCRSVGMAHAEGRNFWMNGSSVHLTAGSMEQAQFLQHAGIETAMCIP

WSDSVLELGTTERVAEDPSLMERIRGFMTEIIPPALPDTSQAQQNRFGGTSITEYSPFSCLSAAGDLPDNLNYQAFDSNS

LNQQLRSVSPSARLATSEMVLVKSDDSQVNYLKSHHCLDTGLLPDPAMLRFPRASRSASDILQMQHQVLPPASSNQFRGR

CRSHSSTTSPSASNLEGRFNTRLSNSPENINTTFSRSRNSRSTTSSSSRRGTPSQGPEFKLENTNTYTGLVPQLPRDDDE

IQKVLTFMQDGGESLGTHREDWQDLYKSSLETENVMRETYLPSSSGVRVEDGETELGSVSWQRTRAVGPILDPRNDRRSP

VLERKAARHCKEIPSLQQGTQNSGAAGQQQPVLTFSGAETSTNTCRGQDAFYLGPLTDQRRVRRVSRIASLGPVNGAHED

AAVNHMMAERRRRVKQKENFTALRKLVPIISKADKASTLGDAIIYLKELQMKIEELKASTTKTENRYKILELSYYNLKKR

NEELESITGDVSRIEYSAILKSSSLQCHGLVGLLCRHPHELILDLNWRVECEELWSSFGYLIQFFKVNPQVNVSYKILSF

RDELEYSATALSIVHCLVLMCLSLPFVQDKRICSWTVSIAY

>Pp060G01060

MTDLNSSLESPGSSVEETPAAANSVATSCEAMMWEAWSTQPSTGDEASTSKLDLLPELVSSSNSRLSFQQSDLLSNMLSS

FHPLSQHSSAGFELSHNRGGSEHSPEFLQEGSSEADTETSSFGDLYINRQTSRNSFLGSIACPLPSNHSDSGKNIRREDL

SNQLGAKSSAPLQLWQSLGPESPSSPLAYHNIGYRHSHGEKWETGSQSHSPWPTVSNTSSTIQLLGGRAAENEVIQVLKS

NDSEISKSLATLQQYGDHGRQLNLNHSSTTNHPEVIYSSKFGPKPSASSHTDVLMSSTNSSFLSIPTAWSTPEYSMSGPS

TRSEQFMNFVRIAQEQNSVPISGPSPILGSYVGCSNRSKSGISRVVSQETPTAKNRLLACEGSSGPAPKRPSYAAHSDSH

ADQAAAMWSSPNLRRSSFPSILTSQAMEIYAIGPALNTNGKPRARRGSATDPQSVYARHRREKINERLKTLQHLVPNGAK

VDIVTMLDEAIHYVQFLQLQVTLLKSDEYWMYATPNTYKGIDLTNSPPQTQRLQSSA

>Pp067G00270

SVRAILFRHASQPIPTLEEIASSRPKRRNVRISKDPQSVAARHRRERISDRIRVLQRLVPGGTKMDTASMLDEAIHYVKF

LKLQLQ

>Pp068G00410

MAENRNDCSTYSSLVCKNKPSPCDLLRGAEHTTEKHVPVTARAYFRERESEGERQRETEKRWCCFLIFNCLLVVVTSHRF

VCCVFPVLLRSFPGHHISAPAYVALDWKLCYAAASTSIFSGDGESDAGSMRFWGSSGSRRVSSCGCSAIAVWSFKTFLED

GLRVWCTRKQCGTDNRCRIVTVEALVGVLLVANDAAYQCFWVAGIFRRPSGWQWSMWHIRLQHCRTMSLYVPERERSGGM

LDAVIPLNDFHRSKKSHFSQLALAPVPDQDTLEVAWNDNGKIDIQGQDSQRSKSPWQINCISDWSSGQEAEAGVTPSKPD

TPKGGNLLETGGNGSNTAPNPVHNIDVDNDEMVSWLQYPLDDPFCSDFFGEIQEANTQLLRESFTHGSTKTLPRTNFPGA

SGNDAGINRETPSDSAMLLGVGRVSGLLPQGGVEAFNKVRSLHSLQQHSSLPRWPQPLTPNSSSGSSLLATNNLTFTKAS

APTLASPSHHMLPPKTQRVAPVLNSQNTPQPSSPGSMNFSHFSRPAAMVKANLHSLVGINCGPPPSTRVKQQHNSPVVRP

IMEACTSTGSSIAESTTAGQGPSGSYQEVKTQPAVLEREQNNGIEESRWPSAPCPSPTKDSDCVVSEGNFRSTPPDQETC

RLSGVTNGAVLASSDKGASHGTQHPDVQEPTITSSSGGYGTSIEPLQKVRTSNKRKCSEREETECQSEDGEDESVDTKHK

PITTGRGSTTKRSRAAEVHNQSERRRRDRINEKMRALQELIPNSNKTDKASMLDEAIDYLKILQLQLQMMSIRTGMTLPP

MVMPPGLQHMQMPQMPQVAAMPSMGMVQMGLGMGMMEQAAQRRTRMPMQSHTGPPLNVCLANTSSMVDVHDLRYQQPGVL

GYNTSMSRQHQPMQTTKGLDLDKCNAYMLQQHQLLFQQQQLQQHQLHQHQQHLQHHQRAPNMGGVQTLGEGCRQCKLNVI

YLFAILVEWNKSLNVKPRSWSLPFHSGIFSRIRVLLGVAMSHHLGSIFRQLCREL

>Pp068G00530

MSLYVPERERSGGMLDAVIPLNDFHRSKKSHFSQLALAPVPDQDTLEVAWNDNGKIDIQGQDSQRSKSPWQINCISDWSS

GQEAEAGVTPSKPDTPKGGNLLETGGNGSNTAPNPVHNIDVDNDEMVSWLQYPLDDPFCSDFFGEIQEANTQLLRESFTH

GSTKTLPRTNFPGASGNDAGINRETPSDSAMLLGVGRVSGLLPQGGVEAFNKVRSLHSLQQHSSLPRWPQPLTPNSSSGS

SLLATNNLTFTKASAPTLASPSHHMLPPKTQRVAPVLNSQNTPQPSSPGSMNFSHFSRPAAMVKANLHSLVGINCGPPPS

TRVKQQHNSPVVRPIMEACTSTGSSIAESTTAGQGPSGSYQEVKTQPAVLEREQNNGIEESRWPSAPCPSPTKDSDCVVS

EGNFRSTPPDQETCRLSGVTNGAVLASSDKGASHGTQHPDVQEPTITSSSGGYGTSIEPLQKDGEDESVDTKHKPITTGR

GSTTKRSRAAEVHNQSERRRRDRINEKMRALQELIPNSNKTDKASMLDEAIDYLKILQLQLQMMSIRTGMTLPPMVMPPG

LQHMQMPQMPQVAAMPSMGMVQMGLGMGMMEQAAQRRTRMPMQSHTGPPLNVCLANTSSMVDVHDLRYQQPGVLGYNTSM

SRQHQPMQTTKGLDLDKCNAYMLQQHQLLFQQQQLQQHQLHQHQQHLQHHQRAPNMGGGPLQ

>Pp068G00970

VRARRGQATDSHSLAERVRREKISERMKYLQDLVPGCRKVTGKAVMLDEIINYVQSLQRQVE

>Pp069G00200

MSLCVPEWDRNNDMLDVVIPSNDFHGPVNWKGEDVRSMKIHFSQLALAPVPDQDTLGVGWNDNEHVGMHGQGCKGNKSQW

DVNYSSGWNSGQETEDGVTPCKPDTPKTGASPEAVANETATEPHPPHNIDVDSDEMVSWLQYPLDDTLERNYCSDFFGEL

PDANTQLLKESFAHGSTKISPRISFSGASGLDGGSNRGTTADAAMLLGAGRAAGFFPQAGVEAFSKVRTIHSPQQHSSLP

RWPQPHNPNSPNGSSECATSYLTSAKVSTSTSASPSSPMLPPRMLLGSPIPNSSNTPQASRPGSMNFSHFSRPAAMVKAN

LHSLAGMNCTPPPCGRFKQQQGLPGCRPIIEACTSTESSIAESTTTGQGPLVSHQEVKTQPAVLEREQINNVEESRWPSA

PGLSPTKDRDCVVSGGDCKSLTPDQETCRLSGVTNGAVLASSEKGASHCTQHLDIQEPTITSSSGRYATSAEPPKEPVTG

TKRKSSEREEPECQSEDMEDESVDTKQKPATTGRVSTTKRSRAAEVHNQSERRRRDRINEKMRALQELIPNSNKTDKASM

LDEAIEYLKMLQLQLQMMSIRTGMTLPPMVMPPGLQHMQMPQMGAIPSMGMVQMGLGMGMMDMAAQGRAVMSMQSHAGPS

LNGNMASTSSMIDPHDLRYQQPGDMDFNTYTAHQHQSMPMTQALNTDKYNAYMLQQHQFILQQQQQHLHQQPQQHHQQAP

NMSGVPPH

>Pp069G00510

MSLKKHARPRETATCGSHPAECELSSLSSTEGAAGRSYHNREQENVAAVHHCKDSNVSGTLTTAQSRDRRRFRYDPPRPG

WPATSDVSPTSQGVGFGGVWAMSEASSGPSTVRHPWAGGVRLRVNGVLPGHSGKKVKKKEQPPKQGFIHVRARRGQATDG

HSLAERARREKISNRMKFLQALVPGCSEVTGKAVMLEEIINYVKSLQRQIEFLSMKLAAVDPRVDTNVEGLLKMEAEHWT

GKNARCSAVQSLHTMQYQVEEEIALLTSDPSELYKLKSSNITSAKERDRDSAPVRFYPPSSIRLRQQLIGCNGCHIANHA

IGDGGMNSVLTTVLPKLGRRQNVTISLYLQK

>Pp069G00550

RGKKVKNKEQPPKQGFIHVRARRGQATNSHSLAERARREKISNRMKFLQALVPGCSEVTGKAVMLEEIINYVKSLQRQIE

FLSMKLAAVDPRLDTNVEGLLKMEVCAVRLVSVPMGVQ

>Pp069G00530

VRARRGQATDSHSLAERVRREKISERMKYLQDLVPGCRKVTGKAVMLDEIINYVQSLQRQVE

>Pp071G00950

MNHLRPKRKRSRAPKQGDEVESQRMTHIAVERNRRKQMNEHLAALRALMPGSYVQKGDQASIVGGAIEFVKELEHLLHCL

QAQKRRRAYNDISTAVIPTSSRIAMPSLDQLQLPAPPIPLLAPASSSLLGMNEIVGEAKSDMASVEVKMVGSDQAMVKIM

APRRSGQLLRTVVALESLALTVMHTNITTVHHTVLYSFHVQISLHCRLNVDEVAAALHQTFSSLHSLQF

>Pp072G00010

MLSKHKLSVEDSTTRDIEEGEGVSVPGMDSWLLVTLARRSSVDEVFMYCGELPLSRSVQKTDANWRITAPVIAWILGEYS

LRAGSRRIRSSGLKIISSGYSVEHVRAGSPRVGKDYDLVKSATDSWLMMIAQMHSNDSVFVVSEAVRYTGIDSRLEGLGM

VCAGMLTIFIAAKVLVYAFFVEELHVVVICCAQSQIRLKPKLAVFELLSSEDADILEANKVVFSEPKSLVRTYCQEAREA

DKVPPTVSINWKENTDPDFHTSYMAHNPRKLDMWNSYQVTGSAKSEEFDKCNGMPRTLSAGLQAAWEQLERTGLHDPLSV

VHYERLDAIGTGTASRHDSGQVGEIQSGYNGIYFDEIPEALPSHHSDFTSINRNPSVTASQLNGASPDLDTEMNSEPEKK

RGRRKFPEGWVASKNLISERKRREKLQKSLLDLRALVPKITKMDKVSILSDAIEHVQDLKQKVEMLENLSTTVEDGSIDQ

ATAECSKSSGSNLEVSEADDEGHNQYHASEDASCSARCDYQSNSSSQDWAMHQVSHTFLAQLDVTKLEHGLYKLNFTCKQ

QPGVLVQLSQAIEAFVIEIVHTNIVVITPTKVTCSFVVKGDMIETFIKIGGQELGIGMAIGNGAKIGKWDWEVEEEI

>Pp072G00020

MCNLRFSLFHCRLEANKVVFSEPKSLVRTYCQEAREADKVPPTVSINWKENTDPDFHTSYMAHNPRKLDMWNSYQVTGSA

KSEEFDKCNGMPRTLSAGLQAAWEQLERTGLHDPLSVVHYERLDAIGTGTASRHDSGQVGEIQSGYNGIYFDEIPEALPS

HHSDFTSINRNPSVTASQLNGASPDLDTEMNSEPEKKRGRRKFPEGWVASKNLISERKRREKLQKSLLDLRALVPKITKM

DKVSILSDAIEHVQDLKQKVEMLENLSTTVEDGSIDQATAECSKSSGSNLEVSEADDEGHNQYHASEDASCSARCDYQSN

SSSQDWAMHQVSHTFLAQLDVTKLEHGLYKLNFTCKQQPGVLVQLSQAIEAFVIEIVHTNIVVITPTKVTCSFVVKMSSW

NVMAVTDVEADIKQLLGQSGLLFS

>Pp074G00230

MMEMGTPNYWDAADPLMVEAFIGGYEIPGYETQDDLASTLGQDLEQNDSVLQRRLHRLVEESSEDWTYGIFWQLSLSPSG

ESMLGWGDGYYKGPKDSDQFEPRKTQTEEHQLQRKKVLRELQALVSCPDDDGTEDVSDTEWFYLVSMCHSFAKGVGTPGQ

ALAFGEYVWLEEADKASYKICTRANLAKMAGIQTILCVPIMNGVVELGSTDAIHERLDVVEYVKMVFQEPTWGLTNMSPI

ISQSQVGKFDTTFMPHYPSIPFDSTSVSGVSSMTLNTDPGLADSESMDFGTRHSHMGKMVSHSGAFGFNGYDHVWGQTNE

FHYNDPLPDDNVERDLGQPMCNILGSLPLQDEKLPLASSPPPKTLDSDSRYSIFQQNNVKKPPQLDHTQTSLPVTERLHP

KPHTSQAFLHHNGSFDVGEMFNPPGHTQTVRSNPPSLDEQLHSPSMPAVEKLPIVEKPTSIYKPESVEKPMPVFKPLPQP

PSPPASKPAVPVPANGLLLAGHLDQECVDTELITMKNNVVEAPKVPRKRGRKPANDREEPLNHVQAERQRREKLNKRFYA

LRAVVPNVSKMDKASLLGDAIAHINHLQEKLQDAEMRIKDLQRVASSKHEQDQEVLAIGTLKDAIQLKPEGNGTSPVFGT

FSGGKRFSIAVDIVGEEAMIRISCLREAYSVVNMMMTLQELRLDIQHSNTSTTSDDILHIVIAKMKPTLKFTEEQLIALL

ERSCQNTGYLRKREGSDRLLQRPDNSPQLQ

>Pp074G00250

METQVPSFWDAGDSAMIEAFMGPAYGIPSSYEVQDDLASTTEKGLELSETVLLRRLHTLVEETSSNWTYGIFWQLSRSPS

GELMLGWGDGYFKGPKENEISEKRIDQGGSEEDQQLRRKVLRELQSLVSNTEEDVSDYVTDTEWFYLVSMSHSFAYGVGT

PGQALATESPVWLTEANKAPNHICTRAHLAKMAGIQTIVCVPTRTGVVELGSTDLISQNMDVVHHIKMVFDEPFWGANRS

QVMAQSLLMDSDATFFPPSPSIMSMGTTSAFASSPSVASRGSTLGKDHESHYRGRNVSVEKIGSSMASTSFDTLDYMWQQ

SDEMQFNDGVSVGTTEKDQGQSRLYYPVLGPPVLVEKLPFSATSLISRTRAAEVKHSSMLQNVEKLASEDQKPSSLPHIK

VHTTHSYPEKTGAGELSQVLSTPDLRQSIEMKLPAQVETRRAPGITGGATKPVAEKAKPVPKPPQQQQTAISGPPASASG

RSSFDQSEHDSFQESEAEISFKESSAVEFSLNVGTKPPRKRGRKPANDREEPLSHVQAERQRREKLNQRFYALRAVVPNV

SKMDKASLLGDAIAYINELTSKLQSAEAQIKDLKGHVVGSSDKSQESLSIARGSMDNSTIDGLSIRPQGSVNSTSISGNA

PSGTKPTIAVHILGQEAMIRINCLKDSVALLQMMMALQELRLEVRHSNTSTTQDMVLHIVIVKIEPTEHYTQEQLCAILE

RSCQPYSCSTKDEGHGLSEKLGSSRRSQ

>Pp077G00810

MNYSQDASVSVSAHEPAQYLQKQLESLLQHLGWSYIALWTFNPQTRNLGWKGGHFRVNSTAVNSHGDTEVWSKELFNSTY

NSSTFPRGRGSVGRAFDERRNVWLTRPSVVQSAGSKELSQFLSHARIETAMFIYWTDGILEIGTCERMYSICKEGLLNTL

WPHQIDNVKLFDNRYRGASSHSREVQLQDANTITAASQHELRVVDDDVINFNFEEEAQWQNLYHPPLDTTNLNQMSQLPS

TSFEDVNMFNAPTVRNEPQLPRQSTTSSISAQSQGGQFKSWQHTRILAPMIQTDRTNQQLLRRCIDLMKTIPNLQQGAER

ERSPVMNNTDSQERRVTSSSSPEAGLPDISIAPDPFDQYVKSWLAQESQIIVSVTQVAPSRGGHRPARGGSRIATMGPIH

AGHDEAAMNHMMAERRRRVKQKENFSALRKLVPIISKADKASILGDAIVYLKDLQRQIEELKESTAETERRYEDLKISYQ

SLEQRNKELELLAGGANMRPARECTLELLSIPTVGLKKEILQLFNIERVNAKHGIEHGSPALKWKATLSMPCSTYDL

>Pp077G00880

MDPASQRHRLGHRSDSHASASSRRGASSHSREVQLQDANTITAASQHELRVVDDDVINFNFEEEAQWQNLYHPPLDTTNL

NQMSQLPSTSFEDVNMFNAPTVRNEPQLPRQSTTSSISAQSQGGQFKSWQHTRILAPMIQTDRTNQQLLRRCIDLMKTIP

NLQQGAERERSPVMNNTDSQERRVTSSSSPEAGLPDISIAPDPFDQVAPSRGGHRPARGGSRIATMGPIHAGHDEAAMNH

MMAERRRRVKQKENFSALRKLVPIISKADKASILGDAIVYLKDLQRQIEELKESTAETERRYEDLKISYQSLEQRNKELE

LLAGGANMRPARLHSMQSF

>Pp083G00740

GLVRHMSLPSTNGGPNSPGLEDNHYGAVPMRTRAKRGCATHPRSIAERVRRTKISERMKRLQDLVPNMDKQTNTSDMLDE

TVEYVKSLQRKVQELSDTVARLKADASQRAKNSNNNNSSN

>Pp084G00090

MNRLVPEWDRSSGMLDDLIPSSGFHGPANGKLDFLRSKSTRFFQVPDQDTLEAGRIDNGHIVMQGQTSKCGKPHWQVNYT

SAWNSEQGTEDGVTPGKPVSPKSDAALEDVVNETATEAHPTHQIDAEHDEMVSWLQYPLDDTLERDYCSDFFGELPDSHI

QLLRESFGQGAAKTLRTPYSGAPGNDGFVNRASTADTAMMLGAGRAAGLLPQAGVEAFSKVRTIHSLQPSSVTRCQQPHP

SSSNGSTVCATANVTTTRAPTSVPTSATPSNPMLPPKTQPAAPTVNTQPSPPNNRPGSMNFSHFSRPAAIIKANLHSLAG

INAAPPPNARFKQQHSHLVKPTVEACTSTGSSIAESTSAGNGQSGLHKDAKVQPLIMEREQSNGTEDSRWQVAPGLSPKK

DRDCVVSGDCKNSPADQETCRPSVVISDAVPASSEKGMSRITHHPDIQEPTITSSSGGYGTSTDRLKEAATSNKRKSNER

EETECQSEDGEDESVDTKKPVTGRGSTAKRSRAAEVHNQSERRRRDRINEKMRALQELIPNSNKTDKASMLDEAIEYLKM

LQLQLQMMSIRTGMTLPPMVMPASLQQHMQMPQIAAMPSMGMGMGMVPMNLGHGMMMDMGVAAQGRAMMPLQSHVGPSLN

GAIASASSMADVHDPRYQTSGVMDPYNAYMTRQHQPMQMTQAVSIDKYNAYMLQQCQLQQHLRQHQQQPQHLQHQQHQQA

PNMNGGPPH

>Pp084G00150

GEDESVDTKKPVTGRGSTAKRSRAAEVHNQSERRRRDRINEKMRALQELIPNSNKTDKASMLDEAIEYLKMLQLQLQVCA

V

>Pp089G00750

MDQRRTSAAQPALPSSLLLSEDLYDLLQGNRFSTTTSTTSASPTTSTLSSGAATTLSDVKSNPNSLQDPLNHFIFDQIAP

APGALKMEPEIGDFMSNSYNWMPTTQLYNVDSLSCLTNYNTNHLPHQLPSTSQFSYSPPQPSVQSSSQIQALRDFMVYGN

LADPIMPTPQFSSAVSSRATTPIGNSLLYQLDDVGTDSLIGISPGVQSKPQDMNLAYRALCSEIYATERSPHRSQFQREN

HILAERQRREEMNEKFSALRAMIPKATKKDKASIVGDTIDYVLELEKRLKHLQACKDTASGSPFIRSLKRKSPSTSANTA

SVHQDSPTDAVTKDCDAPDHRGTNPATTTTSSPSSTSPSREGHSAVNSPSDQVTQESKLQAGKKAAAAEVEVQSLGSRAV

IKIVVERRPGHVLSVLNALEECKVEVMQSNVMTVGESSIHFVTVQLEEGASASTEELVSAILQAINPPKAKGDPPPTC

>Pp090G00170

MAFVSTAGQKQLGPERGTGSPSSLRCNRYRETAPGDFAAQQQRQFHAVQNRHSMAHQQDERSQLHHFTLNEDTKPSTHDF

LSVYDKGSQLAEHHLSSGLSSLLTTHDFLQPLERSGGRSASGDARALREHQARAETTTSRNSTPASAERNFTTASPNYPN

IQVPYVMAPNSAIRMVDEAEVMRQARGYPTASFSSRDGQSGGPESFRGVEVAADPETANARENHAYWSGVRLKFRMPSDL

GFPAGSSAVKSAVVNGHQSISAMTKQWTQGTIEVPKRPARSQSEEDEDDEEELSEWPKGNRKETISYKGDPVQRMEAKNG

EIRGSTPRSKHSATEQRRRSKINDRFQMLRNLVPHSDQKRDKASFLLEVIEYVQVLQEKVQKYETAEQGRHQERLKSMVW

DLCKRGAAGSADEASHINALVFGKSFSSGVGPLDINPVTEADGGVRAKHPSEQAAPTCEVYSTSGPGIHQISNQPACSIH

SCISSFDGEKEHATALTAQPFAVGVPLQQALPLPYALPIIPSRLQSDLDGGRSILGSNSPAKEKMSDLDPVSSRENECAH

ATTPDHQFEVRPRISPPSQADQISGQQSKAEHQIAAGTVVGMIDGAQAGQPSNASASRQPQESAQCKDKAVKPSRHSPEH

RSDQTTPEDGSAASPGPVQPPRIQGGVINVSSVYSQGLLDTLTRALQSSGIDLSQANISVQIDLGVNARAAAADTSAKVA

TTPESSDQSQGRTRPAPVAAEHEPAMKRPKVEKDV

>Pp090G00180

EIRGSTPRSKHSATEQRRRSKINDRFQMLRNLVPHSDQKRDKASFLLEVIEYVQVLQEKVQKYETAEQGRHQERLKSMCK

DKAVKPSRHSPEHRSDQTTPEDGSAASPGPVQPPRIQGGVINVSSVYSQGLLDTLTRALQSSGIDLSQANISVQIDL

>Pp090G00870

MERGKRLHVKHLDPDTMDTRTGRTDTFPMPVTNIDVYNRSGQVIEAGSKSYAPSTDKITMQSEVLFQPAQRFDSRTDSEF

LRKLKENPRAELKRGSSERFHLSNAEMDDVKRKYGWNDTKDSLELIYPDQYHPVMNDKKPSSIKESQDLGIQDWLPECGE

LPDRKLSRPYSMARKYDDASAARSSMHGSQYLAQLAEAHKFSLEPASESNNLAEWVDIPDVKQKLLRSESEPSYALVSPS

EKQTQSCDMEQQLRRAGSISSEEFVEFEVPPGRGRRHNNNTSGLASKNLVSERKRRKKLNDGLYTLRSLVPKISKMDKAS

IVGDSIVYVKELQQQIQSMESEIAEMEENLLSSTGVAAECSGGSRDSTSLESKEPAAGSSSSCEKGTEEAMLGVELNDNT

VLAASEKTSSSADSQGPSQEHSPVVQRKIINMEVAKMEDKTYQLRATCQKGPGILVQLTRALESLDVDILTAHHTSFQEN

MLDTFIVEMRSLDVKEAEHVRKALLDAVAQHGLAVQV

>Pp090G00890

ASKNLVSERKRRKKLNDGLYTLRSLVPKISKMDKASIVGDSIVYVKELQQQIQSMESEIAEMEENLLSSTGVAAECSGGS

RDSTSLESKEPAAGSSSSCEKGTEEAMLGVAKMEDKTYQLRATCQKGPGILVQLTRALESLDVDILTAHHTSFQENMLDT

FIVEV

>Pp093G00920

MMQLAVICKGLSATLCGCPGSPMTPFGTFAGNKDGVSRGMRRRGESEPPRSYFKAGPGEYRCLFKPGMMCKSEEDETGYY

ARRKCYWVREDNIRRTKRRARADQGDVSEAALVIPVAKKKKTRPSEEVFFKSTSMCPQTTLSAREAGRECLRHGSEVNNL

EASFDVVKPSLSDYPFSVHLYTKDCEREFQERARSHANDLERSPSILPKTPTSPEILDHEINVSSQLLSSRPSRPKDNMH

RYNSQNNMTYDDSRYFNGTQESTFEVESEAPEAAGRNMMLNFGGTMDRLDPSASFPGLGELPSNGMEPAVEQNLTGTFTG

DLNSTAYEVGDDHNHLLQEMLQQHSQTQQAQLSDANSQMYKSQTLKANLCQVPATASHGAWEDSLNVLQEMVHLQQQQQV

QQIYNQAQTEAHEFNQSYSADRFRNAPYGSGAKFPGESDLLNLFQFPRSTSTSLIPSFGYTTGIRGKGPWYSTSTISGPV

NTDNRSIGVDPLLAHHSQNSSQGFFHNLSRGNTPEASRHGGPATLVDLDQEREVLSGKNIVYGSKRELGAASAKGEPRGV

NHFATERQRREYLNEKYQTLRSLVPNPTKADRASIVADAIEYVKELKRTVQELQLLVQEKRRAAGDSSGAKRRRSLDATD

TYPGACTPENASNGHLVMQKGNDTFSADGSQLRSSWLQRTSQNGTHVDVRIVHDEVTIKVNQRRGKTCLVFDVISVLQEL

QLDLLQASGATIGEHDVFLFNTKASQMMILNVNTMKCYDLGTYSISQHRTLTSYYLSPKLICRVRKLRRDAHYFSRQTLR

NFSLSLAVSGPRALALALAFALALAPYQNGHETSSRRHLPQFPGSPPKATHHCGKRQNAESPAQEQNQCQRSINIRLQST

ANSRLSVTLNCDLAASLGICVALEGGSNTGDSSNGAGVFIPCERASEQFTGPKVAVDLSYVRLYRNNYHFMEKLKVFSEA

LEAARTAGENHLTNHPERMRSMSEACTIHHSLKINDMLPRCRNRQSFTPGAASPAIQEFLNRGYPRRYISHYSEYPHLPI

QTYFSATPCFILSLPSLGMIFDMIPTTCGDAISICVTEMKASCNQAGLQILAWTVFCQSAEQCAGGGIRMKSQIAAETVT

EVGRTIKGGDSAFIDGPQTHPRQNKSNVEQLDS

>Pp093G00950

MKERNEFYCEREFQERARSHANDLERSPSILPKTPTSPEILDHEINVSSQLLSSRPSRPKDNMHRYNSQNNMTYDDSRYF

NGTQESTFEVESEAPEAAGRNMMLNFGGTMDRLDPSASFPGLGELPSNGMEPAVEQNLTGTFTGDLNSTAYEVGDDHNHL

LQEMLQQHSQTQQAQLSDANSQVPATASHGAWEDSLNVLQEMVHLQQQQQVQQIYNQAQTEAHEFNQSYSADRFRNAPYG

SGAKFPGESDLLNLFQFPRSTSTSLIPSFGYTTGIRGKGPWYSTSTISGPVNTDNRSIGVDPLLAHHSQNSSQGFFHNLS

RGNTPEASRHGGPATLVDLDQEREVLSGKNIVYGSKRELGAASAKGEPRGVNHFATERQRREYLNEKYQTLRSLVPNPTK

ADRASIVADAIEYVKELKRTVQELQLLVQEKRRAAGDSSGAKRRRSLDATDTYPGACTPENASNGHLVMQKGNDTFSADG

SQLRSSWLQRTSQNGTHVDVRIVHDEVTIKVNQRRGKTCLVFDVISVLQELQLDLLQASGATIGEHDVFLFNTKLQALES

ALHLRVGLIPATHLTALEFSYHVSERASSLQVRRLPADGENKTGSDEDRGDKKKLKVFSEALEAARTAGENHLTNHPERM

RSMSEACTIHHSLKINDMLPRCRNRQSFTRVRQFSEFHKMSMMLLTSSGCC

>Pp098G01050

MPLWEDFSTGACFHKLQSFVIFSTVYSLFELEVGLPSARSECASDQIHLGFVGCSICSDERAFSVPSCAIISSLRRVVQL

TEYDIENFSYFSRSLVGRASSLSLQMSLTFRADECNELQSMENWLNAGTSWSEHHGIVDVETTASAPLFNHIAEASSTRG

SPESLFDALRKGKEEIPTHTTFLKLDTVELCKDNPDAVFMKAGSSSMVGSSSNEEEVSVLTNILPSFGVSVSTMADRTPS

LDEPVLCVDAKSPREPLSPGALQLRKSPSRASTKRSFVDASEASWMPVADAQQSRTQHPVKQKQPKNGMLSQEEIQSPPA

AIAYGPPFVGMSTSGEMATLAIPNPTAFLPGSAANVAPLLFSTPLTLPNVPSMEEIVQLRPKRRNVRISKDPQSVAARHR

RERISDRVRVLQHFVPGGTKMDTASMLDEAIHYVKFLQQQLQTLERIGNMSDPRFMTQPGGPMILPQVGASTSMRPFDLN

CAVYQSNPATTMTLPPSHQQPLPWPNTRVSNSFCSSFLDSAQEQFCY

>Pp105G00690

MESTPHRGGVPDPNDVNLLSRSFSMDFASMLESTDGVLHQQIDEYLQGPADHHQFNHDDGTGFSIPYSEPLSFAAASPPF

QSIMSVETLQGSNVMKQSLGRNSHHCGPEIQSKIQYVQMLPISIGAANLCAPHQQIHDMGFLEPSSFERELQAPAQPDMF

YHCSSEPGAVIPGEVSGYEQSCSPDNFRGTTCESIGPQSELGAGNLFQKDQVVKGKRPTDAVGHIIRERQRRDDMTNKFL

LLESILPPAPKRDRATVIKDSIQYVKNLRHRVKNLHQKRSQMRSKLTNVSFLSPTAIMQKKNEKKLLTPTNSQALLQTSV

ASDDIVSCPIHSDEMGKTTDIEKVKVHVDLPHQVVIEMTCRQQPRVQIRLLKTLESMGLDVSRCSVSKIRSHLLFSIIVK

CRRIVDRELDMEAISLDVAE

>Pp107G00940

MPSQSQARQVGLQSTYPTQLNFNALPPSTARAHCRSNAPPSATVVDMESWFPASTGASEFVTRNCGLNSCQQRTEMPVGA

VSDSTGISTGCVFSQLVDFQSDTWWPPITVQAALPTDILTQRNEGQIGCDTKTGALNNSASFSTGNEMESWRRETVNVDV

SDLGLVSEHPEPAMAVVSVAPILVSLSCTKSAGRKESQRLEKGESVSAPTSPSELRLVKEKTNQLGVKKSKGSGKRPVSQ

RENHIWSERERRKGMNCLFTRLRNLLPHPTSKTDKSTVIGEIIKYIQSLQVKLEMLTKKRQQVMAAVLARPGMFVSNNSG

LTLVDHSNFDPSSMTAITALPPPGKESCLQSYLGTNVGLHVCGLNVFITTSSPRGRQGLLQQLLVTIHKHQLDVINATIS

TSSTSVFHCLHCQASQNAEVLNNDLHSALQSIITNFGLPQY

>Pp107G01040

MPSQSQARQVGLQSTYPTQLNFNALPPSTARAHCRSNAPPSATVVDMESWFPASTGASEFVTRNCGLNSCQQRTEMPVGA

VSDSTGISTGCVFSQLVDFQSDTWWPPITVQAALPTDILTQRSMATPLSTNCLFDRYLVAAKLYAYEGQIGCDTKTGALN

NSASFSTGNEMESWRRETVNVDVSDLGLVSEHPEPAMAVVSVAPILVSLSCTKSAGRKESQRLEKGESVSAPTSPSELRL

VKEKTNQLGVKKSKGSGKRPVSQRENHIWSERERRKGMNCLFTRLRNLLPHPTSKTDKSTVIGEIIKYIQSLQVKLEMLT

KKRQQVMAAVLARPGMFVSNNSGLTLVDHSNFDPSSMTAITALPPPGKESCLQSYLGTNVGLHVCGLNVFITTSSPRGRQ

GLLQQLLVTIHKHQLDVINATISTSSTSVFHCLHCQASQNAEVLNNDLHSALQSIITNFGLPQCLCHKHPIKHRSTEDFE

ALLNKLENVHICSAIEDGGHWRCKLVQLKAWNKRWDTDPMSRREKAMHEAHF

>Pp123G00020

MANPEYRGRVAGEENCHLMPARKLQKADRERLRRDHLNEQFAKLAGVLDPIRPKNDKGTILSEGILALKELRAEIARLKS

EQIALRDESRDLTVERCELQEEKTLLETETERLEDLRKQNSENLSALAGWKMDHPGVLTSSQYPCPLPVSSMTKSSPPSK

DIHQDPSSDQTHHAPDSSFLPAAPFSTFLHPAYQTYGVFGSRKSPFMSYNHYSHPGHTTHVERPAARYPAPIHPTPIYPV

KGAQTKSTEPSVVATELQLQTPGLPPSPTHQQSLSGNEQREKRVACATQDRALDVLRDSGIDGTKAASCNDNFAPQLSLS

SNALALQVVQSV

>Pp124G00150

MDETKRNCNWNATKESLDQIDSLLNEIGLAPTKESQDLEFQAWLQNCQHSLDPSELTPNQSLDFPTARKYLDANATKAVI

QESQYFARFAEGHKFSHELDSASSSLAQWIYNSGVKQRLPPSSSDPSSASASHSEVKTTTGSEMGLLRWTGSLSSEDFVE

PEAAQGRGKHQMKSVGLASKNLVSERKRRKKLNDGLYSLRSLVPKISKMDKASIIGDSIVYVQELQQQIQTIEKEIAEIE

EKVSSANCVAEEDSGGSGGSGSTESKEHAAGRGTSLEQVVEVVKPVIELNNTVMAASSSLVDPQDPSPGHSPTVEIQILN

MEVAKLEEQTYQLKTTCQKGLGILVQLTRALESLDVDILTAHHIAFQDNMHDTFIVETRDCSTKKAEHVRKALMDAVAQH

GLTVLASKLLD

>Pp130G00060

MMQYYIDESNIPHTYDLAPPSTATGLLQEFVKTGMEVVDQDSYFSAMTGNLAGLLGNFPAAAAPAMSAEENRQNFALLSG

ITTSASIGEDAASDTVGYGKIVRSEEPYHPWTLTGSMNSSAQPPCAHTGFQDMPEKLNIFIELEQYLEQFRHVEEAGPGQ

GLEELEKRIIHEWSSKQLPLPITTQFPWEIIQEGLQRGDELLEAETIDLTKQSAPMSLLSVQIDDGLDKIAFGCNQNIVA

NIPGGDLHASPLVASSLPNLNNQTISPSLSCMHGTPASTEQVPGRRSSRSAFQVYAPAQSTRLSQHSMSSNRKPMIHYWV

EKFGPKLKSLQVLQPSSSDQSAVGLGNCNVRVGTSYDFSTSISAAHDFRQALVRKKAEQQRRNKFKRSLESLKKIVPTIT

KKDKVAVIYSAIEYIRQLESRISSLKKELEEAAPGFLDMENSATLFARGGVNVSGEQDLGLHYIENPAGPSSSPATTHVV

VEGTEGGETLNLRVEANYHARSLPKLLNVLQDLDVVVVAMDYGCKNERFRATVRVKTSRNVRRSNHELEQTLCRALEVGG

LMPPSRSQQLF

>Pp130G00090

NHMLAERRRRVKQKENFAALRRLVPIISKADKASTLVDAITYLKDLQKEVEELKASKENIEQRYETLDKRCKELEDRNRQ

LVATLSKDQSNSFNNLNSMKSI

>Pp136G00150

MARLRQNSVRRNYWGRSSLAEHINLQSIALADVYGMLSHRQREPSDFIQTLFRDWTFFACQAVGVVSSCLPEFPVVANKM

RNKEEHFECYIMAASSNSWLHQNLLPQHNTQQQGSPHSLFHHSEFPLHVDSTQPGPLQHMNLGHGGHGIDSHPPLHHGMH

STLHMSVELQQQQQQLQHPHPPVQVGSGTSGRAVGPTENGDTASVRKVHKADREKLRRDRLNEQFGELAGVLDPDRPKND

KATILGDSVQVVKDLRSEVKRLKCEQTSLLDESRDLQQEKTELREEKAALKTETEQLQNQLQQRLQGMLPWISMDPTAAL

MGAPPYPYPHPHPMPVPQPPSSAVPSSTPPASEPAPAVPTSQPPVVGPSPFLSLAPPGVYMHPYAIYGARPPERGHPYMP

YPPYSASVGNHSHVERPYAQYPSPMSTMPGYMVQMQPPPPPYAVPGMTLVPPGQQPRGPPHSTYPPTSGPETASLEPQGV

ETSLQLQTPGGSSRPPSASEQPPSQRGEDSNKIPEVADLELTPPGTKEAGDQAVSKPDEMNGGDAISSLTLASKSLQSRT

DDAPLPSRKKLTVATFIIESHFEANWTLLLASRHWSLGGNAAAFDYLYLDAALTSYTLGRLATERTELGGKQPVLTALIL

RRGIL

>Pp140G00150

MGNYQKLSADSASPSAYETISASMHPPKVSSCEGMVWLEGWVSQASTNGEGSSSSTSAFKLPEVVPTLPNTRLSFQDSGL

LSNHWIPSFNTFSQHIPDIAAETLSLEFMQERLETLPEASFEELCMQHNSKPLYLNSIPSPVPNNHPHFDKSRREDFLHY

AKTYAPFHAWQGLRVDRSPSSPLAFHDTLTNSSGDEDTDGAQLRSTWLGKTSSTTIQLLRASASENAWNNGQIKYNNSQK

LAKTDLLSNSNYSLSVVSNQSHGNQSEILGSNRKGKAISRSEDFAFVRGHEPTASSHYEKCMDPNGSFLPMSPIAQVLKY

ATAGPSFVSNSQLQQSKTNFSNETNSTRGLTGNDLDLNFVQKAQERLLAFQPASTASGYSARVEQSRKNSYRFAMDRSSM

SPSRRPNILPQRCPIFSQGNQEAGLAWSPYDTTQSRTTKSKLQCRHLLGTPSQAMDIIAVGPALNTNGRPRAKRGSATDP

QSVYARHRREKINERLKTLQRLVPNGEQVDIVTMLEEAIHFVKFLEFQLELLRSDDRWMFADPFIYNGMDITGSYPHVPS

GLERLNLRG

>Pp145G00290

DSKPKRCKGENDESVKAKAERSCSENSGDSGSPRALKDSNNRNKILSKQDYIHVRARRGQATDSHSLAERVRREKISERM

KYLQDLVPGCNKVTGKAVMLDEIINYVQSLQRQVESLSMKLASVNPGPSTARLDYNFETALNKDMLQ

>Pp147G00560

MSEVLEGEGGGCNARKREWRFSCFEARERSLGRCNRLQQSRGCHQTMNHCVPEWDRSDDMLDALIPSDDFHGPVYGKAEF

VRSRKSHFCQVPVQNTLEAGGNDNGSVKMQGQSSKCNKPQWQANYPTTWSSGQGTEDGVTPGKPVTSKSDALLEAAVNEA

PTEVHPGHHIDVAHDEMVSWLQYPLDDTLERNYCSDFFGELPDSHTQLLRESFGHGSTKTARTSYLGSPGNDSVMNRAST

TDTAMLLGAGRAAGFLPQAGAEAFSKVRTIHSLQPSSVTKWQQPHPNSSNGSNMCATANLATTRAPPSAPPSNPMLPPRT

QPMIPNVNTQTPQPNNKPGSMNFSHFSRPAALMKANLHSLTGMNSVPPPSARFKQQQNQTGKPTVEACTSTGSSIAESTT

AGQRSSGTQQELKTQPGVTEREQSNGIKDCRWQSAPSLSPKKDRDYVVSEGDCKKTFNDQETYRISGVTSDAVLASSEKG

VSHVTQHPDIQEPTITSSSGGCGTSAERPKGFATSNKRKSSEREDTECQSEDGEDESIDTKKPVTGRGSTAKRSRAAEVH

NQSERRRRDRINEKMRALQELIPNSNKTDKASMLEEAIEYLKMLQLQLQMISMRTGMTLPPMVVPGGLQQHMQMPQMPGM

PSMGMGMGMVPMGLSHGMMMDMGVTAQGRGVVPMQSHAGPSLNGSMASASSMVDVHDPRYLASGVIDPYNAYLARQHQPM

QMTQPLNIDKYNAYLMQQHQLQQQQQHHHHQQQHPQHQQHQQTSNMNGGPSH

>Pp147G00720

GEDESIDTKKPVTGRGSTAKRSRAAEVHNQSERRRRDRINEKMRALQELIPNSNKTDKASMLEEAIEYLKMLQLQLQV

>Pp147G00760

MAVKVKGGGGGGVAVMGAEAYKGEEDEDASECAAHGCKLGSWLGRKRPRTTKGEGFEDVRWCPIWDFVVAVEELKFERKT

GQRDWKCRTWNNLKFALEGHDRRSYRLERARESVFWHALLYEHWVIELESSMIFACLLPVCTSFFGPLFLPPQRAGTSEP

HHPSSTIRIQNYFSSRACFKLCEMAGPAGALWSTCDPQPIQQAEIFSGPDNQAGLMSFHVDTPFHWGSEPWALHSRSDDI

ALMSPSLVHDISPYDSVLHLSGVSGDVQDLVCGNPKFRQSGQWGQSEFSYSVQDNMQDLLTNQFIPYNTSSLGLNHLSPN

FTDLDCAPVYNDTKAFGTVTHNRAVPSTNTQSAQHGSSSMVSSNRPITSTASPTTQYGGPRTPSQTTQYGGSSMVTNSME

MFASAAPQGIMTTSGLSGGCNSDLMHLPKRQHAHSLPPTTGRDLTASEVVSGNSISNISGVGSFNSSQKSSASVMMSPLA

ASSHMHKAAAVSEELKMASFNPGPFVPTQKKQQHEQQDTMTSNRIWADKNNLGKISSSPIPIMGFEQSQQQSMSNSSPVT

SLGFEQRQKMSMGSSPSITIIGFEQRQKQPMSSSSPISNMVFEPRQKQPMSSSSPISNIVFEQRQLPTVGSSPPISISGF

EPKKQPSLSNSPPLSNLGFEQRLQPMSNASPISNLPFEQQRQQATMSNTRSAEPDSVESTTKWPLRMDGAIGGCAGLPSS

QKAPVIMQPETGTMKCPIPRTMPSNAKACPAVQNANSVNKRPLTVDDKDQTGSMNKKSMQKFLGPQGCSRLESISALAHQ

KVSQSTTSGRALGPALNTNLKPRARQGSANDPQSIAARVRRERISERLKVLQALIPNGDKVDMVTMLEKAISYVQCLEFQ

IKMLKNDSLWPKALGPLPNTLQELLELAGPEFAGIDGKNTEESSEKPKKSALEVIELDGNQPSAD

>Pp155G00490

MGSASEALMAVLSSDHDYSSVAPVMTPSNSSCTGSSDLFSPPPVSSWLQGSLPGSEQRLGSSQRSKFRGSPLDTLPAPMP

EWPYPRDEKVLVNALQGPCRATNFRSNMTAPHDLNRPRHGGLQRYHSAPSSFLQCLDDLMNEPDVFPQASSSLVDSDVLL

DWADDLTPITERGSQQMDTEKSYPRSGFNDCEQFLSSLTNFSSITPSPQSQNVEPTGSLARQDGLGGFDLPAIQDSIEEN

SVAIYQMPGFLFADGSMSCTSSEKIGAANSTISSAYIPEDSTHLVKEPAVCGGLWNCTTSSGGSVGSSSLRGVEVTMEPP

LGWHGVSSLGGAVTVGEPAPRLEGLIRHSSLPATSRPFSSTFELDDLQADPSMVYLKTLRANRGHATHPRSIAERVRRGK

ISERMKKLQELVPNSDRQTNTADMLDDAVEYVKQLQLQVQELTNTVAELQLLQERLGHPTS

>Pp159G00160

MSKPAGGVQEHLRSIVGPKGWDYAVFWQLHDETRSLDWTGCCCSGAEAAGNDVLVASSSSRFLESSTGCPDVKGFHPDTH

ICSLLASMPSSVSLDSGIQGRIFLGGQPKWVHMDPSMEGQDMAVQTKVCIPVQSGLVELGVANHVTENAALVQYVRGSCG

EPWQSKQGSSSNTALDAASGGHGMMDQQAVKMYYSRHFPTNLENSWPSSHPWEQEDPMLESQLLGGMDQELIQLLGTNVH

FPHHAGPMADSSPRGSGLSKDDGEVKQEIRGDSSDCSDPMEDDEEKGGPRSARRHLSKNLVAERKRRKKLNERLYSLRAL

VPKITKMDRASILGDAIEYVKELQQQVKELQEELLDSKENDMGTAGLGFEEAAVAAEEANLGGAIDIGRCSGKVDSQAVT

IEVIDRKGDHELTQPMQVEVSKMDGRLFSLRIFCEKRPGVFVKLMQALDVLGLSVVHANITTFRGLVLNVFNAEVRDKEL

VGVEQMRDTLFEMASQSNHRSLALAATDPPASLQPSSMSGDRLETITEGSFP

>Pp164G00460

MVRFNYMYPVQEQLEAMTDQHTPSMDSVSSAGEKTSSCIVQQGGNASETSNLWEEWTQGSNGDDSVSTSNFLPELNSSTS

SRLAFHQSDILSTWISGYHPLSQSSLSSEFSHTSDRENHPPAFMQEGLIPSGLILDSDPALTDIYTRSSSSDSLPYPTAR

IMDKALTDHELESAVPLAYEKGCVPPQVLRNLGPLSPSSPLAFQNGLLNPLRDPWDSCPSALPWSNVTTASQTYGQVTTR

TFIPDHSASAIDKLEAVATITAGYGASKPQHTDVFIEPNGTFQSTPAGWAPQFYDGSEATGLLVKPMRAIASLGEAGCGE

ATSEFCTKTKPGLLKGGDTITSPVGSLLGDCKKAESSMKQVWPGKHRLELVELVDGEDTKSSPTQLKRPKHSTDYANVLL

SDHILKGAELRSYFHSGDVGLNASQAMDIIVIGPALNTNGKPRAKRGSATDPQSVYARHRREKINERLKNLQNLVPNGAK

VDIVTMLDEAIHYVKFLQTQVELLKSDEFWMFANPHNYNGIDISDPSSMHSPELESNI

>Pp167G00550

MTPLVPPELRLELQAATRAVKWTYSVFWKPASSNQKTLVWGDGYYNGTIKTRKTIGAKELTPEEFGLQRSQQLRDLYNSL

SDSKTGHQQASKPFALKPEDLAEQEWFFLLCMSCNFAEGVGLVGRAAADGRYAWQCKTNEISTKLFTRALLAKTIFCFPL

MDGVVEFGTTEHKNSTPSQKSQKAENRQKILKEALFRVTRLYDGASEETSASHVLAERRRREKLNDRFVALRELIPNVSK

MDKASILGVAIEYVKELQSQLRALENEDKAATSECTITEESFKPGHVNVRVSMNNDVAIVKLHCPYRQTLLVDVLQSLND

LEFDVCGVRSSISDDILSTVLEAKVLQFCRRFFAIGSCRTIFLFGVCLLIRHVLPFLLHMQLRSASDGSSPTIIEVEKTL

HRAAAGLLKERASASSLQ

>Pp167G00680

MAMAVWWRFGGLDMSTIHRIRTRIVQNEKEMSGSKLEAIKEMTPLVPPELRLELQAATRAVKWTYSVFWKPASSNQKTLV

WGDGYYNGTIKTRKTIGAKELTPEEFGLQRSQQLRDLYNSLSDSKTGHQQASKPFALKPEDLAEQEWFFLLCMSCNFAEG

VGLVGRAAADGRYAWQCKTNEISTKLFTRALLAKKSASIQTIFCFPLMDGVVEFGTTEHVSYLDLPQSHTKILLGFVKTM

AALTFILSSVKARENSANLENEKLLDNLYVCHHKFVLVDSIFTLQGSDSRLYKGENRGKNQSGVLPGRVFSSWKKNSTPS

QKSQKAENRQKILKEALFRVTRLYDGAWKNKVDSSFIFTDRAVEDRTSNLGSQKPVPSSEETSASHVLAERRRREKLNDR

FVALRELIPNVSKMDKASILGVAIEYVKELQSQLRALESNNTQDGTPRQFGTANEDATITNTTREHLECAGVVHVIDEDK

AATSECTITEESFKPGHVNVRVSMNNDVAIVKLHCPYRQTLLVDVLQSLNDLEFDVCGVRSSISDDILSTVLEAKLRSAS

DGSSPTIIEVEKTLHRAAAGLLKERASASSLQ

>Pp173G00390

MPPGGCQGSGYANPGVPAGQSLPGMGARPRVRARRGQATDPHSIAERLRRERIAERMKALQELVPNSNKTDKASMLDEII

DYVKFLQLQVKVLSMSRLGGAGALVNSDPPAEGGNNFAASAGSSGVSNPAQDGLASALTERQVTRMMEDDMGAAMQYLQS

KGLCLMPISLATAISTTNKGPAQANANTGDRQGSAAASNIGKSTAGSSLAGGSKEDGSEAGRVTESTTQDT

>Pp173G00460

MAQQPSTTMMMAMQQQQVHGGGNHHGGMGFHHPGMGGQQGGGGGGSGGGPVMDELMEHMFGMPGGGMFDMAGGRVGPWDY

NVGSGAGKGFGVGGMPSAVGLSKKGNEEVDYGLSEVQIRHHQQQSAGARGESGSGGMPVVREARNGAPDTLTRSVSLGSS

ASEESGPQQGKGDQLMGSMVAPSKHLQQPYGGAGSGVPTLPMNFAPAKAENVMLVGEMDSHNAHGKRFREDEDGRPRPTG

AMPPGGCQGSGYANPGVPAGQSLPGMGARPRVRARRGQATDPHSIAERLRRERIAERMKALQELVPNSNKTDKASMLDEI

IDYVKFLQLQVKVLSMSRLGGAGALVNSDPPAEGGNNFAASAGSSGVSNPAQDGLASALTERQVTRMMEDDMGAAMQYLQ

SKGLCLMPISLATAISTTNKGPAQANANTGDRQGSAAASNIGKSTAGSSLAGGSKEDGSEAGRVTESTTQDT

>Pp182G00030

MTEDMYRKLRNLGGSSITPATIAQQLKISLQTYCPSSLPTDTAQPNESTMNLKPTSRSDLHNPDPFWLLPNFPGDFPDQR

DVNANPVVTSNPQTSTSMWEPVVGNQEFSTQPITAPAFKLSQGDSLTNSPGENAMELPANSSDTAEKKSVGGKRQKSVAS

KNLVSERKRRKKLNEGLFQLRAVVPKISKMDKASIIGDAIAYVRELQKELEEIESEIDDLEQKCTGSIGDDPGSVEEAGT

GENFSSPTSSNLISGVEIQGAEHRVDSNIDKLSANTTQMLFPARLAQKILEVDVARLEEQTYHFRIFCPRGPGVLVQLVQ

AVESLGVQVINSHHTAFQENILNSFIAEMKDPKMETEDVRKTIFSAAAQYGLVQS

>Pp182G00040

MELPANSSDTAEKKSVGGKRQKSVASKNLVSERKRRKKLNEGLFQLRAVVPKISKMDKASIIGDAIAYVRELQKELEEIE

SEIDDLEQKCTGSIGDDPGSVEEAGTGENFSSPTSSNLISGVEIQGAEHRVDSNIDKLSANTTQMLFPARLAQKILEVDV

ARLEEQTYHFRIFCPRGPGVLVQLVQAVESLGVQVINSHHTAFQENILNSFIAEMKDPKMETEDVRKTIFSAAAQYGLVQ

S

>Pp185G00710

VQMRARAKRGCATHPRSIAERVRRTRISERMKKLQDLVPNMEKTTNTADMLDETVEYVKSLQVKVSELQETIA

>Pp194G00400

GKGKVSSEDSKSMEAATEKTTPLKRQKSDTEDVKVVKTEQGSASENSGDSISPRSTLKGATSKPPQDLPKQDYIHVRARR

GQATDSHSLAERVRREKISERMKFLQDLVPGCSKVTGKAVMLDEIINYVQSLQRQIEFLSMKLAAVNPPRLDHNYDLLSK

DM

>Pp194G00610

MALGNGSSSRSFGEGADHYMQETGEQYFSEVDSSDSENEEVPMAFATPVPRGLGKRTYGVFVDDVLAERRLKNAKLDEQL

ASLRSILPGSVLGEEKASVLMDAYQYIMKLQKSVDELTTELVPLSTTSANPNGLLFQEAQDAQSTSSNSICLLYQHPMVE

VKREEGKIEVHIACTNRPGLLVDIMSALESKRITVLHASIACRQNVLFEALSLEVRQPEILKIGHTSYVLENLSMITAAD

TANAKSDKNCQKGTLVLHSGPEPLPTEAEDHVLKEIIAHAIRNDASGNNSL

>Pp209G00050

MSTGSAQGSGYANPGGVPSGQPLPGIGARPRVRARRGQATDPHSIAERLRRERIAERMKALQELVPNSNKTDKASMLDEI

IDYVKFLQLQVKVLSMSRLGGAGALPSLVNNDLPSEGANTFAASAGSSGIPNPAQDGLALTERQVTRMMEDDMGSAMQYL

QSKGLCLMPISLATAISTTGKGSAQATANAGERLGSAAAANTGKSVTDPSSAGGSKEDGSETARVTECATQGT

>Pp209G00080

MAQQPSTTMMMAMQQQQQQHQVHGGGHHHGGMGFQHPGMGSQQQGAGGGGGPVMGEFLEHMFGIPGGGMFDMGAGGRGGS

WDYNVGSAAGKGFGVGGISSGVGLSKKGNEGVDYGLSEVQIRHHHQQQQQQGGGARGEVAMPVGREARNGAAVHGDSLTR

SVSLGSSASEDSGPQQGKSDQLMVSPSHLQQPYGGGGGGSGSGVSPLPMNFPQAKAENVLLVGGMDSHNALGKRFRDDDD

GRPRTTGVMSTGSAQGSGYANPGGVPSGQPLPGIGARPRVRARRGQATDPHSIAERLRRERIAERMKALQELVPNSNKTD

KASMLDEIIDYVKFLQLQVKVLSMSRLGGAGALPSLVNNDLPSEGANTFAASAGSSGIPNPAQDGLALTERQVTRMMEDD

MGSAMQYLQSKGLCLMPISLATAISTTGKGSAQATANAGERLGSAAAANTGKSVTDPSSAGGSKEDGSETARVTECATQG

T

>Pp211G00360

MAIGNNNNTCGGEKGMRGWPFSPVSTRAQNHFGGRGQLGQEGVTQNNNNNDDDDAHLEEVQCTWRHRLPVQSGFPVTTRT

RACWEDEVQVLQNHRVLIFGKRSEMSRHTKKGVLLRSMAFIAAVGHKQLGPDWAIGSPSSLGHNRYGEVVPEESAAQQQR

EFYVVQKRHAMTQQQAERSQPHHFARREDTKPSTHDFLSLYDKGAHQLEIGSSSGLPSSLTTHDFLQPLERSGTRSTYRD

APASREKQARVDTVQSRNSTPASAERNLPTVAPTYSNNQVPYAVNPNSMNRMVNEADALRQGRGYSMVTLSSPDDHNGGP

ESSRGVEVAMDPEVANARDNLAHWPGVSFKFDMPADLTFPTGSQAVKGAVDNGHQSIPAIAKQWTYGIVDVPKRSIRPPS

EDYEDEDEDSSERPKGNRKESISCKGDTAQRMDVKNGDMRGSTPRSKHSATEQRRRSKINDRFQMLRDLVPHSDQKRDKA

SFLLEVIEYIQVLQEKVRKYETTEQGRHQERLKSMVWDICKRGAAGSADEASQLNALIFAKDFNSGVGPVDVKPVTEANG

DVRALRASEQAALTSEVHSTAGINGQQRSSRPACSIHSCVSSLDGEKEHATAPTAQPLAVGVPLQQAIPLPYALSIVPSR

LQSDQDGGRGTPGSNSPAKEKLCDPDPVSSRENGSARANSPDQLFELRPRISPARQAEQISGQQSKAELQTPAATAVDRI

DGAQTGQPSNGSASRQSYGATYYQENVSKPSRHSPARQNGQIAPGDGSAASPCPTQPPPRIQGGVINVSSVYSQSLLDTL

TRALQSSGVDLSRANISVQIDLGTNVATAAADTSAKVATTPESSAQSHQSQGRTRPAPVAAEPAPALKRPKLENHE

>Pp213G00330

MCGKSVAVMDGAASTMIQSHRGTSQASYSSGRSAVDEEISAMLHSGNYSHISQAPLAVSVGLEHFPYSVYQGYPGNQTRP

VPTTQRNTNMPRGEIGDGNAHVGPLLHRGSPPRHSTFNLEEKYVLEAKSEALDANVLRNGATYGTIYENSRPSAMSTTVP

LLHRYHSAPASTYLTQMNEEQIGNTVMSSIFTDGGLTPITENMDVERLGCGNSNSNEFEHYLAPENDFSRRAAFSALHRE

EATPDSFATTFQNSKNAVLCQSSLSSAGLSQPNISMQDRMAENGSEKNSSRNSDDNNHLISGYAASIYSQEDCHWKSPTS

PAKRQRGLDGESETSSGSVEGSTYRKSQQGGLIRHMSLPHSTNGDSSSPGVEDNTFHTVPMRTRAKRGCATHPRSIAERV

RRTKISERMKKLQDLVPSMDKQTNTSDMLDETVEYVKSLQRQVQELSDTVVRLEAAAAQKIFSDFQIRQKYLATPNPYSV

DVTQADATAHRLSGKQQIQVGTVMSGILDHDTISHDV

>Pp226G00320

QDYIHVRARRGQATDSHSLAERVRREKISERMKYLQDLVPGCKKVTGKAVMLDEIINYVQFLQRQVE

>Pp231G00080

MDSCGYGGGWVTPGQASSGIPEFQDVVGSPCLLPDSGQTDAVAFCGTDGILLLRFVEKQKQQPRSSSVTVTVEHRTMPSH

RHELDAGGPALQLRAAALFPNYLLSCTPQAHCRSDSLNSYPLASLVDMDTWLTASGLETTTRNCGSPTALHNYQQQAEKP

MGSVDTNPGIGTGNVFPQIGGYQSDTWWPPINAQMFPRDISTQHGISGTKPGAGNSMTFTTNNYTMDSSWCHNIANTVTT

VANADMCDLNAPGEPPSAIPVASVAPCMAIVAAAGAASTRATKISRKGNQSKGGAVSAPTSPSELRPPREKSLGVQKKWN

GKRPVSQRENHIWSERQRRKGMNYLFSTLRSLLPHPTSKTDKSTVVGEIIKYIESLQVKLDMLTKKRQQVMAARTLSAFH

SIDTLPKAFVSNGLTLVDHSSDPMSMTAITALPPPGSESCLQSYLGSNVGLHVCGLNVFITTSSPRGQRGLLQQLLVTIH

KHALDVINATISTSNASIFHCLHCQASQDAELLNNDLHSALQSVITNFEPQF

>Pp257G00150

MGAASSDIRISSNSGLELEGDSSWKAPKSSRNTSNMDVAKQVVQDTMRAASFCKGVLDEEWYTPETSLMELSSYSSPYGA

QDARSNFSLLDSSLNYDNGNLMANFRPAPSSTLGIGQLESNRILSDLACTGQCSSVGLLSSISPGRHLRRSTTTDSLGSG

LPTSFSQGAVIPSGSLSNITSRNTNTESTNFPPSFSDAYNAPALDLDKTGKECAGDIRDKLEPYQTNKRMESYPPRQQAF

SQKRAASPRSMGGGTVSPTSKSPPRVVTSTSNDSSVDIRDEDSPHVQNFRGAELHSGSDDPNDIGIDGDDHNGKDDDDLD

ESGDGSGGPYEVEEGAGNGTQNNGKSKAKGKRGLPAKNLMAERRRRQKLNDRLYMLRSVVPKITKMDRASILGDAIEYLK

ELLQRINDIHNELEEAKLEQSRSMPSSPTPRSTHHGYPTAVKEECPVLPNPESQPPRVEVRKREGQALNIHMFCARRPGL

LLSTVRALDALGLDVQQAVISCFNGFALDLFRAEAKDVDVGPEEIKAVLLLTAEYGMHSLQ

>Pp270G00110

TSTSSVVSDGKNSKSLEANEQTSLKRQRSGLVKAERSASENSGDSASPRSLKATSKPPQDLSKQDYIHVRARRGQATDSH

SLAERVRREKISERMKFLQDLVPGCSKITGKAVMLDEIINYVQSLQRQIEFLSMKLAAVNPRLDYSYDLLGKDMLQSRSP

>Pp272G00090

MRARAKRGCATHPRSIAERVRRTRISERMKKLQDLVPNMEKTTNTSDMLDETVEYVKSLQMKVKELTETIAQLKAATQMS

P

>Pp273G00100

MYRTRKYSLISCWSSFANLKNKLFHYLEERVHGACTFCNIVADSSVKLGVVLTLFLCVSRAGENHLYTMLETSIGRLLNC

SYSHGALLLQSMSPQLMGSAEQQALVQSFVRLNSGCMLEHSSQQGNGNPNTFFKSTNSSSLVNSTSEMTSPDVDYVGALE

NNLHMNCISSLGGGNNVLIPGLTSMLDRNKNRDGTFMAAHRYYKSFYPMESPGGPLSSGLLPCGKSENSLNKGEARVIAI

ADLQRLLEGDDEDWMVQEEIKMHESSKLVKPQQQSEKHAVFQDSFGGCIEASSSAGFTVTCFDRVTIPSATHPPTPPFRQ

RKGSVGGSPAVDLDNFARMQAILFRQASQLIPTLEDIASSRPKRRNVRISIDTQSVAARHRRERISDRIRVLQRLVPGGT

KMDTASMLDEAIHYIKFLKQQLQTLEQLGIDGCDPGDVALRGGEALQLSSSVRPAFHMNYTSAFQSWPASVTDLGNRSKT

CPQD

>Pp280G00170

MEASTDNSAPLKRHKSDGEDVRAVKAEQASASENSGDSISPRSTLKGATSKRPQDFPKQDYIHVRARRGQATDSHSLAER

VRREKISERMKFLQDLVPGCSKVTGKAVMLDEIINYVQSLQRQVEFLSMKLATVNVPRLDYSYDLLSKEMPMQSRSPETT

LLGPDPLAAPFGDHSQCSPMQSISTPSCHFEALYLRRSMSVPTMMSRFDSAGHFSDLFSQV

>Pp300G00450

MALSTGLPLEFNDAVDYFNYTNQDYSSDLTDTENDEEYACTPRGFQPILPRSCKRTYGVFLESEVVPERRLRGRIHEQLE

LLGAVIPSSCSGEKSSILADAYEYIEKLQRQVEELNYELDMESYLGDDLCHCEDDCSCCEHNLSPSSTERTAESNAGLES

SSGSDCGCSQPTVEIVRTEEGLKIHIECDKRPGLLVEIMELLESRGLNVEQASIACVDQLVFDGISSEIEGNDAEDSRHM

THVNAEDVEASLRSLIADQRQCTSH

>Pp312G00010

MAFMSPAPDQATVFGGLVKKSMSIPSSFNTAAMSPTDSMNSLYSLEKQLVGQPRSNAALSGVPGVSGFSRSATNYWGHAT

STDMQQILGDSASVSPQEALGSGMVSSDWQQLLDAKSSAYKYPPGVSAAATPARGGIMENARIREMLTGRVGGSPVPHKS

WSPLDTVAAGIAVDQSKGGLALHGVDLATSHCLAQFTSDPAFAERAAKFSSFGNGKYPQIALSLPVNDAHCKPRSRSSES

GCKSSRTTCNPNSAKKSTATVAAGMEENTAAAPAAASVLAAPAAADRSCDMDVDGVGNNDKSPTRTASVPELEAIEGASL

VTMDRTHELAEVAAGLEPNNSSSVTEQQQASAASPARSPTGSDDSDRRKRKSSSADKLDVDSKAADVADSQPKRCKGDND

DLVKAKAERSSSENSGDSGSPRAHKENNSSKDHAKQDYIHVRARRGQATDSHSLAERVRREKISERMKYLQDLVPGCSKV

TGKAVMLDEIINYVQSLQRQVENLSMKLASVNPGPSSTRIDHNFETTMNKDMLQSQMSGSLGGSESTTTFGMMQQHQHQP

QPQGHMMNGHCGLDFRAMGTSMDGYLRRSNSAPIRVQSGITSLDSFGDDVSQSIGWDGELQSIVNQMGFSFQGRYGSSPL

DSLQCQLPVGHMKVEM

>Pp371G00200

MCLQEEVCTIENTGMIDVSESRPGRTSVNRMGDDEAPSLQVFYEKNLAVGIYIEVAQGSGTSMIERTLLLLLLLLLLHSW

ADSTNELCRRGGGCCGPQFGHCGHVASIFRAVKLLHGVSVHLKGCQRPLAWTSAPAPTFKHCLIAAPASSGGGGALQNHT

VPDAACAEEQYELIMARTLAIRGPPTVTAPAPRLWGAWARPTARAGRGPKTWIETARALVVSYTSIYMWKGWAPLASMQS

ELMAVVTNVAITGESGYGAAPDNPHATITHPKTSYDSDTRYRILSSRVHEHLNGCLADQLMDIQQGMGVEASTPQLHVED

MPQADRMSVDNDAANLHAFANEFDTGWPTSGNESALVEDLHTRRLQLNGFQSFHNQIFNECAKINFHSNSSPVSAQQLIM

MCNQQKGVQNSRSGSWFSDRIEHPNTMCRPTSDNVFTNRDHASSSSQFDKMQPLGGKIGDNVTHPNCLPSVAQKQSQAPR

LQTELVGCAPISAFYSAEPTCFQSPPQMQSIHNAVSFSLGDEIGQTVPIGANASQENGCATTKQVPNQIDETGNCNARLH

SCEDQGSLDLQRVWCQPSQAITSFQRCGSATTSCGHASGLAPDRSKPGSTRTSEDGGKSSPPVHRTPSGGKHRALTNPKK

GRKQKLPGKTTTQAFLNKAVSQRESHIWSERQRRRSMNQLYTTIRALLPHQSVKTDKATVVMDIINYIRAMQADLEVLSR

RRDQLLAALNLRRQPSQVFSAHGLTCVDHTSDASVLTAVTTLPPPGSVSCLTSFLGNNVAIHICGQHVFVTITSAPQSRP

GLLAQIISTLTNYNLDVLSATVNSRDNTTAYALSVETSQSVESLGDDLHTQLQIVINNFSPSDAKEP

>Pp371G00290

MLTLDNMGLSRGGFRSDLYSRDSFGFGLDASSLEVVEGNVALQLNDFLHNSYTMQQGGVQNQCGSYDGLIARSGSYAGLA

SLGFSQPDIDRGILFDDLPPVVDHENLHDYEPVSNALFSSTSSVELGSSHVPGLPSASLQSQVTTEQRDGGEGLTSRNGL

KRGRSLDNEIDMQPWKKLDKTSRSIGPKGKRMNEQADHILRERQRRDDMTSKFAILESLLPIGTKRDRSTIVDESIEYVK

NLHHRIKELQDRKMLLIQSAATTSKDNTAPSCRKELIMSVQPGSPSKQNEKQTVVPKPPISQEELSRIHSFLRSCLEKVE

VHADLPNQVVIEMVCKPQPRLQSNILQCLECLSLDVKQCSFTKIAHRLICVITAKPQEATAPTSGIVAALKCALQGSD

>Pp377G00250

GKGKTSTSSFASDGKDSKSMEANEQTSLKRQRSGPVKAERSASENSGDSVGPSSLKASSKSVQNLPKQDYIHVRARRGQA

TDSHSLAERVRREKISERMKFLQDLVPGCSKVTGKAVMLDEIINYVQSLQRQIEFLSMKLAAVNPRLDYGFDVLGKDLLQ

LRSP

>Pp386G00050

MDRESTSWSNLPECERELQVRIRSHASDLERNPTSLPETPTSPAILLQENNVSGQLLSSKRVRLIQNMHRYNSPNNMTYD

DSRYFNGNQESTFDVENEAPETAGRNMMLNFGGAMDRLDPSASFPGLGELPSNGMEPAVEQNLTGTFTGDLNSTAYEVGD

DHSHLLQEMLQQHPQSQQAQSTDAHSQAPATASHGSWEDSLHVLQEMVHLQQQQQQQVHQIYNQTQTEHAFNQGYNADRF

RSAAFGGSAKFPGESDLLNLFQFPRSTTTSAIPSFGYTAGIRGKGPLYSTSTISGPVNTGSIYENRGIGYDPLLAHHLQN

SSESFFHNLSRGNTPETSRHGGPNILVDLDQEREDLSGKNVASAYGSKRDHGAASGKGEPRGVNHFATERQRREYLNEKY

QTLRSLVPNPTKADRASIVADAIEYVKELKRTVQELQLLVQEKRRAAGDSSGGKRRRSMDDADNYAGSCTTENASNGHLV

MQKGNDTFSTDGSQLRSSWLQRTSQNGTHVDVRIVHDEVTIKVNQRRGKNCLVFDVIAVLQELQLDLLQASGATIGEHDV

FLFNTKV

>Pp412G00220

MAHGNGSSSHNSGEGAEQHTREIDEEYFSELDSADSENDEVPKAFETNIPRGSGKRTYGVIVNDLHAERRLKNAKLDEQL

SFLRSILPGTTPGEEKASILMDAYQYIMKLQKCVDELNTELIPLSTASANMSAGNLIVGSLQEAPDTQSTRSASVCVSYQ

HPMVEVKREEGKLEVHIACMNRPGLLVDIMGALDSRRITVVHANIACRENAQLEALRLETSSSEQLKHENYNRSFLVQLV

QVEF

>Pp519G00080

MTHIAVERNRRKQMNEHLTALRALMPGYFIQKGDQASIIGGAIEFVRELEHLLHCLQAQKRQRAQSDISNLGNPSICPPA

MPSLDQLHRTLPPLSFINSQVLLTSPATSVANPSSSTPIKPHTGHEIMGEAKSDQASVNVKMVRIDQALVKVLAPRRSGQ

LLRTVMALEGLALTVLHTNITTVHHTVLFSFHVHMGLLCRMSVKEIATVLHGTFSSSPPHSLQ

>Pt00G04250

MDDPTGNPLAAQTNFQFQLQDFIDEANFDRYIDLIRGENEITAFDCDLINGFLVDNQFGLSTGDKFDCDLINHVPTHTSS

AMEQDPNYVPIALPSFDGDMGLEAEEDTDEEDSSGTATTTKKTKKDRSRTLISERRRRGRMKEKLYALRSLVPNITKMDK

ASIIGDAVLYVQELQMQANKLKADIASLESSLIGSDRYQGSNRNPKNLQNTSNNHPIRKKIIKMDVFQVEERGFYVRLVC

NKGEGVAPSLYRALESLTSFSVQNSNLATTSEGFVLTFTLNVKESEQDMNLPNLKLWVTGALLNQGFELLTA

>Pt01G08350

MEEILSSSSSSSLMSFAQETSSTLQQRLQFFLHSRPEWWVYSIFWQASKDASGRLVLSLGDGHFRGNKKYASKESNKQNH

SKFGFNLERKSLFNEDMDMDRLVEGDVAEWYYTVSVTRAFAVGDGILGRAFSSGAFIWLTGDHELQIYDCERVKEARMHG

IQTFVCVSTPSGVLELGSPDLISEDWGLVQLAKSIFGADINAGSVPKQANQESQPQIPNRTVSNFLDFGMFSSPQKERTT

CLENQKESDTRKEPSGQGRSSSDSGRSDSDAGFTENNIGFKKRGRKPSGKELPLNHVEAERQRRERLNHRFYALRSVVPN

VSKMDKASLLADAATYIKELKSKVNELEGKLRAVSKKSKISGNANIYDNQSTSTSTMTNHIRPTPNYMSNNAMEVDVKIL

GSEALIRVQSPDVNYPAARLMDALRELEFSVHHASVSKVKELVLQDVVIIIPDGLVTEEVMRAAIFQRMQN

>Pt01G14220

MTDYRLPPTMNLWTDDNASVMEAFMNSSDLSSLWAPPPQSSASTSTPSAAAQPSEKTMLNQETLQQRLQTLIEGACEGWA

YAIFWQSSYDYSGASVLGWGDGYYKGEEDKGKTRTRNSASSAVEQEHRKTVLRKLNSLIAGPNSVTDDAIDEEVTDTEWF

FLVSMTQSFVNGSGLPGQALFNGSPVWVAGSERLGASPCERARQGQVFGLQTLVCIPSASGVVELGSTELIFQSSDLMNK

VRVLFDFNSLEVVSWPIGTTNTDQGENDPSSFWLTDPETKDGNGGIPWNLNGSDQNKNNHHSSNQSSSSLTDHLGGIHHA

QNHQQQPIHARSLFTRELNFGECSTYDGSSVRNGNSHLTKPESGEILNFGESKRTASSANGNFYSGLVTEENNKKKRSVG

NEEGMLSFTSGVILPSSCILKSSGGTGGDSDHSDLEASVVKEADSSRVVEPEKRPRKRGRKPANGREEPLNHVEAERQRR

EKLNQRFYALRAVVPNVSKMDKASLLGDAISYIDELRTKLQSAESSKEELEKQVESMKRELVSKDSSPPPKEELKMSNNE

GVKLIDMDIDVKISGWDAMIRIQCCKKNHPAARLMSALRDLDLDVQYANVSVMNDLMIQQATVKMGSRFYTQEELRVAIS

TNVGGVH

>Pt01G29430

MEPIGATAEGEWSSLSGMYTSEEADFMEQLLVNCPPNQVDSSSSFGVPSSFWPNHESTMNMEGANECLLYSLDIADTNLY

HFSQVSSGYSGELSNGNVEESGGNQTVAALPEPESNLQPKRESKMPASELPLEDKSRKPPENSKKRSRRTGDAQKNKRNV

RSKKSQKVASTGNNDEESNGGLNGPVSSGCCSEDESNASQELNGGASSSLSSKGTTTLNSSGKTRASKGAATDPQSLYAR

KRRERINERLRILQNLVPNGTKVDISTMLEEAVQYVKFLQLQIKLLSSEDLWMYAPIAYNGMDIGLDHLKLTTPRRL

>Pt02G04200

MGEKFWVKEENRMVESVLGVEACEFLITSASKNILNDLVSPPVNLGVQQGLGKVVEGSHWNYVIFWYASGLKSGGSILVW

GDGICQDPKGGGVVHGSSSGDGKLEGVEKRKVKKCVLRKLHACFNGSDDGSFAASLDEVSDVEMFYLTSMYFTFRCDSAY

GPGEAFKSGRSIWASSMPSCLDHYQLRSVLARSAGFQTVVFLPVKSGVLELGSVKSIPEEHDFVEKAKGLFGASNNAQAK

AVPKIFGRELSLGGSKSRSISINFSPKVEDELVFTSESYAMKATSTNQVYGSTSNGRPSDKSEAKLFPHLNQAIVGFNTE

TVVGGLEQPNDDLSPQGDERKPRKRGRKPANGREEPLNHVEAERQRREKLNQRFYALRAVVPNISKMDKASLLGDAITYI

TDLQKKIGALETERGVVNNNQKQLPVPEIDFQPGQDDAVVRASCPLDSHPVSSIIETFREHQITAQECNVSMEGDKIVHT

FSIRTPSGAAEQLKEKLEAALSK

>Pt02G05410

MAAPLSSRLQTMLQAAVQSVQWTYSLFWQMCPQQGILVWGDGYYNGPIKTRKTVQPMEVSTEEASLQRSQQLRELYDSLS

IGETNQPERRPCAALSPEDLTETEWFYLMCVSFSFSPGAGLPGKAYDRKQHVWLTGANDIDSKTFSRAILAKSAGVQTVV

CIPLLDGVVEFGTTDKVKEDLGFIQHVKSFFSDHHHLPPPKPALSEHSTSNPATSSDHPYLYSPPIPPFYVAADPPANAG

QMNEDDEEEEEDEEDDEDDEEDQESDSEAETSREALLEPCQAQNPLQVAVVEPSELMQLEMSEGIRLGSPDDGSNNLDVD

FPLINPESLMDHQSRADSFRTESARRWAPMLQDNPFSGSLQPSASGSPTLEDLAQEDTHYSQTVSTILQNQTIELAAEPS

LNAYEAYSNQSAFAKWMNLTDHCLNVPVETTSQWLLKYILFTVPYLHSKYREENSPKSRDGDATNKFRKGTPQDELSANH

VLAERRRREKLNERFIILRSLVPFVTKMDKASILGDTIEYVKQLLKKIQDLEACNKQMESEQRSRSVDPPQTITTSTSLK

EQNNGITVVDRARSVGPGSDKRKMRIVEDYTTGRAQPKSVDSLPSPEPMVDVEPEISVEVSIIESDALIELKCGYREGLL

LDIMQMLRELRIETIAVQSSSNNGIFVGELRAKVKENVSGKKLSIVEVKRAIRQIIPHD

>Pt02G05540

SVSCPCLSNWELKTDPPLSLSLGLVHSQISLYTAFYREREREVIVEVELVWSAMNHCIPDWNLEGDLPVSNQKNPIEPDN

ELVELLWRNGQVVMHSQTQRKPNPHVQKHDSPTVRGYDSTLNSSHLIQDDETVSWIHDPLEDSFEKEFCSNFFSELPPPL

SDQISEEKSAKFDASKTSHQQQQLNNNKHPVVPEFPGNPMPPPRIQVQEQNHISVGGFGKAVNVNFSQFSAPLKVGDFRS

SSRQFGGQGSGDFSQGEARECSVVTVGSSNQTPRDRFLSRASGNAIETGTGLSAGPSIDDPRKVISQSERGKGDQTLDPT

ATSSSGGSGSSFARTCKQSAVPSRGQKRKTMDAEESECQSEDAELDSAVANKPAKRSGSTRRSRAAEVHNLSERRRRDRI

NEKMRALQELIPHCNKTDKASMLDEAIEYLKSLQLQLQVMWMGSGIVPVMFPGVQHFMSRMGMGMGMGPPPLPSMQNPMH

LPRVPLVDQSISMTPTQNQAVICQTPVLNPVNYQNQMQNPTFSDQYARFMGFHMQAASQPMNMFRFGSQTVQQNQMMAPP

NSGGGPLSAGTAASDAPPSGKTG

>Pt02G10530

MMDGGESILSSASKTEAKISTESSCKSHKETERRRRQRINAHLSTLRTLLPNPTKTDKASLLAQVVHHVRDLKMKAAGSA

RQYSNNCSSGLEPEENWPYPGEVDEATLSCCGHEEKMIKVSVCCEDRPGLHMDLTRAIKSVRARAVRAEMMTVAGRTKSV

VVMRWDNGSGGEEDVGILKRALNAVVENRASGSGFGQVVQGNKRARVFGLVSDVDRDRN

>Pt02G14330

MSHQCVPSWELDDNPTTAPKVSLRSHSNSGAPYMPMLNYEVAELTWENGQLAMHGLGQPRVPAKPIASTSPSKYTWDKPR

ASGTLESIVNLATCIPQCNKQTFDNSGSDHDFVPWFNHHRASASATMTMDALVPCSKRSDQERTTRVIDSSPAGLGTDCV

VGCSTRVGSCSAPTATQNEVGLLTRKREKVARVPVPAEWSRDQSVNRGATFSKKDSQQVTVDSCERELGVGFTSTTSFGS

QENTSSGTKPCTKTNTADENDSVCHSRPQREAGDEDDKKKGNGKSSVSNRRSRAAAVHNQSERKRRDKINQRMKTLQKLV

PNSSKTDKASMLDEVIEYLKQLQAQVQMVSRMNMQPMMLPMALQQQLQMSMMAPISMGMAGMGMGMGMGMGMGVVDMNTL

AARSNITGVSPVLHPTAFMPMTTWDGSNSHERLPTAAPSATVMPDPLSAFLACQSQPMTMDAYSRMASMYQQLHQQSPAS

NSKS

>Pt02G17210

MGGGAWNDEDKTMVAAVLGSKAFNYLLSNSVANQNLLMVMCGDENLQNKLSDLVDCPNSSNFSWNYAIFWQISCSKSGDW

VLGWGDGSCREPKEGEESEFTRILNIRLEDETQQRMRKRVIQKLQTLFGESDEDNYALGLDRVTDTEMFFLASMYFSFPR

GEGGPGNCYASGKHVWISDALKSGPDYCVRSFLARSAGFQTIVLVATDVGVVELGSVRSVPESIEMVQSIRSWFSTRSSK

LKELRRFLNGSRLAFPGTRNRLHGSSWAQSFGLKQGTPGEVYGSQATANNLKELVNGVREEFRHNHYQGQKQVQVQIDFS

GATSGPSGIGRPLGAESEHSDVEASCKEERPGAADDRRPRKRGRKPANGREEPLNHVEAERQRREKLNQRFYALRAVVPN

ISKMDKASLLGDAISYINELQAKLKKMEAERGKLEGVVRDSSTLDVNTNGESHNQARDVDIQASHDEVMVRVSCPMDSHP

ASRVIQALKEAQVTVIESKLSAANDTVFHTFVIKSEGSEQLTKEKLMAAISFVWSCRGKVRWLSSTIPD

>Pt02G17690

MEELIISPSSPSSPVSLSQETPPTLQQRLQFIVQNQPDWWSYAIFWQTSNDDSGRIFLGWGDGHFQGSKDTSPKPNTFSN

SRMTISNSERKRVMMKGIQSLIGECHDLDMSLMDGNDATDSEWFYVMSLTRSFSPGDGILGKAYTTGSLIWLTGGHELQF

YNCERVKEAQMHGIETLVCIPTSCGVLELGSSSVIRENWGLVQQAKSLFGSDLSAYLVPKGPNNSSEEPTQFLDRSISFA

DMGIIAGLQEDCAVDREQKNARETEEANKRNANKPGLSYLNSEHSDSDFPLLAMHMEKRIPKKRGRKPGLGRDAPLNHVE

AERQRREKLNHRFYALRAVVPNVSRMDKASLLSDAVSYINELKAKVDELESQLERESKKVKLEVADNLDNQSTTTSVDQS

ACRPNSAGGAGLALEVEIKFVGNDAMIRVQSENVNYPASRLMCALRELEFQVHHASMSCVNELMLQDVVVRVPDGLRTEE

ALKSALLGRLE

>Pt02G18030

MKVGFSPHLLRRLGFLGEHNLDLECLDQGFINTESLRFGEERPYFSSPIFDEKMPFLQMLQTVETPPTFPFKEPCFQTLL

KLQHLKKPWNMNNYYMPETESQVQPPELESCVTHDIFDLHSPVKSETRELPNPHSNSCLGGVSPEPAEPYSGSLIPLGTQ

PQTVPNIKTQFSKSTTIITRERRKRKRTRPTKNKEEVESQRMNHIAVERKRRRLMNDHLNSLRSFMPPSYVQRGDQASII

GGAIDFVKELEQLLQSLEAQKRMKEIEAGSTIGISSNQYFTSPPQSDNLAEKGGKCEEKRTVKKKSEAAEIEVTAVQNHV

NLKIKCQRSLGQLARAIVALEELSLTVLHLNISSSQATILYSFNLKLEDDCELGSTDEVAAAVHQIFSSFNG

>Pt02G25280

MNPYVPDFEMDDDYSLPPPPSTHTRPRKPAMQEEEIMELLWQNGQVVMHSQRSQKKSSPPPSELDDAVLPADQLPGTKEI

RSSHDQQQEHHHLFMQEDEMASWLNHPLNDTNFDHDFCADLLYPPTASTASITREAAVTNAAASTVRGATQRMEARSYPA

VSAPRPPIPPVRRAEVVQNFAYFSRHRAGGVSESGRSNSKSVVRESTVVDSCETPTARISETAFARSADNTCGTINGAAV

AGTVSSAPSSNRETMTNPCEMTSTSSPGCSSASAELPALMSPVEDRKRKGREEEAECHSEFIAISITMAHGNQPQSVAIR

QKTQNSADSSKPLQGRRDAANPLKDAEFESADAKKRIRGSMSSKRSRAAEVHNLSERRRRDRINEKMRALQELIPRCNKS

DKASMLDEAIEYLKSLQLQVQMMSMGCSMVPMMFPGFQQYMPPMGIGMGMGMEMGLSRPMMPFPNILAGAPSATPAAAAH

LVPRFPVPPFHVPPIPAPDPSRVQPTNQVDPMLGSPGQQNPNQPRVPNFVDPYQHYLGLYQMHLPGVPRNQAMAQPSTSK

PSTSRVAENPGNHQSGCDGEKMAHAFPSSQAKKQEWKMGKGINGVMLRGCPPLSQ

>Pt03G09220

MTDYRLPPTMNLWTDDNGSVMEAFMNSSDLSSLWAPPPQTSASFSTPAAAAAAQPSDKTMLNQETLQQRLQALIEGARES

WTYAIFWQSSYDCSGASVLGWGDGYYIGEEDKGKGRMKNSASSAAEQEHRKKVLRELNSLIAGPSSVTDDAVDEEVTDTE

WFFLVSMTQSFVNGSGLPGQALFNGSPVWVAGSERLGTSPCERARQGQVFGLQTLVCIPSANGVVELGSTELIFQSSDLM

NKVKVLFNFNSLEVGSWPIGTTNTDQGENDPSSLWLTDPETKDGNAGIPSTTPAHQTANNNNHHSSSSLTDHSGGIHHVQ

NHHSHQQQQQQQIHTQSLFTRELNFGEHSTYDGSTVRNGNSHLMKPESGEILNFGESKRSPSSANGNFYSGLVTEESNKK

KKSPASRGGNEEGMLSFTSGVILSSSGLVKSSGGTGGDSDHSDLEASVVKEADSSRVVEPEKRPRKRGRKPANGREEPLN

HVEAERQRREKLNQRFYALRAVVPNVSKMDKASLLGDAISYINELKTKLQSAESSKEELENQVESMKRELVSKDSSSPPN

QELKMSNDHGGRLIDMDIDVKISGWDAMIRIQCCKMNHPAARLMSALKDLDLDVQYANVTVMNDLMIQQATVKMGNRYYT

QEELKVAISTKVGDAR

>Pt03G12800

MKWLIITLNSNFDMANELRNQERLPDNLKKQLALAVRSIQWSYAIFWSNPTGQPGVLEWADGYYNGDIKTRKTVQSIELN

ADELGLQRSEQLRELYESLSAGEANPQARRPSAALSPEDLTDTEWYYLVCMSFVFDNGQGLPGTTLANGHPTWLCNAPSA

DSKIFSRSLLAKSASIQTVVCFPFMRGVVELGVSEQVLEDPSLIQHIKTSFLEIPYTVTANHSSAKSDKELACATFNREI

HDTKPVPVIRCRELDTLSPDDNSNDQAATDSIMVEGLNGGASQVQSWQFMDDDFSNRVHHPLNSSDSVSQTIVDPVMLVP

FLKDGKVNGQSLQDIQDCNHKKLTALNLQSDDLHYQSVLSCLLKTSHPLILGPNVQNCYQEPSFVSWKKAGLMHSQKLKS

GTPQKLLKKILFEVPRMHVDGLLDSPEYSSDKVVGGRPEADEIGASHVLSERRRREKLNKRFMILKSIVPSISKVDKVSI

LDDTIQYLQELERKVEELECRRELLEAITKRKPEDTVERTSDNCGSNKIGNGKNSLTNKRKAPDIDEMEPDTNHNISKDG

SADDITVSMNKGDVVIEIKCLWREGILLEIMDAASHLHLDSHSVQSSIMDGILSLTIKSKHKGLNAASVGTIKHALQMVA

GNLFSNR

>Pt03G14730

MEEILSPSSSSSLISFAQETSSTLQQRLQFFLHSRPEWWVYSIFWQASKDASGRPVLSWGDGHFRGNKKYSSKVSNKQNH

PKFGFNIERKSLFNEDMDLERLVDGDVAEWYYTASVTRVFAVGDGILGRAFTSGSSIWLTGDRELQIFECERVTEARMHG

IQTFVCVSTPSGVLELGSPVFISEDWSLLQLAKSIFGAEINANPVPKQSNHESQPQISNCNVSNLLDIGLFSSPQTERTS

SLENKKEVFGQGRSSSDSGRSDSDAGFRENHIGFKKRGRKPGGKESPLNHVEAERQRRERLNHRFYALRSVVPNVSKMDR

ASLLADAVNYIKELKRKVNELEANLQVVSKKSKISSCANIYDNQSTSTSTMVNHIRPPPNYMSNNAVEVDVKILGSEGLI

RVQSPDINYPAARLMDALRELEFPVHHLSVTRVKELVLQDVVIRFDDGLVTEEAMRAAIFQRMQN

>Pt04G05570

MFDFCFNSNSSSGSGYMNTFDPFPRSLEGFSDVLQGRPMVSQTLVLDAEMGELVKAPARVGNKGISEAKALAALKSHSEA

ERRRRERINAHLDTLRGLVPCTEKMDKATLLAAVISQVNELKRNALESCKGLLIPTADDEVKVETYFDGTKEGTLYFKAS

ICCDYRPELLSDIRQAVDALPLKMVNAEISTLGNRLKNEFVFTSNRNKNAVDDAEAMQHLTKSIHHALTSVLEKGSASLE

YSPRTTLPNKKRRVTFFDSSSSSS

>Pt04G11290

MVKSAKSQLDEEDEPEDYDSSSYKGEVLKPESKSSEPKANANRSKHSETEQRRRSKINERFQALRNLIPQNDQKRDKASF

LLEVIEYIQFLQEKLQVYEGSYEGWSQEPAKLLPWKIDRASAESLLDHTQVMKNGSAHENSVMLANVHNSIESDMGTAAM

YKALDHPHGPTNPAIPFDVQTPSNVFAAVGRGGLPTQSLQESVSDVENMAYQLQSQLLHGRPCATECSTPNNTLNGQEDL

ASDSLSVNISNAYSQQILNTLTQALQSSGVDLAQTSIGVQIDVGKRENSTTAVAPSSKVNQYLSNQLLVQDGVGSSAENS

EQAHKRPRREKC

>Pt05G00180

MPLSELLYRMAKGKIDFSQEKDPSCSTDLSFVRPENDFGELIWENGQIQSSRARKIQPCSTLPCQNPKIRYKDIGNGTDI

RTGKFGMMESTLNELPMSVPAVEMGVNQDDDMVPWLNYPLDESPQHDYCSEFLPELSGVTVNGHSSQSNFPSFGKKSFSQ

SVRDSRTVSVHNGLSLEQGDVAKNSSAGDTEANRPRTSASQLYLSSSEHCQTSFPYFRSRVSAKNGDSTSNAAHHVVSVD

SIRAPTSGGGFPSIKMQKQVPAQSTTNSSLMNFSHFARPAALAKANLQNIGMRAGTGISNMERTQNKDKGSIASSSNPAE

CTPINSCSGLLKETSSHCLPVLMPPKVDAKPSEAKPAEGFVPAELPEATIPEGDSKSDRNCRQNFCESAIKGVADVEKTK

EPVVASSSVGSDNSVERASDDPTENLKRKHRDTEESEGPSEDVEEESVGAKKQAPARAGNGSKRNRAAEVHNLSERRRRD

RINEKMRALQELIPNCNKVDKASMLDEAIEYLKTLQLQVQIMSMGAGLYMPSMMLPPGMPHMHAAHMGQFLPMGVGMGMR

MGMGMGFGMSMPDVNGGSSGCPMYQVPPMHGPHFSGQPMSGLSALHRMGGSNLQMFGLSGQGFPMSFPCAPLVPMSGGPP

LKTNMEPNACGVVGATDNLDSATACSSHEAIQKINSQVMQNNVVNSSMNQTSSQCQATNECFEQPALVQNNAQDSGVADN

RALKSVGGNDNVPSSEAGEHLQ

>Pt05G05350

MAGNPPPEGLGDDFFEQILAGQPPGYGGEAVGSTSLPMMGLQLGSSAANVGLTRSSANNNMGMMPLGLNLEHHGFLRQQQ

DDGSSSLDTNNSNNINNASPFSITSAGFLGRDSVHMTSLFPTFGQLQIHSSIRSAPPPLGPPQIHQFNSQPTSGAVSAVP

QPPGIRPRVRARRGQATDPHSIAERLRRVRITERVKALQELVPTCNKTDRAAMLDEIVDYVKFLRLQVKVLSMSRLGAAG

AVAQLVADVPLSSVQGEGIEGGANQQAWENWSNDGTEQEVAKLMEEDVGAAMQLLQSKALCIMPVSLASAIFRARPPNAP

TLVKPESNPPS

>Pt05G13970

MGDVYNNETNNINNTCSASSSSRHYHRSSDDISLFLHQILPRSPSSTSSSSFIGPQTLQPFSVPAPFDDPLRSGVILGVD

SSGGGGGALWSGNVRTRLIMSENETDHECDCESEEGLEALIDEISAKPAPPRSSKRTRAADVHNLSEKRRRSRINEKMKA

LQNLIPNSSKTDKASMLDEAIEYLKLLQLQVQMLSMRNGTSLHPMFLPGVLQPSQLSQVRMGFDEEIGSKHMNMTTSVPL

DGEARLQTMLTLPDHCTSSNQASNPIHSETSSGFQSTIQAHFSPFPLRVSSEETWRKDILPGQQLNGLHQKIGILDVLLA

ATKQRTC

>Pt05G20720

MMTLCLGFNTLLKILLKRSSVPIFSRNCLHHFLIKFWKESRLHLMPPHLHSNSISISISDNSTTINLMWFLNFQEIQCPG

NPMPPPRIQVPEKNHAGVGGFGEAVNANFSQFSAPFKGGDFRTSSGQFGGQGSGNSPQGEVRECSVVTVGSSNQIPHDRD

MSRASSNAMGTSTAFSTGPSMDDPRKIVSQSERGKTETLEATLTTSSGGSGSSFGRTCKQSAGPSSSQKRKTIDTEDSEF

QSEAAELDLDSMAGNNPTKRSGSTRRSRAAEVHNLSERRRRDRINEKMRALQELIPHCYKTDKASMLDEAIEYLKSLQLQ

LQVMWMGGGMAPMLFPGVQHFMSRMGMGPPLPSMQNPMHLPRVQLIDQSISMAPTQNSGVMCQAPVLNPVNFHNQMQNPA

FADQYARFMGFHMQAASQPMNMFRFGSQTVQQNQMMAPTSSVGGPLSAGTAVSDAPPSDKVGKSFYSLYLPGKYSSIC

>Pt05G20860

MATPPSSRLQTMLQAAVQSVQWTYSLFWQMCPQQGILVWGDGYYNGPIKTRKTVQPMEVTTEEASLQRSQQLRELYDSLS

IGETNQPARRPCAALSPEDLTETEWFYLMCVSFSFPPGGGLPGKAYARRRHVWLTGANEIDSKTFSRAILAKSARVQTVV

CIPLLDGVVEFGTTDKVQEDLGLIQHVKTFFSDHHHRHLTPPKPALSEHSTSSPATSSHDHPRFHPPPIPPFYVAAEPSA

NAEQIDEDEEEDEEEEEHDSDSEAETSRDDHLEPRQAQNPHQVVAAEPSELMQLEMSEDIRLGSPDDGSNNLDSDFPLTG

PDNSMDHRSRADSYKAESARRWTMLQDNPFSGNLQPSASGPPPLEDLAQEDTHYSQTISTILQSQPVWLAAEPSSIAYEA

RYHQSAFSRWTNRSDHLFHVSVETTSQWLLKYILFSVPHLHSKSREDNSPKSRDGEAASRFRKGTPQDELSANHVLAERR

RREKLNERFIMLRSLVPFVTKMDKASILGDTIEYVKQLRQKIQDLETRNKQMESEQRPRSVDRPQRTSTSDSLKKQKSGV

TVVDRARSLGPLPDKRKMRVVEDSAGGGAKPKTVGALPQPEPVVHKELETSVEVSIIESDALLELECGFREGLLLDIMQM

LRELRIETIAVQSSLNNGIFAGELRAKVKENVNGKKVSIVEVKRAIHKIIPHD

>Pt05G22110

MGDKFWVNGEKGMVESVLGVEACEFLITSASKNLLNDLVSPPVSLGVQQGLVQLVEGFNWNYAIFWHASGLKTGGSILVW

GDGICRDPKGQGIGDGSSSGDGKSEGAEKRKEVKKRVLQKLHMCFNGPDDDNFAASVDEVSDVEMFYLTSMYFTFRCDST

YGPGEAYQSGRSIWALGMPSCLGHYQLRSVLARSAGFQTVVFLPVKSGVLELGSVKSIPEQHDFVEKARSIFGASNTAQA

KAAPKIFGRELSLGSSKSRSISINFSPKVEDELIFTSEPYTMQAMSTDQDYVSTFSGHPSDKSEAKLFPHLNQTIAGFNA

ETLVGGLEQPKDDLSPQGDERKPRKRGRKPANGREEPLNHVEAERQRREKLNQRFYALRAVVPNISKMDKASLLGDAITF

ITDLQKKIRVLETERGVVNNNQKQLPVPEIDFQPRQDDAVVRASCPMESHPVSTIIETFREHQITAQDCNVSVEGDKIVH

TFSIRTQGGAADQLKEKLEAALSK

>Pt06G03710

MQPENCQENSQLYRFLTENGMINVGPYGFPAAMQTLCTSSSTSYHNSNYHFERSVITDMTPEDRALAALKNHKEAEKRRR

ERINSHLDKLRGLLPCNSKTDKASLLAKVVQRVRELKQQTSELPGLESFPSETDEVTVLSGEYSSDGQLIFKASLCCEDR

SDLMPDLIEILKSLHLKTLKAEMVTLGGRIRNVLIIAADKDHSVESVHFLQNALKSLLERSNSSERSKRRRVLDRKLVIQ

>Pt06G07490

MESIENIGEEYQNYWETKMFLQNEEFDSWAIDEAFSGYYDSSSPDGAASSAASKNIVSERNRRKRLNERLFALRAVVPNI

SKMDKASIIKDAIDYIQELHKQERRIQAEILELESGKLKKDPGVDVFEQELPALLRSKKKKIDDRFCDFGGSKNFSRIEL

LELRVAYMGEKTLLVSLTCSKRTDTMVKLCEVFESLRVKIITANITTVSGRVLKTVFIEADEEEKDNLKTRIETAIAALN

DPLSPMSM

>Pt06G13560

MMDEYLNHLISSSSLVDGDVKESSSWVCSEPNQPNAFLPTSLELYQDDKKNSPVSMISSNQSVESLATQDTSSVVLGSES

DYAVDKVLISEQARLQNDCQNCNGNPSPDGMARGNLKFGNTGLQCNGILPTLSSLNYPNQLPIVGDLTSYLSFSEASNAG

CNGREQSEYLRSLKNLQNLSSIPQLWPSQSYEGVSSLPPLMGQDRIEGSGLRGGNLDDDMHIMGKGYMGMDEILRLDKLS

ASPTTEGKEDLQSCPFSSGIAEPNVNMSMNQLSSMPQTTSAAPVEGCNGTGKTRVRARRGHATDPHSIAERLRREKIAER

MKNLQELVPNSNKVDKASMLDEIIEYVKFLQLQVKVLSMSRLGAAGAVIPLLTDGQPEGHNSLSLSPSAGLGIDISPSAD

QIAFEQEVLKLLESDVTMAMQYLQSKGLCLMPIALAAAISSVKASLSGTTSEERKNNGYTSGLVSSSSSITGIDTHPMSN

DNNIATGTLSSKGMIVNGCNEVVKQEVLKNT

>Pt06G14880

MDSPEISSLIFTIPSSSHMPEAHREAPSLLPPILPSTSSPLQHTLQLLQPVLPTPTGNILQQVLQSLQQEPSPQQQAIQS

SQSTPSTTSLLQEAMQPLQAIQSSTSPHQQALQAFALERNIQLPTPESEDAVMTRAILAVLTFPSPSSSSSSHSLPHMHR

VRQGASAFNNYRSALAPKTQTRASLHRHSMLTRVITYYRRLNIERREHMLGGRPSSTQLHHMISERKRREKINESFKALR

SILPPEAKKDKASILTRTREYLTSLKAQVEELTRKNQKLEAQLSKAAVSQVRDSSYERLDVRVTHISESTSEQRIIDLVV

NLRGESPILDTVITRILEFLRQVTDVSLISIEASTHTAESTSFNRVILRLNIEGTDWDESGFQEAVKRVVEDLAR

>Pt06G18660

MRPCSREMQGMNSLLNPSSQIPLQDLQNQQIQNSHFDPNSSSNDDFLEQMLSAIPSCSWADPKSPWDLNPPTNLPFPTNN

NSSSAKPRDLFNETPPSNTDNNNVGFHDNFDESVILASKLRQHQISGGSGAAAAAKMMLQQQLLMAAARGGLSQNDDIDV

SPTQGGDGSMQGLFNGFRAGSMNGTVRASNQSMQHFNHPQGGAMQSPNLGAQGAATTAVMNQPQASGSNGGAPAQPRQRV

RARRGQATDPHSIAERLRRERIAERMKALQELVPNANKTDKASMLDEIIDYVKFLQLQVKVLSMSRLGGAAAVAPLVADM

SSEAGGDCIQANANGGSIARTTNGNQTASTNDSSLTVTEHQVAKLMEEDMGSAMQYLQGKGLCLMPISLATAISTATCHN

RTSGIINSHNPLLQSNGEGPTSPSMSVLTVQSATMGNGVAKDAASVSKP

>Pt06G20210

MYEETACFETNNSIVEGGNDDGFCQVSPFMTGSSTTSSFEESFKLSMEELSNHYHQEESAAAASMEEIQLQHHMAFNNNC

HHLMEQYPTNHHQVLSYDHPSNWDPNTIQFQEMHQVLDQNGNFNATANTPSSLLPDLLNLFNLPRCTSTSTLLPNSSISF

TNPAHKTPSGFMGVDSTSVLFDSNPLAPQFRELVHSLPPHGYGLPAPLFGGGQGGDHVDGLSGGGLSYQDGGHGDGVFEF

TAEMACIGKGIRKSGKVITKHFATERQRREHLNGKYTALRNLVPNPSKNDRASVVGDAINYIKELLRTVEELKLLVEKKR

NGRERIKRRKPEEDGGVDVLENSNTKVEQDQSTYNNGSLRSSWLQRKSKHTEVDVRLIEDEVTIKLVQRKKVNCLLSVSK

VLDELQLDLHHAAGGLIGDYYSFLFNTKINEGSCVYASGIANKLLEVVDRQYASSTSVPAASC

>Pt07G01050

MALQQMHDVQPLSSHSNSAGPCSNQDDFRQALLNSINETTYNNSMAKQLISEEESKLVAAKKHSMAESNRRSRINTQFTT

LRTILPNLIKVNKASVLEETIRCVKELTNTVSELKEIYGGGRLECVFPGGADKLRIGSCEGKGQEVVKVVFSCDDKRKLL

SDVARAVRSVKGKVVRAEMVTMGGRTKCVLWVQGINGNEELEMLRRVLNALTEKPNMARTCNRPKLTLL

>Pt08G07080

MEDHFSPCWPAAPAEANWVQTSAAVYDESFLVPCPSHASASANFQVNGFPSWSIPIQEASENKAASNSKSHSQAEKRRRD

RINAQLGILRKLIPKSEKMDKAALLGSAIDHVKDLKQKATEISRTFTIPTEVDEVTVDCDVSQATNPSSTNKDKDSTFIR

ASVCCDDRPELFSELIRVLRGLRLTIVRADIASVGGRVKSILVLCNKCSKEGGVSISTIKQSLNLVLSRIASSSVPSNYR

IRSKRQRFFLPSHLSQQYT

>Pt08G11200

MRGKGNQNQVEEEYEEDEFGSRKDGPSSSFTVNNNNSSKDGKNSDRANAIRSKHSVTEQRRRSKINERFQILRDLIPHSD

QKRDTASFLLEVIEYVQHLQEKVQKYEGPYQGWSPEPAKLMPWRNSHWRLQSSVGHPQAIKNGYVPGETFPGKLDENNIA

LTPAMLPSTPNLVESDHVACKVLEHQPELGNKAMPLPTPAPIRSVGLVAHPCQLPVSDAQSAECPITSEMLNQQELAIEA

GTINISSVYSQELLNTLTQSLQSAGVDLSQANISVQIDLGKRANRGLTSGTLTSKDPQNPHPTNEMITHLRDASGGEDSD

QAQKRLKTS

>Pt08G11600

MCGLKEEDQEEQTIHNLQNYQEQLLFQYHQQMQQHHQQQSSDIYGGARGSGLIFPEVSPILPWPLPPAHSFNPDHFTSNH

PVRDHDPFLIPPPIPSSYGGLFNRRSPSLQFAYDGTSSDHLRIISETLGPVVQPGSAPFGLQAELSNMTAQEIMDAKALA

ASKSHSEAERRRRERINNHLAKLRSLLPSTTKTDKASLLAEVIQHVKELKRQTSLIAETSPVPTEMDELTVDTADEDGKF

VLKASLCCEDRSDLLPDLIKTLKALRLRTLKAEITTLGGRVKNVLFIAGEEDSSSDSNDHQQQQQPLQYSISSIQEALKS

VMEKTGGDESSSGSVKRQRTNINVLQQQHRSL

>Pt08G18960

MRGLDRAMERLRPLVDSNAWDYCVVWKLGDDPSRIHGEVVISAEPRWLCHATVTTHDSNTLREVAGTQVLIPVIGGLVEL

FAAKHMKKDEKMIESIRAHCHVPVKQEAVTELGYSNSSFNDHRLDSLLEENLPHSCHLLSLIPRTQFLLPLSQPRNSISF

EGSSSGSNPSNEAPSFVSNASQLPQHGHLELSVGKSNHDEKILKQRAGSADCNKKVPKVMRRSERDDYKSKNLVTERNRR

TRIKTGLFALRALVPKISKMDKAAILGDAIDYVGELLKEVKNLQDEIKNAEEEERRASNIELKTSKLEIFQEDHVSSSKI

NQDSSGFVEKKGAEVQLEVDQISKRQFLLKFLCEQRQGGFGRLMETIHSLGLQILDANITTFNGNVLNILKVEADKDIHP

KTLKKSLIELTGNLIQTFGSQI

>Pt09G00560

MDDPTGNPLAAQTNFQFQLQDFIDEANFDRYIDLIRGENEITAFDCDLINGFLVDNQFGLSTGDKFDCDLINHVPTHTSS

AMEQDPNYVPIALPSFDGDMGLEAEEDTDEEDSSGTATTTKKTKKDRSRTLISERRRRGRMKEKLYALRSLVPNITKMDK

ASIIGDAVLFVQELQMQANKLKADIASLESSLIGSDRYQGSNRNPKNLQNTSNNHPIRKKIIKMDVFQVEERGFYVRLVC

NKGEGVAPSLYRALESLTSFSVQNSNLATTSEGFVLTFTLNVKESEQDMNLPNLKLWVTGALLNQGFELLTA

>Pt09G08140

METSSITALYELGMEDPGFTNQWYMNSLDDISLLPLAAAAFGENVHHPFSNQNFNLKTSMDSTPTSINVRPTKQMKTFHL

SDPQSAFSPNFLSFVNPNHANQMGLVKPKEEAVCSKSINNFPSDMVVSQDIFGSQNYVIKGCQGPERISTNTPRLSQSQD

HIIAERKRREKLSQRFIALSAVVPGLKKMDKASVLGDAIKYLKQLQEKVKTLEEQTKRKTMESVVIVKKSHIYVDEGDVN

ASSDESKGPIHETLPEIEARFCDKHVLIRIHCEKRKGVLEKTVAEIEKLHLSVINSSVLAFGTSALHVTFIAQMDIDFNM

SLKDLVKTLRSAFEFFM

>Pt09G08900

MAEGEWSSLGGMYTSEEADFMAQLLGNCPNQVDSSSNFGVPSSFWPNHEPTTDMEGANECLFYSLDFANINLHHFSQGSS

SYSGGSGILFPNTSQDSYYMSDSHPILANNNSSMSMDFCMGDSYLVEGDDCSNQEMSNSNEEPGGNQTVAALPENDFRAK

REPEMPASELPLEDKSSNPPQISKKRSRNSGDAQKNKRNASSKKSQKVASTSNNDEGSNAGLNGPASSGCCSEDESNASH

ELNRGASSSLSSKGTATLNSSGKTRASRGAATDPQSLYARKRRERINERLRILQTLVPNGTKVDISTMLEEAVQYVKFLQ

LQIKLLSSEDLWMYAPIAYNGMDIGLDHLKVTAP

>Pt09G13630

MRYPYQPGNIMNVVQNLMERLRPLVGVKGWDYCVLWKLSDDRRYIELMDCCCAGTEATQNGEELQFPVSAVLPCRDVMFQ

HPGTKSCELLAQLPSSMPLNSGFHAQTLSSNLPRWLNFSSSSDSNVLEETVGTRALIPVPGGLMELFIAKQVPEDQHVID

VVTSQCNFLMEQEAMINSTNMDSSLSIDVNVMSENQSKPFLANENEQEDHHSLNIPYDTSLDRLHMSSSPMNNFMHQFNY

STDETKTKGDLFQGVESGLQDMDDLQKSMMANAESTQMQYMESGLTTKDQHGNDKESIKLENGPSAEYSHSDCNDDEDDA

KYRRRTGKGPQSKNLVAERKRRKKLNDRLYALRSLVPNISKLDRASILGDAIEFVKELQKEAKELQDELEENSEDEGAKN

GNNNNMPPEILNQNGVNLGAYRSDYAVNGFHVEASGISTVSKQNQDSENSHDKGHQMEAQVEVAQIDGNEFFVKVFCEHK

PGGFVRLMEALDSLGLEVTNANVTSNRGLVSNVLKVEQKDSEMVQADYVRDSLLELTRDPPRAWPEMPKASEICCSGMDY

PHHDHHQHHLQNGHMNYNHHHLHHL

>Pt10G04150

MLCPLKQEAMIGHCYSNSSLNKYCLDLFLEENLLLSCHLLSLFPQIQFLHPLTQPNNNLSFEGSSSSSNPSNEASSFVSY

ASQFPQHGHMKLLQRESSEIYPGSESDGYKSKNLVTERNRRTRIKTGLFSLSARVPKISKMDKAAILGDAIDYISELLKD

VKNLRDEIKNAEEEECRASNMELKTSKLETCQKGCMSSTKVNQDPSGFVKKERTEVQLEVDQIGKRHFLLKFLCEKKRGG

FGRLMETIHSLGLQIHDANITTFNGKFLNILKIES

>Pt10G13000

MCGLKEEDQGECSQTIHNLQNYQEQLLLQYHQQMQQHQQQQSSDIYGGARGSGFIFPEVSPILPWPLPPVHSFNPAHFTP

NHPVRDHDPFLIPPPVPSSYGGLFNRRAPSLQFAYDGTPSDHLRIISDTLGPVVQPGSAPFGLQAELSKMTAQEIMDAKA

LAASKSHSEAERRRRERINNHLAKLRSLLPSTTKTDKASLLAEVIQHVKELKRQTTLIAETSPVPTEMDELTVDTADEDG

KFVIKASLCCEDRPDLLPDLIKTLKALRLRTLKAEITTLGGRVKNVLFISGEEDSSSDSNDQHQQQEPLQYSISSIQEAL

KAVMEKTGGDESSSGSVKRQRTNINLLEQQQQQQQHRSL

>Pt10G18670

MEDHYSPCWPAAPAEANWDQTSAAVYDESFLVPCPSHASASANFQVYGFPSWSVPLQEASEDKAASSSKSHSQAEKRRRD

RINAQLGILRKLVPKSEKMDKAALLGSAIDHVKDLKQKATEISRTFTIPTEVDEVTVDCDVSQVTSPPSTNKDKDNTFIR

ASVCCDDRPELFSELITVLKGLRLTIVRADIASVGGRVKSILVLCSECSEEGSVSISTIKQSLNLVLSRIASSSVPSNYR

IRSKRQRFFLPSHLSEQYE

>Pt11G06550

RTMVSQTLMLDAEKGELLKAPARIGKMGISEAKAFAALKSHSEAERRRRERINAHLATLRGLVPCTEKMDKATLLAAVIS

QVKEHKKNALEACKGLLVPMDDDEVKVETYFDGTLHFKASICCDYRPELLSDLRNAIDALPLKTVSAEISTLGSRLKNEF

VLTNRRNKNALDDAGAIQLLTNSIHQTLTSVMEKGSASPKYSPRTKLPNKRRRVTFFDSSSSSS

>Pt12G10490

MAAFSYQHPPLFLDSVILPNITTPIMNMNNSMYWFYDEAGGINSNSFYQVYPPETFHEAPLDVRFHEFSHHDHSSKVSLS

DNETSLTKKQSTGSSTVVDKLETGEQVTQEVTPVDRKRKTTNGSLNSAQSKDVKEVKSKRQKKCRGDMKQEEKRPKAVKK

VPEEPPTGYVHVRARRGQATDSHSLAERVRREKISERMKMLQRLVPGCDKVTGKALMLDEIINYVQSLQNQVEFLSMKLA

SVNPMFYDFGMELDAFMVRPERLSSMSPPLPSLQQCSPIQPTAFADAAAAATTTATPTTSFATANNYPLIDNSTSLLLQG

MRPSAFTTEDSCNLMWDVDERRQKFLSPSGLTSNLCSFH

>Pt12G10600

MLYRLNSNNIWLEDHKEEQDSTTNHHHHNNITNTAAGCGGVMLEGKEEMGSLSTFKSMFEVEDEWYVTNNNSTIHQNHQD

SIKDLTFSPSLVDPDNLLLHQVDSSSSCSPSSSVFNNLDPSQVHYFMHPKPTLSSLLNVVSNNPLEHGFDLSEIGFLENQ

GTNSTTTANVSSLLNRGSGVLGNLGNFTDLSSNSQISIPNLCSDPQFSSSRMLQLPENGPGFNGFRGLDEISGNQLFFNR

SKLLRPLETYPSMGAQPTLFQKRAALRKNLGEVERDKGKREMTQISEEKDKKRKFSSGDDFLEDVSFDGSGLNYDSDEFT

ENTNLEETGKNGGNSSKANSGVTGGGVDQKGKKRGLPAKNLMAERRRRKKLNDRLYMLRSVVPKISKMDRASILGDAIDY

LKELLQRINDLHNELESTPPSSSLTPTTSFHPLTPTPSALPSRIMDKLCPGSLPSPNGQPARVEVRVREGRAVNIHMFCG

RKPGLLLSTMRALDNLGLDIQQAVISCFNGFAMDIFRAEQCKEGQDMHPDQIKAVLLDSAGFHGAM

>Pt13G00130

MPLSELLYRMAKGKTDSSQEKNPACSTDLSFVPENDFGELIWENGQIQYSRARKIQTCNSLPPKIRDKDIGNGTNTKTGK

FGTMESTLNELLAVPAVEVRANQDDDMVPWLNYPLDEPLQHDYCSDFLPELSGVTVNEHSSQSNFPSFDKRSCNQSITDS

HTVSVHNGLNLEQGDVVMNSSAGDIDAKRPRTSASQLYPSSSEQCKTSFPFFRSRDSTKKDDSTSNAVHHVIAPDSIRAP

TSGGGFPSIKMQKQVPAPSPINSSLINFSHFARPAALVKANLQNVGMRASSGTSSMERMQNKDKGSIGLPKETDSHCRPN

MMSSKVEVKPTEVKPAEGSVPAELPEEMSQEGDSKSDRNCHQNFGESAIKGLEDVEKTTEPLVASSSVGSGNSAERPSDD

PTENLKRKHRDTEESEGPSEEIIYVQDAEEESVGAKKPASARAGNGSKRGRAAEVHNLSERRRRDRINEKMRALQELIPN

CNKVDKASMLDEAIEYLKTLQLQVQIMSMGAGMYMPSMMLPPGMPHMHAAHMGQFLPMGVGMGMGFRMGMPDMNGGYSGC

PMYQVPPMHGAHFPGSQMSGPSALHGMGGPSLQMFGLSGQGLPMSFPRAPLMPMSGGPPPKTNREPNACGVVGPMDNLDS

ATASSSKDAIQNINSQVMQNNVANRSMNQTSSQCQATNECFEQPAFAQNNGEGSEVAESGVLKSAGGTDITPSRATGCD

>Pt13G11760

MEPCSWNSARVQAYACEGMSDVFLVNSRLQAETRNGSRSTSSLVLDNERGELVEATVRMERKGVSAEKSIAALRNHSEAE

RKRRARINAHLDTLRSLVPGTSKMDKASLLAEVISHLKELKIQAAGAGEGLLMPLDIDEVRVEQEEDGLCSAPCLIRASI

CCDYKPEILSGLRQALDALHLMITRAEIATLEGRMMNVLVMSSCKEGLGGDSKVRQFLAGSVHKAFRSVLEKFSASQEFS

LKPTLSNKRRRVGLLQPFSSSSSGDLCS

>Pt14G02580

MEDLYGAAAATEPEEISTFLHQLLHNNSSSPSKFMHHALSTPVENGVELLDRHRFSETECGAGVNFSDPDGYYAKEGVGN

AVVSKRGGVSVEDDLGDFSCDSEKGVEVQANTARPRSSSKRSRAAEVHNLSEKRRRSRINEKMKALQNLIPNSNKTDKAS

MLDEAIEYLKQLQLQVQMLTMRNGLSLHPMCLPGALQPMQLPLSGMSFDEGIGLLTTNTLTGIFSANEESSEQNSLNLPT

QCTISNQPITIPSGTNITSSETNFGFEPQIHVNHAPFNLSTSSKEICREGTPQAKLEMNQTTKTSPSGVA

>Pt14G06650

MSHQCVPSWEVDDNRTTAPKLSLRFHSNSSAPDMPMLDYEVAELTWENGQIAMHGLGPPRVPAKPIASTSPSKYTWDKPR

ASGTLESIVNQATCVPQCNKATFDNSTGSDHDLIPWFNHHKASASATMTMDALVPCSNRSDQGRTTHVIDSGPAGLGTCV

VGCSTRVGSCSAPAATQDEDGLLTGKRARVARVPVPPEWSRDQSVNHSATFGKKDSQQMTVDSCEREFGVGFTSTSFGSQ

ENTSSGTNPCTKTLTADENDSVCHSRPQREAGKEDDKKKGNGKSSVSTKRSRAAAIHNQSERKRRDKINQRMKTLQKLVP

SSSKTDKASMLDEVIEYLKQLQAQVQMMSRMNMQPMMLPLALQQQLQMSMMAPMSIGMAGMGMGMGVMDMNTIAARSNMT

GIPPALHPTAFIPLTTWDGSSGHDRLQTTAADPMSAFLACQTQPMTMDAYSRMAAMYQQLHQQPPASNSKG

>Pt14G09970

MKIELGVGGGAWNDEDKTMVAAVLGTKAFNYLLSNSVANQNLLMAMCGDESLQNKLSDLVDRPNASNFSWNYAIFWQISC

SKSGDWVLGWGDGSCREPKEGEESEVTRILNIRHEDETQQRMRKRVIQKLQTLFGESDEDNYALGLDQVTDTEMFFLASM

YFSFPHGEGGPGKCYASGKHMWISDALKPGPDYCVRSFLAKSAGFQTIVLVATDVGVVELGSVRSVPESIEMVQSIRSWF

STRNSSIRAKPMAAAAAAAAAMPAVSEKKDENSPFSNFGIVERVGVPKIFGQDLNSNHGHGHGFREKLVVRKMEERPSWN

AYQNGTRLALPGAQNGLHGSGWAQSFGMKQGTPSDVYGSQATANNLQELVNGVREEFRLNHYQPQKQVQMQIDFSGASSG

PSVIGKPLSAESEHSDVEASCKEERPGTADDRKPRKRGRKPANGREEPLNHVEAERQRREKLNQRFYALRAVVPNISKMD

KASLLGDAISYINELQTKLKVMEAEREKSGSISRDASALDANTNGESHNQAPDVDIQASHDELMVRVSCPLDSHPASRVI

QAFKEAQITVVESKLSAANDTVFHTFVIKSQGSEQLTKEKLMAAFSRESSSLHSLSSTG

>Pt14G10370

MEGLIISPSSSSSLVSLSHETPPTLQQRLQFIVQSQPDRWSYSIFWQASKDDSGQIFLAWGDGHFQGSKDTSPKLSTTNN

SRMSTSNSERKRVMKGIHSLLDECHDLDMSLMDDTDSTDTEWFYVMSLTRSFSPGDGILGKAYTTGSLIWLTGGHELQFY

NCERVKEAQMHGIETLICIPTSCGVLELGSSCVIRENWGIVQQAKSLFVSDLNSCLVPKGPNNPCQEPIQFLDRNISLAD

GGIIAGLQEDDHTIEHGEKRTQERAETKKDNVNKLGQSYVDSEHSDSDFHFVAVNIERRIPKKRGRKPGLGRGAPLNHVE

AERQRREKLNHRFYALRAVVPNVSRMDKASLLSDAVSYINEMKAKVDKLESKLQRESKKVKLEVADTMDNQSTTTSVDQA

ACRPNSNSGGAGLALEVEVKFVGNDAMIRVQSDNVNYPGSRLMSALRDLEFQVHHASMSSVNELMLQDVVVRVPDGLRTE

EALKSALLGRLEQ

>Pt14G10630

MERLQGPINSCFLEEHSLDLGCFDQVFINTESLRFEEEEPHISSPSFEDKMPFLQMLQTVETPPFFPYKEPSFQTLLKLQ

HLQKPWNMNTFYMPETDTQVQPLELESCVTHDIVDLHSPVKSETKEHPNPHSNSCLEGVSPEPAEPNSDSSIPWRAQPQT

VPNMTTHFSESSTLIITRERRKRKRTRATKNKEEVESQRMNHIAVERNRRRLMNDHLNSLRSLMTPSYIQKGDQASIIGG

AIDFVKELEQLVQSLEAQKKIREIETASTAGISPNQYSTSQPQCDLLLEEGGTCEEERTVKKKSEATEIEVAAVQNHVNL

KIKCQRIPGQLLRAIVALEDLGLTVLHLNITSSQATVLYSFNLKLEDNCKLGSTDEVAAAAHQIFSSISG

>Pt14G11140

MDDRGHMELVWENGQVLMRVLPSTSSSCTSYTPHPKKNVSEVENNSDGYTTKRPRLGTGDSILGDFPLIDDRELAKRDKS

SQDDHHPELFSELCETNLNMLLENNENNIYEKNITDAHVVPGYKDANWRPGKASEFAAEVPQLTTASNGQLYQSFLEQHK

ASAPLFHGLPTSKLQQVDSGSDNHSRLQNLSRILRPALPKPSHGSNATRPTSGPGSSRLQQLKSNTDEPPAGCRNLVESG

QMVPTYASKVFKYFNDQQYLMASQIVPIGPIDRSAEASPPDEQSEAVLHNYATTSKRCCDRVFGSTSGSAEKKIKGKPDR

GKSIDQLTATSSICSRGASNDPTSSLERQYEDTEGTAYSSDDLEEEEQVPARGSAGSKRRRATEIHNLSERKRRDRINKK

MRALQDLIPNSNKVDKASMLGEAIDYLKSLQLQVQMMSMGTRLCMPLMMLPTGMQHIHAPLLAQFSPMGVGMDTRLMQMG

VGCSPATFPASGMFGLPAGQMLPMSVSQAPFFPLNIGGHSTHSSVPMPAMSGVASTPLEFMRSAVFPSSKDIIHSNTSAR

K

>Pt15G10420

MAGFSYQHQPLFPDSAFLPSIATPTKNMNNNMYGSFEEAGNMMNTNGFSQIYSPETFHETPSLDVRFHQSSHPDDHSYKV

SLSDNETSLTKKQSTNSSTVVDKLESEHVTQEVTPMARKRKSANGFLNSAQSKDARKVKSKRQNKCSGDMKHEEKKPKVE

KKVHGEPPAGYIHVRARRGQATDSHSLAERVRRERISERMKILQLLVPGCDKITGKALMLDEIINYVQSLQNQVEFLSMK

LASVNPLLYDFGMDRDAFMVRPERLSSMSPPLPSLQHNSPIQPTAFADTASATTATFATEENNYPLIDNSATLFLQGMRP

SDFTTHQDSGYLMWDVDEQRQKFLNPSGLTNNLCSFH

>Pt15G10520

MLSRLNSTSVWLEDHKEEQDSTTNHLHHHHNNINNTTAAGCGGVMLEGREEMGSLSTFKSMLEVEDEWYVSNNNNTIHQT

HQDSIKDLTFSPGLGDPDNLLLHQVDSSSSCSPSSSVFNNLDPSQVHYFMHPKPSLSSLLNVVSNNPLEHGFDLSEIGYL

ENQGTNSAATANVSALLNRGGGVLGNLGNFSDLSSNSQISIPNLCSDPQFSSSRMLQLPENGPGLTSFRGFDENSGNQLF

LNRSKLLRPLETYPSMGAQPTLFQKRAALRKNLGDNGGNLGLLSGIDRDKGKSEMTQISEENDKKRKFSSGDDFLEDVSI

DGSGLNYDSDEFTENTKVEEIGKNGGISSKANSGVTGGVDQKGKKKGLPAKNLMAERRRRKKLNDRLYMLRSVVPKISKM

DRASILGDAIEYLKELLQRINDLHNELESTPPSSSLTPTTSFHPLTPTPSALPSRIMDKLCPSSLPSPNSQPARVEVRVR

EGRAVNIHMFCGRKPGLLLSTMRALDNLGLDIQQAVISCFNGFAMDIFRAEQCKEGQDMHPDQIKAVLLDSAGFHGMM

>Pt16G03540

MQPENCSENSQLYRFLAENGMINVGAYGFPGAAMQTLCTSSSTSYHNNNYQFESSVITDMTPQDRALAALKNHKEAEKRR

RERINSHLDKLRGLLLCNSKTDKASLLAKVVQRVRELKQQTSELSGLETFPSETDEVTVLSGEYSSDGQLIFKASLCCED

RLDLMPELNEILKSLHLKTLKAEMVTLGGRIRNVLIIAADKDHSVESVHFLQNALKSLLERSNSSEKSKRRRILDRKLVI

Q

>Pt16G06850

MYVETACFEPNNSMVEDVTDDGFCHAIPLMAGNSTTNSFEEHLKLSMEEFSSHYPQEESAAAASMEEIQLQHHMAFSNNN

TNHHLMQQYPTQLLSYDHSSNWDPNIIQFQEMHQVLDQNSSFDATANTQSSLPPDLLNLFNLPRCTSTSTLLPNSSISFT

NPAHKAPLGFMGVDNTSARFDPYTLAPQPHLFRELVQSLPPHGYTLPTPLFGGGQGDDHVDGQSGGGLSYQDGDHGDGVF

EFTDEMACIGKGIKKTGKVTKHFATERQRREHLNGKYTALRNLVPNPSKNDRASVVGEAIDYIKELLRTVQELKLLVEKK

RCGRERSKWRKTEDDGGVEVLDNSDIKVEPDQSAYSNGSLRSSWLQRKSKDTEVDVRLIEDEVTIKLVQRKRVNCLLYVS

KVLDELQLDLHHAAGGLIGDYYSFLFNTKINEGSCVYASAIANRLIEVVDRQYASSTTTVPAAGSCY

>Pt17G10170

MVKSTKSHLDEEDEAEDYDSSSYKGEAEKAESKSNELKANANRSRHSETEQRRRSKINERFQALRNLVPQNDQKRDKASF

LLEVIEYVQFLQAKLQIYEGSYEGWSQEPAKLLPRKNYRASAESILGHTQVMKNGSAHENTVMLGNVHNSIKSDMDTAAM

YKTLDHSPGPTNPAIPFEVRTQSRVFAAVGRGGVPTESLQESVSDAENMAYQLQSQLLHGQPCATECITPNNTLNGQEDV

ASDSQSVNISNTYSKQILNSLTQALLSSGVDIAQTSITVQIDVGKRENGTTAVAPSSMVNQYLSNQLIIQDGVGSSVEDL

NQAHKRQRREKC

>Pt18G10950

MQPCSREMQGINSLLNPSSQIPLQDLQNQQNPSQIQNSHFDPNSSSNDDFLEQMLSNIPPCSWPDLKSPWDLTMPINNND

SSNSIAKPRDLSDETAPSNTDNSNLGFHNNFDESVILASKLRQHQISGGGGAAAAAKMMLQQQLLMAAARGVLPQNDVID

GSSFKGGDGSMQGLFNGFGAGSMNGTGQASNQSMQHFNHPQGGAMQAQNFGAQGAATTAVMNQPQASGSNGGAPAQPRQR

VRARRGQATDPHSIAERLRRERIAERMKALQELVPNANKTDKASMLDEIIDYVKFLQLQVKVLSMSRLGGAAAVAPLVAD

MSSEAGGDCIQASADGGSLSRTSNGNQTARTNDSSLTVTEHQVAKLMEEDMGSAMQYLQGKGLCLMPISLATAISTATCH

NRSPAINNNHHALLQSNGEGPASPSMSVLTVQSATMGNVGGDGGAVKDAASVSKP

>Pt18G14160

MESFQNISEYQNYWEMPSMFWNDELTSWEMDQASSQIYDSSSPDGAASASASRNTVSERNRRKKLNDKLYALREAVPRIS

KLDKASIIKDAIDYIQDLQEQETRLQAEIMELESERSEKDKGYEFESELPVLLTSKKTRYDHISDHREPRSDPIEVHQLR

VSSMGEKTLFVSLTCSKAREAMVRICEVFESLKLKIITASVTTVSGMVKKTVLIEM

>Pt18G14180

MEPIENIGEEYQNYWETKMFLQNEELDSWAIDEAFSGYYDSSSPDGAASSAATKNIVSERNRRKKLNERLFALRAVVPNI

SKMDKASIIKDAIDYIQELHDQEKQIQAELSELESGKSKKNQGGFGVYYHQELPVLLRSKKKKIDYQFCDFGGSKISPIE

LLELRVAYMGENTLLVSLTCNKRTDTMVKLCEVFESLGLKIITANITTVSGRVLKTVFIEADEEEKDKLKIRIEAAIAAL

NDPPSPMSM

>Pt19G08900

MEPCSWNSTRVQASDFEGMSDGFLVNTGLQAKTRTGPSSTSSLVLDNERGELVEATVRMERKGVAAERSIAALKNHSEAE

KKRRARINAHLDTLRSLVPGTRKMDKASLLAEVIAHLKELKRQATEASEGLLMPLDIDEVRVEQQEDGLLSAPYVIRASI

CCDCKPGILSDLRQALDALHLIIMKAEIATLEGRMKNVFVMSSCKEGDSGDAKVHQFLAGSIHQAFRSILDKFSASQEFL

LKSTLSNKRRRVDSFKPSLSSSSGDLW

>Pt19G08950

MKNVFMMVWMVCSEDLLGPSAGSVHQAIGILPCHLYSGESHDEARINAHLDTLGILVPCTESTDKASLLAGDINHLKELR

KKAAEGSEGLFMPLDVDEDLSIR

>Sl01G090790

MNQCVPSWDLDDSTVPRKNLIQTQSNSLAVDVPSLDYEVAELTWENGQLAMHGLGPPRANNKPISSYGGTLESIVNQATR

CNDDVPLHLHGKSTVDRNKQSGDEVVPWFNNHNAVAYAPPATGLVAMTKDALVPCSRNTSNSDNQRSVHVPGIDGSTHVG

SCSGATNSRDWTVAPRMRVRPTRREWSSRADMISVSGSETCGGDSRQLTVDTFDREFGTTMYTSTSMGSPENTSSDKQCT

NRTGDDHDSVCHSRDQKEGGDDEDDNDNKKGSKNSSSSTKRKRAAAIHNQSERKRRDKINQRMKTLQKLVPNSSKTDKAS

MLDEVIEYLKQLQAQVHMMSRMNMSPAMMLPLAMQQQLQMSMMGMGMGMGMGMGVAGVFDINNLSRPNIPGLPSFLHPSA

AFMQPITSWDNSNSAPSPPSAAMPDPLAALLACQSQPINMDAYSRMAALYLQFQQPPTGSGPKN

>Sl01G096050

MVTGNMLWSGEDKAMVASVLGKEAFEYLMSGSVSAECSLMAIGNDQNLQNKLSDLVERPNAANFSWNYAIFWQISRSKSG

ELVLGWGDGCCREPKEAEEREVKKILNLRLDDEGQQRMRKRVLQKLHMLFGGTDEDNYAFGLDRVTDTEMFFLASMYFSF

PRGEGGPGKCFGSGKYLWLSDALTSNLDYCARSFLAKSAGMQTIALIPTDVGVVELGSVRSIPESLELLQNIKSCFSSFL

SLVRDKQAAGIAAVPEKNEGNNPRLSNSGAVTERTDGNPKIFGHDLNSGTHFREKLAVRKAEERPWDMYQNGNRMPFVNA

RNGLNPASWAQFSNVKLGKPVELYAPPTPGHNLMNGGREEFRLNNFQHQKPAARMQIDFTGATSRTIVSPAHNVESEHSD

VEASCKEDRAGPVDEKRPRKRGRKPANGREEPLNHVEAERQRREKLNQRFYALRAVVPNISKMDKASLLGDAIAYITELQ

KKLRDMESERELRLGSTSRDAITSEDSPSSEIQIRGPDINIEAANDEVIVRVSCSLETHPLSRIIQIFKEAQINVVESKL

SAGNGTVYHTFVIKSSGSEQLTKEKLLAAFSSESNSLRQLSPVGQ

>Sl01G096370

MEQLAVSSSPMAVAPPPVDVNQVPLGLQQMLQYVVKSQPEWWAYAIFWQTSNDDEGKNFLAWGDGYFQGDGVVINNKGGG

GSSSSLKSQAQSERKKVIKGIQALMDGNGDTDLVDDGDVTDTEWFYVMSLARSFSAGDGSVTGKAFGSDDFLWITGPDQF

QLHYSCERAKEAQIHGIQTLVSIPTSNGVFELGSTQLIKQNLSLVQQVKSLFLCCPPIQFLEKTISFADIGLVTGLQQDD

NDYKLRENSRKPHPVVAKKRGRKPKGGEEDAHMAALNHVEAERQRREKLNHRFYALRSVVPNVSRMDKASLLSDAVSYIN

QLKAKVDELELQLIDHTKKPKIVTESSSADNQSATTSSDDQVIKAANPTAAPEVEVKIVGTDAMIRVQSENVDYPSAKLM

IALQNLQMQVHHASISSVNHLVLHDVVVRVPQGLSTEDELRTALLTSYDL

>Sl01G102300

MAIGKPESGQQKISSTSNLSSFPENDLVELKWQNGQIVMQGQNSSAKKSTVPNNLPSSASGDRDKYTGNSSTSKIGKFGL

MDSMLNDMSLTVPTGELDLIQEDEGVPWLGYPADDSLQQDYCAQLLPEISGVTANEQSGQSVFGLINKRGSSDKMIGDSH

SVPVHNAVNFERRNTSKVSPSSRFSPLSSLPSQKGHASIPTLESGVSDVFSSKNSNTPLSVLGESNQSKASAGDAKSNRI

QKQNMPGNRSNLLNFSHFSRPATLVKAAKLQSSTGGSNISGSPILEAKGKKGEEKVTIGDNHVSAAATENFLTSKKDNFP

HYPTNGVSSQLESRPSGASFHDRSCQAEQSDNAFRDCSSNNDNTHDHFTSAKATKDIADGERNVEHGVACSSVCSGSSAE

RGSSDQPLNLKRKTRDNEEFECRSEDVEEESVGIKKPCAARGGTGSKRSRAAEVHNLSERRRRDRINEKMRALQELIPNC

NKADKASMLDEAIEYLKTLQLQVQIMSVGAGLCVPPMMFPMQHMHGAQMPHFSPMSLGMGMGMGFGLGMLEMNGRSSGYP

MYPMPSVQGGHFPSPPIPASTAYPGIAVSNRHVFAHPGQGLPMSIPRASLGPLAGQPSTGAAVPMNVAREGVPVEIRGAQ

PNLDSKTPVHKNSQIVQNAEASCPQNQTCSQVQATNEVLEKSAQKNDQLPDVIGSAANRLTNRTNVPGNEAGPSL

>Sl02G076920

MNCRNLGEFCENEAKGVVQSLVLDSEKGELVKASGRVEKKIGKSEGKTIAALKSHSEAERRRRQRINAHLSTLRNLVPSS

DKMDKAALLAEVVRQVKQLKETATHDSERFFIPLDSDEIKVEIIAENAIDGTCLFRASVCCEYRTHLLSDLKQTINSLHV

NLVKSEISTLGSRVKNVFLFTNSIHGGGGCATIQARDIFLSSVRQAFSSVLDKVSAFPEYSAYPNKRQRVSCFDSSSLLF

>Sl02G091800

MDNSSLSQWFSRTEEGVFYSNRNNSIDDFTTQKSTISEDQETSELSITPDSARSNSRSYFHKEISKCKASKINFSLITPM

ENFPIGSKRPATDHPQQPKLNKISSSSHRFLSFNNNNNNTDSVFPSHQGCKIEAMDDMYLCDSSDQMFYPYVKSNGCYDA

KDKKGGKKIPSVELQDHIIAERKRREKLSQRFVALSTILPGLKKVDKASILEQAIKHVKDLKEKVQLLEEEKKSVMFVNK

YKVETEEYTSSEENNSGSDLPADIEVRFSDNNVLIRITCARRNAFVLNIHSEIEKLHLTIVQSSMMPFGKQAIDITLVAQ

MEESFCMTLKDVAKHVRMVTGRLMTQA

>Sl02G091810

MDTWWSEMETSNMNDHNFVNQSQAMTQFNEFPFNSKPFSTFTPPPPSNVISNYSYSNSMSTKNQFEKLNTFKVVKHEVPS

GTTINFSSSVNSMDDSDFGDIEAAMGFGAAITTTTDQKKSYNRTSVQAQDHVLAERKRRERLTQRFIALSTLIPNLKKLD

KATVLGDAIQYIKELEEQVKTLEEKNKKCSEEPVIPPAKRPRLVSSCADSSSSDEISSVSTVCTDRSLPDIEVRASDGNI

LIRIYCKKQNGMMKEIFNEVEKLHLSIISCSVMPFGYNTSHITIIAQMDHKLTSNTPNHVANRIRAAMVKEEANSFTA

>Sl02G091820

MDISSATWWSEMDVMNMNELQYIDQTFDDFAFSDNIQVSQIGEIFEEKPALCSSITQNTSTSPPCSSPSVISFSNSNSPS

ATPTTTNAQNYFKNLNTSSLKTEVPSGTTINFSSSNTSSDSDYDDSKQLFQAMGFGAGQSKKMNYSRTPLQAQDHVLAER

KRRERLTQYFVTLSTLIPNLKKLDKASILGDAITYIKQLEEQVKRLDEEANKQPVKRSRLHSNYDNFSTCNENSNKSVVP

EIDVRVSDGNVLIRVCCKKQAGIIKEIFSQVEMFQLTITSSSVIPFGYDTTHITIVAQMDHQLNMATEQVANNIRLSIMK

LINSHK

>Sl02G093280

MANNNVYYHNANFSLTDPDPEPDDISVFLRHILLPSSSSSSSSSNFMALKSNEMQYSSSLPHLMPNNNQQGNLSSMMNSS

ACGIFSSSYGVCNGATTVSSSSVGTIDYDPDEYECESEDGTEDLGAEASVQPPSRNTSKRSRAAEVHNLSEKRRRSRINE

KMKALQKLIPNSNKTDKASMLDEAIEYLKQLQLQVQMLTMRNGLNMYPLGLPRMLQQNQLSHQKVGLCEGNAFTNAKVAG

NLQVNQDASLNAIFNPTENCTETKVTPPITMSNINRSDSAFELESSMNIHLDPFQLSRSTSKEIWREDDLPLYGMNELTT

KTASTGSNLAFSVPLDTDASNLKRSTREACLLRYQFGAVNETNLDCDQLLSQQLYSNF

>Sl03G005350

MADNPPEVYAADDFLEQILAIPSYASLPVTDLTAGASSENSTSGVSQLQQQPLFPLGLSLDNGFADANNTGGFQVKTERE

AMNMGNLYPGLEHLQSHAVCLSVPQVHQVQPFQGHPTSSAIVTIPHQPAIRPRVRARRGQATDPHSIAERLRRERISERI

KALQELAPSCNKTDRAAMLDEILDYVKFLRLQVKVLSMSRLGGTSAAAQVVADIPLQSVEGDTCESHSNQRVWEKWSDSE

TEQEVAKLMEEDVGTAMQYLQSKSLCIMPISLAALIYPTQQSDNQSMVKPEQAAPL

>Sl03G007410

MDGDQNLSDLFDDSECDIFGILEALEGGGGGGGNSGITSKFNDNINNQTATIATTITTTTSDEITGLVSEEGKKRKLISQ

KSTGSCATLQEEETIENKISHITVERNRRKQMNEHLSVLRTLMPCFYAKRGDQASIIGGVVDYINELQQVLQSLEAKKQR

KVYSEVLSPRVLPPQLVPISPRLLTPSPLSPRKPPLSPRMNLPISPRTPQPTSPYKPNANANKPPEPSPTTSSNSSIDSH

VNNELAANSKSAIADVEVKFSGAGANVILKTVSPRIPGQAVKIIAALEQLALEILHVSISTIDGTMLNSFTIKIGIECQL

SAEELAHQIQQTFC

>Sl03G095980

MENSFSSEYCDVNDFLVQNYLPQCSHEGRESASRSHSEAEKRRRDRINAQLSTLRKLIPTWEKMDKAALLGSVVDHVKDL

KDKTAEISNVLNTPTDTDEVSIEHLNEEEDNKGCLIKASFCCDDRPELFSELQRGIKNLQLRMMEADITSLGGRIKCVFM

LSPNDNYVCINSLEKSLKAVLSRIAISPSTSNYRIKSKRQRFFLPPQFS

>Sl03G114720

MELPQPRPFGTEGRKTTHDFLSLYSPVQQDPRPPQGGYLKTHDFLQPLEQAEKTLREEETNVEVATVEKPPPPVAATPSG

EHILPGGIGTFSISYLHQRIPKPEASLFSVAQASSTDRNDENSNCSSFTGSGFTLWDESAVKKGKTGKENSGGDRHVLRE

GGVNTGGVQPTTSLEWQSQSSSNHKHNTTALSSLSSAHQSSPLKSQSFLHMITSAKSAQDDDDDDEDFVIKKEPQSHLRG

SLSVKVDGKGNDQKPSTPRSKHSATEQRRRSKINDRFQMLRGIIPNSDQKRDKASFLLEVIEYIHFLQEKVHKYEESYQG

WDNEPPKLPLSKCHRTTHGVSNLPQRIINASSASLTYAGKFDESIMGISSANPINVQKLEPNISSTGLKDKGQQPGLTNK

PTTVPMHPNTFSFSGTSSTAALYSSKLIADTDKLESKSHSQFSLSRSHMTDYAIPNANPERLELPIESGTISISSAYSQG

LLNTLTQALHSSGVDLTQANISVQIDLGKRANGRVNSSASTVKGDNVSTSNQPIPKSRVTTTREEPDHAFKRRKTS

>Sl03G118310

MGWIDGKEDGGTGSWVNQNNENHQQNNGGFPNENHQLNNGGFTNFQGMVDDGGVDWFMGGGDSNNHHINNNNNNGGGGGG

GSNMQSHISYSTSFTEAENSLLLQPVDSSASCSPVSGNVFNNIDPSQVNFFMPQKSTIPSSLTGLSNNPMDNSFNLGMLN

QAGNGMMNTGYHHLGSPNQMGTNNLSSYTQFSSPNLLQLPQVAGGYSSMGFGANNSANGNTLFLNRSRTHKPLDNFASIG

AQPTLFQKRIAKNLVSNGENLGTEIGQSSSNLTDKKRKSSMNDEFEDVSMDGTLNYDSDEFMDISNKMEDGIKIGDSSNA

ASTVSGADQKGKKKGPPAKNLMAERRRRKKLNDRLYMLRSVVPKITKMDRASILGDAIKYLKELLHDINELHNELESTPA

NNSSLSPATSFHPLTPTASALPSRIKEELVPSPLSSPTGQPARIEVRVREGKAVNIHMICSRKPGVLLSTMKALDSLGLD

IQQAVISCFNGFVLDVFRAEQSNEGQEMHPDQIKAVLMETAGFQGGTI

>Sl04G005280

MRLVLLRRMPHLLAVIPKVYGKSNEKASALRSKHSVTEQRRRSKINERFQILRDLIPHTDQKRDTASFLLEVIQYVQYLQ

EKVQKYEGPYQPWSSEPTKLMPWRNSHWRMQSLPAQPHALKNGSGPESTYLGRFDENLATVTSTMQPNQQNPIESHTSRD

VSFKALDQQNELANKSITTPIPLQAGMKMSVPNNSAFSEPQPRPVSDQCPNTIDALNHDEDDVIDGGRISLSSSYSQGFL

TSLSQALQSTGLDLSKATISVQIDFGKRANQAMTSGPSIAKDDENPTLSGHQHTDHFREASNDEDMNQAQKRLKI

>Sl04G006990

MMNSQEIMDAKALAASKSHSEAERRRRERINNHLAKLRSLLPNTTKTDKASLLAEVIQHVKELKRQTSQIAQTNPLIPTE

INELTVDYCNNEEGNFMIKVSLCCEDRSDLLHDLIKTLKSLRLKTLKAEITTLGGRVRNVLFITRDQQQDNDDTWPINDN

NNDNDDDQMKYCLRSIQEALKEVMEKSNGNDSGNSGSIKRQRTSNNNNIHY

>Sl04G007430

MLARFGVCYVAEIETNNGSGITSRRNEDGLYQYPKASCVEMHKMNERRRRYKIAKKMKVLETLIPNCNKSDRASVLDQAI

QHIQALQHQIQVMSMDRIRGSTLVAAGRNQIMQSTLHFNPYIGAIGYFFNFSNILCSNFSPMLSTGSEFPFLPLAVACNL

LHPGPIMEVFTRGSASVAPLEKRV

>Sl04G014360

MYQFPSFYELGNTCSDSYNNFLHEIITSSSSEMFNNINNLESSSVSPRSMAEAKAIAANKSHSEAERRRRKRINGHLATL

RNLLPNTIKTDKASLLAEAVRCVRELKQTTSELGATTTTTTMSENDDDDDHTTLMTKIMFPSESDELNLSYCNESNNNNT

DNNNDDRNLIIKASMCCEDRPEIMMELRRALSTVEGKIVRAEMSTVGGRIKCILWLEMLENGCKEGLFVQLRRALKVVMD

KANFGPQNMGQDLLGNNKRPRLLGGPINYAT

>Sl04G076240

MLSSLQGCNNVFLQLVDHQKNEKSNTTKKSKSKDAAVTHAVAERKRRERINSHLHTLKKLFPHLPKKDKPRVLTEAVTQL

KELRKNVAQQLELSSLFIPSENDVVIINYCDNINDERTVKTTICCEDRPSLNRDLSSAIQSVQGRVIKAEMATVGGRTKA

ELVVVLGKANGGEKDVGQLKRALKAVVENRALGFGSNVMLGRRFG

>Sl04G077480

MGTKENGSFNCPSTGMNRADSMPNVDPFSGSGWDPLLSLNQKGGFKGSSVVGHNEFVNLPYQSSQFVHYPSDSNLAEMVP

KIPAFGNESYSELVNTFPLQEQLRGANCYANYVKNRGISTEGECQISGEGAVEVSPNGKRKISENHSLSNANKNVEGELQ

KAPSRDSSDCSKEQDGGKRHKTDQNVSSNLRNKQAGKQVKDDSDGGEPPKDNYVHVRAKRGQATNSHSLAERVRRERISE

RMRLLQELVPGCNKITGKAVMLDEIINYVQSLQQQVEFLSMKLATVNPELNFDIDRILSKEMLHQQTSNAALLGLGPGLS

SSLPFPGISHGSFAGIPATTPPFHPLPQNVWDNELQSLLQMGFDSTSSMNNMGPNGRSKLDL

>Sl04G078690

MADPYRTNPHASSSLESEDMSSFFLNFLQGTPASSSATAAAGFYNRSGPAPVAESSSSLNFSDPGRFYAAEFKEGVENVF

ASAGLGECDGMNSANRREFLEDDKVDNFGFSSEECDGLDMPSDPTHPRSSKRSRSAEVHNLSEKRRRSRINEKLKALQNL

IPNSNKTDKASMLDEAIEYLKQLQLQVQILTLRNGLSLYPGYVPGSLQSVQLPSGNEFDGRSFMLSANGGATLPVNREMP

QTAFEISNQNPSGKPTITSHNTENAVALETTIQNHYGLLNHLASSKDMCRDNTLSRLHLDMSCSGNNSSSGVSS

>Sl05G005300

MEKALEWLRPLVDSKNWEYCVVWKFGDDPSRFIEWMGCCCSGANGVDVNVKRENGGKQTFSSLCRDIQVQHPIRTKACEA

LAHFPHSISLYSGIQAEVVTSNEPKWINHAEISNSNLSHELKGTLVLIPVAGGLVELYNSKMIYKDQKTINFIINRFKLG

SEEANSSVAQKEDQVLDFFPYEKSNFCAPLLQYATSFPSSAPHISQVSESSANPSIQGSSTGSIPSNELTLCHSPPDHLS

RNVPLSQSTEGYFEHTELQCSGNLSRMEDTIFPWKQENYIVAGDMFSMGKKRQKGPYQSKNLVTERKRRNRIKDGLFTLR

ALVPNITKMDKVAILGDSIDYINELQEKVKLYKIELNKIEAEVTNNESTPEMVLSDMTEMSKVTGQTNEKTQISVNTTDR

TRMEVEVNQIGAREFLLKVSGSRKPGGFTQLMEAMNYLGLELVNVSCTTSGGEIVSVYIVEANVDRFIDAQKLRSSLIEL

TS

>Sl05G009640

MDDLALIFSHHNNGESSRKQLHKEKEDEKANSLACIISFNNNFENSLSIPKDEANNTISFSNINNIIGAKFGERRSGEQA

LEHLLAERKRRKRISKLFVSLASLIPGLNKMDKASILEGAATLIRQLGERAKEDDHHQSTIGMMTKNNLLPEVEIKSLEK

ELLITILLYKNQQKRNIDEILSVIQRLHLTIKTTNFMPFGTTSMHITVIAQMNDEFCETTDFLAEKLRSLISKV

>Sl05G009880

MLPWSIPPVQPFMNPVHHHDQSFLLPPSPSAYGLFNRNTNTDQQHLRFISDSLVGQVVHHHHHHNQPGSIAPFGLQAELQ

KMSAQEIMDAKALAASKSHSEAERRRRERINNHLAKLRSLLPNTTKTDKASLLAEVIQHVKELKRQTSLISETSLVPTEI

DELTVDNATSDEDGKFIIKASLCCEDRSDLLPDLIKTLKALRLKTLKAEITTLGGRVRNVLFITGDDYYCNNNNNREVDT

CISGDDEDTEMMQQQQQQQPQYCISSIQEALKAVMEKSSGDDSASTSVKRQRTNNINILS

>Sl05G050560

MTMLWSDEDKTMVAAVLGTKAFDYLMSSLVSAECSLMAMGSDENLQNMLSDLVERPNASNFSWNYAIFWQISRSKLGELV

LGWGDGCCREAREGEESELTRILNIRLADEAQQRMRKRVLQKLHMFFGGTDEDNYVSGLDKVTDTEMFFLASMYFSFPRG

QGGPGKCFTAGKHVWLSDVMRSSVDYCSRSFLMKSAGMQTVVLIPTDIGVMELGSVRTIPESLELVHSIKSCFSSFLAQV

RAKQAAPLAAVVAEKKNGNNSVFPSSFPFDQSKENPKIFGQNLESGSTEFREKLALRKPVDGPLEMYRNGNRAPIINTQN

GVRPVSWASFGNVKPGNSVDLYSPQAPPNNLREFVNGGREELRLNSLQHQKPGGMQIDFTNSRPVVSPVPTVESEHSDVE

VSCKEKHAGPADERRPRKRGRKPANGREEPLNHVEAERQRREKLNQRFYALRAVVPNISKMDKASLLGDAIAHITDMQKR

IRDAEYKLEKRGSTSVDAADINIEAASDEVIVRARCPLGTHPVAKVVEAFKETQVSVVESKLAVGNDTVYHTFVVKSSGP

EQLTKEKLMAAFAGESNSL

>Sl06G008030

MNYCVVPDFKMDDDYYDIPSAIFNKKSTIADEEIMELVWQNGGVIMQSQNQRSVRKSNLFPEQSAVEQTVAVSAPLYMQE

DEMNSWLQSPLDDSSFDDFLNTTPSCDAVTSAAAAPPGEIGTSKVEIRPPLVPPCSRPIRCTEGELPHRLQNFGHFSRLS

GEAVLRNGTTSSSGHSVRASTIVDSNETPVAARVSENVTPVTAMNVRGRELTATSMATTSGGREVTMACELALTASTRGS

GGSVSARAGPPQPSHTEADTAAYDRKRKSRESDDNEGQSEDVEYEFADARKQVRSSTSAKKSRAAEVHNLSERKRRDRIN

EKMKALQELIPCCNKSDKASMLDEAIEYLKSLQLQVQMMATGCSMVPMMYPGIPQYMPTMGMNMSMGMDMKMGRNRPLIS

YPPLMPGPAMQNAAAAAQMAPQYPLPAYHLPPFPAPDPSRIPVANQPNSHVGHNISQPRLPNFSDPYYQYFGLQQAQLML

PQNQEVEQLSSSKLNSCIEGSRGNHQSGEHTI

>Sl06G051260

MLRSIQLREMMDDSSCSKYTHETLSSWCHNLQELPLIPTNSCCWSQQFSASRSHSEAEKRRRDRINAQLSTLRKLIPTSE

KMDKAGLLRSVVEHVKDLEGKAKEMSNVLNTPSDIDEVVIEEEDESSNNNNIVVKVSFSCDDRPELFSELNRGLKNLKLT

TMEAKITSLGGRIKCILSLQSINVVCTTHSIKHSLRLLLARIATSPSTSNFRIKSKRQRFFLLAT

>Sl06G064590

MESNYYDSFQGMEYSGNYNYEDESFFEFKPNISQGIYDLSSAYNTNIEEKSGQKSISSNNSSSNSGGFLISFSSNQEEDI

GAMISSENSCQESFLLGENNNNNNNNNNVMYKRSPQQAQDHVIAERKRREKMGDLFISLSKIVPGLKKLDKSSILGDTIE

YMKELQEQVKLLEESKKNTSSSLEHNDSNKEQVLGSNKIKVRIMDKNVLINIHCNKQDGMLGRLLVQMEQLHLSVHDMRI

MPFGPTNLEISLLAQMEDGCCINVEDIVKAIQINILDLVNN

>Sl06G068870

MLSRVNSMQVWDMEGKQEEEKEENFSNKENTTNVELENKQDMELGALSTFKSMLDGTDVDWYHNNMQNHTENICFTQNFT

ELAENSMFLQPVVPVDSSSSCSPSSVSVFNNLDPSQVHYLLAQKAINNNPLDYSFNLGCENGFLEAQGMGGLNKGGFLLA

GGGFHDLSSQNQMGNPNLNSFTQYPSSHLPQNTTTTGFSPLGFVDGSANENSLFLNRSKLLKPLDNFASNGAQPTLFQKR

AALRKNLANTTGGSLGDFGGEIGQNSMNGENERKRKWGSGEELDDVSFDGCTLSYDSDDLTENVTNKVDDTVKNGGNSSN

ATSTVTCGNQKGKKKGLPAKNLMAERRRRKKLNDRLYMLRSVVPRISKMDRASILGDAIEYLKELLQKINDLHNELESTP

PSSSLTQTTSFYPLTPTGPALPGRIKEELYPSSFASPLSSPTGQPARVEVKAREGRAVNIHMFCSRRPGLLLSTMRALDN

LGLDIQQAVISCFNGFALDIFRAEQCKEGQDFHPDQIKAVLLDSAGCHGMI

>Sl06G069600

MSHHTWNFSHQKQEQQVVEKEEEENRYTRGHVHNQQNQVDPMSNKCEVAELTWENGQVAMHRLGSNLSNEQTKHTWGKAG

DTLESIVHQATFQKQHHSYIMGSDGQNQANINREKNVSYGAQQTRGVLKRMRSSDSDPQLYIGGISLEHLNARASAKDND

ITMITWPCNEDSACHGGSENKEEERETKSSNPSKRSRRAAVHNQSERRRRDRINEKMKALQKLVPNASKTNKASMLEEVI

KYLKQLQAQIQLISYAKNMEQQMMMMSLGMQPAHIQMPLLATMGMCSSTTGILNNMTSNLAPAPYQSLIGGRAPLIYPTS

SMPTLFPPFMSPPFATASSIPSTPPQPINAESISPKLTKYAAPPNIAASTSFPFSHPYNAYLPHSMKMEFNNEMAAQYLQ

RGNQENVNIQGQKK

>Sl06G072520

MEANSNSFHVDSVFHVPIKMSGFFEEPNNNITSSSTLPNCVSQFYLQELSVNMSNNVHEISHNEPSHVTNKTNSSSLCST

QSKNVRDGDDGKGQKKRNGNVKREKKTKENKKKAPEEAPTGYVHVRARRGQATDSHSLAERVRREKISERMKILQALVPG

CDKVTGKALMLDEIINYVQSLQNQVEFLSMKLASLNPMYYDFGMDLDALMVKPDQSWSGLEGPLLENTTSNYPHLDSSTS

LMFQQLHLPNSVSQGSGHVLWSVDDQRQKMIINHSELISNNNNLSVPFH

>Sl06G083980

MAEKFFLKGEDKVNMEGVLGSEAVEFFSWSASNHMLTEFTSSRGDLGVQQALCKIVEGSDWTYAIYWQVAKSKSGKSALI

WGDGHCRETKIGQGEGANDSAHQKMMDGNKKKMVLQKIHTCFGGSEDDNIAAKLESVSDVEVFYLTSMYYIFPFDKPSSP

SQSFNSARSIWGSDLKGCLEHFQSRSYLAKLARFETLVFVPLKSGVVELGSVKSIPEDQNLIQMVKTSVVVSNPPQPKAN

TKIFGRELSLGGAKSGPISINFSPKVEEELSFASDSYEVQAALGSSQVYGNSSNGYRSDEGEGKLYKEELDERKPRKRGR

KPANGREEALNHVEAERQRREKLNQRFYALRAVVPNISKMDKASLLGDAIAYITDLQARIRVLDAEKEMVGDKQKQQVIL

EIDFHQRQDDAVVRVGCPLNAHPVSRVLKTFQEHQVVAQESNVSLTENGELVHMFSIRAPGPAAEDLKEKLTAALSK

>Sl07G005400

MNSLLSQQQQSQISLQDLQNGGNGGSTGGVGGLSQHSMGHSHFDPTSSHDDFLEQILSSVPSSSPWPDLSKSWDPHHHLS

SPPHNPSSGEDQPPSNPFHSQFHYDDQASSLLASKLRQHQITSGGGAAAAAKALMLQQQLLLSRTLAGNGLRSPNGASGD

NGLLSLPLNLSNGDQNDGVANPTNDNSVQALFNGFTGSLGQTSNQPQHFHHPQGGSMQSQSFGAPAMNQTPAASGSAGGG

GGSTPAAQPKQQRVRARRGQATDPHSIAERLRRERIAERLKALQELVPNANKTDKASMLDEIIDYVKFLQLQVKVLSMSR

LGGAPLVADMSSEGRGEGNVGRGGNGRASSSSNNETMTVTEHQVAKLMEEDMGSAMQYLQGKGLCLMPISLATAISTSTT

RISNNPLLAPEAGGSTSPTLSALTVQSATAGKDATSLSET

>Sl07G018010

MDNIGDEYKNYWETTMFLQSEELDSYFDEPISSYYDSSSPDGSQSSMASKNIVSERNRRKKLNERLFALRAVVPNISKMD

KASIIKDAIDYIEELHNQERRIRGEISELESGRSSSKKNSNDVEFEQDESFDSKPKRSRRFEMQYGYDSSGSTTRSPPSS

SPVDVLELRVSSMGEKTVVVSLTCSKRTDTMVKVCEVFESLNIKIISANITAFSGRLLKTAFIEADEEERDLLKLRIETA

IASLNDPDSPMSS

>Sl07G039570

MDSIFFLEEGDRTVFLLKIMESFGCTYICLWQYFQPSNTFMSLGGIYNGENVVAQRLFEEYKHSWLIMDNGRIPGLAFKN

NVPYMELKFADLQSHASNPVQLQFYYTTICMGCSIGEIEFGMTSSPQVNLEMGMKNLFPEYFSTRLVLARPQTLLTNIDQ

NRPSPSSSFSLDSPGEYSSLLFNVATTSYVPDAFPEQTVRPVSTSAMPFHQQQPIQTLTQLRGIQFPGVETDDAALTRAY

LAVMTSPSSSSSSHQSRENIDVPITDYHYQKSTAFRRFGPGLGRPSNVQIGTSRTIRRENILRRSIIFFRNLDMMRRKEQ

IQANQRALTSTQVHHMISERKRREKLNDSFQLLRSLLPPGTKKDKASVLASTTEYITCLKDQVEELSKKNEIMLNAQALD

KSSMMKSNDVGDGNDERVVVEIIKNVSSESESRTVELQVSVRSGECNVLDLATRLLEFLKTQDNLSLQSVAANTRPSMVT

HVSLTITIQGSEWDESGFEEAVKRVVDDLT

>Sl07G043580

MGFDHELVELLWRNGEVVLHSQTHKKQPGYDPNECRQFNKHDQPTIRVAGNQTNLIQDDETVAWLNCPIDDSFDKEFCSP

FLSDISTNPHLGEEPDKSIRQSEDNNKVFKFDPLEINHVLPQSHHSGFDPNPMPPPRFHNFGSAQQKHHIVGGDQKGVNF

PPPIRSSNVQLGGKEARSNLMLQDIKEGSVMTVGSSHCGSNQVDTSRFSSSANRGLSAAMITDYTGKISPQSDTMDRDTF

EPANTSSSSGRSGSSYARACNQSTATNSQGHKRKSRDGEEPECQSKADELESAGGNKSAQKSGTARRSRAAEVHNLSERR

RRDRINEKMKALQELLPHSTKTDKASMLDEAIEYLKSLQMQLQMMWMGSGMASMMFPGVQHYISRMGMGMGPPSVPSMHN

AMHLARLPLVDPAIPLTQAAPNNQAAAMCQNSMLNQVNYQRHLQNPNFPDQYASYMGFHPLQGASQPINIFGLGSHTAQQ

TQQLPHPTNSNAPAT

>Sl07G053290

MEFLSSFNGLNEVYGGFQGIIGNGLSSSSSLVLDNESGELVKAMVKPGGKGVNPXXXXIALKNHSEAERRRRERINGHLG

TLRNLIPGTNKMDKAALLAKVIGHIKELRVNAAEATKGVLVPTDIDEVKVEQQAEGSDGATYSVKASLCCDYKHELISDL

RQALDTLPLKTLRAEIATLGSRMVSVFVITEGNEGNTEGTERCQLLITSVRQALRSVLDKFYASEEFSSRSTLSSKRRRV

SLLNSSSSSSLGDFW

>Sl08G008600

MENLNISTSSTPSQPNTLQKTLQYIIHNRQEWWVYAIFWQASKDVNNRLILSWGDGHFRGTKDTTGSTKTGHGQYHQFQK

KFGFNDISETNNNVTDTEWFYMVSMPQCFVADDDLVIRAYTSASHVWLASYYELQIYNCERAKEANLHGIRTIVCISTTS

GVVELGSSDVIQENWEFVQFIRSLFGSNNNMNTTSHLPVNQVTLGDDHKVAKCGSNIIVKQEMTIGNLLSESGISDFEND

DSLTINNVMNGSIKRAKKGDSSHIRREMAMDVHVEAERKRREKLNHRFYALRSVVPYVSKMDKASLLGDAVTYINELKAK

IKNLESKLIEPQKKHILMEQHDSHSASSTIVTDHGANNKSLFSSNGVRNGMEIEVKIIGSEGVIRVQSLDMNYPCTRLMN

AMKEMKFQIYHASISSVKDLMLQDIVIRVPEEFSNEETLKSAIISKLSVMEN

>Sl08G062780

MERLRPIMSLKGWDYCVLWKLSEDQRFLEWICCCCGGAEKNMHGCGQEIFFPDSSTSTCRDVMFQHPTTTACNLLAQVPP

SLALDCGVYAQTLLSNQAKWMNFVPFSESNISNEIMGTRALIPSPLGLLELFSTQQLAEDEKVIEFVSAQCNIYLEQQAM

MNSTFSNGVEENNTSKPFPTEGERDRDDHIKDSQNHYKQRVSPAATSDHLSFDFPLKRKQLDSCSMNFLPPFSTYSTPEV

DNNTGGNMLFDQSTSDMTHFSENRYMSEMDAYLQKQMMRSSSTQAGIDDESIKHDNGRSNSGSDSDQNEEEDDPKYRRRN

GKGPQSKNLMAERKRRKKLNERLYALRALVPKISKLDRASILGDAIEYVMELEKQVKDLQLEVEEHSDDDGTGGGRNSDQ

IHPVVLSHNGTKNRPKSDNGKLTNGSQREISTNSNGSTDPSRKNQDVEENDKLQQMEPQVEVAQLDGNEFFVKVFREHKA

GGFVRTLEALNSLGLEVTNVNATRHTCLVSSIFKVEQKRDNEMVQADHVRDTLLELTRNPSRGWSEMGRASSDNINNNNA

NGTTDYHQHQLHDHHLDNNNQHKQTNSHHFHTHHHH

>Sl08G076930

MTEYSLPTMNLWNNSTSDDNVSMMEAFMSSDLSFWATNNSTSAAVVGVNSNLPHASSNTPSVFAPSSSTSASTLSAAATV

DASKSMPFFNQETLQQRLQALIDGARETWTYAIFWQSSVVDFSSPSVLGWGDGYYKGEEDKAKRKLSVSSPAYIAEQEHR

KKVLRELNSLISGAPPGTDDAVDEEVTDTEWFFLISMTQSFVNGSGLPGQALYSSSPIWVAGTEKLAASHCERVRQAQGF

GLQTIVCIPSANGVVELGSTELIVQSSDLMNKVRVLFNFSNDLGSGSWAVQPESDPSALWLTDPSSSGMEVRESLNTVQT

NSVPSSNSNKQIAYGNENNHPSGNGQSCYNQQQQKNPPQQQTQGFFTRELNFSEFGFDGSSNRNGNSSVSCKPESGEILN

FGDSTKKSASSANVNLFTGQSQFGAGEENNNKNKKRSATSRGSNEEGMLSFVSGTVLPSSGMKSGGGGGEDSEHSDLEAS

VVKEADSSRVVEPEKRPRKRGRKPANGREEPLNHVEAERQRREKLNQRFYALRAVVPNVSKMDKASLLGDAISYINELKS

KLQNTESDKEDLKSQIEDLKKESRRPGPPPPPNQDLKMSSHTGGKIVDVDIDVKIIGWDAMIRIQCNKKNHPAARLMAAL

MELDLDVHHASVSVVNDLMIQQATVKMGSRHYTEEQLRVALTSKIAETH

>Sl08G081140

MAMGHQDQDGVPGNLRKQLALAVRGIQWSYAIFWSTAVTQPGVLKWIDGYYNGDIKTRKTVQAGEVNEDQLGLHRTEQLK

ELYSSLLTSESEEDLQPQAKRPSASLSPEDLTDTEWYFLVCMSFVFNVGQGLPGKTLATNETVWLCNAHQAESKVFSRSL

LAKSASIQTVVCFPYLGGVIELGVTELVTEDPNLIQQIKNSFLEVDYSVILKRPNYVSNDAKNDTNIGSQKPDHNALEND

AYPVEINSPHDSSNGFVANQEAEDSLMVVDGIGETSQAQSWRFMDDNISNGANNSLNSSDCISQNNANCEKLSPLSSGEK

ETKPCPLDRQENDQKKPHLLDHQGDDAQYQAVLSTLLKSSDQLTLGPHFRNMNKKSSFASWKTDIQMPRFGTAQKLLKKV

LLEVPRMHAGVIHKFSRENGKKNSLWRPEVDDIDRNRVISERRRREKINERFMHLASMLPTSSKVDKISLLDETIEYMKE

LERRVQELEARSARRSNDTAEQTSDNCGTSKFNDIRGSLPNKRKACDMDEIEPESSNGLLKCSSADSIVINMIDKEVSIK

MSCLWSESLLLKIMEALTDLHMDCHTVQSSNLDGILSIAIESKSTGSKTLAVGTIREALQRVVWKS

>Sl08G083170

MQDFISTSSSFSSTSLLQKRLHYIIHNRQEWWVYGIFWQASKDANGRLIFSWGDGHFRDLALAKVHNANVSDMEMFYAVS

APNCFLSEDDLIVHAYNSGSYVWLNNYYELQIYNYDRAKEAHLHGIRTLLCISTPHGVVELGSSQVIQENLELVQLIKSL

FGQINDHGFNFVPLGDPMDTKTITMGSDSGNSDESSAMNKDSPKKRARKSTTAKNHVEAERQRREKLNHRFYALRSVVPN

VSKMDKASLLADAVTYINELKAKVEELKAKIEVSTKKLIQKRNCVSSSAVVDGTNINININSSFVDGMEVEVKIIGVEAM

IRVRSPNVNYPCARLMNVLRELEFQIHHATVSSMKEMMQQDVVIRVPHNVTNEEAIKSVILTKLSFA

>Sl09G063010

MNHSVPDFDMDDDFSLPASSGITRTKKSAMAEEEIMELLWQNGQVVMQSQNQRSLKKPHIGNGSGGGGDAVIPSDQAVSR

EIRHVEETTPHHLFMQEDEMASWLHYPLDDPSFERDLYSDLLYPTPTSTFTTAALPRENRTSTFEIRPPPPQPSPAAPIG

TAPRPPIPPSRRTVTENSNRFQNFGHFSRLPKARLEPGQANLSKSPRDSTVVDSNVTPITGQESRVTLIPDNVVAVPGGN

VGCSTVNGSTGTATASTAIREPTTTCDISMTSSPGGSGNSVSASAEPPAPAPAPSHKGAAPTATAAADDRKRKGREMEDE

GQNEDAEFESPDTKKQARGSTSTKRSRAAEVHNLSERRRRDRINEKMKALQELIPRCNKTDKASMLDEAIEYLKSLQLQV

QMMSMGCGMVPMMYPGMQPYMPPMGMGMGMGMGMDIGMNRPMVPYPPLLPGTAMQNAAAAAQMGPRFSIPQFHLPPVPVP

DPSRMQASSQPDPMLNSLVSHNSNQPRLPNFSDPYQQFFGLQQAQVALPQNQAVEQPSNSKSGSSKEVGNPGNHQSG

>Sl09G065100

MEIIQPNSLQLQNMLQNSVQSVKWTYSIFWQFCPKQGVLVWRDGYYNGAIKTRKTVQPMEVTAEEASLHRSQQLRELYDS

LSAGDSNPPARRPSAALSPEDLTESEWFYLMCVSFSFPPPIGLPGKAYSKKHHIWIMGANDVDSKVFCRAILAKTVVCIP

LLDGVVELGTIEKVQEDIGFIHRVKSFFNEPQQAQPPKPALSEHSTSDPAAFSEPHFYFSNTPSSAGICPADQDGRITGE

EENEDEDEDEAEDDEDENDEAELDSDGIAIQSGAGAANPMAAEASELMQLDMSEAIRLGSPDDGSNNMDTDLYLDGISQA

GNTADSFKAETAISWANFQDLQHLPGIPSYDELSQEDTHYSQTVSAVLEHLSNTSSKFASSATIMGSISPDSAQSAFTLW

PVTCSPNLSHCRRHDIGDGSGTTSQWLLKSILFTVPFLHSTKKLSEALSPKSRDAAAADSSAAASRFRKGCTINSCTQQE

ETSGNHVLAERRRREKLNERFIILRSLVPFVTKMDKASILGDTIEYVKQLRKKVQDLEARDRHTEITKKSDEKSGSPIVK

AFPVKGKRRMKSTVEGSIVGAPAKMTGSPPMEEEVLQVEVSIIENDALVELRCPYKEGLLLDVMQVLRELKVEVVAIQSS

LSTGLLLAELRAKVKENIYGRKASILEVKKSINQIIPRVN

>Sl09G083220

MAANQPEGYADDFLEQILAIPPYSGLPVADVGTPSETTSFTSASAVSHLNSAAAAGLQQPLFPLGLSLDNGRDDVGDAGP

YAVKHERDGMNIGNLYAGLEHLQSHAVRHSVPSVHHVQPFQGPPTTSTTVTVPHPPSIRPRVRARRGQATDPHSIAERLR

RERISERIKALQELVPSCNKTDRAAMLDEILDYVKFLRLQVKVLSMSRLGGASAVAQLVADIPLQSVEGDSGESRSNQHI

WDKWSNVDTEREVAKLMEEDVGAAMQYLQSKSLCIMPISLAALIYPTQQPDDQSLVKPEAAAPS

>Sl10G009270

MDEPIFSPSSSQSSLQHRLQYIVKNQTNYCSDWAYIIFWQSSNNRSCLTWGDGHLNMKITNNKDVEWFYLMSLAQSFCVG

EGVVGKCFSSGSLVWLAGDQQFEFCHCERAKEAHYVHGINTFVCIPISSGVLELGSSTMIKQDLNLVQQVKSMFFGYETI

DQFDDFGLFNCLELYGEEAKKGEVVVGTTPHENKAGLKNKTSKKRRREICETQGNHVEAERQRREKLNSRFYALREVVPN

VTKMDKATLLSDAVTYITQLKAKVDELESKLHSNNYHYYYPEMKIKHKMENHDINVVDNQSSITTSRDHTMEIEVKMVGQ

DAMIRVQSENVNYPSTRLMCALQEVELHVYHANISSVNDFMLHDIVVKVPQGLETEDEVKYALLRSLDQQTCS

>Sl10G009290

MDELMVSSSSSSSSFFIPSLLSQTSSNLQQKLQNILKIQTDSWSYAIFWQTTNDDDDGHLFLAWGDGHFHGTKSKTGVQS

SEQSTERKNVIKGIQALICENGDEKVDDDDDDEVTDAEWFYVMSLAQSFSIGDGVPGKAFSTASIIWLTGSQNLQFHTCK

RAKEAHLHGIQTFVCIPTSNGVIEMGSNQLIKENWVLIQQVKSIFNNSIPHIVNCLEQNTNINPKTEELVSVSVSAECND

SDSDCQLLVEKKTPKKRGRKPGATRETPLNHVEAERQRREKLNHRFYALRSVVPHVTKMDKASLLSDAVSYINELKSKVA

ELETQLTRKSKKLKIECTDSFSIDNNSTATTITNSVDQIRHNSFGVHSNLKVEVEVKILGPDAMVRVQSENVNYPSTRLM

RALQDLELHVHHASISSVNDIMLQDIVVKVPIGLSTEDRLKNALIRSIQHQ

>Sl10G018510

MDYEVAELKWEKGEVVMHGLGPPGVPCYYKPLSTPSPTKYTWDDKPHAAAGGTLESIVNQATTHNIDIGDEGGDDDDLVS

WFDDCLPETSMDIVAVVPTSCTNYNQQVPPSTRVASCSGDAEMARVGMGSSFEEISEDFENQEAKNLIGSMVYEGKNNTV

SPGETSLGEERVLTTTSTFKHNKRKTLNNHDSRGQESRDNEDEDEKKRSKISSFSTKRCRVAATHNQSERKRRDKINQRL

KTLQKLVPTSSKTDTASMLDEVIEYLKQLRAQVKAMSMMIHVNMQPPPMMLPNMAFQQQQQQFQMSMMGMARPIDVNALS

SPNITTIPSILHTTAPSNFNNPPIASPGADPLASLVAVRQLSQPMTMDAYSRMAALYQQYLQSNANLGFKN

>Sl10G049720

MEHVVSMSEGDWSSFSGMCFTEEADFMAQLLGNCSFPNELPSNSGYWNIGHESNIGSSGGREHSSFFFPPLSHESHYSSN

SRPILMRNDSSITTERGLMDTNNPIEADEYFANNMEFDANMAEPLLDGKGLQLGRIDYEDHSPTESSKKRVRLHCHVPKN

KRSTKLQTEGKTVEMDMKSKAVLQRQNSMVSCCSEDESNVSLELRRKSRASRGSATDPQSMYARKRRERINERLKTLQSL

IPNGTKVDISTMLEEAVQYVKFLQLQIKLLSSDDLWMYAPIAYNGMDLGLDLRIGNPK

>Sl10G049780

MEPVFEDEWSGMCSTDQEADFMAQLLWQPNNIDNMYFSSYNGCNSSQISFPSLESYYQSHVQSILTRNGSSITKENGMVE

GENTSSPSQVLATYNPIEADDFLNQDVSMESGENTAKVLDSPPESSKKKRLCNLLGDVPKNKRSVNLKKASEMDGKNKAA

LQRQNSISCCSEDESINVPKSRASRGSATDSQSLYARKRREKINQRLRILQSLVPNGTKVDISTMLEEAVQYVKFLQLQI

KLLSSDDLWMYAPIAYNGMDIGLDLKIGTPKS

>Sl10G078380

MQQQQEDFQENLESCNYFSKNKSSSSDNNMMGEYYGFDIEAIKNFCSSSPFYPQHNHVEFQETIESNNNSPESRAKNHKE

AERRRRERINSHLHSLRTLLSCNSKLDKASLLAKVVQRVRELKEQTSQIMQSETNLLFPSETDEITVLSSNDCLADGRLL

IKASLCCEDRFDLIPDLIETLKSLGLSPIRAEMVTLGGRIHNVIVLAVDHKKESNNNNTDHDESSLLFLRDALRSIVQRS

SYGTGERGKRRRVLGQGTRNY

>Sl12G010170

MQAMNSFQSTGENGASSGEHMSHSHFDPSSSHDDFLQQILSSVPSSSPWPEISGDGHPYNFDDHQSTLLASKLRQHQING

GTSAAAAAKALMLQQQLLLSRGIAGNGGSGINGDQNDDGLNSGNDISVQALYNGFAGSLGQTSNQSQHFHHSQAQSFGAP

AASLSMNQTPAASGSAGGAQPKQQKVRARRGQATDPHSIAERLRRERIAERMKSLQELVPNANKTDKASMLDEIIDYVRF

LQLQVKVLSMSRLGGAAAVAPLVADRSSEGGGDCVQGNVGRGGSNGTTSSANNDSSMTMTEHQVAKLMEEDMGSAMQYLQ

GKGLCLMPISLATAISTSTCHSMKPNNPLLLAGGSAINGVGETGGGPSSPTLSASTVQSATMGNGGT

>SM00000G06060

MGEKSRITLRQRLQAAVQSIQWTYAVFWKPCPPPQGELVWSDGYYNGSVKTRKTIIVSRERSPEEHGLQRSDQLRELFEN

LSASGDGSQSSTATRRPTAALSPEDLTDTEWFYLVCMSCTFDPGTGIPGQAFSKGRPVWLCKANEATTKVFSRALLAKTV

VCIPMAEGVLELGSTELSWKEFQRQASISSSQVKSQCQAMLKNVLFRVPHIQSKSVSRKEDDVNTAHAMLERRRREKLND

RFLMLRNMVPFVTKMDKVSILGDAIEYLRQLQRQVADLEQRNKPEDSFPMSTTYKLGPDSSSYKAEIQMQDDFTALEIEC

SFRQGILLDILAALDKLNLDVSTVEARTPDQRTFCASLKAEVSLQAFKVFSFTTPMVSLFYRRKICREQASKI

>SM00000G10450

MYDVPLVCVRRFNFVREVFIPANQEPRRRIIMQHFGATTLYEDHNYFARSSESFTLEGGLDSDNGTGLLDLTDTLEKQLE

ASNLALGEIHSDAPQLSSELHVPSVEFELDDNGQLLHDMLQQQQPANSQQQTTANTTTAASSTATATTWEENVHVLQEML

MQQQHIQVQQQNFFGNSTPPKFRRESELLSLLQLPRCASSSLPGGTISGPFNSVSSKKLSHQGALSSFTTAAEIGDSSHV

HGQQLLPQPSFRHLLHTLPHASDHHLSSKSGIMRPPGLLDLEDRDTSGGHEEGRLFPSLFEKRDFTFGKGAENRGINHFA

TERQRREYLNEKYQTLRSLVPNPSKADRASIVADAIEYVKELKRTVQELQLLVEEKRRGSNKRRCKASPDNPSEGGGVTD

MESSSAIQPGGTRVSKETTFLGDGSQLRSSWLQRTSQMGTQIDVRIVDDEVNIKLTQRRRRNYVLLAVLRSLNELHLDLL

HANGASIGEHHIFMFNTKIMEGTSTFAGQVATKLIDAVDRHITLASSGL

>SM00001G07620

KNLMAERRRRKKLNDRLYTLRSIVPKISKMDRTSILGDAIDYLKELQQRIETVYTDLQSPVMSFASKQKLLFEEELQTSV

TFPMECWEPQVDVQTSGANAISIHMFCEQRPGLLLSTMRALDGLGVDVQEADIKFTNGFQLEIYAEQSTKKLASPEEIKA

VLMHTAGYFNPVMPC

>SM00004G02810

MRPPGLLDYEDRDNSAGHEERRLFPSLFEKKDFTFAKGAESRGINHFATERQRREYLNEKYQTLRSLVPNPSKADRASIV

ADAIDYVKELKRTVQELQLLVEEKRRGSNKRCKASPDDPSATDVESTTAMQQPGGTRVSKETTFLGDGSQLRSSWLQRTS

QMGTHIDVRIVDDEVNIKLTQRRRRNYVLLAVLRSLDELRLDLLHANGASIGEHHIFMFNTKVVLAPSSFLLSLFLFYFF

LFSSALILPR

>SM00006G01050

RPRVRARRGQATDPHSIAERLRRERIAERMKALQDLVPNANKTDKASMLDEIVDYVKFLQLQVKVLSMSRLGSAAAVPSL

VADLPSEGANSLLASTLSRSTGISHDGLASAERQVARLMDEDMGSAMQYLQSKGLCLMPISLA

>SM00012G04610

MGGHQAADKGPSGRGNRKGLPAKNLMAERRRRKKLNDRLYMLRSVVPKITKMDRASILGDAIEYLKELLQRINELHSELE

GPADGGSMGIPPQQQSGALLSPQSFAPCVKEECPASSISPLPLLPGPPTDLQPAKVEVRTRDGKGINIHMFCARTPGLLL

STMRALDDLGLDVQQAVISCFNGFVLDVFRAEQCSDAEIAPEEIKAVLLQTAGCHEAL

>SM00018G02850

MTDLLSLCKSSAQADNAKALAASKSHSEAERRRRERINKHLSTLRTLLPNTAKTDKASLLAEVIERIKELKQQVAEISQF

GPVPSDADELDVDVMESPVDEGGKVLIKASICCADRPSLLTDLVRTLKSLHLRTVKAEMATMEGRTKNVFVMTIKDDAEL

LEPTLACVEEALKSVMEEPSSKENQDDA

>SM00038G0a0580

MSKMTPQEIIDAKALAASKSHSEAERRRRERINNHLNTLRGLLPSTTKTDKASLLAEVIEHVKDLKRQAAEIAEGGPVPT

DVDELKVDTDASSSDGNFVLKASLCCEDRPDLLSDLTKALRTLKLRTLKAEIATLGGRVKNVILIGKDHSDEQGGAAMES

SSDGTGDRPSVSCVQEALRAVIER

>SM00038G0b0590

PKGSKRSRAAEVHNLSERKRRDRINERMKALQELIPNSNKTDKASMLDEAIEYLKLLQHQLQVVCP

>SM00047G00310

GESVDVKKAPPARTSSKRSRAAEVHNLSERRRRDRINEKMKALQELIPNSNKTDKASMLDEAIEYLKMLQLQLQVLSPGS

SKVSS

>SM00055G00880

MVPNSYFRRDANPESMSGALKMEHDGLYWGQQQQIDPLGQNVYPVQASGSSQGTSIQFSASPSSSTHLQALRDFMSQGNS

TGSVPLYSQQLVPLPPPLLVPLGEDSSHFSRSALPDVDMRGQYRGDVFGSEKTPHRTQLQRESHILAERQRREEMNDKFS

SLRAMLPKSSKKDKASIVGDTINYVVDLEKTLKRLQACRAKRKGCHIPKEKSLKSSPSSDPKLEASKTDTVQRLPVQVEV

QALGEQAVVKLVCGKSPKLVLRILTALEQCKVEVLQSNVTTLGDIAVHFFTIELTPGVSATTEELIAALEAAAAAMNSST

S

>SM00059G00730

AESVDTKKPATRPKRSRAAEVHNLSERRRRDRINEKMRALQELIPNSNKTDKASMLDEAIEYLKMLQLQLQVPKIELLHS

SHTFSCSCR

>SM00068G01040

MAKAIVPGLQERLRSLVNGKGWDYVVYWKLNDDQRFIDWVGCCCGGVTAGHGIGADFFSPQPHYLDGGAPCPDISFPHPS

TKTCTILSAMTLSPSIPLDSAGVHAQVLLSGQPRWINFSLESNQHEGVQTKVYIPIQNGIVELGSSSQIAENAMVIQSVK

AKCGDPWQDFQGFQENVDQQGFKSLYKNQEDGLDKHFNGHGDQVHWTSALDTHPWEHDVGFSDNQFSLNIAAPTQLNFLG

QPSKTGGQPSDHFDKQVDCSRPEKQGPPFVQGLQDVPPLAPPNHSFSESPHGSGVSKENSEVKQETRADSSDCSDQVDED

DEKATGRSGRRHLSKNLVAERKRRKKLNERLYSLRALVPKITKMDRASILGDAIEYVKELQQQVKELQDELEDDSQAANN

IPTMTDVCGGGHKHPGSEGITIADVDTNKCALKADDINDKKVEDLTQPMQVEVSKMDAHLLTLRIFCEKRPGVFVKLMQA

LDALGLDVLHANITTFRGLVLNVFNAEMRDKELMQAEQVKETLLEMTSQPEARSSSSLVSESHMGVITIDGGSGSDCSCK

SVEYNNVANFTILEDGYNFKPGVSRQLLDCKDHNKRNAIARWLNMEYGRHEIEDNNPTYSYFVEDLQCFEITSFNGWGHF

RTTNGWYGWSFWAFTGSFIQSSNHLSVDFL

>SM00073G01190

MLVAASFQNQVGGGGAASTSQAGSRNHQGLAATLDHSDNTGQVSSWSHHFVPGNALSFAKLGDGGASSDPNSVKKRYRDE

ELQQMYGAAGQMQIDQQQKHQQDQQQRSQAPLAYDSNAPVAPVNPPPAGISARPRVRARRGQATDPHSIAERLRREKIAE

RMKALQELVPNANKTDKASMLDEIIDYVKFLQLQVKVLSMSRLGGAGATMAPLVADLPLEGAGQELVSSSQLCRQISVNL

SPQDGIALTEHQVARLMEDDMGSAMQYLQSKGLCLMPISLATSISSCSTKVPTTMSLAATAAAAAAAGSSPTTAAIAGAL

QHATLSDSEASAMEGCNGRAATKDFPEQSEEQWLK

>Zm01G02650

MELDEETTFLDELMSLRREAAAAASAQWQAAAAYTGGGGQGMMMGDLLFYGGTTEAAATATSGGGMDQLSPFHQQEPLQV

QAPPHPHSHEEFNFDCLSEVCNPYRGGGVAVITGAEAAAVPGRALAALHDAMVEDETSGHLQGQQYGAGGSPTFVFGGAA

GESSEQTPVVVRGAAGGPHHRSKVHGGGAPSKNLMAERRRRKRLNDRLSMLRSIVPKISKMDRTSILGDTIDYVKELTER

IKVLEEEIGASAEDLDLLNTLKASSSSGSNEMMVRNSTKFDVERRGNGSTRIEICCATNPEVLLSTVSALEVLGLEIEQC

VVSCFSDFGMQASCSQEEEDGNNKRQVLSTDEIKQALFRSAGYGGRCL

>Zm01G10400

MGGHGDHHRRRRQEVEVLVHDDDEDLEHQSRACGGGATSGVVVEQGAGDGGGQAASSSLGSVSAATMAPPQIFCWPPQHH

HNSSNDVGGGQQPPFFPPLPPLPPTPPPFFADLYARRALQLAYDHHHPGGGPSTSSDPLGLYMAGGGSGMMMMPPPFASS

LPFGDFGRMTAQEIMDAKALAASRSHSEAERRRRERINAHLARLRSLLPNTTKVLILIIIDDITHARIE

>Zm01G10410

MRARVTSRTCMLQTDKASLLAEVIQHVKELKRQTSEITEEEACPLPTESDELTVDAGSDEDGRLVVRASLCCDDRADLLP

DLVRALKALRLRALKAEITTLGGRVKNVLLITADDDSSAHDGQREEEEEEAPMSPQRTVASIQEALRAVMERTASAAAEE

SGAAAASTGGAAGLKRQRMTSLSAILENRSI

>Zm01G22080

MNLWTDDNASMMEAFMASADLPTFPWGAPAGGGNSSAAAASPPPPQMPAATAPGFNQDTLQQRLQAMIEGSRETWTYAIF

WQSSLDSATGASLLGWGDGYYKGCDEDKRKQKPLTPSAQAEQEHRKRVLRELNSLISGAAAAPDEAVEEEVTDTEWFFLV

SMTQSFLNGSGLPGQALFAGQPTWIASGLSSAPCERARQAYNFGLRTMVCFPVGTGVLELGSTDVVFKTAESMAKIRSLF

GGGAGGGSWPPVQPQAPSSQQPAAGADHAETDPSMLWLADAPVMDIKDSLSHPSAEISVSKPPPHPPQIHFENGSTSTLT

ENPSPSVHAPPPPPAPAAPQQRQHQHQNQAHQGPFRRELNFSDFASTPSLAATPPFFKPESGEILSFGADSNARRNPSPV

PPAATASLTTAPGSLFSQHTATMTAAAANDAKNNNKRSMEATSRASNTNHHPAATANEGMLSFSSAPTTRPSTGTGAPAK

SESDHSDLDASVREVESSRVVAPPPEAEKRPRKRGRKPANGREEPLNHVEAERQRREKLNQRFYALRAVVPNVSKMDKAS

LLGDAISYINELRGKLTSLETDKETLQTQVEALKKERDARPPSHSAGLGGHDGGPRCHAVEIDAKILGLEAMIRVQCHKR

NHPSARLMTALRELDLDVYHASVSVVKDLMIQQVAVKMASRVYTQDQLSAALYSRLAEPGSAMGR

>Zm01G29230

MQTAIEHACSVVECAATARAAMDMSHYIPDWSSSMGDTFAPLGGEDDDGLIELMWRNGHVVMQAQAPRKPPRPDDDEAAA

AQAQAWFQYPVEERADLFSELFGEAQAAVGGARGEAARQSIRMMPPPPPPPRPAQAPREEKACPGDGGTATATDGAGSSV

LTVVSSLCGSNGNHVQATAPGDVARARDVLMVTSSSTTRSRSCTTKSEQPGPGPGAARRSGKRKHNDATDAEDVGLECEP

AQRTTTAKRRRAAQVHNLSERRRRDRINEKMKALQELIPHCNKADKASMLDEAIEYLKSLQLQLQVVWMGGGIAAAGVHQ

RTMVAAPGRPPHVASLPASAPDLYTRYLAVDHLPPPPLVPPPRTAAAMGLYPRQNPVPATSSPSFRTTENTRKLWQA

>Zm01G30250

MMDRHMEDDSSTFLQWAMNHLQHPAAATAVSAAYQQQDGGAGVGVRISGGAGDQEDSAAAFPSLQALRASQPQAVPGSVR

VRNLTVQVADYGLTNSSSSGDSPGAAMDHDAAAGWSPHTARSRTTGLGGGSNSRPVSWNFSAASAQPADDRGAVGVALPD

APAAVARAQQRAASSAGRRGGGAGPSTAAAPASSPGPVQDHIIAERRRREKINQRFIELSTVIPGLKKMDKATILGDAVK

YVRELQEKVKGLEEEGGAGGSGGIQSAVLVKKQLPPEDDAMASSHGGSGDHGGDGGGMPLPEIEARLSERSVLLLRIHCY

SARGLLVRVISEVEQMQLSITHTNVMPFPASTAIITITAKVEDGFNATVDEIVRRINSALHQHYSSSSEETRG

>Zm01G30260

MMDRHMEDDSSTFLQWAMNHLQHPAAATAVSAAYQQQDGGAGVGVRISGGAGDQEDSAAAFPSLQALRASQPQAVPGSVR

VRNLTVQVADYGLTNSSSSGDSPGAAMDHDAAAGWSPHTARSRTTGLGGGSNSRPVSWNFSAASAQPADDRGAVGVALPD

APAAVARAQQRAASSAGRRGGGAGPSTAAAPASSPGPVQDHIIAERRRREKINQRFIELSTVIPGLKKMDKATILGDAVK

YVRELQEKVKGLEEEGGAGGSGGIQSAVLVKKQLPPEDDAMASSHGGSGDHGGDGGGMPLPEIEARLSERSVLLLRIHCY

SARGLLVRVISEVEQMQLSITHTNH

>Zm01G34530

MDELVCPAASSCSSPSPTSFFAAGHVPELEFVSWDVPEEWMEGTDWFDEPLDGDEGSRSAGNDLSGEPPAPAPKRRRGRK

PGPRTNGPTLSHVEAERQRRDKLNRRFCELRAAVPTVSRMDKASLLADAATYIGELRDRVEQLEAEAKQASAAVTTAVAA

ASHSFAPLQEKLGLEVRMVAGLDAAALRLTTSAARHAPAHLMLALRSLDLQVQHACVCRVGGVTVQDAIVDVPAGLRDER

CLRAALLQRLQPSG

>Zm01G35380

MVATDGEGERSTPAPAPAPALRKERGRSHSEAERKRRQRINAHLATLRTLVPSASRMDKAALLGEVVRYVRELREKASDA

AAGVGLGVIPGEGDEVGVEEEDGCRWRPAGRHHGAGGIGTDADVSQPPPRRVRAWVCCDDRPGLLSDLGRAVRSVSNACP

VRVEIATVGGRTRSVLELEVCDDGDDGSATAAGNGRAVALSTLRAAMRAVLLNRDEHVVAAAGEGYKRPRFSAQIARVQ

>Zm01G38350

MWEGGGSHAHEALLLQAAGSGAADYGHAGPSLLRPWLGPAAASGFSYMAPNHAQPGPLGAEAAVASRFGFGGGGYSDGGV

EQEQFVVIGSAETVPPRHSLQAAGVGGRTTALLPHGPRMVSGLLGTLQAELGRMTAKEIMDAKALAASRSHSEAERSRRQ

RINGHLAKLRSLLPNTTKTDKASLLAEVIEHVKELKRQTSAAARQRHLLLPTEADDLFGGRRGGRRRRQARRAGVAVLRG

PRGPHPRHRPRARRAQPAGPPRRDRHA

>Zm01G46770

MNQFVPDWSNMGDTSRPLGEEDDLIELLWCNGHVVMQSQSHRKVPPRPEKAAAVAAPPAPASVPQEDEGGLWFPFALADS

LDKDIFSEFFCEAPTPAPAAADAAPAASGGGTGTEAGGKSCGGDVPVPAEDDRRGGGGACAVSAGDPCDLMPPPKSTPAS

CSRQQTTMSLANGGDNAGGDLPGLVRAGAEAGASSMLSAIGSSICGSNQVLVQRAACAPGRASASGSGTARGDGSGSAAL

PSAVGSANANAVGGGRGHEASSSGRSNYCCFGAATTTTTTTTTEPASTSNRSSKRKRLDTEDSESPSEDAESGSAAMLAR

KPPQKMTTARRSRAAEVHNLSERRRRDRINEKMRALQELIPHCNKTDKASMLDEAIEYLKSLQLQVQMMWMGSAGIAAPP

AVMFPGVHQYLPRMGVGMGAAAAAALPSMPRLPFMAPQPVVPSAPVSVGPVPAYRGHMPAVGITEPYGHYIGVNHLQPAP

PPPQVQGVSYYPPPLGATAKAVQQAAELHHVPGPGGSIMPAGAAPGVLLPESAQGRGPGTVPCAPPFSSSASVFGLQMGA

LR

>Zm01G47860

MEDSSQFMQWALSTLQHDELPPATPPAAIAYDDNDCNTFSSVPALGYSAASVNSMVPAEPPAQEGHRATAATNSWSSADT

ASVTAAQRDAWSPSSQQNSVNCATPRSSGSSQPVSWDFHSASASASAQLIIKEAQVNSATAARAESAAGGMPVPQPQMVQ

NGSPPTRRASAKMSSASSSAPPCSQDHIVAERKRREKINQRFIELSAVIPCLKKMDKATILSDATRYVKELQEKLKALQQ

GGSCNARGGTESAPVLVKKPRIAAPGDDDKDRGGAPSPSCAPPGAAATTGNALPEIEARISDGNVVMLRIHCEDGKGVLV

RLLAEVEGLRLSITHTNVMPFSACILIINIMAKVAEGFNATADGIVGRLNAVLAAGPTC

>Zm01G47880

MEDSSLFLQWAMSTLRHEQQPAAAVNDDCSSEATFPSLQALREASHAAEMVQELIGEAPANSWSSGDTTDGSIGGNSNSV

PGPAAAAMEHDVWPAASSKTSPARRALSRSSSDTNPPVSWNFSAAASAQLAGSADGMLPEFAPKSALPPDQAYGSPRARR

AGLKSLAGSMSSAAYAQDHIIAERKRREKINQRFIELSTVIPGLKKMDKATILSDATKYVKELHGKLKDLEAGGSNRRKS

IETVVLVKRPCLHAAPAPDDDASPLSASSGTPAETKTQLPEIEARFAENSVMVRIHCEDGKGVAVKVLAEVEELHLSIIH

ANVLPFVEGTLIITITAKVEEGFTVSAGEIVGRLNSALLHNNEHNRCCNIADE

>Zm01G47890

MEDSSLFMQWAMDTLQHENNPAPASAAAVHGGFSEATFPSLKALREASHAAEMVQELIADADVVRHAPNSWNSGDNTTAN

YNVPAGWGFGGATAASALPGSHGMMEAPPAMATRGRPPPGLVYRLPPTRRAGLKSLGSMAAAYAKDHIIAERKRREKINQ

RFIELSTVIPGLKKMDKATILLDATRYLKELQEKLKDLEQRKEAGGGSIETLVLVKKPCLHAAAARDDDGGSSLPASPPA

GTPTEGKRLPEIEVQFSELEKTVAHTCDTIHSFVLCGDTTPSPRHSEVRGGEAAPSAHHANASGGESSPSPALVLDNVDR

RPRVVLAAAPANACDVVSPSPHASGSAEPSRACELAIRSRASVSGKLPHFLRLSRNALKSKPRRKVLLYFKAAAQIFLGV

EVVHGGAVSASDALLRWGWGIITCVGARERTPGMALGSGGGVRALPGGWQINAGAAAVEEGFTVTADDIVGRLSSALLRI

TSIGGCKRP

>Zm01G51370

MATQWFSNMVMDEPSFFHQWQSDTLLEQYTEQQIAVAFGQGEPVDHAVAAALATAMPMQLQQPAAAEQPHHRPRKAAKVN

TSWDSCITEQGSPAADSSSPTILSFGGHTAFAKAEPTTHQPPSCAGYYGAAAAAAKAPKQEMMDAAAMPPFQQARPAKRS

YDDMAAVAEAANAPSANTRPASQNQDHILAERKRREKLSQRFIALSKIVPGLKKMDKASVLGDAIKYVKQLQDQVKGLED

DARRRPVEAAVLVKKSQLSADDDEGSSCDENFVATEASGTLPEIEARVSDRTVLVRIHCENRKGVLIAALSEVERLGLSI

MNTNVLPFTASSLDITIMAMAGDDFCWSVKDIVKKLNQAFKSSF

>Zm01G54900

MSRVNGVTDALSIGPSAGYILFLVIVHILLFFTAYDRSYTLLAIQKVTLLCSACKAPRTDGGDDSQGSSARARQQQHAAV

LRSESGEKTLFSGHSSRSPDEMDGNARSAATNQKKPVVADDDLVELLWHNGSVVAQPQAHQRPAPTYDPDRPGTSGLTGE

ETAAWFPDTLDDALEKDLYTQLWYSTIADAAPQHEGTLPGPTSQPSPSPPVGSSGVESSWAGDICSTFCGSNQVLRTPAR

IRGKDAALQSELPSNTGAHDGTSSSGGSGSNYGGFGLPSDSVHVQKRKGRCRDDSDSPSEDAECEEASEETKPSRRYGTK

RRTRAAEVHNLSERRRRDRINEKMRALQELIPHCNKTDKASILDETIEYLKSLQMQVQIMWMTSGMAPMMFPGVHQFIPQ

MALGMNPGCIPAAQGLSQMPRLPYMNHTLPNHIHLNSSPAMNLMNPLNAANQVQIGHLRDASSHLLHLDGGRAAVVPQVP

GPGPHVHGHQIAQAEEHNKILEVAASTVIPTSKAGQPPTLHRV

>Zm01G57430

MFPVEVSALAAGGAGAGRQAAGDAAGTMLPPFFMGSIWPATAGAAGSEEDEAAAAAAAHDRALAASRNHREAEKRRRERI

KSHLDRLRNVLACDPKIDKASLLAKAVERVRDLKQRAAGVGEAAPAHLFPTEHDEIVVLASGSGAVFEASVCCDDRSDLL

PDLIETLRALRLRTLRSEMATLGGRVRNVLVLARDVVDGGGGAVAGGDDGYGGRAADSAGGATDGGGGDFLKEALRALVE

RPGAAGDRPKRRRVSDMNMQAAA

>Zm02G04290

MEDGSAPRRSTPPTRRSRSAEFHNFSERRRRDKINEKLKALQELLPNCNKTDKVSMLDEAIDYLKSLQLQLQMLVMGKGM

SPVVPLELQQYMHYITADPAQLPPLRPSGQQHRQFQITQANPQRQSNVESDFLSQMQNLHSSEPPQNFLRPPKLQLYTPE

QRGGLPNTSHNTGWISGSSSYNFME

>Zm02G07330

MALSACPAQEELLQPAGRPLRKQLAAAARSINWSYSLFWSISSTQRPRVLTWTDGFYNGEVKTRKISHSVELTADQLLMQ

RSEQLRELYEALQSGECDRRAARPVGSLSPEDLGDTEWYYVICMTYAFLPGQGLPGRSSASNEHVWLCNAHLAGSKDFPR

ALLAKVPEDPDLINRATAAFREPQCPIYSEQPSSNPSADETGEAADIAVFEGLDHNAMDMETAGIAVFEGLDHNAMDMET

VTAAAGRHGTGQELGEADSPSNASLEHITKGIDEFYNLCEEMDVQPLEDAWIMDGSNFEVPSSALPVDGSSAPADGSRAT

SFVAWTRSSQSCSGEAAAVPVIEEPQKLLKKAVAGGGAWANTNCGGGGTTVTAQENGAKNHVMLERKRREKLNEMFLVLK

SLVPSIHKVDKASILAETIAYLKELQRRVQELESRRQGGSGCVSKKVCVGSNSKRKSPEFAGGAKEHPWVLPMDGTSNVT

VTVSDRDVLLEVQCLWEKLLMTRVFDAIKSLHLDALSVQASALDGFMRLKIGAQFAGSGAVVPGMISQSLRKAIGKR

>Zm02G17840

MDMEHQLLHLERFMASSPGFFTVDSGQDPAHFPNGGLFVEQVGATGGVGGDDGGWVEDLMQLGEELFGGRGDANGTDDGM

GDDHYYQQWQCDDGGSPDGGPPPSVSLDGDASPPSGEQEQGTVELASERHPHREAEDGDDDVLGATRKRRDRSKTIVSER

KRRVRMKEKLYELRALVPNITKMDKASIIADAVVYVKNLQAHARKLKEEVAALEARPRSPTGQHSGPAGAGRRRHQQQQQ

ERRRDAGRSAGSGARVTHVGAVQVGEGRFFVTVECERRDGVAAPLCAAAESLACFRVETSSVGGRSGPDRVVSMSTLTLK

GRGQLGDAAAIGEASVKLWMMAALVKEGFRPAATVQIS

>Zm02G18610

MEADSMVAMGCGGFYWQSPPRFLLEPLDLADIVDSSMYVPTNEADEPVSGLCSSNRPAEDYSSSAEGANSCSAAVVPSPP

PPPGVTTTTTRNMDMERTRRRKLNERLYALRSVVPNITKMDKASIVRDAIAHIEHLQEQERRLLAEISVLQSSDDGTAAA

AAVKTEDAAATGGAAYDVDSVPWRKKPRAVPLPSVYFTDNPTSSISSSPPVRILEQVQVSQAGERVAVVSLWCSRGRDAV

GKICLALEPLRLRVVTATITARGDTVFHTLFVETGETGGARLKEAILAALARLNVLANADQVVHELLG

>Zm02G30480

MWDGGVEHGSQEAAHQLLPWLGAAPFSEPAAVAGLGAAMGAYACDGVGGLGHGGVFGFGFDAAVQQQQQQQRAADGSGKA

VVVSGLLGSLQAELGRMTAREMMDAKALAASRSHSEAERRRRQRINGHLARLRSLLPNTTKTDKASLLAEVLDHVKELKR

QTSAMMAATDADADADDEGAGRTQAQAQAQLLPTEADELCVDAGADGAGRLVVRASLCCEDRPDLIPDIVRALAALQMRA

RRAEITTLGGRVRSLLLITADRRADVQRGGGGGGGDGDDDEEEEDEEEGGERAASHRRHECIASVQEALRGVMDRRTASS

CDTSSSGGGGGSIKRQRMNYGAQEQCSV

>Zm02G30550

MQPTTREMQAMAAAGQQISLDDLRAAAGGVHDDFLDQMLGGLPPSAWPELSSAAGGKAPDGGAQAEQMQHQPQHLGGGGG

LYDDSALLASRLRQHQISGGGGGEAVKQMVLQQLADLRQEHHVLLQGMGRSTSTGGSRDGGLLLPLSLGSGGSGGDVQAL

LKAVADSAGGEAAGVFGGSFAGSLHHHQQQQQHFQSHPQQTAPLPGQGFGGGAGASGGVSQPQAGAACGGAAAPPRQRVR

ARRGQATDPHSIAERLRRERIAERMKALQELVPNANKTDKASMLDEIVDYVKFLQLQVKVLSMSRLGGAAAVAPLVADMS

SEGRGGVAVAAGSDDGLAVTEQQVAKLMEEDMGTAMQYLQGKGLCLMPVSLASAISSATCHMRPPVGGPGLGVAAAAAHH

MAAMRLPPHAMNGGAGAGADAVPASPSMSVLTAQSAMAPNGAGGGTDGEGSHSQQQQRHHPKDAASVSKP

>Zm02G31910

MQMDKAALLGEVVRHVRELRGEADAAAAGAAVAVPGEGDEVGVEEGHQHRFCHGGERAARRVRAWVCCADRPGLMSELGR

AVRSVSARAVRAEIATVGGRTRSVLELDVGGRHHDGEGTSTSSRPALQAALRAVLLSREEMLGAECYKRQRFSAHLARVY

DSGAVEVTD

>Zm02G32300

MGIQGNKAATHEHDFLSLYAAAAAAAAAKDAPLLLHDSKTPPPSQGNFLLKTHDFLQPFDQKPGAPPEPSPLPAPAAEIR

HRQQVAVAKQRALPLPLPGGVGTFSISPAPVSVALPSAVVKSEPPPFVLWGQPAATLQPGARGHQQQWALPFAGSLQVSP

PPQQAQPDRKGRGGGGVMDSGSRSSGGAGFDDEDGLTTRREVSSSLQDLAVRVDWKGGSCSDGGTNQGPNTPRSKHSATE

QRRRSKINDRFQILRELLPHNDQKRDKATFLLEVIEYIRFLQEKAQKYEATFPEWNQENPTMLPWSKGQIPGDSPPDPSH

FVRNGSSPGSNFTGKLDDNHTSAAASGALDQAETNHVASGCYRSSETPANITNNAISQSQPQWTGPSPVDDSAVKSEMLN

SQELAIDEGMISVSSTYSQELLISLSHALQSSGVDLSQTSISVQINLGKRAGKRSASAGGVSSNSKEPPADPASSNELGG

HHVTTLGASADGLLPHATKRHKRDNS

>Zm02G33730

MDDLLSPCSSFSPPSPPPFFSHAGNPVIEFASCEVPEQWLLDDVVLDKNEGRYDDVDDLWPVAGRSLSPDSELSEQPLPT

QPVPLPPPQQQQQELTSVTAAPAQQQRPGGKRRGRKPGPRPDGPTVSHVEAERQRREKLNRRFCDLRAAVPTVSRMDKAS

LLADAAAYIAELRGRIARLEADSRRAAAARWVDPVAAAASCGADEAVEVRMLGPDVAAVRATSAAPHAPARLMSALRSLE

LHVQHACVTRVNGMTVQDVVVDVASPLQVQDDDHDGLRAALLQRMQDSAAT

>Zm02G42680

MLPPFGNPLWVPEDMDDQQQHAPPPTPMELLTVPAQGHEEQNLLALASAAAVAGAGCVFSSPAMLDDDWYFDPVAAAAAT

GAQGHLLLAPPGPVPGPGAGSQMFSLFNVGGAATFDVHGFDIGLGTLGGGSGGDLVPLAGAGNTSNSASFSMSLNAGLLA

SSFGGFGTAPAQMPDFGGLGGFDMFNNGAGSSSAAPPPASASLTVPFSGRGKPAVLRPLETFPPVGAQPTLFQKRALRRN

GGGEDYDKKRKAEATAAAAGASSACGGDDAEDDDGGSIDASGLNYDSEDACRGVEDSGKKDGKGSNANSAGDGKGKRKRL

PAKNLMAERRRRKKLNDRLYMLRSVVPKISKMDRASILGDAIEYLKELLRKIEELQNEVESSASPASTASLPPTPTSFRP

LTPTLPALPSRVKEELCPSALPSPTSKQPRVEVRTTREGREVNIHMLCARRPGLLLATMRAIEGLGLDVQQAVASCFNGF

SLDIFKAELCKDGPALLLLPEEEIKSVLLQSAGLHGVAP

>Zm02G43850

MPLHLAAMQGRPAYMQAPVSCPECDCDCSLCCWVRLIRPEADGPLPRDFSSVPFRLVGIMVMKMEHEDNGAIGGTDGTWT

EDDRALGAAVLGADAFAYLTKGGGAISEGLVATSLPGDLQNKLQELVESESPGTSWNYAIFWQLSRTKSGDLVLGWGDGW

CGEPRDGELGAAASAGSDDSKQRMRKRVLQRLHIAFGVADEEDYSPGIDQVTDTEMFFLASMYFAFPRHAGGPGQAFAAG

IPSGFPIRHEAERSPGLAKIFGKDLNLGRPSVGLAVGLSNSKVDERTWEQRSAVGGTSLLPSVQKGLQNFSWSQARGLNS

HQQKFGNGVLIVSNNEGAHRNNGAVDSPSAAQFQLRKAPQLQKLSVVQKTPQLVNQQPMQAHVPRQIDFSAGSSSKPGVL

VTRAGVLDGESAEVDGLCKEEGPPPVMEDRWPRKRGRKPANGREEPLNHVEAEHQRREKLNQRFYALRAVVPNISKMDKA

SLLGDAITYIPDERVALPPLPSRRPCRSGGAHLHGLVVELLLSPSAPPEAKSRRFCSTLGMTNQAVNAAKEAVQRSEDLD

IRIIRLYEVKPKRVQFQKADSVILKNHADIQMQKVKTWKSSANLKKKSLRITEIRLLKEAYGNHWLLRNPIRVEVKTEAK

STSKVAGQQELGPSITPLGLRLELFPTKS

>Zm03G01110

MDDIGSVPFADAGLLDGFYGGSHGHGGDYGLLASQLGAGAGASSTSPAILDGSVPLVDAAASAEEATRRKGDHLQDDKAA

MALKSHSEAERRRRERINAHLATLRTMVPCSDKMDKAAVLAEVITHVKKLKSTAAHIRDRCAAVPADADDVVVELVHGGA

APPSAGGGVLVRATLSCDDGADVFADVRHALRPLRLSVVGSEVTTLGGRVRFTFLITSSTCGDVGAVVVDSVRQALQSVL

DRANSALEFAPRASLLNKRRRVSTFESSSSSSA

>Zm03G01970

MDMDSSSSWLHGYATTNGFMCGGYAASPAEVQCVEDEQQQFLISSQIQHHLNQISMRMNMDDEAAAAVDMHDLLEDLDDP

RRAGAGACSFPSSSSSSSLSLSLPASASLSCSPESSSPPHILGAATAPASGGCNQQYPEVSSHVPLVPPPPTVLASYSNL

HAPAAAPEETSPAARPTGGAFKHYARHLGPTRTTPTKPGTCGQRMFKTAMSVLSKMHVAARYSQQQYYYEAAAAEVAPPP

SVNQLQHMFSERKRREKLNDSFHALKAVLPPGAKKDKTSILIRAREYVRSLEARVAELEEKNKSLESRLAKDGSGCGDDH

DSGSTTKVQVEISRAAANEELCTLKIAVIRSPSPCNMTDVVVRTLQCLKEQIGDGVSLVAMSTSGGGGGGAGPTTTTTGG

KKGSPGDVLTLQIKSPGGTDWEEEPVKDAVAKVVADALTTTPRPPPAAAAAATTVSSCFGEASQLTTSS

>Zm03G04400

MMPGSMAARQPLQGKQELQPYDGRDPSALGAGGSPVLLPRQQESAAPPAVRVAPPPDMSSSSGSGRSATEARALKVHSEA

ERRRRERINAHLATLRRMVPDTRQMDKATLLARVVEQVKLLKRKASEAATTTTQSTPLPPETDEVSIELHTGDAGADRSV

YIRASISCADRPDLVAGLAQAFHGLRLKTVRANMTSLGGRARHVFVLCMEEGWGSAGAGAGASLRSLKEAVRQALARVAS

PETAYGSSPFQSKRQMILESH

>Zm03G15560

MEDGGLVSEAGAWAELGTGGDESEELVAQLLGAFFRSHGEEGRHQLLWSDDQASSDDVHGDGSLAVPLAYDGCCGYLSYS

GSNSDELPLGSSSRAAPAGGPPEELLGAAETEYLNNVAAADHPFFKWCGNGEGLDGPTSVVGTLGLGSGRKRARKKSGDE

DEDPSTAIASGSGPTSCCTTSDSDSNASPLESADAGARRPKGNENARAAGRGAAAATTTTAEPQSIYARKRRERINERLK

VLQSLVPNGTKVDMSTMLEEAVHYVKFLQLQIRLILCEASPQLTDSRNGGDTSNEAHKRNSTDGNNKQTCELDAFIYKSL

EGIQDQEFCNIIELLSVRPPIDAKLKLMKEIAEEHEIGPTYFNGSTLPLPKEKHDETVAASATKLPDEDYESNTGLDSLD

LPEVPKAAICPPSLNHIFNLSFSFSVVIFLAVDVLLDNIRSIDRAEEFAFRAEEDAVWSQVAKAQLREGLVSEAIESFIR

ADDAAHFLDVIRAAEEANVYNDLVKLTDSMPNVADLQNVGDRLYDEELYEAAKIIYAFISNWAKLAVTLVKLKQYQGAVD

AARKANSAKTWKEVDDLEEVSEYYQNRGCFSELIALMESGLGLERAHMGIFTELGVLYARYRSEKLMEHIKLFSTRLNIP

KLIRACDEQQHWKELTYLYIQYDEFDNAATTIMNHSSDAWDHMQFKDVCVKVSNVELYYKAVQFYLQEHPDLINDMLNVL

ALRLDHTRVVDIMRKEDYERLRESVDTHDNFDQIGLAQKLQKHELLEMRRIAAYIYKKADRWKQSIALSKKDNMYKDCME

TCSQSGDRELSEDFLVYFIEQGKKECFVSCLFICYDLIRPDVALELAWMNNMVDFAFPYLLQFIREYTSKVDDLVKDKIE

SQKEERAKEKEEKDLVAQ

>Zm03G22160

MDDSAEVKLVDEITGEGGAAGDWGYLGSDGMGSGSYPAFPFSRDVLSTPTSASLLLSMDPAALFDFNGTFPPSSAAAATA

GSSLSAFHDFSCINPFDDAGHFLGAPPPVPAAAAPQQQGQKGGFFAPLPASDFNDAGMSWDDEDEIDQSVDASSMAISAS

MENAAGAVAGASGAGGGSGRGKKKGMPAKNLMAERRRRKKLNDRLYMLRSVVPKISKMDRASILGDAIEYLKELLQRISD

LHNELESAPSSSLVGPTSASFNPSTPTLQTFPGQVKEELCPGSFPSPTGQQATVEVRMREGHAVNIHMFCARRPGILLST

MTALDSLGLDIEQAVISCFNGFAMDVFRAECADGPGMVPEEIKAVLMHTAGLHNAM

>Zm03G33720

MAWSETDAALFAAVLGRDAAHHLSTTPPHQDAPAASAPELQARLQDLVERGGAWTYGIFWQESCAGGRAVLGWGDGHCRD

GGAPHHDDADRSVARKRALLRLHALYGGGDDEGADYALRLDRVTAAEMYFLASMYFSFPEGAGGPGHALATARHAWATVD

PAPGWYVRASLAQSAGLRTVVFLPCKGGVLELGSAVPVRETPETLRALQTALAVARPPAREECMRIFGQDLSPGGSARAP

RSVDNWAPHPHLAAQATAASALASKEAAAGHKAPEPPRSIDFSKPGKPGHGQAGGEERRPRKRGRKPANGREEPLNHVEA

ERQRREKLNQRFYALRAVVPKISKMDKASLLSDAIAYIQELEDRLRGGGGGGGGCSAARPDSPDVEVKAMQDEVVLRVTT

PLYAHPVSRVFHAIRDAELIVAASDVAVADEAVTHTLVLRSPGPEQLTAETVLAAMSRGMTSATPSP

>Zm03G38000

MDEVWCGLDLQLQADGIHAGPDLHGHDHDPFWSALAECAASFLASDADDTGGLVASAGDVVHAKADGMDTSSFFADNDHH

GLTQRRDEQQQQQPVHSSSSLSSKRSLSIDSGGSSSSTLIPPLDGAAAAAFSPAPPQPQDPFAGDDEAIMLAMMAVISSA

SPSSSESSSPPHRAAGAGAVQPRVHLHGGDDSAGHVTVRTSSLAVAPTSAAARQQDDACMAAGSNNSSQVYHMISERKRR

EKLNDSFHTLRSLLPPCSKKDKTTVLTNAASYLKALEAQVSELEEKNAKLERHVPRDDGGGGTAATAAAVAHRRARVHVA

RAAPGEPQVSVTVMVMVECDIVDLVLRVLECLRWMGGVVSVLSVDADTYSPQAMLKALANIKLHIVDGDCWNEALFHEAM

TKAVHDATSPSSSPSCAAVAPLVAAA

>Zm04G05640

MLPPFGNPLWVPEDVDDQQQHHAPPPTPMGLGPGQGHDEQNLLALASAAAMGAGGIFSSPAVLDDDWYFDPVAAAAAAGA

QGQLLLAPPGPAPDAGSQMFSLFNVGGAATFDVHGFDLGLGGGGGGDLVPFAGAGNASNSASFSLVPAGNAGGFLGSFGG

FGTAPAQMPDFGGLGGFDMFNSGAGSSSAAPPPPASVSLTAPFSGRGKAAVLRPLEIFPPVGAQPTLFQKRALRRNAGEE

DDDKKRKVEAVAAAAGASSGGGGDTVLDDADDDDGGSIDASGLNYDSEDARGVEDSGKKDGKDSNANSTVTGGATGDGKG

KRKGLPAKNLMAERRRRKKLNDRLYMLRSVVPKISKMDRASILGDAIEYLKELLQKINDLQNDLESSPSTASLPPTPTSF

HPLTPTLPTLPSRVKEELCPSALPSPTSQQPRVEVRMREGRAVNIHMLCARRPGLLLSAMRAIEGLGLDVQQAVISCFNG

FSLDIFKAELCKEGPGLLPEEIKSVLLQSAGFHGGVMP

>Zm04G14040

MWEGGGFHGSHEALLLQAAGSGAADYGHGAGPALLPWLGPAAGAAPGFSYMAPHHAQPGPSGAEAAASPFGFGGGGYSDG

GVGGQFGVFGGPETALPAPHGLAAAAGGGTSAMQHGSRMVSGLLGTLQMELGRMAAKEIMDAKALAASRSHSEAERRRRQ

RINSHLARLRSLLPNTSKTDKASLLAEVIEHVKELKRQTSAVLDVEGEEAAAARQRLQLLPTEADDLAVDATEDGEGRLV

VRASLCCEDRAGLIPDIARALAALRLRAHRAEIATLGGRVRNEALRGVMDCKTASSDTSSSSNGGGSMKRQRMSGAHEQG

SL

>Zm04G31030

MQHQGAMNELVSQASFCSPASPPSFFSAAVGHHSMLDFVSCGVPEQWFLGEEALDKPIHDGAEWVAAGGSHDSAGSDLSS

NPPAAGIVLSERAARRRGRKPGPRSDNPGISHVEAERQRREKLNRRFCDLRAAVPTVSRMDKASLLADAAAYIAELRGRV

EQLEAEAKQQVASRKLGGNPAMCPASGGLEEKLEVRMVGRNAAALRLTTASTRHAPALLMGALRSLDLPVHNACVSRVGG

SATVQDAVVDVPAALQDEGCLRAALLHVLQQDESA

>Zm04G36350

MEDGGMCELVIGDHRNLRHRLSGAEDLFSIVGTWEERTNGASGGGGGGGGSSAAAVRAYSQGCTAGTAGAKTAAGTNNSR

RRTGDEEKGGSAPAQKKHKGSSAVSDDEGAAKMSHITVERNRRKQMNEHLAVLRSLMPCFYVKRGDQASIIGGVVDYIKE

LQQVLRSLETKKHRKAYAEQVLSPRPLPAVKSTPPLSPHVAVPMSPRTPTPGSPYKPASGAAATTTGSCRLPHRAAAAAA

AYIGTPTTSSSSSSYSHDQQRHYSTYLPTLDSLVTELAAQAAACSRPAASGGLTRLPDVKVEFAGPNLVLKTVSHRSPGQ

ALKIIAALESLPLEILHVSVSTVDDTMVHSFTIKIGIECELSAEELVQEIQQTLL

>Zm04G40140

MGGGVHHHHPCVAADGDGAGAGPGPASVEAALRPLVGVDAWDYCVYWRLSPDQRFLEMAGFCCSSQFEAQLPALGDLPPS

IQLDSSSAGMHAEAMVSNQPIWQSSRVPELQTGYSSGMVQEPGSSGGPRTRLLVPVAGGLVELFAARYMAEEEQMAELVM

AQCGVPSGGEGGAWPPGFAWDGGASDASRGMYGDAVPPSLSLFDAAGSVAADPFQAVQQAPGAGGGGVDDVAGWQYAAAA

GSELEAVQLQQEQQPRDADSGSEVSDMQGDPEDDGDGDAQGRGGGKGGGKRQQCKNLEAERKRRKKLNERLYKLRSLVPN

ISKMDRAAILGDAIDYIVGLQNQVKALQDELEDPADGAGAPDVLLDHPPPASLVGLENDESPPTSHQHPLAGTKRARAAA

EEEEEEKGNDMEPQVEVRQVEANEFFLQMLCERRPGRFVQIMDSIADLGLEVTNVNVTSHESLVLNVFRAARRDNEVAVQ

ADRLRDSLLEVMREPYGVWSSSAPPVGMSGSGIADVKHDSVDMKLDGIIDGQAAPSVAVGVSEDHYGGYNHLLQYLA

>Zm05G02590

MAGQPPPQGPEDDFFDQFFSMTAGGSYPGATAGGGRAPGDQPFSLALSLDAAAAEASGSGKHADGGKADREAIQLPGLFP

PAFGGGVQPPHLRATPPTQVFHAQQPKQGGAAVGPQPPAPRPKVRARRGQATDPHSIAERLRRERIAERMRALQELVPNT

NKTDRAAMLDEILDYVKFLRLQVKVLSMSRLGGAGAVAQLVADIPLSVKGEASDSGSKQQIWEKWSTDGTERQVAKLMEE

DIGAAMQFLQSKALCMMPISLAMAIYDTQHSQDGQPVKPEPNTPS

>Zm05G25480

MGDGQCELVVDNHCQLRHRLDLSGAEDLFSVIGTWEERTDDGGAGDGGGGGSSAPAMRAYRQISCSAGAVAATRLTGKSR

RRTGEEEEEKGSGGSAPGPAHKKHNKAGSAVTDDDEGAPKISHVAVERNRRKQMNEHLTVLRSLMPCFYVKRGDQASIIG

GVVDYIKELQQVLRSLEAKKHRKAYAEQVLSPRPSAGGVSTSVSAASPRHLAVKSVAPLSPRMAVPISPRTPTPGSPYKA

AASGVAGCCRLPLLPPYMLSSSAAAAAASYSHDQQQHYSTQTTTYLPTLDSLVTELATQQAACRPAAAAAGLALPDVKVE

FAGPNLVLKTVSHRAPGQALKIIAALESLSLQILHVSVSAVDDTMLHSFTIKIGIECELSAEELVQEIQQTLL

>Zm05G31220

MDGFVDPFVPEQAWPQDAMFIGSSWPCAGAGATSLADPAGTYAYLGAQAAPGQGAGFHLQDGSSTALVPPLELHQQFLSA

HLPGDDVTVTQEEGLGFEVNSALVAGMLGPVLSAPCAVSLADSAPVVCSSSNDSSGGSEQSGLPPPPRFLVLGEQPASWP

SAFPRISSLAGEETSRSFGFGAVSDNDLVRGSSCVADVNKYPQIGNARLHVEDDVEFNTGKMLSFAPGLDFGDLQLSQKE

LSGLRHLNSGQQLSSFDATRNPEQSSNEASGGGTGLNAPPFMVPANGAAGNGAPKPRVRARRGQATDPHSIAERLRREKI

SDRMKNLQELVPNSNRTDKASMLDEIIEYVKFLQLQVKVLSMSRLGATEAVVPLLTQSQTENSGGGLLLSPRSGSGRQQQ

ARGGSLPPPSSEVRDGAAFEQEVAQLMESDMTTAMQYLQSKGLCLMPVALASAISGQKGASSAAVQPENGGAKEMLRAVK

PLASPIHGR

>Zm06G13330

MRHASPPQELPAAGREIQAALAPANGNAAGARGGGGSFTALLGLPTPQAMELLLPRTTPPALALAPAAAPAPTPTFPSDP

HLVDRAGRFSTFAPPSPPSPSPTQQPPPPPPAAVAGKRKADPVDRASKGKAAKKGKTAEEKPAAAGGEDEKPAYVHVRAR

RGQATDSHSLAERARREKINARMELLKELVPGCSKVSGTALVLDEIINHVQSLQRQVEYLSMRLATVNPRGDFGGLDSFL

TTECGRIASFNCKNGIDLEQVTWPEMGVHGARQLMQLQQQFWHGDLAHPHQVASQWEKRGDGHPPVFSNSSPSLFGYDLT

SSGKPCTQSYITFLNEFSCLSLTSHITAWLAPQLLSLSLSRQCSLENKVY

>Zm06G18970

MASTASQPFQEGKQHLHLNHGGRALHSAYGGTAAWPASSVTPPLTRLETPASSSKSTAEARQALKIHSAAEKRRRERINA

HLATLRRMIPDASQMDKATLLARVVCQLKDLKKKSAETTQPPLATIPGETNEIAVVCCTGTASTAYERAAATYIRASVSC

DDRPGLHADLAGALRAMRLRPLRADMAALGGRAQCDFVLCREDGAGCRTLKALEEGVRQALAKAAFPETPPYGCNAARSR

RQRLAGSHCVLHGHGHGHGHGHGLHVIGEHGW

>Zm07G03170

MAGQPPPSGASEDDFLEHFFAFPSAASAGAAGGHAGAGVGGDHPFPLALSLDAAAEAKPDRDPVQLAGLFPPVFAGAGGV

HQPHLRGPPPPQMFQAQPKPGEGGMAPQPPAPRPKVRARRGQATDPHSIAERLRRERIAERMRALQELVPNTNKTDRAAM

LDEILDYVKFLRLQVKVLSMSRLGGAGAVAQLVADIPLSVKGEAGDGGGAPQQQQQQHVWEKWSTDGTEKQVAKLMEEDI

GAAMQFLQSKALCMMPVSLAMAIYDTQHPLDGHGHSLKPEPNASS

>Zm07G14220

MAGTGIRGGREGAGWRRAGGGDGTPAAALWWGVVRRLVAGRGEIAGSVFGGSFVGSLQQQQHFQSHHPQQQTAPPLPGQG

FGGGGRGGGRAGAPGGMSQPQAGAAVGGAAAPPRQRVRARRGQATDPHSIAERLRRERIAERMKALQELVSNANKTDKAS

MLDEIIDYVKFLQLQVLSMSRLGGAARSRQGRDGAAAAAGSDGLAVTEQQVAKLMEEDMGTAMQYLQGKGLCLMPVSLAA

AISSATCHMRPPVGGPGVAAGHHMAAMRLPHVTNGGAGADDVSASPSMSMLTTPSAMPNGAGAGVDGKDAASVSKP

>Zm07G16510

MMVPVGSDDETADVAMDRESSPPAATTRSGGTSRSHSEAERKRRQRINAHLATLRTLLPAASRMDKAALLGEVVRHVREL

RGEADAAAAGAAVAVPGEGDEVGVEEGQQRCFCHHGGGERERAAAASARRVRAWVCCADRPGLMSELGRAVRSVSARAVR

AEIATVGGRTRSVLELDVGGQHNGDDAGTSSRPALQAALRAVLLSREDMLGAECCYKRQRFSAHLARV

>Zm07G17090

MGIQGNKAATHEHDFLSLYAAAAAKDAPLLLHDSKAPPPSQGNFLLKTHDFLQPLDQKPGAPPEPSPLPASAAESRHQPQ

QQVVVANQHALPLPGGVGTFSISPAPVSVALPAAVVKSEPPFVLWGQPAATLHPGARGHQQQWALPFAGAGQVRLPPQQA

PPDRKGRGGAGGGVMESGSRSSGGAGFDDDDGLTTRREVSSSLQDLTVRVDRKGGSCSDGGTDQRPNTPRSKHSATEQRR

RSKINDRFQILRELLPHSDQKRDKATFLLEVIEYIRFLQEKVQKYEATFPEWNQENAKMLPWSNMYFRSFWKNAQSKGQI

SGDSPPDPSHFMRNGSSPGSNFTGKLDDNHNIVTSAAASGAQDQVETDHMASGCYRSAETTANFTNNAMSQSQPQWTGPS

PVDDSAVNSETLNNQQLVIDEGTIRVSSNYSQELLNSLTHALQSSGVDLSQANISVQINMGKRAAKRPAAGVSSNSKEPA

DPASSNELGHQLTLSGAGADCLSHATKRHKRSNS

>Zm08G06500

MVMKMEHEDNGAIGGTDGTWTEDDRALGAAVLGADAFAYLTKGGGAISEGLVATSLPGDLQNKLQELVESESPGTSWNYA

IFWQLSRTKSGDLVLGWGDGCCREPRDGELGAAASAGSEDSKQRMRKRALQRLHIAFGVADEEDYSPGIDQVTDTEMFFL

ASMYFAFPRHAGGPGQAFAAGIPIWVPNSERKVVPANYCYRGFLANAAGFRTIVLVPFESGVLELGSTQHIAESSGTVQT

VRSVFAGTSGNKSAVQRHEAERSPGLAKIFGKDLNLGRPSVGLAVGVSNSKVDERTWEQRSAVGGTSLLPSVQKGLQNFS

WSQARGLNSHQQKFGNGVLIVSNNEGAHRNNGAVDSPSAAQFQLQKAPQLQKLSVVQKTPQLVNQQPMQAQVPRQIDFSA

GSSSKPGVLVTRAGVLDGESAEVDGLCKEEGPPPVMEDRRPRKRGRKPANGREEPLNHVEAERQRREKLNQRFYALRAVV

PNISKMDKASLLGDAITYITDLQKKLKEMETERERLLESGMVDPRERAPRPEVDIQVVQDEVLVRVMSPMENHPVKKVFQ

AFEEAEVRVGESKVTSNNNGTAVHSFIIKCPGTEQQTREKVIAAMSRAMSS

>Zm08G22710

MDEVWRGLDLQLQAGDIDGLHPVSDPFWPALAECSASFLAAVDDTACFGIANMDLTAASADAAAGKSDGVDTSAFFAHND

HRGALPMRQDEQQPVYSSSSLSSTRSLSIDSGGSSSTFFPLDYDAVAALPAATFSPAPLQLQQDPFAGDEEAMMLAMLAV

ISSAASTSPSSSESSSPLHRAAAAVQPRLHPHGGDDSASHVTVRSTSLAVAPERTTSAADAAGQQDDACKAAAAAAGSNN

SSQVYHMLSERKRREKLNGSFHTLRSLLPPCPKKDKTTVLMNAASYVMALEAQVSELEDKNSKLQRYVPCGDGGGGGGAT

TATAAHRRARVHISRAASDEQQVSLMVMVMVECDIVDLVLHVLERLRWTSGVSVLSVNADTYSPQALLKALANIKLHIMD

TATAGTRRCSARP

>Zm09G02380

MDFSAGSYFTSWPVNSASESYSLADGSVESYGGEGIMPPSSYFMTARSDHNLKFSVHEQDSTMLPNDQLTYAGARQTDLL

PGETPSRDKLCENLLELQRLQNNSSLPSNLVPPGVLQHNSTPGAFHPQLNTPGLSELPHALSSSIDSNGSEVSAFLADLN

AVSSASALCSTFQNASSFMEPVNLEAFSFQGAQSDSVLNKTTHPNGNISVFDSAALASLHDSKEFISGRLPSFASVQETN

VAASGFKTQKQEQNAVCNVPIPTFTARNQIAVAAMPGSLIPQKIPSWINENKSEGPVSHPSDVQIQPNSVGNGVGVKPRV

RARRGQATDPHSIAERLRREKISDRMKNLQDLVPNSNKADKASMLDEIIDHVKFLQLQVKVLSMSRLGAPGAVLPLLAES

QTEGYRGQLLSAPTNAQGLLDTEESEDTFAFEEEVVKLMETSITSAMQYLQNKGLCLMPVALASAISTQKGVSAASIPPE

Q

>Zm09G02760

MQPSGRAMAGGEGGGGAQQQTDDFFDQMLSTLPSAWADLGAGAAEDLAAQSHFGDDSSALLASRLRQHQIGGGDVKSSAS

HQSQVMLQLSDLHRHGGLGGEESGLFTDRSAPAPEEMEGGFKSPNSAGGDHSLFNGFGVHGAATVQPPFGQSGSMSPESL

GGTAASGGGAPPAGGAASSAGGGAAPPRQLRQRQRAKRGQATDPHSIAERLRRERIAERMKSLQELVPNANKTDKASMLD

EIIDYVRFLQLQVKVLSMSRLGGAAGGMAPLVASMASSEGKSNGSGGGGNTNATTKSGNGGGLRVAEHQVAKMMEEDMGT

AMQYLQGKGLCLMPISLASAISSATSSASLLSRPPPAAAAGGGGGQMHGANGGAAAAATTISSPASANSSG

>Zm09G03640

MNEQGLGGLGGGRGAVQGHGREAMALLQHQQLQQQRRQLEEEDEVRRQMFGGVAAFPAALGHGQQVDYGEDAGGLGDSDA

GGSEPEPPPERTRGGSGGGGGKRSRAAEVHNLSEKRRRSKINEKMKALQSLIPNSNKTDKASMLDEAIEYLKQLQLQVQM

LSMRNGVYLNPPYLSGTIEPAQASQMFAAVGGGNITASSSGAVMPPVNQSSGLQVFDPLNPPRDQPLSFVLPNVDKTIQE

APFHLESSQFHPRPFRMPESSEMMLPGEVVAKHQLTSIQGRVSMPGIGMNPIRQESSTVKADQFDGCSHSKE

>Zm09G15540

MAPLSPTPASASAPVSSSPRSLVPDASAAPMRPASPPQELPAAGGEIQAALAPANASTAGARGGGGSFTALLGLPTSQAM

ELLLPRTTPPALATAPAPTFPSDPHLVDRAARLSTFAPPSPSPSSTSPARPLPAAANAGKRKADPVDRASKGKAAKKGKT

AEEKLAGGDGDDEKPAYVHVRARRGQATDSHSLAERARREKINARMELLKELVPGCSKVSGTALVLDEIINHVQSLQRQV

EYLSMRLAAVNPRVDFGGLDSFLTTECGRIAGFNCKNGIDLEQVTWPEMGVHGARQLMQLQQQFWHGDLTHPHQVASQWE

KRGDGHPPVFSNSSPSLFGYDLTSSGAQQTPASKLKTEL

>Zm09G17300

MNLWTDDNASMMEAFMASADLPAYPWGAPAGGGNPPPPQMPPAMAMAPGFNQDTLQQRLQAMIEGSRETWTYAIFWQSSL

DAATGASLLGWGDGYYKGCDDDKRRHRPPLTPAAQAEQEHRKRVLRELNSLISGGASAAPAPAPDEAVEEEVTDTEWFFL

VSMTQSFLNGSGLPGQALFAGHHTWIAAGLSSAPCDRARQAYNFGLRTMVCFPVGTGVLELGSTDVVFQTAETMAKIRSL

FGGGPGGGSWPPVQPQAAPQQQHAAEADQAAETDPSVLWLADAPVVDIKDSYSHPSAAEISVSKPPPPPPPPQIHFENGS

TSTLTENPSPSVHAPPAPPAPPQRQQQNQGPFRRELNFSDFASNPSLAAAPPFFKPESGEILSFGVDSNAQRNPSPAPPA

SLTTAPGSLFSQSQHTATAAANDAKNNNNNNKRSMEATSLASNTNHHPAAAANEGMLSFSSAPTARPSAGTGAPAKSESD

HSDLDASVREVESSRVVAPPPEAEKRPRKRGRKPANGREEPLNHVEAERQRREKLNQRFYALRAVVPNVSKMDKASLLGD

AISYINELRGKLTSLESDRETLQAQVEALKKERDARPHPHPAAGLGGHDAGGPRCHAVEIDAKILGLEAMIRVQCHKRNH

PSARLMTALRELDLDVYHASVSVVKDLMIQQVAVKMASRMYSQDQLSAALYSRLAEPGSVMGR

>Zm09G24320

MGGHGDHHHHHQEVGVLVDDDGDDEELEHQGRACGGATSGGVVEQGVGDGGSAGHQDAAGMAFEASISSVGSVSAATTMA

PPQILCWPPPPQQQQLQQHHNLGGGHHQSPFFPLLQPPPPPPPPPPQPPFFADLYARRALQLAYDHHHSGGGGGPSTSSD

PLGLYMGHPHHHQQHAGPGMMMMMPPPFASSSSPFGDFVGRMTAQEILDAKALAASKSHSEAERRRRERINAHLARLRSL

LPNTTKTDKASLLAEVIQHVKELKRQTSEITEEACQLPTESDELTVDASSDEDGRLVVRASLCCDDRADLLPDLVRALKA

LRLRALKAEITTLGGRVKNVLLITADDSSAAAGCHDDGGAAAPDDDDDDRQEEAAPVSVSSQQHTVASIHEALRAVMERT

AAASAAEDSGAAAAGLKRQRTTSLSAILENSRSI

>Zm10G09940

MEMGDSFEYYWEMQQYLESEELSLYMGTQDDALSCYDSSSPDGSISNSSWAPAGVAATASEKREGPGGAAAANKNILMER

DRRRKLNEKLYALRSVVPNITKMDKASIIKDAIEYIEQLQAEERRALQALEAGEGARCGGHGHGEEARVVLQQPAAAPAP

VEVLELRVSEVGDRVLVVNVTCSKGRDAMARVCRAVEELRLRVITASVTSVAGCLMHTIFVEVDSDQTNRIQIKHMIEAA

LAQLDDASASPPSVMSYY

>Zm10G11920

MPVSISICRTGPGEELAELLWDRGPALRRAPPPFQPFTCSAAGSSRSQELKRHASDTTKASAFVTAVSVPLGTHDAGSGL

GLAGLPVHDDDDAVPWLHCPVADDGDGDTAPLPPEFCAGLLSEYSEVAAPAPAFHAAATPPAEAAANKLAPPSAAGGGEG

VLNFTFFSRPLQRPQAAAAPAAAAASNPVESTVVQAAANRLRSTPLFSEQRMAWLQPPKAPRTTAAAAAPPPPPLAPLLP

DSRHGETVGTVAQPQPRSQPEARPPDAAAVTTSSVCSGNGGRSQLKRSRHLAADCSVSPDEDLDDEPGATRRSAARSAKR

CRTAEVHNLSERRRRDRINEKMRALQELIPNCNKVDKSSMLEEAIEYLKTLQLQVQMMSMGTGLCMPPAAMLLPAMQQQL

LHHHPMAHFPHLGMGLGFGMGAAAGFDMLPFPCVAAGAHFPCPPGAMFGVPGQAMPSLPAAFAHMYGAGSGAGPAGQTEA

ADAAAPARPGEAEQGDQQVQHAKQT

>Zm10G17090

MDMEDSGQFMQWAMATLQHDEDQAVDYPIDDGRGGATFPSLRALREASSQAAEMIQEPLAAANSRCGGDGGTAAAGNNIS

GTAPRRSSSGSAVIQPLRWNFGAGAAAPPGRDGVPVPAEAAATGSLLPDLAYGPPPTRKQAVLKSVGSIYAQDHIIAERK

RREKINQRFIELSTVIPGLKKMDKATILSDATRYVRDLQEKIKAHEDGGGSNDRGIVESWVLVKKPCVAAPDEDAGSSPS

WDSSGTTAPSPATNPLPEIEARFLNKNVTVRIHCVGVKGVVVRVLAELEELHLSIIHANVVPFHACTLIITITAKVDEGF

TVTAEEIVGRLKTSACINAPAKPLLQ

>Zm10G22550

MALSASRVQQAEELLQRPAERQLMRSQLAAAARSINWSYALFWSISDTQPGVLTWTDGFYNGEVKTRKISNSVELTSDHL

VMQRSDQLRELYEALLSGEGDRRAAPARPAGSLSPEDLGDTEWYYVVSMTYAFRPGQGLPGRSFASDEHVWLCNAHLAGS

KAFPRALLAKSILCIPVMGGVLELGTTDTVPEAPDLVSRATAAFWEPQCPTYSEEPSSSPSGRANETGEAAADDGTFAFE

ELDHNNGMDIEAMTAAGGHGQEEELRLREAEALSDDASLEHITKEIEEFYSLCDEMDLQALPLPLEDGWTVDASNFEVPC

SSPQPAPPPVDRATANVAADASRAPVYGSRATSFMAWTRSSQQSSCSDDAAPAAVVPAIEEPQRLLKKVVAGGGAWESCG

GATGAAQEMSGTGTKNHVMSERKRREKLNEMFLVLKSLLPSIHRVNKASILAETIAYLKELQRRVQELESSREPASRPSE

TTTRLITRPSRGNNESVRKEVCAGSKRKSPELGRDDVERPPVLIMDAGTSNVTVTVSDKDVLLEVQCRWEELLMTRVFDA

IKSLHLDVLSVQASAPDGFMGLKIRAQFAGSGAVVPWMISEALRKAIGKR
