## Supplemental Dataset 2 for "Evolution of vascular plants through redeployment of ancient developmental regulators"

>Aa2051499-Anthoceros_agrestis-A|m.652

MAISRSRTCESEQEQEQGEGQEPEQAAAQLNMSSSLHQSLRGLCLKSGWSYAVMWKLKRR

NRMVLTWEDGFYQSANLCPLNEVQQCTELRKCTGAAIIGNAQQQDPLRQAIAKMSYHVYS

LGEGIIGRVALTAKHQWVFGGGDKGGVVGTGGSRPFGRSSVEYPSGWQNQFAAGIKTIAV

IAVSQGVVQLGSTEIVMEDLDLVAHVKSLFLSSKDLPEILPTTYDTVTRSHGVPVPTAQP

HVAPKAESLAVRSASVATDPDFKSIAPVGLRGPPCPPNMMLCGMPSEELLHQQDGNRRLQ

QLMASVESQRSLEDRSFKSQRVAGPGDEVSQQHRNMQRVFADPCLTGTPRLLSENQFVGS

DAHHVSSGELLSVFGARPQHSIQPLGLSWMPSCENSNIHNQHIMVSGASPTVANGDTLMQ

ASVAHSPGTYSSVPCPPFKGHHLSKSMDESGREALHEVLQDPTHVESGSSWCAGPPLLND

ETTMDVSLTKQLDQSLAGKSCRKRCRPSEAPRPRPKDRQQIQDRVRELREIVPNSKKCSI

DALLERTIKHVHLLQSVARIEGRFKDEVQGEQNGNPNMAVDDGREVVVQTLSQPRQMLVE

MMCEKRGLLLQLADHIRGLGLTIIKGFMEVCCDKIRAQFVVEASRDMHRVEVLLTLMQLV

KSSEMVV

>Sle2000505-Selaginella_lepidophylla|m.905

MSAAGEGLSQALRSLCNRTGWCYAVFWKLKRRSRMVLTWEDSYYEFAPVPAAAAHPFNRK

QDGGGIAEDQIQLAVAKMSFQVYALGEGTIGRVAFSGKHQWVFATGERCMDNGVLAALEN

GWANQFAAGIKTIAVIAVQQQGVVQLGSTQIMSEDLNLIGQIRALFGSLQSMPGALVPDF

SSDGQTGRCLAVPRASKPASTLAAFGSCGRTAMNSSRAVSAVPVEGGVSERMLSLAPKAI

DPGFQHSWGIPCMEQKAQDVRTSANGLYNLRNGFARDAHDFEEDPGLVSGRSMSNKPPAP

VAPARSVDHSTLFLHAAGDAGRLAGFSNDLSYGQSDDVRHLDAGGWPPPPPDSALTVRDY

DPLPTPYGEETTLMRTGPEEGFDYNSLFKAFQKDDLDDFNNLLASLNKEYPVEKQGTSSS

TSCSQFAFGDELSEVVVCHPKCGPSKDELYVSMAADDMKTGKLVLPFDAPATATMMDKGR

DAFVSLSGDGPEQLLDAVVAGASSSSLGCKLPAVLPPKTSVPSVPPLTVGNAVQYLEKQL

SASSSLDRDIALNWPASSASVDVSSSSWTEGSQSKRLEEHSYSKDTSGGAATPVRKQEEA

TRMAGRKRLRPGEAPRPRPKDRQQIQDRVRELRDIVPNATKCSIDALLEKTIRHMKFLQS

VTQHGDKWKVGADVKGERDGADKSSGSSWALEVDGKGSGVPILVENLKQPRQMLVEMLCE

ERGLFWEIAENIRGLGLTILKGVMESRNDKIWARFNVEAAKEVDRLKVVWNLTQLLKPSG

KNGSHSNASFMGSEMSDVCSESFGTRFQPQVASHATAGGW

>Hm2013431-Huperzia_myrisinites|m.916

SAAEMSCGIYPLGKGLIWNVALTGKHQWLHVPEEANCSTMGTRDNGTSLDANPADFYDRF

DVGIKTIAVVAVPHGVVELGSTQKMIEDLNFVSHIQALFGFPCSALAAFPLDFLPKKEGK

PQQVPMSDLITICREKFTTLNGNVLRNCPFNYGRCLNGSNLPVVGLSKKQASQSLHPSRL

FTEDEAVSSPTNVPYNCSTEGDAIRISLSTLSSMSGASCGPSSESLHGCNSLSKMDENLR

NTLLSPSESMHPIWDCLCLGELETSHKYLSADGSRLNFPDDTIEISKSSKLQLEDFADKI

KVFLGSNFSNGASKMIHDRTRYFLKGSSWHMENSNPQFLIKYKLSKILAPIDMTEKERAA

STPGGRIEYIGSSEKFLGSSWSNEGPSLQSFERVSSFYAPHSNMNAEALLKNNVADVTEC

FQLHPKIVTASTCQDLSSKSECVVSSQSTSSVNSCSSNNISNTAANQPKFKVMQTACKDR

QSLKKCAIKKVVTENFGNACVRQSSEKSQFKLIDDDTCIDETESPMAHYQAPIKNKKRSR

PYEKNRQLIKDRLRKLRKIVPNTTRCSIYELLERTIEHMHFLRSVTQHGKKSQDVSESNS

SDPNLPNH

>Hs2016584-Huperzia_squarrosa|m.929

MEATTYPAGWGCQFTAGIKTIAVVAVPQGVVQLGSTYIVMEDMSLVGRIRSLFMTLQNAS

HTHIPYTPEIQSGILHNVCLPFSLPTIEPTTASVGTKGPKNLNFSKFGRDSWAEAPRVEF

NSFWTDLVGVAENGKSSTHPAQQSQFPYGAKASRTDSESSSHHLRTSQIIQSLTSQSSGM

STASNSDAGVLGSLPLGSMFSSSSSSSSSLGHSVSDFTSSLQFLPFDSIGKENLLCLSPL

EDFILPPIISSSSTVTCTEEASGNVVDVSDILAANSLHNTKALPNMLKEEQYNPPIQVLQ

TNTAQNVTSIATNSQLHVARSERLKADCFEVAGPTPFGVAQGWEQFSIDKSTLLPLDKKA

AESIFPVKGGKDSEISFSGVCESFGGSWIGLGALDDIDVFVDSLAKDGCRQIEPFSLNPS

QHDMCDELSEILAPVYKKGSLPDRLEESDENKFLKAFASSVPLQSCTEKGLNQFLLSSSI

QETILSETKGEPLLEAIVAGLTNVPNLLNPNSTTSSLYQTSSSRPNALLNSASVGSIDMG

TPSADQDERKISMTHIPSKGLINGDSVLNGRTYETDTETSIKLHFKQEITETSSTIQVNG

QNKKSQESVVTESLSGKRNEESTKGGRKRVRPGEASRPRPKDRQQIQDRVRELREIVPNG

AKCSIDALLERTIKHMHFLQTITQHGGNWKSGGEKGRQHDSSIVGLEQLDNGASWALELG

GQGMGCPIVVENMSQPGQLRVEMMFDEKGLFLEIADTIRGLGLTILKGVMEARNDKIQAR

FIVEAARDVHRVEVLWSLTQLLQPTTTASSTVTSQGCLV

>Sw2182007-Selaginella_wallacei|m.964

MAEVLKQALRSLCTKTGWCYAVFWKLKRRSRMVLTWEEGHYNALQSGLARSPLEHGLCRQ

TEDQIHLAVVKMSFHVHSLGEGIIGRVAFTAKHQWVYGSHGDKQPGEASGQTSSDKYPDG

WSNQFAAGIKTIAVVAVPQGVVQLGSMQLMPEDLNLVGQIRALFGTLQSVPGAVVSDLTY

EGQARRNAAISRAPTTLTTPQATPPAACFAVGSTMLTKGIGSAQNRAASTGISAESGHGN

YRLADQKATLGFQHDGAQRARAASLWPSGYLKHPSEVSQQFQPLATYDSRIGHTRGQGEG

DPLLDLLRPNSGKPAAADSIQFGPAWSPSSFGVGGFEGLHSSGADTATTGLLDPDEGFDY

NSLFKALQKDDLDDFNWLLATLPDQVPVSSGSNFVLGDELSEALVPQQKNGPGNGYNSSL

LTPPSEDVKAAEKQLQFEPRPSAGYCDILLGLTQKKPEPLLDAVVAGVSVTPDPMKSVRE

MPSAAGGSLLPVQAATVTSPSATCKDSHLPDNTPAAAAAAAAAASAVKWSPTLQVKPAAS

SWAAEACQMKKSEDHTSRDTSSSSRKQEDATRTTGRKRLRPGETPRPRPKDRQQIQDRVR

ELRDIVPNATKCSIDSLLEKTIKHMQFLQSVTQHGDKWKAGGDLKIGGMLERHSMDTGGA

SWAMEVGSKGPCPIVVENLSQPGQMLVEMRCEEKGLFWEIADNIRNLGLIILKGVMESRS

DKVWARFNVEMQASREVDRVQVLWNLLQLLKPNNNSKQHNVGTSAASSFTGSEITDICSD

SLTHHGGIMVSSANW

>Sw2004401-Selaginella_willdenowii|m.980

MAEVLKQALGSLCNRTGWCYAVFWKLKRRSRMVLTWEDGYYEFGTSSALLTGNGRGLLEH

GVCKPDGRVVEDQIQLAVAKMSFHVYSLGEGIIGRVAFTGKHQWIFANGERLECGPTSSE

KFPDGWSNQFAAGIKTIAVVAVQQGVVQLGSTQMMTEDLNLVGQIRALFGTLQSVPGAFV

SEFVPDSRCFPLTKTPSTATRTTSCLGAVGRTSLTNRSMKAMEADQPRNASVTFTQPGIC

STQQLPGWSSADFMTFSPVDQRTVPDMQNSVFQRDAGAGTYGTLVRCSNSFTRDTNLMQA

NNFKQMKFPFDGTPVQMPSTVADGLTRGEQPSLNTSVDACRSSASGFSGSYAYKVPTDNR

QFDTNLWSSSVHSSVFSVGDLELFGEDSGMMQPDEGFDYSSLFKAFQKDDLDDFNNLLAS

ISRDCPGANGPNGSQFAIGDELSEVLVSHQKTGATKEEPLVSTPVPPLNDVKVQPKNVTP

FELPSDIITLDCIPAFGDIFVGLSDNKPEPLLDAVVAGAGTLQQSSANVKESSENSASNE

CSRPMTATANAMQMPKKQPVATSSFDRDIWAEASQLKKSEERTQKDITVSRKEEPARVIG

RKRLRPGESPRPRPKDRQQIQDRVRELRDIVPNATKCSIDALLEKTIRHMKFLQSVTEHG

DKWKAGADIKGLDTSSLESGASWALEFDGKGTGVPILVENLKQPRQMRVEMLCDERGLFW

EIAENIRHLGLTILKGVMETRDDKVWARFNVEAAKEVNRVKVLWNLTQLLKLNPPKGSAP

ANSSYTGSELSDVCSEPLSTNRFQPQVPAHAPGITVSPGTGW

>Ss2010414-Selaginella_selaginoides|m.985

MGEILKQSLRGLCNRTGWCYAVFWKLKRRSRMVLTWEDGYYEFAPVQTGLNVSLLTGNGR

IQQSANLLEHGIYGQDEDQIELAVAKMSFHVYSLGEGIIGRVAFTGKHQWINGDRAAAFE

NSFAQRQQHTAQEKLPDGWENQFGAGIKTIAVVAVPQGVVQLGSLQMIAEDLNLVCHIRT

LFGTLQSVISEYAPDAQSGRAHHGMPGRISAVLAAPKSIGSGLINVSGALATDLGRTLAG

GGISLLSKKECGVQSATASATNNWSAFPSFTAMERKPSVDMQALSFGSEGMMTGLLTNYA

APQQQSEVELFKRPKMHSNASASLMDNLVHEKKARASHISTSASPTFEQSKLAGESGNFG

MNLGVDEFSCEGYDYNSLFKSYPKDDLDELNGLIDSISKEIGIHVTSAGVGGKCTDTVTG

VTGSTGTGSQFTIGDELSEAFTLNNNNNNFISKNGYTKDSLFDMLMSDEPSMKNVDVSSS

AASGLCDILGLTESKGVPLLDAVVAGATTPVAALSAPPSLKHTHGMPTEGDAGTTAAAAA

ASIGGDCEEVNNKWVSSVKVVGTGAAWIERSSASSSKKMEDATSRSNSNGNKKQEEQVAK

VVGRKRLRPGEASRPRPKDRQQIQDRLRELREIVPNATKCSIDALLEKTVKHMHFLQSVT

QHGDKLRDAKMMETSNVGDGKNLDNGASWAMEFGGQGMGCPIVVKNLKQPKQMLVEMMCE

EKGLFLEIADNIKGLGLTIVKGVMECRGDKIWARFIVEASVMEVKRMSVLLSLMQLMELN

LTSNHNNSVVTASDGVGDVYSAYHHHSHHIQHQQQHGMSITVSPSYA

>It2010667-Isoetes_tegetiformans|m.1012

MAIVLQQTLRSLCFKTGWCYAVFWKMKRRSRMVLTWEDGYCEYSNSTLSIPSFDSAPVPA

PANPSLACGSTFHMGGNPLEQGLCGQNGGMEEQIGLAVAKMSYHVYSLGEGIIGRVAFTG

KHQWVFSSGEKAISAAGTVQGRSGGRSNNEKYPAGWTNQFTAGIKTIAVVAVPQGVVQLG

SRETIMEDLNLVGHVKAMFSTLQSVPGAFLSDFVPEAPAGKVQNAYPLNFQMPMVMPMTH

SNTIARGPSLMHAYSNQIASLGEGGQVGKPGNLQQHSLSPSSYPLNLQNSSIPHLQPELA

AQTFQMQQVGPLQNRALPSRGTFEVNSPVTTSDAPIASFWSANCFPPNTFQSLKQKVLSV

NYGIDQERSIPVTAAGGQPGVSTYDENDVQKKSLAGQQFASNYKHCYHVGSALQACKVLS

RRDCSSSTEKDTGMGMSRGVGLSSVSKPTLLMNSRLSDTKPKSKLDPRNFYPLDHSLIDT

GTCTKQENSSFWGVGSRDIDTQVLNHTNPGTFSISNHKLEISVGESLGVDQGCGEDLLDN

DNRQLEAGKGSCLSGESGSNGIGMYPLLKNEHDTLESKFAIGDELSEILAPLYSRIHEKH

LLDDISAASTDVGSKENNSKNVCEKGLDFEHLDGLEYVCMQERSTSESKPEPLLDAIVAG

ITTRSEFATEQPTFLVNGTDQTSGNRDILFGLENVNNLEPILLGKEEIKEKNVLGTVSNV

LGDKLVKQSQYDAQCEQAQKEEFIKLGKKRKRTSELARPRPKDRQQIQDRVRELREIVPN

AAKCSIDALLERTIKHMHFLQSITQNGGKWIAVEGIKDQGESSLGSGDSGASWALQFGGE

GFGCPIVIENLCQPRQMLVEMLCEERGFFLEIAENIRGLGLNILKGMMEVRSDKVWARFL

VEQASRDVQRVEVLWSLMQLFYPASTIENQGLLRTCYNDNDKHLS

>It2013412-Isoetes_tegetiformans|m.1010

MTVVLQQTLKSLCYKTGWCYAVFWKLKRRSRMVLTWEDGYYEHSKPSAVSNFAAMQIGGV

EQNIGRPDGGGMEDQIGLAVAKMSYHVYSLGEGIIGHVAFTGKHQWVFENVDKLDVLENH

GSGRDNYPVGWSNQFTAGIKTIAVVAVPQGVVQLGSNQLIMEDLDLVGHIRGVFGTLQSM

PEVLASDFMSPQFGRTSYATGLPPSLPSGGKTLNSGLMPGQSRASQVALPQAQSQLNLQS

IIADRSFLAPLSVGSGDFPSNIFANKFSRMSTFGSCVKTPFEQDKVLQSLSQTNQLYFQN

KRALEMPYKDDEVKTVATSLQLDVKDRKVLLTSGVLQFPPHTSMHTLDVEDIAVQNGGPA

VKITSSSAINNMVLMPNLSDPHRAGQSPSQCQSNSLWHTSTDHVLDTLETTAVVSPGLPN

LTLDTWENENLTKDVLLLPELSEGIDCDKLVGSSAFESDTFDIDGFFASLTKDCTHTNSL

TTTGSQFAMGDELSEALAPVPKKEAEKEGYLDLLFSPNDMVRSIPPTQMPTASSGQGSGI

LQSSGNSSSLGDILRTGGLGELFHTDVKPEPLLDAVVAGVSVMPSSLSTRGNIYSRDAMS

FPLQRHTSAPLMNFELPQCLPVSAASSSVSTSLSTSKQNDDSGICQSRSNISQGTMKWQS

SMMSAPNSWVEQTKDKHTEERALKEQQPALKTHEETTKNGKKRARPGEPTRPRPKDRQQI

QDRVRELREIVPNATKCSIDALLERTIKHMNFLQSVTQHGGKWKSGELKGLENGSSLLSR

ESLENGASWAFEWGGGQGMGCPIVVENLSQPRQMLVEMLCEEKGLFLEIADNIRGLGLTI

LKGVMEARADKIWARFIVEAIRETHRVEILWSLMQLLQPNSKTSSTVTCQSGLMGSQEVD

VCPADSFPNKVRSLIENR

>Lde2012850-Lycopodium_deuterodensum|m.1016

NVHSSEGSMYPTGLANQFTVGMKTIAVVAVQQGVVQLGSKYMIMEDMNLVGRIRSLFETL

QNGLGNCFSKTSETQNRMLHNVHMPVSLPIIGPLTMPNVVDGTSNLHLTKSGQESLVNVP

RTELNSFWSNSIGMVGNGNNGTHSIHQPQFSIAAKAPSMQVENAFHHSQTPHLIQPFTSF

TSENVPAQNFKSGLAGSLSPNSKFSNPSLDHWAVNEYTSSFKHLPVDSVRKEILQCPSPI

GDYSFSPIMSSSAAIVTCTSEISEKCLDVAEMRAANPPNSKCLMQEDLFNPHARSLRMKN

AAIASSSHLMMQGSESMKKVDPFEVVASTPIKECEGWEKDIVNKNALSSLVETDAENCIP

IRVDLKNSELTFSGLCESSLCGSWLGLDGLDDVDAVVASLAKDGCRNAESASLTPSQPNT

CDELSEVLAPINKKRSSIYKKGALKDGSEGFDENEYLGSFESMHPFQSREEHEISQAHLS

LYLIEETIMSEPKAEPLLEAIVAGLTHVPISACPNTRSSLLQTPTSTPYTGSLTSTHKNL

NSSAMGLPLADPDESRISVTPIPLKSFTSGESILSGRTYETDTEASIKLRPVQELTETSS

SIHVEEGKSKKSQESVITESLSGKKTEESTKGGRKRVRPGEASRPRPKDRQQIQDRVREL

REIVPNGAKCSIDALLERTIKHMHFLQTVTQHSGKWKNGEEVFFEQGREHDSSILGIEQL

ENDASWAVEFGGQGMGCPIVVKNLSQPGQLLVEMMCDEKGLFLEIADTIRGLGLTILKGI

MEARNHKIQARFLVEAARDVRRVEILWALTQLLQASATTSSTVTNQGGLVESQSLDSSPI

ISVFHQDSLAPLCMHQSLH

>Is2076750-Isoetes_sp.|m.1026

MTVVLQQTLKSLCYKTGWCYAVFWKLKRRSRMVLTWEDGYYEHSKPSAVSNFAAMQIGGV

EQNIGRPDGGGMEDQIGLAVAKMSYHVYSLGEGIIGHVAFTGKHQWVFENVDKLDVLENH

GSGRDNYPVGWSNQFTAGIKTIAVVAVPQGVVQLGSNQLIMEDLDLVGHIRGVFGTLQSM

PEVLASDFMPPQFGRTSYATGVPPSSLPPGGKTLNSGLMPGQSRASQVALPQAQSQLNLQ

SIIADRSFLAPSSVGSGDFPSNIFANKFSRMSTFGSCVKTPFEQDKVLQSLSQTNQLYFQ

NKRALEMPYKDDEVKTVATSLQLDVKDRKVLLTNGVLQFPPHTSMHTLDVEDIAVQNGGP

AVKITSSSAINNMVLMPNLSDPHRAGQSPSQCQSNSLWHTSTDHVLDTLETTAVVSPGLP

NLTLDTWENETLTKDVLLLPELSEGIDCDKLVGSSAFESDTFDIDGFFASLTKDCTHTNS

LTTTGSQFAMGDELSEALAPVPKKEAEKEGYLDLLFSPNDMVRSVPPTQMPTASSGQGSG

ILQSSGNSSSLGDILRTGGLGELFHTDVKPEPLLDAVVAGVSVMPSSLSAKGNIYSRDAM

SFPLQRHISAPLMNFELPQCLPVSAASSSVSTSLSTSKQNDDSGICQSRSNISQGTMKWQ

SSMMSAPNSWVEQTKDKHTEERALKEQQPALKTHEETTKNGKKRARPGEPTRPRPKDRQQ

IQDRVRELREIVPNATKCSIDALLERTIKHMNFLQSVTQHGGKWKSGELKGLENGSSLLS

RESLENGASWAFEWGGGQGMGCPIVVENLSQPRQMLVEMLCEEKGLFLEIADNIRGLGLT

ILKGVMEARADKIWARFIVEAIRETHRVEILWSLMQLLQPNSKTSSTVTCQSGLMGSQEV

DVCPADSFPNKVRSLIENR

>Pca2008364-Pseudolycopodiella_caroliniana|m.1036

MALHQTLRSLCSKSGWCYAVLWGLKRESRMLTYEDGYYDSENASATSNWTSLLPIVASYT

SRENNSMYEQDGHLNGREMTGIDDQICLAVAKMPYQVHLLGEGIIGRVALLGKHIWMFGG

EESNNALERNMDSTWWASQFAAGIKTVAVVAVSQGVVQLGSTYKITEDGNLVWRIRALFA

TLQHGYVTCLSNSSASISRALSNVSVPSSSPMKGLMITPPVVDRSSKLGVSERDGLVDIQ

RTPANVVWSNSLCTIANGSNSGHFFDQPHFPAAAEALTPDNERFPHCLQTQKTIQRYDSF

SSDSVLPQNSKLGMLGSSSSSSNLSSSISPHCNRNEFTPSFQPLPVDSKGKYDLQFPSVL

HDFSHSPVAPSSTIVMSRAEETAENHIYTGEMLAAKPLNSKVVLSSKCSVSEEHSGGQSH

LIQPNSAAVVSIPPRQNPGSGKGDTCGIPLQIQKQDLELAFSGVCESLLGRSWLTLDALD

DVDAYVALLAKDISNEVEPLLHTPSQPVMCDELSEALMPMYKMGSETGKTGSPKDQSKEM

TDPSQSCAENGLGHDPLSSSYLFDEAVKGEPLLEAVIAGLTHIPKSSCPNSTRSSMCQSL

NHALENFDGRVGSPLVDQDESKTSVTPTQPKNFNSRGSIPSGSACKLDMEESTKGPSVQE

LTEASSSVHIEGRSKKREESVITESLSGKKSEDASKGGRKRLRPGEAARPRPKDRQQIQD

RVKELRELVPNGSKCSIDALLEKTIKHMHFLQTVTQQGCILKNCQEEVLLENGQDNDGGT

LAREQLENGASWALELGGQGMGCPIVVKNSTQPGQLLVEMICDEKGLFLEIADAIRSLGL

PILKGVLESRNDKIQARFVVEAAEDMHRVQILLALTQLLQPTTASRSAVTSQGGFESPSL

DTNSVISVFCQDSVSSFCLPRGL

>Pca2011661-Pseudolycopodiella_caroliniana|m.1037

MALQQTLRNLCRKSGWCYAIFWRLKRRSRMVLTYEDVYYDSERPLIATKWDNWPSIKGKD

TPSAKSSMYEHEPHLLGRDGAEDQIGLAVANMSYQVYPLGEGIIGRVAFNGKHLWMFGGE

DSNKIVEGNTYSTWWDSQLAVGVKTIAVVPVSQGVVQLGSTYVIPEDMNLVCRIRSLFAT

LQNGYGAYVSKASESKSRLLSSVHFPLSSPFNGLMATPVAVDRSSNSDIFEQWGLLDMQK

SGINSFWNNPLDSFGNGNNGMHSIHQPPFPEAAKVSIMHNDSLSSPSQKYEVIQPFSSFS

SDTVPSHDFKSGMLGCSSSSSKQFSSTSNHWNLHEFTSSFQLISANSLSKDDRLFPSVQG

FSPSPLMPSPPVIVSQTEESSKIHTAMGEMLAENAVNSNVSPSRTCSVLEEHFNACNCLP

QTNDSAVVSVSQFQKLDAGSLNIDPCGVVSTKVRTQEAVERDGLYMGASVSVVEKDVECS

IPLHVDLQDMKVVSNGVCKSLFTGSWFSLDTLDDFDTCLGSLVGKEVEPLSLPSQLIMYD

ELSEVLVPRCKKGSSKDGSEENDKSKLSRLFEYTDPFQLCAEHRLGQGPLSSSYLVDAAI

APELKAEPLLEAVVAGLAQVPKSSSPDSTRKAISEMQCPMLKQFDSSGGVSDESKTTVTP

MLLKSFNSRESVLSGRACELDADASIIVQSVQVPTEGSSSVHIEEGRSATGQESVITEPA

SGKKVEEIPKGGRKRRAPSEPSKPRPKDRQQIQDRVRELREIVPNSAKCSIDALLERTIK

YMNFLQTVTQHGCRWKDYEEEVLLEKVQDDGSDILGRKQVENDASWALELGGQGMGCPIS

VKNLSQPGQLLVQMICDEKGLFLEIADTMRSLGLTILKGVMESRNDKVQACFIVEAAMEV

HRVQILLALTQLLQPTTTTRSAVTTEGGYESQSLDQNPLTSAFLQDSEGPLYLHPRLP

>Dd2013519-Diphasiastrum_digitatum|m.1040

MACMSVRSVCMDMMQLELKTKLVWLSLRXXXXXXXXXXXXXXXXXXVALTGKHQWIFGGE

DDVHASEVNTYPAGWANQFTVGIKTIAVVAVPQGVVQLGSTYMIMEDINLVGRLRSLFAT

LQNGSGTCLSKASETQNRTPHNVHMPVSLPIVGSMTMPAVVDGPPNLHLTKSGQDSLVDV

PRTELNSLWSNSLGVVGNGNNGTHFVHQPQFSIAAKAPRLQIENSFHHSQTPQLVQPFTS

FSSQIAPAPNIKSGLSGSPSPNSKFSSSLLGHWNINDITSSLQLIPIDSVRKENMQCPSL

IEDFTFSPIMSSSAAIVTRTEETSENCIDVAEMLAENPPKSKISPISKCQIQEELFSPPL

KNAAIASSSQLQIPGSESIKVDPFEVVASTPIEQCQGWERDIVNKNTHSSLVEKDAEDSI

PIKVDLKDSELTFSGVCESSLNGSWLGLDVLDDVDAFVASLAKDGYSQVEPASLTHSQPN

MCDELSEVLAPIYKKGSSIYKKGSLKDGLEECDENKFLSSFEFMDPFQSCTEHGLSQALL

SSSYLIEETVMSETKAEPLLEAIVAGLTHVPKPVCPNSTRSSLYQTPTSTSHTGSQTNTH

KNFNSSEMGLPLADRDESRTSVARIPLKSFASGESILSGRTYETDTEASIKLRSVQELTE

TSSSVHVEEGKSKKSQESVITESLSGKKTEESAKGGRKRVRPGEASRPRPKDRQQIQDRV

RELREIVPNGAKCSIDALLERTIKHMHFLQTVTQHGGKWKNSGEEGREHDSSILGMEQLE

NGASWALELGGQGMGCPIVVKNLSQPGQLLVEMMCDEKGLFLEIADTIRGLGLTILKGVM

EARNDKIQAQFIVEAARDVHRVEILWALTQLLQPTATTSSTVTSQGLVESQSLDSSPIIS

VFRPDSLAPLCMRQGLR

>Hs2008802-YHZW-Huperzia_selago-2_samples_combined|m.1054

MNSDLYLIREKEANKSRRRKMAMTLHQALRSLCLKPGWCYAVFWKLKRRSRMVLTWEDGY

CDYVKPASASNLSSLPLTIGNHTSIENNGTQERHGGTGAEDQIGLAVAKMSYHVYSLGEG

IIGRVAFTGKHQWIFGGGENIHALEASTYPAGWASQFTAGIKTIAVVAVPQGVVQLGSTY

IVMEDMNLVGRIRSLFMTLQNGSHTRISNTPEIQSGMLHNVCMPVSLPTFGPTTTPVGTK

ESKNLKFAKFGQDSLAEAPGVELSSFWTDSFGFAENGNSSTHYAHQSQFPNGAKTSRTHC

ESSLPHFRTPQFIQSLTSSSSGMATASNSDSGVLGSLPLDSTFSSSSLGHSVSDFASSLQ

FLPGDSISKENLLCLSPFEDFTLPPIISSSTVTRTEEASGNVVDVSDILAANSLYNTKTL

PNIKYLLKEEQYNPHIQVLQTNTAQNVTTIATNSQLHVARSESLKADSFEVAAPTPFGEC

QGWERFSIDKSTLLSLDEKAVESIIPVKVGGKDSEISFSGVCESFGGSWLGLGALDDIDV

FVASLAKDGCTQIDPFSLNPSQHDMCDELSEILAPVYKKGSLPDGIEESDENKFLKTFAS

SVPLQSCTEKGLNQSLLSSSSLIQETILSETKGEPLLEAIVAGLTHVPKSLYPNSTASSL

YQTSSSRPHTLLHNASLRSFDMGLPAADQDERKISMTHIQSKGLINGDSILSGRTYETDT

EASIKLHFKQEITETSSSVQAKGQNKKSQESVVTESLSGKRTEESTKSGRKRVRPGEASR

PRPKDRQQIQDRVRELREIVPNGAKCSIDALLERTIKHMHFLXRYSAWWQLEEWWRKGSA

T

>Hs2008803-YHZW-Huperzia_selago-2_samples_combined|m.1051

MNSDLYLIREKEANKSRRRKMAMTLHQALRSLCLKPGWCYAVFWKLKRRSRMVLTWEDGY

CDYVKPASASNLSSLPLTIGNHTSIENNGTQERHGGTGAEDQIGLAVAKMSYHVYSLGEG

IIGRVAFTGKHQWIFGGGENIHALEASTYPAGWASQFTAGIKTIAVVAVPQGVVQLGSTY

IVMEDMNLVGRIRSLFMTLQNGSHTRISNTPEIQSGMLHNVCMPVSLPTFGPTTTPVGTK

ESKNLKFAKFGQDSLAEAPGVELSSFWTDSFGFAENGNSSTHYAHQSQFPNGAKTSRTHC

ESSLPHFRTPQFIQSLTSSSSGMATASNSDSGVLGSLPLDSTFSSSSLGHSVSDFASSLQ

FLPGDSISKENLLCLSPFEDFTLPPIISSSTVTRTEEASGNVVDVSDILAANSLYNTKTL

PNIKYLLKEEQYNPHIQVLQTNTAQNVTTIATNSQLHVARSESLKADSFEVAAPTPFGEC

QGWERFSIDKSTLLSLDEKAVESIIPVKVGGKDSEISFSGVCESFGGSWLGLGALDDIDV

FVASLAKDGCTQIDPFSLNPSQHDMCDELSEILAPVYKKGSLPDGIEESDENKFLKTFAS

SVPLQSCTEKGLNQSLLSSSSLIQETILSETKGEPLLEAIVAGLTHVPKSLYPNSTASSL

YQTSSSRPHTLLHNASLRSFDMGLPAADQDERKISMTHIQSKGLINGDSILSGRTYETDT

EASIKLHFKQEITETSSSVQAKGQNKKSQESVVTESLSGKRTEESTKSGRKRVRPGEASR

PRPKDRQQIQDRVRELREIVPNGAKCSIDALLERTIKHMHFLQTVTQHGGNWKNGGEKGR

QHDSSIIGLEQLDNGASWALELGGQGMGCPIVVENMSQPGQLCVEMMFDEKGLFLEIADT

IRGLGLTILKGVMEARNDKIQARFIVEXQGTYTEWKFCGP

>Sk2008662-Selaginella_kraussiana|m.1071

MGEVLKHTLRNLCHKSGWCYAVFWKLKRRSRMVLTWEDGHYDAPNGLENGTVYRQQMEHQ

INLEVVKMSIHVYSLGEGIIGRVAFTGKHQWIYGNQRISENGSSPEKYPDGWSSQFAAGI

NTIAVVAVPQGVVQLGSIQVMPEDLNLVGQIRSLFGTLQSIPGGLVTDLKNPSPLPTVKA

YTTPVTTVTVPNPTFAVPFSGYSKPHDHSTSVPFNTHHQTAYPALDQKAVFQQAATAVRV

GYDSIRVLSDLAVKRPRLANQGQLTMDYHGSQNHHQNSQNHGFMGTSEARNTIPGTNSKL

FDIPGPDTTFRANHGNHFGTSNSIWSPPSQTSSFTGFDSLPQLGEETATTTTTCLLDSDE

NYDYNSLFKAKDGLEDFNNVLAALSKDCGVLLPSNEGFRTSNEGVGFRASNEGVATTGST

NFGFGDELSEVLALLQKNDNHHNVTSFSDNLHLQKNGYNKDHNNGVGATSYSDVMFTATS

SAGGDVKPVDTNVLTARTFELPSQEIDKISGYCDILACLNDNKLEPLLDAVVAGASMASP

CSKPAKESPKPAPFTGTTLVVPPVATAASSGKELEKTAAPQQASSSSLERVPKWSSSALQ

MKAAVSSWASEASNKKEHPVKDSSSASKKPEDASARVVGRKRLRPGETPRPRPKDRQQIQ

DRVRELRDIVPNATKCSIDSLLEKTIKHMQFLQSVTQHGDKWKAGADTKVGGIYEQQQQQ

LSESGATWAMEVGPKGPSPVSVVENLRQPGQMLVEMLCEEKGLFWEIADKIRGLGLTILK

GVMESRNDKVWARFNVEQQPQQQMQASCGVNRCKVLWNLVQLLEPNKHNGGGTSTAAGAA

AAGSAAASFTTAAGSEVTDVCSESFSGFQTQTLAAAHRGITVSSNGW

>Sa2007567-Selaginella_acanthonota|m.1084

MAEVLKQALRSLCTKTGWCYAVFWKLKRRSRMVLTWEEGHYDALQSGLARSPLEPGLCRQ

TEDQIHLAVVKMSFHVHSLGEGIIGRVAFTAKHQWVYGSNGDKQPGEASGQTSSDKYPDG

WSNQFAAGIKTIAVVAVPQGVVQLGSMQLMPEDLNLVGQIRALFGTLQSVPGAVVSDLTY

EGQARRNAAISRAAPTTLTTAATPPAACFAVGTRGIASAQNRNCGGAAPTGFVAGSAHSN

HLPADQKVTFDFQHDGAPKMRAASLWPSGYLKHPNEVSQFQPPPTYDSRIGHSRGQVEGD

PLLDLLRPNSTKPAADGIQFGPVWSPSSFGVGGFEGLQSSGADTAPTGLLDPDDGFDYNS

LFKALQKDDLDDFNWLLATLPDEVPISSGSNFVMWDELSEALVPQLKNGPGNGYNSSMLT

PPPEDVKAEKQLQFEPRPSAGYCDILLGLTQKKPEPLLDAVVAGASVTSDPMKSVREIPS

ASGAGSLLPVQAATVTSSSATCKDSHLPDKTPAAAAAAASAVKWSSSLQVKPAASSWAAE

ACQMRKPDEHTSRDTSSSSKKQDDTTRTTGRKRLRPGETPRPRPKDRQQIQDRVRELRDI

VPNATKCSIDSLLEKTIKHMQFLQSVTQHGDKWKAGGELKIGGMLERHSIDTGGASWAME

VGSKGPCPIVVENLRQPGQMLVEMLCEERGLFWEIADNIRSLGLTILKGVMESRNDKVWA

RFNVEMQASREVDRVQVLWNLLQLLKPNNSKQQNGATPAASSFTGSEITDVCPDSLTHHG

GIMVSSSNW

>Pd2128658-Phylloglossum_drummondii|m.1102

MTKMALALHQKLRDLCCKRGWCYATIWKLPPSNPITLTWGDGDQASTKSLSDSKIASIEV

SEIGTTPSQENGNPLQGHIIGDDATRLEVHIKLPQSKMSSSSLESLIADVALTGKHRWIH

IQERACTSADETAENAANIEQIPNIWSQSHAGIKTIALIAVLEGVIELGSSQKMIEDIRF

VSHILALFNPPQSAPTTSPLHLSRKTLALRSPTGRAVPVSRPMPFCRGAFLVLNGPLKHG

ISSKGSCNNGQFGDGFNSAMSKLSNQRAISNLPLSGCNSTHGPGTKVLHARVEGSPEGPR

GHMEKSSRLDLGGKESAWILTSTWSEGVEYAKPAEELVGSSCSTEAEPLHRFKTMLEPSC

FHLHRKGSSLSGDDVTNLSNVEEYSQFRQNMPGGLDTRGFSSNSDGIVTSRSVLSVNSRS

RDDVSTIIVDQPDFTALQSANNHHESLEEYRIRSVGDQSFRGADVKQLIEGSPIQVKHDT

FCPSTLPPTMSSRKRSGCGKKPTDRQLLRDRLRKLRKIVPDSTRCSIHELLERTIKHMQY

LQRVTQYE

>Ss2003544-Selaginella_stauntoniana|m.1108

MDEPICCRDSTMCLQTIAVISVPQGVVQLGSTQRIMEDLEMVSHVRTMFCTLQRVPGAFF

PDYVNSQAATQGGAFRYMTPIQALTAAPAPNLLTHQPPFPFSCSSRSQVSGGNLLMNAGV

AASNLRDMENLLAGPKNVCDGSSPAMFPSTGTTILDQARHRSRGTNEACSSFVGVGFPCE

ARHDGTSPTTTLSYSNTEESTDEAHGRPPGVGSSGSATELPDLDSLLLPADAFHDRGFDS

LMSRGVEDIDQFFALLPKDSNDYREEDLPKVVPEAGIETQAMADDMNLSVARDHAFGASS

DFVSSMLFPSSPSALQKDEQQPTGVKRARPQEEGKPRSRGRPQILDRVKELRGIVPNAEK

CSIDALLQKTIDHVRFLQAVLQQIGSSSRQEVDNGASLAMKAARFLSKT

>Ss2009233-Selaginella_stauntoniana|m.1105

MSAEGLGQALRSLCNRTGWCYAVFWKLKRRSRMVLTWEDSYYEFATLPPIANALSRSGAG

NGRSPLDMLLLKQDGGIAEDQIQLAVAKMSFQVYSLGEGIIGRVAFSGKHQWVFAAGERS

MESGVPAALEHGWSNQFEAGIKTIAVIAVQQQGVVQLGSTQTMAEDLNLVGQIRALFGAL

QSMPGAFVPEFTPDGQAGRGLSVVPKAMPTYVNKPASTLAAFGSSARTTMSSRPACPVPT

EGEHMLRSAAKVVEPAFQLSTSSWASMVVPSMGQKPPASTGLYCSRSQVFQEDPGHLSRL

ISKAPVESFFNSSETGRLSNFSSELSYGQSIRHFDSSGWPPPDSTLPVSTRDYDQNLPLP

TSFGDETGMMRPGPEEGFDYNSLFKAFQKDDLDDFNNLLASLNKEYAGGQASGSTTCSQF

AFGDELSEVVVSHSKCGPSKEELYVNMPTQPVEDMKPAKLALPFDGATAAMDKTRDAFVN

LSADRPEQLLDAVVAGASTSSLGSRGPTSAKSSLGCAPALTVGNAVQYLEKQLSSSSSLD

RDIGANWPASSVSVDVSSSSWTEGSQAKKSEEHSYKDTSGAAARKQEEPSRMAGRKRLRP

GEAPRPRPKDRQQIQDRVRELRDIVPNATKCSIDALLEKTIRHMKFLQSVTQHGDKWKAG

GDVKGERSAENSSGLENGASWAMELDAKGSGVPILVENLKQPRQMLVEMLCEERGLFWEI

ADNIRGLGLTILKGVMESRNDKIWARFNVEVHAAKEVNRLKVVWNLTQLLRPSGKNTSAP

SFVGSEMSDVCSESFGSRFQPQVVASHGITVSPGSAW

>Pc2067657-Phaeomegaceros_coriaceus|m.627

MVKEILRALCSQGRWSYAVFWRLQRRNRTVLTWEEGFGPLVAQAAGRELQGVGKHEIVAV

NAGDELLRVAVAKMCFQVYYLGEGIIGRVAITGRHQWVFGTGHGVSPGMGSGHIPVRGNL

EKYPGGWDSQFAAGIKTIAVIAVAQGVVQLGSVETILEDLDVVACVRSLFVKLQDTSGGI

FPGHEAQKPVAAPILGSPFGPGWMLSRSSKGPGGLREVPKVSVTHDVFLAEAVPGLGHTE

VEDVSVVSSVSSTELDVYSGLGGSKCSGAELSFLKDAFLPVVGRPSDVCGKALWKNHEPA

GTPTIVRDVSVECKKPRSIQSDHELLGHEVEKGAQALDNDEVTARTCRKRSRLGDNVRAR

PKDRQQIQDRVRELREIVPESNKCSIDSLLERTVRHINFLGSLSQLKGCNAAKKLKDRAS

EGVLHNGNDFSPLIVEDLNYPRHMLVEVVLDKKSLLAEVAENIRGLGLTIVKGMMEVKSE

KVRACFVVETTQKRCGETVLDDAGIHATGAW

>Ld2001113-Leiosporoceros_dussii-B|m.634

MSIMLQKNLREICQKTGWSYAVFWKLKRRNRMLLSWEDGFYQPPLANSLPSADNEPRQVG

NYMQGNHLWPGNTQEDPLCVAVAKMSYHVYSLGVGVIGRVAFTGKHQWVFGGGDKINHLG

GASMEYPSGWQNQFAAGIKTIVVIAVGQGVVQLGSTVTVMEDLDLVGYVRTLFLTGNCHK

PFPPNYLSGARIGHAQNTSTVAMSLPGFSVSEAGRDVVSSWRPALQANPAVVLPSDIMRK

ETFPAVSNDCSQMQIFHQTLLLQVLDDQEMHRLLPSAEPQQRVPAQSSLVTQTFKHENAA

FLAAVNSTSKSLVVDRSMAGNEIANIEGNLTGDLGLGSNLSRAVSGVPALSPMGLSWMCG

CGSSDVDSLHISTSGCTLPPVNGFINSLPNMSGDGSSADCQVVCLTSGYDDTPSPFGRSA

GSGSVDGHTGEGPDEYSTFVPGMPKRERPPLFLTPDHSFRDELSEAFCPSTKKSLEMSLP

GSHTPTSRMEMQGGSVSVSWSPAVLSGDVGQLAHLMAPASSSGEALAQPLLDSGPGTSCS

TSSQTSNVSGFDSTSGSGLWKETKREVSQSCLQVKPMKNWGSGQGLRNERTPEFVSVTKP

DQSAGKTGRKRCRPSEAPRPRPKDRQQIQDRVRELRDIVPNTQKCSIDALLERTIKHVEL

LQCLAQIEGRFKNQFQDQLKSGSNWAVEDGREGACPLVVENLSKPRQMLVEMMCEKRGLL

IQVADYIRGLGLTIIKGVMEARYDKIRAQFVVEASRDMHRVEVLLTLMQLLKPSEQSGGL

SETATSSACLFEPAMCF

>Ld2001114-Leiosporoceros_dussii-B|m.629

MSIMLQKNLREICQKTGWSYAVFWKLKRRNRMLLSWEDGFYQPPLANSLPSADNEPRQVG

NYMQGNHLWPGNTQEDPLCVAVAKMSYHVYSLGVGVIGRVAFTGKHQWVFGGGDKINHLG

GASMEYPSGWQNQFAAGIKTIVVIAVGQGVVQLGSTVTVMEDLDLVGYVRTLFLTGNCHK

PFPPNYLSGARIGHAQNTSTVAMSLPGFSVSEAGRDVVSSWRPALQANPAVVLPSDIMRK

ETFPAVSNDCSQMQIFHQTLLLQVLDDQEMHRLLPSAEPQQRVPAQSSLVTQTFKHENAA

FLAAVNSTSKSLVVDRSMAGNEIANIEGNLTGDLGLGSNLSRAVSGVPALSPMGLSWMCG

CGSSDVDSLHISTSGCTLPPVNGFINSLPNMSGDGSSADCQVVCLTSGYDDTPSPFGRSA

GSGSVDGHTGEGPDEYSTFVPGMPKRERPPLFLTPDHSFRDELSEAFCPSTKKSLEMSLP

GSHTPTSRMEMQGGSVSVSWSPAVLSGDVGQLAHLMAPASSSGEALAQPLLDSGPGTSCS

TSSQTSNVSGFDSTSGSGLWKETKREVSQSCLQVKPMKNWGSGQGLRNERTPEFVSVTKP

DQSAGKTGRKRCRPSEAPRPRPKDRQQIQDRVRELRDIVPNTQKCSIDALLERTIKHVEL

LQCLAQIEGRFKNQFQDQLKSGSNWAVEDGREGACPLVVENLSKPRQMLVEMMCEKRGLL

IQVADYIRGLGLTIIKGVMEARYDKIRAQFVVEASRDMHRVEVLLTLMQLLKPSEQSGGL

SETATSSACLFEPAMCF

>Aa2051930-Anthoceros_agrestis-B|m.638

MAISRSRTCESEQEQEQGEGQEPEQAAAQLNMSSSLHQSLRGLCLKSGWSYAVMWKLKRR

NRMVLTWEDGFYQSANLCPLNEVQQCTELRKCTGAAIIGNAQQQDPLRQAIAKMSYHVYS

LGEGIIGRVALTAKHQWVFGGGDKGGVVGTGGSRPFGRSSVEYPSGWQNQFAAGIKTIAV

IAVSQGVVQLGSTEIVMEDLDLVAHVKSLFLSSKDLPEILPTTYDTVTRSHGVPVPTAQP

HVAPKAESLAVRSASVATDPDFKSIAPVGLRGPPCPPNMMLCGMPSEELLHQQDGNRRLQ

QLMASVESQRSLEDRSFKSQRVAGPGDEVSQQHRNMQRVFADPCLTGTPRLLSENQFVGS

DAHHVSSGELLSVFGARPQHSIQPLGLSWMPSCENSNIHNQHIMVSGASPTVANGDTLMQ

ASVAHSPGTYSSVPCPPFKGHHLSKSMDESGREALHEVLQDPTHVESGSSWCAGPPLLND

ETTMDVSLTKQLDQSLAGKSCRKRCRPSEAPRPRPKDRQQIQDRVRELREIVPNSKKCSI

DALLERTIKHVHLLQSVARIEGRFKDEVQGEQNGNPNMAVDDGREVVVQTLSQPRQMLVE

MMCEKRGLLLQLADHIRGLGLTIIKGFMEVCCDKIRAQFVVEASRDMHRVEVLLTLMQLV

KSSEMVV

>Ph2003035-Paraphymatoceros_hallii|m.642

MIMGTPAEKHSTILRHDMGAMLKETLQALCSQYGWSYVVLWRLQRRNRMVLTWEDGFYQP

SVARGAGRNLHGLKRDIEGQGAWVGDVNAGEDSLRMAVAKMSYHVFYLGEGVIGKAAITG

KHQWVFAGGENGVLNGVGSGQTLVRPRMEKYPAGWDSQFAAGIKTIAVVAVAQGVVQLGS

VETITEDLGMVAHIRSLFLELQDSSGMVVSNHDLEKAFAVPTLGPVLGPDWMLSRSSKSS

GLQEVSAVPAIQDLVSPKAAPQGVEETITDACLFSFDHCVENPVLKTQEPQLEKSPQSVD

NDEVASKTCRKRSRLCENVRPRPKDRQQIQDRVRELREIIPESNKCSIDSLLERTVRHIH

FLDNVTQLRDCNRGKKSKDGAATEVMSHELLHSAWNLDGHSCNPLIVEDLSYPRHMLVEI

LFEKKSLLAGIAENIRGLGLTILKGVMEVKRNKVRACFVVEAAKDTHRLEVFWSLAHFIE

QMEKSTSSSGGECPLGPEMLEEDRV

>Pc2021222-Phaeoceros_carolinianus-sporophyte|m.648

ATYFGLRLTCKRAGLRISVCLVFGWSPERPARGWKCFLPALRSPARCILIRDHALLSLPH

FGLPRIMGMPVEKQSTILRDDMGTMLKEILQALCAQTGWCYAVLWRLQRRNRMVLTWEDG

FYQPVVARGAERNLHGVKRDIEGQGSWIGDVNTGEDSLRMAVLKMSYHVFYLGEGVIGKA

AITGKHQWVFAGGDNGVLNGIGSGQTLVRPRMEKYPAGWDSQFAAGIKTIAVVAVAQGVV

QLGSMETIMEDLGMVAHIRSLFLKLQDSSGLVVSNHDPEKAFAVPSLGPALAPDWMFSRS

SKSSGLQEESAVPAIQDTVSPKAAPEGVDETFTDACLSSLDHCVETPVSKNQEPRLEKSP

QSVDNDEVASKTCRKRSRLCENVRPRPKDRQQIQDRVRELREIIPESSKCSIDSLLERTV

RHIHFLDNVTQLRDCNRRKKSKDGTGTEVKSHELPQSAWNLDGHSCNPLIVEDLSYPRHM

LVEILFEKKSLVADIAENIRGLGLTILKGVMEVKKDKVRACFVVEAAKDTHRLEVFWSLA

HFVEQMGKSTSSSGGECPLGPEMLEENGP

>Pc2141251-Phaeoceros_carolinianus-sporophyte|m.645

MAISRSRTCESEQEQEQGEGQEPEQAAAQLNMSSSLHQSLRGLCLKSGWSYAVMWKLKRR

NRMVLTWEDGFYQSANLCPLNEVQQCTELRKCTGAAIIGNAQQQDPLRQAIAKMSYHVYS

LGEGIIGRVALTAKHQWVFGGVDKGGVVGTGGSRPFGRSSMEYPSGWQNQFAAGIKTIAV

IAVSQGVVQLGSTEIVMEDLDLVAHVKSLFLSSKDLPEILPTTYDTVTISHGVPVPTAQP

HVAPKAQTLAVRSASVATDPDFKSIAPVGLRGPPCPPNMMLCGMPSEELLHQQDGNRRLQ

QLMASVESQRSLEDRSFKSQRVAGPGDEVSQQHRNMQRVFADPCLTGTPRLLSENQFVGS

DAHHVSSGELLSVYGARPQHSIQPLGLSWMPSCENSNIHNQHIMVSGASPTVANGDTLMQ

ASVAHSPGTYSSVPCPPFEGHPLSKSMDESGREALHEVLQDPTHVESGSSWCAGPPLLND

ETTMDVSLTKQLDQSSAGKSCRKRCRPSEAPRPRPKDRQQIQDRVRELREIVPNSKKCSI

DALLERTIKHVHLLQSVARIDGRFKDEVQGEQNGSPNMAVDDGREVVVQTLSQPRQMLVE

MMCEKRGLLLQLADHIRGLGLTIIKGFMEVCCDKIRAQFVVEASRDMHRVEVLLTLMQLV

KSSEMVV

>Mto2053142-Megaceros_tosanus|m.655

PTGPLSTVSSPVSGLPSLSQPQFVRTYRACVHHSGGDTMVKEILRALCSQGRWCYAVFWR

LQRRNRDVLTWEEGFGPLQRDLQGAGKQQIVAANAADDLLRVAVEKMCYKVYHLGEGIIG

RVARTEKHHWVFGTGHGLSPGMGSSHIPMRSEKYPGVWDSQFAAGITTIAVISVAQGVIQ

LGSFETISEDLDVVAYVRSLFVKLQYTPGGAFPGHEAQKPVGVPMLGTACGPGWMLSRSI

RSPGGLREVSKVSVTDDMFAAEAIPGGLGQTEVEDVSVVSSVSSTEVDVFSGLGGSKCSG

ADLSFLKDAFLPVVGRPTDMCGKALWKNHESGATPSPHPNKGVRDASLDCKKPRSIQSDH

ELQDVDEGAQALDNDEVTARTCRKRPRPCENVRPRPKDRQQIQDRVRELREIVPESNKCS

IDSLLERTVRHINFLDNLSQLKGCNVAKKLKDRAGEAVLHTGNDVGPLIVEDLNYPGHMF

VEVVLEKKSLLADVAENIRGLGLTIVQGIMEVKSEKVRACFVVETTQKRCGEMVLNDTMI

HATGAW

>Pc2006372-Phaeoceros_carolinianus-gametophyte|m.658

MGMPVEKQSTILRHDMGTMLKEILQALCAQTGWCYAVLWRLQRRNRMVLTWEDGFYQPVV

ARGAGRNLHGVKRDIEGQGSWIGDVNTGEDSLRMAVLKMSYHVFYLGEGVIGKAAITGKH

QWVFAGGDNGVLNGIGSGQTLVRPRMEKYPAGWDSQFAAGIKTIAVVAVAQGVVQLGSME

TIMEDLGMVAHIRSLFLKLQDSSGPVVSNHDPEKAFAVPSLGPALASDWMFSKSSKSSGL

QEESAVPAIQDTVSPKAAPEGVDETFTDACLSSLDHCVETPVSKNQEPRLEKSPQSVDTD

EVASKTCRKRSRLCENVRPRPKDRQQIQDRVRELREIIPESSKCSIDSLLERTVRHIHFL

DNVTQLRDCNRRKKSKDGTATEVKSHELPQSAWNLDGHSCNPLIVEDLSYPRHMLVEILF

EKKSLLAGIAENIRGLGLTILKGVMEVKKDKVRACFVVEAAKDTHRLEVFWSLAHFVEQM

GKSTSSSGGECPLGAEILEENGA

>Pc2021171-Phaeoceros_carolinianus-sporophyte_1575|m.660

MSSILHQSLRGLCLKSGWSYAVMWKLKRRNRMVLTWEDGYYQPSPLFPLQSAVNEVQQCT

ELRKCTGGAIIGNAQQQDPLRQAIAKMSYHVYSLGEGIIGRVALTAKHQWVFGGGDKGGV

VGTGGSRPFGRTSMEYPSGWQNQFAAGIKTIAVIAVSQGVVQLGSTEIVMEDLDLVAHVK

SLFLSPKDLSESLPPSYSTPTRSPGVTVPSAQPHAAAKAQSATVMPASVTTDPDFKSITS

VGQRGPLCPPNMMLCGMTSEELLHQREGNQRMQQLMASVEPQTSLGNLSFKSQRVPGHGD

KVSQQHKNMQRVFADPCLTGTPRLLPENQFVGSDAHHVSSGELLSVFGARPQHSIQPLGL

SWMPNCETSNIHGQQIMVSGATPTVANGGTLMQGSVAQSPGTYSSVPCPPFKGHHMSKSM

DESGREALHEVLQDPTHVESGSSSWCAGPPLLNDETTMDVVSTKQLDQSSAGKNCRKRCR

PSEAPRPRPKDRQQIQDRVRELREIVPNSKKCSIDALLERTIKHVHLLQSVARIEGRFKD

EVQGEQNGSPKLAADDGREVVVQNLSQPRQMLVEMMCEKRELLLQLADHIRGLGLTIIKG

FMEVCCDKIRAQFVVEASRDMHRVEVLLTLMQLVKSSEMAV

>Ola2014084-Onychonema_laeve|m.267

MNLEGRLKEACLEFGWSYAVFWKPRLSPPPASLLWDDGFTVDYSSSSKLGALLACQGPES

PGRLDKKQLAVALAKMSYDTYHLGQGPVGAVALSGKAQWMFGSSQYKHAGISAAWSLQFS

AGIQTVALIPVAGGVLQVGSPLEISESSKMIARLTLLLAYPDGKAQPATPASTLSALSSS

SAFTSPTAGRCSMLGEEPLALPGREWGPAAPGAPVPGLGQVKLDFFSWGSQSPAGSQASR

VSQLSSSSPGLRSRLQALRQNRARMQLPEQQQQQPKLGQSLGQQQQQELPQLAGVRQPKL

ENLPLNWRLGSFPALPTGQLEKRTGNTSVAAAMPALENLPALETKTSAPFSATRHRASGF

VMGGEGQLSQEASQSMFVDDSLLEDDVDLGALGPLSHLGLPLSPLGGEEGARGKRQKVDH

GYVASFTLENIADLLLGEEHKAMEAKALPAAQLGQQQQQQEQDQQQGEGQEQQGEEHQHQ

HHHMEQQEQQRQQHQHLQLLQHKQPQHRPQLQRNVEGTPGSARAFRESQTSALRSTLPAW

TETFPNSRVTLGTSACTPFDELSQALGVGLTGQLSMRAADISSDVTIPVSPPLRVKEASG

SCSPGGGDSLALGQGPATSPNFSEGRAQGAEVESAQEGLGFGVGKPSEPQPVQEVVHQGD

HPEESGDSRPKSEAPTGCVLPAGASSGSHLLALAGTLAGAAGPSDTLTCSPLPGSGAGED

RALQGACSMLAGIKEKRQREKTAPAASLGEAPASPLGLPSVPGPGPALGALPPEALSNKL

APELGPASAGRARVANAESAGMSVLGQVWGFDGSGMADTDAAGVSVPGGSLGLPAPEGLL

GASTALQAYQRQMRQVLLFQQITAGKLALQPQAQSLALGEARGATLSEQTQALALGQAGR

GTLAEQSQGQALGQSAWGTLAEQVQALTRGHAGSGCPPEQAQTLALRMARQETLAEQAQA

LVRQRLLLGQRPLQLVQGLGRAGANAAAAAAQTRQPREQLQSRALTASPQVGGAPSLERD

SGSLLGQAGLIRLMQGHGPLAGKARSQLGLMSPVAAQQRCKPAPSSALQMLISRHRGVAG

ALAASRPAAAAAASIASAAETNALFGGKVLVGGPLRDALDPSDRLPAFPARGSLINSDQE

LASTMAASLASRVAAGASERQQVLTGILGVPAAAGRDVAGVGGRGLKRGCAGGANHRGGD

GKARPRDRQLIQERLQELRELLPNATKSSIEMLLERAMQRISFLRQATGQQTLILQASDP

PSNTEHVVLGHITVFRPSASLCAIQISVDVPRPIPMDAASALKSLGLLVRRGTLHNGARA

GSNAATFVAEVEDHVECAVVASHVEHFLGEQ

>Ccu2041916-Cylindrocystis_cushleckae|m.332

MQNNVAAMSMVLQQTLQGLCCGTAWNYAVFWKVKRTTRTVLIWEDGFHEFAGPKAFGGKS

GSTLDVDHQTLATSVAKMSYHVYSFGEGLIGRVSFTAKHQWVFAPNDRCVANHLPAQRVP

GRQTLFEKYPAGWQQQFDAGIKTIAVVAVPGGVVQLGSLQLMMEDLKLVSHMRSIFQTLE

CMPGAFLSDLVNNVPTAPGCVQAPFSPRGPTPLPVPGRINSHLLPAPQGNGSQVLPQTQP

MMLPPRVSLSELTKHPPSKQLLGSTAMQQMSQGRVGVTGEGFSMSNQASLQGLGLVWPPS

NVGAQTHGLNQPAGPLNNMRVPLQSLSRRDEQYPQGADIVAASRGVNGKAALASTMCIGN

TNGAIGAPFKDSLYPPTALEVPKRCASAGRRSRSRAIDQIALNVFSSDTPWGQCLPEAQG

PSLQTSPQVVPMAPPVLPPAAPPQLALPVTSFMTGQGPIGRQLRRKPPVEHPLPQSFPVS

LPGPNPGIVQPGATAMKPLPNGDALGRPFPELSVSVSAPFSTELGNESWTSPLEMATMGD

LEALGGLGHGLSEDQLLASLMQESSNSGANNTWLTDEMLKRQFCLQDLKDLTAPLEGPAD

VLGGINNIFGGNYGNVGTAHAQANLLVGGRSVSLLEEADILSSLHLPGFVGGPSQSPLVN

IMDGNRGLQMGELGELGTTLSQAPITEASKRPWDMYMTAKDALPSGELAEALGTGLTGQR

LQRAQGAPNVFSEDLSFLLKDSFVSAAESLSEQGRGQLSANRHAVASGCQVQAGQSGRVE

DLLSDGTRLDLASLEQQATQEVLLRSAGFLPPLNMPAAKSEERGDGVISPPSVGQPLPAP

VNVQAFSAVQNAQSQMIEQRDKHVGDGEPSLIEKIVSANMESLSDLKTPLGAGSSEAELG

AKGAKKAGTLTPRVKKHAPALMSPAGKCASKKVEAKQEVQESETTKGKKRVRGKCSEGSR

PRPRDRQQIQDRVADLRNLVPDSQKLSIDNLFEKTIQHLKYLQGAVQQRQCFKEVEGGSG

LSKRRCLEDALENSNGLDPFSVETLPHMPNHMAIEVLCESQELPLELLDTLHRMNLVVKT

GAMESRDGKLWARYIVEATREVQKMDVMWQLMHIVKGNLDMPPGTGGQSPAGSAILPANH

FAVSQA

>Cco2006162-Cosmocladium_cf._constrictum|m.447

MDLHGALASACVHLGWHYAAFWRARAASRSISLFWEDSYWRLEANDRAPPGTGAQEGNSR

TPALNSGLLLEALSKMTAEVYRPGEGTIGRAALTGKHQWMFAPSSMHSVTGALLPSSFQF

AAGIQTVVVVPVPGGVVQVGSLHVIPESSKLAAQLLLLFSSVELTPLQRTELLQPSAGAP

PPGLASAREAGQAAGVPPVSEIKLEYFSTSASATEHHQGLVPKPQLLEHQKQHHQQQQQR

FLHQQAPALRSRLQDLKLAREERLRQQQQQQQQHKPQTEGEGAQNQSGNYLSSSSSRVPL

RPGPTLRLSAPSRARPPSTLAFGAPPVPYPRLSPATFDELSQALGVGMTGVLPMQPSAAP

QSGPSLPALESEMLEELPGATGLLPPLVAQGCSMEAMRALAGPSPPLSPMPGLESTSHSP

GLCSGSPAGLAQPEGCQQMVPPLPSGPQGYLAAIPVANLEARGSSLDPGMGPTVSGGHRD

SRGAVGQPEGAAAALPSAQREILEQARQVELGTAEFSSTGSKAGELAAMEEVALKGARLM

VEGLMRKRQKTGDWQSLHWPHRPDRASQLALGETHKSPLQEPTGVASRALSACHQSASPL

PAAVGSPSATFRLLGPSLAEQRQAAVYAALAAPLSNVPEPYSLLRCGMEPAGSGFLRPSL

GSAGECSGSCELNLGTLSAPLQSSLLGVAWEKQQQSGAASALGLLPRPGPGGEGMKKDRE

IPTHLPNIWLHQHSSEAMRQSGAAPAELPECTGAVTSSSALMGLGFLQQQNLGQQPIGAR

VGRDAAGPGSSELWNPEEPTRAVKERFEVDRRRDSLFGTTAATLPLPLDCSIGQLPMGAA

SALGPQEVFEGSKRLGLSVPGGWAPAEGVLGAPELACSIAASLVPPSRSAYDPVDRESPR

TEGGGVVTVPSGAGSLPLGQPKGGRMRDTKARPRDRQLIQERLEELRSLLPSATKLSIEV

LLERTIQRICFLRQATHQQREIHKAMSVPRSLGRERPSAEAGPVLAILLTPALLAIRIQW

PSRRSAPVDAASWLSRIGFVVCKGATRSATVCTAEKASREHLEAAETLFVVENKGDCTCL

EAAISVHQEVERSG

>Po2002964-Planotaenium_ohtanii|m.457

MADVLQQTLQGICVGGGWDYAVFWKLTKRNRMVLIWEDAYCVNPSSSQPDNFSSLLPPFE

NSAGENREASGEDPLSWMMEANMKRKEKALLSTHNNVDYKALSFLIAKTSYQVFSYGEGV

IGRVAFTGKSQWVFAPASLRPGGKGEDASSALPSSFSSEKDSNNPFVNSSLKGSAGSRKG

SAFDPSPNVWSEHFAAGIRTIAIISVAEGVIQLGAGIDIPEEMAFVHQIRSLFSQLQKVP

GAFLSDSLNASFPSSAPASLSPLHPLVGHKRKPSFPPPPPSLLSPLSSSPGLPPDSNFSR

SHLLRQQQLQSLRRVSGVVQMDRAPAGWQQQRPYPATVSPQLPPSSQASRGSPAQRHGAL

PPSPAPSLSVPMSPAFTPAFSANCTPPPGLGPSPSISPFLSDQRPSRGAFVAPSGFSLPA

APSVTTHQREACASPVPRYGSAPRLSDDLSSSVRDALVSTLACKRASTSVMEGVQPKRNV

QIQMAQLQQLQRSRERRKQITSGAAGAGRQSQNHSFALPLPSNATREPGERELQTVEKAE

VLLAVASPGTGSHLEISTSVGCDAPLTPTTFLRAVNSQSLSPSFLPEATAHCARPALARE

VKVEESCNAPRQPPAPGALPSVDTSQVEDRALANVPTSESAYQLVEISTPQVGASPFALP

EPRFIESRRPPIGLDEIFSAPSRNYLSKRADESIPALTSDSQIARRDEARHEDGADLSVR

ENSAMPLQEENPMLDFSFLDEDALPSSPSFLIPRSPPALSQAFCDELSRALGNNMRGQIA

LEVAMQEEDKEDDEKFFPAARAHSQDDFAEVASRTSLQIAEDQRAPALFGVWKDCPLVSP

DPKDNLSGQRSSAKTAFAPKLERDSVSPVAPASRVAITHRPHGSSLAGSQLIESRDAPSP

VFLPLAPSSSPPSSSSSSDSHPNATMLRSSQSLSPTAPAPRGPSETNSSEDERDNVRSLR

PTXPPPPPPGGARSNSKRLRINIPAFPPLGECEGDSRRAANSTQRIAREGDISARSANAS

VPPLRSEGERDAKRARIGFHDLSGGTEEEPFSLEDLERQIALKALHRASSRKLLHHSLPP

SHTAASGQYFASSPPTSHSSLPQLVASPATSSAATHRVPATTWLDGSSHPTRLPPFAPVS

FSATPLGRPSAALEAVSRLRGSSVASAADPPLSLAPLPSSPVAPGHYTAVPRAPSLRGAV

EGALPFAMQPPILSSLPPPRLLSCSSNTSLLSASLASPPLYFSPSFSHAAQTVQQQALAA

VVPFSPSLCAPSLPTSAAPAIPPPSPSFQSPLLAVIQAKDRATLLRENECLIATVSNSLS

ASHLAKATAVAGGSYATEGGRNEKKGGAATGRDSGSTAKGKRSDGAVTAGGEDSPGLSQC

FEGAGKGGTAQSQSKPRPRDRQLLQDRVKELRGLIPGSEKLSIDSLLDCSIRHLVFLSSA

CNALRPVREKTENSAPLSLSSLARTTVGGISFLPVEPESQMKTTEKVLCLPGPAALSPAI

PPPRLFLLSFSGRPLAQRREGEGREGSGEGTSSNVESARALSSAAHPLPSNMDGSSLDQG

AATADVNSAFLSLLRRFDCSKFHVINTHCNRAWAKEGKDGEEIGGKVLAADSHQVTSDLW

RIQVLLGPKPIEGGEEGMKAGQEGSESASLDTPSDSPAHADDRDKESGGQIMSQLASCVR

NCFQVSCD

>Sc2002532-Staurodesmus_convergens|m.500

MDLDTGAVLPSTLTKLGAACALNKQHLANSLAKMSYEVHRPGEGPVGRTALSKKHMWVFA

PSAPHQAGAGLPASWSLQFAAGIQTIALVPVPGGVLQLGSNFLIPESSKMVARFVLLLTP

SEKKAVGRVACQPVTIGDLLGFSPCGVSHETPAVSPPPAPSPALLPGLSQVQLRFFSNAP

AAPSAHPVGGQGRPSGLRERLQSLQQARQRGLQRPGPGVQGQPALRQGQQPSSASRRAGG

EQPKLPQEGQGNQAPKQLPEGQERQLLLLLQRRMGMQVEPPVGQEQQLLLLAQGGQGHGG

GQLLAEGPKGREHEALLGRKDELKVGLFAALAGAGKELPQEQRLLHGQQLLTSLGAKRER

DDDADGGNARTAGIKGGMERPEETERAVAAGEEDAVKKPRIVSPPLCVEERQDDSSPLER

GMNVDVTRGDGGALSSLIPGASRPATVPRPATLSLQCIADLLFDDGKAVPGDFTASGLPS

PIDLVDAVSAVSPVNQLVRVQGGANPEASPREAFLGPRLKAAGALPASFDELSQALGVGL

SGCGSAVWKNFNLNGQLIGRGQPPSEMSCRVEEVTDAEEKDALPGAPKAPDARACCAAGA

PAADGTVAEGASPTARWGIVGQSPTVGVKLEGPQVLESAEGGDFTGAKDRAASPDVGAGS

FSLLMEESPSHSPPRGHPFTAEPGMLGGGGGSESAAMQASGPEASRGPGSSSETNVDLKP

PLAPVVGGEVQGLPMAGVDPIAAMEQMLAHGTRRQGAASGAAQRGPMLHLQKRLKMGGGE

STSDSSSLSPLPCPPALPALVSPALASFLSRRQRISGEACEALLEARDGEVQDGRKSTAG

ALLEAGGGVAPALNSSDLGLSGGQHLELSAEVQEMLRLQEIQRLRARLAVEVCPPPGALQ

GLDGLGPARLPQDLQAQAVHQTSSQALAAAILRNRARQAAVLQSVRASALAPPLGVPGVP

ALPGADALGPYAAMQLSLQRQLSSLTGVGGGSSIADAQGLLNQHSLRLQGVGSVLPSAQQ

LAAEQCLINRGMNRVPGEGLASLAGSASWGALEPAVGPTASGEAMLQNARLLALTQRSIF

QGPSSGPAAGDASGALGEAQGKAGSATLPTGQQLACSIAAALGRGLQEGALKGAGAAAAA

GAGGAAGDADKARGKAEDDGGAAGGGARGKGGEGRVRPRDRQLIQERLQELRELLPCATK

LSIEMLLERAMQRICFLRQATSQHKEIAQAECTAQGPAIGDGATTVCQLTPAVIKIKVTW

RPRVVVPLDSGSQLKDMGLVVLKGRLHCLRQPPPEDDAEDAMAVFIAEGSQSSCEEVTLK

MESLLEPL

>Mb2009311-Mesotaenium_braunii|m.555

MASSMMLRQNLQDLCKGTGWNYALFWKLRQSGSKEERFFWEGCYHEFEAAAPGLYHDASG

RLLRGREGSDYKGLCDTMSKMIYHMYTLGAGLMGQVAFKKSYHVAYAPDCLGGGTVTQFP

HGWHEQFTAGIKTIAVAYVPGGLLQLGSLDELRFTELDFKHVTMVFSALQHELPSALAPG

FPVEGLPRSLPCAAATTMGGLGLQYGGLLASHPPIPSGPLFNADGGVSALQRHSAMVSAG

VVPVPDGSAGLSGPLQGLSLPGGRLPDHIVNPRRITLEEAMRGKPVGVRMGSVLEPGPGS

LYSSQAPPGDYHMLTRTAGNPSLHTLPAFQALQQQPPLVGSQAPLAAGVPLSSTPGVEMS

GPGHGACRVEADTVPVSRGAGRWRASRSQCVRAAPIESIQLAALKGTGVSGASVPIPVTN

GTPPLSSQTLASLEDAQSRAAAVLSFGSPPQGWGSFQSQSPLEVPTFQGGAQLAIDLSAL

PQPAVMPYPSLPFADLSRQQLHPSCAATGDGSLSLEQEDGPSCPVDSLHDWDANALPLAF

SSAQLGSAEVGADIGGDLSAPGAVAYEAEWRGPGGDRVPACSSVDLFPESSGGGGGAPTG

PSTLVSSCIINVERDAGVKLLQHGSLGGLQVPPLGAGGDGVTKGGGPSQLPLLQSGSSML

ASVQSLPTGKEFERSPWAPFCADSGERRPLVQQEHGGQQDCSHPDMYLYSAREEEGEQLR

PQGFGGCLPADLASMPLLLPGGGGFDDELANAVVRFTGSFQGSSANYIDDNFDISLLLST

EQQLLLQQQQQQQQQQQGFSYGGKGALSTGQDFPGTGFSASLSTAAGPLDPLPSITPQVI

PLAGQRAFMHESLYAQLQENLQPVKSVSSLPLPGATTQHESRPHSPLLQPIGGGLGGHSF

LSNGRAVSQATPPDYQQTAVYTADNKLALGVVPPPPPPPMLPSPPFQGGPQLCNPQQVQS

AADPLARYLCPMERTGVSGQLGDALFASTIPSAAGPMSSSKVPGLDPVQLKEIATLLDAQ

QGDPVAEYDREETRGRKRRAGAAVLCPQEKAERRRPRPRDRQQIQDRTAELRGLIPQSDK

CSIDVLYGRTLDHLKFLDQAMAGHQRFREVEEQMMSSEAASPGAGAGTQSAGGILRLCHV

DGSGEKQAYVKFLCDSQLLSWEVPDSLEGLGLEICKVAMECSGGVMTMSLFVEAKEGSVD

LKALTAHLEQVRPTPAGLAVLPAGC

>Tl2080512-Takakia_lepidozioides|m.1219

LHRTLRGLCFKTGWTYAVFWKLKRRNRMVLTWEDGYYEYPKLPVMESSPLLEGSHALLPG

GQYPPGEGADHLMGGAAAAEDRIGLAVAKMSYHVYSLGEGIIGRVAFTHKHQWIFGNAEK

PGCGDGTTGSRTSSRPILEKYPAGWQHQFSAGVKTIAVVAVPQGVVQLGSTQLIKEDLSL

VAHVRALFGALQSVPGAFLSDYVPELQAGQVQGVYPLGMPMHHTSKMSTTSMLQVQAGQV

NSLSRVAMGPVGSSPLAMQNSVKMKNGPSALTGISPISSLKQMHMHSQYQNSSSDSSFHT

NGTVHVPQQQQASPLLGQSHSWQVAMSGGSHLGTSGFVPTPPHECHPWRNFGPVSGENTS

VASSLVELCAMTGINTSVHPTAWACYEGTGLNLAEQNEAKSQGQLQSIAEKLHSEFRELQ

PNFNRPVRIQQASHMEEKGNFVGQVPVNLPVLRSGESLLFPESLGSMDTAVVNPRDSVLH

PWTMVGVPNKRDEERVLPSEKQGLSSLPVSQLSSGHFSPSRGLPPTTPYDLLSMLPEHSD

VPKSKGLSLTSSLQYNGPARGFDFPTLDTPDSVRLPPLGVCLDAMGHEEQLVRKWGKPTE

VVTADGLLPLTGNTKYPELTFGGDLCGAMVGSPQLGTDDLKAYIASLTLAHCQQQDQTSR

LSQFAISDELSMALCPSLKRGTERILENSLARDDTEKFGTSLDGLPAPFDSRLLLDRVRC

GSGLRDEMQENSGPNVPSLDEVESKPIQDATLAVAVSVSGITNIPTSSMCYNIDSHCWNQ

LQPHCKTGHQVNLEVQSTSRLNLDDCDSNPMQLSRLSTEILHRPSALLSLPMQDETRLFD

QPQVTHSEGAERMWPRAGQLHQSKILASNCHREGSMKWFETHSLLKAGVSAQCTDSLKDK

DEEGSTTESSGAKMQEEVGKLGRKRARSGENPRPRPKDRQQIQDRVQELREIVPNVTKCS

IDALLERTIKHMHFLQSVTQQGHKWKQAGEFKGHDMESNLVTTQNDLESGASWALEVGGQ

GIGCPIIVENLSQPHQFLVEMLCEENELFLEIADNIRGLGLTILKGIMKARNDKIWARFM

VEALKEVHRLDVLWSLTQLLHPDTKMG

>Ba2070572-Buxbaumia_aphylla|m.1162

MMMETPLASFWDADSAMIEAFMGSYGISSYEAQDELASTGEKGAELGDTIWLERLHLLVE

NAPVNWTYGIFWQLSPSPAGELILGWGDGYFKGPKENELNDLQQVRRGGSEEDQQLRRKV

LRELQALVSNSEDDISDDVTDTEWFYLVSMSHSFARGVGIPGQALATNRHVWLIEANKAP

NHCTRAPLAKMAGIQTIVCVPTKSGVVELGSTDLITEDWEVVEHIKMVFEDSLWGANRSQ

TMTQSLDTSYIPPSPSIMSLTTTSVLTPSPSIASRGSNPGKDHEAHYTGRGALIDKVGSS

MTSTNLDNLEYLWSRSEDAQFNDMGSANTVDKDRGHSGMHYTPRKPPMQEEVHPTSASSI

LSRNRVPEGKHTSTSPNLNRFSFEEQRPSLPYVKVHPSFTYSQNPGVGEVSQAINLSGPR

QSAGLKPPMLEEKVHVPVMNGGDVRHLAENPKNASKPSQQHQSMMSGPPVSGSGRSGFDQ

SEHDCQESEAEVSFKESPVDFSLNVGTKPPRKRGRKPANDREEPLSHVQAERQRREKLNQ

RFYALRAVVPNVSKMDKASLLGDAIAYINELQSKLQMAEGQIKDLKSRSAVAFDKPQDSL

SLGRGSISNPTKETLNLRPQGSGNNLLGHAKGGVDDKKPSISVHILNQEAMIRINCMKDS

YVIAHMMMILQDLRLEIRHSNTSVMQDMVLHIVIVKIDPAEHYTQEKLRTVLEKSCLSYR

SATNEEDRGHFEKLGSSRQSNSTLSLTP

>At1G04840

MKSLSVIFKPKSSPAKIYFPADRQASPDESHFISLIHACKDTASLRHVHAQILRRGVLSSRVAAQLVSCSSLLKSPDYSLSIFRNSEERNPFVLNALIRGLTENARFESSVRHFILMLRLGVKPDRLTFPFVLKSNSKLGFRWLGRALHAATLKNFVDCDSFVRLSLVDMYAKTGQLKHAFQVFEESPDRIKKESILIWNVLINGYCRAKDMHMATTLFRSMPERNSGSWSTLIKGYVDSGELNRAKQLFELMPEKNVVSWTTLINGFSQTGDYETAISTYFEMLEKGLKPNEYTIAAVLSACSKSGALGSGIRIHGYILDNGIKLDRAIGTALVDMYAKCGELDCAATVFSNMNHKDILSWTAMIQGWAVHGRFHQAIQCFRQMMYSGEKPDEVVFLAVLTACLNSSEVDLGLNFFDSMRLDYAIEPTLKHYVLVVDLLGRAGKLNEAHELVENMPINPDLTTWAALYRACKAHKGYRRAESVSQNLLELDPELCGSYIFLDKTHASKGNIQDVEKRRLSLQKRIKERSLGWSYIELDGQLNKFSAGDYSHKLTQEIGLKLDEIISLAIQKGYNPGADWSIHDIEEEEKENVTGIHSEKLALTLGFLRTAPGTTIRIIKNLRICGDCHSLMKYVSKISQRDILLRDARQFHHFKDGRCSCGDYW

>At1G05750

MGLLPVVGITSPALITHKNHANPKIQRHNQSTSETTVSWTSRINLLTRNGRLAEAAKEFSDMTLAGVEPNHITFIALLSGCGDFTSGSEALGDLLHGYACKLGLDRNHVMVGTAIIGMYSKRGRFKKARLVFDYMEDKNSVTWNTMIDGYMRSGQVDNAAKMFDKMPERDLISWTAMINGFVKKGYQEEALLWFREMQISGVKPDYVAIIAALNACTNLGALSFGLWVHRYVLSQDFKNNVRVSNSLIDLYCRCGCVEFARQVFYNMEKRTVVSWNSVIVGFAANGNAHESLVYFRKMQEKGFKPDAVTFTGALTACSHVGLVEEGLRYFQIMKCDYRISPRIEHYGCLVDLYSRAGRLEDALKLVQSMPMKPNEVVIGSLLAACSNHGNNIVLAERLMKHLTDLNVKSHSNYVILSNMYAADGKWEGASKMRRKMKGLGLKKQPGFSSIEIDDCMHVFMAGDNAHVETTYIREVLELISSDLRLQGCVVETLAGDLLNA

>At1G06150

MGYTLQQILRSICSNTDWNYAVFWKLNHHSPMVLTLEDVYCVNHERGLMPESLHGGRHAHDPLGLAVAKMSYHVHSLGEGIVGQVAISGQHQWIFSEYLNDSHSTLQVHNGWESQISAGIKTILIVAVGSCGVVQLGSLCKVEEDPALVTHIRHLFLALTDPLADHASNLMQCDINSPSDRPKIPSKCLHEASPDFSGEFDKAMDMEGLNIVSQNTSNRSNDLPYNFTPTYFHMERTAQVIGGLEAVQPSMFGSNDCVTSGFSVGVVDTKHKNQVDISDMSKVIYDEETGGYRYSRELDPNFQHYSRNHVRNSGGTSALAMESDRLKAGSSYPQLDSTVLTALKTDKDYSRRNEVFQPSESQGSIFVKDTEHRQEEKSESSQLDALTASLCSFSGSELLEALGPAFSKTSTDYGELAKFESAAAIRRTNDMSHSHLTFESSSENLLDAVVASMSNGDGNVRREISSSRSTQSLLTTAEMAQAEPFGHNKQNIVSTVDSVISQPPLADGLIQQNPSNICGAFSSIGFSSTCLSSSSDQFPTSLEIPKKNKKRAKPGESSRPRPRDRQLIQDRIKELRELVPNGSKCSIDSLLECTIKHMLFLQSVSQHADKLTKSASSKMQHKDTGTLGISSTEQGSSWAVEIGGHLQVCSIMVENLDKEGVMLIEMLCEECSHFLEIANVIRSLELIILRGTTEKQGEKTWICFVVEGQNNKVMHRMDILWSLVQIFQPKATNSLHLYRQSQILYMNAFANVHSLRVPSHHLRDFSASLSLAPPNLKKIIKQCSTPKLLESALAAMIKTSLNQDCRLMNQFITACTSFKRLDLAVSTMTQMQEPNVFVYNALFKGFVTCSHPIRSLELYVRMLRDSVSPSSYTYSSLVKASSFASRFGESLQAHIWKFGFGFHVKIQTTLIDFYSATGRIREARKVFDEMPERDDIAWTTMVSAYRRVLDMDSANSLANQMSEKNEATSNCLINGYMGLGNLEQAESLFNQMPVKDIISWTTMIKGYSQNKRYREAIAVFYKMMEEGIIPDEVTMSTVISACAHLGVLEIGKEVHMYTLQNGFVLDVYIGSALVDMYSKCGSLERALLVFFNLPKKNLFCWNSIIEGLAAHGFAQEALKMFAKMEMESVKPNAVTFVSVFTACTHAGLVDEGRRIYRSMIDDYSIVSNVEHYGGMVHLFSKAGLIYEALELIGNMEFEPNAVIWGALLDGCRIHKNLVIAEIAFNKLMVLEPMNSGYYFLLVSMYAEQNRWRDVAEIRGRMRELGIEKICPGTSSIRIDKRDHLFAAADKSHSASDEVCLLLDEIYDQMGLAGYVQETENVY

>At1G08070

MMLSCSPLTVPSSSYPFHFLPSSSDPPYDSIRNHPSLSLLHNCKTLQSLRIIHAQMIKIGLHNTNYALSKLIEFCILSPHFEGLPYAISVFKTIQEPNLLIWNTMFRGHALSSDPVSALKLYVCMISLGLLPNSYTFPFVLKSCAKSKAFKEGQQIHGHVLKLGCDLDLYVHTSLISMYVQNGRLEDAHKVFDKSPHRDVVSYTALIKGYASRGYIENAQKLFDEIPVKDVVSWNAMISGYAETGNYKEALELFKDMMKTNVRPDESTMVTVVSACAQSGSIELGRQVHLWIDDHGFGSNLKIVNALIDLYSKCGELETACGLFERLPYKDVISWNTLIGGYTHMNLYKEALLLFQEMLRSGETPNDVTMLSILPACAHLGAIDIGRWIHVYIDKRLKGVTNASSLRTSLIDMYAKCGDIEAAHQVFNSILHKSLSSWNAMIFGFAMHGRADASFDLFSRMRKIGIQPDDITFVGLLSACSHSGMLDLGRHIFRTMTQDYKMTPKLEHYGCMIDLLGHSGLFKEAEEMINMMEMEPDGVIWCSLLKACKMHGNVELGESFAENLIKIEPENPGSYVLLSNIYASAGRWNEVAKTRALLNDKGMKKVPGCSSIEIDSVVHEFIIGDKFHPRNREIYGMLEEMEVLLEKAGFVPDTSEVLQEMEEEWKEGALRHHSEKLAIAFGLISTKPGTKLTIVKNLRVCRNCHEATKLISKIYKREIIARDRTRFHHFRDGVCSCNDYW

>At1G09190

MEIERKLLRLLHGHNTRTRLPEIHAHLLRHFLHGSNLLLAHFISICGSLSNSDYANRVFSHIQNPNVLVFNAMIKCYSLVGPPLESLSFFSSMKSRGIWADEYTYAPLLKSCSSLSDLRFGKCVHGELIRTGFHRLGKIRIGVVELYTSGGRMGDAQKVFDEMSERNVVVWNLMIRGFCDSGDVERGLHLFKQMSERSIVSWNSMISSLSKCGRDREALELFCEMIDQGFDPDEATVVTVLPISASLGVLDTGKWIHSTAESSGLFKDFITVGNALVDFYCKSGDLEAATAIFRKMQRRNVVSWNTLISGSAVNGKGEFGIDLFDAMIEEGKVAPNEATFLGVLACCSYTGQVERGEELFGLMMERFKLEARTEHYGAMVDLMSRSGRITEAFKFLKNMPVNANAAMWGSLLSACRSHGDVKLAEVAAMELVKIEPGNSGNYVLLSNLYAEEGRWQDVEKVRTLMKKNRLRKSTGQSTICDVSV

>At1G09410

MKSQILLRRTYSTTIPPPTANVRITHLSRIGKIHEARKLFDSCDSKSISSWNSMVAGYFANLMPRDARKLFDEMPDRNIISWNGLVSGYMKNGEIDEARKVFDLMPERNVVSWTALVKGYVHNGKVDVAESLFWKMPEKNKVSWTVMLIGFLQDGRIDDACKLYEMIPDKDNIARTSMIHGLCKEGRVDEAREIFDEMSERSVITWTTMVTGYGQNNRVDDARKIFDVMPEKTEVSWTSMLMGYVQNGRIEDAEELFEVMPVKPVIACNAMISGLGQKGEIAKARRVFDSMKERNDASWQTVIKIHERNGFELEALDLFILMQKQGVRPTFPTLISILSVCASLASLHHGKQVHAQLVRCQFDVDVYVASVLMTMYIKCGELVKSKLIFDRFPSKDIIMWNSIISGYASHGLGEEALKVFCEMPLSGSTKPNEVTFVATLSACSYAGMVEEGLKIYESMESVFGVKPITAHYACMVDMLGRAGRFNEAMEMIDSMTVEPDAAVWGSLLGACRTHSQLDVAEFCAKKLIEIEPENSGTYILLSNMYASQGRWADVAELRKLMKTRLVRKSPGCSWTEVENKVHAFTRGGINSHPEQESILKILDELDGLLREAGYNPDCSYALHDVDEEEKVNSLKYHSERLAVAYALLKLSEGIPIRVMKNLRVCSDCHTAIKIISKVKEREIILRDANRFHHFRNGECSCKDYW

>At1G11290

MSSQLVQFSTVPQIPNPPSRHRHFLSERNYIPANVYEHPAALLLERCSSLKELRQILPLVFKNGLYQEHFFQTKLVSLFCRYGSVDEAARVFEPIDSKLNVLYHTMLKGFAKVSDLDKALQFFVRMRYDDVEPVVYNFTYLLKVCGDEAELRVGKEIHGLLVKSGFSLDLFAMTGLENMYAKCRQVNEARKVFDRMPERDLVSWNTIVAGYSQNGMARMALEMVKSMCEENLKPSFITIVSVLPAVSALRLISVGKEIHGYAMRSGFDSLVNISTALVDMYAKCGSLETARQLFDGMLERNVVSWNSMIDAYVQNENPKEAMLIFQKMLDEGVKPTDVSVMGALHACADLGDLERGRFIHKLSVELGLDRNVSVVNSLISMYCKCKEVDTAASMFGKLQSRTLVSWNAMILGFAQNGRPIDALNYFSQMRSRTVKPDTFTYVSVITAIAELSITHHAKWIHGVVMRSCLDKNVFVTTALVDMYAKCGAIMIARLIFDMMSERHVTTWNAMIDGYGTHGFGKAALELFEEMQKGTIKPNGVTFLSVISACSHSGLVEAGLKCFYMMKENYSIELSMDHYGAMVDLLGRAGRLNEAWDFIMQMPVKPAVNVYGAMLGACQIHKNVNFAEKAAERLFELNPDDGGYHVLLANIYRAASMWEKVGQVRVSMLRQGLRKTPGCSMVEIKNEVHSFFSGSTAHPDSKKIYAFLEKLICHIKEAGYVPDTNLVLGVENDVKEQLLSTHSEKLAISFGLLNTTAGTTIHVRKNLRVCADCHNATKYISLVTGREIVVRDMQRFHHFKNGACSCGDYW

>At1G13410

MITSLRDSSLLVAESRELITHAKCSTESLNQMFLFGMLCLMGVIASANKVFCEMVEKNVVLWTSMINGYLLNKDLVSARRYFDLSPERDIVLWNTMISGYIEMGNMLEARSLFDQMPCRDVMSWNTVLEGYANIGDMEACERVFDDMPERNVFSWNGLIKGYAQNGRVSEVLGSFKRMVDEGSVVPNDATMTLVLSACAKLGAFDFGKWVHKYGETLGYNKVDVNVKNALIDMYGKCGAIEIAMEVFKGIKRRDLISWNTMINGLAAHGHGTEALNLFHEMKNSGISPDKVTFVGVLCACKHMGLVEDGLAYFNSMFTDFSIMPEIEHCGCVVDLLSRAGFLTQAVEFINKMPVKADAVIWATLLGASKVYKKVDIGEVALEELIKLEPRNPANFVMLSNIYGDAGRFDDAARLKVAMRDTGFKKEAGVSWIETDDGLVKFYSSGEKHPRTEELQRILRELKSFNILRDEEHFM

>At1G27740

MDVFVDGELESLLGMFNFDQCSSSKEERPRDELLGLSSLYNGHLHQHQHHNNVLSSDHHAFLLPDMFPFGAMPGGNLPAMLDSWDQSHHLQETSSLKRKLLDVENLCKTNSNCDVTRQELAKSKKKQRVSSESNTVDESNTNWVDGQSLSNSSDDEKASVTSVKGKTRATKGTATDPQSLYARKRREKINERLKTLQNLVPNGTKVDISTMLEEAVHYVKFLQLQIKLLSSDDLWMYAPLAYNGLDMGFHHNLLSRLM

>At1G50270

MIELKTLLDLPLHFLHLKQIHCLLLTSPIFYTRRDLFLSRLLRRCCTAATQFRYARRLLCQLQTLSIQLWDSLIGHFSGGITLNRRLSFLAYRHMRRNGVIPSRHTFPPLLKAVFKLRDSNPFQFHAHIVKFGLDSDPFVRNSLISGYSSSGLFDFASRLFDGAEDKDVVTWTAMIDGFVRNGSASEAMVYFVEMKKTGVAANEMTVVSVLKAAGKVEDVRFGRSVHGLYLETGRVKCDVFIGSSLVDMYGKCSCYDDAQKVFDEMPSRNVVTWTALIAGYVQSRCFDKGMLVFEEMLKSDVAPNEKTLSSVLSACAHVGALHRGRRVHCYMIKNSIEINTTAGTTLIDLYVKCGCLEEAILVFERLHEKNVYTWTAMINGFAAHGYARDAFDLFYTMLSSHVSPNEVTFMAVLSACAHGGLVEEGRRLFLSMKGRFNMEPKADHYACMVDLFGRKGLLEEAKALIERMPMEPTNVVWGALFGSCLLHKDYELGKYAASRVIKLQPSHSGRYTLLANLYSESQNWDEVARVRKQMKDQQVVKSPGFSWIEVKGKLCEFIAFDDKKPLESDDLYKTLDTVGVQMRLPDELEDVTAES

>At1G56690

MKRLKLILRRTYLTSTGVNCSFEISRLSRIGKINEARKFFDSLQFKAIGSWNSIVSGYFSNGLPKEARQLFDEMSERNVVSWNGLVSGYIKNRMIVEARNVFELMPERNVVSWTAMVKGYMQEGMVGEAESLFWRMPERNEVSWTVMFGGLIDDGRIDKARKLYDMMPVKDVVASTNMIGGLCREGRVDEARLIFDEMRERNVVTWTTMITGYRQNNRVDVARKLFEVMPEKTEVSWTSMLLGYTLSGRIEDAEEFFEVMPMKPVIACNAMIVGFGEVGEISKARRVFDLMEDRDNATWRGMIKAYERKGFELEALDLFAQMQKQGVRPSFPSLISILSVCATLASLQYGRQVHAHLVRCQFDDDVYVASVLMTMYVKCGELVKAKLVFDRFSSKDIIMWNSIISGYASHGLGEEALKIFHEMPSSGTMPNKVTLIAILTACSYAGKLEEGLEIFESMESKFCVTPTVEHYSCTVDMLGRAGQVDKAMELIESMTIKPDATVWGALLGACKTHSRLDLAEVAAKKLFENEPDNAGTYVLLSSINASRSKWGDVAVVRKNMRTNNVSKFPGCSWIEVGKKVHMFTRGGIKNHPEQAMILMMLEKTDGLLREAGYSPDCSHVLHDVDEEEKVDSLSRHSERLAVAYGLLKLPEGVPIRVMKNLRVCGDCHAAIKLISKVTEREIILRDANRFHHFNNGECSCRDYW

>At1G60060

MEEHLNPLAVTHLLQHTLRSLCIHENSQWVYAVFWRILPRNYPPPKWDGQGAYDRSRGNRRNWILVWEDGFCNFAASAAEMSSGEGSGGGGGSAAYGNSDFQQYQGLQPELFFKMSHEIYNYGEGLIGKVAADHSHKWIYKEPNDQEINFLSAWHNSADSYPRTWEAQFQSGIKTIALISVREGVVQLGAVHKVIEDLSYVVMLRKKLSYIESIPGVLLPHPSSSGYPFINASPSDTWHFPGVAPPHQQPEHQFYHSDHNHRFLIGHHNQPQAVGGAAPPLPLSMKITPSMSSLEALLSKLPSVVPPATQPGYYPFHHSAKEEMSQEEQNDAFRVERNDLVGEGSNNHNHNNNYNSNNDIYNYSNNCSNNNYDRENKIGGFLSEDY

>At1G62914

MLAKISSSAKRFVHRSLVVRGNAATFPLSFSFCRRRAFSGKTSYDYREVLRTGLSDIELDDAIGLFGVMAQSRPFPSIIEFSKLLSAIAKMNKFDLVISFGEKMEILGISHNLYTYNILINCFCRCSRLSLALALLGKMMKLGYEPDIVTLNSLLNGFCHGNRISDAVALVDQMVEMGYKPDTVTFTTLIHGLFLHNKASEAVALIDRMVQRGCQPDLVTYGAVVNGLCKRGDTDLALNLLNKMEAAKIEANVVIYSTVIDSLCKYRHEDDALNLFTEMENKGVRPNVITYSSLISCLCNYGRWSDASRLLSDMIERKINPNLVTFSALIDAFVKKGKLVKAEKLYEEMIKRSIDPNIFTYSSLINGFCMLDRLGEAKQMLELMIRKDCLPNVVTYNTLINGFCKAKRVDKGMELFREMSQRGLVGNTVTYTTLIHGFFQARDCDNAQMVFKQMVSVGVHPNILTYNILLDGLCKNGKLAKAMVVFEYLQRSTMEPDIYTYNIMIEGMCKAGKWKMGGIYFVASALKE

>At1G63650

MATGENRTVPDNLKKQLAVSVRNIQWSYGIFWSVSASQPGVLEWGDGYYNGDIKTRKTIQAAEVKIDQLGLERSEQLRELYESLSLAESSASGSSQVTRRASAAALSPEDLTDTEWYYLVCMSFVFNIGEGIPGGALSNGEPIWLCNAETADSKVFTRSLLAKSASLQTVVCFPFLGGVLEIGTTEHIKEDMNVIQSVKTLFLEAPPYTTISTRSDYQEIFDPLSDDKYTPVFITEAFPTTSTSGFEQEPEDHDSFINDGGASQVQSWQFVGEEISNCIHQSLNSSDCVSQTFVGTTGRLACDPRKSRIQRLGQIQEQSNHVNMDDDVHYQGVISTIFKTTHQLILGPQFQNFDKRSSFTRWKRSSSVKTLGEKSQKMIKKILFEVPLMNKKEELLPDTPEETGNHALSEKKRREKLNERFMTLRSIIPSISKIDKVSILDDTIEYLQDLQKRVQELESCRESADTETRITMMKRKKPDDEEERASANCMNSKRKGSDVNVGEDEPADIGYAGLTDNLRISSLGNEVVIELRCAWREGILLEIMDVISDLNLDSHSVQSSTGDGLLCLTVNCKHKGTKIATTGMIQEALQRVAWIC

>At1G64625

MGSEYKHILKSLCLSHGWSYAVFWRYDPINSMILRFEEAYNDEQSVALVDDMVLQAPILGQGIVGEVASSGNHQWLFSDTLFQWEHEFQNQFLCGFKILIRQFTYTQTIAIIPLGSSGVVQLGSTQKILESTEILEQTTRALQETCLKPHDSGDLDTLFESLGDCEIFPAESFQGFSFDDIFAEDNPPSLLSPEMISSEAASSNQDLTNGDDYGFDILQSYSLDDLYQLLADPPEQNCSSMVIQGVDKDLFDILGMNSQTPTMALPPKGLFSELISSSLSNNTCSSSLTNVQEYSGVNQSKRRKLDTSSAHSSSLFPQEETVTSRSLWIDDDERSSIGGNWKKPHEEGVKKKRAKAGESRRPRPKDRQMIQDRIKELRGMIPNGAKCSIDTLLDLTIKHMVFMQSLAKYAERLKQPYESKLVKEKERTWALEVGEEGVVCPIMVEELNREGEMQIEMVCEEREEFLEIGQVVRGLGLKILKGVMETRKGQIWAHFIVQAKPQVTRIQVLYSLVQLFQHHTKHDDLLS

>At1G66470

MALVNDHPNETNYLSKQNSSSSEDLSSPGLDQPDAAYAGGGGGGGSASSSSTMNSDHQQHQGFVFYPSGEDHHNSLMDFNGSSFLNFDHHESFPPPAISCGGSSGGGGFSFLEGNNMSYGFTNWNHQHHMDIISPRSTETPQGQKDWLYSDSTVVTTGSRNESLSPKSAGNKRSHTGESTQPSKKLSSGVTGKTKPKPTTSPKDPQSLAAKNRRERISERLKILQELVPNGTKVDLVTMLEKAISYVKFLQVQVKVLATDEFWPAQGGKAPDISQVKDAIDAILSSSQRDRNSNLITN

>At1G74630

MTIAIHHCLSLLNSCKNLRALTQIHGLFIKYGVDTDSYFTGKLILHCAISISDALPYARRLLLCFPEPDAFMFNTLVRGYSESDEPHNSVAVFVEMMRKGFVFPDSFSFAFVIKAVENFRSLRTGFQMHCQALKHGLESHLFVGTTLIGMYGGCGCVEFARKVFDEMHQPNLVAWNAVITACFRGNDVAGAREIFDKMLVRNHTSWNVMLAGYIKAGELESAKRIFSEMPHRDDVSWSTMIVGIAHNGSFNESFLYFRELQRAGMSPNEVSLTGVLSACSQSGSFEFGKILHGFVEKAGYSWIVSVNNALIDMYSRCGNVPMARLVFEGMQEKRCIVSWTSMIAGLAMHGQGEEAVRLFNEMTAYGVTPDGISFISLLHACSHAGLIEEGEDYFSEMKRVYHIEPEIEHYGCMVDLYGRSGKLQKAYDFICQMPIPPTAIVWRTLLGACSSHGNIELAEQVKQRLNELDPNNSGDLVLLSNAYATAGKWKDVASIRKSMIVQRIKKTTAWSLVEVGKTMYKFTAGEKKKGIDIEAHEKLKEIILRLKDEAGYTPEVASALYDVEEEEKEDQVSKHSEKLALAFALARLSKGANIRIVKNLRICRDCHAVMKLTSKVYGVEILVRDRNRFHSFKDGSCSCRDYW

>At2G01510

MLAKQTPLTKQMLSELLRASSSKPKQLKKIHAIVLRTGFSEKNSLLTQLLENLVVIGDMCYARQVFDEMHKPRIFLWNTLFKGYVRNQLPFESLLLYKKMRDLGVRPDEFTYPFVVKAISQLGDFSCGFALHAHVVKYGFGCLGIVATELVMMYMKFGELSSAEFLFESMQVKDLVAWNAFLAVCVQTGNSAIALEYFNKMCADAVQFDSFTVVSMLSACGQLGSLEIGEEIYDRARKEEIDCNIIVENARLDMHLKCGNTEAARVLFEEMKQRNVVSWSTMIVGYAMNGDSREALTLFTTMQNEGLRPNYVTFLGVLSACSHAGLVNEGKRYFSLMVQSNDKNLEPRKEHYACMVDLLGRSGLLEEAYEFIKKMPVEPDTGIWGALLGACAVHRDMILGQKVADVLVETAPDIGSYHVLLSNIYAAAGKWDCVDKVRSKMRKLGTKKVAAYSSVEFEGKIHFFNRGDKSHPQSKAIYEKLDEILKKIRKMGYVPDTCSVFHDVEMEEKECSLSHHSEKLAIAFGLIKGRPGHPIRVMKNLRTCDDCHAFSKFVSSLTSTEIIMRDKNRFHHFRNGVCSCKEFW

>At2G02980

MAISSASLISSFSHAETFTKHSKIDTVNTQNPILLISKCNSLRELMQIQAYAIKSHIEDVSFVAKLINFCTESPTESSMSYARHLFEAMSEPDIVIFNSMARGYSRFTNPLEVFSLFVEILEDGILPDNYTFPSLLKACAVAKALEEGRQLHCLSMKLGLDDNVYVCPTLINMYTECEDVDSARCVFDRIVEPCVVCYNAMITGYARRNRPNEALSLFREMQGKYLKPNEITLLSVLSSCALLGSLDLGKWIHKYAKKHSFCKYVKVNTALIDMFAKCGSLDDAVSIFEKMRYKDTQAWSAMIVAYANHGKAEKSMLMFERMRSENVQPDEITFLGLLNACSHTGRVEEGRKYFSQMVSKFGIVPSIKHYGSMVDLLSRAGNLEDAYEFIDKLPISPTPMLWRILLAACSSHNNLDLAEKVSERIFELDDSHGGDYVILSNLYARNKKWEYVDSLRKVMKDRKAVKVPGCSSIEVNNVVHEFFSGDGVKSATTKLHRALDEMVKELKLSGYVPDTSMVVHANMNDQEKEITLRYHSEKLAITFGLLNTPPGTTIRVVKNLRVCRDCHNAAKLISLIFGRKVVLRDVQRFHHFEDGKCSCGDFW

>At2G03880

MKSVMSKIKLFRPVVTLRCSYSSTDQTLLLSEFTRLCYQRDLPRAMKAMDSLQSHGLWADSATYSELIKCCISNRAVHEGNLICRHLYFNGHRPMMFLVNVLINMYVKFNLLNDAHQLFDQMPQRNVISWTTMISAYSKCKIHQKALELLVLMLRDNVRPNVYTYSSVLRSCNGMSDVRMLHCGIIKEGLESDVFVRSALIDVFAKLGEPEDALSVFDEMVTGDAIVWNSIIGGFAQNSRSDVALELFKRMKRAGFIAEQATLTSVLRACTGLALLELGMQAHVHIVKYDQDLILNNALVDMYCKCGSLEDALRVFNQMKERDVITWSTMISGLAQNGYSQEALKLFERMKSSGTKPNYITIVGVLFACSHAGLLEDGWYYFRSMKKLYGIDPVREHYGCMIDLLGKAGKLDDAVKLLNEMECEPDAVTWRTLLGACRVQRNMVLAEYAAKKVIALDPEDAGTYTLLSNIYANSQKWDSVEEIRTRMRDRGIKKEPGCSWIEVNKQIHAFIIGDNSHPQIVEVSKKLNQLIHRLTGIGYVPETNFVLQDLEGEQMEDSLRHHSEKLALAFGLMTLPIEKVIRIRKNLRICGDCHVFCKLASKLEIRSIVIRDPIRYHHFQDGKCSCGDYW

>At2G13600

MATKSFLKLAADLSSFTDSSPFAKLLDSCIKSKLSAIYVRYVHASVIKSGFSNEIFIQNRLIDAYSKCGSLEDGRQVFDKMPQRNIYTWNSVVTGLTKLGFLDEADSLFRSMPERDQCTWNSMVSGFAQHDRCEEALCYFAMMHKEGFVLNEYSFASVLSACSGLNDMNKGVQVHSLIAKSPFLSDVYIGSALVDMYSKCGNVNDAQRVFDEMGDRNVVSWNSLITCFEQNGPAVEALDVFQMMLESRVEPDEVTLASVISACASLSAIKVGQEVHGRVVKNDKLRNDIILSNAFVDMYAKCSRIKEARFIFDSMPIRNVIAETSMISGYAMAASTKAARLMFTKMAERNVVSWNALIAGYTQNGENEEALSLFCLLKRESVCPTHYSFANILKACADLAELHLGMQAHVHVLKHGFKFQSGEEDDIFVGNSLIDMYVKCGCVEEGYLVFRKMMERDCVSWNAMIIGFAQNGYGNEALELFREMLESGEKPDHITMIGVLSACGHAGFVEEGRHYFSSMTRDFGVAPLRDHYTCMVDLLGRAGFLEEAKSMIEEMPMQPDSVIWGSLLAACKVHRNITLGKYVAEKLLEVEPSNSGPYVLLSNMYAELGKWEDVMNVRKSMRKEGVTKQPGCSWIKIQGHDHVFMVKDKSHPRKKQIHSLLDILIAEMRPEQDHTEIGSLSSEEMDYSSNLLWDNAM

>At2G14760

MEAMGEWSTGLGGIYTEEADFMNQLLASYEQPCGGSSSETTATLTAYHHQGSQWNGGFCFSQESSSYSGYCAAMPRQEEDNNGMEDATINTNLYLVGEETSECDATEYSGKSLLPLETVAENHDHSMLQPENSLTTTTDEKMFNQCESSKKRTRATTTDKNKRANKARRSQKCVEMSGENENSGEEEYTEKAAGKRKTKPLKPQKTCCSDDESNGGDTFLSKEDGEDSKALNLNGKTRASRGAATDPQSLYARLKQLNKVHCMMVQKRRERINERLRILQHLVPNGTKVDISTMLEEAVQYVKFLQLQIKLLSSDDLWMYAPIAYNGMDIGLDLKLNALTR

>At2G20540

MAFHGIREVENYFIPFLQRVKSRNEWKKINASIIIHGLSQSSFMVTKMVDFCDKIEDMDYATRLFNQVSNPNVFLYNSIIRAYTHNSLYCDVIRIYKQLLRKSFELPDRFTFPFMFKSCASLGSCYLGKQVHGHLCKFGPRFHVVTENALIDMYMKFDDLVDAHKVFDEMYERDVISWNSLLSGYARLGQMKKAKGLFHLMLDKTIVSWTAMISGYTGIGCYVEAMDFFREMQLAGIEPDEISLISVLPSCAQLGSLELGKWIHLYAERRGFLKQTGVCNALIEMYSKCGVISQAIQLFGQMEGKDVISWSTMISGYAYHGNAHGAIETFNEMQRAKVKPNGITFLGLLSACSHVGMWQEGLRYFDMMRQDYQIEPKIEHYGCLIDVLARAGKLERAVEITKTMPMKPDSKIWGSLLSSCRTPGNLDVALVAMDHLVELEPEDMGNYVLLANIYADLGKWEDVSRLRKMIRNENMKKTPGGSLIEVNNIVQEFVSGDNSKPFWTEISIVLQLFTSHQDQDVITNNNALAFIGIV

>At2G22070

MDAPVPLSLSTLLELCTNLLQKSVNKSNGRFTAQLVHCRVIKSGLMFSVYLMNNLMNVYSKTGYALHARKLFDEMPLRTAFSWNTVLSAYSKRGDMDSTCEFFDQLPQRDSVSWTTMIVGYKNIGQYHKAIRVMGDMVKEGIEPTQFTLTNVLASVAATRCMETGKKVHSFIVKLGLRGNVSVSNSLLNMYAKCGDPMMAKFVFDRMVVRDISSWNAMIALHMQVGQMDLAMAQFEQMAERDIVTWNSMISGFNQRGYDLRALDIFSKMLRDSLLSPDRFTLASVLSACANLEKLCIGKQIHSHIVTTGFDISGIVLNALISMYSRCGGVETARRLIEQRGTKDLKIEGFTALLDGYIKLGDMNQAKNIFVSLKDRDVVAWTAMIVGYEQHGSYGEAINLFRSMVGGGQRPNSYTLAAMLSVASSLASLSHGKQIHGSAVKSGEIYSVSVSNALITMYAKAGNITSASRAFDLIRCERDTVSWTSMIIALAQHGHAEEALELFETMLMEGLRPDHITYVGVFSACTHAGLVNQGRQYFDMMKDVDKIIPTLSHYACMVDLFGRAGLLQEAQEFIEKMPIEPDVVTWGSLLSACRVHKNIDLGKVAAERLLLLEPENSGAYSALANLYSACGKWEEAAKIRKSMKDGRVKKEQGFSWIEVKHKVHVFGVEDGTHPEKNEIYMTMKKIWDEIKKMGYVPDTASVLHDLEEEVKEQILRHHSEKLAIAFGLISTPDKTTLRIMKNLRVCNDCHTAIKFISKLVGREIIVRDTTRFHHFKDGFCSCRDYW

>At2G22410

MNISKAKLLLLPPPLTPKLNRSLYSHSQRRTRSLPHHRDKPINWNSTHSFVLHNPLLSLLEKCKLLLHLKQIQAQMIINGLILDPFASSRLIAFCALSESRYLDYSVKILKGIENPNIFSWNVTIRGFSESENPKESFLLYKQMLRHGCCESRPDHFTYPVLFKVCADLRLSSLGHMILGHVLKLRLELVSHVHNASIHMFASCGDMENARKVFDESPVRDLVSWNCLINGYKKIGEAEKAIYVYKLMESEGVKPDDVTMIGLVSSCSMLGDLNRGKEFYEYVKENGLRMTIPLVNALMDMFSKCGDIHEARRIFDNLEKRTIVSWTTMISGYARCGLLDVSRKLFDDMEEKDVVLWNAMIGGSVQAKRGQDALALFQEMQTSNTKPDEITMIHCLSACSQLGALDVGIWIHRYIEKYSLSLNVALGTSLVDMYAKCGNISEALSVFHGIQTRNSLTYTAIIGGLALHGDASTAISYFNEMIDAGIAPDEITFIGLLSACCHGGMIQTGRDYFSQMKSRFNLNPQLKHYSIMVDLLGRAGLLEEADRLMESMPMEADAAVWGALLFGCRMHGNVELGEKAAKKLLELDPSDSGIYVLLDGMYGEANMWEDAKRARRMMNERGVEKIPGCSSIEVNGIVCEFIVRDKSRPESEKIYDRLHCLGRHMRSSLSVLFSEYEITNNMMNSSLLTPSSSSSSHIQTPSTTFDHEDFLDQIFSSAPWPSVVDDAHPLPSDGFHGHDVDSRNQPIMMMPLNDGSSVHALYNGFSVAGSLPNFQIPQGSGGGLMNQQGQTQTQTQPQASASTATGGTVAAPPQSRTKIRARRGQATDPHSIAERLRRERIAERMKALQELVPNGNKTDKASMLDEIIDYVKFLQLQVKVLSMSRLGGAASVSSQISEAGGSHGNASSAMVGGSQTAGNSNDSVTMTEHQVAKLMEEDMGSAMQYLQGKGLCLMPISLATAISTATCHSRNPLIPGAVADVGGPSPPNLSGMTIQSTSTKMGSGNGKLNGNGVTERSSSIAVKEAVSVSKA

>At2G27230

MGVLLREALRSMCVNNQWSYAVFWKIGCQNSSLLIWEECYNETESSSNPRRLCGLGVDTQGNEKVQLLTNRMMLNNRIILVGEGLVGRAAFTGHHQWILANSFNRDVHPPEVINEMLLQFSAGIQTVAVFPVVPHGVVQLGSSLPIMENLGFVNDVKGLILQLGCVPGALLSENYRTYEPAADFIGVPVSRIIPSQGHKILQSSAFVAETSKQHFNSTGSSDHQMVEESPCNLVDEHEGGWQSTTGFLTAGEVAVPSNPDAWLNQNFSCMSNVDAAEQQQIPCEDISSKRSLGSDDLFDMLGLDDKNKGCDNSWGVSQMRTEVLTRELSDFRIIQEMDPEFGSSGYELSGTDHLLDAVVSGACSSTKQISDETSESCKTTLTKVSNSSVTTPSHSSPQGSQLFEKKHGQPLGPSSVYGSQISSWVEQAHSLKREGSPRMVNKNETAKPANNRKRLKPGENPRPRPKDRQMIQDRVKELREIIPNGAKCSIDALLERTIKHMLFLQNVSKHSDKLKQTGESKIMKEDGGGATWAFEVGSKSMVCPIVVEDINPPRIFQVEMLCEQRGFFLEIADWIRSLGLTILKGVIETRVDKIWARFTVEASRDVTRMEIFMQLVNILEQTMKCGGNSKTILDGIKATMPLPVTGGCSM

>At2G29760

MAIFSTAQPLSLPRHPNFSNPNQPTTNNERSRHISLIERCVSLRQLKQTHGHMIRTGTFSDPYSASKLFAMAALSSFASLEYARKVFDEIPKPNSFAWNTLIRAYASGPDPVLSIWAFLDMVSESQCYPNKYTFPFLIKAAAEVSSLSLGQSLHGMAVKSAVGSDVFVANSLIHCYFSCGDLDSACKVFTTIKEKDVVSWNSMINGFVQKGSPDKALELFKKMESEDVKASHVTMVGVLSACAKIRNLEFGRQVCSYIEENRVNVNLTLANAMLDMYTKCGSIEDAKRLFDAMEEKDNVTWTTMLDGYAISEDYEAAREVLNSMPQKDIVAWNALISAYEQNGKPNEALIVFHELQLQKNMKLNQITLVSTLSACAQVGALELGRWIHSYIKKHGIRMNFHVTSALIHMYSKCGDLEKSREVFNSVEKRDVFVWSAMIGGLAMHGCGNEAVDMFYKMQEANVKPNGVTFTNVFCACSHTGLVDEAESLFHQMESNYGIVPEEKHYACIVDVLGRSGYLEKAVKFIEAMPIPPSTSVWGALLGACKIHANLNLAEMACTRLLELEPRNDGAHVLLSNIYAKLGKWENVSELRKHMRVTGLKKEPGCSSIEIDGMIHEFLSGDNAHPMSEKVYGKLHEVMEKLKSNGYEPEISQVLQIIEEEEMKEQSLNLHSEKLAICYGLISTEAPKVIRVIKNLRVCGDCHSVAKLISQLYDREIIVRDRYRFHHFRNGQCSCNDFW

>At2G31280

MGSTSQEILKSFCFNTDWDYAVFWQLNHRGSRMVLTLEDAYYDHHGTNMHGAHDPLGLAVAKMSYHVYSLGEGIVGQVAVSGEHQWVFPENYNNCNSAFEFHNVWESQISAGIKTILVVAVGPCGVVQLGSLCKVNEDVNFVNHIRHLFLALRDPLADHAANLRQCNMNNSLCLPKMPSEGLHAEAFPDCSGEVDKAMDVEESNILTQYKTRRSDSMPYNTPSSCLVMEKAAQVVGGREVVQGSTCGSYSGVTFGFPVDLVGAKHENQVGTNIIRDAPHVGMTSGCKDSRDLDPNLHLYMKNHVLNDTSTSALAIEAERLITSQSYPRLDSTFQATSRTDKESSYHNEVFQLSENQGNKYIKETERMLGRNCESSQFDALISSGYTFAGSELLEALGSAFKQTNTGQEELLKSEHGSTMRPTDDMSHSQLTFDPGPENLLDAVVANVCQRDGNARDDMMSSRSVQSLLTNMELAEPSGQKKHNIVNPINSAMNQPPMAEVDTQQNSSDICGAFSSIGFSSTYPSSSSDQFQTSLDIPKKNKKRAKPGESSRPRPRDRQLIQDRIKELRELVPNGSKCSIDSLLERTIKHMLFLQNVTKHAEKLSKSANEKMQQKETGMQGSSCAVEVGGHLQVSSIIVENLNKQGMVLIEMLCEECGHFLEIANVIRSLDLVILRGFTETQGEKTWICFVTEVGSRITQFMKEIPKQIKSQNSKVMQRMDILWSLVQIFQPKANEKG

>At2G42920

MSPTILSFSGVTVPAMPSSGSLSGNTYLRLIDTQCSTMRELKQIHASLIKTGLISDTVTASRVLAFCCASPSDMNYAYLVFTRINHKNPFVWNTIIRGFSRSSFPEMAISIFIDMLCSSPSVKPQRLTYPSVFKAYGRLGQARDGRQLHGMVIKEGLEDDSFIRNTMLHMYVTCGCLIEAWRIFLGMIGFDVVAWNSMIMGFAKCGLIDQAQNLFDEMPQRNGVSWNSMISGFVRNGRFKDALDMFREMQEKDVKPDGFTMVSLLNACAYLGASEQGRWIHEYIVRNRFELNSIVVTALIDMYCKCGCIEEGLNVFECAPKKQLSCWNSMILGLANNGFEERAMDLFSELERSGLEPDSVSFIGVLTACAHSGEVHRADEFFRLMKEKYMIEPSIKHYTLMVNVLGGAGLLEEAEALIKNMPVEEDTVIWSSLLSACRKIGNVEMAKRAAKCLKKLDPDETCGYVLLSNAYASYGLFEEAVEQRLLMKERQMEKEVGCSSIEVDFEVHEFISCGGTHPKSAEIYSLLDILNWDVSTIKSGFAELFDATTRIGFTYLAEK

>At2G44880

MLRHAIETNVQIFTKFLVISASAVGIGYARKLFDQRPQRDDSFLSNSMIKAYLETRQYPDSFALYRDLRKETCFAPDNFTFTTLTKSCSLSMCVYQGLQLHSQIWRFGFCADMYVSTGVVDMYAKFGKMGCARNAFDEMPHRSEVSWTALISGYIRCGELDLASKLFDQMPHVKDVVIYNAMMDGFVKSGDMTSARRLFDEMTHKTVITWTTMIHGYCNIKDIDAARKLFDAMPERNLVSWNTMIGGYCQNKQPQEGIRLFQEMQATTSLDPDDVTILSVLPAISDTGALSLGEWCHCFVQRKKLDKKVKVCTAILDMYSKCGEIEKAKRIFDEMPEKQVASWNAMIHGYALNGNARAALDLFVTMMIEEKPDEITMLAVITACNHGGLVEEGRKWFHVMREMGLNAKIEHYGCMVDLLGRAGSLKEAEDLITNMPFEPNGIILSSFLSACGQYKDIERAERILKKAVELEPQNDGNYVLLRNLYAADKRWDDFGMVKNVMRKNQAKKEVGCSLIEINYIVSEFISGDTTHPHRRSIHLVLGDLLMHMNEEKYNW

>At2G46510

MNMSDLGWDDEDKSVVSAVLGHLASDFLRANSNSNQNLFLVMGTDDTLNKKLSSLVDWPNSENFSWNYAIFWQQTMSRSGQQVLGWGDGCCREPNEEEESKVVRSYNFNNMGAEEETWQDMRKRVLQKLHRLFGGSDEDNYALSLEKVTATEIFFLASMYFFFNHGEGGPGRCYSSGKHVWLSDAVNSESDYCFRSFMAKSAGIRTIVMVPTDAGVLELGSVWSLPENIGLVKSVQALFMRRVTQPVMVTSNTNMTGGIHKLFGQDLSGAHAYPKKLEVRRNLDERFTPQSWEGYNNNKGPTFGYTPQRDDVKVLENVNMVVDNNNYKTQIEFAGSSVAASSNPSTNTQQEKSESCTEKRPVSLLAGAGIVSVVDEKRPRKRGRKPANGREEPLNHVEAERQRREKLNQRFYALRSVVPNISKMDKASLLGDAISYIKELQEKVKIMEDERVGTDKSLSESNTITVEESPEVDIQAMNEEVVVRVISPLDSHPASRIIQAMRNSNVSLMEAKLSLAEDTMFHTFVIKSNNGSDPLTKEKLIAAFYPETSSTQPPLPSSSSQVSGDI

>At3G02330

MAESLRLLHMTRSVVSFNRCLTEKISYRRVPSFSYFTDFLNQVNSVSTTNFSFVFKECAKQGALELGKQAHAHMIISGFRPTTFVLNCLLQVYTNSRDFVSASMVFDKMPLRDVVSWNKMINGYSKSNDMFKANSFFNMMPVRDVVSWNSMLSGYLQNGESLKSIEVFVDMGREGIEFDGRTFAIILKVCSFLEDTSLGMQIHGIVVRVGCDTDVVAASALLDMYAKGKRFVESLRVFQGIPEKNSVSWSAIIAGCVQNNLLSLALKFFKEMQKVNAGVSQSIYASVLRSCAALSELRLGGQLHAHALKSDFAADGIVRTATLDMYAKCDNMQDAQILFDNSENLNRQSYNAMITGYSQEEHGFKALLLFHRLMSSGLGFDEISLSGVFRACALVKGLSEGLQIYGLAIKSSLSLDVCVANAAIDMYGKCQALAEAFRVFDEMRRRDAVSWNAIIAAHEQNGKGYETLFLFVSMLRSRIEPDEFTFGSILKACTGGSLGYGMEIHSSIVKSGMASNSSVGCSLIDMYSKCGMIEEAEKIHSRFFQRANVSGTMEELEKMHNKRLQEMCVSWNSIISGYVMKEQSEDAQMLFTRMMEMGITPDKFTYATVLDTCANLASAGLGKQIHAQVIKKELQSDVYICSTLVDMYSKCGDLHDSRLMFEKSLRRDFVTWNAMICGYAHHGKGEEAIQLFERMILENIKPNHVTFISILRACAHMGLIDKGLEYFYMMKRDYGLDPQLPHYSNMVDILGKSGKVKRALELIREMPFEADDVIWRTLLGVCTIHRNNVEVAEEATAALLRLDPQDSSAYTLLSNVYADAGMWEKVSDLRRNMRGFKLKKEPGCSWVELKDELHVFLVGDKAHPRWEEIYEELGLIYSEMKPFDDSSFVRGVEVEEEDQWCYC

>At3G04750

MCFVLLLRRGFRLFGTECGSKTTKWDPVQSLQLNHQSLVLLENCNSRNQFKQVLAQIMRFNLICDTFPMSRLIFFSAITYPENLDLAKLLFLNFTPNPNVFVYNTMISAVSSSKNECFGLYSSMIRHRVSPDRQTFLYLMKASSFLSEVKQIHCHIIVSGCLSLGNYLWNSLVKFYMELGNFGVAEKVFARMPHPDVSSFNVMIVGYAKQGFSLEALKLYFKMVSDGIEPDEYTVLSLLVCCGHLSDIRLGKGVHGWIERRGPVYSSNLILSNALLDMYFKCKESGLAKRAFDAMKKKDMRSWNTMVVGFVRLGDMEAAQAVFDQMPKRDLVSWNSLLFGYSKKGCDQRTVRELFYEMTIVEKVKPDRVTMVSLISGAANNGELSHGRWVHGLVIRLQLKGDAFLSSALIDMYCKCGIIERAFMVFKTATEKDVALWTSMITGLAFHGNGQQALQLFGRMQEEGVTPNNVTLLAVLTACSHSGLVEEGLHVFNHMKDKFGFDPETEHYGSLVDLLCRAGRVEEAKDIVQKKMPMRPSQSMWGSILSACRGGEDIETAELALTELLKLEPEKEGGYVLLSNIYATVGRWGYSDKTREAMENRGVKKTAGYSSVVGVEGLHRFVAAEKQNHPRWTEIKRILQHLYNEMKPKLDCLDLLEIEIK

>At3G05240

MMKKHYKPILSQLENCRSLVELNQLHGLMIKSSVIRNVIPLSRLIDFCTTCPETMNLSYARSVFESIDCPSVYIWNSMIRGYSNSPNPDKALIFYQEMLRKGYSPDYFTFPYVLKACSGLRDIQFGSCVHGFVVKTGFEVNMYVSTCLLHMYMCCGEVNYGLRVFEDIPQWNVVAWGSLISGFVNNNRFSDAIEAFREMQSNGVKANETIMVDLLVACGRCKDIVTGKWFHGFLQGLGFDPYFQSKVGFNVILATSLIDMYAKCGDLRTARYLFDGMPERTLVSWNSIITGYSQNGDAEEALCMFLDMLDLGIAPDKVTFLSVIRASMIQGCSQLGQSIHAYVSKTGFVKDAAIVCALVNMYAKTGDAESAKKAFEDLEKKDTIAWTVVIIGLASHGHGNEALSIFQRMQEKGNATPDGITYLGVLYACSHIGLVEEGQRYFAEMRDLHGLEPTVEHYGCMVDILSRAGRFEEAERLVKTMPVKPNVNIWGALLNGCDIHENLELTDRIRSMVAEPEELGSGIYVLLSNIYAKAGRWADVKLIRESMKSKRVDKVLGHSSVETMFMSIVTVPSATSKVQQIKTLISVACTVNHLKQIHVSLINHHLHHDTFLVNLLLKRTLFFRQTKYSYLLFSHTQFPNIFLYNSLINGFVNNHLFHETLDLFLSIRKHGLYLHGFTFPLVLKACTRASSRKLGIDLHSLVVKCGFNHDVAAMTSLLSIYSGSGRLNDAHKLFDEIPDRSVVTWTALFSGYTTSGRHREAIDLFKKMVEMGVKPDSYFIVQVLSACVHVGDLDSGEWIVKYMEEMEMQKNSFVRTTLVNLYAKCGKMEKARSVFDSMVEKDIVTWSTMIQGYASNSFPKEGIELFLQMLQENLKPDQFSIVGFLSSCASLGALDLGEWGISLIDRHEFLTNLFMANALIDMYAKCGAMARGFEVFKEMKEKDIVIMNAAISGLAKNGHVKLSFAVFGQTEKLGISPDGSTFLGLLCGCVHAGLIQDGLRFFNAISCVYALKRTVEHYGCMVDLWGRAGMLDDAYRLICDMPMRPNAIVWGALLSGCRLVKDTQLAETVLKELIALEPWNAGNYVQLSNIYSVGGRWDEAAEVRDMMNKKGMKKIPGYSWIELEGKVHEFLADDKSHPLSDKIYAKLEDLGNEMRLMGFVPTTEFVFFDVEEEEKERVLGYHSEKLAVALGLISTDHGQVIRVVKNLRVCGDCHEVMKLISKITRREIVVRDNNRFHCFTNGSCSCNDYW

>At3G11460

MIVVTSFVRNSAVAAVASTPWNVRLRELAYQSLFSESISLYRSMLRSGSSPDAFSFPFILKSCASLSLPVSGQQLHCHVTKGGCETEPFVLTALISMYCKCGLVADARKVFEENPQSSQLSVCYNALISGYTANSKVTDAAYMFRRMKETGVSVDSVTMLGLVPLCTVPEYLWLGRSLHGQCVKGGLDSEVAVLNSFITMYMKCGSVEAGRRLFDEMPVKGLITWNAVISGYSQNGLAYDVLELYEQMKSSGVCPDPFTLVSVLSSCAHLGAKKIGHEVGKLVESNGFVPNVFVSNASISMYARCGNLAKARAVFDIMPVKSLVSWTAMIGCYGMHGMGEIGLMLFDDMIKRGIRPDGAVFVMVLSACSHSGLTDKGLELFRAMKREYKLEPGPEHYSCLVDLLGRAGRLDEAMEFIESMPVEPDGAVWGALLGACKIHKNVDMAELAFAKVIEFEPNNIGYYVLMSNIYSDSKNQEGIWRIRVMMRERAFRKKPGYSYVEHKGRVHLFLAGDRSHEQTEEVHRMLDELETSVMELAGNMDCDRGEEVSSTTREHSERLAIAFGILNSIPGTEILVIKNLRVCEDCHVFLKQVSKIVDRQFVVRDASRFHYFKDGVCSCKDYW

>At3G12770

MSEASCLASPLLYTNSGIHSDSFYASLIDSATHKAQLKQIHARLLVLGLQFSGFLITKLIHASSSFGDITFARQVFDDLPRPQIFPWNAIIRGYSRNNHFQDALLMYSNMQLARVSPDSFTFPHLLKACSGLSHLQMGRFVHAQVFRLGFDADVFVQNGLIALYAKCRRLGSARTVFEGLPLPERTIVSWTAIVSAYAQNGEPMEALEIFSQMRKMDVKPDWVALVSVLNAFTCLQDLKQGRSIHASVVKMGLEIEPDLLISLNTMYAKCGQVATAKILFDKMKSPNLILWNAMISGYAKNGYAREAIDMFHEMINKDVRPDTISITSAISACAQVGSLEQARSMYEYVGRSDYRDDVFISSALIDMFAKCGSVEGARLVFDRTLDRDVVVWSAMIVGYGLHGRAREAISLYRAMERGGVHPNDVTFLGLLMACNHSGMVREGWWFFNRMADHKINPQQQHYACVIDLLGRAGHLDQAYEVIKCMPVQPGVTVWGALLSACKKHRHVELGEYAAQQLFSIDPSNTGHYVQLSNLYAAARLWDRVAEVRVRMKEKGLNKDVGCSWVEVRGRLEAFRVGDKSHPRYEEIERQVEWIESRLKEGGFVANKDASLHDLNDEEAEETLCSHSERIAIAYGLISTPQGTPLRITKNLRACVNCHAATKLISKLVDREIVVRDTNRFHHFKDGVCSCGDYW

>At3G14730

MTVSFLRIEIQNCSAGILRFLPRNPDLFAAIKPSSALASLYSTVSGQIEENPKRYEHHNVATCIATLQRCAQRKDYVSGQQIHGFMVRKGFLDDSPRAGTSLVNMYAKCGLMRRAVLVFGGSERDVFGYNALISGFVVNGSPLDAMETYREMRANGILPDKYTFPSLLKGSDAMELSDVKKVHGLAFKLGFDSDCYVGSGLVTSYSKFMSVEDAQKVFDELPDRDDSVLWNALVNGYSQIFRFEDALLVFSKMREEGVGVSRHTITSVLSAFTVSGDIDNGRSIHGLAVKTGSGSDIVVSNALIDMYGKSKWLEEANSIFEAMDERDLFTWNSVLCVHDYCGDHDGTLALFERMLCSGIRPDIVTLTTVLPTCGRLASLRQGREIHGYMIVSGLLNRKSSNEFIHNSLMDMYVKCGDLRDARMVFDSMRVKDSASWNIMINGYGVQSCGELALDMFSCMCRAGVKPDEITFVGLLQACSHSGFLNEGRNFLAQMETVYNILPTSDHYACVIDMLGRADKLEEAYELAISKPICDNPVVWRSILSSCRLHGNKDLALVAGKRLHELEPEHCGGYVLMSNVYVEAGKYEEVLDVRDAMRQQNVKKTPGCSWIVLKNGVHTFFTGNQTHPEFKSIHDWLSLVISHMHGHEYMTVDD

>At3G15240

MVGSEHHHHHQQHHHQNHQQQQQRSKEALGMVALHDALRTVCLNTDWTYSVFWSIRPRPRVRGGGNGCKVGDDNGSLMLMWEDGYCRGRGGTEGCYGDMEGEDPVRKSFSKMSIQLYNYGEGLMGKVASDKCHKWVFKEQTESESNASSYWQSSFDAIPSEWNDQFESGIRTIAVIQAGHGLLQLGSCKIIPEDLHFVLRMRHTFESLGYQSGFYLSQLFSSNRTTSSSNTPLMASNHPLLPTQTQPLQPTLPHYNWSGTSQRPMMPQTSLPTYQPHMTMPFPVMPHSNKEQDPDVKWPTGLSFFNALTNNVNAKLLFDSEGLGDKTDHQNQSQEQSNSESQANPSEFLSLDSHHRNMSFLEMILRNPLKSPFNSELSIFKALLMSTITESISNDYSRFISILGVCKTTDQFKQLHSQSITRGVAPNPTFQKKLFVFWCSRLGGHVSYAYKLFVKIPEPDVVVWNNMIKGWSKVDCDGEGVRLYLNMLKEGVTPDSHTFPFLLNGLKRDGGALACGKKLHCHVVKFGLGSNLYVQNALVKMYSLCGLMDMARGVFDRRCKEDVFSWNLMISGYNRMKEYEESIELLVEMERNLVSPTSVTLLLVLSACSKVKDKDLCKRVHEYVSECKTEPSLRLENALVNAYAACGEMDIAVRIFRSMKARDVISWTSIVKGYVERGNLKLARTYFDQMPVRDRISWTIMIDGYLRAGCFNESLEIFREMQSAGMIPDEFTMVSVLTACAHLGSLEIGEWIKTYIDKNKIKNDVVVGNALIDMYFKCGCSEKAQKVFHDMDQRDKFTWTAMVVGLANNGQGQEAIKVFFQMQDMSIQPDDITYLGVLSACNHSGMVDQARKFFAKMRSDHRIEPSLVHYGCMVDMLGRAGLVKEAYEILRKMPMNPNSIVWGALLGASRLHNDEPMAELAAKKILELEPDNGAVYALLCNIYAGCKRWKDLREVRRKIVDVAIKKTPGFSLIEVNGFAHEFVAGDKSHLQSEEIYMKLEELAQESTFAAYLPDTSELLFEAGDAYSVANRFVRLSGHPGTKNWKSTVR

>At3G16890

MRGFASSASRIATAAAASKSLNASTSVNPKLSKTLNSSGKPTNPLNQRYISQVIERKDWFLILNQEFTTHRIGLNTRFVISVLQNQDNPLHSLRFYLWVSNFDPVYAKDQSLKSVLGNALFRKGPLLLSMELLKEIRDSGYRISDELMCVLIGSWGRLGLAKYCNDVFAQISFLGMKPSTRLYNAVIDALVKSNSLDLAYLKFQQMRSDGCKPDRFTYNILIHGVCKKGVVDEAIRLVKQMEQEGNRPNVFTYTILIDGFLIAGRVDEALKQLEMMRVRKLNPNEATIRTFVHGIFRCLPPCKAFEVLVGFMEKDSNLQRVGYDAVLYCLSNNSMAKETGQFLRKIGERGYIPDSSTFNAAMSCLLKGHDLVETCRIFDGFVSRGVKPGFNGYLVLVQALLNAQRFSEGDRYLKQMGVDGLLSSVYSYNAVIDCLCKARRIENAAMFLTEMQDRGISPNLVTFNTFLSGYSVRGDVKKVHGVLEKLLVHGFKPDVITFSLIINCLCRAKEIKDAFDCFKEMLEWGIEPNEITYNILIRSCCSTGDTDRSVKLFAKMKENGLSPDLYAYNATIQSFCKMRKVKKAEELLKTMLRIGLKPDNFTYSTLIKALSESGRESEAREMFSSIERHGCVPDSYTKRLVEELDLRKSGLSRETVSAS

>At3G22690

MAMLGNVLHLSPMVLATTTTTKPSLLNQSKCTKATPSSLKNCKTIDELKMFHRSLTKQGLDNDVSTITKLVARSCELGTRESLSFAKEVFENSESYGTCFMYNSLIRGYASSGLCNEAILLFLRMMNSGISPDKYTFPFGLSACAKSRAKGNGIQIHGLIVKMGYAKDLFVQNSLVHFYAECGELDSARKVFDEMSERNVVSWTSMICGYARRDFAKDAVDLFFRMVRDEEVTPNSVTMVCVISACAKLEDLETGEKVYAFIRNSGIEVNDLMVSALVDMYMKCNAIDVAKRLFDEYGASNLDLCNAMASNYVRQGLTREALGVFNLMMDSGVRPDRISMLSAISSCSQLRNILWGKSCHGYVLRNGFESWDNICNALIDMYMKCHRQDTAFRIFDRMSNKTVVTWNSIVAGYVENGEVDAAWETFETMPEKNIVSWNTIISGLVQGSLFEEAIEVFCSMQSQEGVNADGVTMMSIASACGHLGALDLAKWIYYYIEKNGIQLDVRLGTTLVDMFSRCGDPESAMSIFNSLTNRDVSAWTAAIGAMAMAGNAERAIELFDDMIEQGLKPDGVAFVGALTACSHGGLVQQGKEIFYSMLKLHGVSPEDVHYGCMVDLLGRAGLLEEAVQLIEDMPMEPNDVIWNSLLAACRVQGNVEMAAYAAEKIQVLAPERTGSYVLLSNVYASAGRWNDMAKVRLSMKEKGLRKPPGTSSIQIRGKTHEFTSGDESHPEMPNIEAMLDEVSQRASHLGHVPDLSNVLMDVDEKEKIFMLSRHSEKLAMAYGLISSNKGTTIRIVKNLRVCSDCHSFAKFASKVYNREIILRDNNRFHYIRQGKCSCGDFCLTDDDLEVLKGCLDLGFGFNYDEIPALCKTLPALELCYSMSQKNLDDKHTPSLQLPIGRSLVPSCDNPDDVKARLKCWAQARRRRFQKGIGRSEFN

>At3G23330

MSSSKALIKTLIKNPTRIKSKSQAKQLHAQFIRTQSLSHTSASIVISIYTNLKLLHEALLLFKTLKSPPVLAWKSVIRCFTDQSLFSKALASFVEMRASGRCPDHNVFPSVLKSCTMMMDLRFGESVHGFIVRLGMDCDLYTGNALMNMYAKLLGMGSKISVGNVFDEMPQRTSNSGDEDVKAETCIMPFGIDSVRRVFEVMPRKDVVSYNTIIAGYAQSGMYEDALRMVREMGTTDLKPDSFTLSSVLPIFSEYVDVIKGKEIHGYVIRKGIDSDVYIGSSLVDMYAKSARIEDSERVFSRLYCRDGISWNSLVAGYVQNGRYNEALRLFRQMVTAKVKPGAVAFSSVIPACAHLATLHLGKQLHGYVLRGGFGSNIFIASALVDMYSKCGNIKAARKIFDRMNVLDEVSWTAIIMGHALHGHGHEAVSLFEEMKRQGVKPNQVAFVAVLTACSHVGLVDEAWGYFNSMTKVYGLNQELEHYAAVADLLGRAGKLEEAYNFISKMCVEPTGSVWSTLLSSCSVHKNLELAEKVAEKIFTVDSENMGAYVLMCNMYASNGRWKEMAKLRLRMRKKGLRKKPACSWIEMKNKTHGFVSGDRSHPSMDKINEFLKAVMEQMEKEGYVADTSGVLHDVDEEHKRELLFGHSERLAVAFGIINTEPGTTIRVTKNIRICTDCHVAIKFISKITEREIIVRDNSRFHHFNRGNCSCGDYW

>At3G26630

MAAPSPSSPPNLLSPPPFRSPEASYFLRTCSNFSQLKQIHTKIIKHNLTNDQLLVRQLISVSSSFGETQYASLVFNQLQSPSTFTWNLMIRSLSVNHKPREALLLFILMMISHQSQFDKFTFPFVIKACLASSSIRLGTQVHGLAIKAGFFNDVFFQNTLMDLYFKCGKPDSGRKVFDKMPGRSIVSWTTMLYGLVSNSQLDSAEIVFNQMPMRNVVSWTAMITAYVKNRRPDEAFQLFRRMQVDDVKPNEFTIVNLLQASTQLGSLSMGRWVHDYAHKNGFVLDCFLGTALIDMYSKCGSLQDARKVFDVMQGKSLATWNSMITSLGVHGCGEEALSLFEEMEEEASVEPDAITFVGVLSACANTGNVKDGLRYFTRMIQVYGISPIREHNACMIQLLEQALEVEKASNLVESMDSDPDFNSSFGNEYTDGMNETNETPSQHQIMFTKWDTGRFMTSSLPVRAPSWVSSRRIFEERLQDLPKCANLNQVKQLHAQIIRRNLHEDLHIAPKLISALSLCRQTNLAVRVFNQVQEPNVHLCNSLIRAHAQNSQPYQAFFVFSEMQRFGLFADNFTYPFLLKACSGQSWLPVVKMMHNHIEKLGLSSDIYVPNALIDCYSRCGGLGVRDAMKLFEKMSERDTVSWNSMLGGLVKAGELRDARRLFDEMPQRDLISWNTMLDGYARCREMSKAFELFEKMPERNTVSWSTMVMGYSKAGDMEMARVMFDKMPLPAKNVVTWTIIIAGYAEKGLLKEADRLVDQMVASGLKFDAAAVISILAACTESGLLSLGMRIHSILKRSNLGSNAYVLNALLDMYAKCGNLKKAFDVFNDIPKKDLVSWNTMLHGLGVHGHGKEAIELFSRMRREGIRPDKVTFIAVLCSCNHAGLIDEGIDYFYSMEKVYDLVPQVEHYGCLVDLLGRVGRLKEAIKVVQTMPMEPNVVIWGALLGACRMHNEVDIAKEVLDNLVKLDPCDPGNYSLLSNIYAAAEDWEGVADIRSKMKSMGVEKPSGASSVELEDGIHEFTVFDKSHPKSDQIYQMLGSLIEPPDPGELVAVR

>At3G46790

MFLSHPPQVIQPTYHTVNFLPRSPLKPPSCSVALNNPSISSGAGAKISNNQLIQSLCKEGKLKQAIRVLSQESSPSQQTYELLILCCGHRSSLSDALRVHRHILDNGSDQDPFLATKLIGMYSDLGSVDYARKVFDKTRKRTIYVWNALFRALTLAGHGEEVLGLYWKMNRIGVESDRFTYTYVLKACVASECTVNHLMKGKEIHAHLTRRGYSSHVYIMTTLVDMYARFGCVDYASYVFGGMPVRNVVSWSAMIACYAKNGKAFEALRTFREMMRETKDSSPNSVTMVSVLQACASLAALEQGKLIHGYILRRGLDSILPVISALVTMYGRCGKLEVGQRVFDRMHDRDVVSWNSLISSYGVHGYGKKAIQIFEEMLANGASPTPVTFVSVLGACSHEGLVEEGKRLFETMWRDHGIKPQIEHYACMVDLLGRANRLDEAAKMVQDMRTEPGPKVWGSLLGSCRIHGNVELAERASRRLFALEPKNAGNYVLLADIYAEAQMWDEVKRVKKLLEHRGLQKLPGRCWMEVRRKMYSFVSVDEFNPLMEQIHAFLVKLAEDMKEKGYIPQTKGVLYELETEEKERIVLGHSEKLALAFGLINTSKGEPIRITKNLRLCEDCHLFTKFISKFMEKEILVRDVNRFHRFKNGVCSCGDYW

>At3G49142

MKRINVDLHLLHFPKFRKFQSRKVSSSLPKLELDQKSPQETVFLLGQVLDTYPDIRTLRTVHSRIILEDLRCNSSLGVKLMRAYASLKDVASARKVFDEIPERNVIIINVMIRSYVNNGFYGEGVKVFGTMCGCNVRPDHYTFPCVLKACSCSGTIVIGRKIHGSATKVGLSSTLFVGNGLVSMYGKCGFLSEARLVLDEMSRRDVVSWNSLVVGYAQNQRFDDALEVCREMESVKISHDAGTMASLLPAVSNTTTENVMYVKDMFFKMGKKSLVSWNVMIGVYMKNAMPVEAVELYSRMEADGFEPDAVSITSVLPACGDTSALSLGKKIHGYIERKKLIPNLLLENALIDMYAKCGCLEKARDVFENMKSRDVVSWTAMISAYGFSGRGCDAVALFSKLQDSGLVPDSIAFVTTLAACSHAGLLEEGRSCFKLMTDHYKITPRLEHLACMVDLLGRAGKVKEAYRFIQDMSMEPNERVWGALLGACRVHSDTDIGLLAADKLFQLAPEQSGYYVLLSNIYAKAGRWEEVTNIRNIMKSKGLKKNPGASNVEVNRIIHTFLVGDRSHPQSDEIYRELDVLVKKMKELGYVPDSESALHDVEEEDKETHLAVHSEKLAIVFALMNTKEEEEDSNNTIRITKNLRICGDCHVAAKLISQITSREIIIRDTNRFHVFRFGVCSCGDYW

>At3G62470

MAAAPWLHLSRRRSSTQTSLRSFLICSSSFDSDEFISSQSRVIGGRGEEEVRFSGALFSRMIHSSTYHPYRQIPLPHSSVQLLDASLGCRGFSSGSSNVSDGCDEEVESECDNDEETGVSCVESSTNPEEVERVCKVIDELFALDRNMEAVLDEMKLDLSHDLIVEVLERFRHARKPAFRFFCWAAERQGFAHDSRTYNSMMSILAKTRQFETMVSVLEEMGTKGLLTMETFTIAMKAFAAAKERKKAVGIFELMKKYKFKIGVETINCLLDSLGRAKLGKEAQVLFDKLKERFTPNMMTYTVLLNGWCRVRNLIEAARIWNDMIDQGLKPDIVAHNVMLEGLLRSRKKSDAIKLFHVMKSKGPCPNVRSYTIMIRDFCKQSSMETAIEYFDDMVDSGLQPDAAVYTCLITGFGTQKKLDTVYELLKEMQEKGHPPDGKTYNALIKLMANQKMPEHATRIYNKMIQNEIEPSIHTFNMIMKSYFMARNYEMGRAVWEEMIKKGICPDDNSYTVLIRGLIGEGKSREACRYLEEMLDKGMKTPLIDYNKFAADFHRGGQPEIFEELAQRAKFSGKFAAAEIFARWAQMTRRRFKQRFMED

>At3G62890

MSKGAAIIAYANPIFHIRHLKLESFLWNIIIRAIVHNVSSPQRHSPISVYLRMRNHRVSPDFHTFPFLLPSFHNPLHLPLGQRTHAQILLFGLDKDPFVRTSLLNMYSSCGDLRSAQRVFDDSGSKDLPAWNSVVNAYAKAGLIDDARKLFDEMPERNVISWSCLINGYVMCGKYKEALDLFREMQLPKPNEAFVRPNEFTMSTVLSACGRLGALEQGKWVHAYIDKYHVEIDIVLGTALIDMYAKCGSLERAKRVFNALGSKKDVKAYSAMICCLAMYGLTDECFQLFSEMTTSDNINPNSVTFVGILGACVHRGLINEGKSYFKMMIEEFGITPSIQHYGCMVDLYGRSGLIKEAESFIASMPMEPDVLIWGSLLSGSRMLGDIKTCEGALKRLIELDPMNSGAYVLLSNVYAKTGRWMEVKCIRHEMEVKGINKVPGCSYVEVEGVVHEFVVGDESQQESERIYAMLDEIMQRLREAGYVTDTKEVLLDLNEKDKEIALSYHSEKLAIAFCLMKTRPGTPVRIIKNLRICGDCHLVMKMISKLFSREIVVRDCNRFHHFRDGSCSCRDFW

>At4G00050

MSQCVPNCHIDDTPAAATTTVRSTTAADIPILDYEVAELTWENGQLGLHGLGPPRVTASSTKYSTGAGGTLESIVDQATRLPNPKPTDELVPWFHHRSSRAAMAMDALVPCSNLVHEQQSKPGGVGSTRVGSCSDGRTMGGGKRARVAPEWSGGGSQRLTMDTYDVGFTSTSMGSHDNTIDDHDSVCHSRPQMEDEEEKKAGGKSSVSTKRSRAAAIHNQSERKRRDKINQRMKTLQKLVPNSSKTDKASMLDEVIEYLKQLQAQVSMMSRMNMPSMMLPMAMQQQQQLQMSLMSNPMGLGMGMGMPGLGLLDLNSMNRAAASAPNIHANMMPNPFLPMNCPSWDASSNDSRFQSPLIPDPMSAFLACSTQPTTMEAYSRMATLYQQMQQQLPPPSNPK

>At4G00120

MENGMYKKKGVCDSCVSSKSRSNHSPKRSMMEPQPHHLLMDWNKANDLLTQEHAAFLNDPHHLMLDPPPETLIHLDEDEEYDEDMDAMKEMQYMIAVMQPVDIDPATVPKPNRRNVRISDDPQTVVARRRRERISEKIRILKRIVPGGAKMDTASMLDEAIRYTKFLKRQVRILQPHSQIGAPMANPSYLCYYHNSQP

>At4G00480

MSLTMADGVEAAAGRSKRQNSLLRKQLALAVRSVQWSYAIFWSSSLTQPGVLEWGEGCYNGDMKKRKKSYESHYKYGLQKSKELRKLYLSMLEGDSGTTVSTTHDNLNDDDDNCHSTSMMLSPDDLSDEEWYYLVSMSYVFSPSQCLPGRASATGETIWLCNAQYAENKLFSRSLLARSASIQTVVCFPYLGGVIELGVTELISEDHNLLRNIKSCLMEISAHQDNDDEKKMEIKISEEKHQLPLGISDEDLHYKRTISTVLNYSADRSGKNDKNIRHRQPNIVTSEPGSSFLRWKQCEQQVSGFVQKKKSQNVLRKILHDVPLMHTKRMFPSQNSGLNQDDPSDRRKENEKFSVLRTMVPTVNEVDKESILNNTIKYLQELEARVEELESCMGSVNFVERQRKTTENLNDSVLIEETSGNYDDSTKIDDNSGETEQVTVFRDKTHLRVKLKETEVVIEVRCSYRDYIVADIMETLSNLHMDAFSVRSHTLNKFLTLNLKAKFRGAAVASVGMIKRELRRVIDFREPICDVPLSLHQVFRVFVCKVCQSLVGIFDNVVSSSSTKPRSILIHNSWAICIFH

>At4G01030

MYRFLGLTIHGGLIKRGLDNSDTRVVSASMGFYGRCVSLGFANKLFDEMPKRDDLAWNEIVMVNLRSGNWEKAVELFREMQFSGAKAYDSTMVKLLQVCSNKEGFAEGRQIHGYVLRLGLESNVSMCNSLIVMYSRNGKLELSRKVFNSMKDRNLSSWNSILSSYTKLGYVDDAIGLLDEMEICGLKPDIVTWNSLLSGYASKGLSKDAIAVLKRMQIAGLKPSTSSISSLLQAVAEPGHLKLGKAIHGYILRNQLWYDVYVETTLIDMYIKTGYLPYARMVFDMMDAKNIVAWNSLVSGLSYACLLKDAEALMIRMEKEGIKPDAITWNSLASGYATLGKPEKALDVIGKMKEKGVAPNVVSWTAIFSGCSKNGNFRNALKVFIKMQEEGVGPNAATMSTLLKILGCLSLLHSGKEVHGFCLRKNLICDAYVATALVDMYGKSGDLQSAIEIFWGIKNKSLASWNCMLMGYAMFGRGEEGIAAFSVMLEAGMEPDAITFTSVLSVCKNSGLVQEGWKYFDLMRSRYGIIPTIEHCSCMVDLLGRSGYLDEAWDFIQTMSLKPDATIWGAFLSSCKIHRDLELAEIAWKRLQVLEPHNSANYMMMINLYSNLNRWEDVERIRNLMRNNRVRVQDLWSWIQIDQTVHIFYAEGKTHPDEGDIYFELYKLVSEMKKSGYVPDTSCIHQDISDSEKEKLLMGHTEKLAMTYGLIKKKGLAPIRVVKNTNICSDSHTVAKYMSVLRNREIVLQEGARVHHFRDGKCSCNDSW

>At4G02750

MEINKFRALSRRAQQLHYTSLNGLKRRCNNAHGAANFHSLKRATQTQIQKSQTKPLLKCGDSDIKEWNVAISSYMRTGRCNEALRVFKRMPRWSSVSYNGMISGYLRNGEFELARKLFDEMPERDLVSWNVMIKGYVRNRNLGKARELFEIMPERDVCSWNTMLSGYAQNGCVDDARSVFDRMPEKNDVSWNALLSAYVQNSKMEEACMLFKSRENWALVSWNCLLGGFVKKKKIVEARQFFDSMNVRDVVSWNTIITGYAQSGKIDEARQLFDESPVQDVFTWTAMVSGYIQNRMVEEARELFDKMPERNEVSWNAMLAGYVQGERMEMAKELFDVMPCRNVSTWNTMITGYAQCGKISEAKNLFDKMPKRDPVSWAAMIAGYSQSGHSFEALRLFVQMEREGGRLNRSSFSSALSTCADVVALELGKQLHGRLVKGGYETGCFVGNALLLMYCKCGSIEEANDLFKEMAGKDIVSWNTMIAGYSRHGFGEVALRFFESMKREGLKPDDATMVAVLSACSHTGLVDKGRQYFYTMTQDYGVMPNSQHYACMVDLLGRAGLLEDAHNLMKNMPFEPDAAIWGTLLGASRVHGNTELAETAADKIFAMEPENSGMYVLLSNLYASSGRWGDVGKLRVRMRDKGVKKVPGYSWIEIQNKTHTFSVGDEFHPEKDEIFAFLEELDLRMKKAGYVSKTSVVLHDVEEEEKERMVRYHSERLAVAYGIMRVSSGRPIRVIKNLRVCEDCHNAIKYMARITGRLIILRDNNRFHHFKDGSCSCGDYW

>At4G08210

MVMDLKLIAAGLRHCGKVQAFKRGESIQAHVIKQGISQNVFIANNVISMYVDFRLLSDAHKVFDEMSERNIVTWTTMVSGYTSDGKPNKAIELYRRMLDSEEEAANEFMYSAVLKACGLVGDIQLGILVYERIGKENLRGDVVLMNSVVDMYVKNGRLIEANSSFKEILRPSSTSWNTLISGYCKAGLMDEAVTLFHRMPQPNVVSWNCLISGFVDKGSPRALEFLVRMQREGLVLDGFALPCGLKACSFGGLLTMGKQLHCCVVKSGLESSPFAISALIDMYSNCGSLIYAADVFHQEKLAVNSSVAVWNSMLSGFLINEENEAALWLLLQIYQSDLCFDSYTLSGALKICINYVNLRLGLQVHSLVVVSGYELDYIVGSILVDLHANVGNIQDAHKLFHRLPNKDIIAFSGLIRGCVKSGFNSLAFYLFRELIKLGLDADQFIVSNILKVCSSLASLGWGKQIHGLCIKKGYESEPVTATALVDMYVKCGEIDNGVVLFDGMLERDVVSWTGIIVGFGQNGRVEEAFRYFHKMINIGIEPNKVTFLGLLSACRHSGLLEEARSTLETMKSEYGLEPYLEHYYCVVDLLGQAGLFQEANELINKMPLEPDKTIWTSLLTACGTHKNAGLVTVIAEKLLKGFPDDPSVYTSLSNAYATLGMWDQLSKVREAAKKLGAKESGMSWII

>At4G09820

MDESSIIPAEKVAGAEKKELQGLLKTAVQSVDWTYSVFWQFCPQQRVLVWGNGYYNGAIKTRKTTQPAEVTAEEAALERSQQLRELYETLLAGESTSEARACTALSPEDLTETEWFYLMCVSFSFPPPSGMPGKAYARRKHVWLSGANEVDSKTFSRAILAKSAKIQTVVCIPMLDGVVELGTTKKVREDVEFVELTKSFFYDHCKTNPKPALSEHSTYEVHEEAEDEEEVEEEMTMSEEMRLGSPDDEDVSNQNLHSDLHIESTHTLDTHMDMMNLMEEGGNYSQTVTTLLMSHPTSLLSDSVSTSSYIQSSFATWRVENGKEHQQVKTAPSSQWVLKQMIFRVPFLHDNTKDKRLPREDLSHVVAERRRREKLNEKFITLRSMVPFVTKMDKVSILGDTIAYVNHLRKRVHELENTHHEQQHKRTRTCKRKTSEEVEVSIIENDVLLEMRCEYRDGLLLDILQVLHELGIETTAVHTSVNDHDFEAEIRAKVRGKKASIAEVKRAIHQVIIHDTNL

>At4G13650

MNKYIWLVRLWHSKEEPMFLRSVSSSFIFIHGVPRKLKTRTVFPTLCGTRRASFAAISVYISEDESFQEKRIDSVENRGIRPNHQTLKWLLEGCLKTNGSLDEGRKLHSQILKLGLDSNGCLSEKLFDFYLFKGDLYGAFKVFDEMPERTIFTWNKMIKELASRNLIGEVFGLFVRMVSENVTPNEGTFSGVLEACRGGSVAFDVVEQIHARILYQGLRDSTVVCNPLIDLYSRNGFVDLARRVFDGLRLKDHSSWVAMISGLSKNECEAEAIRLFCDMYVLGIMPTPYAFSSVLSACKKIESLEIGEQLHGLVLKLGFSSDTYVCNALVSLYFHLGNLISAEHIFSNMSQRDAVTYNTLINGLSQCGYGEKAMELFKRMHLDGLEPDSNTLASLVVACSADGTLFRGQQLHAYTTKLGFASNNKIEGALLNLYAKCADIETALDYFLETEVENVVLWNVMLVAYGLLDDLRNSFRIFRQMQIEEIVPNQYTYPSILKTCIRLGDLELGEQIHSQIIKTNFQLNAYVCSVLIDMYAKLGKLDTAWDILIRFAGKDVVSWTTMIAGYTQYNFDDKALTTFRQMLDRGIRSDEVGLTNAVSACAGLQALKEGQQIHAQACVSGFSSDLPFQNALVTLYSRCGKIEESYLAFEQTEAGDNIAWNALVSGFQQSGNNEEALRVFVRMNREGIDNNNFTFGSAVKAASETANMKQGKQVHAVITKTGYDSETEVCNALISMYAKCGSISDAEKQFLEVSTKNEVSWNAIINAYSKHGFGSEALDSFDQMIHSNVRPNHVTLVGVLSACSHIGLVDKGIAYFESMNSEYGLSPKPEHYVCVVDMLTRAGLLSRAKEFIQEMPIKPDALVWRTLLSACVVHKNMEIGEFAAHHLLELEPEDSATYVLLSNLYAVSKKWDARDLTRQKMKEKGVKKEPGQSWIEVKNSIHSFYVGDQNHPLADEIHEYFQDLTKRASEIGYVQDCFSLLNELQHEQKDPIIFIHSEKLAISFGLLSLPATVPINVMKNLRVCNDCHAWIKFVSKVSNREIIVRDAYRFHHFEGGACSCKDYW

>At4G14820

MTLPPPIASTAANTILEKLSFCKSLNHIKQLHAHILRTVINHKLNSFLFNLSVSSSSINLSYALNVFSSIPSPPESIVFNPFLRDLSRSSEPRATILFYQRIRHVGGRLDQFSFLPILKAVSKVSALFEGMELHGVAFKIATLCDPFVETGFMDMYASCGRINYARNVFDEMSHRDVVTWNTMIERYCRFGLVDEAFKLFEEMKDSNVMPDEMILCNIVSACGRTGNMRYNRAIYEFLIENDVRMDTHLLTALVTMYAGAGCMDMAREFFRKMSVRNLFVSTAMVSGYSKCGRLDDAQVIFDQTEKKDLVCWTTMISAYVESDYPQEALRVFEEMCCSGIKPDVVSMFSVISACANLGILDKAKWVHSCIHVNGLESELSINNALINMYAKCGGLDATRDVFEKMPRRNVVSWSSMINALSMHGEASDALSLFARMKQENVEPNEVTFVGVLYGCSHSGLVEEGKKIFASMTDEYNITPKLEHYGCMVDLFGRANLLREALEVIESMPVASNVVIWGSLMSACRIHGELELGKFAAKRILELEPDHDGALVLMSNIYAREQRWEDVRNIRRVMEEKNVFKEKGLSRIDQNGKSHEFLIGDKRHKQSNEIYAKLDEVVSKLKLAGYVPDCGSVLVDVEEEEKKDLVLWHSEKLALCFGLMNEEKEEEKDSCGVIRIVKNLRVCEDCHLFFKLVSKVYEREIIVRDRTRFHCYKNGLCSCRDYW

>At4G16835

MLSRFNIHQPFKRCKFRFFLRSIGNPDTILVESCSSSSCSSPEPSLVRSDYLTKPSDQDQIFPLNKIIARCVRSGDIDGALRVFHGMRAKNTITWNSLLIGISKDPSRMMEAHQLFDEIPEPDTFSYNIMLSCYVRNVNFEKAQSFFDRMPFKDAASWNTMITGYARRGEMEKARELFYSMMEKNEVSWNAMISGYIECGDLEKASHFFKVAPVRGVVAWTAMITGYMKAKKVELAEAMFKDMTVNKNLVTWNAMISGYVENSRPEDGLKLFRAMLEEGIRPNSSGLSSALLGCSELSALQLGRQIHQIVSKSTLCNDVTALTSLISMYCKCGELGDAWKLFEVMKKKDVVAWNAMISGYAQHGNADKALCLFREMIDNKIRPDWITFVAVLLACNHAGLVNIGMAYFESMVRDYKVEPQPDHYTCMVDLLGRAGKLEEALKLIRSMPFRPHAAVFGTLLGACRVHKNVELAEFAAEKLLQLNSQNAAGYVQLANIYASKNRWEDVARVRKRMKESNVVKVPGYSWIEIRNKVHHFRSSDRIHPELDSIHKKLKELEKKMKLAGYKPELEFALHNVEEEQKEKLLLWHSEKLAVAFGCIKLPQGSQIQVFKNLRICGDCHKAIKFISEIEKREIIVRDTTRFHHFKDGSCSCGDYW

>At4G18750

MAMLVTNLSSSSFCFFSSPHLQNQKEIRSGVRVRKYVIFNRASLRTVSDCVDSITTFDRSVTDANTQLRRFCESGNLENAVKLLCVSGKWDIDPRTLCSVLQLCADSKSLKDGKEVDNFIRGNGFVIDSNLGSKLSLMYTNCGDLKEASRVFDEVKIEKALFWNILMNELAKSGDFSGSIGLFKKMMSSGVEMDSYTFSCVSKSFSSLRSVHGGEQLHGFILKSGFGERNSVGNSLVAFYLKNQRVDSARKVFDEMTERDVISWNSIINGYVSNGLAEKGLSVFVQMLVSGIEIDLATIVSVFAGCADSRLISLGRAVHSIGVKACFSREDRFCNTLLDMYSKCGDLDSAKAVFREMSDRSVVSYTSMIAGYAREGLAGEAVKLFEEMEEEGISPDVYTVTAVLNCCARYRLLDEGKRVHEWIKENDLGFDIFVSNALMDMYAKCGSMQEAELVFSEMRVKDIISWNTIIGGYSKNCYANEALSLFNLLLEEKRFSPDERTVACVLPACASLSAFDKGREIHGYIMRNGYFSDRHVANSLVDMYAKCGALLLAHMLFDDIASKDLVSWTVMIAGYGMHGFGKEAIALFNQMRQAGIEADEISFVSLLYACSHSGLVDEGWRFFNIMRHECKIEPTVEHYACIVDMLARTGDLIKAYRFIENMPIPPDATIWGALLCGCRIHHDVKLAEKVAEKVFELEPENTGYYVLMANIYAEAEKWEQVKRLRKRIGQRGLRKNPGCSWIEIKGRVNIFVAGDSSNPETENIEAFLRKVRARMIEEGYSPLTKYALIDAEEMEKEEALCGHSEKLAMALGIISSGHGKIIRVTKNLRVCGDCHEMAKFMSKLTRREIVLRDSNRFHQFKDGHCSCRGFW

>At4G18840

MVEKDIYMCAEIIIRPQAYNLRLLQKENLKKMSVCSSTPVPILSFTERAKSLTEIQQAHAFMLKTGLFHDTFSASKLVAFAATNPEPKTVSYAHSILNRIGSPNGFTHNSVIRAYANSSTPEVALTVFREMLLGPVFPDKYSFTFVLKACAAFCGFEEGRQIHGLFIKSGLVTDVFVENTLVNVYGRSGYFEIARKVLDRMPVRDAVSWNSLLSAYLEKGLVDEARALFDEMEERNVESWNFMISGYAAAGLVKEAKEVFDSMPVRDVVSWNAMVTAYAHVGCYNEVLEVFNKMLDDSTEKPDGFTLVSVLSACASLGSLSQGEWVHVYIDKHGIEIEGFLATALVDMYSKCGKIDKALEVFRATSKRDVSTWNSIISDLSVHGLGKDALEIFSEMVYEGFKPNGITFIGVLSACNHVGMLDQARKLFEMMSSVYRVEPTIEHYGCMVDLLGRMGKIEEAEELVNEIPADEASILLESLLGACKRFGQLEQAERIANRLLELNLRDSSGYAQMSNLYASDGRWEKVIDGRRNMRAERVNRSLDVA

>At4G21065

MSPFSETSVLLLPMVEKCINLLQTYGVSSITKLRQIHAFSIRHGVSISDAELGKHLIFYLVSLPSPPPMSYAHKVFSKIEKPINVFIWNTLIRGYAEIGNSISAFSLYREMRVSGLVEPDTHTYPFLIKAVTTMADVRLGETIHSVVIRSGFGSLIYVQNSLLHLYANCGDVASAYKVFDKMPEKDLVAWNSVINGFAENGKPEEALALYTEMNSKGIKPDGFTIVSLLSACAKIGALTLGKRVHVYMIKVGLTRNLHSSNVLLDLYARCGRVEEAKTLFDEMVDKNSVSWTSLIVGLAVNGFGKEAIELFKYMESTEGLLPCEITFVGILYACSHCGMVKEGFEYFRRMREEYKIEPRIEHFGCMVDLLARAGQVKKAYEYIKSMPMQPNVVIWRTLLGACTVHGDSDLAEFARIQILQLEPNHSGDYVLLSNMYASEQRWSDVQKIRKQMLRDGVKKVPGHSLVEVGNRVHEFLMGDKSHPQSDAIYAKLKEMTGRLRSEGYVPQISNVYVDVEEEEKENAVVYHSEKIAIAFMLISTPERSPITVVKNLRVCADCHLAIKLVSKVYNREIVVRDRSRFHHFKNGSCSCQDYW

>At4G21300

MSISSVAKRFAPAIAPYKKSLPLRNSSRFLEETIPRRLSLLLQACSNPNLLRQGKQVHAFLIVNSISGDSYTDERILGMYAMCGSFSDCGKMFYRLDLRRSSIRPWNSIISSFVRNGLLNQALAFYFKMLCFGVSPDVSTFPCLVKACVALKNFKGIDFLSDTVSSLGMDCNEFVASSLIKAYLEYGKIDVPSKLFDRVLQKDCVIWNVMLNGYAKCGALDSVIKGFSVMRMDQISPNAVTFDCVLSVCASKLLIDLGVQLHGLVVVSGVDFEGSIKNSLLSMYSKCGRFDDASKLFRMMSRADTVTWNCMISGYVQSGLMEESLTFFYEMISSGVLPDAITFSSLLPSVSKFENLEYCKQIHCYIMRHSISLDIFLTSALIDAYFKCRGVSMAQNIFSQCNSVDVVVFTAMISGYLHNGLYIDSLEMFRWLVKVKISPNEITLVSILPVIGILLALKLGRELHGFIIKKGFDNRCNIGCAVIDMYAKCGRMNLAYEIFERLSKRDIVSWNSMITRCAQSDNPSAAIDIFRQMGVSGICYDCVSISAALSACANLPSESFGKAIHGFMIKHSLASDVYSESTLIDMYAKCGNLKAAMNVFKTMKEKNIVSWNSIIAACGNHGKLKDSLCLFHEMVEKSGIRPDQITFLEIISSCCHVGDVDEGVRFFRSMTEDYGIQPQQEHYACVVDLFGRAGRLTEAYETVKSMPFPPDAGVWGTLLGACRLHKNVELAEVASSKLMDLDPSNSGYYVLISNAHANAREWESVTKVRSLMKEREVQKIPGYSWIEINKRTHLFVSGDVNHPESSHIYSLLNSLLGELRLEGYIPQPYLPLHPESSRKVYPVSRFIEKEMRDPDKV

>At4G30980

MNSSSLLTPSSSPSPHLQSPATFDHDDFLHHIFSSTPWPSSVLDDTPPPTSDCAPVTGFHHHDADSRNQITMIPLSHNHPNDALFNGFSTGSLPFHLPQGSGGQTQTQSQATASATTGGATAQPQTKPKVRARRGQATDPHSIAERLRRERIAERMKSLQELVPNGNKTDKASMLDEIIDYVKFLQLQVKVLSMSRLGGAASASSQISEDAGGSHENTSSSGEAKMTEHQVAKLMEEDMGSAMQYLQGKGLCLMPISLATTISTATCPSRSPFVKDTGVPLSPNLSTTIVANGNGSSLVTVKDAPSVSKP

>At4G33880

MEAMGEWSNNLGGMYTYATEEADFMNQLLASYDHPGTGSSSGAAASGDHQGLYWNLGSHHNHLSLVSEAGSFCFSQESSSYSAGNSGYYTVVPPTVEENQNETMDFGMEDVTINTNSYLVGEETSECDVEKYSSGKTLMPLETVVENHDDEESLLQSEISVTTTKSLTGSKKRSRATSTDKNKRARVNKRAQKNVEMSGDNNEGEEEEGETKLKKRKNGAMMSRQNSSTTFCTEEESNCADQDGGGEDSSSKEDDPSKALNLNGKTRASRGAATDPQSLYARKRRERINERLRILQNLVPNGTKVDISTMLEEAVHYVKFLQLQIKLLSSDDLWMYAPIAFNGMDIGLSSPR

>At4G33990

MKFGTFSLPRQIPTCKGGRFTRVLQSIGSVIREFSASANALQDCWKNGNESKEIDDVHTLFRYCTNLQSAKCLHARLVVSKQIQNVCISAKLVNLYCYLGNVALARHTFDHIQNRDVYAWNLMISGYGRAGNSSEVIRCFSLFMLSSGLTPDYRTFPSVLKACRTVIDGNKIHCLALKFGFMWDVYVAASLIHLYSRYKAVGNARILFDEMPVRDMGSWNAMISGYCQSGNAKEALTLSNGLRAMDSVTVVSLLSACTEAGDFNRGVTIHSYSIKHGLESELFVSNKLIDLYAEFGRLRDCQKVFDRMYVRDLISWNSIIKAYELNEQPLRAISLFQEMRLSRIQPDCLTLISLASILSQLGDIRACRSVQGFTLRKGWFLEDITIGNAVVVMYAKLGLVDSARAVFNWLPNTDVISWNTIISGYAQNGFASEAIEMYNIMEEEGEIAANQGTWVSVLPACSQAGALRQGMKLHGRLLKNGLYLDVFVVTSLADMYGKCGRLEDALSLFYQIPRVNSVPWNTLIACHGFHGHGEKAVMLFKEMLDEGVKPDHITFVTLLSACSHSGLVDEGQWCFEMMQTDYGITPSLKHYGCMVDMYGRAGQLETALKFIKSMSLQPDASIWGALLSACRVHGNVDLGKIASEHLFEVEPEHVGYHVLLSNMYASAGKWEGVDEIRSIAHGKGLRKTPGWSSMEVDNKVEVFYTGNQTHPMYEEMYRELTALQAKLKMIGYVPDHRFVLQDVEDDEKEHILMSHSERLAIAFALIATPAKTTIRIFKNLRVCGDCHSVTKFISKITEREIIVRDSNRFHHFKNGVCSCGDYW

>At4G35130

MAATLLSQCYRIYNSDACKCVSSENHQTTGKRNGNRNLEFDSGISKPARLVLRDRYKVTKQVNDPALTRALRGFADSRLMEDALQLFDEMNKADAFLWNVMIKGFTSCGLYIEAVQFYSRMVFAGVKADTFTYPFVIKSVAGISSLEEGKKIHAMVIKLGFVSDVYVCNSLISLYMKLGCAWDAEKVFEEMPERDIVSWNSMISGYLALGDGFSSLMLFKEMLKCGFKPDRFSTMSALGACSHVYSPKMGKEIHCHAVRSRIETGDVMVMTSILDMYSKYGEVSYAERIFNGMIQRNIVAWNVMIGCYARNGRVTDAFLCFQKMSEQNGLQPDVITSINLLPASAILEGRTIHGYAMRRGFLPHMVLETALIDMYGECGQLKSAEVIFDRMAEKNVISWNSIIAAYVQNGKNYSALELFQELWDSSLVPDSTTIASILPAYAESLSLSEGREIHAYIVKSRYWSNTIILNSLVHMYAMCGDLEDARKCFNHILLKDVVSWNSIIMAYAVHGFGRISVWLFSEMIASRVNPNKSTFASLLAACSISGMVDEGWEYFESMKREYGIDPGIEHYGCMLDLIGRTGNFSAAKRFLEEMPFVPTARIWGSLLNASRNHKDITIAEFAAEQIFKMEHDNTGCYVLLLNMYAEAGRWEDVNRIKLLMESKGISRTSSRSTVEAKGKSHVFTNGDRSHVATNKIYEVLDVVSRMVGEEDIYVHCVSRLRPETLVKSRSNSPRRHSVRLATCFGLISTETGRRVTVRNNTRICRKCHEFLEKASRLTRREIVVGDSKIFHHFSNGRCSCGNYW

>At4G37380

MASSPLLATSLPQNQLSTTATARFRLPPPEKLAVLIDKSQSVDEVLQIHAAILRHNLLLHPRYPVLNLKLHRAYASHGKIRHSLALFHQTIDPDLFLFTAAINTASINGLKDQAFLLYVQLLSSEINPNEFTFSSLLKSCSTKSGKLIHTHVLKFGLGIDPYVATGLVDVYAKGGDVVSAQKVFDRMPERSLVSSTAMITCYAKQGNVEAARALFDSMCERDIVSWNVMIDGYAQHGFPNDALMLFQKLLAEGKPKPDEITVVAALSACSQIGALETGRWIHVFVKSSRIRLNVKVCTGLIDMYSKCGSLEEAVLVFNDTPRKDIVAWNAMIAGYAMHGYSQDALRLFNEMQGITGLQPTDITFIGTLQACAHAGLVNEGIRIFESMGQEYGIKPKIEHYGCLVSLLGRAGQLKRAYETIKNMNMDADSVLWSSVLGSCKLHGDFVLGKEIAEYLIGLNIKNSGIYVLLSNIYASVGDYEGVAKVRNLMKEKGIVKEPGISTIEIENKVHEFRAGDREHSKSKEIYTMLRKISERIKSHGYVPNTNTVLQDLEETEKEQSLQVHSERLAIAYGLISTKPGSPLKIFKNLRVCSDCHTVTKLISKITGRKIVMRDRNRFHHFTDGSCSCGDFW

>At5G01310

MDDFNLRSENPNSSSTTSSSSSSFHRHKSETGNTKRSRSTSTLSTDPQSVAARDRRHRISDRFKILQSMVPGGAKMDTVSMLDEAISYVKFLKAQIWYHQNMLLFINDHETTSSCTYSPGAGEFGPKLFGYDDDYAPIMDTYSQGVPLTVADSKYTPWFGSVDDEQEHVTYFKYRRATRHALRGHCNCIIGETEEFADQREKMEVQIEESGKNQTSPESIEADKAKQIVVLLIGPPGSGKSTFCDTAMRSSHRPWSRICQDIVNNGKAGTKAQCLKMATDSLREGKSVFIDRCNLDREQRSEFIKLGGPEFEVHAVVLELPAQVCISRSVKRTGHEGNLQGGRAAAVVNKMLQSKELPKVNEGFSRIMFCYSDADVDNAVNMYNKLGPMDTLPSGCFGEKKLDTKSQPGIMKFFKKVSALPASSSNEATNTTRKADEMTANVRVSPVKLGSADIVPTLAFPSISTADFQFDLEKASDIIVEKAEEFLSKLGTARLVLVDLSRGSKILSLVKAKASQKNIDSAKFFTFVGDITKLRSEGGLHCNVIANATNWRLKPGGGGVNAAIFKAAGPDLETATRVRANTLLPGKAVVVPLPSTCPLHNAEGITHVIHVLGPNMNPNRPDNLNNDYTKGCKTLREAYTSLFEGFLSVVQDQSKLPKRSSQTAVSDSGEDIKEDSERNKKYKGSQDKAVTNNLESESLEDTRGSGKKMSKGWNTWALALHSIAMHPERHENVVLEYLDNIVVINDQYPKARKHVLVLARQESLDGLEDVRKENLQLLQEMHNVGLKWVDRFQNEDASLIFRLGYHSVPSMRQLHLHVISQDFNSDSLKNKKHWNSFTTSFFRDSVDVLEEVNSQGKANVASEDLLKGELRCNRCRSAHPNIPKLKSHVRSCHSQFPDHLLQNNRLVARAET

>At5G03800

MSTVNHHCLLNFPHIPPSIPPNHRPKLLSSLSLYRKPERLFALSASLSLSPATIHECSSSSSSSSSSFDKEETEDIESVIDGFFYLLRLSAQYHDVEVTKAVHASFLKLREEKTRLGNALISTYLKLGFPREAILVFVSLSSPTVVSYTALISGFSRLNLEIEALKVFFRMRKAGLVQPNEYTFVAILTACVRVSRFSLGIQIHGLIVKSGFLNSVFVSNSLMSLYDKDSGSSCDDVLKLFDEIPQRDVASWNTVVSSLVKEGKSHKAFDLFYEMNRVEGFGVDSFTLSTLLSSCTDSSVLLRGRELHGRAIRIGLMQELSVNNALIGFYSKFWDMKKVESLYEMMMAQDAVTFTEMITAYMSFGMVDSAVEIFANVTEKNTITYNALMAGFCRNGHGLKALKLFTDMLQRGVELTDFSLTSAVDACGLVSEKKVSEQIHGFCIKFGTAFNPCIQTALLDMCTRCERMADAEEMFDQWPSNLDSSKATTSIIGGYARNGLPDKAVSLFHRTLCEQKLFLDEVSLTLILAVCGTLGFREMGYQIHCYALKAGYFSDISLGNSLISMYAKCCDSDDAIKIFNTMREHDVISWNSLISCYILQRNGDEALALWSRMNEKEIKPDIITLTLVISAFRYTESNKLSSCRDLFLSMKTIYDIEPTTEHYTAFVRVLGHWGLLEEAEDTINSMPVQPEVSVLRALLDSCRIHSNTSVAKRVAKLILSTKPETPSEYILKSNIYSASGFWHRSEMIREEMRERGYRKHPAKSWIIHENKIHSFHARDTSHPQEKDIYRGLEILIMECLKVGYEPNTEYVLQEVDEFMKKSFLFHHSAKLAVTYGILSSNTRGKPVRVMKNVMLCGDCHEFFKYISVVVKREIVLRDSSGFHHFVNGKCSCRDLW

>At5G06540

MSNIVLNTLRFKHPKLALLQSCSSFSDLKIIHGFLLRTHLISDVFVASRLLALCVDDSTFNKPTNLLGYAYGIFSQIQNPNLFVFNLLIRCFSTGAEPSKAFGFYTQMLKSRIWPDNITFPFLIKASSEMECVLVGEQTHSQIVRFGFQNDVYVENSLVHMYANCGFIAAAGRIFGQMGFRDVVSWTSMVAGYCKCGMVENAREMFDEMPHRNLFTWSIMINGYAKNNCFEKAIDLFEFMKREGVVANETVMVSVISSCAHLGALEFGERAYEYVVKSHMTVNLILGTALVDMFWRCGDIEKAIHVFEGLPETDSLSWSSIIKGLAVHGHAHKAMHYFSQMISLGFIPRDVTFTAVLSACSHGGLVEKGLEIYENMKKDHGIEPRLEHYGCIVDMLGRAGKLAEAENFILKMHVKPNAPILGALLGACKIYKNTEVAERVGNMLIKVKPEHSGYYVLLSNIYACAGQWDKIESLRDMMKEKLVKKPPGWSLIEIDGKINKFTMGDDQKHPEMGKIRRKWEEILGKIRLIGYKGNTGDAFFDVDEEEKESSIHMHSEKLAIAYGMMKTKPGTTIRIVKNLRVCEDCHTVTKLISEVYGRELIVRDRNRFHHFRNGVCSCRDYW

>At5G08305

MLKSSSSLVAKSILRHQCKSMSELYKIHTLLITLGLSEEEPFVSQTLSFSALSSSGDVDYAYKFLSKLSDPPNYGWNFVIRGFSNSRNPEKSISVYIQMLRFGLLPDHMTYPFLMKSSSRLSNRKLGGSLHCSVVKSGLEWDLFICNTLIHMYGSFRDQASARKLFDEMPHKNLVTWNSILDAYAKSGDVVSARLVFDEMSERDVVTWSSMIDGYVKRGEYNKALEIFDQMMRMGSSKANEVTMVSVICACAHLGALNRGKTVHRYILDVHLPLTVILQTSLIDMYAKCGSIGDAWSVFYRASVKETDALMWNAIIGGLASHGFIRESLQLFHKMRESKIDPDEITFLCLLAACSHGGLVKEAWHFFKSLKESGAEPKSEHYACMVDVLSRAGLVKDAHDFISEMPIKPTGSMLGALLNGCINHGNLELAETVGKKLIELQPHNDGRYVGLANVYAINKQFRAARSMREAMEKKGVKKIAGHSILDLDGTRHRFIAHDKTHFHSDKIYAVLQLTGAWMNLDVDYDDQDNHCFCS

>At5G15300

MIRRQTNDRTTNRRRPKLWQNCKNIRTLKQIHASMVVNGLMSNLSVVGELIYSASLSVPGALKYAHKLFDEIPKPDVSICNHVLRGSAQSMKPEKTVSLYTEMEKRGVSPDRYTFTFVLKACSKLEWRSNGFAFHGKVVRHGFVLNEYVKNALILFHANCGDLGIASELFDDSAKAHKVAWSSMTSGYAKRGKIDEAMRLFDEMPYKDQVAWNVMITGCLKCKEMDSARELFDRFTEKDVVTWNAMISGYVNCGYPKEALGIFKEMRDAGEHPDVVTILSLLSACAVLGDLETGKRLHIYILETASVSSSIYVGTPIWNALIDMYAKCGSIDRAIEVFRGVKDRDLSTWNTLIVGLALHHAEGSIEMFEEMQRLKVWPNEVTFIGVILACSHSGRVDEGRKYFSLMRDMYNIEPNIKHYGCMVDMLGRAGQLEEAFMFVESMKIEPNAIVWRTLLGACKIYGNVELGKYANEKLLSMRKDESGDYVLLSNIYASTGQWDGVQKVRKMFDDTRVKKPTGVSLIEEDDDKLMMRYLLSSEPESRSRGRIN

>At5G24352

MTLQAPFLGQGIVGEVASTGNHQWLFSDTLFQNQFLSGFKTIAIIPLGSSGVVQLGSTQKILESTKILEQTTRALQETC

>At5G37570

MIQRLSHPSLLSLETLFKLCKSEIHLNQIHARIIRKGLEQDQNLISIFISSSSSSSSSLSYSSSVFERVPSPGTYLWNHLIKGYSNKFLFFETVSILMRMMRTGLARPDEYTFPLVMKVCSNNGQVRVGSSVHGLVLRIGFDKDVVVGTSFVDFYGKCKDLFSARKVFGEMPERNAVSWTALVVAYVKSGELEEAKSMFDLMPERNLGSWNALVDGLVKSGDLVNAKKLFDEMPKRDIISYTSMIDGYAKGGDMVSARDLFEEARGVDVRAWSALILGYAQNGQPNEAFKVFSEMCAKNVKPDEFIMVGLMSACSQMGCFELCEKVDSYLHQRMNKFSSHYVVPALIDMNAKCGHMDRAAKLFEEMPQRDLVSYCSMMEGMAIHGCGSEAIRLFEKMVDEGIVPDEVAFTVILKVCGQSRLVEEGLRYFELMRKKYSILASPDHYSCIVNLLSRTGKLKEAYELIKSMPFEAHASAWGSLLGGCSLHGNTEIAEVVARHLFELEPQSAGSYVLLSNIYAALDRWTDVAHLRDKMNENGITKICGRSWISR

>At5G37800

MSLINEHCNERNYISTPNSSEDLSSPQNCGLDEGASASSSSTINSDHQNNQGFVFYPSGETIEDHNSLMDFNASSFFTFDNHRSLISPVTNGGAFPVVDGNMSYSYDGWSHHQVDSISPRVIKTPNSFETTSSFGLTSNSMSKPATNHGNGDWLYSGSTIVNIGSRHESTSPKLAGNKRPFTGENTQLSKKPSSGTNGKIKPKATTSPKDPQSLAAKNRRERISERLKVLQELVPNGTKVDLVTMLEKAIGYVKFLQVQVKVLAADEFWPAQGGKAPDISQVKEAIDAILSSSQRDSNSTRETSIAE

>At5G40405

MSRIGKHPAIALLDSGITFKEVRQIHAKLYVDGTLKDDHLVGHFVKAVALSDHKYLDYANQILDRSEKPTLFALNSMIRAHCKSPVPEKSFDFYRRILSSGNDLKPDNYTVNFLVQACTGLRMRETGLQVHGMTIRRGFDNDPHVQTGLISLYAELGCLDSCHKVFNSIPCPDFVCRTAMVTACARCGDVVFARKLFEGMPERDPIAWNAMISGYAQVGESREALNVFHLMQLEGVKVNGVAMISVLSACTQLGALDQGRWAHSYIERNKIKITVRLATTLVDLYAKCGDMEKAMEVFWGMEEKNVYTWSSALNGLAMNGFGEKCLELFSLMKQDGVTPNAVTFVSVLRGCSVVGFVDEGQRHFDSMRNEFGIEPQLEHYGCLVDLYARAGRLEDAVSIIQQMPMKPHAAVWSSLLHASRMYKNLELGVLASKKMLELETANHGAYVLLSNIYADSNDWDNVSHVRQSMKSKGVRKQPGCSVMEVNGEVHEFFVGDKSHPKYTQIDAVWKDISRRLRLAGYKADTTPVMFDIDEEEKEDALCLHSEKAAIAFGIMSLKEDVPIRIVKNLRVCGDCHQVSMMISKIFNREIIVRDRNRFHHFKDGHCSCNGFW

>At5G41315

MATGQNRTTVPENLKKHLAVSVRNIQWSYGIFWSVSASQSGVLEWGDGYYNGDIKTRKTIQASEIKADQLGLRRSEQLSELYESLSVAESSSSGVAAGSQVTRRASAAALSPEDLADTEWYYLVCMSFVFNIGEGMPGRTFANGEPIWLCNAHTADSKVFSRSLLAKSAAVKTVVCFPFLGGVVEIGTTEHITEDMNVIQCVKTSFLEAPDPYATILPARSDYHIDNVLDPQQILGDEIYAPMFSTEPFPTASPSRTTNGFDQEHEQVADDHDSFMTERITGGASQVQSWQLMDDELSNCVHQSLNSSDCVSQTFVEGAAGRVAYGARKSRVQRLGQIQEQQRNVKTLSFDPRNDDVHYQSVISTIFKTNHQLILGPQFRNCDKQSSFTRWKKSSSSSSGTATVTAPSQGMLKKIIFDVPRVHQKEKLMLDSPEARDETGNHAVLEKKRREKLNERFMTLRKIIPSINKIDKVSILDDTIEYLQELERRVQELESCRESTDTETRGTMTMKRKKPCDAGERTSANCANNETGNGKKVSVNNVGEAEPADTGFTGLTDNLRIGSFGNEVVIELRCAWREGVLLEIMDVISDLHLDSHSVQSSTGDGLLCLTVNCKHKGSKIATPGMIKEALQRVAWIC

>At5G43175

MENEAFVDGELESLLGMFNFDQCSSNESSFCNAPNETDVFSSDDFFPFGTILQSNYAAVLDGSNHQTNRNVDSRQDLLKPRKKQKLSSESNLVTEPKTAWRDGQSLSSYNSSDDEKALGLVSNTSKSLKRKAKANRGIASDPQSLYARKRRERINDRLKTLQSLVPNGTKVDISTMLEDAVHYVKFLQLQIKLLSSEDLWMYAPLAHNGLNMGLHHNLLSRLI

>At5G43790

MTSPSTSKNHRCLNLISKCKSLQNLKQIHAQIITIGLSHHTYPLSKLLHLSSTVCLSYALSILRQIPNPSVFLYNTLISSIVSNHNSTQTHLAFSLYDQILSSRSNFVRPNEFTYPSLFKASGFDAQWHRHGRALHAHVLKFLEPVNHDRFVQAALVGFYANCGKLREARSLFERIREPDLATWNTLLAAYANSEEIDSDEEVLLLFMRMQVRPNELSLVALIKSCANLGEFVRGVWAHVYVLKNNLTLNQFVGTSLIDLYSKCGCLSFARKVFDEMSQRDVSCYNAMIRGLAVHGFGQEGIELYKSLISQGLVPDSATFVVTISACSHSGLVDEGLQIFNSMKAVYGIEPKVEHYGCLVDLLGRSGRLEEAEECIKKMPVKPNATLWRSFLGSSQTHGDFERGEIALKHLLGLEFENSGNYVLLSNIYAGVNRWTDVEKTRELMKDHRVNKSPGISTLN

>At5G44230

MTVAHSHRFSTAVNPINISLLSKQLLQLGRTSNNSGTFSEISNQKELLVSSLISKLDDCINLNQIKQIHGHVLRKGLDQSCYILTKLIRTLTKLGVPMDPYARRVIEPVQFRNPFLWTAVIRGYAIEGKFDEAIAMYGCMRKEEITPVSFTFSALLKACGTMKDLNLGRQFHAQTFRLRGFCFVYVGNTMIDMYVKCESIDCARKVFDEMPERDVISWTELIAAYARVGNMECAAELFESLPTKDMVAWTAMVTGFAQNAKPQEALEYFDRMEKSGIRADEVTVAGYISACAQLGASKYADRAVQIAQKSGYSPSDHVVIGSALIDMYSKCGNVEEAVNVFMSMNNKNVFTYSSMILGLATHGRAQEALHLFHYMVTQTEIKPNTVTFVGALMACSHSGLVDQGRQVFDSMYQTFGVQPTRDHYTCMVDLLGRTGRLQEALELIKTMSVEPHGGVWGALLGACRIHNNPEIAEIAAEHLFELEPDIIGNYILLSNVYASAGDWGGVLRVRKLIKEKGLKKTPAVSWVVDKNGQMHKFFPGNLNHPMSNKIQDKLEELVERLTVLGYQPDLSSVPYDVSDNAKRLILIQHTEKLALAFSLLTTNRDSTITIMKNLRMCLDCHKFMRLASEVTGKVIIMRDNMRFHHFRSGDCSCGDFW

>At5G46760

MNGTTSSINFLTSDDDASAAAMEAFIGTNHHSSLFPPPPQQPPQPQFNEDTLQQRLQALIESAGENWTYAIFWQISHDFDSSTGDNTVILGWGDGYYKGEEDKEKKKNNTNTAEQEHRKRVIRELNSLISGGIGVSDESNDEEVTDTEWFFLVSMTQSFVNGVGLPGESFLNSRVIWLSGSGALTGSGCERAGQGQIYGLKTMVCIATQNGVVELGSSEVISQSSDLMHKVNNLFNFNNGGGNNGVEASSWGFNLNPDQGENDPALWISEPTNTGIESPARVNNGNNSNSNSKSDSHQISKLEKNDISSVENQNRQSSCLVEKDLTFQGGLLKSNETLSFCGNESSKKRTSVSKGSNNDEGMLSFSTVVRSAANDSDHSDLEASVVKEAIVVEPPEKKPRKRGRKPANGREEPLNHVEAERQRREKLNQRFYSLRAVVPNVSKMDKASLLGDAISYINELKSKLQQAESDKEEIQKKLDGMSKEGNNGKGCGSRAKERKSSNQDSTASSIEMEIDVKIIGWDVMIRVQCGKKDHPGARFMEALKELDLEVNHASLSVVNDLMIQQATVKMGSQFFNHDQLKVALMTKVGENY

>At5G53900

MVGSGVGGGDRSKDAVGMMALHEALRSVCFNSDWIYSVFWTIRPRPRVRGGNGCKIGDESGSLMLMWEDGFCGGGRSEDLCLETDIEGHEEDLVRKAFSKMSIQLYNYGEGLMGKVASDKCHKWVFKEPSESEPNLANYWQSSFDALPPEWTDQFESGIQTIAVIQAGHGLLQLGSCKIIPEDLHFVLRMRQMFESIGYRSGFYLSQLFSSNRTATPSSSTVPNQIPQSQGFNWGSHSPLLPSPSFQNQLPASARFGFLQDNNVPPQMLPPMEEHEDDIKWPNGLSLFNALTGRADEASRLLFNQEQNPMNVENQNEFLNLEGHHPNKFRRSYTLPARMDSSSSSTSLDQQQPLEFRNNNSGSNSGLFPDVMETFLR

>At5G56310

MIQRINALSLSSGLNWFVTSLKIHGNNLKTLKQSHCYMIITGLNRDNLNVAKFIEACSNAGHLRYAYSVFTHQPCPNTYLHNTMIRALSLLDEPNAHSIAITVYRKLWALCAKPDTFTFPFVLKIAVRVSDVWFGRQIHGQVVVFGFDSSVHVVTGLIQMYFSCGGLGDARKMFDEMLVKDVNVWNALLAGYGKVGEMDEARSLLEMMPCWVRNEVSWTCVISGYAKSGRASEAIEVFQRMLMENVEPDEVTLLAVLSACADLGSLELGERICSYVDHRGMNRAVSLNNAVIDMYAKSGNITKALDVFECVNERNVVTWTTIIAGLATHGHGAEALAMFNRMVKAGVRPNDVTFIAILSACSHVGWVDLGKRLFNSMRSKYGIHPNIEHYGCMIDLLGRAGKLREADEVIKSMPFKANAAIWGSLLAASNVHHDLELGERALSELIKLEPNNSGNYMLLANLYSNLGRWDESRMMRNMMKGIGVKKMAGESSIEVENRVYKFISGDLTHPQVERIHEILQEMDLQIQSKV

>At5G58010

MENGNGEGKGEFINQNNDFFLDSMSMLSSLPPCWDPSLPPPPPPPQSLFHALAVDAPFPDQFHHPQESGGPTMGSQEGLQPQGTVSTTSAPVVRQKPRVRARRGQATDPHSIAERLRRERIAERMKSLQELVPNTNKTDKASMLDEIIEYVRFLQLQVKVLSMSRLGGAGSVGPRLNGLSAEAGGRLNALTAPCNGLNGNGNATGSSNESLRSTEQRVAKLMEEDMGSAMQYLQGKGLCLMPISLATAISSSTTHSRGSLFNPISSAVAAEDSNVTATAVAAPEASSTMDDVSASKA

>At5G59200

MISSLAAITGGPSTFRRDPDSNTLRLSRRKTLISVLRSCKNIAHVPSIHAKIIRTFHDQDAFVVFELIRVCSTLDSVDYAYDVFSYVSNPNVYLYTAMIDGFVSSGRSADGVSLYHRMIHNSVLPDNYVITSVLKACDLKVCREIHAQVLKLGFGSSRSVGLKMMEIYGKSGELVNAKKMFDEMPDRDHVAATVMINCYSECGFIKEALELFQDVKIKDTVCWTAMIDGLVRNKEMNKALELFREMQMENVSANEFTAVCVLSACSDLGALELGRWVHSFVENQRMELSNFVGNALINMYSRCGDINEARRVFRVMRDKDVISYNTMISGLAMHGASVEAINEFRDMVNRGFRPNQVTLVALLNACSHGGLLDIGLEVFNSMKRVFNVEPQIEHYGCIVDLLGRVGRLEEAYRFIENIPIEPDHIMLGTLLSACKIHGNMELGEKIAKRLFESENPDSGTYVLLSNLYASSGKWKESTEIRESMRDSGIEKEPGCSTIEVDNQIHEFLVGDIAHPHKEAIYQRLQELNRILRFKENQIDIIMGF

>At5G66520

MNVISCSFSLEHNLYETMSCLQRCSKQEELKQIHARMLKTGLMQDSYAITKFLSFCISSTSSDFLPYAQIVFDGFDRPDTFLWNLMIRGFSCSDEPERSLLLYQRMLCSSAPHNAYTFPSLLKACSNLSAFEETTQIHAQITKLGYENDVYAVNSLINSYAVTGNFKLAHLLFDRIPEPDDVSWNSVIKGYVKAGKMDIALTLFRKMAEKNAISWTTMISGYVQADMNKEALQLFHEMQNSDVEPDNVSLANALSACAQLGALEQGKWIHSYLNKTRIRMDSVLGCVLIDMYAKCGEMEEALEVFKNIKKKSVQAWTALISGYAYHGHGREAISKFMEMQKMGIKPNVITFTAVLTACSYTGLVEEGKLIFYSMERDYNLKPTIEHYGCIVDLLGRAGLLDEAKRFIQEMPLKPNAVIWGALLKACRIHKNIELGEEIGEILIAIDPYHGGRYVHKANIHAMDKKWDKAAETRRLMKEQGVAKVPGCSTISLEGTTHEFLAGDRSHPEIEKIQSKWRIMRRKLEENGYVPELEEMLLDLVDDDEREAIVHQHSEKLAITYGLIKTKPGTIIRIMKNLRVCKDCHKVTKLISKIYKRDIVMRDRTRFHHFRDGKCSCGDYW

>Atr00029G01160

MVAGWSHQCLFLIKRCTTIKQVHQIHSLMITTGLSHCNFAMSKIIHFCAVSDPKNLEYALSLFNQVTNPTNFIWNTMIRGFSISQNPQKAILIFTKMLQKSLSPDKHTFPFVLRACVNSKQGNVIYTHVLKNGLVHDTFVCNSLIAMYSKCDALDCAYRVFDETPQRDVVTWTALIDGYVRANRATMGLDLFAKMRLVGIEPDEITMVSVLCAIGLVGALRLGRCVHAHFIEPKKVIYDSILGCALLDMYAKCQDIDSARKIIDRYMETHMAPTVDSNENYSSGNALFGNTELVGMHPLCQIASN

>Atr00038G00550

MFKISLSHDSFAVNQFVSACSSTGSMDYAISVFAHLQKANIFVWNSMIKGFVHWHSYQKAISMYKKLLVSPVSATSYTFSSVINAGTQVLGLSVGESIHGQALKWVLNSQVFVGTALIDLYSNRREI

>Atr00058G00590

MRPPLHSPFALSPNKLIEEALFSLLDHCSTHNHIREAHARIFVLGLHQNNFLAAKILGACSSVSAINHAILAFKHASNPTICLYNTLIRALAQNNLPFETIDLYTAMRRNSLPPDNFTYPFVAKACAALSALSLGQAIHAQALTYGLLRDPYISNSILDMYWKYAFSIGRVMEYYDLLLFSTRQASPSLGPL

>Atr00062G01890

MFKISLSHKSFAVNQYVSACSSTGSMDYAISVFAHLQKPNIFVWNSMIKGFVHCHSYQEAISMYKKLLVSSVSATSYTFSSVIKACTQVLGLSLGESIHGQALKLGLNSHVFVGTALIDLYSNCSEVRNARKVFDAMEARDAVTFTTMISX

>Atr00081G00180

MGAPSCLLSPPLSSPQQLPLSISKLRHTNSIKDHPTLFLLQSCTSMKQLKQIHAQFFKTGLHNDQFALSKIIEFCSISPFGDLDFAHLLFGTISEPNQFIWNTLIRGHSLSSTPMDSLSLYIAMLLSLTPPNNYTFPFLLKSCACELAVEEGRQIHGQIIKFDLELDPFIHTSLINLYAKFGDLGCACSVFYSSPRSDPVSWTALIDGFSSNGHLNEARKLFDQCPVKDVVSWNAIISGYTQSGAHDEALNLFEKMVGDGFRPNVSTMASVLSACASSGSIEIGRWVHSWIQENRLGSNPNLVNSLIDMYSKCGSIKIAHHLFDEMPERDRVSWNVMIGGCAHAGFYKEALSLFRRMQLFEEPNEVTFLTILPVCAQLGALDLGKWIHAYIDKDMKNPIFAKNAEG

>Atr00149G00160

MSNFSLLQHIERCTHMNQLRLIHAKMIRNHEIEDVLKVSRVIAFCALSNYGSLDYARRIFALIKSPNIYIWNTIIRGYVQSSNPSEAFALYKKLLSKGLIPNNYTFPFLLKACTQLVYLNIGSSIHASTIKYGFEDSDAFIQTALAMYTVGGIVRPFHSFKPCCGEPHG

>Atr00009G02220

MASGLQNQRGVQVNLLRRRLGSAVQGLQWSYGIFWTLSSQHQGVLEWGDGYYNGDIKTRKTVQPMELTPEEMGLQRSLQLRELYASLSMGESSQQARRPCAALSPEDLTDTEWFYLVCMSFTFNPGQGLPGRTLSRGQHIWLCNAHNADSKFFSRSLLAKSASIQTVVCFPVPNGVLEMGVTELVTEEPTSIQRMMTFFLDLTKPACSEQSHSSPQNSDHDYDDLDFAKIGHENPGKDTCESETQHFNYYQDDTHNAFSFDISYLPEEGVEFEKEGIKELNGVCEDFKTRSPDFSEHGSSQQLDDLYMPDGITNTCQVQSCPFIEEENGADGSPNNSDCISQTFVEASPKGEILKPFMQDAQDSWDGTPLNYGVNGADNSHYSRTLSSILQRGENFKFTEAPRRLSIKSCSSKSSFLLWKKGSELPRRQIEMPQKMLKKVLFIVGRFRSENSPMTQECDVGKFGVWKKDVEDLNHSHVLSERRREKLNEKFLALRLLDPSISKIDKASLLGDTIEYLKELERRVKELEARGTRKQADISERTSDNYAITEKYSGNKRKASVLDPAQLDSSRVPLDGIEVTILERDISVEMCCPWRESLLLEIVQGLSKLQLDAHSIKSSTNNDMLTLSLKAKLRRKVQTTAGELKQTLTRIVGWG

>Atr00012G03230

METPLRQLLKGFCHDSEWQYAVFWKLKHRSRMLLTWEDGYYNFPKPPCNIQDTTTNAFFNSIGGADYSSDAIDGRVRHSVRDPIGAAVANMSYLVYALGEGIIGQVAFSGRHYWAFAEKVFNGEGNSQFVPEYPSEWQFQFAAGIKTIVLIPVVPHGVVQLGSLKLLMEDLKLVDHVKSSFNMLQNKAGAFFPDPVHCSSNKNNPDPVSSSFDSISQNSFASSAIYPSISRGIQAENLVENSAAPLVSNSFTYFLNQVVKSELTSFQIHHKPLNDFQDLILGEEMGHLAMRQKPVEELPDQNIYEDSLFNFCGQSDSNIMQGSSLSSLTQVVDQDSLLKQSMRSASCKDQEQNGEDYLWALSFPAESELHKVLKPVFSNMGSTDAASTDSSTQTATMSELIEPLVGEFDAWLRSEGSSEHLLDAVVANALSTGAQSCNSSSTLLGGSCLTESNGGGSGSIADDSISDPWSGYLGFVQGSRGTSVRSPSGLSSKAMSTMVEGERKEVFSCSHSKKLIEPSKLTKRRAKPGESCRPRPRDRQQIQDRVKELREIVPNGAKCSIDALLERTIKHMIFLRNVTSHADKLKLCSKVADNKQRPLLVGRSNSDQRGASWALDLGSQTGVCPVVVENLDHPGHMLVEMLCEEDGLFLEIAQVIRNLGLTIIKGLMEARADKFWAHFVVEGPRGIQRMDVLWQLMQLLQPKSPSTQLQANVLHVM

>Atr00019G01200

MHRHSTKTKGISPLSAKCLLHHERESFTLREKAYKVMEDQGSQAFSLEVSYLLQQTLRSLCSLDNSHWLYAVFWRILPRNYPPPKWDSQGAMLDRTRGNRRNWILVWEDGFCNFMASPRERRENWPTNAVSTSLFPNSHTSTYMEPQNFFKMSHEVYNYGEGLIGKVAADNSHKWIFREQPQDQETASISPWQTFVDPQPRIWEAQFQSGIQTIALIAVKEGVIQLGSLLKVMEDLNYIAILKKKFSYLHSIPGVLLPHPSTNLQTNTTHPHTCQSHLQNRPAFYPLLSPQPNTNPPKQTFSPNFLSPQTTLIAPSMTSLESLLSKLPSIQMPNPNPNPNPSLNHIPNPNLSHIPNLNNSISSSTLTYYPLDYGLKEQEEEEDQKEWRSMGLSLECNGPGGKLKEKIESDDNCSNQFGGMDFDLAGEGDITEDCASSLLMG

>Atr00164G00270

MEPATIVDAFWSSNVGAQFSTCCGEASSMGLGDEAEFMSLLLANYPFMGSQAGQEETELNVRVPTSWPTTNMVENEVPIENGAYFSSESLNPSSYYWYQGGLMNTPSECDGIPNISSLKNGVFPMIQSSIPKEFSGCYSVDDQNTSPSLQTISDRPLEGALCEREMMGTENLKELRERKPIKIVTPKRKFEDSSFNDGAPVTDAAFITGEDSQSEETTQIEPQKRAQVSGDVYKSGKTVHSKKTQKLRTDDEEENNTMNVQSSCSYSSEDDSNASQDQNTTTSSGKGSTALNINGKTRAGRGSATDPQSLYARKRRERINERLRILQNLVPNGTKVDISTMLEEAVQYVKFLQLQIKLLSSDDLWMYAPIAYNGMDLGLDLKIS

>Atr00025G00260

MVPTPSSSSKPNWSMATAETPALSSAAAQLQRLLQLAVQSVQWTYSVFWQICPQQGVLIWGDGYYNGAIKTRKTVQPMEVNAEEVCLQRSQQLRELYDSLSAGETNQPSKRPCAALSPEDLTESEWFYLMCISFTFPLGIGIPGRAASRRHYIWLTGANEEDSKVFTRAILAKVVPVQTIVCIPVMDGVLELGTTERVQEDSSMVQQLKALFMDDQGQLQPQKPVHSEHSTSKPATSTDSFPYHRDHHRPHRQHHHQQHQHHQQPHDQQQEQPPLPWQAVEIEEEGDGESESESEREMQTKDIQVGLSPEGCSDPNPINMTAHTSLTNPPCQEQHLPMPTNNLLLDDTNPWPLLHDDISIGLPSSGATMHRDDMVATQEDGHYSKTVAAVLQRNNTNNNAAPPFLVTYSCQFPKSEDTVFSKWKWNNQAATTTSQGGGGQQWLLKYILFSVPFLHSKYRDENSPKARGDGESGSRLRRGVTTPQDELSANHVLAERRRREKLNERFIILRSLVPFVTKMDKASILGDTIEYVKQLRKRIQDLESRNRQMEINLKTRASVSSETQKLSSTKDRTNSNTSAVLTTAQTLNDRSRTMALDKRKRHILEGARTKMAAGGCITDVQVSIIESDALLELQCPYRNRLLLEIMQTLNELHFETQSVQSSSDNGVLIAEFRAKVKENPNGEKVTLVEAKQAIHHILENC

>Atr00030G01080

MGLSLREALKSFCSEGGWSYAVFWKHDCRNQMILMLGEGYYEFGNHPAVSECATNADPDMLLQEWESFLNPLELHSSLVKQQGEDQLRAVVGKMVDQIHVIGEGFVGHVALAGKHQWIFRTGGNSMGPADFKSVSEVASGWKEQFLVGIQTIAVISVSPHGVVQLGSTQMIAENLELVSHVRSVFIRLGRVPGALPYMQKPWSQKLELHSSLGETTPSDFDRNFNAKGVFPPLIADIWNKPMLKTIASESNYQSQPSFNALSCDSIPACASASPLTSNSTGLSPTQYYNNSLQKAFVSVNPSNSVTVQNGMVATGDAQVIMSGGSGPIFPSNNKFPLYDISSISGDRSLVMGSGAVCSTSSFMEQELLTGVQNQGLENRTTASSSTRNGMDSIFDLSFERNRQLGSVHGPGSPKFGNINGTDITQMQTMTGLSSQEENRLFQSPWLSLAGSGGNNPVSCYGDMPLTEISHSVQKGLVGKSNSGKRSPESSHLSYDENSANISANNCLEFVNNGDVKVTDAPLRHCSVDELFDSLGFDLGLNEFQFNWENTFEQGEDSKHCNSSTGISTISDPALDAFNKGRIGDAFFSNTKSDNLLDAMIADIHSVTNEESDENLSCKTALTKLGSSFSSISEQKQVRLVDLSQAPLDGDSKRSSAIFNPKFSVDKLTDSQTDTVSRSPLSNICVENGQSIKCEETSISHAKKLEEPLKVNRKRARPGESTRPRPKDRQMIQDRVKELREIVPNGAKCSIDALLEKTINHMLFLQSVTKHVDKLKHTGEPKMISKEGGLILKDNYEGGATWAFDVGSQPMICPIIVENLNAPRHMLVEMLCDERGFFLEIADIIRGLGLTIVKGVMEACNNKIWARFAVEANRDITRMDIFLSLAPLLEQTTKSSTGPKCASSVSRVFNDTPQTSIPVMGLAKTLH

>Atr00069G00240

MSSPVKETLKSLCNNYGWSYALLWKFKYLDPMLVMCEDAYYASPEDQLEVSNCSSLTTIKPRPSHSFSKDIIEAHVKCETDNEIGSMVDKMLHQVHVVEECLIGHVAHTGKHQWVFQDSSTEKGSLAGYADQQAHFQNVTALGHQFSVGIKTIVVIDVASLGVVQLGSIKKIMENKEFIAHVRSLFLQVNSVQEVLSASAQKASNPTIHAIATQRTSASETPSRISSSKRDCAVPLSFGPFKEIDGKHSITLTQPSNSGIIKPQKGIEHQISSSRVTCTSAHVDIISGYPNMESNMRNLRLNLTSQLPTSKTEAQVIIPYPNLNLLQDMPHFNPNPKMDMVQSASCLASEANCYASLALLEQELRSRMKRQEDVNMFSAITTDSCLSQNPLPNSQAESIPSLSTYGTVGSFDAGNDTCILESDRPYLYPGKHVDAQISSSPFQGIECTGSGNSPKVSEDLRQRSNKPSLSRQNLGNIGSSTWTPGQPEQVGDSSMDSLGNTISQCLGMDSLFGSIESQDQNWLRDMLTQNIPQSSSLTSSHQLKENPKLNAHNSDGVRETSSNGPMLPFGVDELFGTLGCDLRNSSEQARWDNVLLPTAEGNLPNSSNGVSTNFSELDLSIVPENGDFSENRSGDLLEAVVASVSSFSGKSSCTTLTRSGNTSLCSSQVRSSVISRENSMVLGSEGTVFTDWNLGKAKSDIQSTGLSISRVTSWIEDGQSMKKESAIIGQPNKADEPPKVSRKRARPGESTRPRPKDRQQIQDRVKELREIVPNGAKCSIDALLERTIKHMLFLQSVTKHADKLRQASEPKMIGNESGLVLKDYLDGGGGATWAFEVGCQSMVCPIIVEDLNPPRQMLVEMLCEERGFFLEIADIIRGFGLTILKGVMESRNDKIWAHFAVEANRDMTRMEIFLSLVQLLEQAAKGTVSSSLQLPRVIGSGSHLFTSFQSPIPPAITLAERFH

>Atr00088G00400

MEENISHGLPLTLTHLLQQTLRSMCVHENSPWVYAVFWRILPRNYPPPKWDVNGGGCDRSRGNRRNWILVWEDGFCDFSATISDTAEMRTSDCASSSLYTTSEFQHSRGLQPELFFKMSHEIYNYGEGLIGKVASDHSHKWIFKEPQDHEINFLSSWHSPADSHPRTWEAQFQSGIQTIALIAVREGVIQLGAANKVMEDLNYVVLLRKKFSYLGSIPGVLLPHPSSAAFPLQGDGGAPSVPHNWPFQPSIAPDSNTCHPEYHDYLSSMNPNFHGSGIKIMPSMTSLEALLSKLPSVEPSLHSASSYCGPASSYCGPSPLFASNTQGIDKMKEEENDEYGNGGESSTAVHLFRHR

>Atr00094G00110

MVGSDRCKEAVGMMALHEALRNVCMSSDWTYSVFWTIRPRPRSRGGNGCKVGDDNGSLMLMWEDGFCRARGFNGGGGAECVEECLDGDDPVRKAFSKMSIQLYNYGEGLMGKVASDKCHKWVFKEPSECEPNISNYWQSSFDALPPEWTDQFASGIQTIAVIQAGHGLLQLGSCKIIPEDLHFVLRMRHTFESLGYQSGFYLSQLFSANRNNGASSSSNPTKQMVRPPVFNWGQRPPSSANPSPPPPPPPATNFNISPVFLLPHQQCPTSDELESDIKWPNGLSFFTALTGGSDDTKLLFGPEGMVGDQHTTATAPPPLEGCGVGEYLSLEGQQARSGDARLRKADKFKRSLTLPARVASSSSSVEHHHGSHVEPGIYSDIMETSWID

>Atr00024G03270

MTLARESKATQEANNLIMFQAYPFHSSTYVSSAGAPPLHPSSLYLGNTFEAQIPFAVTDPTELSNCTDLVMGGSSSSSSEDLEGLGAMLSFTPGDNHSYNRTPWAYPCNSPLIFEQGGGVGDHFSSSIPVESFRLFSSGDSFFRGEEKERERSLCLQHKRPYPGEEIKVQASKRVCGYSKAMDVDMDMDMDIDMGGDCIPSLLFRKPTKQRHVPSKDPQSIAAKNRRERISERLKILQDLVPNGTKVDLVTMLEKAISYVKFLQLQVKVLATDEFWPAQGGQAPEVGQVKEAIEAILSSHNSKASSSSPSNPTM

>Cm05G00470

MYTRFGQCLVPVNQAGGCCTAVSLGHGAHQRLTLQRTSRHMLGFCVLPQWRLNARVRSACRLVRNKAFQAHQRVARYMLRVSHASVTTQGEQSPWTAKDKSVKEANDSPYKHTVSLPETSFSQRANSTEREPQIQRFWEENRIYEGLYEDNSRNAEFLLHDGPPYANGSLHMGHALNKILKDIINRYFILCGKRVRFVPGWDCHGLPIELKAVQSARSNHREALDPLQIRLLARKFALEAIEEQRKGFKRYGVWGTWNAPYLTLDHQYEAAQVRIFGEMFLKGYIYRGRKPVYWSPSSRTALAEAELEYPDVHVSRSAYVAFEVVDAGKLRGIISELEFDSFRVAIWTTTPWTIPANRAVAVNPNLPYSIVALEDEQGKPCTLLVVAQDLVETLEQKLHRPLSNKKIVPGSELVGMTYRHPLYPMEVYRVVAGGDYITTESGTGLVHTAPGHGLEDYNVGLRESLEVFAPVDDVGRFTSDAGPELAGLPVLQEGNARVLELLRSSNALLLEEKYEHKYPYDWRTREPTIVRATEQWFASVRQFRDKALEALQSVQWVPPAGENRIRGMIESRDDWCISRQRPWGVPIPVFYDEDTGEAVISMEIIQYVSEIFAKHGSDAWWTLPIEQLLPASIIQTGRRLRRGLDTMDVWFDSGSSWASLSKSHGEGSSSSWLPSVVADLYLEGSDQHRGWFQSSLLTSVAVRGYAPFRAVLTHGFVLDEKGLKMSKSLGNVVDPSEIINGGQNQKKQPALGADVLRLWVASVDYMSDVLIGPTILKQTADAYRKIRNTLRFLAGNVPRWDETCPVSVNALNDANLPSLDRYILRRAAAVLKDIEDAYKNFTFNRIFQSVLRFCVADLSNFYLDIAKDRCYVAAPDEPRRQTCQATMWAILMDMARALAPILPHTVEDLWQCLRKDGKLPSGAPISIFQNGWYSYVPQILSDEYEDKKWTIVREVRELVNKALETARTNGLIGSSLEARVIIYPHARHVEEALTSFDPSNGVDDLRYFFITSQVDVVRCSGNGESPGAASETHVEVQKAYGSKCARCWNFSETVGALHEHSQVCERCVRSLKHLGVERV

>Cm10G00530

MLNGSTIGRPSSAYPGASHIAEATTPGEIGGPVEETELNGYLSANRKVAQATPLVQNQVHGRSTAHAFDEHIHVGSPPVQNGISQLNGGNFGPLVTASHANDGVAFEFHPSGIFIPSSRNSNQQGPPGAFAPEWAPCARTWTAAPAQTVAGAVYTYPHRNPAAWANSADGRYFSRSLGRGLSPGHSNAIPIHGVQKLGHTASPLKQESIEPGCLTDYYDRNRVAIQLTGKLIASGQLESLLEQSPEDEHVTVWHPATGRTVAGNAAPYRRNLDAWLEKNPGWVEKSFEEKSSKRRRASRRSRAAMAAFSSLSTANDHVRDILKEVDQKVREALRERRRTGTPEEAMMVAETVPIASEASFQDSFTTTVMNYVLKDWTSDEVVRLVELIVRCSEELAHLGHGVDSDDECCDLAELAEHGSIEANITESFLNCRKHRSRGPCSVAMPLRTCAAQREVAPGSGGGVGGGSGSGGGQLVPLDMDDIQFVPSPVRNETPCEIHPHSTDTNSAWRKIFEALPSKHPSSVILRTHILFSQVLLAAAHRGTLTNLIQSPSPTSSTSLGRTHLEHLNRALSSSHDSSYGAEQLAGFASLGLHAHSPSVSPATGLSCSAPTESNGTPRSMFRDVRVTVWDPTTGKTISGNAAPCWRNLEIWMKEHPGWCVKPEDELSASRRSKHKRARELGIPCSSSPFPASVSGFETALAEIRTRSDRLPGSRCAVAAGQADYSSMAHSLPDEHDCIEGLLMMHAGRPLGNARAEVADLRASPYGKPPLPPQPLLLTTSSDQSSTPETMHANGETA

>Cm10G02200

MDVSAIAESVRRKLDERIPRTEWTEEERTWYKEDKKRRKEAEAALKKHVDLEGNVPSPSEEAVLSAQLESLKMDADAPAFGRYTLKALTTSSCGDDSVRRTWVEAGTLGVAHPACATNQEVWLRGRVQTVRAQGSKVAFLVLRHQYATVQVVLVADSVPSVTRDMVRWLARSGEVTPESVVDICGILVPAQVKSCTQNDVEVHARRIYLVSRARVPLPFEIADASRRADEDGPHVHRDTRLDHRFMDLRVPAHQAILRLQSGVSTLFREWLVAHGFTEIHTPKIIGGSSEGGAEVFRLRYFDENACLSQSPQLYKQMAICGDMRRVFEVGPVFRAENSNTYRHLTEFVGLDIEMEIQESYHEILDVLEQLFQHIFEGLEQRFGAELSSIQRQYPFQPLRYKRGRGLRLTHRQAVDLLREHGEQVDDLVDFTSEQEKALGEIVSKEFQTDFYVIDRFPAKCRPFYTMPSPDDSRYSNAFDVFMRGEEIVSGAQRIHDVDLLRQRITEKGLDPATLQSYVEAFQYGVFPHGGAGIGLERVVMLYLGIGNIRECSMFPRDPKRLYP

>Cm11G00970

MPSTGLIQAAFVGLVQLPVQVRARGFDVPGKQRRCFPCNQRVLRFRTVHRQRGIIRELRACAHRVERSRATTSLMPPQEVDAALASTLRASGVSASPEYLRNSAFHIPDYETYCEMYTRSLRDPTGFWLEMARNNFYWRDPNAITPDSVYRYNFDVRQGPIQVEWFFRAETNICYNALDRHVEQGLGNRVALYCEANDMDDMDTHRAITYAELLDMVKRIARVLRRQGVRRGDTVTIYMPMVPELPAAMLACARIGAVHNVVFGGFSAEALAGRLLDARSAVILSCDGIRRGPKRIELKATVDEAIEICQTRDPSFHVSCQLVLRRLGESMLPNLTMQAGRDVDLRVAMEEAVSGPDDAIEWMKAEDPLFLLYTSGSTGTPKGVLHTTGGYMVNAALTFKYSFNYQPGDVYFCTADCGWITGHSYVVYGPMLNAATQVLFEGVPTWPDPGRLWAIVDKYQVTHLYTAPTAIRSLKKAGDEYVKRYKRTSLKILATVGEPINPQVWEWYHQVVGDSRCNVVDTYWQTETGAHVVISPPVRGFQQKPGSAGVPFFGIRLAVLDDRGKEIPFVPGQESEGYLCIRDPWPSCLRTIFGAQARFENGYFKPFPGYYMTGDGCRRDGDGYIWITGRVDDVINVSGHRIGTAEVESALVRHPAVAEAAVVGMPHEIKGECIACFVILADGYTPSDTLRNELVQTCRKEIGAFAAPERIIWTPALPKTRSGKIMRRILRKISEQGRDLDRSVLGDLSTLADPSAVDAVIQSFLS

>Cm11G01580

MLNGSAGISYSGRRNLRSNVPLDAGLLLLLLVHKVPNMDQVAACFRAAYADAVGSTPLLASAEERAVFVKNSLEFALKVEAQSRDVTAAGAEQQRTEIDRARQRLWFFCVGAAELLKEPGAQFPSPSLVTAAQQALRQRQQHWCNRLAAASRLSADTTLSTADEVDSLIALVRIGLTVLGLFVCLSILGEREHWRGPFLCELWQQGRAVYEALERQKASSAALRTTASFLQSYWLDALCTATWWIADLSNHAAATAWTLDTSPSSLLQIRDSASAQCSVWNPLHCSFTQPCATIRDQHWNCRGCSALQASSWTPHQCTFLLTGRSFRDQHWYRCSTCAFREDEGVCTSCALICHAGHEVSYVRFSGFFCDCGANAAATTPRTKCLLCTLVRSEETSSTTSSSEALHTSTETPRGAERRPAKAPELRDEHLLWMRPPSVSPLVELDSTKACFYHVFMEIAQLHGSSDRCPVRDASGSGETAPAGVLLAKAITEHLQSTFEWLRDSVLFRFLQSPTPGEIPLETNAQSQSVVQMPGTEQAPRKAFLPALHRAAAAGDESPRTATCNARDVREIQQDTCQVRGAGTYIQRATWKPEAVQVLTRVRHVDTNQGQSTLGMGVFWQLLPCGKQEARNPRELHWHHEPVLAFVDGLFLVTLAASEHHVAFLSDEADIRRSSASVPVHPSSEPLERAPLAGCACTGTVRSTRPTALDARRWIRRPLGFCAGELGFAPSCPRRGSLQRLLALNRETSTLRLYCVGSTVADVDGSLHAQSCASESPSRPVAAAAHPVVEPDKVRSVTLPTSGQSFTASWGTTDAAAAGDGTTVVHSAEALVGASGSRPAPAGPSANTATLVNAVGVSADAVVSDQLAEYPSNTRSRMQAPARRSTIRAHRAQRVLQRWREYHQVHAKRCLRLLLRPVPGPGAASAEGGTAYGAAEVATVPKTTPATLSLEQPERSSTGTMTFAAPAEASQFGARRATSTDTTSSQHPGRLYLLLERQCLETGEPIQEASWLDAYAIDWLLCLGRSRSSLEVLSIEYACWCWFCRWIREAHGEAAASAADTTSRPAGPDGTETCLTVSEAPSGTVSNDEEPFLWLSFADMDEALAARQPCTLRPAPCTEAAAENARLLLEQTTGSESQILRWLPDAAPEHDPWHMRERKRFRLMFSNARDGVAEGFVASSATTARDIWHSEVTEATLAGPVTAWTYGRVFFDASDRNRSPPGRLLLFLVLSNQHLVVYEYVAEARAASSVDIETSGLRQVASLELPLFDTTPCMHTLANAPEEKPSSPSATSTSTPARVLGLFYEAAQQELMLYWSNGHWSRWKLTWCREPGLDAAPEPDIGSRSIRNGHRSLSSERLLPFMPQLELTSVNSGCLFEGKAIVAEREALEDVQRVPVRCPHGCCCTSSSCSLMLLRSARTLYLLWWQREKRCWMATRQGAPYSSVSDVRVRQAGANSGAETAAQNTSMATQADSREQLAILCSLDAGTQDALLPERARRMDRPIPERCRQQRLIQATGWAQLEATACESASAFLIDVIMLLEDGTVLGTQTRLSCDVQISWNALEGSERPDRQPVSAVDSWKQQKHPEGVRYSCSGDVRLSQPHERDLSTAAARFAHLYWNHSTAEAVWLPASPEDGVAAADDEDEATPPDVASLPETPVEDHRGVRGNPVLQVRYPWIHCATSVAYQMTCSRSVSPWLGRPQGNSVPSALLLADQRNSNGGLVGFFELLDRVPSGMLAWTGDMWQLGLRSMTETQSWLEQWPSDSMELLSAPIRAFTYRSSAEQRPFRREDDAPGTRLTVLAYRVHSTSSSGAAREKRTAGLLQMPQRCEDTPLPLAADALLGQEQGQHHQLVFHVSNVGDNDASWALLGIRMAWLQPNRSGVQGVATPTQLEPQLVELLVQGDHEPATVPERGTSAPWTSRLSLELLPDPWVNPVPPEQGTWRWMEYCWDDPVVLRRNPCNRQVVLTIRAPAMDAAEASAWLVLGTIEVYGVRLCDSRWPNLLGFLEWRRKLLRQLDSAGNTRPRAEARKGEVSALQSQATTRPGTALSSTCPRLADKDEIGKALHAQDRLAPALSVSGTAVSSGIPVGNALPAADDGIATTVEQRSERSRMEINASSAEVASSSAAVSSGTPLLPAQHIALCERDKLRSTEAAHSAAVQDIASADNIEDTAAGRIVWRACWMLAAILAITGDNATTGLQLVDGSLQHRLKSFPGASDAAEDANSPQMKHPSGYLDRHWFGTWRALWKLLSTRKYCWQKRLMFAQSLQETGDEHAALEQLLHAETHQARRFLRLDDTENTLLLQQQARKCFGCWSADQDQVPAAWTRDLVCWTVDSLDTCSDQHQGREALLWAIQGLEWLYEACASWTTTTTTTTATTAAHSTTRSDQGHPLSIDAASVCSSQETLVLSIDLARVYAETAAAWLSAFDTEGAGSASAPNEEDESSQRDRPTFTERNLWTGFLVAAKRLGDAWCSSSRCWQQRPSRHANEGSSSELDFSAVFSWLFEPFITEAAPNVLTRFLQVWLTDVDKLIGVDATHPSAFDYLDTWLRACSHHCALHSSEQTECRARMQAAPSALHGRCALPWRRLPPERLRLILRASLLCVSDQNHWWSLLRLINNVVTSASHMDEALRTAEHFAALLSDEEAVVESSTRAMDSWSTLEVFWIVSNTVAPHMMDSMSLFETATGPGATFEALAAGRQRECSQRRETHHDIALSVWIRCFQQQWVPDHWSDCTAWLRAYFFLVLAASCLWFTVGNGNDLSGSADATTTCIAAAAAAAATSSTCSRDTTTARMRTSTVVDDGDDDATTVSPASTFTATDPATARMRPCEAGVASACSPDSLVEASNSGNEPGWNVAATRTLIESVELALWDWYGAGLCFGYLEQPPAALTDLVEVLIPSRDVVSAAFQLVAVRQTLAALSTVDNGASALCDACCDPAGQTQRKTSTYWNRWQRQQRVQQAVLWISAVRERELCTDLGLATLLPTLVTALKHDDRYSAGLLLQSLLTLGSELAHWQQISLSLRLTWAWQLLRLLLVEPELRHWAQDALVIVLERLLCDLMDAPDPGAWAATGQQSPACAGVCDASEASMLLVLLRKYVALYRPRALCKALPTLADWVTLLAAHLHITAELVGLERFLMQATFPGSTTAVTERSCGREAGATPSNNQEQVSWPLSWLLASPVCVDCFHEAQGAGLTLQAAGVHGARSSASLQTAQLSGSTCLGGASSAAAVAGTFVKRQPLSTLATCVRFDDDRIQATWTSPKRLHCLGVRLRHTRRSLATILGFRVLVRSLGTALDAAEVPGASATRAPANWRCLCELVYRQDGALLGVLCFLDEWPEPLLGADNALATGLALGLTRQTGRCCVPETQICPRCARLVSERGGVCPACRENVYQCRACRFIDYGRLDALLCRECGNSRLMRCDIQVETSDVDQEGTLLEALMHWRFRHTTGTSSSSISNTGSGGGTETAPGGNASAPAVSAETHEPKAAAAAAAAAWIGARERRTALVAWCLHAPTEALLPVACARCAQRRLPRLWTMFAEAVGRAESASTDQWSQVFEPLHRALLVLFSSGGFIDGERATQQLPVTLGSLPQPARDAWLRILEEALVASESGPRNEHGEDARSRCEFQCDLGQAYLHLLTQSFACSGAFGADRLLQGAEATSATTMLGGMQQRIVIPATLFNSTADYLAAVYSQRWCVRLFQRRWCRGQSPWRRPCPEQSCSEACIWRALFHPYAPGVRTWALRVLAQRPGYWRWLFHHRILLFGCWQALRHGCVDMLDAYWALWSWLWHHERAWVDFWDSTGGLELLCTVIAFAAGGVPSGADSKTAIACGCRRNAWVPSLAEAVIHACLERIQEAVVLRGRLQWTRAALDCLLCAFEALYVVADTRMSLAAQLALLIQRQAEREPLLAHWQARWMTQRLERLTMSAGTVSCLHTGATVLERNEGNAYDR

>Cm11G01790

MFGAVALRSACRLWLRSTQCTGPWAEASRYRSANAERCHKLQNVSLYQLGKRCTSTWNNGDQHPSVCFTGDVLSLQAGLNTRFTADRASSALHSRFGSVRGASSLTPPRSASRGLAQGQAGVQNGAVAIVLLSGGLDSATTLAIARSQGFICKALSFDYGQRHRCELEAARRVARAQQVTGQHWIRLDLGSIGGSALTAPDIPVPKAANVDDVWEHTPETIPLTYVPARNTIFLAYALALAEVQGSYDIFIGANAVDYSGYPDCRPEFFHAFERVAELGTRAGVAPSDARRFRIHAPLLQMPKTEIIRTGLALGVDYSITHSCYDPLDESGKPCRRCDACRIRHTAFRELGLEDPQLSRFGAGSAAAPRTAKSE

>Cm12G03390

MGAELNQFTNHVQNKSVVELDALIIGAGVAGLYQLHKLRSAGLNALAVDTAEDVGGTWYWNCYPGAKFDSECYVYQYLFDEKLYKDWSWSERFPGQPEIERWLHYVTDRLGLRPYLRFCTTVTAAEYDEKRGRWIIKTDRGQTYDARFLIGCCGMLSAPLENRFPGQERFRGRIFHTARWPREPIDLNGKRVGVVGVGATGIQVIQTIADKVGQLKVFVRTPQYVVPMKNPKYNPDDVNSYKKRFEELRELLPTTASGFEYVIKHRWADLTPEQRLAVLEETYNDGSLRAWLGGFDEIFTDPQISEEVSEFVREKMRARLKDPKLIEILVPTDYGFGSRRVPLESGYLEVYHRDNVEAICVRNNPIVEVVPEGIRLADGTVHELDIIILATGFDASSGAFIRIDIRGRDGRSLKEEWSRDICSALGLMVHGYPNMFLTGAPLSPSAALCNMTTCLQQQVEWITDCIQYIQAKGHNVIEPTASFQEAWVRHHDDLANSTLFSKTNSWYTGTNVEGKPRRMLSYTGGVGKYRQKCAEVAAGGYEGFAVA

>Cm15G01460

MLSDCSAFVVVPRFCYRLHVGASPPGALQSSRRRVARLRERSAGTLPWARATFCKAGCRIGSSSCSVLLRWKSCFLAGEERSHLRQLGSLVQQPGALRQHVSMSLFSTPAQVSSYEIIESYPHDPSAFTQGLVYAEAPIHSPDEAEWTMLQTSTEVFYESTGLVGESSLRCVHKLSGKVLKCVQVPAPHFAEGLALVPRPTRMLFQLTWLTRTAFLYDGVSLKQVREIPYDLEGWGATYDSNRHCIYTSNGTDRIHIVDADTFEILDQVRVRSSPYKFSRSCPTLWYLNDLEMIPDTDELWCNVFFSDYVAVVDPASWELKRWIDLRGLLLPEHCLPCHEVDVLNGLAWDRNRQLMYVTGKRWPRMYAIRERP

>Cma2000460

TTLFNLLCPPCPELFEHTRRRTLLITGRKEEEEYQKKKRRRRKGEAMAASAVVAGSPDFMAGFQLCDDDHAFLEAFIGSYDDPLWPVSSTPAENAPAPSSSSTALETALPAGTAGGAAASDNSFQQRLQFVVENCSPAWTYAIFWQLTLSDKGEQVLGWGDGFFNSREEHQQTIVSEVHQQLRRRILRELQSLIGGDGAAAAGFDALETDVTDTEWFYLVSMMYSFLLGTGTGPGKAFASSGHVWLRGPQSRDCPRAQLAQRFGIRTILCIPIANGVVELGSTDLIQEDPALVHSISSVFSNRYRPHPNPLLLPSSSYTKLYASSVPSHSDAFSSGQSSYDTNLVSSFAMPNSTLPSHSFGPGLPQPNPSFASKGRASTQNLSFAPPNDRSRLSWPSSIAKNMRIGDEDRFPALSELNKASVSNESSLLSQTSEVGDSRLLLADLHSRKQSLWFPEISHSIGSHPNNGHFSKAATAQMYNKNVASAPKGASQKPVEIAAKFHLPISVQSFSQSTVSSGDFISASSTSMKRGPEMRDSMQFLDVPAKSASHIHPAVNLSQLGASTQVLCQQQAPSQGMVNGAARRDDLLGMHVTGPIPSSIESEHSDMETX

>Cma2000461

TTLFNLLCPPCPELFEHTRRRTLLITGRKEEEEYQKKKRRRRKGEAMAASAVVAGSPDFMAGFQLCDDDHAFLEAFIGSYDDPLWPVSSTPAVNAPAPSSSFTAPEAAMAGTAGGAAASDNSFQQRLQFVVENCSPAWTYAIFWQLTFSDKGEQVLGWGDGYFNPLEEHQQTKVSEVDQQLRRRILRELQALIGGDGAAPVGFDALETDVTDTEWFYLVSMMYSFSLGTGTPGKAFASSSHVWLRGPQSCDCPRAQLAQRFGIRTILFIPIGNGVVELGSTDLMQEDHALVHSISSVFSNRYRPLRNAFLLPSSYYTNPSASCVPSHSNAFSPGQSGSATHHVSPFAMPSSKLPSHSFGPGLPQPNPSFASKGRASTQNLSFAPPNDRSRLSWPSSIAKNMRIGDEDRFPALSELNKASVSNESSLLSQTSEVGDSRLLLADLHSRKQSLWFPEISHSIGSHPNNGHFSKAATAQMYNKNVASAPKGASQKPVEIAAKFHLPISVQSFSQSTVSSGDFISASSTSMKRGPEMRDSMQFLDVPAKSASHIHPAVNLSQLGASTQVLCQQQAPSQGMVNGAARRDDLLGMHVTGPIPSSIESEHSDMETX

>Cma2000462

TTLFNLLCPPCPELFEHTRRRTLLITGRKEEEEYQKKKRRRRKGEAMAASAVVAGSPDFMAGFQLCDDDHAFLEAFIGSYDDPLWPVSSTPAENAPAPSSSSTALETALPAGTAGGAAASDNSFQQRLQFVVENCSPAWTYAIFWQLTLSDKGEQVLGWGDGFFNSREEHQQTIVSEVHQQLRRRILRELQSLIGGDGAAAAGFDALETDVTDTEWFYLVSMMYSFLLGTGTGPGKAFASSGHVWLRGPQSRDCPRAQLAQRFGIRTILCIPIANGVVELGSTDLIQEDPALVHSISSVFSNRYRPHPNPLLLPSSSYTKLYASSVPSHSDAFSSGQSSYDTNLVSSFAMPNSTLPSHSFGPGLPQPNPSFASKGRASTQNLSFAPPNDRSRLSWPSSIAKNMRIGDEDRFPALSELNKASVSNESSLLSQTSEVGDSRLLLADLHSRKQSLGFPVISHSIGSHSNNSHFPKAPTAQMYNKNVASAPKGGSQRPVEFAAKLHLPSFSQSTISSGNFMSASSSSMQRGPERRDCMQVLDVPAKTASHINPAVNLPQLGAX

>Cma2000463

TTLFNLLCPPCPELFEHTRRRTLLITGRKEEEEYQKKKRRRRKGEAMAASAVVAGSPDFMAGFQLCDDDHAFLEAFIGSYDDPLWPVSSTPAVNAPAPSSSFTAPEAAMAGTAGGAAASDNSFQQRLQFVVENCSPAWTYAIFWQLTFSDKGEQVLGWGDGYFNPLEEHQQTKVSEVDQQLRRRILRELQALIGGDGAAPVGFDALETDVTDTEWFYLVSMMYSFSLGTGTPGKAFASSSHVWLRGPQSCDCPRAQLAQRFGIRTILFIPIGNGVVELGSTDLMQEDHALVHSISSVFSNRYRPLRNAFLLPSSYYTNPSASCVPSHSNAFSPGQSGSATHHVSPFAMPSSKLPSHSFGPGLPQPNPSFASNERASTQNLSFGPPNDRGQFSWPSSIAKNIRIGDEDRFTALSELNKASISDESSLLSQTSEVGDSRLLLADLHSHKQSLGFSRHFPNAPTAQMYNKNVASELNKASVSDESSLLSQTSEVGDSRLLLADLHSRKQSLGFPVISHSIGSHSNNSHFPKAPTAQMYNKNVASAPKGGSQRPVEFAAKLHLPSFSQSTISSGNFMSASSSSMQRGPERRDCMQVLDVPAKTASHINPAVNLPQLGAX

>Cma2000464

HL*AVVAIIY*LGI*VVEEEEKEEEEREAMSASAAAAVAGSPDFMAGFQLCDDDHAFLEAFISSYDDPLWPVSSTPAVNAPAPSSSFTAPEAAMAGTAGGAAASDNSFQQRLQFVVENCSPAWTYAIFWQLTFSDKGEQVLGWGDGYFNPLEEHQQTKVSEVDQQLRRRILRELQALIGGDGAAPVGFDALETDVTDTEWFYLVSMMYSFSLGTGTPGKAFASSSHVWLRGPQSCDCPRAQLAQRFGIRTILFIPIGNGVVELGSTDLMQEDHALVHSISSVFSNRYRPLRNAFLLPSSYYTNPSASCVPSHSNAFSPGQSGSATHHVSPFAMPSSKLPSHSFGPGLPQPNPSFASKGRASTQNLSFAPPNDRSRLSWPSSIAKNMRIGDEDRFPALSELNKASVSNESSLLSQTSEVGDSRLLLADLHSRKQSLWFPEISHSIGSHPNNGHFSKAATAQMYNKNVASAPKGASQKPVEIAAKFHLPISVQSFSQSTVSSGDFISASSTSMKRGPEMRDSMQFLDVPAKSASHIHPAVNLSQLGASTQVLCQQQAPSQGMVNGAARRDDLLGMHVTGPIPSSIESEHSDMETX

>Cma2000845

KACTSLAAFKEGKQIHGGVIESGFELDEGVGSALIDMYAKCGSLMEARAVFDKLQKSFLLPWNVMITGYAQHGDGQQALGLFHQMLQQKIEPNNVTCLNMLKACSSIAHLEHGKLVHGDIVESGFESSTAICNTLIDMYSKCGSAEDACRVFENLPSRDVVSWNAMISGLALPHSQQALCLFEQMQQEGTKPNNITYVNILKACCTTEGSLDQGKWIHTHVIENGFESDLIVGSSLMDVYAKCGSLKDARRIFDKLPKRDVVACNAMIAGYAQHGQGEEALQLFQQMLHEGKKPDDATLVSILKACSSVTDVDQGKSIHTLILRSEFKLDVAVGSTLMDMYIKFGNIREARRLFDKMPKKDVVTWNALFTGYVQHGQPEEALELFQQMQNEGTEPNNITYVSLLKACSSIVALEKGKRVHGHIIESRLELDDFVGNALVDMYAKCGHLDGARKVFHEIPKRSVVAWSAMSAGYALRSNYKVALQYFEDMQREGLKPDDVAFVNLLSACSHKGLVQEGCRHFKSMIDDHGITPVLEHYTCMADLLSRAGRLIDAEAILKSMPCQSNVVGWMSLLSNCRTHGKVEIGRRCFDRIVAIDYKYAAAYVVMSNIYSDAGMQKEADKIEEMREHAQAWKMPGKALVEVESKLHAFMVGDQSHPRSR

>Cma2001431

SSRRMPRVSASREVPPFFRSTIVELFVYLNAKLGAPPTSH*EPAHTVEYAVNI*ATSVRNLTLGQLE*QMQSGHGISMSPYDSHTGLAQSQGAVGVVTGAPKPRVRARRGQATDPHSIAERLRRERIAERMAALQELVPNSNKTDKVSMLDEIIEYLKFLQLQVKVLSMSRLGGAAAVAPLVADLPAEGKNSLAAAALGQNAGLPQDAMTAAEHEVASLMEEDMX

>Cma2001432

QVDECREFLHRGRSLRSFEVPSSSCSYISTQSLELRRPRIENLHTQWSTR*IFKQHL*EI*LLDN*NSRCNQVTVFQCHHMIVTQV*HNLKVLLELLLGHXTGAPKPRVRARRGQATDPHSIAERLRRERIAERMAALQELVPNSNKTDKVSMLDEIIEYLKFLQLQVKVLSMSRLGGAAAIAPLVADLSAEGKSSLAAAALHQNAGLPQDAMMAAEHDVASLMEEDMGTAMQFLQSKGLCLMPISLATAX

>Cmaa2001433

RERIAERMKALQELVPNSNKTDKASMLDEIIEYVKFLQLQVKVLSMSRLGGAAAVAPLVADLPAEGKNSLAAAALGQNAGLPQDAMTAAEHEVASLMEEDMX

>Cma2004939

SPLGLQVTLNCYPNYIYRPPNLHTSPLGLQVTLNFFLRHSKKSLHRAYPHP*PKPLQKVWKAILRHSRILCREHIHRQQLHVWNKKMES*KGFKGRNTMTRVAGPWRWHAADDVERRRWRWDGYRYARERRLI*GFGAGQAPRLLFCYSLSLMEPALVSTQTIAPSESQLKDCFSWLSWKHPDSVRCCDPGSASNSAQSSSSSCRLQDCLEVLESMDKEGTLNMRKIKHAKFFNACKKHGAVKEAFRFTQLIKRPSMSTYNQLLSVCARAKDIQGAFEVLATTEKAGLKADCILYTTLISACAKAGKVDSLFQVFHEMVNKGVEPNVHTYGALIDGCARVGQVGKAFGVYGIMRSKKLKPDQAIFNSLITACGRSGALARAFDVLSEMTAEPIHLEPNHVTIGALIKACTQAGQVDRAMEVYAMMAEKRIQGSVEVYTEAVHACSKIGDLDTAFAIYKDLKDSLIQPDEVFFSALIDVAGHAAKLDACFEFLEEMKQLKLDVGAVVYSSVMGACSNTGNWEKALEVYREMQTASLSPTISTMNALLTALCGGGQLKEAKRILQEMRHTGITLNDITYTILFGACEQAGDIDFSFDLHKCATLEAIIPTVGMCESIIGLCLQVIWKLSRGPTPTSSLGTMQSISFCERCTSWALSVYRKGVATGVVPTMKMFSQLLGCLRLPRTVDDNVLYNYAFAEHSNHKQLSLIDGFGVYDPRALALFEEAAALGIVPTFSYTGGPIMIDTSLMPVFVAEVCILTVLKGFKHRLAAGARLPAMTIILGITDTELVTPDSGQKKVTLISRTSQAVAGLLRRLRLNFHGYESSGKLKVTVQAIKHWLHPRPHKLPMLQLTSGPRSSFLGKKIVEQQRMLRYVESSRWSTHSSQQCNNIDAAQHLPPNNKDVQMHHPVSNGVHALESVKSDETVRIPSAKEGVLST*E*IKPQMGSA*LVRGLAQAGN*FCLCNHLACLRIEPWGQGPCSCQCCMHIFPGISTNQHVGIRNAKSFLIV*FLCD*TPGQWSQVHA*NQALECGQAKISLILNKTKLSLTTNKKKKTCXKNNDWFV*F*AFFVVLSTQETIHKKKTSVNKQRFX

>Cma2007900

IRWEILQTVEKLLNAQITPMLPLRGSITASGDLVPLSYLAGVLTGRPNSKAVTAEGKVVSGSEALKMVGVEKPFELQPKEGLAIVNGTAVGAGLASMVCFDAHILSLLSVVTSAMFCEVMQGKPEFTDPLTHRLKHHPGQIEAAAAMEWILDGSAFVKAASKLHGVDALKKPKQDRYALRTSPQWLGPQIEVIRFGTQLIQREINSVNDNPIIDVSRDLALHGGNFQGTPIGTAMDNIRLALAAIGKLMFAQFSELVNDFYNNGLPSNLSAGPNPSLDYGLKGGEIAMASYTSELEYLANPVTNHVQSAEQHNQDVNSXWG*YPQGRQRRPSKC*RS*APPTWSGYARRWT*DISKRICRRL*RGWCREWPNRRSRRGKMECCCHRASVKMSCLKWWQMSTSLHMLMIRPAQDTL*CRGCARCWLSTPSGIPLTSEMRPRQS*AGSPSSKTNCVPSCPPSLSMSGQLTTKAARPSPTRSRSVALSHSINLSEPSWAPNCSRAPATSHPDKILKLSLMPSRKERSCSPYLNAWKAGPRLQSNAKCRNNFLP*TRVQCVMVSSFVSIMAFSLHMA*CIPVP*LKFSSYRWFGIFNX

>Cma2007902

IRWEILQTVEKLLNAQITPMLPLRGSITASGDLVPLSYLAGVLTGRPNSKAVTAEGKVVSGSEALKMVGVEKPFELQPKEGLAIVNGTAVGAGLASMVCFDAHILSLLSVVTSAMFCEVMQGKPEFTDPLTHRLKHHPGQIEAAAAMEWILDGSAFVKAASKLHGVDALKKPKQDRYALRTSPQWLGPQIEVIRFGTQLIQREINSVNDNPIIDVSRDLALHGGNFQGTPIGTAMDNIRLALAAIGKLMFAQFSELVNDFYNNGLPSNLSAGPNPSLDYGLKGGEIAMASYTSELEYLANPVTNHVQSAEQHNQDVNSXWG*YPQGRQRRPSKC*RS*APPTWSGYARRWT*DISKRICRRL*RGWCREWPNRRSRRGKMECCCHRASVKMSCLKWWQMSTSLHMLMIRPAPPTPSPRGCARSSSSMPSATPPTSSMRLLLSFPASPSSKTSSAPSFLPSSSMSGLPSTTTARPSPTRSAAAAPSLCINLSGNSSALASLRAPVTSPLAKTSRTSSTPSLTASSLPPCSNASTAGLPANAD*PTQCILH*VI**DFNILSYPVHL*ICNFMQLAMQTLLHNTCEGNIIVLPSMKK*GLITN*YICX

>Cma2007903

CIVFYVSTFPCA*VIHLFLTMVAAADMASSPCQLVQPPPRPLSLDGDRLIDPFNWVQSANALGGSFLDDLKSMAQTYFESKEIKIEGKTLTIAQVAAVARRGEVIVTLDEEAAKERVDESSMWVQNKIMKGCDVYGVTTGYGATSHRRTSQGIELQRELIRFLNAGIFGKNEGNSLPVDASRAAVLVRTNTLMQGYSGIRWQILETIEKLLNANITPKLPLRGTITASGDLVPLSYLAGLLTGRPNSKAVTAEGKVVSGQEALKMVGVEKPFELQPKEGLAIVNGTAVGAGLASMVCFDAHVLGLLSVVTSALFCEVMNGKPEFTDPLTHRLKHHPGQIEAAAAMEWILQGSAFVSDAAKLNGVDALKKPKQDRYALRTSPQWLGPQIEVIRFATQLIQREINSVNDNPIIDVSRDLALHGGNFQGTPIGTAMDNIRLALAAIGKLMFAQFSELVNDFYNNGLPSNLSAGPNPSLDYGLKGGEIAMASYTSELEYLANPVTNHVQSAEQHNQDVNSXWG*YPQGRQRRPSKC*RS*APPTWSGYARRWT*DISKRICRRL*RGWCREWPNRRSRRGKMECCCHRASVKMSCLKWWQMSTSLHMLMIRPAQDTL*CRGCARCWLSTPSGIPLTSEMRPRQS*AGSPSSKTNCVPSCPPSLSMSGQLTTKAARPSPTRSRSVALSHSINLSEPSWAPNCSRAPATSHPDKILKLSLMPSRKERSCSPYLNAWKAGPRLQSNAKCRNNFLP*TRVQCVMVSSFVSIMAFSLHMA*CIPVP*LKFSSYRWFGIFNX

>Cma2008861

QQGTALPIWHDLWMCVRARARVCVVLCRYCAVA*SLHC*RSCRSHRVTSLHSFLTRSKLPAPWQNHAQTICFCKVSFSLLFLCSGSAFLLMMMEQVMPFLGFHSARPFFFSQAMLHR*VMPILGFYFKSQGHIDVV*GSSLARFHWC*GITEARVSCKLGFHSGLKLGSRVPISRRQRLKLGFRARQGFTQFMVAFRPEFHHNQARVSFWLGFSSHPG*GFSILRLWFQPGSHYTQARVSFRQVFVILNPGGQGFSQASFHHTQPRRSGLH*ARHSSQGFTQARARV*YIHTLEFRSVYLTASGEER*HDRAWEVLFLCLLGKEEQV*VAGSLDGVHAVAVSGASEEGENKAGYSS*AGGLVKRSVLH*IAVMAMMHLQQTLKSLCLKTGWCYAVFWKLKRRSRMMLTWEDGYYDYPKSSADTCNSHGNGSLETGGNPLDQGGSGDLEGQIGLAVAKMSYYVYSMGEGIIGRVAFTGKHQWVFHDGENRMEAAPGGILNSQSVPEKYPDGWQNQFAAGIKTIAVIAVPQGVVQLGSTKLITEDLKWIDHIKSVFGALQTAPPPLRSDLLSEGQGGRLAYSMPFMSAASTSTAVRRPITLQQSQCSLRNVEDVLQWRKYGLQASPKGPSNIQGLWSMGGTAFVAGAMPQELQHQQVVKMLAEAQRSKHASTYPGPQSSQMRPTVPHFDLPLNCNQKILSRDEMRSRHPRDSDEKQLRENEVEQCKVSSLMGLMTGGVSTGLHNEFKGNAGLSTCLRGQVDEDLSSNQEDVCLSKRFAMYSMSNTKKDEGILGLSSSNGAGHQKQVDTETPLHMPAHNVGATYMLKQHLPSFNHSSSSSSFLRCDVADKHCNSFSVKDNMANQECSNDVSQTAVASCRLPAKQSTSNFHANDGFAATSKDSVYSIACLNAWDRERLARSESMVSEADPNDCSFGSQSDTSTLDELEKLLASFSREQSWDCSFAIGDELSQALGPAFKKGGIDKTFEELRGLGGDQKIEAIGENLLGKAYVRQDEMVQKWGSSFVNDNLNKTVLGINFPEFKSEPLLDAVVGNASSSYRSVCSNADGSFSDKTMFSNLTSGMPVLNTSKERESKIEACNSSLSGKHMQAAPHHEAVNDVHTTGVVKNAQGHQNFCVSKDSPAKVVLHSWADEGLXSLKSDSQSKKSEELSKAGKKRSRSEGSTRPRPKDRQQIQDRVRELRDIVPDGSKCSIDALLEKTIKHMTFLKSVTRHADQLKKNGELKDDLESGASWALDLGGQDTEYPILVKDLNQPRQMLVEMMCEERGLFLEVADIIRSLGLTILKGVMESQNDKIWAKFVVEANRDVQRVDVLVSLLQRLHLDNNSSSSSMTMASQSAPIRPQRGVDSSSSQTICGIEQSCFSI*PFYERM*QVQVTSSGLCLASGLV*LASEHKALYTTARETQQAGLIARP*AATPSL*QQRRAGCYV*GA*YWPVRLQQFNLKRQASFSVQKCEYCSKHVFNEVGPILGALERKS*HRVYAVLYEAM*VNYEDCTLLPCFSFGRRGLEFVMMPNSTCHRLDQITLHVYLKGFNWHLTPSLVAQS*GHMLWKMYGSKSGECINILGPPWISALGLFLLGVYHVQKL*HYGASMHTCLALLL*SIFSQRALQCQ*NEHRVTYFTWALLLMLERAVCARMSVFFTSMHMCTPX

>Cma2008862

ECSNDVSQMAVASCRLPAKQSTSSSHASDGFAATIKDSVCSIACPNVWDRESLARSESMISEAELNVCSLSTQSNTSTLDELDKFLASFSREKSGNCDCSFAIGDELSEALGPAFRKGGIDKTFEELPPLGGDQKIEANGENLLGKAYVRQDEMVQKWGSSFVNDNLNKTVLGINFPEFKSEPLLDAVVGNASSSYRSVCSNADGSFSDKTMFSNLTSGMPVLNTSKERESKIEACNSSLSGKHMQAAPHHEAVNDVHTTGVVKNAQGHQNFCVSKDSPAKVVLHSWADEGLXSLKSDSQSKKSEELSKAGKKRSRSEGSTRPRPKDRQQIQDRVRELRDIVPDGSKCSIDALLEKTIKHMTFLKSVTRHADQLKKNGELKDDLESGASWALDLGGQDTEYPILVKDLNQPRQMLVEMMCEERGLFLEVADIIRSLGLTILKGVMESQNDKIWAKFVVEANRDVQRVDVLVSLLQRLHLDNNSSSSSMTMASQSAPIRPQRGVDSSSSQTICGIEQSCFSI*PFYERM*QVQVTSSGLCLASGLV*LASEHKALYTTARETQQAGLIARP*AATPSL*QQRRAGCYV*GA*YWPVRLQQFNLKRQASFSVQKCEYCSKHVFNEVGPILGALERKS*HRVYAVLYEAM*VNYEDCTLLPCFSFGRRGLEFVMMPNSTCHRLDQITLHVYLKGFNWHLTPSLVAQS*GHMLWKMYGSKSGECINILGPPWISALGLFLLGVYHVQKL*HYGASMHTCLALLL*SIFSQRALQCQ*NEHRVTYFTWALLLMLERAVCARMSVFFTSMHMCTPX

>Cma2009894

RCSLLLLLWTVGGLPCKLAAPLVVLINPNPIPTTSSSGYLISHLLAL*EKRKSLPRIWLWWHITHTQSLFRFCLKLSLQPWR*PAELPASVCLREVLGFPEQNI*GF*GFHIEXXXXXXXKQRFFRVYLLS*ANFVGFSSLHH*KVHLDYPSSHKQKILGFPSSSKLKIIGFASWPKVKFLSLCIAKDRASDQCCNFFLS*RGKSNTPTC*FSVQTELP*IN*RSTCRPSVSPVLLARVRCRSAAFLESGDGEL*EHA*SDQSSAVLGYM*RYSVILYTFKQHISYLFKAQKPVTALRRRNPNCFEEETLAALGGDSKEGMGVNLPFSLSAPPWKRRAILRRKLAGRASCRAS*LFHKKRGGSVRISPQPCHYRLFSPGLKSL*RGLPASKKQVPAMALHQTLRSLCSKSGWCYAVFWKLKRRSRMVLTWEDGFYEYQSSGICGEINCLETGGNPLNQGSSGHEASYSEGQIGLAVAKMSYCVYFMGEGIIGRVAFTGKHQWFFGGGNNSMLGTAPGNATSTRCMALEKYPSSWNIQFAAGIKTIAVIAVPHGVVQLGSTQMIMENMEWVDQVKSAFGALQNVPGTFLSDLVSEGQGRKAVISPLEMPAGSPSVRVGRFNPGQFSSGSIGDVRGTGLQSLVYGSQGRSSAVKGSSNMQKPATAVEAAEALRAMPQKSEMLQVIEMGMGAKGLRPAINHSGSQISEAFSAVPQVDVLSVINQPISFLSEDMRIGHYGQTRDTQLKNSHVQPSLEAFGRGLGPEADADVSTCLHSDIFPEVSCNQEQSSINAMLVFNRILDSKSCLTSVPISGNGPQQPEVAPSSSLAPYGFNSGVSVIASNQTETACNLAGAGQDSLSSFAGKTVGDEEYLAGSTFGDVPQPQITYPDGECLNPGSKDPVPHSGRVGNKNLAAGGWGDFENYLASVAQEQSRKWGSFAVGDELSQALGPAFKQGHDKEMWEEVLLPLGTDQKKVAVEQQTLGRTCRLSDQSVADKWGSSFVKDYLHFLESKSEPLLDAVVANASSSHHSISTIADDSFSCRTFSGLDSEISALNTTKETGSKRARCLSSQDCLSQGYMQAVPSDCEASVDTQTLTKVNNDPMNFNMSTNGSPLKTVLSSWAEDMQSLKSDSTQTSQSKNPEDLTKATRKRARPGESTRPRPKDRQQIQDRVRELREIVPNGSKCSIDALLEKTIKHMHFLKNVTLHADNLKKNGELKDDSESGASWALDVGGEETGCPILVKNLNQPRQLLVEMFCEEKGHFLEIADIIRSLGLAILKGVMESRSGKIWARFVVEANRDVRRVDIMMSLMQLLHVNNNSSSSMTMASQSGPFRSQQGVDSSSSQTFCGFQQPSLHA*LVYSLS*IPLPSPDFLRPQLVEVLRVGLKFSWGLP*LASERFYLNSLTIKQVLIHQFLLLITYSLIREGKGCHG*SLCPSHMLLRIGVMAATLHILLLQILVGLMAPLSTMCASIPPVLMSTSSPFCSSRNTPGFAMYNML*SKCRYYNVSSRWLLIMVCSPDYLQRVFSMSKKWX

>Cma2009895

TPSPTSSRRF*GFQAEDLGCPQQKISGFLSRRFRVSIFGLS*RFRVSRASILIRGEIL*GLSFELSKFF*GFPLYTTRKSILIIHLDVSRKF*GFHLHLS*SL*GLHLDLS*NQGKKIQVTSAVFLPKPAISSCLEGENPTAFLASFLCKRSFNKNNWLFRPSVSPRPSR*GLVQICSFLGFGDGEL*EHARSDQSSAVLGYM*RYSVILYTFKQHISYLFKAQKPVTALRRRNPNCFEEETLAALGGDSKEGMGVNLPFSLSAPPWKRRAILRRKLAGRASCRAS*LFHKKRGGSVRISPQPCHYRLFSPGLKSL*RGLPASKKQVPAMALHQTLRSLCSKSGWCYAVFWKLKRRSRMVLTWEDGFYEYQSSGICGEINCLETGGNPLNQGSSGHEASYSEGQIGLAVAKMSYCVYFMGEGIIGRVAFTGKHQWFFGGGNNSMLGTAPGNATSTRCMALEKYPSSWNIQFAAGIKTIAVIAVPHGVVQLGSTQMIMENMEWVDQVKSAFGALQNVPGTFLSDLVSEGQGRKAVISPLEMPAGSPSVRVGRFNPGQFSSGSIGDVRGTGLQSLVYGSQGRSSAVKGSSNMQKPATAVEAAEALRAMPQKSEMLQVIEMGMGAKGLRPAINHSGSQISEAFSAVPQVDVLSVINQPISFLSEDMRIGHYGQTRDTQLKNSHVQPSLEAFGRGLGPEADADVSTCLHSDIFPEVSCNQEQSSINAMLVFNRILDSKSCLTSVPISGNGPQQPEVAPSSSLAPYGFNSGVSVIASNQTETACNLAGAGQDSLSSFAGKTVGDEEYLAGSTFGDVPQPQITYPDGECLNPGSKDPVPHSGRVGNKNLAAGGWGDFENYLASVAQEQSRKWGSFAVGDELSQALGPAFKQGHDKEMWEEVLLPLGTDQKKVAVEQQTLGRTCRLSDQSVADKWGSSFVKDYLHFLESKSEPLLDAVVANASSSHHSISTIADDSFSCRTFSGLDSEISALNTTKETGSKRARCLSSQDCLSQGYMQASDCEASVDTQTLTKVNNDPMNFNMSTNGSPLKTVLSSWAEDMQSLKSDSTQTSQSKNPEDLTKATRKRARPGESTRPRPKDRQQIQDRVRELREIVPNGSKCSIDALLEKTIKHMHFLKNVTLHADNLKKNGELKDDSESGASWALDVGGEETGCPILVKNLNQPRQLLVEMFCEEKGHFLEIADIIRSLGLAILKGVMESRSGKIWARFVVEANRDVRRVDIMMSLMQLLHVNNNSSSSMTMASQSGPFRSQQGVDSSSSQTVCGFQQSSRHA*PFYLLPLLPSPLTF*GPIL*RCCEWVLNFLGACHDWHQS*RHEATLHILLLQIVVGLIGTTFTNVRLHSS*TLLLFVPPSKQAQEKGLGFEMYNIQRSKCRYYNVSSRCLLTMVCSHDNIYKEFSEENN*SEGLGLGCKEVWDSVRFPSVWLA*T*AMQIILVTDAKL*IALSKIYRAL*RWATPLCSMVKFYLLLA**LVAPRAIADYSMLEHSRAPEGRALKIQMSCEPAMWAKQLLYLERMNIHRCKWCLSLGF

>Cma2009896

RCSLLLLLWTVGGLPCKLAAPLVVLINPNPIPTTSSSGYLISHLLAL*EKRKSLPRIWLWWHITHTQSLFRFCLKLSLQPWR*PAELPASVCLREVLGFPEQNI*GF*GFHIEXXXXXXXKQRFFRVYLLS*ANFVGFSSLHH*KVHLDYPSSHKQKILGFPSSSKLKIIGFASWPKVKFLSLCIAKDRASDQCCNFFLS*RGKSNTPTC*FSVQTELP*IN*RSTCRPSVSPVLLARVRCRSAAFLESGDGEL*EHA*SDQSSAVLGYM*RYSVILYTFKQHISYLFKAQKPVTALRRRNPNCFEEETLAALGGDSKEGMGVNLPFSLSAPPWKRRAILRRKLAGRASCRAS*LFHKKRGGSVRISPQPCHYRLFSPGLKSL*RGLPASKKQVPAMALHQTLRSLCSKSGWCYAVFWKLKRRSRMVLTWEDGFYEYQSSGICGEINCLETGGNPLNQGSSGHEASYSEGQIGLAVAKMSYCVYFMGEGIIGRVAFTGKHQWFFGGGNNSMLGTAPGNATSTRCMALEKYPSSWNIQFAAGIKTIAVIAVPHGVVQLGSTQMIMENMEWVDQVKSAFGALQNVPGTFLSDLVSEGQGRKAVISPLEMPAGSPSVRVGRFNPGQFSSGSIGDVRGTGLQSLVYGSQGRSSAVKGSSNMQKPATAVEAAEALRAMPQKSEMLQVIEMGMGAKGLRPAINHSGSQISEAFSAVPQVDVLSVINQPISFLSEDMRIGHYGQTRDTQLKNSHVQPSLEAFGRGLGPEADADVSTCLHSDIFPEVSCNQEQSSINAMLVFNRILDSKSCLTSVPISGNGPQQPEVAPSSSLAPYGFNSGVSVIASNQTETACNLAGAGQDSLSSFAGKTVGDEEYLAGSTFGDVPQPQITYPDGECLNPGSKDPVPHSGRVGNKNLAAGGWGDFENYLASVAQEQSRKWGSFAVGDELSQALGPAFKQGHDKEMWEEVLLPLGTDQKKVAVEQQTLGRTCRLSDQSVADKWGSSFVKDYLHFLESKSEPLLDAVVANASSSHHSISTIADDSFSCRTFSGLDSEISALNTTKETGSKRARCLSSQDCLSQGYMQAVPADCEASVNTQTLSKVNNDPLNFNMSTNG

>Cma2009897

RCSLLLLLWTVGGLPCKLAAPLVVLINPNPIPTTSSSGYLISHLLAL*EKRKSLPRIWLWWHITHTQSLFRFCLKLSLQPWR*PAELPASVCLREVLGFPEQNI*GF*GFHIEXXXXXXXKQRFFRVYLLS*ANFVGFSSLHH*KVHLDYPSSHKQKILGFPSSSKLKIIGFASWPKVKFLSLCIAKDRASDQCCNFFLS*RGKSNTPTC*FSVQTELP*IN*RSTCRPSVSPVLLARVRCRSAAFLESGDGEL*EHA*SDQSSAVLGYM*RYSVILYTFKQHISYLFKAQKPVTALRRRNPNCFEEETLAALGGDSKEGMGVNLPFSLSAPPWKRRAILRRKLAGRASCRAS*LFHKKRGGSVRISPQPCHYRLFSPGLKSL*RGLPASKKQVPAMALHQTLRSLCSKSGWCYAVFWKLKRRSRMVLTWEDGFYEYQSSGICGEINCLETGGNPLNQGSSGHEASYSEGQIGLAVAKMSYCVYFMGEGIIGRVAFTGKHQWFFGGGNNSMLGTAPGNATSTRCMALEKYPSSWNIQFAAGIKTIAVIAVPHGVVQLGSTQMIMENMEWVDQVKSAFGALQNVPGTFLSDLVSEGQGRKAVISPLEMPAGSPSVRVGRFNPGQFSSGSIGDVRGTGLQSLVYGSQGRSSAVKGSSNMQKPATAVEAAEALRAMPQKSEMLQVIEMGMGAKGLRPAINHSGSQISEAFSAVPQVDVLSVINQPISFLSEDMRIGHYGQTRDTQLKNSHVQPSLEAFGRGLGPEADADVSTCLHSDIFPEVSCNQEQSSINAMLVFNRILDSKSCLTSVPISGNGPQQPEVAPSSSLAPYGFNSGVSVIASNQTETACNLAGAGQDSLSSFAGKTVGDEEYLAGSTFGDVPQPQITYPDGECLNPGSKDPVPHSGRVGNKNLAAGGWGDFENYLASVAQEQSRKWGSFAVGDELSQALGPAFKQGHDKEMWEEVLLPLGTDQKKVAVEQQTLGRTCRLSDQSVADKWGSSFVKDYLHFLESKSEPLLDAVVANASSTHHSISTIADDSFSCRTFSGLDGEMSALNTIKETGSKRARCLSSQDCLSQGYMQAVPADCEASVNTQTLSKVNNDPLNFNMSTNG

>Cma2010579

FMGEGIIGRVALTGKHQWVFGGGDNNMLGTSPGNATRAQCVLEKYPSGWDIQFSAGIKTIAVIAVPQGVVQLGSTQMIMENMEWLDQVKSAFGALQNVPGAFLSDLVSEGQGGKAVISPSGMHAGSPSARVGRFNPGQFSSRSIRDIHGTGLQSLVYGSHGRSGTVKGSSNMQKSSTVVEAAEALQAMPHKSEMLQVIX

>Cma2010594

GDCDAGVSHLLSHQETLSFAVALSLAVAVACLSFRQRISQKMMMLSNLLFKHRAESP*LQAWRTRWILNLLPSLLIQVIKGLYAFQ*WRNRRSLLNTTALARLRDSRLGLFSPISKVIMGWDDDQALLEAFIGSQYEEALWVDSQQSAIAPASAATATASAAVGSNSLEDNLQQRLQSVVENFPQSWTYAIFWQLTHSSHGQQVLGWGDGYFNPKEGEQSASTRHVSDADLQLRRRILSELQTLIGQSSGDDPITAGLDFLDADVTETECFYLVSMWYSFPLGSGTPGLAFATSRYMYKWLTGANQTQNRDCPRAELALRFGIRTMLCVPTLHGVVELGSTELINENVNFVQFVKNSFSSSEDPSNQQPFFLQADLSCFPSASGHGSFPAPVLNNSFPTSSNALSSLSYVSSGLSSSQISRASDPSKLQTDSCIAERRPPTTSGAEAEDLFSKDLPFPDMMFMGIGDKPMPMTWFSPLHTFPGVDEDMLVAMSSDLKSESSVLTQRSEVGDSAKYFESKRAVTADMQSSNYAAKCPDVQSYSNFPPGNLPDLLWKXASDVWQYCCQGGCWVRRRGSKGPRGTELQQFCKSSSTSELWVVKQVAGST*SYSGFQSCRGRRSEVPKRSWSAGSGIGLQVWPQWPSRNAEIRSRHEVRREFEL*SACQELWGCRNFRPPIQRREVQRGFATGRQPSAPNKWSLALQCGIRAFGCGGLF*GCRMQRRSPGEEAPQAGQEARQWQGGAIESCG

>Cma2010595

GDCDAGVSHLLSHQETLSFAVALSLAVAVACLSFRQRISQKMMMLSNLLFKHRAESP*LQAWRTRWILNLLPSLLIQVIKGLYAFQ*WRNRRSLLNTTALARLRDSRLGLFSPISKVIMGWDDDQALLEAFIGSQYEEALWVDSQQSAIAPASAATATASAAVGSNSLEDNLQQRLQSVVENFPQSWTYAIFWQLTHSSHGQQVLGWGDGYFNPKEGEQSASTRHVSDADLQLRRRILSELQTLIGQSSGDDPITAGLDFLDADVTETECFYLVSMWYSFPLGSGTPGLAFATSRYMWLTGADQTQNRDCPRAELAHRFGIRTMLCVPTLHGVVELGSTELINENSNFVQFVKNSFSPSEDPSNQQPFFLQADLSCFPSASHGFPSPVPNSFSTSFNTLSSSTYVSGLASSQISRTSAPSKMQTDSFIAERRPPTSGTEVEVSQGGMFSKDLPDMMFMGNGDKSMPMTWFSPLQKFPGVDEDLLVTMSSDLKSESSVLTQRSEVGDSAKYSETKRASITADLQSSNYAAX

>Cma2010596

GDCDAGVSHLLSHQETLSFAVALSLAVAVACLSFRQRISQKMMMLSNLLFKHRAESP*LQAWRTRWILNLLPSLLIQVIKGLYAFQ*WRNRRSLLNTTALARLRDSRLGLFSPISKVIMGWDDDQALLEAFIGSQYEEALWVDSQQSAIAPASAATATASAAVGSNSLEDNLQQRLQSVVENFPQSWTYAIFWQLTHSSHGQQVLGWGDGYFNPKEGEQSASTRHVSDADLQLRRRILSELQTLIGQSSGDDPITAGLDFLDADVTETECFYLVSMWYSFPLGSGTPGLAFATSRYMYKWLTGANQTQNRDCPRAELALRFGIRTMLCVPTLHGVVELGSTELINENVNFVQFVKNSFSSSEDPSNQQPFFLQADLSCFPSASGHGSFPAPVLNNSFPTSSNALSSLSYVSSGLSSSQISRASDPSKLQTDSCIAERRPPTTSGAEAEDLFSKDLPFPDMMFMGIGDKPMPMTWFSPLHTFPGVDEDMLVAMSSDLKSESSVLTQRSEVGDSAKYFESKRAVTADMQSSNYAAKCPDVQSYSNFPPGNLPDLLWKXASDVWQYCCQGGCWVRRRGSKGPRGTELQQFCKSSSTSELWVVKQVARST*SYSGFQSCRGQRS*VCAKRSWYAGSGIGLQLCIKGPSRNAELRSRHEALREFEL*SACQELWGCKNARNFRQPVQRREVQREFATGRHPSASNEWSHTLQCGIRAFRCGGLF*GC*MQ*GSPGEEASEEGQEACQWKGGAIESCGGGAPEEREAQPALLCLEGRGTECVEDGQGIIAQRRGIIHPRAQMQATGS*DRKQESNCSVEDQQEGSLFSC*SW*PFHLDSRHYQKQGSGSFPSRWRQSDCEGSFSGRQGSDDQSRQPERKPPGGQGYGGLARSAASSSSCKCVHRPRHNSSVHTSENERPVLLHRRPAHSCDFWKRRLDSG**KILAILHACLHL*PPFT*DLNTAYYNPCT*IYVLLDYCVLR*NX

>Cma2011585

EYRYCAG*TD*LIDRWMDRLPICFVFLGLLIALICQVSIETQNL*ALLGPHNYSPQKKSC**IIIKRCLRKAWRS*PLLACTHNHCMLKRSFKYWR*CFFSAGDGQGETWAFFLGRLVRLHWPLFDRSEAFFDRFFFVRKAMSFCVPDWDGNGFLGDFSSNTRGKWEAARPISMESPFVPDDEFEELCWEDGQLMLVMPSQNNRPPHKQLSSWAASVSPLNKPVSRDQVDASLYMGDIATNHDETLDAVVDDGDGSRVAGFPPAISDDEMIAWLQYPLDDELRVDDSDPVLHSGHHAVSYLPSLPFQKSDSVHRNIGAMQRERPSIGITKGKQAATTTNTDLAQAIARRGSQTESFSASFGNSTSALTTDGATSDPKPGTQYLEQTIFSSKAASLLPPLVGSPYCSQQSDPQKILQSISRPVTDMKPATGVPNWPASLKHVKHIDRGNVEVCMSTDSSIESTITGRSLQEPLGDRSGGLTGPNVSHCMTATTSTLHQLLSVKGPDAAACIDDFAVCSETVVADIPSGSKLFCSADANQTVEKGSGAFESTDTSCAAGSGNTTATNGGKVDSNGSYGKRKVKDLDDSESQSEDLGKQPPLSQATSTKRGRAAEVHNNSERKRRERINEKLKALQELIPNANKTDKASMLGEAIEYLKALQSHILVMSYRTGITIPPMLISHGLPHPHVPPHAGIGFGAGMGVKTGVMDMSAATSTCSMPKALALGCLIPSVTIPALSTPATNVATLPQNGLSRPVVGYSTNYFPFSTFGAQPFLVHTNQLQNPPATTSHKTPSLHRHTHQPSQLLSTDMFNPSMQEWQQQQQHMHKQ**HMQQLGQHCMVQ*RCIAAGRVFFLMETDISKPQKVARRSTISMARCRTVDHLLLGQSSTCKIEYVVKP*YLRRPESVSXTHTHTHTHTGHICFVPLEGFCCIIREERATFVVLLAAL*SPCSSN*E*GALKSVLETLKTVFKFFRYKQCLEKDSTNTIFEMMACQCGNDIGIGMYQRRFHVKGIVMNSL

>Cma2011586

EYRYCAG*TD*LIDRWMDRLPICFVFLGLLIALICQVSIETQNL*ALLGPHNYSPQKKSC**IIIKRCLRKAWRS*PLLACTHNHCMLKRSFKYWR*CFFSAGDGQGETWAFFLGRLVRLHWPLFDRSEAFFDRFFFVSKAMSFCVPDWDGNGFLGDFSSNTRGKWEAARPISMESPFVPDDEFEELCWEDGQLMLVMPSQNNRPPHKQLSSWAASVSPLNKPVSRDQVDASLYMGDIATNHDETLDAVVDDGDGSRVAGFPPAISDDEMIAWLQYPLDDELRVDDSDPVLHSGHHAVSYLPSLPFQKSDSVHRNIGAMQRERPSIGITKGKQAATTTNTDLAQAIARRGSQTESFSASFGNSTSALTTDGATSDPKPGTQYLEQTIFSSKAASLLPPLVGSPYCSQQSDPQKILQSISRPVTDMKPATGVPNWPASLKHVKHIDRGNVEVCMSTDSSIESTITGRSLQEPLGDRSGGLTGPNVSHCMTATTSTLHQLLSVKGPDAAACIDDFAVCSETVVADIPSGSKLFCSADANQTVEKGSGAFESTDTSCAAGSGNTTATNGGKVDSNGSYGKRKVKDLDDSESQSEDLGKQPPLSQATSTKRGRAAEVHNNSERKRRERINEKLKALQELIPNANKTDKASMLGEAIEYLKALQSHILVMSYRTGITIPPMLISHGLPHPHVPPHAGIGFGAGMGVKTGVMDMSAATSTCSMPKALALGCLIPSVTIPALSTPATNVATLPQNGLSRPVVGYSTNYFPFSTFGAQPFLVHTNQLQNPPATTSHKTPSLHRHTHQPSQVCRQKIKACMMPYILGHAISILYVERV*SMLLVHGCVA*R*RVWPPRVRTWVQYPPVDTGCWTHYS*IIYIPQNIKM*KIHAS*TIVLCSAFLQLLSTDMFNPSMQEWQQQQQHMHKQ**HMQQLGQHCMVQ*RCIAAGRVFFLMETDISKPQKVARRSTISMARCRTVDHLLLGQSSTCKIEYVVKP*YLRRPESVS*HTHTHTHTHRAHLFCAIGGLLLYYKRGKGNLCSVTGSFIISLFF*LRVRSLEICIGDSQNCF*IL*VQAMPRKGFHKHDLRNDGLPMWQRHRHWHVPKAISRQRNCNEFP

>Cma2015959

TDTEWFFLVSMMCSFPIGTGTPGQAFASSQYAWLNGADQLRGRDCMRAELAQRFGIRTILCVPTQNGVVELGSTEVIGEDLAFLNIIRHSFDPLLSNMDMPYFSSPMRSFTDVNPALVAEFGYLNNSGNEVSAGDSGQTKMSSSGMMMQDLQLGATVEKAGQFLGNGTHGDPLHSLLPQDSPYSDMMFPGNESERLLLPMIWQQSSFSQPAVAASEDGRALTTANQSHDPQLLQKAEFEDVTKRRGMSPDFSHYDHYVKQLKDDSQASGFKARECHSYKETTKVESSSEFMKPVAVQNYSHSMVGEMQSQSQTLPQNYTPFVKTAHIDNFNQFGMSTVPQTYPHHSVKVTEMQSHSQMSTMAAGPQIYPNGIKTAEMQSHSQVGMTIVPQSFTKTSEVQSYSQPVRAPETESHFDGGKAPDLFNHDHFEKETLGTLSHSQAAPASAIKTSDSARKSPDHIVPMRSHVQSYNQTSKLWGLDVQTYSLKPADSTSSYQQDVKAVELLTFDKRSTELNKQSSDEKEGGLLQLDESVAAQVTGAVRSSVESEHSDVEASFKEADCSQSVVEKKPRKRGRKPANGREEPLNHVEAERQRREKMNQRFYALRAVVPNVSRMDKASLLGDATSYIEELRAKVHGLEIEKKKLLLRLDGGKADRMMSHGGSFKNSGQGDPPARTSPTDRKSMNSLACPHGKVAINVHFLQGREALIQVESSRENFPVAKLMLALQELQLEVHHATAAVVQGMLFQKIIVIMKTPSYITEQQLTAALSTRAVDCNCC**YCLIIMMGQDSLSILGWSGHIILKLDGWHKCVKALRGWQLLLLVIVTVVRAPSHGSPMVLIRPCQP*AL*WGGTIM*KLGECC*RRKVKSSVHY*IMSVGVIFPRIYHPSCEI*PHHQGPLQDLFSDRPCHNSMFCALYYCIISG*AVIICLFKHV*YMAIHVFSIR**TLELGYTTSGSVL*GTCSFQSLQACRLSGH**VVYVDYHMIAQKSHDSCFYMFLHVQRA

>Cma2015960

TDTEWFFLVSMMCSFPIGTGTPGQAFASSQYAWLNGADQLRGRDCMRAELAQRFGIRTILCVPTQNGVVELGSTEVIGEDLAFLNIIRHSFDPLLSNMDMPYFSSPMRSFTDVNPALVAEFGYLNNSGNEVSAGDSGQTKMSSSGMMMQDLQLGATVEKAGQFLGNGTHGDPLHSLLPQDSPYSDMMFPGNESERLLLPMIWQQSSFSQPAVAASEDGRALTTANQSHDPQLLQKAEFEDVTKRRGMSPDFSHYDHYVKQLKDDSQASGFKARECHSYKETTKVESSSEFMKPVAVQNYSHSMVGEMQSQSQTLPQNYTPFVKTAHIDNFNQFGMSTVPQTYPHHSVKVTEMQSHSQMSTMAAGPQIYPNGIKTAEMQSHSQVGMSAVCQSLTKTSEVQSFSQPVRASETKSNMDGGKAPDFFNLNRFEKKTLGTLSHSQEAQTSAIKASDSARKSPEHKVPTMSHVQSYKQTSKLWGLDVQTYSLKPADSTSSYQQDVKAVELLTFDKRSTELNKQSSDEKEGGLLQLDESVAAQVTGAVRSSVESEHSDVEASFKEADCSQSVVEKKPRKRGRKPANGREEPLNHVEAERQRREKMNQRFYALRAVVPNVSRMDKASLLGDATSYIEELRAKVHGLEIEKKKLLLRLDGGKADRMMSHGGSFKNSGQGDPPARTSPTDRKSMNSLACPHGKVAINVHFLQGREALIQVESSRENFPVAKLMLALQELQLEVHHATAAVVQGMLFQKIIVIMKTPSYITEQQLTAALSTRAVDCNCC**YCLIIMMGQDSLSILGWSGHIILKLDGWHKCVKALRGWQLLLLVIVTVVRAPSHGSPMVLIRPCQP*AL*WGGTIM*KLGECC*RRKVKSSVHY*IMSVGVIFPRIYHPSCEI*PHHQGPLQDLFSDRPCHNSMFCALYYCIISG*AVIICLFKHV*YMAIHVFSIR**TLELGYTTSGSVL*GTCSFQSLQACRLSGH**VVYVDYHMIAQKSHDSCFYMFLHVQRA

>Cma2015961

TDTEWFFLVSMMCSFPIGTGTPGQAFASSQYAWLNGADQLRGRDCMRAELAQRFGIRTILCVPTQNGVVELGSTEVIGEDLAFLNIIRHSFDPLLSNMDMSYFSSPMRSFTEVNPPLVGTDFGYLNSSGNELSAGDSGQSQTKMSSSG

>Cma2015962

TDTEWFFLVSMMCSFPIGTGTPGQAFASSQYAWLNGADQLRGRDCMRAELAQRFGIRTILCVPTQNGVVELGSTEVIGEDLAFLNIIRHSFDPLLSNMDMPYFSSPMRSFTDVNPALVAEFGYLNNSGNEVSAGDSGQTKMSSSGMMMQDLQLGATVEKAGQFLGNGTHGDPLHSLLPQDSPYSDMMFPGNESERLLLPMIWQQSSFSQPAVAASEDGRALTTANQSHDPQLLQKAEFEDVTKRRGMSPDFSHYDHYVKQLKDDSQASGFKARECHSYKETTKVESSSEFMKPVAVQNYSHSMVGEMQSQSQTLPQNYTPFVKTAHIDNFNQFGMSTVPQTYPHHSVKVTEMQSHSQMSTMAAGPQIYPNGIKTAEMQSHSQVGMTIVPQSFTKTSEVQSYSQPVRAPETESHFDGGKAPDLFNHDHFEKETLGTLSHSQAAPASAIKTSDSARKSPDHIVPMRSHVQSYNQTSKLWGLDVQTYSLKPADSTSSYQQDVKAVELLTFDKRSTELNKQSSDEKEGGLLQLDESVAAQVTGAVRSSVESEHSDVEASFKEADCSQSVVEKKPRKRGRKPANGREEPLNHVEAERQRREKMNQRFYALRAVVPNVSRMDKASLLGDATSYIEELRAKVHGLEIEKKKLLLRLDGGKADRMMSHGGSFKNSGQGDPPARTSPTDRKSMNSLACPHGKVAINVHFLQGREALIQVESSRENFPVAKLMLALQELQLEVHHATAAVVQGMLFQKIIVIMKTPSYITEQQLTAALSTRAVDCNCC**YCLIIITVAR*LIHLALVRPHHTEA*WAAQMC*GLERLGMVTVARLPSHGAPTVLIRPCQPRAL*QGWHHYVKALRVLLKKKGEVSCTLQSYER*CHFSSLAAKSDPIIEAHFRICCQRYLFISSRHILCMP*QHVLCHSHILLYYIRMSSHYRVLCLFKQVRTHV*YMAIHVCSIR**ILELGYKAKAYVML*LCCLRHVLNSGKCSRVDNGKMGNSPSGKPPLNLEFKAI*ETQCADX

>Cma2016024

SRLLIWMLQESFKLICICDDKFMAIDQSICTSASSGGGVRAMRKMLGDSTGDRCSCCRKKREESSEDDGAQLGGKARQRWASSEVRKALGWHAGANLILDPSGSAMANSEISAVKIMSSARV*ERRRQWRPWLGFRMP*PIISVTHTSWKVGGRGCLPSRNAVCVGCTIHTMTCNCAATVLLAEGNNISSCWSHGQAPPSFCCSRKANSLIAIHTNSTAGSKSSFAKVHTSINPTTVAENPQTKSEQAQRSGRETRAQLLQRCISTTTLLEAKHIHEHLVKTELVPDIFLENTVINMYVRCGSIAHAGQVFENMPKRNVFSWTLILAGYEKLGRPDEAVKLFWQMERQGVSADNVTFVSVLKACTRVASLEQGKKIHSHIIKCGIKSNGFVFNALVDMYAKCGGMVDAVHVFDQMDEKDVVSWNAIISGCVQSGHHTIALNFFQQMKSEGLKPDEVTLVCILKSCGSLMGVEEGGQMHVLILESGYESNPYIGSTLVDFHAKSGHMDIACQVFNRMLERDVVSWNAMIAGYANHGHGEDAFWAYRGLLQEGVKPDEVTFMNILNACVGPGFLEHGRRVHAHAIEVGKELDVSVGNSLIDMYAKCQSLEDAVLMFGKLPNRNTVSWNTMITGYSQVGCSEEAVKLFDIMQKEHVKPNEVTFVSGLHACSLLAALEEGKTIHAQAVYAGHESSTRVGNTLIDMYSKCGRLEDALQVFDTLPNRDVVSWNVIITGYVQHGHSLTAIEMLQEMQQEGLKPNEVSFINGISACAGMASLQHGLLIHADAVKAGYEADIFVGSTMVDMYVKCESMHDAWDVFDKLPEKVSASWNTIIAGYGDNGGEEEALRLFYIMQTDEVIPDDFTFVSILNACASLCALEEGKCIYCQYVKAGYSVNIFVGSALVDMYAKCGHVEDARHVFDEVPDQDLFLWTVMIAGYAQHGHHQEVFELWEKMQREGVKPDGIIYVCVLSACSHVGLVDKGLQLFNSMTQDHGISPTIDHFACFVDLLGRAGCLNEAEHFIKKLPVQPDDAVWRTLLGACRSHGNLELAELASKRLLKLKPNDSAAYVYLSNTYAAAGSLST*GFWTRCTPLQTPPKPL*LMLC*TFSNKSY*N*QAHKRLVACYCRPFLKLPS**AIAGPKATCMYIPH*LVILPNQLLLDWRLLL*S*S*LCVAWSX

>Cma2017862

LKACSIIGAADQGRKLHALAVFRGLDSNVFVGNTLIDMYCKFGSLNDAFTVFESLASRDVITWNALIADYSLHGHSDEAFHLYKQMQLEGFKPDLVTYISLLKACSVMSALQEGRELHNQIIQIGLDLDLSISNTLIDMYSKCGNLDDACMVFVRLSKQDVVTWNALMSGYVQHGHGPEALRLFQKMQHEGMEPNQVTLVIMLKACSSVAALGQX

>Cma2021686

DSIEPDKFIFSSILKACSSVEVLDEGKLVHDHITRSGLESDVIVGSALIDMYAKCKSMQDACTVFYSLPDKNVVSWNALIAGYAQHGHGRPALQLYEKMKGNGIKANKVTFLCVLKACGCIIALEKGRAIHNKLIKNGFESDVFIQX

>Cma2023385

LEVLSDGYIRNVFSSELDEFDDFIASLMREQEKQSGGVSSRPAGPTVLEEGVSEALKKSPQKASFWDPLRMQGPLEPTLAKEQTIILSSKPKEETTVSCKKADSTDLHWDRCAKMPKLESPGTNPETDTKMGYQQQAVVLPASKQDEPLKVGKKRARPGENARPRPKDRQQIQDRVHELRDIIPSSAKCSIDALLEKTIKYMNFLQTVTQDAGKWKQSSKLQDDEGNSGSGTIEGGASWAVELGGQGRGCX

>Cma2024951

KPRQLLVEMLCEEKGIFLEIAESIRGLGLTILKGVMEVRNNKIWARFVVEAVQDVHRVQVLWSLMQLLESNTRTSSNVTNPPCG*RTPLII*QLKPLYPHTSNAFPRVEALQQLNILRCYEWVI*LAMISIVYDDLSPLNGL*HNAFFFCFCQPFFLYYVIHLQGSCCNHIHGTWFAFHKPYIKGVCX

>Cma2024952

KPRQLLVEMLCEEKGIFLEIAESIRGLGLTILKGVMEVRNNKIWARFVVEAVQDVHRVQVLWSLMQLLESNTRTSSNVTNPPCG*RTPLII*QLKPLYPHTSNAFPRREALQQLNSRRHIITDILRCCEWVI*PAMISIVYDGLSPLNVL*HFVCQLFFLYKVIHLQGSHISGFCLPHTSFFEDSVALLTTLTLLDYNHTYFKLH

>Cma2027618

NLLCEDYFEDMQSLTSLFEDSELSVRNLSAAGDCVENLLNTELDDFDDFLASLTREEEKESDMFSELGTGFASMGTLTSNQNVGSLATEDVGVFAKREKDHANELRGSNAASGHVDVTQAKPEPHLATLSDDRVASNWSAIEETLLCKEVATDRFVKTRTMKAVSFPCLRHTDGDAEMGCQDAVLPASRQDGTVRKGQKRSWPGEASRPRPKDRQQIQNHVRELRDIIPNAAKCGIDTLLEKTVEYMNLLQMVSHDTGKWTQRSKLQNDDCGNVIGHGPDGTAVDASWAVELGGQGRGCPIIVENLNQPRQMLVEMLCEEKGFFLQIADSIRRLGLTILKGVVEARDDKIWARFVVEAVQDVHRVQVLWSLMQLLEPNTRCSSDITNPPHDLKRIH*LPA*PLNF*KPCAVCFGGTKHHGTLTLGAYNPLLEPKVL*VGLFDLQ**ASSAYVLGLCVVLPPH*GCDIMYSFPW*P*TGYAFTKKLL*WHSLDLVKRIHYVTTX

>Cma2032445

KKRKGDRKRRGSQPRPKDRQQIQDCVRELRDIIPNGSKCSTDSLLEKAIKHMLFLRRVLHHSDLLNQSHDFQIEKRRDASGAFEFG

>Cma2041653

QMLVEMMCEERGVFLEIADIIRSLGLAILKGVMES

>Cma2053128

KKSEELSKVGKKRSRSGESTRPRPKDRQQIQDRVRELX

>Cma2080164

PTGWQDQFAAGIKTIAVIAVPQGVVQLGSTQTISEDLKFVGHVK

>Cma2084136

VGEGIIGRVAFTNKHQWILSSSDTVATAEGTPSSKCSGRTQPEKYP

>Cma20G01660

MLACPGAARGERRTVSDDAPSPRRDQRTVSRRNIRRVGPAARASPSRSMHRGVAIPATVQPRSYLCIVLRGARMVEPSVVWWCLVEALKRRLLRQQRWQLWLLVVGLFLLCTIRPRMQSVLDAASKEPGSLLDPNPMYIPAWDAQTGAVLTAFDPQRHLLPFHFIACSSDDWGRIADSVPLFRDSHARQEFLAVASEELKRSLRQTDWSMATVETVRDLQRLHEFLLRLQEAAGEPRQRFVLSPMWVVGGPDIPAMQNQLDIKREGSTGPLRERSSDAQLWVDARVQRAAATPERRGRMRLVQRSPRERWRNSRQRLHETHAEYPQPGVGQRSTDPVGETRRAGVTLQRPDSLKRFTGQSSTWGLLTRPVVPTDAQAVHAAPTGFPYRSMYMCRLSRNGASILPGRDLIESIEVAEWYRKLWYDRVWAPQFHGSAHIHPHRWLEALERTRPLRRQRSTARTRIIRSAYDSLREGYVYAGNITALRSEFADGAWSPASLRKGIETFAGFWGYRPRVMSSPHNTWTPSFVQHLVASHAVFGIDAGTGQCATLVIPNATQRSVSCIDREPWDTFAGRAARLDARMPLIQERIAALRPGFTSLAWHAQNALSATTSPRVAAAHLEHLETFVHWVQRQNHTVFVTSNELHQIRLRGWSLEVWPDAFVYRNFLPYDILVLVPCLEDMFSGASPWEAAPLRVVRLDQRLIRAPISQSSPELTQCRPWSALMADSQHGHALPRSLRYIQRQKQKRKQLAATPAFRGCRCQRIRLPSFMDDGVLLLPGNSHWIEVRRLSHE

>Cma2105

LEKTIKHMLFLKSVSLHAGQFKKNGELKDDLQSDASWALDLGGQDTKYPILVKDLDQPRQMLVX

>Cma2112385

LEKTIKHMLFLQNVTSHADNLKKNGELKDDVESGASWALELGGEETGCPPILVKNLNQPRQLVVEMFCEEX

>Cma2118195

EAISKS**AQSMDSSGGYPVVSYLLQQTLRRLCTDSDWIYAVFWRILPRNYPPPKWESESSVLDRSKGNKRNWILVWX

>Cma2127205

QRGYATLYGHIAGKGMQEMVS**FGSSEKEGERLTFQQ*KSLLSAMASMLQKTLKGLCYDTGWCYAVFWRLKRQMRTVLTWEDGFHEPFKSSEX

>Cma2133381

GLHAFDQGQLIHDQIVENGFDSDVVVGTTLVDMYARFGSLDDAHKVFSRLPNRNVVSWGAIVAGYVQQGHGLPALELYSKMQQDSIKPDKVMLLCVLKACASAGAMAQGKX

>Cma2149983

RTWEAQFKSGIETIAIVAVKEGVIQLGSLKKTSEDLNFVLFLQRKFNYLQSIPGVYAVHPVLMQHATSAKKAVAAVAVGDSWPPEEKRQHATAEYWPLDQHVHHLQQQHHQQQKLMAVVRVAAGENISGNIPSSLDVTDCPVSHSTANQHLPHQLDALPSAHHLMPAGLQAHSSANQHLPHQLDPSPSDHNVMQASLQALLSRLPSVTSPPPSAANSSSSSPTFQQPGHPYFAQQAPFLHQLPVATHHHHALTRPSSIPHHLNYPSGPKQDQLAGFSPSSTPQLLYSQPLSDQDLDPTAFLKGIIS*QMP*CTITAVSLHILFETHESFFHQX

>Cma2151988

QSESFYGTPYNLIRSRCCCLHSHFFPLALPTHAFLTPSTPSQILI*LLILIKI*TIFNGT*PWLDMKLPPPSSLVVVSIFLDNPCLFVWPFSDIITFSYQIMERVGLPLVNHLLQHTLRSLCGEAQWVYSVFWRILPRNYPPPKWDNQGGPFDRNKGNKRNWILVWEDGFCDFVACSRDVLKESQEASKEIDEEGEATTTTTSLQPELFFKMSHDVYNYGEGLMGKIAADNSHKWIFRDPSEQDISFLSPWQGSVDSYPRTWKAQLKAGIQTIAIVAVGEGLLQLGSTKKVIEDLNFVIMLQRKFNYLQSIPGVFAQHRMGEAKKGPTTIVENYEVEGQSEKLNLNLRYTNPTIRKTLFLQDNTQQQPQQLSGVKRPLDAEHDPSYKLPLPLAAPLNQSPNSVAVVPSMSSLQALLSKLPAVTSSPSCRDPTSANMDPSVACIPSRRALCPTLNATESAPLMSKSFLYAAARSGDGKCTADNDNTNALIMGRPAGVPPTPAGDWPDAHRTLLRPQPDAAFTLSDEFEAYNSMFGEMIS*SHFTITPKLTSLYVTYTSYVCNSLGVYFAST

>Cma2152236

ERERESALER*ASPSSCALGFFVSTA*TCACGQNPQRTHF*TNAFAFCFLLLLPRLQMQTWLLAFCFFLSFLLLPVCCVGFSTSAFEL*RTETQMSVQESSFQAKKCAKGSFVQRETLICQGSSAQVETLNCPNGQGSTV*VQTLHRIVFLCCLGSMLFSGNRPIRKATLTYSKQIFRRKRAEKQGSLS*FHLPVSVSYSTYLPQNAALSHRPNLPYLNHQTDYYHHYH*SSHLLIDLVGILLQITLHFGALF*RKQSKPSTD*AVIKQ*SSEPTIS*HKQ**ANAQFLKHFWLKGERTHHR*GGVW*SPHFLGAIKSVFKSKPSID*AEFCLIKGCLRYFSAD*GVLRKPNSGIRKATFA*RVS*GEENSQSGFAHCHCMLLGRHSQFWVFLSEVGAFGFHLHQQQMLFM*TWVKKHRFYVSIHTEI*QVFYKGFFAKQFS*KQTWFICSFGKKELQRCGIFPVFLE*GKGRCCR*RQHLLMRV*YARHMMLGQFGLF*TGFYIALTMASVLQQTLRGLCCDTGWCYAVFWKLKRQFRTVLTWEDAFYERSKSQEVAKLNSNPAAMSIPVSLGAFGQQMSGGLADFEDPIGLSVAKMSYHVYSVGEGIIGRVMFTNKHQWVFSTSDKVTTADGVPNSKRNTRTQPEKYPA

>Cr01G13260

MRVLSSKRKPRSRPTLPPSPRLPSGPGAVAQAKLRWRRAPARRLRLCSFWTRPRRRRWQPHRCWTWGKLIRRPSASVKTPYVADIQLLPPPPPSAHPDGEAGKDAAAAGAGVTEHGRVGELVLAHAPAMDCAGMVTPGATVYMSKSAPRAPGVPPAATSHAIQLVEDQRPSGTAAVVGYHPQLAERLAAELVRRRLLEAALGAGRVVGLESQRTFGNTRVDYVLRMEDGSRMLLEVKNVVCADFPDVPGGVPPGRPAVGVYKVPVPSGCLDDYSPTALFPHGAQKPKIKVVSDRAIKHVTELTALHKHGRELPEAGDDSGSSSGQGKGEGEGSAGEGRLRCAVLFLVNRSDCAAFRPCHEADPLFAQVLKAAEEAGVKLVAYDVVWRGGAAYAGRHLPVVFGPGVTSAYDPARLAEVLHFNATDPRKNWKSTKAKAAGKAEAAVKGGKAAGGAKAASRRGGKKAKKGESVSDEDSE

>Cr03G04920

MPVHGAPSDAPPPPNISVGAALQNLKASLSQQVQSSLSGKHTPTHGEQLKSQYKQAMSDNVQRRLLGRADNYHNTTSSLDASDSEGAGSGLQSGNLTRTRLMGALGSDNGGTVPGSPLRGASPAPTSRAAALLAGGGGSMARSAMRVSGTGSGAGDEDVRDAPQSAASSRRTSMDVQRPPPVPAPAPVPKPAAAPSPPPPIVPVPATGFKPPAGLTSPAVSSPLPAPVPATGFKMPMAASPAPAVAATSLSPAASFRSDAPALSPSPGSGGIGWKFGMDSGGSVNLGGPDTDGGGGKGEGGLSPFASGQSLQTSNSFAPPPEPSGRSNSPPRADPRMVLNSYQSMQPSRPAATMPLRTVETVRLRTEPAAMQLPPGFALPPGLMGAAAAGGLGGGFGFGGGFGHGMGGGSSGVGFGTPSNLGQPPREQSRDSGNDSPETFSPMLGVQRPPPGRRRSDGSDNSDVTPIPPAGMSSTPSIPVGSFSFGINAAVFSSGPSMDSATAGGGGDEHSPAPAASSTSRHSLSQAAHALHGLQTVTEEDAASEDSPPKPVMGWKIGAVSEVESCAYAIDVTEEPPPPPPPSAVGFSPPPAFGFSPPPAQGSSLQPATASGPLSTLGWQPPPVATLPHVPVSPGAAVVPEHTSFRLSDGASMPGSDGGLRSDRGLQVEGREKLPWEDSTDDDEELQPAPLLRQSSSLRRLPPPIVAVSASGGMAPVSPGAVQLQPSPGAVTVMSSSLAPVQEADSEGRDSSVRLPKLQDDEVGAFPSYSENDVRPSLVLPAVNAASSSGQSAPPPTPAAVATLAAVAAPPAAANSSSQLQRQTSTLSSRSSGVVAATTPTGAHPPGDVSITITDADIARSGGVVPALLEIVRDRFRAKGMSNLAQFAAGSGTGSMAGGNSHGALGDEAHGAGTPQSAALAGETLARRIMARVDTFHSARRGGGPEDLVAALQSVAEHDEGGPRGVSEDGSSDRAVPVFRRTSTAASSGAAGASSAAGTSTGATAAAAVAAPGLSPTNSGSYAMLMAHSQQAAQQKVAATAIAAAAAAAAAAGGAQAAAITGVAPVTAASTVGMPPNLSRGPSIRIDSVRQALAPPIESPGAAASAASSHAATAATSAPVRAIRAPPAGGAAASLAHAQSAGTLTSSPPLEPYTSSSLSPRPAAVGTPSSPASTSKVETATQTVPPPVADGLPPALGTALDYFASTVVPVSGGTKGRPRNAAAEDERALPSDMPAVRTAADFFKGAANRATINTSAMAVPPRDTPAGISPAAGAAGVGAVAAASAPPGRVRTARDAMLNSSSATTPARPQRQSVSGRSGVASSGGGGVLTRAGSAGRRVSAAGRVVIDDEPVDLATYPSRSVSNRHVSAAASPGVAPSALRTSAHGGPGPSGPSDVGVDGLAKQQQLQLQHQALMQYQQQGYDRQALTREMLLHLAHMSPSRHPFETTTDEEREEEERRARRHARRTRSPPYPSTRGNGDSASAGQQQQPQQQPPTQPSSARRPAPAVGMASVPPVPWPRHQAANAPAGLALNPRISGGVEQLQARFDAEAAAAAGGRGFAAGGLPPRLADVLAAEAGLGPAAESPAVAAAPGDGIPVPPLSLDAPDMPEPGGGDTASAAGLSGVLLAGGGLSGSLPLPHETFFISSPRWRANDSDGLAGTGEVFLASGAGGKKAAGSGRGGAVSMLGRGRISATGGSGPEYTEDGGVSGVLAQRSRRSTTVTNTTSWTSSSDAEEAQLQGRHVDGVSGGGKGRRPDEAAGPRVQQSGGKQAAQLPVRSSAGARSSSSRGVNLERADTAGSGTAGGEHRPRLSATGRRRAAYWADEVSPLYGGPDAPYTDPDGLDREPSAASRGGAASPRGSSAAASVLSARRYVAVARAHRSVSEANSSKRLGAGAVLEGDDEDEEDEDEEAGGNARAGGADDVEEEDYDGPQDLQLDEEERQEQQPQQRRQAGTRAKPALAASRGDGAGAVGSQLRWGVADDEAGMGSTRAMASSSEHDEDEDGHEHSGRAPRDSTDSGGLTPDGTRVRQSLQRLLDMRAASATSPSRPAASSAKEQSPFAGRGPAPPSAAGTDSPAGLTGGSDGRSNSRSPSVTATAGGTTDADARDREQLLRRKGRAEGTSRTAAQPTATASAGATTSAVQRALLGNGAGAGAGNAHADGSADGASSDDAPPEDPHELVSPKYTQDMAEEEYGWDVATGDADVDNIASPRGRRPAAATTPRGGGSSRGRRGGRAARPPPPPLPYDPAEVPGAGTHDASPSYTRELEQEELRDTMAELIGSGGGGGGTREPPPPVMGTAAAVAAAVAALQADQEPSFSLSAPPAPGSRFGSAPGTAAAAVSAALAAAGSGSPAQHSPRGRHTVGGLPAAAAASTQSLTHTALASDGFNLALTNLLSGLGPVTLSSRASASSSSAAGMAGATSGGAGSASTPGKPIATQTSVGGPAVAVSTQTSLVAADSQQRQLLQGLQALEEQARRAAQATAALSTSASARGSPSASQPVADVAASEPDPELRRLLLAAWSKWYLRRLSADSGAGHERDSPAGGSAGGASDGQEAATEAATAAQARLQALTAALSNGGMDAREEDPSSSPRMQAAASASRLHMAFTRWLGAARSQRSAKAAARAPYQHAASSIPGLAVGANTNIAPPYPTSPSPAAPTYIGAFQHQHPQPGPGSLTYGSYLPTGTRQLAQPHAHQGSRAPAPAPSPPPVATSWPAALRTISSPPPQSAAAAAYQRAGMVAASAAGLRAGASPALPLASAGALSSGAGSVVGEALATNTPTVSRASGVRQVVSMLQAPGQAQPAHRHQPLATQAYTRVGYQGSAPAPPAAAAAAGAAAPVRRGGYPALGLLGTPAVPSAAAPAPGPASDLEAAVAQHLLASLVSGSGSGSDSVIVAGMQLATGRALLPATLASGTRGFALGGTLGGTPPAGGSTGGGLDARPRPASTGRLGPGASRPLAAASPAAAAGGAGGGGILGGRAAAAHSGLASSLGAHRPASPVAARRVPGATSSGADNIRLAPAANLGGLNLAYYGVGK

>Cr03G08450

MPQVVVSHAGCSLELLAVAPSGAAASISVQPIPAGEVVADMRALRDSGGNTDASSARVGAAASPAGAAAAAAAAPADQLAVLWGSGRLAVMVYDHACSRFVAVQEVALCRPGTQPRHTPRLLAVSPSGRHGLAAAALQDRVMFLPPRRQLVAAAAAASTTAAAAVAAVPRVGTAAAAAPAAADAAAATQAAAALGAPAPAAAGREGQDPDGGDDGGGEGEGGSPLLADAPVHYHHWVTRGRGGRQPVMTAGPVPGAAPRDLRRAVADALAAFQPRVGEVEHRPPPPPGPVAADAEAAAEAGAGPAGEVAGVVAAVARLAAAAAGDAPDRAQAVEGEAVAVAAAAGVDAVMERVAPLAAGEEAAPPPPPPAPVAPPAPAAPLPQLAGAAPGPGSGPAAGALASPPPAASQAAPTAAAVDAEALDLLLLPGCGAIWDLALLEWEGECEATGGAAGAVGGSGSGSGSGSRRGSGVGGVVRIAVLAHRSSTDCGELLLLHWSRAHAALLCSLVVRLDGGGGGVERGPGGAAAAKREHERDREREQGEEREGEGERERSGWAGRRGRAGADSGGPPPLGTAHLLRVTPHGEQPAGGLVVLGTHGALYVQAGLLHALSAAAPAHPHQHTAAPGAAVAGAGAAAAEAPSGAAPLLAHQQSHHRPQSHPLGSAAGAADVSGGASPARAAAADAAAAVAAAKPTPPPPPRQAPELFLADLLRTRHRRSSSVPAVAAGGPGAAGSSGAGTSSREACAPCLPVAAAAAATAAAAAAATELQIEPPSTAAGRGWSFLRFARTRNLAPASPFAAAAAALGMSLFAATAAAKAAAAATARAAVTGTGGAAAAAAGGGPGATETGDAGDTNADAVATTAVAGRSSGVAALLHRYGYVSASGDVGAAGADRGGDQRGDGLDGDRASWAAGGDSMEEGKGRQQAGGSIRDRPKSTQEMIVGLRAAIQKAAAAARVGARGRPEDVDAAMRVVVGPMVVKISVGRKEPKPLRQAAPVLGVERMPLPEPTQPGHILAAAAWLQPAALPVPAPAPTAAANGGPAAVTTAGASASAGRVSVSRLLLCSATGGVLCLTVAASQQPEHQALQGGAATGSATATRAPAATATSASLAAPAPASDGGVGGVGVEGMEVDEGAADSVTAQHDNTGAAVVSAAQLEAAMRGLALVRLGALPCALLGDAAGEATAIAAATASAQRRRREDPGSSGGDEEGFVEGQPGLRVPRALAELTSLTGTAEAGGAANSNGSSCGTGPDGGGSSRGGTGLLVWAEEGGDVLFLRHRLLPQPQQLLQWLADGEQAAWRRPRPYRSQAAAAAGCAADAAGAPRSPSFSWAARGARPAGQGGAAAAAGAAGSRLRLLPGAAELAAAPAAASSLAAAAAVLMGMDVGPPPPPAPLPLPHGGKSGGGPQPAADAAGHGDAGQGVSAAPARAQPAAGPTAALAQAKPHPKSRRSPQAQVRAPAHAPAPMCWALKFRLAGRVPQPRPVVDFVVSEAQAAQAHGSRPLQVTGAVLELPSAGEGGSAHSVFAPASPPTRRSLADGLAGAGSVSWGAPTHSAAAAAAQQQLPALSWEPPLEVLRGGQSVMVPVVGRVPSPGATPTVLAPVVPAPPAGSLVVLSFLSGSRALASAPGAAIAGAATAVDGGGGGWGELRDVTEGLQLRHWEPTVAAGLLAEGVLAQPRSTWPSPKSSRFEEQAAVAAAPLQAQLAAGLGAGGGGAGAGAGVGTGAGGVGPGAHDAPGTPMSVGQAASMFLPTSAGSIGANDWQQVQGAAGSAGAAGAPAAAGRGRISASGFSTDVPPADGGGGGGARAAAAAAAGASAMVRAASTQAFSKVGASNDDGLDGGGGEEAEAETRLGLGLALGDVDDSLLSGGGGGGDMAAQRSAAGTPAPVMMQTTADTPGPEADMEVSMMGPAGAMTQTGGAAAVSSGGGWGCHPGAAVRRVGHVLSTDALQTAAGSEPLSIEALAVEAVEEVELPYAGGAAWWSAGGGGGGGGGGREVGLAAVAPGCVALMRQQQRCITVLGVLGRHQLQQRRAEAAAAAATAASAGQAPGAGTAAGLVRSLSNGSSRDGGGDVDGGGLRGGSSRKRPRGGLVLVELGSLPPLEHEPSCMALRQLPPAWVPPPPRRSASRRSLAAAVAAASAAAAAAAAAPSLSAQAVPATGGGLAGGSAPPSLPPLPPRMGSSGSGGMGAAGQGAGISGRLQVAPANASSNVAPAAAAWRYQLLVGDHAPSIQLFLLQGPTGGGGGGGGGGALLTVTHLAAISTAPPPSLPLPRTPAGPPPAPVPESVLFLSPEPAGAAAAAAAAAGTVVSTPRAGSRKALSGSGAGGAGSSAVVASAAGGGARAGRGRRLRHAAGEPSGAALSSSAVVSAAAAEADAAAELTVCLVGLRSGGVAAFSYGPPPAPTAQLAPRAQVQQGQATGDAGGCGGGGSAMDVEVDGGGTEATAGPSEYVLCRLWHTQLEQVPLRLVALAPQRPWQALAVGTSLHAVSVAQPSGRLTTRQLLPPGGTWGDHGASAGGAGAVPPAAAAVEAVAAVPLFCPLPAAAVRSRKAGGSRRARSPQPQRPPPGVGSGPGTDPATATAAAAAAAAPNAPGPSATAAAASELTLVALTDGSWRLLSLQFLSRAGRQWQSVAGLGPAGRHALPPPPGTWPIGRPQRVLRVPPLEQPPLPPSVPAASTQAGAAGASPLASRPRSAAVGRSRAPTASGEAAAATAEDSGAGPSGSQAPGVAAAADAAADTAMDGSGPGPGGSRVVPALSAAAAATAAISTAATTPPSSTSSYIVLLAEDPAAPPRPPPTAAADAPAWAAAPRVTGAEWLHVLDAASGRQLLSYDIATATGGFLATCVEAWQVGPEAGVQPAAAPNGAAAAAAAAAAGDARRRSGSGHGLQGLLGPWDGEAGGGAAAAGFGEWAFEAQAGEEARGAAAGAVGAGAGAQAREQDAFRAHLAILRRLAAAQEDEEDEDVIGAGDAAGAAGPAGAAGHRDRNRNRGGQQEWGDAMDADVWDVDMPDGDAAAGAAGAAGAGEGDVEAGMALGAWRRRLQDHFGGAAQQQQQQQQQQQQQQQPPRRIRVPEADLAGAAEAAAAAEAGAHAGGAAAGAAAAAGGRQEPAAAGEAAAGVEDEAAAGEQDAAFRAAEEALEAAEAAALEAGAELVAFRQRLDEAVAQLDVRRAEARQAAEAAEALRGLPQGGLAEQVVEAAEAARQDVAAQEGRLGALEAMLEERFRALQAAEVGAAAGRRRLLRVLLHEARDEVQEVVVAAAAAAAAADRRRRRLVELEEQLLRGAAAAAAAGAAAGAADPAGAAGNGRDADANEDADADAAAAGAAAGDGGARQDAVVLLRLLEELRHQFGGLPQGLRLPPPPPRMRGRLGALEAAEMAAAAGAAGLPAGPAARGTAAATVPPLPPPVPRGEPATLLAVGVGSQMGGEVMLLRLLPPALPPSPPAPGLAPDTEGDDGDDDGADIKGTARAGASADADMSDAADERAPGWQQPLTPHPPLAVHAQQPRQGLQQQDADAQPSAPRLVVAFRLLLPKPVAAVCPFGSQNLVVAAGRRLLFFRLRGGRLHRTGWHATRNTATALAACPARRLLAVTDGATGVVLYSAILAKAGRGGGGGTGGAGGAGGGARLGPIIALRLVAADAVPRPITSLVLLPAPGAGAAAVGCTTPASAQQLLLEQEQERHVLQQRERRRRRRRRLGLGLDGGRADGRAAHGDAGGMDMDVDMDVEPEVELEDGGMEGAGGRRGRGAAQRLDASDSEGDDEEEEVRRDVREGPGAGRGEAEGAEDGVREDGDRERDGEGGPRGPRHAAAGAAGGSADDEAGDGAADRPAGVAVDDPSRMPVVAATDTCGRLLLLAPEPASATPCRNLVPVAAVQSGDVGLRLRAHVAEPPPGCGAAGSTATAPPASLVLAGLSGAVQQLRPCPDAAVPLMALVERALAAHTAAATANAAAASRNSTTAAAAGGSAAAAADDAVGSDAWWQWWCPPGAHPSGVLNGDSLTALLDLPPRQRAAALARAPRRALRAALRAATSDR

>Cr04G00720

MAENQAPPKGARVDDATRRELRRELHQACQDVHRQKEVLCDISTNRLVEVIRHSETLLDKAEGIQKAKEVAVSAEIFRNLADCNNELANKLVQKQGRGPMDLIVKLRAAFVHTANPQVDGAEDPTAFNWGELGAFVEHLRRPARGASCMFGPMLVQPKERKAPAQRQKRQAPAELVRPEEKDCMVEGSEEKQDTDRNMEVMWGVIKAQNSPLVGFPELVLNAASFAQTVENMFTLSFLVRDNRVALVEDEERGWCVQPLEGGKAKGAQQPPQQQQQHAPQLQFIMSFHLKDWEAMKCYVRPSECLMPHRDKAAQHERDKAAAAAEERATKKARQHERAA

>Cr05G00660

MPGQNTSWLADDLELRRFEHRSTGTLPVSLMSDVLLTTIPNPLASHVLPLLSQHLGKGKDGPAPWKWYQYQLVTSRPAGPGSPAGKEGLAMSETGSQRSGADASDAASSKDDASIIKAKRARIRSAGEDGSDLLEQKKSLQEGVHAAMYMLARSKRSDNWKMAVFKIVLEGLIPFLVVFNPTNDWGIKTSNPLWQALRWLLPRSPIARIWGYDTYIKIFYVFVALVYISVGCIAALTLAMRRQEHSKALKRFAAGIQILTDVVFSLFYVAVFDYMTFLFNCHFSGHGLHSHNYFTGVQCFKGPHLIHMAVAGVTCVVFFLCTAFMLVGACDLNPVAQGLMSSPAAATRVKVLIFKAAYVLCVNVLYSMVKVQSVAMLVCAVLITYFNFAAIPFVRAYINPYWVGDWAAVSYTCVIYCLFKFNKNAHLPSVEHDMTMLVLYGVFPTYFVAAGLTWLYIRWRMSKTAAFVGVDPTIKLKKVYSFGSPEEVELLSRGMRHWDSDGVILEDAAQLGEMVIKCGLATYPGHVGLLILNANFLMEVKRDGPAARTQLQLAGKGNPSMIQRYQIFSTIENSKRLKDGQSGQLDLQSYVEFKRNYRAVIRVHKAALGAQRDFWSLMQKGHVRVSTVEAIMRAMDEAAETAHQVYKRVLERYPSNGKLLRCYGKFLEDVRNDPVAASRAYTEAARNGGGDGLLSLDLQIQGSDKPDFLTSMDLHEDACMVINHEGNIMMVNSCITNLLGYAKTELEGNNVSMIMPQPFSGRHAGYMQRYIQGGEPHILDSVRDVVALHRDRVVFPLQLCVTKLSGIGTDSIFLGVMRPVPLDAHNVRAWVAPNGLVLCTDPQFASLTGLTSEEMVGLTVQSMCVDGTAIEDLLDYCKNAPYEELASGSIVRRFNLVNRHIHQVPVEVKITPGGTDTQRLFVFNAHRTDGNMDGLMVVDTKGAVTFATWDVAAMLGYPLKKFLTMKLDQLLPQPFAAMHARHLREHPATIPPTSCRAGALVSLLNSNGAQEDVASGHTKHVVCVRKVDAGSPSDMYGDKRLTLTCTMDGKVLSVDQPTSLLFGFTSGSVVGSNLADCIDVFAEWREKAGAHQTELLLLSLLDKEAEMPGTSWRVKVHGPEDEHNLLPNIDKRGPKRAVDELGGNRGNAAFDLALGPEQAAGGDGGMMDSSQDHLRTLAKIVLWRRDLLCGTVELDPKLVVRRADVATGLIVGLPPSALARMPLYKLLEVPRGVGWDDLMVSKETAGKRGKKSALKGGTGQTVSPVAAFEGPHPDCGTMRIFVQGVTGGGMNRIAAMLHPDTQFTVIAEDAAEARGDGDGAGKAGEAPDVSDVDELPPRDNASMKDSKLSSSSTSDDDDGSADGKVAPGSEEAPDASVADAAEAARMHMAAHNQSEFVAQWVRTLTRQGTTTADAAGERDHIPASRAGTKDLMPVPPPHEHGSAGLRAALGGNSKKKVSIKKMLELQPVAEEAAGGEGSGSDGGSDKGLSHAERSNISAQGGKPPSSGGDDASSVNDDDEATSQGLGQAFTDASSAVDVDISTDTRRARLLKRLTRLLAGVRLAAPLGAMRSRTLWLLLGMMVAHVAVYVIFTSLISSQFVVSVKAAIAEFCTRPGVAAESVCELGLNSTIGDLRAAIDDLKTYHQGIYLGGVTPRKIPEPTAYHIWTDRTNSFDAYMDTHPPRTANETLGAWQLGNRVIAAGRELLYWAGALAGNISSLRQYKLLIFASPVALFAGYTRSLDYLSLGGTETGVLIGGKKGAGAGLKRGSHGRMRINGKELVPSWKALLMFLLPLVCWEITLVAIGAVSFVKLSGMQGPLASLNMASHVIYRYTRVRLIAFLLVAADTSDWKDHYRGMLVKEVSNLENEYDTLMYGGISITQAGSVFNKPVPASTFASV

>Cr06G12880

MRPARPAFPLALALALLAFVIRAGATLGVGITEPVLNKPLPSGFRVTCFDGQELIFTNDNYTNGVLVAADAYAYPVGQCRRCPAGTATMDGFRCIPCPSGYWSDEGARECIPCPVGTITVSTPKGTSGPSRTYDQIQRLNAGAGSCRKCPPGYFQPALAGTVCLPCPSGFTSTLGAEGCSPCQEGTFHGDGWQLAANGAYGARGMRNASGGTGGREASVVELADITGIVPTAGVEYTVLPNTCIACPRNTYQPLKAQAATGSLGLGAFSACRRCEDGWWSPPGSAYCQPCPAGSYRNSYFDGTVQLSNAAQSLTYNLANETAPNTLASCFLCPKGTFAPDPGASVCQPCPAGTHAAATGSTGCNRCGAGTNSLYGLRGQQLSWQSNIATGPTAAYYTYTISGFDKALYDRNFWLAGKGEPCAYNLPGYYTDVEGLPVQLPCKPGYFSPEPYTDKTKCFTCTFGTFNEEFAQPICKACWPGSFASQRAMTKCEITLPGYFTNPNPVARNATYDLTVLTTTAPNMTELVAGQSAPTPCGLGYYQPEYEKSYCLACANGTYADVVGLRTCKDCQAGSRCKAGFYADKPIGATACRSCPRGYYGPYEAAYSADGFTPEGPRGCFKCNFDTYTNRGAMTYCNNCTDLLLSTGNAVPTCTETTGAMRCKPCSMLINRVENRTTIF

>Cr07G08900

MSLLRAGRVVLDALVFRDQEGMLTRLVRSFSAGVILALALVHIIPEAVEEMAGLGGIEYPLGGTCVLFGVALMVFLEHLAHIMHGPHSHAPAADSAAAAFTALPSSCTDIEAGATPCGAAKRATAQTSSNCEADPSGVLASDSSVPMNTSPAATQAASGSLRLKILAYMFELGCVFHSFIIGISLGVNTTDLVEVRALLIALSFHQFLEGVSLASVVLRGGFSTLKGAIMILTYSLTCPVGIAVGMAIASSYDAESERARGVQGTLNGVSGGMLMYISLVQLVAEDMGRFVPGSPSGGASARLLSFLALFLGAGSMCILAVWS

>Cr08G00050

MPITARPPISAAAGASGTAGSLPARVTGPGRQNQGHNAQHVPVARAPTLASARSTTRTSAASGTTVMTVTDRVRKSIERRSMDARPQEQGEPLEPLEQVPEDPLERLTADLAALSMQHAAAVAAAVAITTASASSSGTSSSAPALPAASSGRNGRPQGPSGRYGSSSASGPAPVHASYNNSGAPSSSPNALQQQQSQSGSADSRRCDLGDTGLPAQLLALRDPACPVSLPPPTGLPSSCLPLDNWKLDKLVLSLSANKATWRRSLLLFEWLKAAGHQLDDRLCTTLIRVCSDHGDAVSALAVYDWMTGSTASGGAALEATAYTYTAAMRAALAGGLTDRALSIWNEAWRRHSAGRLQLDCRLCITYLELCTRLGLTDQALAMYAAMRAAPAGSRMAPTVHAYTAAMRAATEGGRWYRALDIWADMRSAGCEPTGHAYSAAISACAAAGDWRRAVALFDEMTGPGGIRPDVVSCTALITALAAGGEADRAEAVVAWMLSNSVRPNARTYTALMAALGNAKRWARAVEVLGRMQTPEWGGVQPNAYTYSALLKSLGEHGQWQLAEAVFSSIERQVLGPAAGSAPPAAAALSLASALVPAASPPPSPLAAMIAEASATAAAAAAAVASAAGSAAVAAAIRPPASPCASESSSSAPSQWTRPSQLNGLSLDLHSPTSTGGAVEAGPSRRSFSLFSHPPAPSSSTATSCAPSSTGTSGRSSFSADSAVPSDVYSVASTSAAEAWRSTLDHASAAALAGQLAAMAAASGCGAPGPLATVPEGELVTFPSIDAEEASTSAATTLAPGVYTPGVLRAGVAPPPPPPPPPMSSSAQQQQQQQSASTSGNSASSSSSGGVLNEVVCGALMLAYERAGKWQEAVAVLLRALNLGITPNTVMYNTAISAAGKAGQLEIAEKLYGKVRQPDAVTHETMIAAYGMAGLPERAEAVFSAMTSAGLRPRDFAFCGLIAAHSLAGNWEGAMRVRARMRRAGVQPSVHVYNALLAACERAGQPDRALELLGAMRREGVEPNTLTANLLQLVGRQGVRSVESQQQLAASVSAALAAYGALLMQTGLF

>Cr08G01240

MQGHKNAVTSVCFSPDGRSLVSGSEDKTLRVWDAASGECKATLSGHSSAVTSVCFSPDGRSLVSGSEDKTLRVWDPASGECKAMLSGHSSAVISVCFSPDGRSLVSGSDDKTLRVWDAASGECKATLSGHSSAVISVCFSPDGRSLVSGSDDSTLYVWDAASGKCKAKLGGHSGNVVSMCFSPDGRSVVSGSYDGSFSVRDAASGECKATLWMDHRRDRINVVSSVCFSPDGRSVVSGIQDKMLRVWDAASGECKATLSGHRSAVTSVCFSPDGRSLVSGSEDKTLHVWDPASGEYMANTKFWGRSRAVSSVCFSPDGRSLVSGSHEETTLRVWDAASGESKAKLYIRKNSINSSEVSSVCFSPDGRSVVSGSLDSTLRVWSAVFWDGKPPLIISGHSSAVTSVCFSPDGRSLVSGSEDKTLRVWDPASGECKATLSGHSSAVTSVCFSPDGRSLVSGSEDKTLRVWDPASGDCKATLSGHSSAVTSVCFSPDGRSLVSGSDDKTLRVWDAASGDCKATLSGHSSAVTSVCFSPDGRSLVSGSFDCTLRVWDVASRECKAKLSGHSSAVTSVCFSPDGCSLVSGSHDETLRMWHARFMLPVSTRLHSVC

>Cr10G02760

MNAALVAILILALGHAQGLAPLPSHSLGHRNEWSARRMLDEDADEGTNLDDSDSGEGPSLADAVAARVVNSTCAPGCSEHGVCNEELGRCDCPRHFVGPDCQTPNPEVSDVCAKYGFTVSQCRGASPCFNSCNGRGRCVAGVCHCLPGFWGMDCALSRGADGSVQLLEGQGYVPRKDSIKIYVYELPPNVTSWFNIKRLDRPLHLLFWQRLMSAGLRTVNGDEADYFFIPLNTRTLMAPEQAAWILPYIRNTWPYWDRDNGHRHLIIHTGDMGLHELPLGLRRKMNETLSNITWLTHWGLHTYHPIGTWFPAHRPGKDIVIPVMITTPGFQLSPLNPAVAEKAAKRGRPYTREQTFFFAGRICGDRKPPDPLTHECAPKRTDYSASVRQRVYFHHHNRTGFKVLTGTSKYMQEITSHKFCLAPTGGGHGKRQVLVALMGCIPVTITDGVYQPFEPELPWADFSVPVAEDDIPRLHEVLEALPPEQVEQMQSRLHCAAQHMFYSSSLGAIIGEDGRYDAFETMIEILRVRKAHPDVAPDKYVEVDERFRQFANCELGGPDPKTLCAQGMDRQHPDMPTCAECHRQVSARRTGSSFFSWAGGLSPHQTCSKYGCPST

>Cr10G04160

MAEQTGADSAAAAEQLALLLSSANLGSEGGTNGAWLTQTDILTTDNLDFEGVLNGGGGRQGKAGISQQQREADPFLDSAQELESVYSSHMSSAQDIMSSMHSQEPNLFLSTDPNNINAAGGHGDGPCDPAPRSLSASLAHDAIGEAGTSAGASHRSGGPNARDGAQASPRKAQRGQGYRRLWQQVTKVSKGQKLQDNLGLAFSDMTVEDLLKIVLKLAPQESAVEAIKQGLVYLDSSATAALLKELAKQGYLKRAVEIFDWLRSLAPGDELSSLCDLYTYTTMISQCGSHQQLRRALELVAEMRSRGIDCNVHTYSALMNVCIKANELDLAQDVYKQMLEEGCSPNLVTYNILIDVYVKRCQWEEAVRVLDTLEKQAEVRTYNTVISACNKSGQPEQALKVYEKMLAAGVKPSATTYTALISAYGKKGQVEKALDIFRDMIRRGCERNVITYSSLISACEKAGRWEMALELFSKMHKENCKPNVVTYNSLIAACSHGGHWEKASELFEQMQTQGCKPDSITYCGLITAYERGGQWRRALKAFEQMQSQGCHPDAAVFNSLMEVLWQSGVLLAQAKALQLWTLANRSGHFRIYTNSKQDSNVLQYSTVAFTSGAAVVTVVRWVSELKNKLLKDGAGFFRDKVVFTLHKSKQNRSEQPSARIFEAVSLLLAGNASPFAMHLFDQTITLEAASPQLTAWLRSAAFNDFVHIQQSQQLKRISLDLLLSEDVAIAARCIEAFSAVKRYEQVFSINPALCNQALLAQRPQIVALALKYGAAFGLKDDTIYDALQLFDRVSCSGGHINVAAWPLMLCSCLLLAARQVEAPAMWPPLEQVTLLTGFGTDVMVAMERNVLLWLGHDISTISPMRVIQLYLERLGHYLPDFKGIDRITKDLQTLVLKVACSPVVGLRPSLVGAAALVVVRRARGMVPAWPSVLQTMTDYASNDGELAACVMHMEALMQQ

>Cr12G05740

MADERSYYQATYQGHQQQYSGQQHHGGRGGGPGPGATGPPGAPGPQPPAVGWGGQQIQEPSGYRVPWFGRSRSKYGGRFGGKKGDPGPPGAPPRYGGPGGAGPSNRGAEDALSVEEMAVIIQKLGQNEELPEVIFRALHHVDSRAVALLLKDLSRLGKDRRAMELFDWLRSANERSPLRQLCDVYSYTATISLCIYSQDVDRAMELMNEMRQRNIERNVHTFTALMNVCIKCGKLPLALEIYNNMRAANCMPNVVTYNTLVDVYGKLGRWERAIHVLDLMKQEGVEPVLRTYNTLIIACNMCNQPREALAVYQRLLSDGYTPNSTTYNALISAYGKTMQLGKALEVYQEMLRQNMERSVITYSSLISACEKAGQWETALRIFNEMQQDNCVPNTVTYNSLVTACAQGGQWEKATEVFEQMTAHGCTPDVVTYTALISAYERGGQWQKALQAFGKMCMQGCKPDAIVYNAIIDTLWETGIIWAQGRALQLFLTAVQQGHFRQEPVVRRAGRVEVNLHAMTAGVAMVCLYCWLLEIKRIAAEKGLAGLPQTLAIVTDMGKASREQGNCIVKEAVAAMMSFWDAPFRLVQDGAYASVLEATSAQVVEWVTSMPFSAQLASLFPITDSSKMTEDMYAVREQQVLQECQEAFAAVQHFENTHGLVLSNMSTGYVQQRPVLISRLLDLSSKLGLRDEVVHDAVLLMDRTASQAKQVAEDVLPLVGVAALVIAAKQGDSSDRVPTNAEIEQITELPPTSVAKMEWNIRGLLADDISAISTMRCLKVYLERLGYRYLDKQDVYGMAGFAIMLAVESLYDLSLLNCRPSVVAAAILYAERRLRGAVPFWPSMLSKMTGYHDMSTPELTVAVRVAQKLSRKLVYAQMYKAQLIQLAGQAGPIAGPVSVAPELITGPPSASAAAAAAGGSMVAPGMRGPPPPPPMGGALGAAGVMMPPPPPVMMQTAMQGGLVQPTGWPPQARFDPLAAQHAAQAGGQAGGVGPSGVGQQADDGATITALTQAMQGLTTGGRVMQPDWQRLGGPVDDGSEGPGR

>Cr12G16140

MTTCTYEHRCDCPKNRTGDDCVDIADTDTLTSRCRRNYYPSVKECTTSELSCLNNCNKRGTCVSGWCHCKPGFFGADCSLSLDAEGKPELLAGTGYATRAKRPWVYVYELPPDLTTWTNTKRLDRSTHIHFYQRLLGSGARIADGDKADWYYIPIRQRMTADSRFLSEAVAYISATYPWWNRTGGSRHFVIHTGDLGADETQLGARLQAPNITWLTHWGLTMDKVFSGWKKAHRPDKDVVIPVFLTPGHFKHFGLERTPLHPLMDKQERTTTFFFAGRICGDRKPPKTGSWPNCGPRSPGYSAGVRQLVHHHHWDRPGFKVVLHEPNYGAALGSSKFCLAPLGGGHGQRQIIVSFMGCLPVCIADDVYEPFEPQYNWTQFGVRPAESDIPELHTILESVSAKEYAAKQRALRCAAQHFVYSSIVGGLFGEDGRYDAFETTLEVLRVKAEHPDAAPETYRQIDEDFDAFMDCRDPPSFSRINCSTVGESTG

>Cr13G03250

MAVQCHQGCTLHGVCNEELGRCDCPRTRAGAACEVQLTGEPLKQRCKSMGFSGISSCTSEKNTCPNNCLRRGTCVAGFCLCQPGFFGNDCGLSMVPAGPNGTRTPVVLSDRGYTPRAAGPRVYIYDLPPELTTWRNDDRLDRWTTRHFLEMLTATGARVGDPAAADWFYLPVRLRSSSDGHVLRRALEYVQAAQPWFNATGGKDHFVLAVGDMGRLESERGPLSANVTFVSHWGLYRSKAEQLQSPHWRASHRNATDIVLPVYLTLRKLQKFGILGSRHHPKFATVAPPDVRERNGPLFWFAGRVCQDSSPPRTDVWPNCPKAMGYSAMTRQAVYFHHWNRTGFAVLRGDKQYAKHMLTAKFCFGPMGGGHGQRQFQAALAGCVPVVIGDGVLEAWEPYLDWNDFGVRVAEADIPRLHTILGAIGPEEYARKVRSLRCAAQHMAFSSVTGAYMGESGRFDAFETLLAVLAARARHPDTPPELLRQADPPPAGGLRCVGAGDLAACPRPWA

>Cr13G05580

MYTDADQLIGSGMNVEAYAKSQQRLVATTLPPDAQRPQQFTAGLFRRSWQQFEAQLDRCHTFSCPSCRRKPHVLVGDATAITMSVDLYMGTPVTDRIVLPPSDMGKRAHKRADRCWCPEEKGRAARRGYSSAPRGDGKDVYGQKVAGWAGVGDSAKQHDSPAAVGAMEASLPAADQSSGLTAGFLRVASNATHQWQRAPAGSLLPVAAFLRKLRKLAVLCGEGYDGDTLFLVDRFHWKNHTGCSHAYNLSLYQDLQGLNSEVAEQVNSLLQPYKPMVSQMRQDNFMSCLRFVLGDRNSRRISFL

>Cr14G02650

MQPTSLTGRQQQQLGLLGLAGRTQPDMLEAARVAALLNGQGQFNLQTSGLGSGALSGQSLNLSLGSLPTSSLQQTSQQAPMQQSSQLGLPDQLALLSGFPAALFPQQYGSGDRDLQLGGLRNVGKTKSSDSRSSSAYASRHQAAEQRRRTRINERLELLRKLVPHAERANTACFLEEVIKYIEALKARTLDLESQVEALTGKPVPKSLALPTGMPSVLAGGSTSADNTNASPRMVGAATSSQGGPAGSLPSGQPGAGGAGAGSLASPSTTPPPTMTAQQASQQLSLMQSGGQAGGSQGLPSQLTLPSGGAGAGLLSAAQQSLLGFPQSGGLSLSGAGLSLGGSGLGHGTSGISLTQFAGNLQAAAAAAAAASHGAGSQSHSQSQSQHSGLSLGSHHVTASQLNELQAMQMMQSLQQHHNQHAAAAAVVAAAGGGGGSRPGSTFHPTNNKAFLHFNEDAYAFSGKPELSLPARSLLGAAAASAATPSTSLQLTTVQLPADSNTLLQVEMARKAASGSPVSSEESGVPLKKRKVLVL

>Cr14G03470

MSTPRLLRHVSNLREEESLLTQMVREATAAYEHVLGLVQAVRHMPSEARVHMVLESLLRQQAVGCTIPHTHSDSVLESRLLCQAAAHLKALRELRVQLGAVQSQLHAALDGLYDRTAASAVAARAHSGSSTVVAAAAAESSGAQPEAPEQLSSPSHQQQWQRSSTIPGSPSSSPQASSSAASSARTVIIRDGDEENDATYDSGFAASAGGGSSGGRYEQYKQCPQHLLAAELVLTSPRQLGRRRREANSSACSRRAAAAAVRSLFGNNHGAAAGVSARVNSGGAGGAASSDSSVFDLISLSSACPSPPLPSWGCSAATVAAGSIRHQRHAAVLCSKEGRQGPEQQLQPMDQMQESWSFADAASSMSCDLVARDVYHPKSGRHLDLSPSRQSPAAAPPATIAATSSNMVRSGPSLHAVSVMLPGGGVLPANYQLTPEAKAWFEDLRKRAEEAEAQQLAMEMASPTGSGGSSSRIDLVAADSVLQQSAAPSAAAAGLCDEFCNQHYQPSEGGDFDFPSPPPPPAYEESWQHPTLMAGGAVAGRRGDSATEACCTGDVDTQTGSGPSWTAHTTASGYGTGGMAACWPAAAGATGVMACGAAGHVQLDSDGYGYCWKRLEQEATAMFADEFEGFNDIPLTDRSGAPDASDEQQDEEEEEEQEWPPRDQEVLDALIAEGASNAYASEVVQGVPFTYYTKAVFTPRKWN

>Cr16G00520

MWQSGQVSSQNTSQRTPGTNIQHGAHVSQSQQDALLGTSPGGLGGAQLQGWRILPAGGSGAPAASQQFATAGGLLLQAQYLPVAVSLPPGLHQTGHGLSGAGLQQQPPALLSPAQYSLGTAAVQTIPQAVPPQQWGAAQTPSAWAAAPGPPTAQQQLVHQQQLSHAAAQLQQQLHLHAHAHPQLIQQQQQQLPRGQPAHEAVASRLDGLDGRAVAALMKQLSTNGLAAAAWDIFDWLRGLEPGVDLGRLLDVYLYTAMISLCSSNRRDLDVALSLSREMAARGVPRNVHTFSALMNVCIKAGQHQAALDVWRELQEAGCRPNVVTFNTLIDVYGKTGQWAEALKVLARMKEEGMAPVTRTYNTLMIACNSSNQWHEALSVYAQLVAAGQAPNTTTYNALITAYSKAGRLEKVLETLREMEAAGCERTVITYSALISACERNGQWELALQMFGQMLREGCNPNIITYNSLITALAAGAQWERAADVFAKLQRQGCEPDVVTYTALINAYEKGGQWRRALQAFKRLLQQGCRTDHVVFAAITDVLWDSGTASAQARAARIYRAAVGSGAIKPVTAAPPAAAGSSTATTSTAGTTSNNSSGAVPQALMALALSGGAAPPPGAASGGSATGSATAATGSTAGGCLEPLQEVHLGAVASGSSVAAFFCFLAEQRDRAIESGPGALPPRVVLRFARSRGGRDAGPLLLDALQAWLAARGAPFKLMQDFGPVPAQRLEAAGPDLAAWLTGDVAALELAPFSSVGAGAGAAAAPGGGAGGGNALLQAALVGVDEESDHDAALEASCRDAFAEVRSFEATHCLSVAAMGPAYLARRRELVANLLSVFEAVFAGGAVGAADGAAGAGAGGGGGATGGASGGGGGGLPRVAVHDAVLLLDRFMSSGPALQDNLLTMASVTALRMLAVSELAAATAASAAAADHAAAAEAELEGLLARVTALPTLGFQSMEVNVRSALAGDTAAISAARCAAVYLRRLGAAPEAAGEEQLMALLDAALLDDEFLNYRPSATAAAALYAHRLRRGDTPPWPTAAAALTGYADLQHPELAAAVGAVQR

>Cr16G00700

MAMPRHGRRPARCSSNRASQWLLLVLGVALAASPRFLLLVEAATYTLTLDKLGPTVNPTTSDAVTFTATVASPDSTTAVFFTLDYGDGVTAETTRTTTGALSTTPTANLVSAGTYTVTYASIGTKFVTLRLYDSAVAPGVLLASKTVPIYVEDSTLTATLLQSGVPRLNLAFSGFKGRVSSSTANRADMWATIQLDTAPGVFESSRIFIGIAPTASTNYDFVIPDQVYNLEGAKTTVLRIYDAPVGGTLLRTFTPAAANAVYVVDPSKYVLTLTVGPTSVTTADQVTFTQTTTEVSYSASATSPILQWRFNWDDPSVVETPLAYPDALTAASNFPTTATAVSSAAASTFRYTSTGSKNARLRLYDGANNVIAEKIVVITVSNAGYTLALAKTTADPVTTDDTIAFSAGAKHLSSTSQVWWTIDYGAGESSPRTALTMTNVGAAAPNAIASLSNQYTSGGTKLATLRIYDRDGVGANTGLLLASTTVTFTVTPVLYALESAVEPFSPIATVAAKWSFRIQRSKATPAGVTESIKCAFFGADTGTAPADLAAWLTAANGAGGLTATILPSSIPSDIISFTRTYAAAAASLQGKLQCFIGSTPLWDPYYPTPVFQVLAAAPTYTLSASVTPAVVPVDTATLWTYNIIRSVPVPAGGPSLPILCSFWDGKTGAAPTTDAGWAALAGSANGKGTSMAPGSTTATCSFTPSYSTTGTATPTLQLIQNSFALDAATTVGFLSPVYTAPAFATVTAASYTISSYLNPVTPVAGGAAAVWRIVITRNAAVTASAKTLTCQMPDNGQGGSPADVTADIAVGGTTTVCVFSIAGYTTATPGPYFATVNVVDGAVTTSHITKNFTVLASGTTAPTYAVTSVVSPATPVKVSTPVTYTFTITRTTAVPAGGIPQPIICEFFNGEGTAPASAAAYWRVSTTIPDADTVVAVMAPGETTTTCTFTTYYTTVSAGGFTAKLMVFGESATAAPLLTSLSVTPSQLLAAVHSFATPMVVAAAVVAVESTTISPNYNPTTPYTNIPTYFTFTLLRDPPVPPSASSGVQFACALYTGQNVNPASAPSAITDAVYKTFTDVTTAVATDANYFADQQLRVVTMAPGTGRVSCTFPTLYAAAGPFSPKFFVFEYASSTVGANALAVADTVTSLTSFTTQAAPTFITGPTNVPQRVPLPKGFRTTCFDGYELIFSNDNYTNGVRVAVDAYPYPVGQCRKCPGGTATMDGYRCIPCPSGYWSNEGARECTACPAGTIAKPAALTARAKYSIDPTTYHFVTHLAMGPESCKKCPKGYFQPNIAGTVCLPCPSGFVSTSGATGCTACSEGTYHTDGVGTTTPGEATSLDTTDTFGSIYPIIPNTCRQCPANTYLPLRGQAAIASMNLAAVSSATPCRPCEDGTWSKAGAAGCQKCPPGTYRNTWFSGQLGSPFITADGVPVATTLTELGSGCSQCPPGTYAPTFGMSVCLPCPAGTFASAPGATACQQCKPGTNSLMGDRTQQMALVVTNAANDFPALRAYTISGMVAGPAYAKPIVTGPDTNFFMAGKSETCSTNLPGYYTDVDGLPIQLPCKPGTFMPFDTATANLLDTGLTVDGTQCYTCQTGTFNDEFSQPVCKACWSGSFASKRGLPTCEIAQPGTFTNVAAAANATFNTATLIPTGLVKGAQAPTPCGMGYFQSSAETTTCTACAVGTYADQAGLAACKPCQPGRYQNSIGQRVCKPCDMGTYSRYGGELCTKCPAGTVASKTGSSQCTPCAAGFYANAPDSATSCRACPRGYYGPYSGAYADNLGDEFEGPRGCYKCPYDFFADRPGVRQCTACPPLDLGGGNLVEQCTEDLGSQRCKPCSLLSKPKTARTEQSPPPPSPSPPPPPPPSPRPPSPNPPSPRPPSPAPPSPNPPPTSPPPSPPPSPPPPRPPPPPPPPPSPPPPNRSPPPPPPASSAINPGGGVNQNGDPVGHRRAILSLMEDEDAEAAQEEQAIVDVDAEMQPQDDE

>Cr22G00420

MGQQIPSQGWQPEPCFPLSRGGQAVPANMLPLHAAAARTRRRLGPDAPINAVVRDVPALIAPLGSMLPWLAAGAGAGAGAGAPSGRKDSAAAGAGATPAMGPQPRSFMVVRLESEQATGQVAPGVSAASVSAGADAGTRGSASGSGEKGEGGSVPAEAAAAAAEAQAAAVPAAVGAASETPSEAPAAERKEGSRGGAATQQASSPVDQPTEAVPAALEAPRGPALSSAPPSLEAAPPAAAATPAPVQRKKGLALRPLLTGPTQDQVAALWHFVLAAITAPPPPPPPPAAATAAPGGEGQAADGGQPGGGAAGDDDDDDDDDDDDEDYDEDDDDDLDDEFAELGLTGGRRRRRNAAPVTRAPLMTAAARAAADVAAAALVLDVLGSNWAGLPEGGLAAGLPYRLVAAAAAAAGGGVDGAAAALGLAAMATEWPASLLRVYLDCTHSGESAGAGAEDDPQGVAATGGLEASRQALALALGWQPPPAGRPSTQQLLTYAPWWYENDLDVEEEADSPGQQQHQQPYGLGSYPVVGSVPLLSPGQTAGVGPVAAPASPAPAAASAAAASPYPSPYKAPSSTALLSAAAAAAAAATAAAARTPGALPVPFPTATAAGMPPGAATAAAARLAVALAPGRGPADPALLAAMVQPRSRRGPAPLSRRSAQPFRRPDDQMVLPPGPLNVARWELVQAPTALPWGAGAMWPLGPHLEPAGQQPQQPQQPQQPQQPQQPQQPLDGVESPAAGVVGSGGAAYVRSPGVVGVAGAGAADAGTGTAAVAPTPAQRHQPGAGAGAGAVDSPVVMEEVQAFETQPEVDGGGKARGAWRSSGGGGGSSGGGGAAAVGPAVVVPTAEDSDPEFPGALSAQASSVGSAATAPAGGLASRRTLSLQRHDSVGGSGAGSGLVTAEASAASIGAASVIAASIATAAMSTAEEDAWVQQERMRRQAQGPGGQEQEPLLPGQVQADGSPPPQQQQQQQQSAVLGLERLPPDLAAAFLQAAAGAKRHGHLRTVQPHGYHTAPSHTGAATYLPQHSQPLQPSQPYFSQSQADPYYGTEAPYGGYAVTTGPHPPAPLQRRPPNAASRWAAVLGMGGPGPAAAASRAEDARIGAYQDFVGVTALGRPVPRSAAGYGVGSGYGGVFPGPGFTGQYS

>Ed2002255

GKESRKRARPGESGRPRPKDRQQIQDRVRELRDIIPNGSKCSIDSLLEKTIKHMLFLQSVSCHADVLKQSRDIKANTGRGTTAALEFHGHGTEX

>Ed2002292

KPAGIQTILCVPAGNGVVELGSTELIPEDPKFIQHIKHSFSHGIWEQAENPTHSNNNHHSVVPSSLTTTPSPLNGHSVEHAAQKLRFVTGNRTVGSFMPPDMGNPGKTAPIVEENDRFRPLVVAPFAPPNPFFTVTAKVFQGNSWHPRQDLEMRKGVDLSKTHQGIKITHQDAKXNLEPRFASGLGGLHSIARIPPPPHLVERDSGKAPESEXXXXXXXXXXGGGGEEPLNHVEAERQRREKLNQRFYALRAVVPNISKMDKASLLGDAIAYIQELQENLKELEIEKEDLQARCAASQSTDVSSNCARIGESDMEVRVEDRDAIIEVCCPKETHPMAKVMLALQNLRVEVRNSSISLAKDSVVHTLSVRIESSGSSSFSKNQLLEAMTGTQGLSSDPDGRIIKM*IR*GFSVNCLLLAHACTAFVKFHYCQKDFPRNKGGNIRGFK*REERTEDKGRGGAVVGCFHHSGRVA*GSAYFCCKLMG*VSVDTYWIMPYASA*LVVSFCSAMTYRMSFSHLFG*LGQLYDVARLATCHSLICSFHTGI*YATACHMFTHFCA*YKPVYGVA*LASCHMSFTHLFGQSGQVYVVAWLATFHSDLLTQSG*LX

>Ed2004062

LHGHGMECPIRVENLKQPYQMLVEMLCEERGLVLEIADTIRGLGLAIFXXXVIESQTGKVWARFIVEATADVHRVDIVLALMHLLQPEGKEWRN*MFLCVF*NCGCFPIGGIYCLATLFIHMRLR*RQMLGQKCVWVAPTWLRXLS*TKSIHLSLHKMDANLAIYRNTRCVIKRKIYAFLFKGGMSSMGMQWWHFL*TRSPFSLLTGRIGI*NKGLRYAFTSY*FSLVLLRRGGILRLWNFVKQALAYCETNIEAFMYLWEHGPCGILVGAVSCVGT

>Ed2008279

ATKKSRATEVHNLSERRRRDRINEKLKALQELVPNSRKTDKASLLEDAIQHMKMLKAQLQMMYARTGMDMPQMLISPGMQQLQMSPFPPPFGFTPLFPPPPPPFTMTMGTPRHHITPRHHKPRHHPKTPP*ICKEIQKQ*TPLCNVVPYYRERFVHYLFKTHLKKKKNHILRKALPRHRKVPRHCKPRRIPKHRLNI***NRKTINSVVX

>Ed2008741

QHGRGENEGEAEEEEGREEG**EGGGRRRGRLC**RGCRRC*CWIFPSIP*RIRASLPYW*AQSTPCRKGKEVSRAIWLQ*RNNIASNPQWGXLNGVTSVDVSLTDGETEDLASLRVPPPPWTRSRHSCNGTEDSGKEAAAAASSSKRKRNHGDGASHRQTVTTGRPKRSRAADIHNESERRRRDRIKEKMRELQELIPNANKMDKASMLDEAIEYLKFLQAQIQMMSMRTGMGVVSPMMLSMGMQHPQMPPVSQMGLGIGMGMMDASTMVHSFKGPNSSYMPNVFHPQPTPFSTSETQYGLQSIGAMDNAVLQQQYILHLQNSLFQPQLLQGNNHGNVAMKKF*KFPLPSKWK*ASIAYVFAKGLVVELFLSILSSSSQFSTCGVTTCNRVIPYKLSIWLLSIAMIESTMVPCFSGFLCPCSIYLVLLSEQ*RDSCCHQNGNMQLYPTCLQRALLLNSLWILSS*YSQCSECGGTMCNQVIAYRFHLIELLR*V*DWKVFGSFLWQMIEKNHSALSCWFPLLVPRLASCYWINNEEIHVVX

>Ed2009884

GG*TLEGRRGFD*RKGKVV*MDLSCSTNCGSGNSSLPLLSCLLQTTLRSLCTQGTNWVYAVFWRILPRNYPPPKWDTAESILDRSSGNKRNWILVWEDGYCDFSACAEDSVNTAHRSSSSSSSSNNNNNYNNNLYYNQNQNNNDNNNYHYPCDNNANRQWSFLQPELFFKMSHEVYNFGEGVMGKIAADNCHRWFSKESTENETLSPSSWQGVHDLQPRTWEAQFRSGIETIAVVAVKEGLLQLGSLRKVEEDLNFVIFVQRKFNYLQNISTYYAITCSSSSSPITTSTPPSFQQKDNNYYYYNNQCPLVPRKTHKENLATTNISQKVDEEDEEEQLTHFEPTASSGYLRQIATSSKAGPVAGGNNQQLINPSRGHVLANRNHSLPVAPRLGMPLYPVQQQNSPEESEAPYSAFWHDLMG*PSVHMCNNHTPLPYVPLRHIACLYNTTSLPCLVYILLIYFPIKFCSSKX

>Ed2010640

FSCMSALSRIDSSMSDISPKLELAAATLKVSNDHRYPSSFLQGFEKLEKLPCLPITMVATNSEEKDTPKSNSASILFSALMEESQGGFKSETSQQSTHTKKSEESGKGSSRKRARPGESGRPRPKDRQQIQDRVRELREIVPNGLKCSIDSLLEKTIKHMLFLQGVTRHADVLKQSGDIKSDTEGGTSGALELCGHEIGLPIIVENLKQPCQMLVEMLCEERGFFLEIADTIRGLGLTILKGIIESRSDKVWARFIVEATMEVRRVDIVFALTQLLRPPGTSQSDLGASVPLVQRTSNGHTNVVGGFQQTSSSSSSSLPLLHANTMLP*ISVFRQSFLFIFPPLSPLNQRQKKGHRNQFYPSLLLTMLCC*MMKRRLASLSLFCSQGDVFNGFAMAFPLNMCPFSLNSRRIGLFMYSRTWPFRTHTVSLWYWQRGDPVLQLQNTRPFRATRVVLFGIFRRGDPVLEVCTGILNALAFEPRHFNIYWKNX

>Ed2010667

RIVPRNFPPPKWETIGGGGGMPDRSKANKRNWILVWEDGYCDFEAITSSSNEEEEGLQPRLFFKMSHEVYNYGEGIVGKVAADTGHKWVYRDPPQMPAVAYPSPPLSQAPWQNAIDPLPRIWEAQFKSGIQTIVLIAVREGVVQLGSTNKVSEDMNIAMELQSNLHFLQTIPFLPSSSTTPSFPLPLVQEQPGLDQLYWNTYNPIASPLPFATGIDDNRNLAGKEEGSLERCLSWYARISTNETNTDTNTNTIMNTITNTNTNPTYHPLQYEQQHALFGNPPPPPPPLINAHAHPSSHTDFLNPSPPLSALLQHPRQSLQHAPGDPAHCHVYYPANVANTTLAADLDPPRVPPNTNMNTGTDVNANANTECIATYVPSVSNNTRYLDSFTALPLPSTLHDPELPKTGK*LAVQGFDLAS*AKKNLWRFG

>Ed2035799

RNWIIVWEDGYCDFDACMRETMMISSVDEEEEEDDGVASSIINNTSLNPELFFKMSHDVYNYGEGLMGKIASDNSHKWIYRDPPTNEMSSVLSPWHTSIDPHPRTWDAQFKAGIQTIAVISVGEGLLQLGST

>Ed2036153

HDVYNYGEGRLMGKTASDNSHKWIYKEPFESDNLAFPNDPHPRAWEAQFSCGIETIAVVAVKEGLLQLGSMKKVVEDLNLVIFLQRKFHALLTIPRRLEEESVSASWIPCHVDCHMDCESMPRGAGFERGCCCVQASSQASP

>Ed2036285

EMLCEERGLFLEIADTIRGLGLTILKGVIESRPDNVWACFIVEPNNSAEESGCLPFK*SSCTFLYLR*FQCSLWNLTSLVK*HFL*YSLRPLCLRYM*IQDYSRQVLEQRFALD*HWSFSPLNTCF*E*QNLLNSPHLTFCQ*FN

>Ed2036976

EVANVVEQMQKEEILMDIFSYTSYIKACCKAGDMQKAVDLMEEMKIMGVEPNMKTYTTIIHGWAWVSYPEKALEYFGKMKAAGLSPDKAVYKCIMACLLSRAAVAYDTVSLPLQAISDEMYARGMCLDLLTAKHWAKLMRQAKERNVELTRALERLFPPSWDEHINTNDX

>Ed2037454

KGNKRNWILVWEDGYCNFPACAKPFNNNNNSNRKLGNFLQPELFFKMSHEVYNYGEGLMGKIAADNSHKWIYKEPTENDPSFPSSWQGALDPQPRTWQAQFDSGIETIAVVAVKEGLLQLGSLKKREEDLNFVIYLQRKFNYMHSIPGLYAMLSPSNEAPNPNHNYRPDNYDPNPDYYDPNQPPRGSHTWT

>Ed2040719

PFCVLVSLLYLFYREGTPLPEPIWAPLSANKDDTLRICQWTREAGSYNRGGSLLILMVYVACWFLSAVLETCSNRRVATSFETSLLNWLRQLR*FELWV*MEEYIEQIFSTPTWVDLNGGTRAPWDFNNAPGVETGALVGNGGSEESMPSLGQATSVASTWRQPYIAGIEPPVSLGVLGQPKAENLSPEEGASNGSHLMGKRSRDEEEDRCGPQENMFGTFMGQPQSARTGALQSMHQLQPMPGVSASAYGNHASLVQSQASGATIAAPARPRVRARRGQATDPHSIAERLRRERIADRMKALQDLVPSANKTDKASMLDEIIEYVRFLQLQVKVLSMSRLGGAGASMPCITELFTEGYSDATMGASNGVQTPSQDGIATTEKHVARLMEDNMGSAMQYLQSKGLCLMPVSLVNKMPTRPPSASGVQDISSSPPSNAVLPTTLLPPLNTNCDGATNDLSKGVRNPPTEHQVHDHLTSPSTSAKSGGNNSKEEPVRRG*PLRVLEWLSRELICLPILCSHRKGVC*SLTVY*VSLTI*N*LNTT*SQLQYRRPPIRX

> Kn001060270

MSLVLQQTLRGLCHNTPWSYAVFWKLKERARIVLTWEDGFYDYAKRNQAPAGEGDEVDTELLGLAVAKMSYHVYSYGEGIIGKIAFSGKHQWVFMGASVNTDEKHPVGWENQFMAGVKTIAVIAVPQGVVQLGSTRVVQEDLALVNHVRGLFTTLRSIPGAFLSDLMAEGPAGAQFNLAAFQTPQWAPPPRPQTRPPQPTLSQNPIQEGPVAATVVPNPSHYFARPPPAGPLPAQSPRGGPGGRPPYPPIAPRGAWMQPNQQMGAGFYTQVTPQLVAEGSPYGVPHPYGVPISAPVYSNPPTPTEGGGSQGGFSMGGMARSQRKGRRRGGETASSGGSRGVPEQTVSEARSSDSTQPRMQRQLSTGRKEPSLGGGYELRGSEVEGVSLTPPSLSDSQLEFARRLSPPGEMPTNSGLMWGLLVQSRPSTSYLRHSVGSLPDRYQTLLMGPHHMGGQPSPPPGRLSLLAPNPIQVQQARVELPPQAILEKGGYFRTHLSKGHVSEKEASPRGDTGCSTVSNAQPDVHRPPLDTRSEGGSTVGESDFPALGKLPEPGHVQSNFSGGSSSLSTQVSAPAPELGKRLPALPNLRVSTGFTKMLSEGQKPTLESVEAYRMQQQAQLESLEKAGLKGTEVYNMWEEIMKENYRVGLALAGFNGEQDAPVTAPLSVQPSPAFPPSSLSNFDNTPATIPLLGRQRSGALQGIPSGGPMQAVSETPLISCGATEQIQNMHLSSGSRSPDPLEVKSDRTDQPEPRIPEGVPAGRVSRTSVRGGGLPRGGPLTGAETVLELAAMYSHSPFLAAVNTVANDELGADGIFRGATNVGLSRMMPTPGLSPTPTDLDLPASETPSPTVDPPPGPSEAFKPRSSPPVPLFFANARTPSPFAGGRPGVPLFNNETIASLPTSSVWAIALKGSGEAAARRRAGLAPPSEPSMSHQGSAATQPMSYSMQACASSDVQANFPSTPSAVRVTSALDVTPPNEAESPRSRSPEPGEMTVMSAGAALESRGGGADFADSPGSRLEHAISAPAGWETPSVSDVSKRTVSKRKGDEFDDRAEASDATGEGGGESERGGARGRKKGRLRAPPDTFSPHELTSTGRPRPKDRQQIQKRLAELRGLVPNTGKLPIDGLLERTIKHLEFLRSVKSEKPVKKELSLPPLLPLLKFDEPQSASEILPGLTANPRKRPALRGSSSLQNPEVNAKLRPERPAFSEASVLNPPSLKPSNSTLNLLNNPPQSPELNLSLLANPHLFSSLQHSLGEPLRTEGPVDTWTRQNAPGGGPGLGRKSSVEQLLSSLHSEPVIVEVLKDPRQLRIEVLCKDESFYAELSGFLHGLGLQVKRSVLQTVANGQQWARFVVETVKPMDRLEVSVPLRRFIELKTGGFEGHLESRAGEAGGSGTQLSLLPPLDLDFGAHFPSSHWQGLEGVLEHDLNMYLDDP

>Mp2002970

PPPTTTTTMPPATTTTTTAAAICKTEPIPPTSAIRSSAIPTTITCSVLACAACKKLPHPFQVSELEQERL*PGPRP*EAAEAKDQVDPNRLLPLPRHGGSHT*VASHLCHWV*DKQNQKRFFVVTPEKRIFLGKGLVMRTRCP*ERLIQGTMFAMHRKDFSPPSPVAKSPRTKCLAQLGNSPISREMVPRLKGCKDKDKRCPRTGAPR*RRRRNPIRIRVWPGGGGPPXRPRVRARRGQATDPHSIAERLRRERIAERMKALQELVPNSNKTDKASMLDEIIEYVKFLQLQVKVLSMSRLGGAGAVAPLVADLPSEGPSSYVSATIGRSNGAPGPSQDGLALTERQVARLMEEDMGSAMQYLQSKGLCLMPISLATAISSTNSRSQGGGAGAGAGAAGTQQQQQGSGDRHRSESGAAG

>Mp2005369

AVVLQHPRTWEAQFKAGIQTVAVISVREGLVQLGSLKKVVEDLNFVILMQRKFNYLQSIPGVFVPHPSPHPSIKRNNSLDGSPPSEAGCWAGNNNNNKDPASMIGFFTRSNSDPFRLTAHSKQLGSGVLGIKRPNEDVPLTLGTGRDFPFLSPAINFSGSPRFNCGQLNRAPECPSPPKALNTGLSGNHSLLPSMSSLHNLLSKLPSVTASENGSPLSSSPSVLSNLSLSNQLQHSQSFSRQPLSPSSSSDSPGLVGSLHSHGGMMGDQDPSPRSSPPHQSCANGSNGMVDSAAGGGGGGGGGAGAGSRHDNGRSVSETSSSQRQPHQQQSSWHNSEYAEQQQQQHHQHQHHHHAQQQQQQQQASGMSDQHHHPHHRPGGNKERLSSFLDTFDPDALSDLGLHDELLENSETYNSFLSEIIS*PPPRSRSGTFFLLTVSSWLYSSTHHDIDSAQT*TEIATRGCVSEGEGGRAIAHEGGESSQSSGSCSTISI*IMYL*DLALDIHHPIPCLPVSSSLMHARTLAAQKDIGRRRASVTFLNRRLSKEVAL*M*IDSG*HADACLMRRDRPNPDEKFKKIIPLFQWKSCLAVSSYANRHPWQLTVNCHPCTRNKRGNIM*EENLS*LTMQLFQARALYISMRHTLAGSRPSTQFVTIKTSPDS*CKGLMRRHTIIPNSFICPDCASICISPVSVSSCAPVWYCGTVVRLSSTLSCFGAFISPCPPMSWSAFSGEHMLFPLSLSHVYTHTSEDIPRCSLARPRPVVKLGQSNKLNMARGTDRPSFRLL*GPH

>Mp2008574

QAAFSSSVLKVTVLEQRIFVKNDCGTTSFGSNSVIRFATDFRCFLRRFAAYWGLPLCRRLESC*MRPVSFMKTRKRFGLQ*SPQA*NVCYDTMAGAGKRNSSTLIEVPKATLRGQLQAAVQSIKWTYTAFWQQSKQKEVLVWGDGYYNGAIKTRKTVQAAELSQEELGMQRTRQLRDLYETLSTAGDGNQSTTRRPSAALTPEDLTETEWFYLLCMSCSFSPENGLPGKALAQGRHLWLTETNEASPEVFTRSLLAKTARIQTVLCIPLCKGVVEFGTTELVREDPALVQHIKTFFAEQFKPVGSEQSSSGPQHQGDTLLQQQQQQKPGRSGANVDQHGASAAWLSTSSPWASVNVSHQDSVVTFSHEAARDSAHGHGHGLQDDAVAMLVDDQKSMGASAKSWDLLSDHIDG

>Mp2008660

YRSSSFSLSHPLSVSNSLSRSPLVSSSSTVYLNNNKNTSSQQQMSYDKSYSQFS*ISSRRQEITATRQALTYIHSNQCG*VKTTAV*KKKQKIQNIKLFKMGQGEQLRELVRSMGWSYAVLWSIPPLMTELVWTDGWYEASNLKVNVERLFNSSYKTCSFAPGYGYVGKVFTQGQHIWVTGDAVHRQSTTSGQATFFSSAGIQTLLCFPWFNGVVELGAEGLVPQNDDLLQQIRRFLSNTPASKLQQQQQHLRGIPAGDDSCHSSRFSSLSPSQGPLTSQGLSSDDAEGGGLSDTTLNQPVDSCPSLSMEDRMRQHFRLTAMSGIDGSCYWAQGPAAVDAEILLPAKSSSMSQVTGAAAGGLHHQGAHSGITGMIGADWRLDFGSDIHQLTSSPTSTLQQAVFTPLPGTEHEEFESSSFLVASSQLNQTAKSLYQQQQHLQQMAAKSPTSSGVEISSNQQQHLRGFLSTKKTGTPAFKPWKGQKQVIGVKRQKSQSNQVLLQRSIKMVHQISLLNQKKDEKKRTEQMALAMRRAPTEAREYHSCHDEAAINHMLAERKRREKQKENFSALRALIPFVSKTDRASILGDAIQYVKQLRSRVQELESINRELEAQIPNSQRKPNSSV*WIDGLID*LIMRSRSHKPPCHIQLPPVVDVQSLSARPYAPMLVTMSQANCTLPFFGSSWQNFGVSREVQIIRRD*MTIDDNPPVHRIALLFITEGTVVAASEPAPFATPTLPTGTLEHLEA*NLLSHLYRSPTRTSHCCSLVCLSNPIPGVYHVQSNFEYTSKEIRILIKA

>Mp2010146

GREPAREG*GGREEAGRQEDIESHIWVQLSRLGV*AESSQQ*HHSPLVVQLS*ARQEEGEREGERVSER*RGGVLKSQGSSGGR*EGHHRRKGASLKESKIHCCRRESEPEHSVA*HCIALHAVAAIARIRGIFAPASLWVGI*ARES*RPRDRDSRGSAAGV**FRGERRKFGELGLKTEKAKGDERKDGLLKPCCDRMRRT*VEPQHLVGALPSHLPRMVVDRKFFRNFSAAVPSFP*SCAFLRAS*RHRIDN*TLE*LSRLSSGTSSEVKRSAVQCSAFPAPGSH*SPTTLLSALLPSEPICNGHHPRHGRWGLTVFRPTVIRSGLFQVLERHQNGKPVPRRLDHHDQQQRQLQQWVVRCQLYSGDGGPGEVPRQSALEVQSGRCSGINGHGSGQMRFFHRDLHSAQSVRAALGRCHRDLRLGRGHGAHK*DAIRKLESGSPQRRIAGSEQARHVPLPFGNFRRQLARRRSPGEQYRELHERWARLERRPRRRRIRRGRLHERGAAVAVKYYSELPASAPAAQQDWHGIARLRRQRPICRSGRQ*QQRLLAQRNGEPAR*SDRNGGECCRRPELVTKYGAGKSGHCSARAAGCEGERKAAADIGHGFEVEARGSGSRRERELEQQEIEVEPAVVADRLARSRSPVSGPXPGLPVPFLAPPLAASLNGMNVNPSSAPFLMSSNVRSIYQMQQPVLAGLGAPADLTKVAGPSAKGGLMANMAPHHLLFGPNFPAIANLDPAAIENSRPKRRNVKISKDPQSVAARHRRERISDRIRVLQRLVPGGTKMDTASMLDEAISYVKYLKLQVQTLESCGNAGFDPRMPYPAYYMPVRSAQGDCSTMSQISPGGGSADLGIYGAGLGQSTMKFPQFCH*SGRSSSSS*GLSLCTIIRLEEADAEDDACFDFLAPFALFRLSGSLSLALSSRDGEIQSAKQSEQVLLMLMMMMEETRMVTVVPL*SL*IEALLQHPPDRTAFLGRHHVAPSIHTPERRSMLIFESRRSIASDRGSWFSFIQRGFFICKSSTHLCRLIGVATTTREIF*PFTFVQIQLDRFIIIIS*PPSTTRGARLDTAPLRSFGVSVVSSAVAQEQTAWVLLHLVSLDSPCTSTPPKSALPFS*AYDITRVQKADCS**VYVVKLKCELQFSLVLILLAGITIEDVLICHT*PFLVQLPPX

>Mp2033447

CSVCDARMLGILEVWSKPRIST*VVLVVLRGRIGHRCMMERQGLPMLNHVLQHTLRGLCCDTQWVYAVFWRILPRNYPPPKWENEGGVMDRSKGNKRNWILVWEDGFCDFSACTAGGGX

>Mp2035313

RVEQSRAVI*HIC*WCRGISLECAGSSYGRVREHQFAVPCFQEVLCEVRPLLEVLSSRRDAESYWIFWCT*ILNER*GGEIYRQGAMAMVLQQALRGLCYKSGWSYAVFWKLKRRSRMVLTWEDGFYEFSKPSAVSSFDSMQTTGGFNMMNSGRPGTDPLAPDVPGAEDEIGLAVARMSYHVYSLGX

>Mp2035326

AGWQQQFTAGIKTIAVVAVPQGVVQLGSTQIVMEDLTLVSHVRGLFGTLQSVPGAFLSDYVPDSQGGRVSVPCPVVMPLPMLIPMGATNETTRSANVTHLHPGQLSMNNVPQIPTTAVVSGSYDMMNLCKASVPARAHLSNPFRTSPVNPLSLQTSRIMPSPHVNKDSTSRNNSLLSTVSMAVGNRK

>Mp2035742

NDTNHKRLHLDTSPNVQTTVAPTTLKRSSSWASSLTSTCSKTVDAMNPGPLPYTKSNSRHAVGPALNTNLKPRARQGSANDPQSIAARHRRERISERLKTLQDLVPNGSKVDLVTMLEKAINYVKFLQLQVKVLTTDDYWPSNEKGAAPCTALPNFEDMEGALKVLVAQAEKEANSNLSNTTSNNGGNSSSSSQENKSPNQADNNSS*QLM*ETPNNRGFE

>Mp2039322

APLGSGNWNCPPSMPSVAQSKTGGLFDRRLHADHRLRHQLQTSLPVCGVKSYQDTVLTTTKGPLQQTSICVTSSPITTGGGVDCKPARTFGQEAFRAMESCNTESTLKTAGFTGGALDTKGLREQSQLYNRNHGELLQGQSRRSSPNVEMSNVEMPSKACMTAPTEGLLSHAGKWAHTVTTGPHSEVSKVAVNNVSQSDDKRKTTTYSQVREDEKVCKQVYVQSPPKFADVAGFSNQSVPISKASGSSLQGIQVCTEGGRNVQVRSQISDSAWNESPKAEWDCFSGLLNPFPIGDELSQALRPASLRECDRDIFSNLHPVHVPDDVRENSSVTTVGETKLLTDSGEGKCDRNTAMDYDIFQAISDAVLMDSRSEPLLEAMVAGVSSAPSAPSPPKVGSCSHEPNLSGGNSTLSGEPESMRQRNDTYRSGSSGDTLEYDTASGISPCKTEMEYEKFASCGIDFQRESMLKASQNSMRMVGDEGQKRTDDELAPYDNVLPAKRQEEFGKVANSRKRPKLGDTLRPRPKDRQQIQDRVRELREIVPNACKCSIDSLLERTIRHMHFLQSVTQHGGNWTNFGGLKEEEDAECLSVDCKAYPDMQGGEKEKSEVGPNCDLKFDKPNNGCPIVVENTNQPRQMRVEMLCEERGLFLEIADTVRGLGLTILKGAMESRSGKTWARFVVEATSRDIHRVEVLWYLMQLIQPTSALLPSYGRPSNFADPGCLYGAAHDSTSGSMTSSFPHMLRPSVPLQMTAR*

>Ol03G00770

VIVIGGLDIARAIASVQAVIEREMASRDELTRRTSATLDIGPRARERDAQEHPLVSHPGAMLQSLSVRSSDDTRVRGNEARGGQKSALAAKTDASSERRHVSPDSALETTGTFGSFESEDGSLEDDDNRMDAANNKRVESMTDFMNSARESQTSAQTMASADLTRSGSRQQIDRLLDEAMSMFHRDGPLVNPLERCASTSSMEMRMRFVCEAKGFDYGLFWEFDEDKKSLTCVSRVCIPDASGISLFVDTSFTMFKRFSMGFGMPGRIGYTGNYEWHEDIRQLPAWSFQRKRQAEQANIRTVIGVPLDDGIVEFGTRSVADHNVTSVQYIQKLCQVSKK

>Ol03G05350

MFPIFVVSMLGTFLIPATIARLASAATKTERGSGEGKGDAASSKKQTQTISTEISDVERTNLWYTLGWIVMIALSVYITRTPLQEKRFDPYDILDLRVGASTKEIKSAYRKLSLKYHPDKNPDPAAAVYFAESIAPAYKTLTDDVARENYEKYGHPDGKQSTKLGIALPEQLFGKGGMAPVMLIVLVVGGIMLPLFIAMCSIRKMNKFGGNNVLKQTQVNYARMLKPVLALSKVPETLAVAHEFIETPFLDGQDAAVSQLLKDYKNEYESKDQKLMKRLPTIIKAHMLILTQTSRRAASLPPVLSADAKKLVLTLPRLIEELLKIAAMPINRAGHSYARPQISVMEFYQCFTQGVPLSSRKRDEDGNASLLQLPHFSTENLNGVAKKCKSLHALMKLSSEDRKKLLIGARFSEAATKDVERQLAVIPRVTTFEAKISVDDDDDTIMEGDFVTAKLKIKIGRSGGPLGGALPPLPFCAADRTEGWRVFVYDQSTNTLLASSTLKQRDVEKAERGDEPLDLSVQFVGLPSGMYNVGVSLMSDYWIGVDAKVSCMMKVLKPTATNVAAREAKSASRSDVTKPNAEVEEEESSESDYSDDDEDDYSDEDYPSDETGTSESDDEARARIDAMRRPEQTKNTVKEGATAAKPASVPQRKTSTFGQKAAAARPSIPGQAVVQQTPATEKPVVAEEVKPDTVDNVD

>Ol04G00550

MESIDAPAQGERIAWDVVASAVNPEGGKFKFTTSTLNFAIKELGERGSFDRAHALYLWMSRKQGRYAPNEFTYVSLASAAKTLTQTRTVQNLWQRGIAENDESLICNEIASAVIAALNRVSDWSGAYQVFRDMGDKGKPRNLYTYTAVLTALRDEAKPDEALAVLNEMAREPGVQPTSLAFSLTLTAFDNCRRWIEGNAVAKRIKKYDVRPDATLMHAIITMAGRAGDMAHANEVFDAMRNSTMIVTTYTFNALLGGYARYGDWEGCTEVYDEMKRSKIQPDSYTFTQLISAAERSGEYIAADGVWTEMLRNRIIPHTVMCGAYIHCLGCQGRDLEAEAVMEKMRNYWDVPRNAAVYNALIGAHVRSGEVTRGLSVLDDMQRIDGLMPTEITFAVLIRACQESALHKRAEGLEGMRASLANAGQLIQDLSGASTSTAKA

>Ol04G04170

MTRDGTFESLGTKASTTTTTTTSGRGDALREAQMDVVLPPLRAAMDSGSASVIAAALGAVQVLISRGLRDESEPSGARNHAGEIVDAICGAAEVRDEAVELQVLKGILTAVSSRTFEVHDRALLRVVRTCYNIYLSSKSEVNQNTAKATLTQMLTTVFHRLEADDPHASAPTIVVADLLRPIGSEAEVDGVTAMSAAVQSFVNKVTTDMNSVGSFNYFSDPDAVVKSDAIEHEITESEFDNDTAPMTPNAVTQSLDAFSPGAMTPARTSGTEQASELETDAFLVFRSLCKLSKKPGSDVNGVALVRSKVLSLQLLKIIIENAGDAFSSSSRFADAMREYLCDAIVSNATPNVPEAYQLACSIFLTLLTRYRAYLKAEIGFFFPMLLLKPLELVEGAPLSAYNQRATLVKGFQIICADSQLMVDLFVNYDCDLDSQNVFERCVLSLVRIAQGVDVSQASGPEAARESVLKLEALECLTTLVASLDDWVRVQSGGDASTSDSQHDVVEESESGFSTPLKTSSPADLGDSIAKLKADKQEFQEGITLFNKKAKKGLAYLQSIGRLGTSHNEIAEFLRTTPGLDKTVVGDYLGERDDPMLQVMHAYVDALDFTSLTLDDAIRKFLEGFRLPGESQKIDRLMEKFAERYHKLNPEVYKSADTAYVLAFSVIMLNTDAHNPQVKNKMTKEGFVRNNRGIDDGQDLPSEVLEDLYDRIVNNEIKLKEPAEVALSAAEKKDKNNFSARLGMDVLFSLMSGKREEETIQIDTADLISQVRARAATTKGFLTVVEAGCAKPMLELIWNPILSLLGTAFEDSESVSVISNCLECFRRVISVTSTLGMQETRDTFIASLTKLTSLHHAHSMRTKNVIAVKTLVRVAIENGNDLGDMWTTILACVSRYEHLYALASGFNDSSLFSESGYSRDDDAQKQARPRLFRRSISSDRALKSPLAPQSSNVNVRDDSSSTVEVEQKFDLLGLDGLNPPDRAVLEQLHPDELDHLFHASVNLSGDAIVGFVRSLCELAIEETSSNHPRAYALGKIVEVASFNMDRIRFIWARVWQVLSDFFVKVGCSPNLQISMQVVDSLRQLAMKFLSRTELANYSFQNEFLRPFVIVMRQSPAVEIRELIIRCVSQMVQARVAHIKSGWKSMFMVFTTAAADESSQIVALAFQTIERIIREHFHYIIETDTVAFTDCVNCLVAFTNSEAGSEVCLNALAFLRFCALKLAEGALGDLEETAATEKQLATDGVVEVTQMKSTVTTTCFTDADAHTYFWFPLLAGLSELTFDPRAEIRTSALEVLFDTLKFHGGSFAPGFWSRVYGRILFPIFDHVRADIMPSTRTIGGDVEYEVAAEDIDDWLYGTCTRCLELVVDLAVQFHEPIVEAGVMPDLLELLCGLASRSHEQLAACGVVAFKRLLINGASSIKEREWHQCMEALKKAFGETTPDFDVFIRG

>Ol06G01040

MTLGELIDRAGELEAKVDGTRANESEYFYIFRELVRCKRLHDSVDLLKHMKERGVKELGRRVSHRDFFSACRSLRVVSVGFEFVDVIESGDIRPYNMLVHACATAGDLQAATLAIEKMKNAGFEPDLQAYTTLLGACSKCGDVERAFEVYAELKRAGFEPNEKTYGSMIDAISRDLATSLKGSRKRRVDSEHVRSTLQSCFMIFEEIKTTNMKLDKIVMNSLLTVCARAAVVPSVRKEACEKVAMVHDEMIERGFELDSYAYQALICCALAEKNYTRAFEYFDEMHDAGINGTTEVYTVMIRAYGKLGKADKAKLIWYAMLEDNIIPDQMSYATMMRLALLDEDDDFCDELMTSMRRNRVRPGPELYSTLTGVAARQGDASQVEEIMQNAKKRGVVAPIECYNSLIAAHARADRPDLAVEAAGKLEAAGYELDAISYEGLIFAYAFARDVEEASNMFERLLESGIRPTFPTFNCLVAAHARSGDMDEACRLVSVMKQHGYVEDSITWRELLLGSVQSGDIEAAWKMYKESRASGNADSERALNTILGQTLVHIRSLTDMKNRSNGKPNEFGSFDDEGDYIAQEWTERAVAAFHEATLAGIKPRVETLSTMLACLRPPSTDEQNAAEYSEVARAVSHETSSHEDAARYYPSQALIMYEEAQGLGIVPKFSRDDEDFVYDIREFPPAAAEVMLLTWLRVVRRRTDAHGLDATIPTMTIRVRADEEVVRMIKEQHMDRIDHSLGRLCKTGERLLTLLRRLRINYGGGLQEGTIELSGHALGRWLQGFVPGDFGNHTGSVFSEHSLSGGVRDQAMRIRANSFGSKDDDVWTPSKMRQAAFNIHDYYGNDDDDPSDFGARPFYPKNWVSQSYVSSYDEDDDDATDLERILGSRK

>Ol06G04110

MARGHARAERGKKARDGDGDGEASGEANARARETNWIASARARRGETILTKTQVAKLAGKLPYGLEFLGVDGPIGHVSREGALYGTSEEEARRWSARAAVALGPTGDDFLPDFGLGVVGVGPSTSKRGMTKKGKANGVGMAARAHAALAADPFAALVVDVEDPVRPPSRGIGGIGVGGGTTSQSKRARAEDRSKRVALRKRLIELENLVNNLGKRATDLVERRDELSLIAQTANASVNEIDAKRKVSSVSRGVSAIRQKLACKPDQQHGFNTLRYRCLLEIVHKQCLSAVRQLMAHKWGFPFSAPVDPDALGLPTYREIITEPMDLGTIKKLIENGGKYVMAEEVDADVRLTFANAMKFNNEGTDVHTMACGLLDEWEPKWEAIKQRIADVEACVLVERDMAVAKNEAAQRRADVVSKEKECAKASEALDLVSMQLREVETQVLALMRPLQREDRLDLASDLRCLPESLRSGAKDIIAANTTGWSAQAHLEDIDAHNEITLHLLARYTKTMNRNRLAVVAGWCGNATPEHLLEKLKQEQDVSADTVNFVIGQPLDDATSGLGIRSIDHMQDEFDFDPSAFNDLLNGVPLDVDGDAIDIDIGAGDHMDLDL

>Ol16G00630

VGAYVSEGRVEEAERVLREMTAEGVRAGPRLFNTLITGYGREKNLRGVEASSMAMRTLGVTPNQATWGAKVNAYVSCDRLDLAMDTLEQGVRFSPRVERRPGVQAYTALVQGLAHSGRVVEADELLRRMARDGVKPNVYTYSTLIDGLAKSAQIGLAETALAEMRRAKIKPSVVTYNSLLKGVVRGIGKADRTDEVLRRAREMFDRMRDDGVPPDLVTYNTLIDACINARAPAEAWNILREISESGLKPDVVTYTTLLKYFVQVGDDSATQWVIAELETDPQVVEDVGVYNCLINAYARQGDMCRAVETLEGMKAKNITPNVSTYGSILEGYIRLGNVGEAFKVYNLCVKSAGLAPDARMRKSLIYGCGLHGMSDIA

>Ol21G00370

MKTTPNCAPNIRSYNGVISAATRKKHFPGAMWAWEEIETANLQPTMITYGAMLAAGAAADDVDVRWSEDLFAQALESGACGNAGNDHMVTSMLQTYARGVALEQIERDVAMERGESVVQALIEDAQWDERRAESTPNGRVWSALITLCARCGRAARAIEVLKIMISTRSHRVGHEWLHLTYALTSALEASKEGIEYFARVQREIEQSPAMVRDCTGVRNGLIATHLHFGDFKSAFKVYDDFKNDIFKYRKAHEENWNARARRGLERQLPDTITYNSLIYACADDDVKAMGLYHDMVSNGINPTVRTYVALIVALSRSKRGSKVTEAEKIFKAAIDDGVTPNEFLFTALMDAQVKGNRPLSAFETYARMIEADVNCTTVTFGCALQACCYVEDVEESVERAYSVLRDMTERDVQMNDWCSNTFLRVISRAGRIEEMLEEVKKTVRRKGKLEQETLEAIIRALCSAGYVERANRFISMMNSRNLEPREQTFKEFIVASSRDGFVDWAWESYKRFTRLGHKLDAGTRSALVTVLSVASTSPDPDDAELLLARAIGVFEAAFKRADEDNGPQVLDVIDAEARCALIVAMARSEKLDRALDIWRDSPKAQSFSKARKHTSSEGNDYIGDVRAMYECLIEVCCHEDRIDDALEVFDHLKDAGVRVSTVTLAFLESSCRRCRVEEWRMFDVCAQMRAQVEQKNEGRLAKPTKMSHHVRDDGNIASELATDGLGGEQKTSAWRKNVD

>Os01G02110

MMAAQASSKRGMLLPREAVLYDDEPSMPLEILGYHGNGVGGGGCVDADYYYSWSGSSSSSSSSVLSFDQAAVGGSGGGCARQLAFHPGGDDDDCAMWMDAAAGAMVENTSVVAGGGNNYCHRLQFHGGAAGFGLASPGSSVVDNGLEIHESNVSKPPPPAAKKRACPSGEARAAGKKQCRKGSKPNKAASASSPSPSPSPSPSPNKEQPQSAAAKVRRERISERLKVLQDLVPNGTKVDLVTMLEKAINYVKFLQLQVKVLATDEFWPAQGGKAPELSQVKDALDAILSSQHPNK

>Os01G10090

MRALSPALPNSYTLPLALRAAASPRVASAVHAHALHLGLHAQHDVAGQILAAYSRLGRAADARRVFDAMPPGRTTFHWNALISAYSSGCDPDAARDAFARMAAAGARPDAVTWTALLSAHARSGKHADVLQLFGEMQRSGCEGNAESMAVALSACPYAGDLALAKGKAIHGCGVVKGLMHGYLFVTNSLICMYGKLGEMDDAKKAFRDATAKNTVTWNTLITSYAAARLCDKALDVLAQMEQIGGTVAPNVVSWSAVIGGFASSGDTDRALELFRRMQQQWLSPNVVTMATVLSACVDLLALRLGRELHGHAMKAELDRHSLVENGLINMYAKCGKVSGARKVFDGMKTRDLISWNSMLAGYGMHGLCDEALALFTDMAGATVEPDGVTFVAVLSACGHAGRVTEGRRLFDRMVRAHKISPSMEHYTCMVYLLGRAGLLRDASELVETMPVRPDLCVWGALLNSCRIHGDAAMAEATIANVLQSEDQSTGNHMLITNLYAMCGMWDESKKVRVMTKEAGLRKNPGQSWIEVDNKVVAFAAGSAPPNLTGAEDVFGMLDDLYAEMEDEQR

>Os01G25270

MRSAGTISQQLTRYAAAQALLPGAHLHASLLKSGSLASFRNHLISFYSKCRRPCCARRVFDEIPDPCHVSWSSLVTAYSNNGLPRSAIQAFHGMRAEGVCCNEFALPVVLKCVPDARLGAQVHAMAMATGFGSDVFVANALVAMYGGFGFMDDARRVFNEADSERNAVSWNGLMSAYVKNDQCGDAIQVFGEMVWSGIQPTEFGFSCVVNACTGSRNIEAGRQVHAMVVRMGYDKDVFTANALVDMYMKMGRVDIASVIFEKMPDSDVVSWNALISGCVLNGHDHRAIELLLQMKYSGLVPNVFTLSSILKACSGAGAFDLGRQIHGFMIKANADSDDYIGVGLVDMYAKNHFLDDARKVFDWMFHRDLILCNALISGCSHGGRHDEALSLFYELRKEGLGVNRTTLAAVLKSTASLEAASTTRQVHALAVKIGFIFDAHVVNGLIDSYWKCSCLSDANRVFEECSSGDIIACTSMITALSQCDHGEGAIKLFMEMLRKGLEPDPFVLSSLLNACASLSAYEQGKQVHAHLIKRQFMSDAFAGNALVYTYAKCGSIEDAELAFSSLPERGVVSWSAMIGGLAQHGHGKRALELFGRMVDEGINPNHITMTSVLCACNHAGLVDEAKRYFNSMKEMFGIDRTEEHYSCMIDLLGRAGKLDDAMELVNSMPFQANASIWGALLGASRVHKDPELGKLAAEKLFILEPEKSGTHVLLANTYASAGMWNEVAKVRKLMKDSNIKKEPAMSWIEVKDKVHTFIVGDKSHPMTKEIYAKLVELGDLMSKAGFVPNVDVDLHDLDRSEKELLLSHHSERLAVAFALLSTPPGAPIRVKKNLRICRDCHVAFKFISKIVSREIIIRDINRFHHFRDGTCSCGDYW

>Os01G45840

MRYLYFHGNASQARPIRHHLLAYLDACASRAHLAELHGRLVRAHLTSDSFVAGRLIALLASPAARHDMRYARKVFDGMAQPNAFVWNCMIRGYSSCEAPRDALAVFREMRRRGVSPDNYTMAAVVSASAAFAGLKWRSNGDAIHALVRRIGFTSDVFVMSGLVNYYGAFRSVEEASKVFEEMYERDVVSWTSMISACAQCGHWDKVLKMLSEMQAEGIIPNKVTIISLLSACGQTQAVDEGRWVYNQPWCCP

>Os01G64560

MAGAAPLRDSLRRLCTDVGWSYAVFWRATRAADSQRLKLVWGDGHYERAAGAPSISGFEAMDLLLKEKAAALRSGTGRGGGGGEGHAADGAAGHSHDRVDALVHKAMAQQVHVVGEGVIGQAALTGLHRWIVHDIVDECEEEDEVLLEMKGQFCAGIQTIAVIPVLPRGVIQLGSTKMVMEEAAFIDHVRSLFQQLGSSTAVVPCGSFVQDSIMRTPFHKSLGVPTSSHSEDLAGGGNTYNDDMINHQFRHQKSPASTIQSFNPVQQFYAGPTFCRPVTIASRCDLFQPDHGSTFTLNSQSEDNRSTALLKNSVSHSKTSNDAFSHAFNPLNEPNVSISGRRECVSIEQHGSCRNGEMEITIGRTASSSCTGKTNIINKVDDLLSQDCLVGCQASNATSVNRKFQTMSIVDNTKLQDGSYAIPHAALVDSTQYSDCFQSLLGTIQGSSSSNSNAIHVDTSHNAVHGKSNFCPLGDRNAANSSDLAELLASPIPLELTGGNDLFDVLQLQQKPNGSNNSEVNNRQSMPYGSEQAVKSLIGCVDDDFTGLITEADPDQLLDAIVSKIITGHKQNVDTSASCSTTVAGFDRPLHSDCHLYTTGPSSGPIFCNFASVAPVAIKTEGPAAGSRQSSSSIDKSAGCSQTQESYKSQIRLWVENNHSVGSDSLSTGQASDSLSTGQCKRSDEIGKSNRKRSRPGESARPRPKDRQMIQDRIKELREIVPNSAKCSIDTLLEKTIKHMLFLQNVAKHADKLKGSGEPKIVSHEEGLLLKDNFEGGATWAFEVGTRSMTCPIIVEDLNPPRQMLVEMLCKERGIFLEIADQIRGLGLTILKGVMEVRKDKIWARFAVEANKDVTRMEIFLSLVHLLEPSTGSSILSAGVENTSLPRDSFFPSSIPASGFSNCL

>Os01G72930

MPPPPPFPSLDAFYLHLLRACTSLRHAAAVHAHIARAHPAASLFLRNTLLAAYCRLGGPLPARRLLDEMPRRNAVSFNLLIDAYSREGLAPLSLETLARARRAGVDVDRFSYAAALAACSRAGHLRAGRAVHALAILDGLSSGVFVSNSLVSMYSKCGEMGEARRVFDVAEERDDVSWNSLVSGYVRAGAREEMVRVFAMMRRGGMGLNSFALGSVIKCCSGRGDGTMDIAEAVHGCVIKAGLDSDVFLVSAMIDMYAKKGALVEAAALFRSVQEPNVVMFNTMIAGFCRTETVIGKEVASEALTLYSEVQSRGMQPTEFTFSSVLRACNLAGYLEFGKQIHGQVIKYTFQEDDFIGSALIDLYFNSGCMEDGFRCFRSSPKHDIVTWTAMVSGCVQNELHEKALSLFHESLGAGLKPDLFTISSVMNACASLAVARAGEQIQCFATKSGFDRFTVMGNSCVHMYARSGDVDAATRRFQEMESHDVVSWSAVISCHAQHGCARDALHFFDEMVDAKVVPNEITFLGVLTACSHGGLVDEGLRYYETMTKDYGLSPTIKHCTCVVDLLGRAGRLADAEAFISNSIFHADPVIWRSLLASCRIHRDLERGQLVANRIMELEPTSSASYVILYNMYLDAGELSLASKTRDLMKQRGVKKEPGLSWIELKCGVHSFVAGDKSHPESSAIYTKLEEMLSRIEKLATTDTEISKREQNLMNCHSEKLAVALGMIHLPQSAPIRVMKNLRVCRDCHSTMKLISKSENREIILRDPIRFHHFRDGSCSCADYW

>Os02G01610

MTHAAHALCNRYAAILSSAAGDGGRTGVRVAGAVHCLILKTFLQAPPTFLLNHLLTAYAKSGRLARARRVFDEMPDPNLFTRNALLSALAHSRLVPDMERLFASMPERDAVSYNALITGFSSTGSPARSVQLYRALLREESVRPTRITLSAMIMVASALSDRALGHSVHCQVLRLGFGAYAFVGSPLVDMYAKMGLIRDARRVFQEMEAKTVVMYNTLITGLLRCKMIEDAKGLFQLMVDRDSITWTTMVTGLTQNGLQLEALDVFRRMRAEGVGIDQYTFGSILTACGALAALEEGKQIHAYITRTWYEDNVFVGSALVDMYSKCRSIRLAEAVFRRMTCRNIISWTAMIVGYGQNACSEEAVRAFSEMQMDGIKPDDFTLGSVISSCANLASLEEGAQFHCLALVSGLMRYITVSNALVTLYGKCGSIEDAHRLFDEMSFHDQVSWTALVTGYAQFGKAKETIDLFEKMLANGLKPDGVTFIGVLSACSRAGLVEKGCDYFDSMQKDHGIVPIDDHYTCMIDLYSRSGRFKEAEEFIKQMPHSPDAFGWATLLSSCRLRGNMEIGKWAAENLLETDPQNPASYVLLCSMHAAKGQWTEVAHLRRGMRDRQVKKEPGCSWIKYKNKVHIFSADDQSHPFSSRIYEKLEWLNSKMAEEGYKPDVSSVLHDVADADKVHMISHHSEKLAIAFGLIFVPQEMPIRIVKNLRVCVDCHNATKFISKITGRDILVRDAVRFHKFSDGTCSCGDFW

>Os02G16650

MASRAPCLLAARGIASSPHLARRLKQTENEIVQMFRTPSPRNEDAVAALSPRYTNSVRVLDERFIRILKIFKWGPDAERALEVLMLRVDHWLVREVMKTDVGVNVKMQFFRWAAKKRNYQHDTSTYMALIHCLELVEQYGEMWKMIQEMVRSPICVVTPMELSQVIRMLGNAKMIGKAITIFYQIKARKCQPTAQAYNSMIIMLIHEGQYEKVHELYNEMSNEGHCHPDTVTYSALISAFCKLGRQDSAIRLLNEMKENGMQPTAKIYTMIISLFFKLDNVHGALSLFEEMRYMYCRPDVFTYTELIRGLGKAGRIDEAYHFYHEMQREDCKPDTVVMNNMINFLGKAGRLDDGLKLFEEMGVSHCIPNVVTYNTIIKALFESKSRVSEVFSWFERMKGSGISPSPFTYSILIDGFCKTNRIEKAMMLLEEMDEKGFPPCPAAYCSLIDALGKAKRYDLACELFQELKENCGSSSARVYAVMIKHLGKAGRLDDAINLFDEMSKLGCTPNVYAYNALMSGLARACMLDEALTTMRKMQEHGCLPDINSYNIILNGLAKTGGPHRAMEMLTNMKNSTIKPDAVSYNTVLSALSHAGMFEEAAELMKEMNALGFEYDLITYSSILEAIGKVDQE

>Os02G35660

MGGFAYPFTPSPAWSRDAVFAGSPWAAGGVSSLADALVSYGAVDDEEAAFLGKTAASSPSTARLHEQQQLLLEAELLRHGDGLGFAAMDDDGGAAMLGALEPCAMPLTDSGGPPVICSSSSNDSSGSEHSAAMPAGGGFLVGEQQQHVPPAAYAAGGVLPSMAAGEETPQSFGFGSLFNGDLLQEATVSKYHHHQQQQQLGVVPSSQPHHLNDDIDFNTGKLMSFASGQQHVTPSIDSLQIDQKEFSSGLHHLNLSSLISGPLASFNATQSHRQPAEACGGKNGGAAPFVNLSEVLPKGNGSGSAGNGAPKPRVRARRGQATDPHSIAERLRREKISDRMKDLQELVPNSNKTNKASMLDEIIDYVKFLQLQVKVLSMSRLGAAEAVVPLLTETQTESPGFLLSPRSSSGERQAGAGAVTGGLPGDQPELLDGGAMFEQEVVKLMEDNMTTAMQYLQSKGLCLMPVALASAISAQKGTSSAAVRPEKKKNGDGDGGGDEEDVKGEFDAPRRPPVGRPKEMRSRV

>Os02G45170

MAESLGLGALCRGGGWCYAAIWRSDRRDPRLLTIGEFHSEDGTRNVVEKMLNQVHVVGEGIIGRALVSGECQWISDTSFSFAQTSDADNQDLFQGYTWWQHQFLCGIKTIAVIPIADLGVAQFGSMQKISECLEFLDQVKGIFCQREIVPWDLSAEEIQRNVLPYHQQFQLSSLSSADGLTNIKTDPENKKLLENSASVESLRSLASFSSKYSQSSSNGFTSYESCNSMNPHIVAMPVNSKSINTVRAFNSTGKLLQHNIGSENPLQIKFCQHPDSNLASATDVFLSLNNLPRIENEISCPPNKLGYCIQSEKPYSFQSSFSSCFSVGDELKPILFDSATSFVQNDLMQEFNLTGFTSQADSAVHELPKQILGETATGALYSDRKSNNGSSDLLDGTIFDPFVQEWCDNNALLEGNTPHFGATTADSVTEHASSYPLSVEERSLFSESVFEELLGVSGNVNTDAPGDSAVVMAGDPLVGLVSGCQLPTYTLQDSLSVCKPQQEPSLDFPSGSDTSEHVPNGSSKMIPLSLGALSMDDCCSLNTAHSKVSQVKRPEEVKVVKKRARPGESTRPRPKDRQQIQDRVKELREIVPNSAKCSIDALLDRTIKHMLFLQSVTKYAEKIKQADEPKMISNKDSGAVLKENSSGVVLKDNSSAGSNNGGATWAYEVAGRTMVCPIIIEDLSPPGQMLVEMLCEERGFFLEIADTIRGFGLTILKGLMELRDGKIMARFLVEANKNVTRMDIFLSLVQLLQQNSLNRSSDQISKVIRNGVPSFAEHQQSPISVPVGLADR

>Os02G48060

MRMALVRERAMVYGGGCDAEAFGGGFESSQMGYGHDALLDIDAAALFGGYEAAASAGCALVQDGAAGWAGAGASSSVLAFDRAAQAEEAECDAWIEAMDQSYGAGGEAAPYRSTTAVAFDAATGCFSLTERATGGGGGAGGRQFGLLFPSTSGGGVSPERAAPAPAPRGSQKRAHAESSQAMSPSKKQCGAGRKAGKAKSAPTTPTKDPQSLAAKNRRERISERLRILQELVPNGTKVDLVTMLEKAISYVKFLQLQVKVLATDEFWPAQGGKAPEISQVKEALDAILSSSSPLMGQLMN

>Os02G50280

MDSCRLLHPYPQPLPLPPPPPPPTSTPRPTHLQWGVLRRRRRRHHFLRCVAASAATLQKELTVPRTPTAAQSPGPVNPPTLFDRMPERSVATVSAADNLLDEMSRTCGAGQRGRPLEAPPRDGGGKSASAAIVALAHAGRHAEVVELFCRMRRGGVPVSRFVLPSVLAACAGLRDIGMLRAVHALVIKCGLCQHVIVGTALVDGYTDFGLVDDARKAFDEITDANIVSWSVLIGGYARSSRWEETLDAFSAMRRAGVLPNDSVLVMAIQACGALGRLVHGKQLHGLAVVLGFDRNATVWNCLMDMYGKCGDIDSCKMVFETMIGRDQVSWNTLISSYARVGLCEEALDMIVQMQESGYIVDRFTLGSGVTACARLADIDSGRAFHGYLVRRLLDTDVIQGSALVDMYGKCHNMELAHIVFDRMDERNYVSWDALLSGYVENEQVDLALEIFRQMGCANIKYNQHNFANLLKLCGSQRYKEYGRQIHGHAIKTINKMNVVLETELIDMYAKCGCIEVARLLFLRMNERNLISWNALLSGYAADGQPVATINIYRQMELACIRPDKYTLAGLLSLCRYQGLLHYGRQIHAHLIKMGSEMNVVMQTILVHMYIKCMRQQDAENVCIMIEERNSYVLDAFSKVYGDDYLI

>Os02G50900

MEEQLNPLAVTQLLQHTLRGLCTQGDSQWVYAVFWRILPRNYPPPKWDLQGGVYDRSRGNRRNWILAWEDGFCNFAASACDQEDTPAAAGYTDYAAAGHEVKGLQPELFFKMSHDIYNYGEGLVGKVAADHGHKWVSQEANEHEINLVTSWNNPADSHPRTWEAQFQSGIKTIALIAVREGVVQLGSMKKVAEDLSYVVALRRKFGYLESIPGVLLPHPSSAAFPGAGGLQDAAWAPSPTMDLYDPYYGAHAAAAQMHHIVPSMSSLEALLSKLPSVGPTAAPGAIRGAIGGGSVAKEELDDAMDAAGNGGGESTSAATTPLVPYYVDVAKPDEGF

>Os02G55020

MPAWCSKGSGAMARGRGRWLLPTRLLNVCLAALCRGGSLAAAESVLVDAIRLGLPPDVVTYNTLLAAHCRAAGLEAGLVVMGRMREAGVEPDAVTYNSLIAGAARRGLPIHALDLFDEMLRSGIAPDSWSYNPLMHCLFRSGHPEDAYRVFADMAEKGIAPCDTTYNTLLDGMFRAGYAMNAYRMFRYLQRAGLPVSIVTYNTMINGLCSSGKVGYARMVLRELGRTDHAPNIITYTAVMKCCFKYGRFEQGLDTFLSLLDRGYISDVYPYCTVISALVKKGRLGEANNYCDLMLQNGSRLDSVCYNTLIHMRCQEGKLDDAFELVSMMEDGGLESDEYTFAILVNGLCKMGHIEAAEKQLFYMEIKGMQSNVVAYNCLVDALCKFQEVDAAIRLLQCMKLKDDFTYTSLVHGLCRVGRYHMASKFLRICLHEGNNVLASAKRAVIAGLRSSGFKNDLRKVRVALNMAKLLRP

>Os03G10770

MEDSEAMAQLLGVQYFGNDQEQQQPAAAAPPAMYWPAHDAADQYYGSAPYCYMQQQQHYGCYDGGAMVAGGDFFVPEEQLVADPSFMVDLNLEFEDQHGGDAGGAGSSAAAAAAATKMTPACKRKVEDHKDESCTDNVARKKARSTAATVVQKKGNKNAQSKKAQKGACSRSSNQKESNGGGDGGNVQSSSTNYLSDDDSLSLEMTSCSNVSSASKKSSLSSPATGHGGAKARAGRGAATDPQSLYARKRRERINERLKILQNLIPNGTKVDISTMLEEAVHYVKFLQLQIKLLSSDDMWMFAPIAYNGVNVGLDLKISPPQQQ

>Os03G27390

MGEKVNPWCHWSNPPWTESSANNLHPPDVSLDNTNSVALPTYLNSDGYIYSGVAASMPSIAASVTDRPVSFSSRFVTTLVPSVGLSTAETLRKRPLVFFHNVNNTFTVGPLLSKGTLDTVPELQGSNETNVTDVGAQNTECMHENTEEIDALLCSDSDEGCLKVQELNNRVRKYPMQNDTMSVESVASAGASQPAKKRRLSSGTDRSVVDTASSARPDHSVDQKHLSHDDDAQSCCIGEVESDHQFALREGEEAEGDDGPDDRKRRRERIQETVAALRKIVPGGIAKDATAVLDEAICYLKYLKLKVKTLGAVSL

>Os03G42100

MESGGVIAEAGWSSLDMSSQAEESEMMAQLLGTCFPSNGEDDHHQELPWSVDTPSAYYLHCNGGSSSAYSSTTSSNSASGSFTLIAPRSEYEGYYVSDSNEAALGISIQEQGAAQFMDAILNRNGDPGFDDLADSSVNLLDSIGASNKRKIQEQGRLDDQTKSRKSAKKAGSKRGKKAAQCEGEDGSIAVTNRQSLSCCTSENDSIGSQESPVAAKSNGKAQSGHRSATDPQSLYARKRRERINERLKILQNLVPNGTKVDISTMLEEAMHYVKFLQLQIKLLSSDEMWMYAPIAYNGMNIGIDLNLSQH

>Os03G43470

MARACSSPSSPSPSPASSRPLLPSSASISAFLASHPALTLLHTQCASMAHLRQLHAALVKSGLARDPIAASRAVAFCAGDGRDAAYAARLVRHHPRPNAFMWNTAIRALADGPGPGAAVALFVDMLGSPTPPERRTFPSLFAAYARLGRAGDGAGLHGMVVKLGLGGDAYVRNSVIAMYASRGAADEAIALLARCEAFDAVACNSAIVALARAGRVDEARAVFDGMPARTVATWSAMVSAYSRDSRCHDAVELFSAMQAEGVEPNANVLVSVLGCCASLGALEQGAWVHAYIDKHDVAMNALVVTALVDMYCKCGDIRKAREVFDASRSRGQAKLSSWNSMMLGHAVHGQWREAAALFSELRPHGLRPDNVTFIAILMAYGHSGMADEAKAVLASMASEHGVVPGVEHYGCLVDALARAGRLREAEGAIAAMPVAPDAAVWGALLSGCRLHGDAEAAARAAREAVRCDPRDSGAYVLAASALARGGEARRGAAVRGRMREEGVGKVPGCSMIEVDGVVHEFVS

>Os03G55550

MAVDWIWERRRREEEYNHQMSQDELQQPGQVQWTPAPEEKSEIAVQFFTAPYPCQNGQLDHGEHHALGGIGACSSVHWQPDRGTCYWPPPLSGDGGGGSGSGSSGTGEGSYIGERCYYVGEPDVPIGLNLLVGDNDGAGVVLRDAAPQAKRRTQAGHGGDLGRQKKKARVSDKRNQESMQSGSCSDNESNCSQVNRRKVDRVAGGGNGKVPARRRSATIAQSLYARRRRERINGRLRILQKLVPNGTKVDISTMLEEAVHYVKFLQLQIKVEVQIVCHDQMLSSDELWMYAPIVYNGMDLGIDLNISPPR

>Os04G35610

MARPHHTGIHLVSHLRASAPLADLLRSAPGLRAARAAHARALRSPFAGETFLLNTLLSAYARLGSLHDARRVFDGMPHRNTFSYNALLSACARLGRADDALALFGAIPDPDQCSYNAVVAALAQHGRGGDALRFLAAMHADDFVLNAYSFASALSACASEKASRTGEQVHALVTKSSHGSDVYIGTALVDMYAKCERPEEAQKVFDAMPERNIVSWNSLITCYEQNGPVDEALALFVRMMKDGFVPDEVTLASVMSACAGLAAGREGRQVHTRMVKSDRFREDMVLNNALVDMYAKCGRTWEAKCVFDRMAIRSVVSETSMITGYAKSANVGDAQAVFLQMVEKNVVAWNVLIATYAHNSEEEEALRLFVRLKRESVWPTHYTYGNVLNACANLANLQLGQQAHVHVLKEGFRFDSGPESDVFVGNSLVDMYLKTGSISDGAKVFERMAARDNVSWNAMIVGYAQNGRAKDALLLFERMLCSNERPDSVTMIGVLSACGHSGLVKEGRRYFQSMTEDHGIIPTRDHYTCMIDLLGRAGHLKEVEELIENMPMEPDAVLWASLLGACRLHKNIDMGEWAAGKLFELDPDNSGPYVLLSNMYAELGKWADVFRVRRSMKHRGVSKQPGCSWIEIGRKVNVFLARDNIHPCRNEIHDTLRIIQMQMSRMSIDAEIADDLMNFSSEACG

>Os04G47080

MEETPLPSGKNFRSQLAAAARSINWTYAIFWSISTSRPGVLTWKDGFYNGEIKTRKITNSMNLMADELVLQRSEQLRELYDSLLSGECGHRARRPVAALLPEDLGDTEWYYVVCMTYAFGPRQGLPGKSFASNEFVWLTNAQSADRKLFHRALIAKSASIKTIVCVPFIMHGVLELGTTDPISEDPALVDRIAASFWDTPPRAAFSSEAGDADIVVFEDLDHGNAAVEATTTTVPGEPHAVAGGEVAECEPNSDNDLEQITMDDIGELYSLCEELDVVRPLDDDSSSWAVADPWSSFQLVPTSSPAPDQAPAAEATDVDDVVVAALDSSSIDGSCRPSPSSFVAWKRTADSDEVQAVPLISGEPPQKLLKKAVAGAGAWMNNGDSSAAAMTTQGSSIKNHVMSERRRREKLNEMFLILKSVVPSIHRVDKASILAETIAYLKELEKRVEELESSSQPSPCPLETRSRRKCREITGKKVSAGAKRKAPAPEVASDDDTDGERRHCVSNVNVTIMDNKEVLLELQCQWKELLMTRVFDAIKGVSLDVLSVQASTSDGLLGLKIQAKFASSAAVEPG

>Os04G58980

MERRRMIADLLRASARGSSLRGGVQLHAALMKLGFGSDTMLNNNLIDMYAKCGKLHMAGEVFDGMPERNVVSWTALMVGFLHHGEARECLRLFGEMRGSGTSPNEFTLSATLKACGGGTRAGVQIHGVCVRTGFEGHDVVANSLVVMYSKGRWTGDARRVFDVIPSRNLATWNSMISGYAHAGQGRDSLLVFREMQRRHDEQPDEFTFASLLKACSGLGAAREGAQVHAAMAVRGVSPASNAILAGALLDVYVKCHRLPVAMQVFDGLERRNAIQWTTVIVGHAQEGQVKEAMCLFRRFWSSGVRADGHVLSSVVAVFADFALVEQGKQVHCYTAKTPAGLDVSVANSLVDMYLKCGLTGEAGRRFREMPARNVVSWTAMINGVGKHGHGREAIDLFEEMQEEGVEADEVAYLALLSACSHSGLVDECRRYFSRICQDRRMRPKAEHYACMVDLLGRAGELREAKELILSMPMEPTVGVWQTLLSACRVHKDVAVGREVGDVLLAVDGDNPVNYVMLSNILAEAGEWRECQGIRGAMRRKGLRKQGGCSWTEVDKEVHFFYGGGDDAHPQAGDIRRALREVEARMRERLGYSGDARCALHDVDEESRVESLREHSERLAVGLWLLRDGTGDDGGGGGGEVVRVYKNLRVCGDCHEFLKGLSAVVRRVVVVRDANRFHRTPRVFFLDCLLPATLAVAPSLMDHPAKVMMGWALIAKYHLPQPSSSDQAQTGPYRLVFALYVSTARPDCRQSPSDMAPGKQRGKAKGAPPPPAAPNAAAAGGFPACLRLMPPSTVAISIQAKPGSKLATITEIGDEAVGVQIDAPARDGEANAALVDFISSVLGVKKREVSIGSGSKSREKVVLVQDATLQGVFDALKKACASA

>Os05G36350

MPPPPSSSAPWQLEDAVMARLRACVTFRDLLRVHGHVVRLRISQSSYLATQIVHLCNAHRRVTHAARVFAQVRDPNLHLHNAMIKAYAQNHQHRDAVAVYIRMLRCPTSPPDGHAGGDRFTYPFLLKACGGTAALELGKQVHTHVVRSGCDSSAIVQNSLIEMYTRAGDLALAHKVFDEMRERDVVSWNMLISAHARLGQMRKATALFNSMPDKTIVTWTAMVSGYTTVGDYPGAVDAFRSMQTEGFEPDDVSIVAVLPACAQLGALELGRWIYAYCKRHGMLTSTHICNALMEMYAKCGCIDQALQLFDGMADKDVISWSTVIGGLAAHGRAHEAVWLFTEMEKEGKVRPNVITFVGLLSACSYAGLVDEGLSHFDRMNDVYGVEPGVEHYGCVVDLLGRSGQIRRALDLVRDMPVPADAKVWGSLLSACRSHGDVDTAVLAAERLVELEPDDVGNLVMLANVYAAARRWSDVASTRKAIRSRSMRKTPGCSLIEVGNVVREFVAGEGLSSELGGLAGVLDILASHLADDEEDIDFADSDCTVYANLAND

>Os05G49920

MPPPPARTHPNPPLLHLLASHRAPQPLPLTPAHGHLPPRKRPRGVGSAAAPPPPRAAASAEATYSDRSAALRALCSHGQLAQALWLLESSPEPPDEGAYVALFRLCEWRRAVDAGMRACARADAEHPSFGLRLGNAMLSMLVRFGEIWHAWRVFAKMPERDVFSWNVMVGGYGKVGFLEEALDLYYRMLWAGMRPDVYTFPCVLRTCGGIPDWRMGREVHAHVLRFGFGDEVDVLNALVTMYAKCGDIVAARKVFDGMAVTDCISWNAMIAGHFENHECEAGLELFLTMLENEVQPNLMTITSVTVASGMLSEVGFAKEMHGFAVKRGFAIDVAFCNSLIQMYTSLGRMGDAGKIFSRMETKDAMSWTAMISGYEKNGFPDKALEVYALMELHNVSPDDVTIASALAACACLGRLDVGIKLHELAQNKGFIRYVVVANALLEMYAKSKHIDKAIEVFKFMAEKDVVSWSSMIAGFCFNHRSFEALYYFRYMLGHVKPNSVTFIAALSACAATGALRSGKEIHAYVLRCGIGSEGYVPNALLDLYVKCGQTSYAWAQFSVHSEKDVVSWNIMLSGFVAHGLGDIALSLFNQMVEMGEHPDEVTFVALLCACSRAGMVIQGWELFHMMTEKFSIVPNLKHYACMVDLLSRVGKLTEAYNLINRMPIKPDAAVWGALLNGCRIHRHVELGELAAKVILELEPNDVAYHVLLCDLYTDAGKWAQVARVRKTMREKGLEQDNGCSWVEVKGVTHAFLTDDESHPQIKEINVVLHGIYERMKACGFAPVESLEDKEVSEDDILCGHSERLAVAFGLINTTPGTTISVTKNRYTCQSCHVIFKAISEIVRREITVRDTKQLHCFKDGDCSCGDIGYG

>Os06G09370

MDYSAGSYMWPGNSGSENYNFVDGSSESYAEEGSLPPSGYFMGAGSDRSLKITENERNPTMLANGCLPYNTQAHPLSGQILPKGELPNNLLDLQQLQNSSNLRSNSIPPGVLQCNSTSGTFDAKLDTPGLAELPHALSSSIDSNGSDISAFLADVHAVSSAPTLCSAFQNVSSFMEPVNLDAFGFQGAQNVAMLNKTSLPNGNPSLFDNAAIASLHDSKEFLNGGSIPSFGTVLQALGAGGLKAAQQEQNIRNIPLPTFTSGSHLAVTDAQGPPLPSKIPPLIHDHNSEYPINHSSDVEPQANSAPGNSANAKPRTRARRGQATDPHSIAERLRREKISERMKNLQVLVPNSNKADKASMLDEIIDYVKFLQLQVKVLSMSRLGAPGAVLPLLRESQTECHSNPSLSASTISQGPPDMPDSEDSSAFEQEVVKLMETSIISAMQYLQNKGLCLMPIALASAISNQKGMAAAAAIPPEK

>Os06G10820

MEFDMAMDMMSQEQLMHIISQLDSALASSPSPSTSPSASPPRQSPAAHVPVPPGLLNTTMVSTSRAQAAPSAPLHPVAATAAVQSSSRGIMYTTTRQGVIDAAEEEEAAAPRPRRRNARVSSEPQSVAARLRRERVSQRMRALQRLVPGGARLDTASMLEEAIRYVKFLKGHVQSLERAAAALHMHGGHAAAAGFAGDAGDAVYSCPSYYA

>Os06G12740

MDEQLSPVAVTHLLQHTLRSLCTSGDDSQWVYAVFWRILPRNYPPPKWDLPGGAYDRTRGNRRNWILAWEDGFCNFAATSAACGDGAAAAYAAAECEETKQVGVAGGGLQPELFFKMSHDIYNYGEGLIGKVAADHSHKWVFKEPQEQEINLISSWNNPADSHPRTWEAQFQSGIQTIALIAVREGVVQLGSMKKVAEDLSYVVALRRKFGYLESIPGVLLPHPSSAAAAFPGGPPDAAGWPAGMMVSPPVPPELYVDPYGGAAAGAVPPPSMQIMPSMSSLEALLSKLPSVVPAAAAPSPPPGSSSMPPTGAAAASSAPPKEEAAEDDYVHCHGMDMATSSTNGGGESTGGAPLPSSYFVNVGVKPSEGF

>Os06G30090

MAMVAGDEAMSVPWHDVGVVVDPEAAGTAPFDAGAGYVPSYGQCQYYYYYDDHHHHPCSTELIHAGDAGSAVAVAYDGVDGWVHAAAAATSPSSSSALTFDGHGAEEHSAVSWMDMDMDAHGAAPPLIGYGPTAATSSPSSCFSSGGSGDSGMVMVTTTTPRSAAASGSQRRARPPPSPLQGSELHEYSKKQRANNKETQSSAAKSRRERISERLRALQELVPSGGKVDMVTMLDRAISYVKFMQMQLRVLETDAFWPASDGATPDISRVKDALDAIILSSSSPSQKASPPRSG

>Os07G01130

MQCHAAAAAAAATVPAYCALIRPLASAGRVDAVDAAVASARSRLSPATIHPLYVASIRAYARAGRLRDAVDAFERMDLFACPPAAPAYNAIMDALVDAAYHDQAHKVYVRMLAAGVSPDLHTHTIRLRSFCLTARPHIALRLLRALPHRGAVAYCTVVCGLYAHGHTHDARQLFDQMLHTHVFPNLAAFNKVLHALCKRGDVLEAGLLLGKVIQRGMSINLFTYNIWIRGLCEAGRLPEAVRLVDGMRAYAVPDVVTYNTLIRGLCKKSMPQEAMHYLRRMMNQGCLPDDFTYNTIIDGYCKISMVQEATELLKDAVFKGFVPDQVTYCSLINGLCAEGDVERALELFNEAQAKGIKPDIVVYNSLVKGLCLQGLILHALQVMNEMAEEGCHPDIQTYNIVINGLCKMGNISDATVVMNDAIMKGYLPDVFTFNTLIDGYCKRLKLDSALQLVERMWEYGIAPDTITYNSVLNGLCKAGKVNEVNETFQEMILKGCHPNPITYNILIENFCRSNKMEEASKVIVKMSQEGLHPDAVSFNTLIYGFCRNGDLEGAYLLFQKLEEKGYSATADTFNTLIGAFSGKLNMHMAEKIFDEMLSKGHRADSYTYRVLIDGSCKTANVDRAYMHLVEMIKKGFIPSMSTFGRVINSLTVNHRVFQAVGIIHIMVKIGVVPEVVDTILNADKKEIAAPKILVEDLMKKGHISYPTYEVLHEGVQSTIYCLEYGKQRASDPKCRKTTEGCVEINVDGAQLRIMLTNLKVSPVWQKVSPQDNIFICELRILTWGDVYVDKVITEIKGDLYDSPIDSKNQIVMSTLYNNDQYQSYPLCPIEAALLSMSSHTYSLGEELIGKVALTGQHCWISSYEFSSTFMYKYHEDWQLQFAAGIKTILFVPVVPHGVLQLGSLDLVPESSTSVALIKDLFYKLYDASISGSPSGTGFGYSNTGRQPAAMLPMDSPDVVPHNFFRSIKSSAQLLNNDHLSLLHAFPVLEFASTEDSIVSIYDTSLTACAVEPLDGNDSDIWTNVHEELSQFTRCNTASEADKANISYMDKLINSDSKMSCRSVSHAEDPGYGNIDHFILTEMERENQEHINNYTSVNDYAVTSNPSFHSELHKTLEPISREEREDCMWHIRYRQQESTSSALLQENGNKAGFDKQCENNDYAELLLDAINDQVIWASNSESSHSTDSPVSCATQIQKDDHVPRLDESSVPNFPGGQDFSLISIDEGFMSCTMTGSSLTETNKAILVEEDFISDPIEGMHRETVEIKGRCRKTGLHRPRPRDRQLIQDRMMELRQLVPNTSKVQLLALFDVSPSTKDVLFICLFVLSFGNEEANGLLTVSSFYKLKQCSIDSLLDKTIAHMQFLQCVSEKADKMVCEEYGVFLEIAHVMKDLEVTILKGLLESRSDKLWARFVIQNDLHCETEGVVVITTTTEIQKRLIRIDVMHLTGFTRLRPDADSIPTDAPVAEAEMELSSTIIHLPWVCMLIIVGLFNNSMLGSGLIE

>Os07G39940

MAQFLGAHGDHCFTYEQMDESMEAMAAMFLPGLDTDSNSSSGCLNYDVPPQCWPQHGHSSSVTSFPDPAHSYGSFEFPVMDPFPIADLDAHCAIPYLTEDLISPPHGNHPSARVEEATKVVTPVATKRKSSAAMTASKKSKKAGKKDPIGSDEGGNTYIDTQSSSSCTSEEGNLEGNAKPSSKKMGTRANRGAATDPQSLYARKRRERINERLRILQNLVPNGTKVDISTMLEEAVQYVKFLQLQIKLLSSDDTWMYAPIAYNGVNISNIDLNISSLQK

>Os08G01640

MSRRVLSAPDGTKVSRFNLMKLQGRQEAAAAGTDSHHAHHAFDELLHRPTTSSIVDLNRALSDAARHSPAVAISLFRRMVMVARPKVPPNLITYSVVIDCCSRVGHLDLAFAALGRVIRSGWTAEAITFSPLLKALCDKKRTSEAMDIALRRMPVLGCTPNVFSYTILLKGLCDENRSQQALHLLHTMMVADDTRGGYPPDVVSYNTVINGLLREGRQLDTAYHLFDQMLDQGLSPDVVTYNSIISALSKARAMDKAAVVLVRMVKNGAMPNRITHNSLLHGYCSSGKPNDAIGVFKRMCRDGVEPDVFTYNTLMGYLCKNGRSMEARKIFDSMVKRGHKPNSATYGTLLHGYATEGSLVKMHHLLDMMVRNGIQPDHYIFNILIGTYTKHGKVDDAMLLFSKMRRQGLNPDTVTYGIVMDALCMVGKVDDAMAQFGRLISEGLTPDAVVFRNLIHGLCARDKWDKAEELAVEMIGRGICPNNIFFNTLLNHLCKEGMVARAKNIFDLMVRVDVQRDVITYNTLIDGYCLHGKVDEAAKLLEGMVLDGVKPNEVTYNTMINGYCKNGRIEDAFSLFRQMASKGVNPGIVTYSTILQGLFQARRTAAAKELYLWMIKSGIKFDIGTYNIILLGLCQNNCTDDALRIFQNLYLIDFHLENRTFNIMIDALLKGGRHDEAKDLFASLLARGLVPNVVTYWLMMKSLIEQGLLEELDDLFLSLEKNGCTANSRMLNALVGKLLQKGEVRKAGVYLSKIDENNFSLEASTAESLVLLVSSGKYDQHINAIPEKYRPVVKTRAV

>Os09G07780

MRSAGTISQQLTRYAAAQALLPGAHLHANLLKSGFLASLRNHLISFYSKCRRPCCARRVFDEIPDPCHVSWSSLVTAYSNNGLPRSAIQAFHGMRAEGVCCNEFALPVVLKCVPDAQLGAQVHAMAMATGFGSDVFVANALVAMYGGFGFMDDARRVFDEAGSERNAVSWNGLMSAYVKNDQCGDAIQVFGEMVWSGIQPTEFGFSCVVNACTGSRNIDAGRQVHAMVVRMGYEKDVFTANALVDMYVKMGRVDIASVIFEKMPDSDVVSWNALISGCVLNGHDHRAIELLLQMKSSGLVPNVFMLSSILKACAGAGAFDLGRQIHGFMIKANADSDDYIGVGLVDMYAKNHFLDDAMKVFDWMSHRDLILWNALISGCSHGGRHDEAFSIFYGLRKEGLGVNRTTLAAVLKSTASLEAASATRQVHALAEKIGFIFDAHVVNGLIDSYWKCSCLSDAIRVFEECSSGDIIAVTSMITALSQCDHGEGAIKLFMEMLRKGLEPDPFVLSSLLNACASLSAYEQGKQVHAHLIKRQFMSDAFAGNALVYTYAKCGSIEDAELAFSSLPERGVVSWSAMIGGLAQHGHGKRALELFGRMVDEGINPNHITMTSVLCACNHAGLVDEAKRYFNSMKEMFGIDRTEEHYSCMIDLLGRAGKLDDAMELVNSMPFQANASVWGALLGASRVHKDPELGKLAAEKLFILEPEKSGTHVLLANTYASSGMWNEVAKVRKLMKDSNIKKEPAMSWVEVKDKVHTFIVGDKSHPMTKEIYSKLDELGDLMSKAGYIPNVDVDLHDLDRSEKELLLSHHSERLAVAFALLSTPPGAPIRVKKNLRICRDCHMAFKFISNIVSREIIIRDINRFHHFRDGTCSCGDYW

>Os10G35790

MADDGRCPPDVVSYNTIIDGLFKEGDVDKAYITYHEMLDRRVSPDAVTYNSIIAALSKAQAMDRAMEVLTVMVMPNCFTYNSIMHGYCSSGQSEKAIGIFRKMCSDGIEPDVVTYNSLMDYLCKNGKCTEARKIFDSMVKRGLKPDITTYGTLLHGYASKGALVEMHDLLALMVQNGMQLDHHVFNILICAYTKQEKVDEVVLVFSKMRQQGLTPNAVNYRTVIDGLCKLGRLDDAMLNFEQMIDKGLTPNVVVYTSLIHALCTYDKWEKAEELIFEILDQGINPNIVFFNTILDSLCKEGRVIESKKLFDLLGHIGVNPDVITYSTLIDGYCLAGKMDGAMKLLTGMVSVGLKPDSVTYSTLINGYCKINRMEDALALFKEMESNGVNPDIITYNIILHGLFRTRRTAAAKELYARITESGTQLELSTYNIILMDFAKTNSLMMHFGCFRTYV

>Os10G40920

MGKCAARQRQWRWPLLHRSPRPTPPPPHGLHPPRRALAEHARMPAEQQQPPPVAPARVTFSRVFQSCAQAGREALAAGRAAHARMVVSGFVPTAFVSNCLLQMYARCAGAACARRVFDAMPRRDTVSWNTMLTAYSHAGDISTAVALFDGMPDPDVVSWNALVSGYCQRGMFQESVDLFVEMARRGVSPDRTTFAVLLKSCSALEELSLGVQVHALAVKTGLEIDVRTGSALVDMYGKCRSLDDALCFFYGMPERNWVSWGAAIAGCVQNEQYVRGLELFIEMQRLGLGVSQPSYASAFRSCAAMSCLNTGRQLHAHAIKNKFSSDRVVGTAIVDVYAKANSLTDARRAFFGLPNHTVETSNAMMVGLVRAGLGIEAMGLFQFMIRSSIRFDVVSLSGVFSACAETKGYFQGQQVHCLAIKSGFDVDICVNNAVLDLYGKCKALMEAYLIFQGMKQKDSVSWNAIIAALEQNGHYDDTILHFNEMLRFGMKPDDFTYGSVLKACAALRSLEYGLMVHDKVIKSGLGSDAFVASTVVDMYCKCGIIDEAQKLHDRIGGQQVVSWNAILSGFSLNKESEEAQKFFSEMLDMGLKPDHFTFATVLDTCANLATIELGKQIHGQIIKQEMLDDEYISSTLVDMYAKCGDMPDSLLVFEKVEKRDFVSWNAMICGYALHGLGVEALRMFERMQKENVVPNHATFVAVLRACSHVGLFDDGCRYFHLMTTHYKLEPQLEHFACMVDILGRSKGPQEAVKFINSMPFQADAVIWKTLLSICKIRQDVEIAELAASNVLLLDPDDSSVYILLSNVYAESGKWADVSRTRRLLKQGRLKKEPGCSWIEVQSEMHGFLVGDKAHPRSGELYEMLNDLIGEMKLSGYEPDSASFVEVDEEGSAPEHCLELLVANLL

>Os11G06010

MAVGDALRRLCEEVGWSYAVFWKAIGAADPVHLVWEDGYCGHASCPAGSDPSEALPTDVGCAAAADTMTMCSLVNKVMASQVHVVGEGTVGRAAFTGNHQWIIHGTANDHGIPSEVAAEMSYQFRVGIQTIAIIPVLPRGVLQLGSTGVVLENKSFMTHAKKLCSQLNNRSSMAVSSSVKNSSSQQGRSRPLHGASNVQSTENRSKLFSQFPVTCEQYNHPDTMAVSGSTSLNACMNGSLLKIAQLNGQAVREHIVYSKPDVRFIQQVYRDGQLGSNAQSIAMSSDLISSSLRSVQKQPLLMNNISQLEYGDGAETSADLRKNVLLKPPVCLDPFIHDRNINISHGITEVSNVINDHGNFDFLSGGARVVRANLCTSATSQVLDRRSHSVSGMLLHREPIVSCEVPQSSEFSTKMGSLERGSFQISSAPSSESDVQISNGLNTSISRENQLSVSNHICQDQKINGVNDLSATLSTERMNNMDGCKPPGLSLERTSPLFMEQSVENDLFDILGPQFHHLCHNAGADLVPWTDAKPESSDRDVPESSIHADSAPLFSSRDNELYSGIFSLTDTDQLLDAVISNVNPAGKQSSDDSASCKTSLTDIPATSYLCSKEMKQCGSSGVPSVLIKNESAQFIKQPCLAENAEDGCLSQNNGMHKSQIRLWIESGQSMKCESASASNSKGLDTPSKANRKRSRPGESPKPRPKDRQLIQDRIKELREMVPNGAKCSIDALLEKTVKHMLFLQSVTKHADKLKDSTESKILGNENGPVWKDYFEGGATWAFDVGSQSMTCPIIVEDLDRPRQMLVEMICEDRGIFLEIADFIKGLGLTILRGAMEARKSKIWARFTVEANRDVTRMEIFLSLVRLLEPNCDSSGAAENANNVNMPLGLVHQPVIPATGRIQ

>Os11G25920

MVGSGAAGGGGGGGGGGDHARSKEAAGMMALHEALRNVCLNSDWTYSVFWTIRPRPRCRGGNGCKVGDDNGSLMLMWEDGFCRPRVAECLEDIDGEDPVRKAFSKMSIQLYNYGEGLMGKVASDKCHKWVFKEPSECEPNIANYWQSSFDALPPEWTDQFASGIQTIAVIQAGHGLLQLGSCKIIPEDLHFVLRMRHMFESLGYQSGFFLSQLFSSSRGTSPSPSSFPLNRRRRRRTRGTAARRTWRST

>Os11G41640

MDARCANIWSSADARSEESEMIDQLKSMFWSSTDAEINFYSPDSSVNSCVTTSTMPSSLFLPLMDDEGFGTVQLMHQVITGNKRMFPMDEHFEQQQKKPKKKTRTSRSVSSSSTITDYETSSELVNPSCSSGSSVGEDSIAATDGSVVLKQSDNSRGHKQCSKDTQSLYAKRRRERINERLRILQQLVPNGTKVDISTMLEEAVQYVKFLQLQIKLLSSDDTWMFAPLAYNGMNMDLGHTLAENQE

>Os12G06330

MAVGDALRRLCEEARWSYAVFWKAIGAADPVHLVWEDGFCGHASCSAGSEASEAGCESGGAVCTLVRKIMASQVHVVGEGTIGRAAFTGNHQWIVHETANDHGLRSEVVAAEMNNQFRAGIKTIAIIPVLPRGVLQLGSTSVILENISSVQQYKKLCCQLNNRSSMVASASAKNDLSQKVQSRSLHGLPSIHPYEQCYGHDARALSSSTSANTGRNTSLLKVAQRNDQAIREQVLYAPDMRFRQQLPYSDRRVDINTHSSAMSSGFISSISASVEKYPLLTNNIGQVEHGNMEESSGPRNVLLKSLSCRNPVVHENTNTSLFHGGDEVPAFLNSHGSFDFLQAGPRVVEANLYNNGTSSQVLDQRCSSTSGMAGYKPSVSYKFPHSAQFIVKMENPRRQSFQDPAAPSSGSDVQVSSGLKTTTRQFNPEHMCQNKKTNEVNDSSAAVSTQDVKNMDRHKILDISNERTSSFVMDPSTENDLFDIFGTDFHQLHRSLDGDLSWNTAKPQSSDRDAPESSIYLDSSPAFGAQEDEFSYSGIFSLTDTDQLLDAVISNVNPGGKQISGDSASCKTSLTDIPSTSYCGSKETKQCKSSGAPPLLIKNELAVSNFVKQPCFLEKAEDGCLSQNNGVQKSQIRLWIESGQNMKCESVSASNSKGLDTANKANRKRSRPGESPKPRPKDRQLIQDRIKELRELVPNGAKCSIDALLEKTIKHMVFLQSVTKHADNLKDSNESKIHGGGENGPLLKDYFEGGATWAFDVGSQSMTCPIIVEDLDRPRQMLVEMLCEDRGIFLEIADFIKGLGLTILRGVMEARKNKIWARFTVEANRDVTRMEIFLSLMRLLEPSCDGGGGGVGDNPNNVKIPPGIVQHPVIPATGHLR

>Os12G06335

MAVGDALRRLCEEARWSYAVFWKAIGAADPVHLVWEDGFCGHASCSAGSEASEAGCESGGAVCTLVRKIMASQVHVVGEGTIGRAAFTGNHQWIVHETANDHGLRSEVAAEMNNQFRAGIKTIAIIPVLPRGVLQLGSTSVILENISSVQQYKKLCCQLNNRSSMVASASAKNDLSQKVQSRSLHGLPSIHPYEQCYGHDARALSSSTSANTGRNTSLLKVAQRNDQAIREQVLYAPDMRFRQQLPYSDRRVDINTHSSAMSSGFISSISASVEKYPLLTNNIGQVEHGNMEESSGPRNVLLKSLSCRNPVVHENTNTSLFHGGDEVPAFLNSHGSFDFLQAGPRVVEANLYNNGTSSQVLDQRCSSTSGMAGYKPSVSYKFPHSAQFIVKMENPRRQSFQDPAAPSSGSDVQVSSGLKTTTRQFNPEHMCQNKKTNEVNDSSAAVSTQDVKNMDRHKILDISNERTSSFVMDPSTENDLFDIFGTDFHQLHRSLDGDLSWNTAKPQSSDRDAPESSIYLDSSPAFGAQEDEFSYSGIFSLTDTDQLLDAVISNVNPGGKQISGDSASCKTSLTDIPSTSYCGSKETKQCKSSGAPPLLIKNELAVSNFVKQPCFLEKAEDGCLSQNNGVQKSQIRLWIESGQNMKCESVSASNSKGLDTANKANRKRSRPGESPKPRPKDRQLIQDRIKELRELVPNGAKCSIDALLEKTIKHMVFLQSVTKHADNLKDSNESKIHGGGENGPLLKDYFEGGATWAFDVGSQSMTCPIIVEDLDRPRQMLVEMLCEDRGIFLEIADFIKGLGLTILRGVMEARKNKIWARFTVEANRDVTRMEIFLSLMRLLEPSCDGGGGGVGDNPNNVKIPPGIVQHPVIPATGHLR

>Os12G32400

MECSSFEAICNESEMIAHLQSLFWSSSDADPCFGSSSFSLISSEGYDTMTTEFVNSSTNVCFDYQDDSFVSAEETTIGNKRKVQMDTENELMTNRSKEVRTKMSVSKACKHSVSAESSQSYYAKNRRQRINERLRILQELIPNGTKVDISTMLEEAIQYVKFLHLQIKLLSSDEMWMYAPLAFDSGNNRLYQNSLSQE

>Os12G39850

MEGGGLIADMSWTVFDLPSHSDESEMMAQLFSAFPIHGEEEGHEQLPWFDQSSNPCYYSCNASSTAYSNSNASSIPAPSEYEGYCFSDSNEALGVSSSIAPHDLSMVQVQGATEFLNVIPNHSLDSFGNGELGHEDLDSVSGTNKRKQSAEGEFDGQTRGSKCARKAEPKRAKKAKQTVEKDASVAIPNGSCSISDNDSSSSQEVADAGATSKGKSRAGRGAATDPQSLYARKRRERINERLKTLQNLVPNGTKVDISTMLEEAVHYVKFLQLQIKLLSSDEMWMYAPIAYNGMNIGLDLNIDT

>Os12G40590

MMSFPYSSGDLGEATTAAAAAVDMITLDQMFRDYDASTGDDLFELVWESCGGGEIDSGAGEVQPAGVPCCRRLLPGSSPEPTSEDEMAAWLSTIVTGSGGGGGDDVAAGGDHQDPAVKKPDGEPLTEKMDKKLPTRTEERRRVKHKARRNPGYAETHGLTEKRRRSRINEKFKMLQRLVPGCDKCSQSSTLDRTIHYMKSLQQQLQAMYPTMVRPAAVYPVVQPPPAFAAGGPPAASQGGLHRHRRLMVVVSVHHRCRCFRLGRQ

>Pe2000963

PVCPFAYDSVLRLSNERLSV***YTLNI*PVTIHDGELGLVIAICNSLQIVVAALG*SVIYIYIYSSVLLIRMDGGGLPVVNYLLQHTLRSLCSDSQWVYAVFWRILPRNYPPPKWDTEGGIMDRSKGNKRNWILVWEDGYCDFSGCSRAANSLPNRCSTTISGEGPEEMMTESFQPELFFKMSHEVYNYGEGLMGKVAADNSHKWVYKEPPENDINFLSSWHGSLDPHPRTWEAQFKSGIQTIAVVAVREGLIQLGSLHKIVEDLNFVIFLQRKFNYLQSIPGVFAIHPPLSFDLRRSKMASSPSQENPSCEGFNNHGDQHDKICPSPAKDHSHPHTQLVGMKRPPDQFIDMSKQNTMPVLYGSCLSPNKAINAGWNSRQASINNNTIVLPLMQPSMSSLQALLSKLPSVTSAEQTSSQFPASLEPVRASSLNARHPVTNPLETSEASSFQKSCLTLNSTAHISTSPSAHADFADSKPHVRHQEATCSNSSSHHGLCPNFLDSCSNPLQFHEPHGDLETSDDAYHSLLNQIIS*FFHPDMNF*A*L*LFFGILPSIFSLQQHYMRIGIWRVARISVRLLFNIV*GEFMSFVDRSRKAIRIFMTILLFFSVNVRVIIQARQTILLESSMQAWFLSDVWCAIKYIAVIIR*

>Pe2006376

VNCNIYISLMQACTNIKQLNGIYALMYRTGFHDNIVLVNRLVRLYVMYNRMDDARRVFDKIHKPNIFLWNMMIRGYATNGFCEEASNSITKWRK*VYRRTLSRLILCSRPVLGCCL*EKG

>Pe200907

1IILHMI*AIDLGGRENTATGYFTQIREIGTLTVNLRSICISRFGMGTSAFAGAVSP*TCGHCEMAGTNMIYPNFQQPHPTRGTPCGHCEMAGTNMIYPNFQQPHPTRGTPGGTSLSCSSALCLCVPYSHRFHRHYTPTTKTCNNSSRHDTFTKSHNNFSRYHRFVSANTSLYLLFCHVGAVDIYQRKYPTKNGFNPSKSLRAKASANALVVQQEDTVNKICKAIEEDQWECLQDNFLNSMAQDLNPDAVVKVLNLQTDAQNALRFFQWADKQEGYDHNTDAYFTMIDILGRAKMFTELQSLLQKMQTQGCEITRSMLHSFVMSYGRSGRFKESLEAFNLMKEMGYEPGLIDTAYNSVLVSLVKNKKLDMAENLFAQMIHNGVSCNNLTYTSMIQCFFLKEKMEDAMKLLDDMIQNNYAPDVVTYTIVISALCKRKMIEQAYGVLQKMRENGCEPNIYTYNALIQGLCAVRRPEEALELVTLMEQGGVPPNIYTYTILIHGLCKVRRLDRAKEMFNEALARGLKPNRVTYNTLLNGYCRGSRLIEAMDILKEMHQNDCTPDRVTYTTLIQGLVQGNQLPDALRMHDEMENKGYDVNFDTLNILARGLARVGNHKDASIFYRRMKDRGFAYSASDYYLAIHCLSTAGEMEEAQALLYEMINKGYSPNLTTYNTMIKGFCRQGRLDDADAMLNFMIENGIGPXXXXXXXXXQGRTQDADQLYATALERGVVLNPKPVIQEPDELPEGLVRSQEELVRGCDA*GLPFCILTCCFSSVVG*RWITLNKDA*LILVPE**LTTQLNELILLWK*IVDSFASTKYCILSRSLLCLP*MSLFRFFFTYFFFRSLISLFKLYCRKTLGLQKIVDYQQLSTLDIS*I**LSAVYSYFPHVI*VVAQTRWDGS*KRFIL*QKNX

>Pe2036703

SIDALLLRTITYIKFLHTITQHGGRWSYKGKIKIARNEAMTVLSEESKHGASWVVDLGAH

>Pe2039132

PAGWKNQFAAGIKTIAVIAAPYGVVQLGSKQTILENLELVSHIKALFRNLHNVVGAFSRDSIPELLGGRIHGTX

>Pe2044419

NGHIWARFVVESVRDIHRLEVLWSLTHILESVANKRENTGISN*GA*AWT*NTEIVPGYFDLPRSIFPCGG*KR*QVLFSHSLHIKKHLYEFCQ*MIWVYIFLYQCX

>Pe2045356

SLCSNSDWNYAVFWKLKHHLNRMILTWEDGYYESTKPPGISNLSTLTAPDSLLKGWDTGGNQLDYGSHSSDGCSMEDQIRLFMTQMSHQTYSFGEGIIGHSASTGKHRWVFQENCX

>Pe2050082

SPGSINAKPKQNSGLSSVLTNILKKSLKIH*SESLYC*YS*HTRKMVGSSDRAKEALAMMALHEALRNVCVNSEWTYSVFWTIRPRPRSRGGSGNGCKVGEENGSLMLMWEDGYCRSSKPRVNRVPGPFNGNGCMDKLVKSSSFDCLEELDADDPVRKTFRKMSIQLYNYGEGLMGKVASDKCHKWVFKEPTECETGISNYWQSSFDAHPPEWTD

>Pe2050436

RRCGCIPTVETYNVLVYGLVQKKQIESALKVIEDMVLAGISPNERTYTTLMQGYASSGDIGRAFEYFTKIKNEGLRLDELSFAALLKACCKAGRMQSALAVTKEMSASGIQRNTYIYNILIDGWARRGDVWEAADLMQKMIQDGISPDIHTYTSFVNACSKAGDMNRAMAALEEMASLGMKPNIKTHTALINGWALASYPDKALMCFEEMKQAGLEPDQAVYHCLMTSLLSRG

>Pe2050967

LV*MAGAGAATLPVPSLALLPSNQNPHHFFSSNTFYRHSNSRIPLKKSSRQYQFIGKSQRDSITFGSELKPKPKPTSQGQEEYNDVSLDICEKIRKLCRQEKFQEALDIFQLMQHRGIEVDIDTYALLLQGCIVTKSIEDGRRVHAHMFESGVESDIYLSNKLLIMYTKCGKMSDACQLFEEMSERNSVAWNAMIAGYTQHGQSKQALELFCQMRKDGVEVDKYTFSTVLRACARVAVIEAGEQVHACIVKLDFHKNVYV

>Pe2051696

DFTERENRSATTGLDGRQGPLLTNYARSQPGQQNTGLQVLQSSQGNSLQQQGNRSVISQDQLAATAGAVLNGAPRTRVRARRGQATDPHSIAERERRERIAKNLKSLQELVPNANKTDKASMLDEIIDYVKFLQLQVKVLSMSRLGGAGAVAPLIADVPAEGSGSLASAALGQAGGSLSQDGLAFEQNVVKLMEKDMTAAMQYLQNKGLCLMPIALATAISSSTGKPLLGSGVTNSVDSTSDRQSSGNQPASLTVDSCAGALGMGSAGFGSDFLLKENTVEKRGRELVPTAESNGAEFGIISX

>Pe2054114

RMAGR*GRKYGNLFIYGILRIEIDRPLGMGRGSEGIVWDS*WDRFLLS**DQPLMTENPIISLEGWVCLLMCEMEETQGLPVATHLLQHTLRSLCMGSQWVYAVFWRILPRSYPPPRWDAENGIYDRSKGNKRNWILAWEDGFCDFPECARAAMESNTGPGLQPELFFKMSHEVYNFGEGLMGKVAADNSHKWIFRDPLEHEINFLSPWHNSIDPHPRTWEAQFKSGVQTIAVIAVREGLVQLGAIQKVIEDLNFVIHLQRKFNYLLAIPGVFLPHPSSSLQSKTTDDQAGSWQLQGLVGGESSSGRHLEKCFSWNGTTMMMDQNRHNATEFMQQQMSSGAGPVLIDTLQQQLPVTPSMSSLQALLSKLPSVTPSQYTNYNNISSPNVLARSLSIPRHHYSSELTPTSMLMEPQQFQLLPPAAAASRSHDFIEEFHDDLTHNDETDGAAACTGMDRCYLSEVVD*TRHV*ACVMFFSFITFQLVSPPSNLVX

>Pe2055480

NLLSLRESSKVSEAAFGELLPVTDEYGHNGFDDFDKYLKSLMKGQDNSNSSISLPAADEIFDVLGPVCRNGQDNVVWDDILLPIGDGSSANFSSYSFGVSAGTATPSDPSIDRSLQGGNFSERKSEHLLDAVVANIRAPVNQSMDENMSTEHTVSKLSSAPSLYSASVKADSCLTGPLIGCEQKQSEFAGHSQSRDDADIKNLPNSSFTNEFQRERAVKRSLHDSSLKFQPSSWAEDFHSAKCENPGLAQIKKSDEPVKVNRKRAKPGESTRPRPKDRQQIQDRVRELREIIPNGTKGSIDALLERTIRHMVFLQSVTKHAEKLKQSGESKVLDKDGGFLSKGNLESGASWAVEVGGQTMGCPLIVENLSQPRQLLIEMLCEERGLFLEIADMIRGLGLMILKGVMEARSDKIWARFVVEANRDVQRMDILWSLMQLLQQNTKSSLSLTHQPSVVTHQGMWYPNVQCVSATSDASHL*HGYWIMYMYSCRTCAARGC*LDLDRCTQSSTKCMIL*CYLTQLLQPSIGRENVSARRQLYPLSWVVNFSKDSQLHFF*TISHLVAK*VVLQMQDFCILKVFLPIHVGIFLPFLS*WASEYLTKLSTAVMTTFVLLLCIL*TNINAMDPAV*PALEMLFLPCKCLIAYF*Q

>Pe2056043

ITSKD*YY*FLVTGHFLITLSCHVSISNMALPMSSWK*DGIVL*LELREYFNITVCVASSWQVDSLKFVMDDFLDQILSSSSSWMDMSGGQANGMSGDSMGPFQETLKHPTIPSDTLHRLVESNTVAGGPTSLQLNTSVPVTRQGSTSQFLTAVGGPSRTAVSLAELASAGSSSSEASGFQQALADSHPPAPVWSESYTVTSALPAAVGQGKIEGFTLKEETVQGDGHLLGKRSHSEDKMLARENRSGDTEHDDLQGTLLTSYTGPQGGQTVLPRTPGSQSHQQNLSTQTIQSGEGTSLQQYGNHSAVSQLQSGAGGGNAANGAARPRVRARRGQATDPHSIAERLRRERIAERMKSLQELVPNSNKTDKASMLDEIIEYVKFLQLQVKVLSMSRLGGAGAVAPLIADVPSQGSGGVMSTALGQASGPLALSQDGLAFEQEVARLMESNMTSAMQYLQNKGLCLMPIALATAISSSSGKPILATVPGTGVDSSEKQNSDMQSIVLPFSSSSTTMGLRASASGSDAPINENSMNKASIEKVVAEKSNGISPGLSSDVPKVGSQNREELLNRAQ*FLCPQE*ACECAKAVGEIF*YAIF*QIGYGS*SIQLKWFTCLGSCEISIGAKYVMYK*TPVLMRIIGGTLLVQGCQPFYLHVPFRQISGILSIYSNLNMTGADHRKDLLLRYQNGSGIHFPVSLCFDLWISRYLSKLSAIG*SCFNLLQQLLKSIFSIPRSILSL

>Pe2056101

PIISISAQAASECKSFKFSKQAAGFESRDCEIIRRGRIPLRLDILASLVNFLVDMQPSTSMLGTGPVQGVSSVTACLSSSPQVGLQEQMHSQQHQINNQQQHFSSQFDQTNQPHQNMDDFLEQMLFMPQWSDVAGGKSPWEFNPNNAPQGNNASNNNNSPQSLQSNTQKLFTMSLMPPGIGSQGLNSQENDGHAGEHMQYSYDQSPFLANRLRQHQLSGNSSINQTPNQVSQGINETNTSPGRSMVLQLSTGNASASQLLASMGNSPRGTAVMTRSPSTGGSCNGSDGGMLPLPLSLGQAGKSGDMNEPGSREEIEASFKSVNNARDSNLGGLFQPFAVSPRGVRPTGQNFHAQPGQVPLQGYGGMPQPQHQNPPPTGVGAAPPVRPRVRARRGQATDPHSIAERLRRERIAERMKALQELVPNSNKTDKASMLDEIIDYVKFLQLQVKVLSMSRLGGAGAVAPLVADIPAEGANAPQGTRTNGSQNSSPDGLALTERQVAKLMEEDMGTAMQYLQGKGLCLMPISLASAINNSGSRPQAPSTPSLQGLLASNVNDRQVLEPSSNSALSALTSTSMTIQPSISSPGSGITEQGDNAHKAGNGTRNPKDSREANSVTKSNGIGPALSKNAVKGEDEPQRRTGQ*QHHLANGNCTILLTPDCMKLFKHTKFVFHMLQQFLNSRSMGEHSFKKSPKHTSNFPFL*SLVFIFIDG*KN*ASLSK*GTRPTLMMPYLHRLT*I*QYVLLDPSMKINTX

>Pe2056957

SAKAIGLCEHNRTG*STCVLFHDSNHSLTFNPSAKSHPSSFLIASIKFGLRNLIRLAS*QLQLNLTFKVWNWYQWQLGFELGSENTGFLGRQAVFMFSFRQWACWFILWRFTELGEEVGTVRRMVVE*H*IYLASKAEYA*FDRGK*VRFLQGFCS*LDMVQELKGIDLWNFQYEGL*ISVLIMLWNAWHILGIEG*S*QGIFGAGADFRVLSWVFLSFSTVERIDLVIYCME*RSRCPRPYKAMALVDFTEVPRTRLRQQMQAAVQSIQWTYSVFWQFSHPEGLLVWMDGFYNGGIKTRKTVQPMELSPEELCAQRSLQLRELFESLSAGETNPPTRRPCAALSPEDLTESEWFYLMCMSFTFAPGVGLPGRALAKRHHVWLSQANEADSKLFSRAILAKSARIQTVACIPLADGVLEIGSTELIREDIGLIHQVTSLLADHSKPVCSEQSTSNPHSENASRALPPDQLSMQPPDGIVFEEQNVKATEYNSDNAHENAHIPMQVSGLQSEKTVTEFGQNGSEVMQLDMSPEDCSNDIGSELQVTGGNSCMQVKHSTNAWPDISHGLQSSGTQQYLDQGSDDESGHYSKTVSTILQQQRPSQWTETTTLQLVRGNINPQDITQRCAFSCWTGNGGVSVQKSMNPQWVLKYILFSVPNLHSRDRDESSPKLREGENGCRVRRSGQDDITVNHVLAERRRREKLNERFIILRTLVPFVTKMDKASILGDAIEYVKQLRRRIQDLEVRSKQMEAELKKSAEPRKQLSTTSQEKIVRQKSGGTTNNNNVDQELVSSCRNFSDHSDQQQYKISRFEKRKIRVLERSEPLTIIDDCSTDVQVSIIEHEALVELQCPWREGLLLDIMQTLSNLLFETHSVQSSVVNNTFVAKIRAKVKAANIGEKPTITKVKNAVLECIPSRC*IWNFICKISAVLMGLFPC*CDLRSLE*EEILVM*ARFGKYIWISACRYFMETSSAKEKYEYQDLA*ALWRFGFCDDCVSSVMLFTLILIFQVTDLTPPLQYSYTS*CNVQVLDAMLRCQILSSKNIFWKSWALFNRDL*YDDCYHPLEFAMIMTITST*SILILI

>Pe2057083

MMLTWEDGYCESGNSSAVTNASGHVVPSLLLNTMGNGCEGASGQDKWSVGDQIGPVVAKMSYQVFSLGEGIIGHVAFTSKHQWVFGERNCTNEAHSAVSVFENINPEYPTGWQNQFSAGIKTIAVVAVMPHGVVQLGSTQIIMENLEFVHHVKKLFGTLQNVPGAFLPDGIQDAVSGKIQAPIFPMMPISITSPGNLASSGAIKATSFQKQSNQVTTKFPSLGVLDRFVSPVDLLEKNKSLPAQTLPLPSMGSFDSSVEMKNRVNLNNILTHITNARGTEMGEKRILHSASKEAYPFKRQLQANRSLSVASKHDTQQPLFSGSGKTLMEKLQFQGNMASDCQEECILNPLLGSDEIANVGNLCQDNFMISVGADGLLESSLFNSGRSELTDLKQMDVQPPIPIFQGHAGHSGYCSSPSSLGTCNTNKTEMETTPMHVPMADAGNISYNVLSCSANIRTSACDLRESSDASWRLTADHTETKPLHGIGNSLMPSLGHDGLSMSNTLGINHRSMLGINYQPWHKGFLSDLPGLIQTNSVHRLNTDKKCSESFLPGFGKISAIPADCDKPGGNGDSDNCSYWPSKEQDDNAHAFMHVPSGAELLEALGPSFKKGTDKGMWNEMLFHGQDVCSSNLGTGNFGISHRSDYHSELDVSQSSDVSVVNAPEGWFFSETKSEDLLDAIVANVCPAPNRGTVDNISCRTVCSKITNSNSLPCASVMTDSISAGQLSVCDQKQSVLNISPPREYADLENDQNSNFSSEVHDDVGLKKSSNESSTCAQLKSLVSEVRCQNRGLPQRKKLEEPIKVNRKRSRPGESSRPRPKDRQQIQDRVKELRDIVPNGAKCSIDALLERTIKHMVFLQSVSKYADKLKQSGESKVLDKEGGMLATNNVEGGATWAFELGGSTVQCPLIVENLNQPRQMLVEMLCEERGLFLEIADIIRGLGLTILKGIMEVRNDKIWAHFIVEANRDVHRVDILWSLMQLLQPNTKSTTNTINQSFLVRSQGIDNVSQTYDAFQLSPMSTLLTNGAVSGS*NLCLP*LASVHYYCLYSVM*VI*VVESCSYPAL*QIT*VQKMIYSSYKWEE*ERSYF*RLIYHSSDCNLFNNLSLLTCFLWQG

>Pp00002G00970

MPSSLAEVQGAKLQAKAKTIAHGNESSGTFDNVVTKRWKYEDGGTSAESRATSGRSASTSGLVRHKLKGSLGRSIGRRSKRNLNADEASVDDARTSQGLVRINGSSTTALENCSLDLMISEIRSSERLEEVGSRSIGLARLSSKAIGQSRPYKSIKQDLMSQIRAEDGSGLSGVEAEQIAASPRSQDGRSDGTSTSTFVELLADAAVGSGDSSVVATDCAIGGNGTIDGVDIFSTDIGSGPLANESRKLSVQADNLGMNKNLERVSVLDNGSAEQVFPREQTKIISDAKKSAILNIHKQLSSLTSRKGGTSLVDRGEKIVVRKTTTWRDAVKLPDTIEEVLPVDSKRDPAVYLFLYEKFLEAGRLRDCVAVLESMDEHLILNMKKVNSYEFYSACKKRRALKEAFKFSRLVRRKSLRNFNMLLSVCAHAQDASSACRVLDMVKMAGLQADCIFYTTLISACAKASRIDLMFKFFNEMEIQGIEANVQTFGAMIDGCARAGDVPKAFGIYKKMLNQEVEPDRVIFNTLITACGRAGAFLRAFEVLADMRDAPQPIALDHITYGALIAACSRAGEVERALEVYKRMRSSKVSGTTECYTAAVHACSHKGYLNIALSIYDDMREDGVQPDEVFFCAMMDVAGHAGKIDVAFAILQEMKNIGTKPSPVTYNTLMVACSKVDDAENAMRVYEEIKALGLRPIRWSGAGRELDNAFKVLEELRNSGVTPNKDTYNILLEACERYNGRTVVLRGYNIMS

>Pp00006G00590

MAEVLSSTASILVAFFQVESSGSSREEMCPVAVPSSVASSCERLIWEGWTAQPSPVEESTTSKLLPKLLPELETSSYSALTLQQPDALSSILSVLHPFSHYSSASLELARNPDWSLKSSNPLRESSSEAGIRTSSFEGLYSGQHTTKKIHLGVIPYHLSEDQRQCAVSPPENECRLLSANSSGSLHWWHSIGPESPSSTLAFHNIGIQHSTFEKCEPRGQSHSSWPAASGTSPTVQYFHAHSADNEGVEVVKQDDSQISKALATYQPHGDHSLVLNSDRIASTTSHSEDPCGPKPGRRPAASYDTEMILSPSESFLTTPNMLSTLECVISGASNISDQYMNFVREPQEQRLSSISDLSLIPDSHADPHSIGFISGTFRTDSHGTGIRKNRIFLSDEESDFLPKKRSKYTVRGDFQMDRFDAVWGNTGLRGSSCPGNSVSQMMAIYEFGPALNRNGRPRVQRGSATDPQSVHARARREKIAERLRKLQHLIPNGGKVDIVTMLDEAVHYVQFLKRQVTLLKSDEYWMYATPTSYRSKFDDCSLVPGENN

>Pp00019G00310

MEKESSFNALRELVIGVEWTYRIIWKLELNQRSLICYQSYFNEREATEVDQAQEFQLVYCSRQFATSRPGYAYNAWKQQSSSWWTPQLMQQVPTDENRDLFLRKAGIQTLVCFPGRTDDGIGYCVELATKAQILETDTLRSFIQGFLYSSIAPHIQEEKVLRKVGGRIEGFPQASRTVTLPMEPVEHNSPWAMTVLQANANLGQPSVGIQELKYVDFPIHNSQLDLGSIDLPDIIGRDSPNRDPSTAFLFSTEWMTAGGKEIAGEGGIHLENLAGTPIDLQLRKKIAIENLDKILMEIPDEQLQASPSLQTNGGILDTTMTSPGSSFHQNSPTGIGKVGQVPPGHSSKSLFRPRRKSSFARLSSSSKRQPMIHNWAKNLAPKIVELQREAQPLTLPDFNAYESPGYSPHEHYQAVAHKLAERDRRFKFNQRMQTLISIVPIISKKDKVSVLTSTIAYIHQLTSRISRLEKGEELPAQPQAPTMETPPPLDAAGVSTLERVNQQTFVSDEAGAGRSSPSWPSVLVDEDDDDSTLIIKLEASNNNISLIQLLNLLLELDLRIRTLDYVCTDGRFSAKTRVKNSKTSNKEIQVTLQRFLRIKE

>Pp00022G01720

MAKRCVSRQVKGIIRLIASNGVERNPGIWVSSQLHHFGSSDIKDDLLSSSTPSHQLFLSGIRHTHYSGFIFCRNRLFKHASTLPEFPEEPGVNPDPQCEKTPLDRPGDVLAQDTLPNVEISPSSVGNVAKMLTIVKNLPWTSSSEKALSQFRGTLHPDQVSTVLWRLEGREEVETGLKYFYWAKAQNGFKHSITSFNTMLVVLASWGILDPLETLLKEMVAEGRPLRPNTLVKLITAYGRGNKSGDAFDLFNQAESFACSPTVHAFTKLIDILVNSGEFERAELVYKKLVQKGCQLDRFAYNVLIRYFGRSGQLDSAMEMFREMKIKGSEPDEYTYGFLVNALGKAGRVQEARSFFDAMLERGLTPNIPTYNLLMDAFRKVGQLDMALGLFAEMKRRGFQPSVVTYNILLDALCSAGRVGAARKLFHKMTGDGCSPDSYTYSTLVNGLGKSGRVEEAHKVFREMVDRGVAVDLVNYNSLLATLAKAGNMDRVWKLMKEMSRKGFHPDAFSFNTIMDALGKANKPDAAREVFARMVESGCKPDLISYNILIDSYARFGDAAQARQMLEEMVEAGFIPETKTYNSLIHWLATDGQVDEAFAVLEEMETAGCRPDVVTYNRLMDMLGKRGENQRAARLFQQMKDKGVEPDTLSYAVRIDGLAFDDRLDEALVLFKDMKAVGCPVDKAMYRILIRAAHRAGDTELEAQLKHESQFMPVESRLTSQKAKTK

>Pp00026G00840

MYVDVLQRCFKHKDLTSVKQVHDCILKSGMDQNPYVANKLMRVYIRCGKVQDARHVFDKLVKKNVFNWTTMIGGYAEHGRPADAIEVYNQMRQEGGRPNEVTYLSILKACACPVGLKWGKEIHAHISHGGFRSDVPVQTALVNMYAKSGSIKDARLVFDEMAERNVITWNVMIGGLAQHGFGQEAFSLFLQMQEEGFVPDSTTYLSILTATACSSAGALGWVKEVHRHAVKAGFDSDMRVCNALVHVYSKSGSVDDARLVFEGMLDRDVISWSAMIGGLAQNGCGHEAFSLFLKMQREGVIPNVTTYVSILTASASAGALEWVKQVHNHARKAGLGSDFRVCNALVHMYAKSGSIDDARLVFDQMSVRNVFTWNAMIGGLAQHGCGQEAFSLFLRMRREGVVPDAITYMSILNASASTGALGWVKEVHRQAVQAGLDSDVRVGNALVHMYCKTGSISDARLMFDGMVERDVITWTAMISGLAQNECGQEAFSLFLQMQREGFIPVATTYASILNVCTSTGLAMVEEI

>Pp00031G00300

MNPRSGGVDTHAENLLLERLGILVDRLNWSYGVIWTLNPRTRVLDWTRGYFQVNTQVGTASSAPRSNWDAQLFYTTYKSCSFAPGNGAVGRASVDGRRFWLTGDSVGQNAGTMEQSQFLRCAKINSTEFSALRMNSATTMICIPCIDSVLEMGTHIHVRENFQVMDQIHEVLSNDLLGVLPLQHTTSEHACSWDTLMFQADSHLGFHQDSAANNPSSLALQQPSSEMMSRSISEMSLRFSEDLNRPFQSEFNLSDQVWRSAISSNLSLNSDIFMHGQQDTQLDSAIDKVTTENLSTLHSHGGTPSTFLMPNPNAPSNPAHAEADFHVSQQVNQRIMTQSASHGLMVSQADRYRPWSGAATSSSVSSQGDSSARLSASPEPQQPQIHHNPLPNSQNSTLDRTSSSGTLRSLLNQNLRRTTSCGDQYEIQKAFSLFEAQEMQEHWRQQAGQQPELAGAQAATMNDKVTALKHDPGSLQRSTDDPPNLLKRSSNVSLVNQVSKVAGVLLRRKSLSALEVLKAVDTYHRETADKSLNLVQNIEEEPTGLATTVTENITPNLRGRRRWGMEHASSSRGQGPIHVGHDEAAVNHMMAERRRRMKQKENFTALRRLVPTISKADKASTLIDAITYLKDLQNKIQEMKASKEDINQRCETLENKCRELEDRNQQLVAMLSINHPSSSNQLDTLKSELLQSFFPSELKTSKAAGNPTYSISSDYHM

>Pp00048G00740

MSLTFRADDCNELPSMENWLNAGADWNEHHGIVDVDATATAPLFYHTAEASTTSGEDSLFDALKKGQEAISTHPTFLKLDTLELCKQNPDAVLVRTGSSSMVDSSSNEVDVSDHANSLPSTGMLVSNLLDRTGSLDESGLYMEADSPRGPLSPAALQLGKSPKGVAIKRSFEDASEASWMPAAEATHIRARKSAKQQLPKNVALSQEDLQASAAAVPYPPFVAMASSGEMAPLAIRSGVAPAFVPASAANVPPLLFPTPLTLPNVPSMEEIAQSRPKRRNVRISKDPQSVAARHRRERISDRVRVLQHFVPGGTKMDTASMLDEAIHYVKFLQQQLQTLEQIGNMSDPRFMAQPGGPMMMPQMGAMNAVSPLDLNCAVYQSYPGTSVVHPASQPPSQWPNTRPTSPFCSQGFQDSAQEQFCH

>Pp00051G01180

MMMMRGVLLQRALRTASVCDRALIFTSAVHQSCPAVEELATSESGHVEELQSGPTGLNDMNRQCMDSRYAVRGTTRHGTPGRTRAQSKPASELIQDPRIMMLFKLVRERKFKELQAAHEQMMKEEAESDETGLDVYFYSRLMTQCVRHAGNETALPVFELMTEAGVTPNVVPYTIVIRALMDLADNEKAHHSGQQGLNEVAKSETLASKAVELLHEMRGRGIRPNELTYKPIMSWLPARKRDAQFQELKNLMAEDGVAFDGVFKFYEIRLALKVGDLKKAERLYIAAKKENAKIASSLDGGSSEQLNCSFVSYILGSSKAKKLEETLGLWRDVEEQCRINAPPEVFNILIDSCCKHGHVSWGLDLLDEMQQRGTKLSPYAFNPFICDFARWGMYEEAFEMKAAMGRLGVQPSVVTYSTLVNCCVKLGDMEQAYNLLAEMKQVGIQPNAHCYNPLIMGFGSQARLDRALEVLREMLSAGVQPDSYTYSMLIFACSMVRNEDKAVELFEEMLQRGVQPNAGIYSAMASVFARCGKLERSIEMVKEIERRGEVVGTKAKSAILAGLSLAGRLGEALALYGALKREGAFPEAYAAGILLVAVGKAGDLDRMFNIFEDCRKENLWTKLTYQQRAEFLNVRCINVVLGCIRHNQLGRALEFLRKVKDENIADVAVLFDKIFLHISNGGRDANEMCWLDVDDGFAVVAAMRELGLSPSRMALEALLDGCASMNDSEQAQRVVSEMEKEGLALNVFSQIRLFRAYVAAEDEEKAIELLQQIDPYDWQDVHVQFILQQTLQPHLEAAQDSPEAAQAPETLPGVRQKLAELIDLYPYRQFMDD

>Pp00055G00040

MTLSLRPDDRSALRTMEKWSNSGAGWSNHQHPMLESSISPLFNYTQSHNGLQSQSMFLHPMGAAEQQALVQAFVQANVGCMMDQPSQSGCGNLDSLLKSSNSSSMVDSSASEMTSPDDDCTGAPENNLHMNCIPSLGGGPVSPVLTSMLDRTASNGASLMAAHPQSKNDFFRMRSPGGPLSPGASACAESEDSLHGGEACAPAMTGLKRRSFEGDDGDWMVDQMHLSQKVEKQHHQQQPEKALAFQESFGGYNVEVASTVGLTATYSDSLTIPSLMQPSPQPFLRSSGNCGAASPVDLDEFASMRAILFRHASQPVPSLEEIASSRPKRRNVRISKDPQSVAARHRRERISDRIRVLQRLVPGGTKMDTASMLDEAIHYVKFLKLQLQTLEQIGNNGCDPRSFLEQGGATEELANLVRPFDLNCTAAFQTWPPQTTTMVNHGDTSPQGHNSTEWACEANQDDCPM

>Pp00060G00320

MDTTQQGLPTLNYLLQHTLRRLCTESQWVYAVFWRILPRNYPPPQLDTEGGMIDRSKGNKRNWILVWEDGFCNFSACAAGIGMDNGAFSSFPQSAQEPGRGSDEDCEAMNPELFFKMSHEVYNYGEGYHFPFLHAATFALRTPYVLRRASSIASKISYNFCQIPVCERLKIILWILMFGSLMGKVAADNSHKWVYREPVEHEISFFSPWHGSLDPHPRTWEAQFKSGIQTIAVVAVQEGVLQLGSTKKIMEDLNFVLQMQRRFNYLQSIPGVFVPHPMSFAGKKRSNGDGSPPSDRGHWIAVPDCDKLASNPHMSACSINQFLSPRIGQQLWRSSSDTIFLGVKRPSDTEPLSFNFNEYPYVNNGESFLHRPECPSPPKSLNTGLASPQSGGSPTLSNLLPSMSSLQALLSKLPSVTRTERDSNSCIGGAVPTPSGFDGTAGSSCRPPVSRVVVNKIADERAASLKPAAAGDNESAHRDAHMHLKSTSKTENEPLHDQKHGVTRAATSISSLSSRNSEVVSIEQASDDTEGHQNGTSHCKPDCHKSFCSSTFLDTFDNLADFSTLHDSLIEAGDSYNSFLSEIYR

>Pp00060G00870

MTDLNSSLESPGSSVEETPAAANSVATSCEAMMWEAWSTQPSTGDEASTSKLDLLPELVSSSNSRLSFQQSDLLSNMLSSFHPLSQHSSAGFELSHNRGGSEHSPEFLQEGSSEADTETSSFGDLYINRQTSRNSFLGSIACPLPSNHSDSGKNIRREDLSNQLGAKSSAPLQLWQSLGPESPSSPLAYHNIGYRHSHGEKWETGSQSHSPWPTVSNTSSTIQLLGGRAAENEVIQVLKSNDSEISKSLATLQQYGDHGRQLNLNHSSTTNHPEVIYSSKFGPKPSASSHTDVLMSSTNSSFLSIPTAWSTPEYSMSGPSTRSEQFMNFVRIAQEQNSVPISGPSPILGSYVGCSNRSKSGISRVVSQETPTAKNRLLACEGSSGPAPKRPSYAAHSDSHADQAAAMWSSPNLRRSSFPSILTSQAMEIYAIGPALNTNGKPRARRGSATDPQSVYARHRREKINERLKTLQHLVPNGAKVDIVTMLDEAIHYVQFLQLQVTLLKSDEYWMYATPNTYKGIDLTNSPPQTQRLQSSA

>Pp00067G00140

MTLSLRPDDRGALRSMEKWSNSEIGWGNYQHHPMLETSMNPLLNYSQSQSVFPQLMGAADQQSLMEAFAQANAGCMLNQPSQGGGDSLDTLLKSSNSSSMVDSSTSEMTSPDNYCVGAPENNLHMNCISSLGGGPVSPVLTNMLDRTASTGACLVAAHQQSKSEFFRMESPGGPLSPGAPPCGESEDSLHGGEASMTGLKRRFDGGDKNWMVDQMHGSRKQQKPQLQQSEKALGFQDSVGGFVDAASTAGMPGKYSDLATVPSVMQPPPLSFLPSIAAGGASSGVDIDKFASVRAILFRHASQPIPTLEEIASSRPKRRNVRISKDPQSVAARHRRERISDRIRVLQRLVPGGTKMDTASMLDEAIHYVKFLKLQLQTLEQIGNNCCDPRSFPQQGSAAEELASLVRPFDLNCTAAFQTWPIPPATMGNHGDTSTKWGTEANQATFSDGRM

>Pp00074G00230

MMEMGTPNYWDAADPLMVEAFIGGYEIPGYETQDDLASTLGQDLEQNDSVLQRRLHRLVEESSEDWTYGIFWQLSLSPSGESMLGWGDGYYKGPKDSDQFEPRKTQTEEHQLQRKKVLRELQALVSCPDDDGTEDVSDTEWFYLVSMCHSFAKGVGTPGQALAFGEYVWLEEADKASYKICTRANLAKMAGIQTILCVPIMNGVVELGSTDAIHERLDVVEYVKMVFQEPTWGLTNMSPIISQSQVGKFDTTFMPHYPSIPFDSTSVSGVSSMTLNTDPGLADSESMDFGTRHSHMGKMVSHSGAFGFNGYDHVWGQTNEFHYNDPLPDDNVERDLGQPMCNILGSLPLQDEKLPLASSPPPKTLDSDSRYSIFQQNNVKKPPQLDHTQTSLPVTERLHPKPHTSQAFLHHNGSFDVGEMFNPPGHTQTVRSNPPSLDEQLHSPSMPAVEKLPIVEKPTSIYKPESVEKPMPVFKPLPQPPSPPASKPAVPVPANGLLLAGHLDQECVDTELITMKNNVVEAPKVPRKRGRKPANDREEPLNHVQAERQRREKLNKRFYALRAVVPNVSKMDKASLLGDAIAHINHLQEKLQDAEMRIKDLQRVASSKHEQDQEVLAIGTLKDAIQLKPEGNGTSPVFGTFSGGKRFSIAVDIVGEEAMIRISCLREAYSVVNMMMTLQELRLDIQHSNTSTTSDDILHIVIAKMKPTLKFTEEQLIALLERSCQNTGYLRKREGSDRLLQRPDNSPQLQ

>Pp00074G00250

METQVPSFWDAGDSAMIEAFMGPAYGIPSSYEVQDDLASTTEKGLELSETVLLRRLHTLVEETSSNWTYGIFWQLSRSPSGELMLGWGDGYFKGPKENEISEKRIDQGGSEEDQQLRRKVLRELQSLVSNTEEDVSDYVTDTEWFYLVSMSHSFAYGVGTPGQALATESPVWLTEANKAPNHICTRAHLAKMAGIQTIVCVPTRTGVVELGSTDLISQNMDVVHHIKMVFDEPFWGANRSQVMAQSLLMDSDATFFPPSPSIMSMGTTSAFASSPSVASRGSTLGKDHESHYRGRNVSVEKIGSSMASTSFDTLDYMWQQSDEMQFNDGVSVGTTEKDQGQSRLYYPVLGPPVLVEKLPFSATSLISRTRAAEVKHSSMLQNVEKLASEDQKPSSLPHIKVHTTHSYPEKTGAGELSQVLSTPDLRQSIEMKLPAQVETRRAPGITGGATKPVAEKAKPVPKPPQQQQTAISGPPASASGRSSFDQSEHDSFQESEAEISFKESSAVEFSLNVGTKPPRKRGRKPANDREEPLSHVQAERQRREKLNQRFYALRAVVPNVSKMDKASLLGDAIAYINELTSKLQSAEAQIKDLKGHVVGSSDKSQESLSIARGSMDNSTIDGLSIRPQGSVNSTSISGNAPSGTKPTIAVHILGQEAMIRINCLKDSVALLQMMMALQELRLEVRHSNTSTTQDMVLHIVIVKIEPTEHYTQEQLCAILERSCQPYSCSTKDEGHGLSEKLGSSRRSQ

>Pp00093G00570

MVARVEDCFLGREQGFGLSKFSWKWSQGSNQVAQCLRDGDVQHSNSFYDASRSGDLPTWSSNPSGMQGFTKGLHNVHYGVQPHHFLNATVVPESDQNDSVLNSNHGSSYVAARNQVANAITLAPGRAGVNSLAPGLARNPSFESINNTGSCSTPSFGVGSPETVFPAHPSDLGSRDSSDMMLRNRDQLTVGAYPESKVQPQLSDSQSNDSLSYGSQDAIVNCVDDFWSPQEAIDAVLPDFRLEAMPDEISDTGACGLSFSDGQLGSLVSTHHGHSTGDTGPYNYFRRGSELAASLSTMAHAAPAPDFCQSFLVGGVDDSENPMIGNSVNLSTRPSQLSIPPKLSSQAVFNGSPDHPGRLCWSHQPWSSSQHTSGSGSLDYIKQHNITSPTSLAGSSPSSPTTPLDQHLFASSTLQGTSPVGNISSVLKADEIQKKLAELIGANSNKRSFSAMLGTCAAGTAYGGSSASGGKETSLSLPSLGQGGANTDKFDAAALSALLYSNSPADQDFRSNFASPPAPKQRRRHGTATDPQSIAARTRREKFTDRIRILQSLVPNGERLDTVHMLSQTFEYVRFLQHKVWDLYNNKDSISEVKCEKWKEFVDATTGQVIV

>Pp00098G01090

MSLTFRADECNELQSMENWLNAGTSWSEHHGIVDVETTASAPLFNHIAEASSTRGSPESLFDALRKGKEEIPTHTTFLKLDTVELCKDNPDAVFMKAGSSSMVGSSSNEEEVSVLTNILPSFGVSVSTMADRTPSLDEPVLCVDAKSPREPLSPGALQLRKSPSRASTKRSFVDASEASWMPVADAQQSRTQHPVKQKQPKNGMLSQEEIQSPPAAIAYGPPFVGMSTSGEMATLAIPNPTAFLPGSAANVAPLLFSTPLTLPNVPSMEEIVQLRPKRRNVRISKDPQSVAARHRRERISDRVRVLQHFVPGGTKMDTASMLDEAIHYVKFLQQQLQTLERIGNMSDPRFMTQPGGPMILPQVGASTSMRPFDLNCAVYQSNPATTMTLPPSHQQPLPWPNTRVSNSFCSSFLDSAQEQFCY

>Pp00111G00580

MLNHQLQHTLRMLCTEMQWVYAVFWRILPRNYPPPQWDNEGDSMDRSKGNKRNWLPLRERKMRRILVWEDGFCNFSTSAAGAVKDVRGALPFFQAAQPWPQGSDHEREGMSPDLFFKMSHDVYNYGEGLVGKVAADSSHKWVYRKPEEHKFSFLSQWHSSLDPHPRAWEAQFKSGIETIAVVAVQEGVLQLGSNKTILEDLNFVLHTQRGFNNLQSVPRVYLPHPNQCGRKKQLKNGGFQPFDPNYWMAPSDCDRYTSIPHLTPRSIDQFLSPQIGPQLWFSSSGTTVLGVKRPPESEPSSLSFKKPSYMNGSGGFKHQPECPSPPKSLGTGHRNLQGGAGLPILNVLPSMSSLQALLSKLPSVIRIEGESTYTGAISTASGFDVGNTLVSSNCPLVNRPAMNYSFQVSRAVDDSNDHMQRMIFSDSETGASQEIISTNTSSSLSSSNSDMVSGEHASDETNGRWTTSMSNLKDSHNFSSSYLDTFDNILDFGTHMNSLEIGDCYNSFLNEIYTYWGCICHRFVPPAAVGASTTAPTWLVDELSL

>Pp00112G00410

MRQDRRDIPQGFRVPLNVGDILRENSERALQCLVFNESVIVHQSSCTLNHQLRLFDHRPLAVRANSERAKSLRYEACVEEKKLQTYMDSRPGLPPVNPMLLQHILRGLCTDSHWVYAVLWRILPRNYPPPQILVWEDGFCNFSACAGPSCDSRAGSSSFLSVRQSPSNDGNSEAMNPELFFKMSHEVYNYGEGLMGKVAVENSHKWVHREPLENEMSFLSPWQTSLDPHPRTWEAHFKSGIQTIAVVSVQEGVLQLGSTIKIVEDLNFVLYMQRKFNFLYSISGGLVPTSVVSGGSRPSNGLPLEGGHSYGQLSRRSIDQFLQPWVRPHDLWRPTTNSTFAGMKRPSEYVEPLSVTEYSYLDGSGFEQCRPECPSSPKALNAGHTSPLLSPSSSMPAVVPSMSSLHALLSKLPSVKTLDAENETSRLLTSCALPSNRLEGNFTCNLPPMPQPVSKIHSERAQAKEAISAGKRVVIIGELMASGIEMSPPNSGSDLTHSTVGEGMEEQQNQSTMARENYLYANSTFLETFDKLNGYETPESLDNGDSSYSSFLNEICS

>Pp00140G00140

MVWLEGWVSQASTNGEGSSSSTSAFKLPEVVPTLPNTRLSFQDSGLLSNHWIPSFNTFSQHIPDIAAETLSLEFMQERLETLPEASFEELCMQHNSKPLYLNSIPSPVPNNHPHFDKSRREDFLHYAKTYAPFHAWQGLRVDRSPSSPLAFHDTLTNSSGDEDTDGAQLRSTWLGKTSSTTIQLLRASASENAWNNGQIKYNNSQKLAKTDLLSNSNYSLSVVSNQSHGNQSEILGSNRKGKAISRSEDFAFVRGHEPTASSHYEKCMDPNGSFLPMSPIAQVLKYATAGPSFVSNSQLQQSKTNFSNETNSTRGLTGNDLDLNFVQKAQERLLAFQPASTASGYSARVEQSRKNSYRFAMDRSSMSPSRRPNILGNQEAGLAWSPYDTTQSRTTKSKLQCRHLLGTPSQAMDIIAVGPALNTNGRPRAKRGSATDPQSVYARHRREKINERLKTLQRLVPNGEQVDIVTMLEEAIHFVKFLEFQLELLRSDDRWMFADPFIYNGMDITGSYPHVPSGLERLNLRG

>Pp00143G00220

MGNLLVGVAQIWPCRSFVQEGKHQETVHLVSAVGGTTKKIPTRHGFEAGVGYHEPDVAATSITQRVYRPAVGYAVFWRILPRNYPPPQWDTEGGIMDRSKSNKRNWILVWEEGFCNFPAYSLARTDSEASRASSSFLSTQQSHHGNDSEEPGAISPELFFKMSHEVYNYGEGFMGKVAVDNSHKWVYRDPAENEISFFSPWQGSLEPHPRVLDAQFKSGIQTIVVVAVQEGVLQLGSTQKIMEDLNFVLYMQRKFNLLLTVPGLFVPPQQASTSGDGTRRTIDDYNGNWSVPSGKEFEGSSTAQPNDFQYLSPREAWSFTTAGTKSVPPILEPYAFSNGNLEACLTTCSSPTKAHNTGHTSSHGSTSPTMPPTLIPSMSSLQALLSKLPSVTPVEDPEGATRLHPSYSFPVANFPSNWPPISRQGLGMPPHAHDLSADTSRSQNSSLILESFDNVGDFGIQKAVENGDSSYSSFLNEICS

>Pp00147G00560

MSEVLEGEGGGCNARKREWRFSCFEARERSLGRCNRLQQSRGCHQTMNHCVPEWDRSDDMLDALIPSDDFHGPVYGKAEFVRSRKSHFCQVPVQNTLEAGGNDNGSVKMQGQSSKCNKPQWQANYPTTWSSGQGTEDGVTPGKPVTSKSDALLEAAVNEAPTEVHPGHHIDVAHDEMVSWLQYPLDDTLERNYCSDFFGELPDSHTQLLRESFGHGSTKTARTSYLGSPGNDSVMNRASTTDTAMLLGAGRAAGFLPQAGAEAFSKVRTIHSLQPSSVTKWQQPHPNSSNGSNMCATANLATTRAPPSAPPSNPMLPPRTQPMIPNVNTQTPQPNNKPGSMNFSHFSRPAALMKANLHSLTGMNSVPPPSARFKQQQNQTGKPTVEACTSTGSSIAESTTAGQRSSGTQQELKTQPGVTEREQSNGIKDCRWQSAPSLSPKKDRDYVVSEGDCKKTFNDQETYRISGVTSDAVLASSEKGVSHVTQHPDIQEPTITSSSGGCGTSAERPKGFATSNKRKSSEREDTECQSEDGEDESIDTKKPVTGRGSTAKRSRAAEVHNQSERRRRDRINEKMRALQELIPNSNKTDKASMLEEAIEYLKMLQLQLQMISMRTGMTLPPMVVPGGLQQHMQMPQMPGMPSMGMGMGMVPMGLSHGMMMDMGVTAQGRGVVPMQSHAGPSLNGSMASASSMVDVHDPRYLASGVIDPYNAYLARQHQPMQMTQPLNIDKYNAYLMQQHQLQQQQQHHHHQQQHPQHQQHQQTSNMNGGPSH

>Pp00147G00590

MAGPAGALWSTCDPQPIQQAEIFSGPDNQAGLMSFHVDTPFHWGSEPWALHSRSDDIALMSPSLVHDISPYDSVLHLSGVSGDVQDLVCGNPKFRQSGQWGQSEFSYSVQDNMQDLLTNQFIPYNTSSLGLNHLSPNFTDLDCAPVYNDTKAFGTVTHNRAVPSTNTQSAQHGSSSMVSSNRPITSTASPTTQYGGPRTPSQTTQYGGSSMVTNSMEMFASAAPQGIMTTSGLSGGCNSDLMHLPKRQHAHSLPPTTGRDLTASEVVSGNSISNISGVGSFNSSQKSSASVMMSPLAASSHMHKAAAVSEELKMASFNPGPFVPTQKKQQHEQQDTMTSNRIWADKNNLGKISSSPIPIMGFEQSQQQSMSNSSPVTSLGFEQRQKMSMGSSPSITIIGFEQRQKQPMSSSSPISNMVFEPRQKQPMSSSSPISNIVFEQRQLPTVGSSPPISISGFEPKKQPSLSNSPPLSNLGFEQRLQPMSNASPISNLPFEQQRQQATMSNTRSAEPDSVESTTKWPLRMDGAIGGCAGLPSSQKAPVIMQPETGTMKCPIPRTMPSNAKACPAVQNANSVNKRPLTVDDKDQTGSMNKKSMQKFLGPQGCSRLESISALAHQKVSQSTTSGRALGPALNTNLKPRARQGSANDPQSIAARVRRERISERLKVLQALIPNGDKVDMVTMLEKAISYVQCLEFQIKMLKNDSLWPKALGPLPNTLQELLELAGPEFAGIDGKNTEESSEKPKKSALEVIELDGNQPSAD

>Pp00161G00230

MDTKQGLPTLNHLLQHTLRRLCTESQWVYAVFWRILPRNYPPPQWDSQGGMDRSKGNKRNWILVWEDGFCNFSACAAGVAKDTRGGPPSYPAAQSKQRGNSDECEAMSPELFFKMSHEVYNYGEGLMGKVAADSSHKWVYREPVEHEISFLSPWHSSLDPHPRTWEAQFKSGIQTIAVVAVQEGVLQLGSTKKIMEDLNFVLHMQRKFNYLQSIPGVFVPHPMSCAGKKRPNGDGSPPSDRNNWMPMVGSDYDRPTSNSHMTARSIDQFLSPRIGPQLWRSPSDTTVLGAKRPPDSEPLILNFKDYPYKNGSDGFHHRPECPSPPKSLNTGHNSPQGATSPTIPNLLPSMSSLQALLSKLPSVTPTEGESTCTSTMPTTSSFDGSTTFVSSSRPPVSRPAVTKPPVVDSRPNPSTESPSIENGNGHMPEVVLAKTETEPLQEKIGAAVSSSPSSGNSDVASVEHTSDDTEGHEATGTSKFKDSHKFSSSFLDTFDNLADFGAHLESLETGDSYHSFLNEIYR

>Pp00163G00450

MSQSLRANDCSEFPDMEVWLNARTKWSENHDVHAESSVSVPLSFHSDETLITGAGSLFEDSRKEDVTLRPFLKLETMGLCVENQNARLLSARSAGMVDSSCNEEDISEQASSVALSVIPSGTVSAADFLDQAPRVDEVANLYMEADSPHGLLFSTGLQLETTSRTASHKRVLDDSSEASCMQVADVQSQSRKSPKVQHGPVLLQDDGLQVPVDYVPFVAMAKPSQAHFAISTSATFLPTSPGNVVPAVLFPTPLTLPNVPSMEEIAQTRPKRRNVRISSDPQSVAARHRRERISDRVRVLQHFVPGGTKMDTASMLDEAIHYVKFLQQQLQAMERISRGGSVMMPEVGYNYNAGRVLDLDYGSHETHPSKSMAPPASQQSVPTRLDKTWATNPLHSQGLRDPVLQQFCH

>Pp00164G00360

MVRFNYMYPVQEQLEAMTDQHTPSMDSVSSAGEKTSSCIVQQGGNASETSNLWEEWTQGSNGDDSVSTSNFLPELNSSTSSRLAFHQSDILSTWISGYHPLSQSSLSSEFSHTSDRENHPPAFMQEGLIPSGLILDSDPALTDIYTRSSSSDSLPYPTARIMDKALTDHELESAVPLAYEKGCVPPQVLRNLGPLSPSSPLAFQNGLLNPLRDPWDSCPSALPWSNVTTASQTYGQVTTRTFIPDHSASAIDKLEAVATITAGYGASKPQHTDVFIEPNGTFQSTPAGWAPQFYDGSEATGLLVKPMRAIASLGEAGCGEATSEFCTKTKPGLLKGGDTITSPVGSLLGDCKKAESSMKQVWPGKHRLELVELVDGEDTKSSPTQLKRPKHSTDYANVLLSDHILKGAELRSYFHSGDVGLNASQAMDIIVIGPALNTNGKPRAKRGSATDPQSVYARHRREKINERLKNLQNLVPNGAKVDIVTMLDEAIHYVKFLQTQVELLKSDEFWMFANPHNYNGIDISDPSSMHSPELESNI

>Pp00167G00680

MAMAVWWRFGGLDMSTIHRIRTRIVQNEKEMSGSKLEAIKEMTPLVPPELRLELQAATRAVKWTYSVFWKPASSNQKTLVWGDGYYNGTIKTRKTIGAKELTPEEFGLQRSQQLRDLYNSLSDSKTGHQQASKPFALKPEDLAEQEWFFLLCMSCNFAEGVGLVGRAAADGRYAWQCKTNEISTKLFTRALLAKKSASIQTIFCFPLMDGVVEFGTTEHVSYLDLPQSHTKILLGFVKTMAALTFILSSVKARENSANLENEKLLDNLYVCHHKFVLVDSIFTLQGSDSRLYKGENRGKNQSGVLPGRVFSSWKKNSTPSQKSQKAENRQKILKEALFRVTRLYDGAWKNKVDSSFIFTDRAVEDRTSNLGSQKPVPSSEETSASHVLAERRRREKLNDRFVALRELIPNVSKMDKASILGVAIEYVKELQSQLRALESNNTQDGTPRQFGTANEDATITNTTREHLECAGVVHVIDEDKAATSECTITEESFKPGHVNVRVSMNNDVAIVKLHCPYRQTLLVDVLQSLNDLEFDVCGVRSSISDDILSTVLEAKLRSASDGSSPTIIEVEKTLHRAAAGLLKERASASSLQ

>Pp00169G00600

MRNERHCGPDREDVSNTHQPRPTETDRATYVALLQNCTRKRLLPEAKRIHAQMVEAGVGPDIFLSNLLINMYVKCRSVLDAHQVFKEMPRRDVISWNSLISCYAQQGFKKKAFQLFEEMQNAGFIPNKITYISILTACYSPAELENGKKIHSQIIKAGYQRDPRVQNSLLSMYGKCGDLPRARQVFAGISPRDVVSYNTMLGLYAQKAYVKECLGLFGQMSSEGISPDKVTYINLLDAFTTPSMLDEGKRIHKLTVEEGLNSDIRVGTALVTMCVRCGDVDSAKQAFKGIADRDVVVYNALIAALAQHGHNVEAFEQYYRMRSDGVALNRTTYLSILNACSTSKALEAGKLIHSHISEDGHSSDVQIGNALISMYARCGDLPKARELFYTMPKRDLISWNAIIAGYARREDRGEAMRLYKQMQSEGVKPGRVTFLHLLSACANSSAYADGKMIHEDILRSGIKSNGHLANALMNMYRRCGSLMEAQNVFEGTQARDVISWNSMIAGHAQHGSYETAYKLFQEMQNEELEPDNITFASVLSGCKNPEALELGKQIHGRITESGLQLDVNLGNALINMYIRCGSLQDARNVFHSLQHRDVMSWTAMIGGCADQGEDMKAIELFWQMQNEGFRPVKSTFSSILKVCTSSACLDEGKKVIAYILNSGYELDTGVGNALISAYSKSGSMTDAREVFDKMPSRDIVSWNKIIAGYAQNGLGQTAVEFAYQMQEQDVVPNKFSFVSLLNACSSFSALEEGKRVHAEIVKRKLQGDVRVGAALISMYAKCGSLGEAQEVFDNIIEKNVVTWNAMINAYAQHGLASKALGFFNCMEKEGIKPDGSTFTSILSACNHAGLVLEGYQIFSSMESEYGVLPTIEHYGCLVGLLGRARRFQEAETLINQMPFPPDAAVWETLLGACRIHGNIALAEHAANNALKLNARNPAVYILLSNVYAAAGRWDDVAKIRRVMEGRGIRKEPGRSWIEVDNIIHEFIAADRSHPETAEIYAELKRLSVEMEEAGYFPDTQHVLHDLGKAHQETSLCTHSERLAIAYGLIKTPPGTPIRIFKNLRICGDCHTASKFISKLVGREIIARDSNRFHSFKNGKCSCEDYW

>Pp00173G00460

MAQQPSTTMMMAMQQQQVHGGGNHHGGMGFHHPGMGGQQGGGGGGSGGGPVMDELMEHMFGMPGGGMFDMAGGRVGPWDYNVGSGAGKGFGVGGMPSAVGLSKKGNEEVDYGLSEVQIRHHQQQSAGARGESGSGGMPVVREARNGAPDTLTRSVSLGSSASEESGPQQGKGDQLMGSMVAPSKHLQQPYGGAGSGVPTLPMNFAPAKAENVMLVGEMDSHNAHGKRFREDEDGRPRPTGAMPPGGCQGSGYANPGVPAGQSLPGMGARPRVRARRGQATDPHSIAERLRRERIAERMKALQELVPNSNKTDKASMLDEIIDYVKFLQLQVKVLSMSRLGGAGALVNSDPPAEGGNNFAASAGSSGVSNPAQDGLASALTERQVTRMMEDDMGAAMQYLQSKGLCLMPISLATAISTTNKGPAQANANTGDRQGSAAASNIGKSTAGSSLAGGSKEDGSEAGRVTESTTQDT

>Pp00208G00430

MLHQGAGRARGGLPSLLLGRRFDLWASFPWSFSSTSDDPSSRQSEENSQTIDELWEYFAGPSQWWDNRIHKRNPRSPDLKHKVTGKALWIDGCFTPEWVKFQPVAQVGLQSYATCTTKCTEGGAQGKRQPKGNDGKLATACESARVLGRIAQGGINMHVQTANTLSEAIVVLMNRLQRGLITDSFMYVEVLKRCLKQKDLMAAKQVHDCIIKSRMEQNAHVMNNLLHVYIECGRLQEARCVFDALVKKSGASWNAMIAGYVEHKHAEDAMRLFREMCHEGVQPNAGTYMIILKACASLSALKWGKEVHACIRHGGLESDVRVGTALLRMYGKCGSINEARRIFDNLMNHDIISWTVMIGAYAQSGNGKEAYRLMLQMEQEGFKPNAITYVSILNACASEGALKWVKRVHRHALDAGLELDVRVGTALVQMYAKSGSIDDARVVFDRMKVRDVVSWNVMIGAFAEHGRGHEAYDLFLQMQTEGCKPDAIMFLSILNACASAGALEWVKKIHRHALDSGLEVDVRVGTALVHMYSKSGSIDDARVVFDRMKVRNVVSWNAMISGLAQHGLGQDALEVFRRMTAHGVKPDRVTFVAVLSACSHAGLVDEGRSQYLAMTQVYGIEPDVSHCNCMVDLLGRAGRLMEAKLFIDNMAVDPDEATWGALLGSCRTYGNVELGELVAKERLKLDPKNAATYVLLSNIYAEAGKWDMVSWVRTMMRERGIRKEPGRSWIEVDNKIHDFLVADSSHPECKEINESKDKVIEKIKAEGYIPDTRLVLKNKNMKDKELDICSHSEKLAIVYGLMHTPPGNPIRVFKNLRVCTDCHGATKLISKVEGREIIVRDANRFHHFKDGVCSCGDYW

>Pp00209G00080

MAQQPSTTMMMAMQQQQQQHQVHGGGHHHGGMGFQHPGMGSQQQGAGGGGGPVMGEFLEHMFGIPGGGMFDMGAGGRGGSWDYNVGSAAGKGFGVGGISSGVGLSKKGNEGVDYGLSEVQIRHHHQQQQQQGGGARGEVAMPVGREARNGAAVHGDSLTRSVSLGSSASEDSGPQQGKSDQLMVSPSHLQQPYGGGGGGSGSGVSPLPMNFPQAKAENVLLVGGMDSHNALGKRFRDDDDGRPRTTGVMSTGSAQGSGYANPGGVPSGQPLPGIGARPRVRARRGQATDPHSIAERLRRERIAERMKALQELVPNSNKTDKASMLDEIIDYVKFLQLQVKVLSMSRLGGAGALPSLVNNDLPSEGANTFAASAGSSGIPNPAQDGLALTERQVTRMMEDDMGSAMQYLQSKGLCLMPISLATAISTTGKGSAQATANAGERLGSAAAANTGKSVTDPSSAGGSKEDGSETARVTECATQGT

>Pp00257G00150

MGAASSDIRISSNSGLELEGDSSWKAPKSSRNTSNMDVAKQVVQDTMRAASFCKGVLDEEWYTPETSLMELSSYSSPYGAQDARSNFSLLDSSLNYDNGNLMANFRPAPSSTLGIGQLESNRILSDLACTGQCSSVGLLSSISPGRHLRRSTTTDSLGSGLPTSFSQGAVIPSGSLSNITSRNTNTESTNFPPSFSDAYNAPALDLDKTGKECAGDIRDKLEPYQTNKRMESYPPRQQAFSQKRAASPRSMGGGTVSPTSKSPPRVVTSTSNDSSVDIRDEDSPHVQNFRGAELHSGSDDPNDIGIDGDDHNGKDDDDLDESGDGSGGPYEVEEGAGNGTQNNGKSKAKGKRGLPAKNLMAERRRRQKLNDRLYMLRSVVPKITKMDRASILGDAIEYLKELLQRINDIHNELEEAKLEQSRSMPSSPTPRSTHHGYPTAVKEECPVLPNPESQPPRVEVRKREGQALNIHMFCARRPGLLLSTVRALDALGLDVQQAVISCFNGFALDLFRAEAKDVDVGPEEIKAVLLLTAEYGMHSLQ

>Pp00273G00090

MLETSIGRLLNCSYSHGALLLQSMSPQLMGSAEQQALVQSFVRLNSGCMLEHSSQQGNGNPNTFFKSTNSSSLVNSTSEMTSPDVDYVGALENNLHMNCISSLGGGNNVLIPGLTSMLDRNKNRDGTFMAAHRYYKSFYPMESPGGPLSSGLLPCGKSENSLNKGEARVIAIADLQRLLEGDDEDWMVQEEIKMHESSKLVKPQQQSEKHAVFQDSFGGCIEASSSAGFTVTCFDRVTIPSATHPPTPPFRQRKGSVGGSPAVDLDNFARMQAILFRQASQLIPTLEDIASSRPKRRNVRISIDTQSVAARHRRERISDRIRVLQRLVPGGTKMDTASMLDEAIHYIKFLKQQLQTLEQLGIDGCDPGDVALRGGEALQLSSSVRPAFHMNYTSAFQSWPASVTDLGNRSKTCPQD

>Pp00282G00340

MVARVEECFLGREQAFGTSKQPWKWSVASNLVAQGVKDGEATNINMLYAASHNDMLHDASRTGDLSSWTNYRSDMQGSSKGLFGVHHSSLQHHQSLNSDTVPEPVQNDSILMNSNYLSSYFMAQRQAASAITLAPGRAGVNTLAPGLARNPTSESIKNIGSCSTPSFCVGSPSPSTVFPSHPSDLGSRDSSDMLLLNLEQQAVDTYSESRVQPQLSDPSQSKNSLSYSSGEERDAIVNYVDSFWSAQGANGDAILPELRFDAMLDDTADTGACGLSFPVVEMGPLLPAREIGHHSNPRSVQRGSAMAASLSAMTHAASAPDFCQSYLVGGVGTAENAIVGNRTTHGNKPSQLAIPAKQSSVAAVNTSSDHPARLCWSDESWFGSQYPVGSGSLESSKQPDIKSPTSFAGSSPSSPTTPLDIHLFASSTLQGPSSARSTSSVLKAEEIQKRLTEFNVDGSNTNKRSFSTMQSTSHGGNPLGRSSIPGAKDILSRVHGGVNTDKFDAAALSALLYSNNRGNEDFRSNVVSPPAPKQRRRHGTATDPQSIAARTRREKFTDRIRILQGLVPNGERLDTVHMLSQTFEYVRFLQHKVWDLYNNKDSMSEVKCEKWKEFIDASTGQVIV

>Pp00306G00390

MDTNQQGLPSLNHLLQHALRRLCTESEWVYAVFWRILPRNYPPPQWDTEVDMMDRSKSNKRNWILVWEGGFCNFSACATGLGNDGRPGSLCFQGRQQPQGSDEPCEAMSPELFFKMSHEVYNYGEGLLGKVAADNGHKWVYREPVEYDISFLSPWHGSLEPHPCTWEAQFKSGIQTIAVMAVQEGVIQLGSTKRIVEELNFILHLQRRFNYLQSIPGVLVPHPFSCASRKRLIGNGPPPAGRGHRMAIPDYDKPTSNLNLPVRSVDQFMSSRIGPQLWRSSSDTTVLGLKRPLDSDPLPLCTDHPYMNTGEGFHHQRESPSPMKSLNTGLTSAQGGASPTIPNLLSSMSSLQALLSKLPSVKPTEMDSNNCTSVVTATSSGFDATAGISGRPPVSRSVVNKMAKERAASSKETAAAENGYVPGDSQTHLNPASKTKAEPVQDQNIDIAGAATTSTSSPSSGNSDVVSAEHASDVDTEVHQTGTSECKSDSQKAFCSSTFLDTFDNLADFGTLAEAGDSYNLFLSEMYSG

>Pp00307G00320

MASGAGFLSAEAKAASAGVGVGVVVGNHGSAGLKGSASSSRGGRWRPTTRQGGHHGGDAWVSQRVAVTGNNRAGWGHQQPGHQQQTQSASAAATTAGGLNAEFCGRRSTRLVSKFHHGRQKLYMGKHSPEADLAAQEILSAPSTPLNSTALLLEKWSHQLVGLEDFPYLLRELGNRGEWERALQGYEWMVQQVHLRSEWSKLASIMISTLGRLGKVEIALDVFNRAQKAGFGNNVYAYSAMVSAYGRSGRCREALKVFQAMKKAGCKPNLITYNTIIDACGKGGVDLKQALDIFDEMQKEGVEPDRITFNSLIAVCSRGGLWEDSQRVFAEMQRRGIEQDIFTFNTLIDAVCKGGQMELAASIMTTMRGKNISPNVVTYSTMIDGYGKLGCFEEAISLYHDMKESGVRPDRVSYNTLIDIYAKLGRFDDALIACKDMERVGLKADVVTYNALIDAYGKQGKYKDAACLFDKMKGEGLVPNVLTYSALIDSYSKAGMHQDVSNVFTEFKRAGLKPDVVLYSSLIDSCCKCGLVEDAVVLLQEMTQAGIQPNIVTYNSLIDAYGRYGQADKLEAVKANMPNSVQKIGERSMEVVRKPPPSQQNASDHTGVLAAVSVFHEMQQFGLKPNVVTFSAILNACSRCASLQEASVLLEQMRFFDGWVYGIAHGLLMGLREQVWVEAQRLFDEISRMDYATGAAFYNALTDVLWHFGQRQGAQEVVVAAKRRQVWENAWWRSEQQFCLDLHLMSVGAAQAMLHVWLLDLRALVWDGHALPRVLSILTGWGKHSKVAGVSTVKRAVELRLQEIKAPFQVGRYNEGRLVCAGHIVREWLGDPRTSKLLMLQDAIVKPNLQWNRLPDLGSSLKMPQLAEVAT

>Pp00327G00290

MVQLYMSSVEEQRETMVQPYVSSMDSGSTSGRQTPSCVVQQGSNTFETSNLWEEWTQASNGDDTVSTSNFLPEISSFTSSRLSFQQSDSLTTWMSGFPPLSQTALSPDLSHSSDPVDHPPAFMQEGLGPGDSILDYSPALTEMYPKSSSKHNSSDCLPYPAASAPDKKMTDHELGSAISLAYDRGTVSRQLLRALGPLSPSSPLALQNGLQNPLGDPWDASPSAMPWPMATTGHAYGPGATRTSIPDHLANAINHLEGIAPSSASHASKPRHTDIFIAPNGTFDSTPGGWTPQYYDGSVTTDESVKAMKLIASLREAGHAEATIGFCTESKPSFLRGGDRTTSPVDSFFGKCVGAKTSIKQACSGKHPLELEEIVDSENSELNPTQLKRSKLFENHPNALWSDQSMNGRELRSYSHLVGSSLTASQPMDIIAIGPALNTDGKPRAKRGSATDPQSVYARHRREKINERLKSLQNLVPNGAKVDIVTMLDEAIHYVKFLQNQVELLKSDELWIYATPNKYNGMDISDLSDMYLQELESRA

>Pp00391G00130

MDSRQGLSTMNPMLLQHALRSLCTDQRWVYAVFWRILPRNYPPPQWDTQAGIMDRSKTNKRNWILVWEDGFCNFPACSTAGTANEGLRGSSSSFLSTRQLPRGNDADRPDTMNPELFFKMSHEVYNYGEGLMGKVAVDNSHKWVYREPLENEMCFLSPWQGSLDPHPRTWDAQFKSGIQTIAVVAVQEGVLQLGSTQKIVEDLIFVLYMQRKFNPLHTVPSLFVPSQPASTAAGEGKRRTNELSSAICSNGGYHHGQWSMPEYERALPMPKGHVNGAQFLTPREPWRNSAAGTKRPSEAEPCAYTSGSFDPSRPECPSLPKALNTGHASPQGSTPQSIPPAFRPSMSSLQALLSKLPSVMCLEAVASTGLHPGNGFAVANFTTNRSPFPLPGPVASSSNGHDLPASVARSQNASQFLESFDHLGEFGIQEALENSDSSYNSFLNEICS

>Pp00421G00080

MDVRQGLPTMNHLLQHTLRRICTESQWVYAVFWRILPRNYPPPQWETDGGMDRSKGNKRNWILVWEDGFCSFSACAADAAKHTRRGLPFIQATQPQQRGSDDECEAMSPELFFKMSHEVYNYGEGLMGKVAADSSHKWIYREPVEHEVNFLAPWHSTLEPHPRTWEAQFKSGIQTIAVVAVQEGVLQLGSTKKIMEDLNFVLHLQRKFNYLQSIPGVFVPHPMSCAGKKRSTSDGSPPSDRYNWMTTPEVDRPTSISHLTARSIDQFLSPRIGPQLWRSSSDTTVLGVKRPPESEPLSLSFKDCSYMNGSDSFHHRPECPYPPKSLNTGHTSPQGGVSPTIPNLLPSMSSLQALLSKLPSVTPTEGEGTITGAAPTTSGFETGNTFVSNSRPPVSRPAVTKPVVADSRSNSSKVSPSIEIGNGHLEQVVLTKVETEPLQGKIGAVASSSPSSGNSDVASAEHASDDTEGHQTSGTSKFKDSHKFSSSFLDTFENLADFGTHLDVLESGDSYNSFLNEIYS

>Pp00543G00050

MDSTMQLLNSSNVSSELIDGQSGRGIIFSSFRLNEAQVQRISVGSTVLSGGQTRRSQLYSLSISGCPKGEGHKYLPSAHVCANASVDGAAEQSKNVPTAKDAVALLKIRVQQGIVIDSFSYVDILQRCLKQEDILLAKQVHVCIIKSGMEQNLYVANKLLRVYIRCGRLQCARQVFDKLVKKNIYIWTTMIGGYAEYGYAEDAMKVYSQMRREGGQPNEITYLSILKACCSPVSLKWGKKIHAHIIQSGFQSDVRVETALVNMYVKCGSIDDAQLIFDKMVERNVISWTVMIGGLAHYGRGQEAFHRFLQMQREGFIPNSYTYVSILNANASAGALEWVKEVHSHAVNAGLALDLRVGNALVHMYAKSGSIDDARVVFDGMVERDIFSWTVMIGGLAQHGRGQEAFSLFLQMERGGCLPNLTTYLSILNASAITSTGALEWVKEVHKHAGKAGFISDLRVGNALIHMYAKCGSIDDARLVFDGMCDRDVISWNAMIGGLAQNGCGHEAFTIFLKMQQEGFVPDSTTYLSLLNTHVSTGAWEWVKEVHKHAVEVGLVSDLRVGSAFVHMYIRCGSIDDAQLIFDKLAVRNVTTWNAMIGGVAQQKCGREALSLFLQMRREGFFPDATTFVNILSANVGEEALEWVKEVHSYAIDAGLVDLRVGNALVHMYAKCGNTMYAKQVFDDMVERNVTTWTVMISGLAQHGCGHEAFSLFLQMLREGIVPDATTYVSILSACASTGALEWVKEVHSHAVNAGLVSDLRVGNALVHMYAKCGSVDDARRVFDDMLERDVYSWTVMIGGLAQHGRGLDALDLFVKMKLEGFKPNGYSFVAVLSACSHAGLVDEGRRQFLSLTQDYGIEPTMEHYTCMVDLLGRAGQLEEAKHFILNMPIEPGDAPWGALLGACVTYGNLEMAEFAAKERLKLKPKSASTYVLLSNIYAATGNWEQKLLVRSMMQRRGIRKEPGRSWIEVDNQIHSFVVGDTSHPESKEIYAKLKDLIKRLKAEGYVPDTRLVLRNTDQEYKEQALCSHSEKLAIVYGLMHTPYRNPIRVYKNLRVCSDCHTATKFISKVTGREIVARDAKRFHHFKDGVCSCGDYW

>Pt00G09760

MHPPNNQQCKPILTLASTLISMAEACNSMSRLKQIHAHSLLAGLHDHSIILAKMLRFAAVSPSGDLAYAQRLFDQLPHPNTFFYNTLIRGYAKSSIPSYSLHLVNQMRQNGVDPDEFTFNFLIKARSRVRVNINRNLPLVVECDEIHGAVLKLGAWQVFNSMSRKSLVTWNSMISACANNRNPEDAFGLFSRMFNYGVAPDGVTFLAVLTAYAHVGLVDEGYRLFESMQRDHGIEARIEHYGCVVNMLGQAGWLEEAFELITSMPLPSDVVWGVLLAACRKHGDVYMGERVVKKLLELKPDGGYYTS

>Pt01G01950

MTVYYVTKVRSHHFIPKPLLNPFFSCLSLHSHSFSTHKSNPTSWNTTHTYVLSNPLLSLLENCKSFSQLKQIQAQMILTGLILDGFASSRLISFCAISESRNLDYCIKILNNLQNPNVFSWNAVIRGCVESENPQKGLVLYKRMLTRAGCRPDNYTYSFLFKVCANLVLSYMGFEILGQVLKMGFDKDMYLYNGIIHMLVSVGESGLAHKVFDEGCVRDLVSWNSLINGYVRRRQPREAMGIYQQMITEQVKPDEVTMIGVVSACAQLESLKLGREIHRYIEESGLNLKISLVNALMDMYVKCGDLEAGKVLFDNMRKKTVVSWTTMIVGYAKNGLLDMAGKLFHDMPEKNVVAWNAMIGSCVQANLSFEALELFREMQLSNMKPDKVTMLHCLSACSQLGALDTGMWTHNYIKKHNLSLDVALGTALIDMYAKCGNMTKALQVFNEMPRRNSLTWTAIIGGLALYGNVNDAIFYFSKMIDSGLMPDEITFLGVLTACCHGGLVEEGRKYFDQMKSRFNLSPQPKHYSCMVNLLGRAGLLEEAEELIKTMPMEADAMVWGALFFACGIHRNLLIGERAASKLLDLDPHDSGIYVLLANMYREAGKWEEAQNIRKMMMERGVEKTPGSSSIEVNGIINEFIVRDKSHPQSEQIYECFNLINKTIGVCYV

>Pt01G08350

MEEILSSSSSSSLMSFAQETSSTLQQRLQFFLHSRPEWWVYSIFWQASKDASGRLVLSLGDGHFRGNKKYASKESNKQNHSKFGFNLERKSLFNEDMDMDRLVEGDVAEWYYTVSVTRAFAVGDGILGRAFSSGAFIWLTGDHELQIYDCERVKEARMHGIQTFVCVSTPSGVLELGSPDLISEDWGLVQLAKSIFGADINAGSVPKQANQESQPQIPNRTVSNFLDFGMFSSPQKERTTCLENQKESDTRKEPSGQGRSSSDSGRSDSDAGFTENNIGFKKRGRKPSGKELPLNHVEAERQRRERLNHRFYALRSVVPNVSKMDKASLLADAATYIKELKSKVNELEGKLRAVSKKSKISGNANIYDNQSTSTSTMTNHIRPTPNYMSNNAMEVDVKILGSEALIRVQSPDVNYPAARLMDALRELEFSVHHASVSKVKELVLQDVVIIIPDGLVTEEVMRAAIFQRMQN

>Pt01G08600

MGSSVLKQKLKSLCCSNGWSYGVFWCFDQINSMLLTMEDAYYEEEMGAVVNSMLSEAHILGEGIVGQAASTGKHQWIFSDASDGGWDSAASIGGQDIFQDDSEIHRQFSSGIKTIAIISVESHGVVQFGSTLKILERDEFLDQTKRLFGEMENVDGLTSKANSPSSLKSESYDLNEWFDSFCNGNIMPMLGGNCNELTEMAYSSMDVTQSSAFTSDAEQDRMNPLCLDSSLPTNQLNTDVTTEAQMIFSSHPSAQFQQVSSQSPSMNKITTQTPCTSTWISGDSNLTSWESKFQSEMVVQDSTTVFSTERSMNNQHSGPSIHVTERETSLNRFPVEFNPDDLTIDLSKSGVTDNILEWFAPSPEHSISGTAAIMNGNLSQSGGATSASSGLIGDLLVHIPSKQPATSAQSSVTETYFSSGKEKSVSVTGAENDLFEGLGLVFRGGQTGHCWEDMMMPVARSGQITASTGVSECISELDVGSKVGPQKGLFSELGLEELLDSVSNSSYVTKYSIDDQLSNAKRRRVENSLVSSDKLQLVNASYPTSSRMMQPAYNLDKTKNLPSKQEVFPKSQVSLWIDDSYSVNTGSSGLPKPEELAKPTKKRARPGESTRPRPKDRQQIQDRIKELKQIIPDGAKCSIDALLDRTIKHMLFLQSVTKYAERLKQADEPKLIGQENRLLLKDNTTSSGGATWALEVADQSMVCPIIVEDLSQPGLMLIEMLCEDRGFFLETADVIKGFGLNILKGLMESRENKIWARFIVEANVHITRVEVFWYLLQLLERTGTSVMDSTKQPSNSMHGRIPELSSYQLPALPCPVSLTETIQ

>Pt01G10360

MATKLHNQERLPGNLKKQLAIAVRSIQWSYAIFWSISARQPGVLEWGDGYYNGDIKTRKTIQSIELDEDELGLQRSEQLRELYESLSVGEASPQARRPSAALSPEDLTDTEWYYLVCMSFIFDIGQGLPGTTLANGHPTWLCNAHSADSKVFSRSLLAKSASIQTVVCFPFMRGVIELGVTEQVLEDPSLINHIKTSFLEIPYAVAAKNSSARSEKELACATFNRETLDTKPIPVIGCGELDITSPNRNSNDQPAADLIMVEGLNGGASQMQSLQFMDDDHSVHHSLNSSDCISQTIVDPVKVVPILKNVKVNNQNLLDVQDCNHTKLTSLDLQKEDFHYQSVLSCLLKTSNPLILGPDVQNCHQESSFVSWKKAGSVHTHKLKSGTRQKVLKKILLEVPRMHVDGLLDSPEYNSNKVVVGRPEADENGASHALSERKQREKLNKRFMILKSIVPSISKVVDKVSILDETIEYLQELERKVEELGSNRELLEVLTKRKPQDTAERTSDNYGSNKIGNGKHSLTNKRKAPDIDEMEPDINHNVSKDGSAESITVSVNKEDVLIEIKCRWREGILLEIMDVASHLHLDSHSVQSSTMDGILSLTIKSKHKGLNATSIGTIKQALRRVAGKC

>Pt01G13540

MIGTPVLQKIRFLSLPEAPTQTTGLSLKLKEQECLSLMKRCKNMEEFKQVHAQVLKWENSFCASNLVATCALSDWGSMDYACSIFRQIDQPGTFEFNTMIRGYVNVMNMENALFLYYEMLERGVESDNFTYPALFKACASLRSIEEGMQIHGYIFKRGLEGDLFVQNSLINMYGKCGKIELSCSVFEHMDRRDVASWSAIIAAHASLGMWSECLSVFGEMSREGSCRPEESILVSVLSACTHLGALDLGRCTHVTLLRNIREMNVIVQTSLIDMYVKCGCIEKGLSLFQRMVKKNQLSYSVMITGLAMHGRGMEALQVFSDMLEEGLKPDDVVYLGVLSACNHAGLVDEGLQCFNRMKLEHGIEPTIQHYGCIVHLMGRAGMLNKALEHIRSMPIKPNEVVWRGLLSACKFHHNLEIGEIAAKSLGELNSSNPGDYVVLSNMYARAKRWEDVAKIRTEMARKGFTQTPGFSLVQVERKIYKFVSQDMSHPQCKGMYEMIHQMEWQLKFEGYSPDTSQVLFDVDEEEKRQRLKAHSQKLAMAFALIHTSQGAPIRIARNLRMCNDCHSYTKLISVIYQREITVRDRNRFHHFKDGTCSCRDYW

>Pt01G19180

MDSLGWDVDSQVLTNSSSLWSNQQHDGVDLGEDIFHQIQELQKTQTSAPQLNSGSERHQEVTRLVANTVLAKSSGAGTICNWGDASAQELYSSSLISKPFSVTCMADFSMGGQPRINTNGLQNSKACVSTGSLESLDCLLSATNSNTDTSVEDDGISMIFSDCRNLWNFAPNSSAAVSSGESENNTCNPGNKEMHCPVSELDETVSHCSSDQYGKNRDCSQTKPVSTKRSNDHCSELKMGLKHPFFDILQSECSNQEGGFRLISDNPPKSKKPRSDKRPSSSNINFQQPSSSISSSIEEPDPEAIAQMKEMIYRAAAFRPVNLGLEVAEKPKRKNVRISTDPQTVAARQRRERISDRIRVLQGMVPGGSKMDTASMLDEAANYLKFLRSQVKALENLGHKLDSVNCPQPTNIAFSSLPFNHSFPLQNHFPFQNPNHIHPSQG

>Pt01G21690

MQTIAVIPVCPHGVLQLGSSLAILENIGFVNNVKSLILQLGCVPGALLSDNHMEKEPTERIGMPISCGMALPVCFSGNYKVPSSTPSLADSCNQQIISSQEASRIVGQPSCSQTRQVQDDQHATSSAIHIPNATGILAKSCDDFREPKITSLMKPDNPFMGQLANGVVGAEVIPSNPGAWLNHQTASSSSRLGFNHRPIISQSNTNSSILKLLEQQIFSDVGAQNHVSHYKNESNGLTMAHPRTNEGHFLTSTGGSHISGQLPSEVGTKRRANSNLCSLLKPQKLADINHSSTLLAGGGTQNVGSSRAEDDHLSGLLDQSSASGILSGGSNLEYPHTDVKPTKKEATTMEKKIEGDLFQALNVQLTQPGEHVYLGENVLGSVNDCLMSASGSQNTVTVNAKREEPCAQPPSGDDLYDILGVEFKNKLLNGKWNNLLGDEPCVKTQDMVKDASTFMSIREANSDLFSLTGGVSDSNMFSDLGTDHLLDAVVSKAHSAAKQSSDDNVSCRTTLTKISMPSFPGGSATYGRIGMCDQVQRELISLPKRAGTIASSSFRSGCSKDDVGTCSQTTSIYGSQLSSWVEQGHNAGHDCSVSTAFSKKNDETSKPNRKRLKAGENPRPRPKDRQMIQDRVKELREIVPNGAKCSIDALLERTIKHMLFLQSVTKHADKLKQTGDSKLLNKESGLLLKENFEGGATWAFEVGSQSMVCPIIVEDLNPPRQMIVEMLCEERGFFLEIADLIRGLGLTILKGVMETRNDKIWARFAVEANRDVTRMEIFMSLVQLLEQTVKGSAPPVGALENGNTMVHHTFPQATSIPATGMPSSLQ

>Pt01G29430

MEPIGATAEGEWSSLSGMYTSEEADFMEQLLVNCPPNQVDSSSSFGVPSSFWPNHESTMNMEGANECLLYSLDIADTNLYHFSQVSSGYSGELSNGNVEESGGNQTVAALPEPESNLQPKRESKMPASELPLEDKSRKPPENSKKRSRRTGDAQKNKRNVRSKKSQKVASTGNNDEESNGGLNGPVSSGCCSEDESNASQELNGGASSSLSSKGTTTLNSSGKTRASKGAATDPQSLYARKRRERINERLRILQNLVPNGTKVDISTMLEEAVQYVKFLQLQIKLLSSEDLWMYAPIAYNGMDIGLDHLKLTTPRRL

>Pt01G31650

MKSLHALFKPPSSPTKTTTSSSSTATTASKTRKPATSSSSSNAPIPSPNPPEITTTISKTSENKPKSSLSALFIPPTTPTEAHFISLIHGSKTILQLHQIHAQIIIHNLSSSSLITTQLISSSSLRKSINHSLAVFNHHKPKNLFTFNALIRGLATNSHFFNAIFHFRLMLRSGIKPDRLTYPFVLKSMAGLFSTELGMAIHCMILRCGIELDSFVRVSLVDMYVKVEKLGSAFKVFDESPERFDSGSSALLWNVLIKGCCKAGSMKKAVKLFKAMPKKENVSWSTLIDGFAKNGDMDRAMELFDQMPEKNVVSWTTMVDGFSRNGDSEKALSMFSKMLEEGVRPNAFTIVSALSACAKIGGLEAGLRIHKYIKDNGLHLTEALGTALVDMYAKCGNIESASEVFGETEQKSIRTWTVMIWGWAIHGHSEQAIACFKQMMFAGIKPDEVVFLALLTACMHSGQVDIGLNFFDSMRLDYCIEPSMKHYTLIVDMLGRSGQLKEALRFIERMPMNPDFVIWGALFCACRAHKKTKMAKFALNKLLKLEPTHTGNYIFLSNAYAALGQWEDAERVRVLMQNRGVHKNSGWSCIEVEGQVHRFVSGDHDHKDSKAICLKLEEIMAGAVKQGYIPGTEWVLHNMEQEEKEDVLGSHGEKLALAFALICTSPGMTIRIVKNLQVCGDCHSLMKYASKISQREIMLRDMKRFHHFKDGSCSCRDHW

>Pt01G39780

MVGSGGADRSKEAVGMMALHEALRSVCLNSDWTYSVFWTIRPRPRVRGGNGCKVGDDNGSLMLMWEDGFCRGRVGDCLEEIDGEDPVRKAFSKMSIQLYNYGEGLMGKVASDKCHKWVFKEPTECEPNISNYWQSSFDALPSEWTDQFESGIQTIAVIQAGHGLLQLGSCKIIPEDLHFVLRMRHTFESLGYQSGFYLSQLFSSTRNTSSSSSIPTKQSAIPTRSTQPLFNWGQRPLPSAAPSLLSSQNFQNPAARLGFPQAKDEPHMFILPHSSETRMEEMMGEHENDIKWPNGLSFFNALTGRADDAKLLFNPESLGNKGDRNHHPHILEGKSPNPNSDASNMNNAGGMNPNEFLSLDCHPDSARKMENKFKRSFTLPARMTSSSSTSVDHHQHHPVEYRNPESGVYSDVMETFLE

>Pt02G04060

MGADLHNTLRSLCFNTDWNYAVFWKLKHRARMVLTWEDGYYENCEQHDAFESKCFSQTQEKLHGGHYTRDPLGLAVAKMSYHVYSLGEGIVGQVAVSGKHQWIFADKYAASSFSSHEFSDGWQSQFSAGIKTIVVVAVVPYGVVQLGSSNKVIEDVNLVTRIKDVFFTLQDSSVRHVSGPLQHSMKNALCPKTAAGLRNKQVLEISTPTNDESIKLLHLRSNASYLDHQSQLGMNIISDQMYGGETNVWKDLGRRSEHNVTMHSNSFMKDKVNPSDLILPNDKLGADLAGIPADLFDATICESDGTNLYPKLVLDAPESSNITLKKDLEKKLDHQAESTHFNASDTFFKFSAGCELLEALGPSFINRCMPFDYQAGKSEAVNGFEMPEGMSSSQMTFDFGTENLLEAVVGNACHSGSDVKSEKSSCKSVQSLLTVEKMPEPSIQTKHIFNSAGYSINPSSVVEEDAQNFSNSTEVFGGMSSKGFLSTCTSICTEQLDKHAEPAKNSKKRAKPGEKFRPRPRDRQLIQDRIKELRELVPSGSKVRHVPCKGPVSVSVLRGLGNLLHMFTHGNFKFSILHVVGSDFWSLLSMHQKGTDASKYEQGSSWAVEVGGHLKVSSIIVENLNKNGQMLVEMLCEECNDFLEVAEAIRSLGLTILKGITEVHGEKTWICFVVEGQNNRTMHRMDILWSLVQILQPKTTN

>Pt02G05410

MAAPLSSRLQTMLQAAVQSVQWTYSLFWQMCPQQGILVWGDGYYNGPIKTRKTVQPMEVSTEEASLQRSQQLRELYDSLSIGETNQPERRPCAALSPEDLTETEWFYLMCVSFSFSPGAGLPGKAYDRKQHVWLTGANDIDSKTFSRAILAKSAGVQTVVCIPLLDGVVEFGTTDKVKEDLGFIQHVKSFFSDHHHLPPPKPALSEHSTSNPATSSDHPYLYSPPIPPFYVAADPPANAGQMNEDDEEEEEDEEDDEDDEEDQESDSEAETSREALLEPCQAQNPLQVAVVEPSELMQLEMSEGIRLGSPDDGSNNLDVDFPLINPESLMDHQSRADSFRTESARRWAPMLQDNPFSGSLQPSASGSPTLEDLAQEDTHYSQTVSTILQNQTIELAAEPSLNAYEAYSNQSAFAKWMNLTDHCLNVPVETTSQWLLKYILFTVPYLHSKYREENSPKSRDGDATNKFRKGTPQDELSANHVLAERRRREKLNERFIILRSLVPFVTKMDKASILGDTIEYVKQLLKKIQDLEACNKQMESEQRSRSVDPPQTITTSTSLKEQNNGITVVDRARSVGPGSDKRKMRIVEDYTTGRAQPKSVDSLPSPEPMVDVEPEISVEVSIIESDALIELKCGYREGLLLDIMQMLRELRIETIAVQSSSNNGIFVGELRAKVKENVSGKKLSIVEVKRAIRQIIPHD

>Pt02G11920

MNTQAMEAFRDGELWNFSRMFSMEEPDCTPELLGQCSFLQDTDEGLHFTIPSAFFPAPESDASMAEDESLFYSWHTPNPNLHFDSQESSNNSNSSSSVFLPYSSHESYFFNDSNPIQATNNNSMSMDIMDEENIGLFMPLFPEIAMAETACMNGDMSGDKTGDLDDNLKPAANDVLAKGLQLKRKLDVPEPIANTLDDMKKKARVTRNVQKTRKVGQSKKNQKNAPDISHDEEESNAGPDGQSSSSCSSEEDNASQDSDSKVSGVLNSNGKTRATRGAATDPQSLYARKRRERINERLKILQNLVPNGTKVDISTMLEEAVHYVNFLQLQIKLLSSDDLWMYAPLAYNGIDIGLNQKLSMFL

>Pt02G15940

MAHVVQSQQAILEKLRKQLAIAVRSVQWSYAIFWSLSTRQKGVLEWGGGYYNGDIKTRKVQATELKADKIGLQRSEQLRELYKSLLGGDAGQQAKRSSPALSPEDLSDEEWYYLVCMSFVFNPGEGLPGRALANKQTIWLCNAQYADSKVFSRSLLAKSASIQTVVCFPYLEGVMELGVTELVTEDPSLIQHIKASLLDFSKPDCSEKSSSAAHNGDDDEDPMSTKISHEIVDSLVLENLYTPTDDIELEQEGINDLHGNLREEFKRNSPDDCSDGCEHNHQTEDSMHEGLNGGVSQVQSWHFMDDEFSDDVLDSMNSSECISEAVVKQGKAVLSSKEKNVTRLQSQVFQEGNHTKLSSFDLGADDDLHYRRTVCVIMKSSSQSIENPCFRSGDHKSSFFSWKKRAVDGVMPRVQQNMLKKILFAVPLIYGGHSLRFDKENGGTDCLKKLEGCETCKEHYKSDKQRVNDKFIVLRSMVPSISEIDKESILSDTINYLKQLESRVAELESCKGWIDHEAGHRRSYMDMVDQTSDNDDIKKIDNGKRSWVNKRKALDIDEAELELDGVSPKDGMPLDLKVCTKEKEVLIEIRCPYREYMLLDIMDEINKLQLDVHSVQSSTLDGIFALTLKSKFRGAAVAPAGMIEQALWKIAGKT

>Pt02G17210

MGGGAWNDEDKTMVAAVLGSKAFNYLLSNSVANQNLLMVMCGDENLQNKLSDLVDCPNSSNFSWNYAIFWQISCSKSGDWVLGWGDGSCREPKEGEESEFTRILNIRLEDETQQRMRKRVIQKLQTLFGESDEDNYALGLDRVTDTEMFFLASMYFSFPRGEGGPGNCYASGKHVWISDALKSGPDYCVRSFLARSAGFQTIVLVATDVGVVELGSVRSVPESIEMVQSIRSWFSTRSSKLKELRRFLNGSRLAFPGTRNRLHGSSWAQSFGLKQGTPGEVYGSQATANNLKELVNGVREEFRHNHYQGQKQVQVQIDFSGATSGPSGIGRPLGAESEHSDVEASCKEERPGAADDRRPRKRGRKPANGREEPLNHVEAERQRREKLNQRFYALRAVVPNISKMDKASLLGDAISYINELQAKLKKMEAERGKLEGVVRDSSTLDVNTNGESHNQARDVDIQASHDEVMVRVSCPMDSHPASRVIQALKEAQVTVIESKLSAANDTVFHTFVIKSEGSEQLTKEKLMAAISFVWSCRGKVRWLSSTIPD

>Pt02G17690

MEELIISPSSPSSPVSLSQETPPTLQQRLQFIVQNQPDWWSYAIFWQTSNDDSGRIFLGWGDGHFQGSKDTSPKPNTFSNSRMTISNSERKRVMMKGIQSLIGECHDLDMSLMDGNDATDSEWFYVMSLTRSFSPGDGILGKAYTTGSLIWLTGGHELQFYNCERVKEAQMHGIETLVCIPTSCGVLELGSSSVIRENWGLVQQAKSLFGSDLSAYLVPKGPNNSSEEPTQFLDRSISFADMGIIAGLQEDCAVDREQKNARETEEANKRNANKPGLSYLNSEHSDSDFPLLAMHMEKRIPKKRGRKPGLGRDAPLNHVEAERQRREKLNHRFYALRAVVPNVSRMDKASLLSDAVSYINELKAKVDELESQLERESKKVKLEVADNLDNQSTTTSVDQSACRPNSAGGAGLALEVEIKFVGNDAMIRVQSENVNYPASRLMCALRELEFQVHHASMSCVNELMLQDVVVRVPDGLRTEEALKSALLGRLE

>Pt02G21460

MVACLSSTLPDALIISSRRRSPFLNSHYHTSSTCWPKFGSAEALLLLQNCTSFNHLKLVHGKIIRNALSANQLLVRKLIHLCSSYGRLDYAALLFHQVQEPHTFTWNFLIRTYTIHGYSMKALLLYNLMIRRGFPPDKFTFPFVVKACLASGSIRKGKEVHGLAIKTGFSKDMFLYNTLMDLYFSCGDEGYGRKVFDKLRVRNVVSWTTFIAGLVVCGDLDAARRAFDQMPTRNVVSWTAIINAYVRNQRPHEAFELFWRMLLANVKPNEYTLVNLLKACSELGSLKLGRWIHDYALKNGFDLGAFLGTALIDMYSKCGSLDDARQVFREMQIKSLATWNAMITSLGVHGYGEEALSLFTKMEEANVRPDAITFVGVLCACLQTDKVREGDMYFKYMREHYGITPVVEHYTCMIELYSRANLLNALDELVKGMPRELSDDVAAAWIKSRLIDDIDDTENSLEHQGEELQYWETRTEHLSQYQLQGFKWDVG

>Pt02G21490

MANRAEDDDPSIIQVYTCTSLIKQCGTSLLSLKTFHASMLKSHLHRNLHFLTNLIAQYASLGSVSYAYSLFSSTPSADLFLWNVMIRGLVDNSHYHHAILLYKQMLRLGIQPDNFTFPFIIKACSCLRHFEFGIRVHQDVVKFGYQSQVFISNSLITMYGKCDKYELSRQVFDEMPDKNAVSWSAIIGACLQDDRCKEGFSLFRQMLSEGSRPSRGAILNAMACVRSHEEADDVYRVVVENGLDLDQSVQSAAAGMFARCGRVEVARKLFDGIMSKDLVTWATTIEAYVKADMPLEALGLLKQMMLQGIFPDAITLLGVIRACSTLASFQLAHIVHGIITTGFFYNQLLAVETALIDLYVKCGSLTYARKVFDGMQERNIITWSAMISGYGMHGWGREALNLFDQMKASVKPDHITFVSILSACSHSGLVAEGWECFNSMARDFGVTPRPEHYACMVDILGRAGKLDEACDFIERMPVRPNAAVWGALLGACRIHLNVDLAEMVARALFDLDPHNAGRYVILYNIYTLTGKRKEADSIRTLMKNRGVKKIAGYSVIEIKNKLYAFVAGDRSHPQTDLIYSELERLMDRIRQEGYTPDINFVLHDVDEETKESMLYLHSEKLAIVFGLLNLGPGSVIRIRKNLRVCGDCHTATKFISKVTGREIVVRDAHRFHHFKNGACSCRDYW

>Pt02G22030

MANRAEDDDPSIIQVYTCTSLIKQCGTSLHSLKTFHASMLKSHLHRNLHFLTNLIAQYASLGSVSYAYSLFSSTPSADLFLWNVMIRGLVDNSHYHHAILLYKQMLRLGIQPDNFTFPFIIKACSCLRHFEFGIRIHQDVVKFGYQSQVFISNSLITMYGKCDKYELSRQVFDEMPDKNAVSWSAIIGACLQDDRCKEGFSLFRQMLSEGSRPSRGAILNAMACVRSHEEADDVYRVVVENGLDFDQSVQSAAAGMFARCGRVEVARKLFDGIMSKDLVTWATTIEAYVKADMPLEALGLLKQMMLQGIFPDAITLLGVIRACSTLASFQLAHIVHGIITTGFFYNQLLAVETALIDLYVKCGSLTYARKVFDGMQERNIITWSAMISGYGMHGWGREALNLFDQMKASVKPDHITFVSILSACSHSGLVAEGWECFNSMARDFGVTPRPEHYACMVDILGRAGKLDEACDFIERMPVRPNAAVWGALLGACRIHLNVDLAEMVARALFDLDPYNAGRYVILYNIYTLTGKRKEADSIRTLMKNRGVKKIAGYSVIEIKNKLYAFVAGDRSHPQTDLIYSELERLMDRIRQEGYTPDINFVLHDVDEETKESMLYLHSEKLAIVFGLLNLGPGSVIRIRKNLRVCGDCHTATKFISKVTGREIVVRDAHRFHHFKNGACSCRDYW

>Pt02G22060

MVACLSSTLPDALIISSRRRSPFLNSHYHTSSTCWPKFGSAEALLLLQNCTSFNHLKLVHGKIIRNALSANQLLVRKLIHLCSSYGRLDYAALLFHQVQEPHTFTWNFLIRTYTIHGYSMKALLLYNLMIRRGFPPDKFTFPFVVKACLASGSIRKGKEVHGLAIKTGFSKDMFLYNTLMDLYFSCGDEGYGRKVFDKLRVRNVVSWTTFIAGLVVCGDLDAARRAFDQMPTRNAFELFWRMLLANVKPNEYTLVNLLKACSELGSLKLGRWIHDYALKNGFDLGAFLGTALIDMYSKCGSLDDARQVFREMQIKSLATWNAMITSLGVHGYGEEALSLFTKMEEANVRPDAITFVGVICACLQTDKVREGDMYFKYMREHYGITPVVEHYTCMIELYSRANLLNALDELVKGMPRELSDDVAAAWIKSRLIDDIDDTENSLEHQGEELQYWETRTEHLSQYQLQGFKWDVG

>Pt02G23960

MNHSSSMLSPVAGCSALPPNSGTKPKNTPNRRRHFISLLQNCKHNNQIPPIYAKIIRNHHHQDPFVVFELLRVCSNLNSIGYASKIFSHTQNPNVYLYTALIDGLVLSCYYTDGIHLYYQMINSSLVPDSYAVTSVLKACGCHLALKEGREVHSQVLKLGLSSNRSIRIKLIELYGKCGAFEDARRVFDEMPERDVVASTVMINYYFDHGIKDTVCWTAMIDGLVRNGESNRALEVFRNMQREDVMPNEVTIVCVLSACSELGALQLGRWVRSYMDKHRIELNHFVGGALINMYSRCGDIDEAQRVFEQMKEKNVITYNSMIMGFALHGKSVEAVELFRGLIKQGFTPSSVTFVGVLNACSHGGLAELGFEIFHSMAKDYGIEPQIEHYGCMVDLLGRLGRLEEAYSFIRMMKVAPDHVMLGALLSACKIHGNLELAERVAKSLVACKNADSGTYILLSNAYSSSGKWKEAAEVRTNMREEGIEKEPGCSSIEVNNEIHEFLLGDLRHPQKEKIYKKLEELNQILRLEGYTPATEVVLHDIEKSEKEWALAIHSERLAICYGLISTKPLTTLRVVKNLRVCNDCHLTIKLISNITRRKIVVRDRNRFHHFENGVCSCGDYW

>Pt02G25880

MYAVMVKTNTNQDCYLMNQFISALSTFNRMDYAVLAYTQMEIPNVFVYNAMIKGFVQSYQPVQALELYVQMLRANVSPTSYTFPSLIKACGLVSQLRFAEAVHGHVWRNGFDSHVFVQTSLVDFYSSMGRIEESVRVFDEMPERDVFAWTTMVSGLVRVGDMSSAGRLFDMMPDRNLATWNTLIDGYARLREVDVAELLFNQMPARDIISWTTMINCYSQNKRFREALGVFNEMAKHGISPDEVTMATVISACAHLGALDLGKEIHYYIMQHGFNLDVYIGSALIDMYAKCGSLDRSLLMFFKLREKNLFCWNSVIEGLAVHGYAEEALAMFDKMEREKIKPNGVTFVSVLSACNHAGLIEEGRKRFASMTRDHSIPPGVEHYGCMVDLLSKAGLLEEALQLIRTMKLEPNAVIWGALLSGCKLHRNLEIAQVAANKLMVLEPGNSGYYTLLVNMNAEVNRWGEAAKIRLTMKEQGVEKRCPGSSWIEMESQVHQFAASDKSHAASDEIYSLLAELDGQMKLAGYVPELCEAETVEQRRSSISHEKGNVPVNPGCFGANTEASILWTWYQRSGCDRRHLLYRRTFYGALYMIALAAGSGGTKPNASTMGLINLMIFNLKKTKSYHSLIALGYALPTMGVAVSIIVFLVSTPFYRNKLPPVSLFTRMDQVIVASVRKWKVPVPGDPKQLHELSLDEYTGSGKFGIAYTSSLGFLDKAAVESGSRSPWMLFPVTQVEETKQMIKVLPVWAATFIPSTILAQVHTLFIKQGTVLDRSMGPHFEIPPACLAAFVTISMLISLAIYDRYFVLMARHYMKRPRGTTLLQRMGIGFMLHVIVMITACLAERKRLSVAREHNIISKNEVVPLSIFILLPQFVLMGVADNFVEAAKIEFFYDQAREGMKSLGNSYIATFLGIGSFLSSFLLSTVSKITKKHA

>Pt03G00750

IVGQAASTGKHQWIFSDASDGGWNSAASIGGQDIFQTIAVISVESQGLVQFGSTQKGSTLKADRVNLTKFPTRLHL

>Pt03G03160

MPFSLGFEIHLPSTKHHLHSSFLKKMISMACSQSSPVTENPPLSLFETCKSMYHLKQIHSRTIKTGIICNPIIQNKILSFCCSREFGDMCYARQLFDTIPEPSVFSWNIMFKGYSRIACPKLGVSLYLEMLERNVKPDCYTYPFLFKGFTRSVALQLGRELHCHVVKYGLDSNVFAHNALINMYSLCGLIDMARGIFDMSCKSDVVTWNAMISGYNRIKKDVISWTAIVTGFVNTGQVDAARKYFHKMPERDHVSWTAMIDGYLRLNCYKEALMLFREMQTSKIKPDEFTMVSVLTACAQLGALELGEWIRTYIDKNKVKNDTFVGNALIDMYFKCGNVEMALSIFNTLPQRDKFTWTAMVVGLAINGCGEEALNMFSQMLKASVTPDEVTYVGVLSACTHTGMVDEGKKFFASMTARHGIEPNIAHYGCMVDLLGKAGHLKEAHEIIKNMPMKPNSIVWGALLGACRIHKDAEMAERAIEQILELEPNNGAVYVLQCNIYAACNKWDKLRELRQVMMDRGIKKTPGCSLIEMNGIVHEFVAGDQSHPQTKEIYGKLNKMTSDLKIAGYSPNTSEVFLDIAEEDKENAVYRHSEKLAIAFGLINSGPGVTIRIVKNLRMCIDCHHVAKLVSKVYDREVIVRDRTRFHHFRHGSCSCKDYW

>Pt03G08710

MTINHKYYKHSLSLNKFLKSKKNLSKTPPKLFPNDPTPQQITQTISNSSNAPTLDLNHPILQKLEQSCTNIKQFNQIHTQLTVLGLFQHPFAASRYIKKLCACLNSVSHCVSLYNHIEEPDAFMCNTIMRSFVNVNDPFGALRFYYEKMIAKWVLPNHYTFPLVAKVCADIGSLREGQKVHALVVKFGFELDLFVRNSFIRFYSVCGRTSDARMVFDNGFVLDLVSWNSMIDGYVKNGELGLAREIFDEMYERDIFTWNSMISGYVGVGDMEAARGLFDKMPSRDVVSWNCMIDGFARIKDVSMAAKFFDEMPLRNVVSWNVMLALYLRCKKYSDCLRFFDMMVGGDFVPDEASLVSVLTACAELKMLDQGKWVHSYMKDNGIKPDMLLSTALLTMYAKCGAMDLAREVFDKMPEKSVVSWNSMIIGYGIHGHGDKALEMFREMEKGGPMPNDATFMSVLSACSHSGMVWNGWWYFDLMHRKYRIQPKPEHYGCLVDLLGQAGLKEPSEDLTRKTHTEVEPTLWGDLLSACRAHCISEPGEILAKQLIKLFPNHVVPYLLLSNTYVAEGRWDDVENLRMTLKNKTLNSKMVFTSRRHSMLSEAAAQIKLSNIDSIRT

>Pt03G09220

MTDYRLPPTMNLWTDDNGSVMEAFMNSSDLSSLWAPPPQTSASFSTPAAAAAAQPSDKTMLNQETLQQRLQALIEGARESWTYAIFWQSSYDCSGASVLGWGDGYYIGEEDKGKGRMKNSASSAAEQEHRKKVLRELNSLIAGPSSVTDDAVDEEVTDTEWFFLVSMTQSFVNGSGLPGQALFNGSPVWVAGSERLGTSPCERARQGQVFGLQTLVCIPSANGVVELGSTELIFQSSDLMNKVKVLFNFNSLEVGSWPIGTTNTDQGENDPSSLWLTDPETKDGNAGIPSTTPAHQTANNNNHHSSSSLTDHSGGIHHVQNHHSHQQQQQQQIHTQSLFTRELNFGEHSTYDGSTVRNGNSHLMKPESGEILNFGESKRSPSSANGNFYSGLVTEESNKKKKSPASRGGNEEGMLSFTSGVILSSSGLVKSSGGTGGDSDHSDLEASVVKEADSSRVVEPEKRPRKRGRKPANGREEPLNHVEAERQRREKLNQRFYALRAVVPNVSKMDKASLLGDAISYINELKTKLQSAESSKEELENQVESMKRELVSKDSSSPPNQELKMSNDHGGRLIDMDIDVKISGWDAMIRIQCCKMNHPAARLMSALKDLDLDVQYANVTVMNDLMIQQATVKMGNRYYTQEELKVAISTKVGDAR

>Pt03G12800

MKWLIITLNSNFDMANELRNQERLPDNLKKQLALAVRSIQWSYAIFWSNPTGQPGVLEWADGYYNGDIKTRKTVQSIELNADELGLQRSEQLRELYESLSAGEANPQARRPSAALSPEDLTDTEWYYLVCMSFVFDNGQGLPGTTLANGHPTWLCNAPSADSKIFSRSLLAKSASIQTVVCFPFMRGVVELGVSEQVLEDPSLIQHIKTSFLEIPYTVTANHSSAKSDKELACATFNREIHDTKPVPVIRCRELDTLSPDDNSNDQAATDSIMVEGLNGGASQVQSWQFMDDDFSNRVHHPLNSSDSVSQTIVDPVMLVPFLKDGKVNGQSLQDIQDCNHKKLTALNLQSDDLHYQSVLSCLLKTSHPLILGPNVQNCYQEPSFVSWKKAGLMHSQKLKSGTPQKLLKKILFEVPRMHVDGLLDSPEYSSDKVVGGRPEADEIGASHVLSERRRREKLNKRFMILKSIVPSISKVDKVSILDDTIQYLQELERKVEELECRRELLEAITKRKPEDTVERTSDNCGSNKIGNGKNSLTNKRKAPDIDEMEPDTNHNISKDGSADDITVSMNKGDVVIEIKCLWREGILLEIMDAASHLHLDSHSVQSSIMDGILSLTIKSKHKGLNAASVGTIKHALQMVAGNLFSNR

>Pt03G14490

MGLVLKEKLKSLCCSNGWSYGVFWCFDQRNSMLLTMEDAYYEEEMGVVVNNMLSEARMLGEGIVGQAASTGKHQWIFSDASDGGWNSAASIGGQDIFQDDSEIHRQFSSGIKTIAVISVESQGLVQFGSTQKILESEEFLGQTKRLFGKMENINGLTSNSDSPSSLNCESYDLNEWFDSFCNGNITPMLGDNCNELMEIAYSSMNFTQPSAITSVVEQDRMIPLCLDSSHPTNQLKTSEAQMILSCNPKTQSQHLSSQSPSMNKTTALTPCTSTWSNAGSNLTSLESKLGYEMVVQDSPTVFSTERSMSNLHSAPSIHVTEGELSEREMSQNRFPLEFKPDDFPTDLSNSCVVDNILEWFAPSPEHSISGMAPMMNGNLSQPGGVTPASPGLIGDILVDIPLKQPATLAQSSVTESYLSNGKEKCASITGTENDLLEGLGLVFGGGQARHCWEDIMVPVASSGHTTASTGISECISELDVDSKVGPRKGLFSELLDSVSNSNYVTKSSSDDQLSNAKRRRVENSSVNGNQLQLVNASCPTSSRVMQPAYNFDKTKNLLSKQEMFPKAQTVLWIDDSYSVNTGSSGLTKSKKPEEPAKANKKRARPGESTRPRPKDRQQIQDRIKELKQIIPDGAKCSIDALLDRTIKHMLFLQSVTKYAEKLKQADEPKLIGQHNRLLPKDNSTSSGGATWALEVADQSMVCPIIVEDLSQPGLMLIEMLCEDRGFFLEIADVIKGFGLNILKGLMESREDKIWARFIVEVAHQNLDFKANMQITRVEVFWSLLQLLERTGASVMDSTNQPSNVMHGRIPELNSYQLPALPCPVSLTETIQ

>Pt03G14730

MEEILSPSSSSSLISFAQETSSTLQQRLQFFLHSRPEWWVYSIFWQASKDASGRPVLSWGDGHFRGNKKYSSKVSNKQNHPKFGFNIERKSLFNEDMDLERLVDGDVAEWYYTASVTRVFAVGDGILGRAFTSGSSIWLTGDRELQIFECERVTEARMHGIQTFVCVSTPSGVLELGSPVFISEDWSLLQLAKSIFGAEINANPVPKQSNHESQPQISNCNVSNLLDIGLFSSPQTERTSSLENKKEVFGQGRSSSDSGRSDSDAGFRENHIGFKKRGRKPGGKESPLNHVEAERQRRERLNHRFYALRSVVPNVSKMDRASLLADAVNYIKELKRKVNELEANLQVVSKKSKISSCANIYDNQSTSTSTMVNHIRPPPNYMSNNAVEVDVKILGSEGLIRVQSPDINYPAARLMDALRELEFPVHHLSVTRVKELVLQDVVIRFDDGLVTEEAMRAAIFQRMQN

>Pt03G22390

SLVSSSSILISLIAHSSHLKHIHQTHAFMLLRALDTDNLLLSRFIHACSSLGFYSYAYSLFTSITHAPDIYLYNNIIKALSSSPTHPKASIFLYNNIQLAGLRPDSYSFPFALKAVTRFSSIQTGRQLHSQSIRFGLHSDLHVLTAFVQMYSSFGSGCICDARKMFDGMSMSTGDVALWNAMLNGYAKHGDLCNARDLFERMPQRNVISWTALITGYAQANRPHDAIALFRRMQLENVEPDEIAMLVALTACARLGALELGEWIRHYIDRLGLLTTNIPLNNALIDMYAKSGDIKSALQVFENMNHKTIITWTTMIAGLALHGLGTEALEMFSRMERARVKPNDITFIAILSACSHVGLVQTGRWYFNRMISRYGIEPKIEHYGCMIDLLGRAGHLKEAQTLLAQMPFEPNAVIWGSLLAACNTHGDPELGELALQHLLELEPDNSGNYALLSNIYASRGRWNESRVVRKVMWDAGVKKMPGGSLIEVNNRVHEFIAGEISHSQFDRIQEVLSKINRQLGLSQHFEKESGALLELG

>Pt04G08890

MALAKDRMGSVQTCPYNGNVMGDFSSMGSYGFDEYQKVAFYEEGNSTFEKTSGLMIKNLAMTSSPSSLGSPSSAISGELVFQATDHQAEEAHSLISFKGIGFDNIMHNNGSLLSFEQSNRVSQTSSQKDDYSAWEGNLSYNYQWNEMNPKCNTSPRLMEDFNCFQRAGNFISMTGKENHGDWLYAESTIVADSIQDSATPDASSFHKRPNMGESMQALKKQCNNATKKPKPKSAAGPAKDLQSIAAKNRRERISERLKVLQDLVPNGSKVDLVTMLEKAISYVKFLQLQVKVLATDELWPVQGGKAPDISQVKEAIDALLSSQTKDGNSSSSPK

>Pt04G12550

MQISVPVRAPTWVSTRRIFQQKLQDLHKCTSLNHIKQVHAQILKQNLHQDLYVAPKLISAFSLSQEMTLAINVFKQIPDPNVHLYNTFIRACVQNSHSLLAFETFFEMQRNGLFADNFTYPFLLKACDGQSWLPLVKMIHNHLEKYGFFQDLFVPNSLIDSYCKCGLLGVKSAMRLFKVMDERDVVSWNSMIRGLLKVGELSEACKLFDEMPMKDAVSWNTILDGYVKAGEMNKAFGLFESMPERNVVSWSTMVSGYCKAGDMEMARMLFDRMPVKNLVSWTIIVSGYAVKGLAKDAIRSFEQMEEAGLKPDDGTVISILASCAESGLLGLGKRVHTSIERIRYKCSVNVSNALVDMYAKCGQVDRALSVFNGMSKKDLVSWNCMLQGLAMHGHGEKALQLFSIMRQEGFRPDKVTLVAVLCACVHAGFVDEGIRYFNNMERDYGIVPHIEHYGCMVDLLGRGGRLKEAYRLVQSMPVEPNVVIWGTLLGACRMHNAVGLAEEVLDCLFKLEPSDPGNYSLLSNIFASAGDWSSVANVRLQMKNFGIQKPSGASSIEVDDEVHEFTVFDKSHPKSDKIYQMINRLGLDLKRVHVVPK

>Pt04G20590

MRFSAIVSGSKLPNWILRIKESSANGKWQEVVSHYHEIKKAGIQTVDVSVFPPILKAWSFLSHRHGKSLHACLIKQGFDSFTSIGNSIMGFYIRCGDFDIAVDVFNSMRRSRDSVSWNILIHGHLDNGALVAGLWWFTNARVAGFEPNISTMVLVIQACRILGTKHDGLILHGYLIKSGFWAISSVQNSLLSMYVDADMECARELFDEMHEKDVIAWSVMIGGYLQWEEPQVGLQMFRKMVLVPGIEPDGVVMVSVLKACASSRDVCTGRLVHGLVIHRGFDCDLFVENSLIDMYSKCKDAGSAFKVFNEISQRNNVSWNSMLSGFVLNENYSEAQSLISSMRKERVETDEVTLVNILQICKYFVHPFHCKSIHCVMIRRGSEANELVLSALIDAYAKCYLIEIAWEVFARMRRRDVVSWSTMISGFAHCGKPDEAIAVYQEMDRDLVKPNVITIINLLEACSVTAELKRSKWAHGVAIRQGFASEVTVGTAVVDMYSKCGEILASRRAFDQLALKNIVTWSAMIAAYGMNGLAHEALALFAEMKRHGLKPNPVTTLSVLAACSHGGLVEEGLSLFKSMVQELGLEPGFEHYSCMVDMLGRAGKLDTAIEVIKAMPDNLKNGASIWGSLLSACRSYGLTELGKEAISRVLELEPSNSAGYLVASSMYAADGLWDDAARIRVLAKEKGVKVVAGYSLVHIDNKACRFVAGDGSHPRSDEIFSMAQQLHDCIKIDEKKEGNTWLAVIECLT

>Pt04G23700

MMMMMSMSMNMSSSSSSVQVQMHVEPFHFEKCQSMSQLRQYHSQIIRLGLSSHNHLIPPLINFCARASTSDALTYALKLFDSIPQPDAFLYNTIIKGFLHSQLLPTNSILLLYSHMLQNSVLPNNFTFPSLLIACRKIQHGMQIHAHLFKFGFGAHSVCLNSLIHMYVTFQALEEARRVFHTIPHPDSVSWTSLISGYSKWGLIDEAFTIFQLMPQKNSASWNAMMAAYVQTNRFHEAFALFDRMKAENNNVLDKFVATTMLSACTGLGALDQGKWIHEYIKRNGIELDSKLTTAIVDMYCKCGCLEKALQVFHSLPLPCRWISSWNCMIGGLAMHGNGEAAIQLFKEMERQRVAPDDITFLNLLTACAHSGLVEEGRNYFSYMIRVYGIEPRMEHFGCMVDLLGRAGMVPEARKLIDEMPVSPDVTVLGTLLGACKKHRNIELGEEIGRRVIELEPNNSGRYVLLANLYANAGKWEDAAKVRKLMDDRGVKKAPGFSMIELQGTVHEFIAGERNHPQAKELHAKVYEMLEHLKSVGYVADTNGVLHGHDFDEEEDGENPLYYHSEKLAIAFGLSRTKPGETLRILKNLRICEDCHHACKLISTVFDREIIVRDRTRFHRFKMGQCSCQDYW

>Pt05G01100

MKLSRLNYLSKPFKIPTFTLKPTLTLPILETHLQKCQNIKQFNQILSQMILSGFFKDSFAASRLLKFSTELPFININQSYQIFSHIENPNGFICNTMMKGYMQRNSPCKAIWVYKFMLESNVAADNYTYPILFQSCSIRLAEFDGKCIQDHVLKVGFDSDVYIQNTLINMYAVCGNLSDARKVFDGSSVLDMVSWNSMLAGYVLVGNVEEAKDVYDRMPERNVIASNSMIVLFGKKGNVEEACKLFNEMKQKDLVSWSALISCYEQNEMYEEALILFKEMNANGIMVDEVVVLSVLSACSRLLVVITGKLVHGLVVKVGIETYVNLQNALIHMYSSCEEVVTAQKLFSESCCLDQISWNSMISGYVKCGEIEKARALFDSMPDKDNVSWSAMISGYAQQDRFTETLVLFQEMQIEGTKPDETILVSVISACTHLAALDQGKWIHAYIRKNGLKINIILGTTLINMYMKLGCVEDALEVFKGLEEKGVSTWNALILGLAMNGLVDKSLKTFSEMKEHGVTPNEITFVAVLGACRHMGLVDEGHRHFNSMIQEHKIGPNIKHYGCMVDLLGRAGMLKEAEELIESMPMAPDVSTWGALLGACKKYGDNETGERIGRKLVELHPDHDGFNVLLSNIYASKGNWVDVLEVRGMMRQHGVVKTPGCSMIEAHGRVHEFLAGDKTHPQNEHIEHMLDEMAKKLKLEGYAPDTREVSLDIDEEEKETTLFRHSEKLAIAFGLIAIDPPTPIRIVKNLRICNDCHTAAKLISKAFNREIVVRDRHRFHHFKQGSCSCMDYW

>Pt05G01940

MSRACREIERNILRLLHGRETRTQLREIHAHFLRHGLNQLNQILSHFVSICGSLNKMAYANRIFKQTQNPTIILFNAMIKGYSLNGPFEESFRLFSSMKNRGIWPDEYTLAPLLKACSSLGVLQLGKCMHKEVLVVGFEGFSAIRIGVIELYSSCGVMEDAEKVFDEMYQRDVIVWNLMIHGFCKRGDVDMGLCLFRQMRKRSVVSWNIMISCLAQSRRDSEALGLFHDMLDWGFKPDEATVVTVLPICARLGSVDVGKWIHSYAKSSGLYRDFVAVGNALVDFYNKSGMFETARRVFDEMPRKNVISWNTLISGLALNGNGELGVELLEEMMNEGVRPNDATFVGVLSCCAHAGLFERGRELLASMVEHHQIEPKLEHYGCMVDLLGRSGCVREAYDLIRIMPGGAPNAALWGSLLSACRTHGDVELAHLAVKELIDLEPWNSGNYVLLSNMYAEEERWDKVANVRGMMREKNVKKTPGQSVIG

>Pt05G20260

LANGPIDLRQRIHSAVFPAQKRDNAAFDKNCTSMKDLQKIHAQLIKTGLAKDTIAASRVLAFCTSPAGDINYAYLVFTQIRNPNLFVWNTIIRGFSQSSTPHNAISLFIDMMFTSPTTQPQRLTYPSVFKAYAQLGLAHEGAQLHGRVIKLGLENDQFIQNTILNMYVNCGFLGEAQRIFDGATGFDVVTWNTMIIGLAKCGEIDKSRRLFDKMLLRNTVSWNSMISGYVRKGRFFEAMELFSRMQEEGIKPSEFTMVSLLNACACLGALRQGEWIHDYIVKNNFALNSIVITAIIDMYSKCGSIDKALQVFKSAPKKGLSCWNSLILGLAMSGRGNEAVRLFSKLESSNLKPDHVSFIGVLTACNHAGMVDRAKDYFLLMSETYKIEPSIKHYSCMVDVLGRAGLLEEAEELIKSMPVNPDAIIWGSLLSSCREYGNIEMAKQAAKRVNELDPNESSSFILLSNVYAAHNHFEEAIEQRLSLKEKQMDKEPGCSLIEVNGEVHEFVAGGRLHPRSKDIYHALDDLGLTLKEMG

>Pt05G20860

MATPPSSRLQTMLQAAVQSVQWTYSLFWQMCPQQGILVWGDGYYNGPIKTRKTVQPMEVTTEEASLQRSQQLRELYDSLSIGETNQPARRPCAALSPEDLTETEWFYLMCVSFSFPPGGGLPGKAYARRRHVWLTGANEIDSKTFSRAILAKSARVQTVVCIPLLDGVVEFGTTDKVQEDLGLIQHVKTFFSDHHHRHLTPPKPALSEHSTSSPATSSHDHPRFHPPPIPPFYVAAEPSANAEQIDEDEEEDEEEEEHDSDSEAETSRDDHLEPRQAQNPHQVVAAEPSELMQLEMSEDIRLGSPDDGSNNLDSDFPLTGPDNSMDHRSRADSYKAESARRWTMLQDNPFSGNLQPSASGPPPLEDLAQEDTHYSQTISTILQSQPVWLAAEPSSIAYEARYHQSAFSRWTNRSDHLFHVSVETTSQWLLKYILFSVPHLHSKSREDNSPKSRDGEAASRFRKGTPQDELSANHVLAERRRREKLNERFIMLRSLVPFVTKMDKASILGDTIEYVKQLRQKIQDLETRNKQMESEQRPRSVDRPQRTSTSDSLKKQKSGVTVVDRARSLGPLPDKRKMRVVEDSAGGGAKPKTVGALPQPEPVVHKELETSVEVSIIESDALLELECGFREGLLLDIMQMLRELRIETIAVQSSLNNGIFAGELRAKVKENVNGKKVSIVEVKRAIHKIIPHD

>Pt05G22250

MGMGTDLHDTLRSLCFNTDWNYAVFWKLKHRARMVLTWEDGYYDNCEQHDALENKCFRQTQENLHGGHYPRDPLGLAVAKMSYHVYSLGEGIVGQVAVSGKHQWIFADKHVTNSFSSYEFSDGWQSQFSAGIRTIVVVAVVPYGVVQLGSLNKVSEDVNLVTHIKDVFFALQDSTVSHVTSPSQHGMKNALCLKTAAELKNKQEVLEIPTPTNDESIDLLNLKSNASYLDHRSQLGMNIISDRMFGGETSVWKDLGRGSEHNTTMHSNSFMRENVSLSDLVLPNEKLGADLAGFPADLFDSTICDRDKSDSINLRPNVVLNAPESSDITFKRDLEKKLDHPAESTHFNSSDTFFKFSAGCELLEALGPSFLNRCMPFDYQTRKSEAGNIFEMPEGMSSSQMTFDFGSENLLEAVVGNVCHSGSDVKSEKSGCKSVQSLVTAEKLPEPSIQTKHIMNSAGYSINQSSVVEEDVHNLSNSTEVCGGMSSKGFSSTCPSTYSEQLDKRSESAKNSKKRAKPGENCRPRPRDRQLIQDRIKELRELVPNGSKCSIDSLLERTIKHMLFLENITKHADKLNKCAEPKMHQKGTEASNYEQGSSWAVEVGGHLKVSSIIVENLNKNGQMLVEMLCEECSHFLEIAEAIRSLGLTILKGITEVQGEKTWICFVVEVCLNLLDELSVTRGTMISFSTSELYIKAMSSNIYRSEPSLSNRLQCCLFFCFLISFAF

>Pt06G06870

MSFKKKKFTLPPIIKALDKLPSPPPQPPLPPLPKPPPTTHLAPLPTLTLTPTSNHTNLLHFLNSHLTKIQPLTPQNLLHFFKTKLHHHPHFSHYDFHIFNWVSTIDSFSHDHQTFEWMARTLAITNRLEELALLLQFMSSNPCPCSEGIFSCPRIEPIFQFCINAYCKARKLDDAFLAFECMRKLIDGRPSVVVYNILINGCVKCGEHDRAIGVYDRMLKDRVKPDVFTFNILISSYCRNYMFELALELFREMKEKGCSPNVVSFNTLIKGFFRERKFEEGVKMVYEMIDLGCEISSVTFEILVDGLCKEGQASEACGLLIDFTRKGVLPRKFDSFGLVDMLCKKRMADRALEVLDELWRNGNIPSMISCTTLIEGLRKSGRREEAFGLMERMLKENIVPDIMTFNCLLHDLCNEGRTVDGNKLRLLASRKGLDVDEMTYDILVSGCIREGKRKEGEALVDEMLDKEFIPDLATYNRFIDGLSKTRSSAQ

>Pt06G09000

MGTTDLRQLLESLCNNSDWNYAVLWKMRYGSPMILTWEDGYFDCPKPREPLQTISSDVYCNGGNDLASSLRDASASNANFGGHQIELVVADMLHLQYPLGEGVVGEVAYTGDHFWLSFNNIFSCEMSKNLVPEFPEEWLLQFASGIKTILLVPVLPHGVLQLGSFDEVAEDIQIVAYIKGRFNDLHSTRENAVPLTLKREFKAQSTLISCPVEQLNATSAISISQVKSEDSNYSIPVNSVKLHKDEQPEVFKCESKNNSLSPIFADVSPPSESLSASQPGMVESKIFELSYLMDELQAYSDCNEYNVGWFGEPLDGMMNTYPTADMVEQSSGGMDANDVYHKNRQSFLSFPKGSELHKVLGPPFLSQTNEKTWEPSLLVEDSCKSSNFIFSEDHSARIEPSLFAREGEVEFLLEPVAGNSYSSSDNASSNRSHSLKSSEMLSGHLLATSQNQFQTRTLVGDDLAPWNHLASVCISGSGNTDTTAALDSMMSTIFDQEQQEKDQSYKHPWKGQKMSNVARRRARPGENQKPRPRDRQLIQDRVKELRELVPNGSKCSIDGLLDQTIKHMQYLRSVTDQAEKLRQWVHQEVADRKNCRLSETNVNIQSGKSWAFEFGNDLQICPIVVEDLAYPGHLLIEMLCNDRGVFLEIAQVIRSLDLTILKGVMESRLSNTWAHFIVEACKGFHRLDIFWPLMQLLQRKRSSISGKI

>Pt06G10260

MDYHSTTNPSSSGSSSAPKNGREKRTGGKKSNGGVKLSTDPQSVAARERRHRISDRFKILQSLVPGGTKMDTVSMLEEAINYVKFLKNQILLHQTIMNSVDDERSLDYHLPAGSASLPTEQPSYLDSNLASVVHPSSSLPCPDSYFQADENYTHYDAFDSKNYYF

>Pt06G10570

MSSLILNKSAHIKIRLFQGFNSLKHLKHVHAALLRLGLDEDSYLLNKVLRFSFNFGNTNYSHRIFHQTKEPNIFLFNTMIHGLVLNDSFQESIEIYHSMRKEGLSPDSFTFPFLLKACARLLDSKLGIKLHGLVVFDDIPEKNVAAWTAIISGYIGVGKCREAIDMFRRLLDMGLRPDSFSLVRVLAACTRIGDLRSGEWIDDYITKIGMVRNVFVATSLVDFYVKCGNMERACSVFDGMLEKDIVSWSSMIQGYASNGLPKEALDLFFKMLNEGFRPDCYAMVGVLCACARLGALELGNWASNLMDRNEFLGNPVLGTALIDMYAKCGRMDSAWEVFRGMRKKDIVVWNAAISGLAMSGHVKAAFGLFGQMEKSGIEPDGNTFVGLLCACTHAGLVDEGRQYFNSMERVFTLTPEIEHYGCMVDLLGRAGFLDEAHQLVKSMPMEANAIVWGALLGGCRLHRDTQLVEGVLKQLIALEPSNSGNYVLLSNIYSASHKWEDAAKIRSIMSERGIKKVPGYSWIEVDGVVHEFLVGDTSHPLSEKIYAKLGELVKDLKASGYVPTTDYVLFDIEEEEKEHFIGCHSEKLAIAFGLISTAPNDKIRVVKNLRVCGDCHEAIKHISRFTGREIIVRDNNRFHCFNDGSCSCKDYW

>Pt06G13560

MMDEYLNHLISSSSLVDGDVKESSSWVCSEPNQPNAFLPTSLELYQDDKKNSPVSMISSNQSVESLATQDTSSVVLGSESDYAVDKVLISEQARLQNDCQNCNGNPSPDGMARGNLKFGNTGLQCNGILPTLSSLNYPNQLPIVGDLTSYLSFSEASNAGCNGREQSEYLRSLKNLQNLSSIPQLWPSQSYEGVSSLPPLMGQDRIEGSGLRGGNLDDDMHIMGKGYMGMDEILRLDKLSASPTTEGKEDLQSCPFSSGIAEPNVNMSMNQLSSMPQTTSAAPVEGCNGTGKTRVRARRGHATDPHSIAERLRREKIAERMKNLQELVPNSNKVDKASMLDEIIEYVKFLQLQVKVLSMSRLGAAGAVIPLLTDGQPEGHNSLSLSPSAGLGIDISPSADQIAFEQEVLKLLESDVTMAMQYLQSKGLCLMPIALAAAISSVKASLSGTTSEERKNNGYTSGLVSSSSSITGIDTHPMSNDNNIATGTLSSKGMIVNGCNEVVKQEVLKNT

>Pt06G18660

MRPCSREMQGMNSLLNPSSQIPLQDLQNQQIQNSHFDPNSSSNDDFLEQMLSAIPSCSWADPKSPWDLNPPTNLPFPTNNNSSSAKPRDLFNETPPSNTDNNNVGFHDNFDESVILASKLRQHQISGGSGAAAAAKMMLQQQLLMAAARGGLSQNDDIDVSPTQGGDGSMQGLFNGFRAGSMNGTVRASNQSMQHFNHPQGGAMQSPNLGAQGAATTAVMNQPQASGSNGGAPAQPRQRVRARRGQATDPHSIAERLRRERIAERMKALQELVPNANKTDKASMLDEIIDYVKFLQLQVKVLSMSRLGGAAAVAPLVADMSSEAGGDCIQANANGGSIARTTNGNQTASTNDSSLTVTEHQVAKLMEEDMGSAMQYLQGKGLCLMPISLATAISTATCHNRTSGIINSHNPLLQSNGEGPTSPSMSVLTVQSATMGNGVAKDAASVSKP

>Pt06G20750

MHQFMQGNPLSLTSPSMPLNLSSSNPSISLLIKSCKTIPHLHQFHAHIIHKGLEQDHFIIAHFLSISTSVSHSTSIFNRLLNPSTFLYNILLKIFSKNSQFIETFSLFYRMKQSEYALPDKYTYPLLIKVCSNELRLKEGEIVHGSAIRCGVSDDVYVGSSLISFYGKCKEILSARKVFDEIPERNVVSWTAMVAGYASVGDLENAKRVFERMPERNLPSWNAMISGLGKAGDLSGARKVFDEMVERNVVSFTVMIDGYAKVGDMASARALFDEAPEKDVVAWSALISGYSRNEQPNEAVKIFFEMVSMNVKPDEFIMVSLMSACSQLGNSDLAKWAHVLAALIDMHAKCGNMEKAVKLFQDMPSRDLIPCCSLIQGLSIHGRGVEAVELFNRMLDEGLIPDTVAFTVILTACSRGGLIEDGWHFFDTMKNKYSVVPSPDHYACMVDLLSRAGQLRAAYDLLKSMPLKPHACAWGALLGACKLHGDVELREEVANRLLELEPEKAGSYVLLSNIYASANQWLDVSIVRDEMKERGIRKIPGCSYIFTEA

>Pt06G23120

MILSRNLISLSQTKQAHARIIVSGLAGKASLMGHILSFLATFPSSPFDYSLSIYRTIKNPNVFASNNMIRCFAKSDLPLQSLVLYSSVLRNCVRPNNYSFTFLLQACSKGLGLVEGVQVHGHVLKLGFGEDVYVRNALIHLYSSCCRTESSKQVFDESPHHCDVVTWNAMLAGFARDGQVSVVQKLFDEMPERDVISWNTMLMAYVHNGKLGEALECFKRMRESGLVPDEATLVTMLSASAQLCLLEHGQSIHSIIDSLSLPMTISIGTALLDMYAKCGCIEQSRLLFENMPRRDVSTWNVMICGLASHGLGKDALTLFERFLNEGLHPMNVTFVGVLNACSRAGLVKEGRHYFQMMTDSYGIEPEMEHYGCMVDLLGRAGLVFEAIKVIESMAISPDPVLWAMVLCACRIHGLAELGEKIGNRLIELDPTYDGHYVQLASIYANSRKWEDVVRVRRLMAERNTSKVAGWSLIEARGKVHRFVAGHREHEQSLEIQKMLEIIETRLAAAGYVPNVSPVLHDIGEEEKENAIKVHSERLAIAFGLLVTGPGSCIRIVKNLRVCWDCHEVTKMISRVFEREIIVRDGSRFHHFKEGKCSCLDYW

>Pt06G24530

MATLTLPNIFFSSLSPSIHKPPTLNPKTSHSVLRPHWIIDLLKSCSNIREFSPIHAHLITANLIHDPEITSQVLAFLLSVNNLDCAHQILSYSHEPESIIWNTLLENKLKEGCPQEVLECYYHMVTQGVLLDISTFHFLIHACCKNFDVKLGSEVHGRILKCGFGRNKSLNNNLMGLYSKCGKLKEVCQLFEKMTHRDVISWNTMISCYVLKGMYREALDLFDEMLVSGVLPDEITMVSLVSTCAKLKDLEMGKRLHLYIVDNKLWIRGSLLNCLVDMYSKCGKMDEAHGLLSRCDESEVDVVLWTTLVSGYVKSNKIDKARQLFDKMNERSLVSWTTMMSGYVQGGYYCESLELFQQMRFENVIPDEVALVTVLSACVHLEDFDLGRSVHAFIVTYGMLVDGFLGNALLDLYAKCGKLDEALRTFEQLPCKSAASWNSMLDGFCRSGGVDKARDFFNKIPEKDIVSWNTMVNAYVKHDLFNESFEIFCKMQSSNVKPDKTTLISLLSSCAKVGALNHGIWVNVYIEKNEIGIDAMLGTALIDMYGKCGCVEMAYEIFTQIIEKNVFVWTAMMAAYAMEGQALEAIDLYLEMEERGVKPDHVTFIALLAACSHGGLVDEGYKYFNKLRSFYNIIPTIHHYGCMVDLLGRVGHLEETVKFIERMPIEPDVSIWSSLMRACRSHHNVELAEQAFKQLIEIDPTNNGAHVLLSNIYADAGRWDDVSKVRTKLHETGVPKQPGFTMIEQNGVVHEFVASNLVSADILCMLQDIERRLLVKQELSDTTSQHSERLAVAFGLINNQENSPIRVVNSVRMCRDCHSVMKLISQAYDREIVIRDNYRFHRFTDGHCSCKDYW

>Pt06G27120

MVARGCQPNVYTYTTIINGLCKTGEAAEAAGLFKKMEEAGCQPDVVTYSTLIDSLCKDRLVNEALDIFSYMKAKGISPDIFTYTSLIQGLCNFSRWKEASALLNEMTSLNIMPDIVTFNVLVDTFCKEGKVLEALGVLKTMTEMGVEPNVVTYNSLMYGYSSCTQVVEARKLFDVMITKGFKPVVFSYNILINGYCKAKRIDEAKQLFNEMIHQGLTPNKVTYNTLIHGLCQLGRVREAQDLFRNMRTNGNLPNLRTYSILLDGFCKQGYLGKAFRLFRAMQSTHLKPNPVMYTILVHAMCKSGNLKDARKLFSELFVQGLQPNVQIYTTIIFGLCKEGLLDEALEAFRNMEEDGCPPDEICYRVIIRGFLQHKDESRAVQLIGEMRDRGFVADARTRLSEVG

>Pt07G07310

MLPRNPNLNLKRTLITFLDKCKSMLQLNQLHALVITFGLSQDDLFMSRIVSFSALSDSNNTDYSYRALLNLQDPTIFEWNSVIRGYSKSKNPNKSISVFVKMLQVGIYPDHLTYPFLAKATSRLLRKELGVSIHGHVIKSGFEIDRFVANSLIHMYGSCGDIVYARKVFDGTPVKNLVSWNSMVDGYAKCGYLDLARGLFDLMPERDVRSWSCLIDGYAKSGNYGDAMAVFEKMRTSGPKANEVTMVSVLCACAHLGALDKGRMMHQYLVDNGFELNLVLRTSLIDMYAKCGAVEEAFAVFRGVSLRKSDVLIWNAMIGGLATHGLVKESLDLYTEMQIAGIKPDEITFLCLLSACAHGGLVKQASYFFEGLGKNGMTPKTEHYACMVDVMARAGQVAEAYQFLCQMPLEPTASMLGALLSGCMNHGKLDLAELIGKKLIELDPEHDGRYVGLSNVYAIGRRWDEARIMREAMERRGVKKTPGYSFLEMSGAHHRFIAHDKSHPSSEQIYTMLSFIVSQMQFGVLKEGQEHCLYGIEGI

>Pt08G07080

MEDHFSPCWPAAPAEANWVQTSAAVYDESFLVPCPSHASASANFQVNGFPSWSIPIQEASENKAASNSKSHSQAEKRRRDRINAQLGILRKLIPKSEKMDKAALLGSAIDHVKDLKQKATEISRTFTIPTEVDEVTVDCDVSQATNPSSTNKDKDSTFIRASVCCDDRPELFSELIRVLRGLRLTIVRADIASVGGRVKSILVLCNKCSKEGGVSISTIKQSLNLVLSRIASSSVPSNYRIRSKRQRFFLPSHLSQQYT

>Pt08G10650

MSISSSTPYLSPVPSKTTALNSQQKQHLRKPDHAISLLQNCKNPKDLIQLHTLLIKTSLIKEKYAFGRLLLSFASFDNLGSLNYAQKLFDTVDIPRNSFMYTTMIKAYANFGNPREAFAFYSRMLCDQRYVYPNDFTFTYVFSACSKFNGVFEGKQAHAQMIKFPFEFGVHSWNSLLDFYGKVGEVGIVVRRVFDKIEGPDVVSWNCLINGYVKSGDLDEARRLFDEMPERDVVSWTIMLVGYADAGFLSEASCLFDEMPKRNLVSWSALIKGYIQIGCYSKALELFKEMQVAKVKMDEVIVTTLLSACARLGALDQGRWLHMYIDKHGIKVDAHLSTALIDMYSKCGRIDMAWKVFQETGDKKVFVWSSMIGGLAMHSFGEKAIELFAKMIECGIEPSEITYINILAACTHSGLVDVGLQIFNRMVENQKPKPRMQHYGCIVDLLGRAGLLHDAFRVVETMPVKADPAIWRALLSACKLHRNVELGEQVGRILIKMEPQNDMNYVLFSNVYAAVNRWDISGKLRREMKVRGMQKNPGCSSIELNGAVHEFVSRDHSHPQSQVIYELLHILTNHMVQEDHEPMMTIMAENQGIR

>Pt08G12140

MQQQRCLRKIANILPSFTTKSPHRILEEKFISLLQSCKTLKVLHQIHSQIITHGFEHEDYIAPKIISGYGSLKKMENAHKVFDQIPEPNASIWNAMFRGYSQNESHKDVIVLFRQMKGLDVMPNCFTFPVILKSCVKINALKEGEEVHCFVIKSGFRANPFVATTLIDMYASGGAIHAAYRVFGEMIERNVIAWTAMINGYITCCDLVTARRLFDLAPERDIVLWNTMISGYIEAKDVIRARELFDKMPNKDVMSWNTVLNGYASNGDVMACERLFEEMPERNVFSWNALIGGYTRNGCFSEVLSAFKRMLVDGTVVPNDATLVNVLSACARLGALDLGKWVHVYAESHGYKGNVYVRNALMDMYAKCGVVETALDVFKSMDNKDLISWNTIIGGLAVHGHGADALNLFSHMKIAGENPDGITFIGILCACTHMGLVEDGFSYFKSMTDDYSIVPRIEHYGCIVDLLGRAGLLAHAVDFIRKMPIEADAVIWAALLGACRVYKNVELAELALEKLIEFEPKNPANYVMLSNIYGDFGRWKDVARLKVAMRDTGFKKLPGCSLIEVNDYLVEFYSLDERHPEKEQIYGTLRTLTKLLRSSGYVPGLMELDERN

>Pt08G18960

MRGLDRAMERLRPLVDSNAWDYCVVWKLGDDPSRIHGEVVISAEPRWLCHATVTTHDSNTLREVAGTQVLIPVIGGLVELFAAKHMKKDEKMIESIRAHCHVPVKQEAVTELGYSNSSFNDHRLDSLLEENLPHSCHLLSLIPRTQFLLPLSQPRNSISFEGSSSGSNPSNEAPSFVSNASQLPQHGHLELSVGKSNHDEKILKQRAGSADCNKKVPKVMRRSERDDYKSKNLVTERNRRTRIKTGLFALRALVPKISKMDKAAILGDAIDYVGELLKEVKNLQDEIKNAEEEERRASNIELKTSKLEIFQEDHVSSSKINQDSSGFVEKKGAEVQLEVDQISKRQFLLKFLCEQRQGGFGRLMETIHSLGLQILDANITTFNGNVLNILKVEADKDIHPKTLKKSLIELTGNLIQTFGSQI

>Pt08G21700

MPCYHQARAVHIIQQLKNCKDFIFAISTHANLFKFGLLNDTITTNHLLNSYLRFRRIQYAHHLFDEMHEPNVVSWTSLMSGYVNMGRPQSALWLYTKMSESEVSPNGFTLATVINSCSILADLKTGKMVHAHVQILGLQGNLVVCSSLVDMYGKCNDVDGARMVFDSMSCRNVVSWTAMIAGYAQNGKGYEALEVFREFSSYMMERPNHFMLASVINACASLGRLVSGKVTHGAVIRGGYELNDVVASALVDMYAKCGSFLYSEKVFRRIRNPSVIPYTSMIVGAAKYGLGKLSLNLFEEMTDRKVMPNDVTFVGILHACSHSGLVDEGLRLLNSMHEKHGVMPDVRHYTCVVDMLSRVGRLDEAYKLAKSIRVNPNEGALLWGTLLSSSRLHGRVDMAVEASKWLIEYNQQVAGAYVTLSNTYTLAGEWENAHSLRTEMELVGVHKEPGCSWIEIKDSIYVFYAGDLSCERGDEVISLLRELERRMMERGCVGGSTGLVFVDVEQEVKEKIVGLHSERLALAFGLISIPKGVTIRVMKNLRICSDCHEAFKLISKIVERDFVVRDVNRFHHFKDGSCTCKDFW

>Pt09G01770

MGLLLREVLKTLCCVNQWCYAVFWKIGYQNPKLLIWEECHSESTLCSVSPSTSGTENLVLPFREREGYLGSEVHSSQFGVHEGNRLRLLINKMMANNQVIIVGEGIIGRAAFTGNHEWILANNYCKDAHPPEVLNEAHHQFSAGMQTIAVVPVCPYGVLQLGSSLAIPENIGFVNIVKSSILQIGCIPGALLSDNHMENESTERIGIPISCGMPLPVCFSGNYKVPNSTPYLADNFNPQIISSQAASRIVSRPSCSQPRQIQDNQLATSSAIHIHNVTKTLAKSCDDFCEPKIIPVMKPDNPFMGQLPNGVVGAEVVPSNPGAWLNQQTDSRPEFNHQPITSQSDANNNIIKLLDRQIFSDGGARNHVGHNKNESDSLTMSHVRTNGGLFLTSPGGSHISGQLPNEMGGQTRPHSIPCSLLKLQKLADINHSSTFLAGVGIQNAGSSRAEEVHLSSLLGRFSASGILSGSSNHEYHPTDVKPTKNEIPAMEKKVDSDLFQALNIPLTQPGEHIYLGEKILGPVNDCLKNASGSQNTVIVNAMLDEPCAQLPSGDDLYDILGVGFKNKLLNDQWNNLLREEACVKTQDMVKDALAFTSIREANSDIFSLNEGISDSNMFSDMGTDLLDAVVSRVHAAAKQSSDDNVSCKTSLTKISTSSFPSGSPTYGSIGMADQVQSELISLPGKAGTIASTSFRSGCSKDDAGSCSQTTSIYGSQLSSWVEQGHNALHDSSVSTAFSKKNDGTSKPNHKRLKPGENLRPRPKDRQMIQDRVKELREIVPNGAKCSIDSLLERTIKHMLFLQSVTKHADKLKQTGDSKLINKEGGLHLKDNFEGGATWAFEVGSRSMVCPIIVEDLNPPRQMLVEMLCEEKGFFLEIADLIRGLGLTILKGVMEARNDKIWACFAVEANRDITRMEIFMSLVQLLEQTVKGSAGPVGALENGDMMVHLAFPQTTSIPATGMPSGLQ

>Pt09G03760

MWAFQSPKTTLLPSNATFLPRPSLKPPICSITLNPTASTADNNKLIQSLCKQGNLTQALELLSLEPNPAQHTYELLILSCTHQNSLLDAQRVHRHLLENGFDQDPFLATKLINMYSFFDSIDNARKVFDKTRNRTIYVYNALFRALSLAGHGEEVLNMYRRMNSIGIPSDRFTYTYVLKACVASECFVSLLNKGREIHAHILRHGYDGYVHIMTTLVDMYAKFGCVSNASCVFNQMPVKNVVSWSAMIACYAKNGKAFEALELFRELMLETQDLCPNSVTMVSVLQACAALAALEQGRLIHGYILRKGLDSILPVISALVTMYARCGKLELGQRVFDQMDKRDVVSWNSLISSYGVHGFGKKAIGIFEEMTYNGVEPSPVSFVSVLGACSHAGLVDEGKMLFSSMHVAHGICPSVEHYACMVDLLGRANRLEEAAKIIENMRIEPGPKVWGSLLGSCRIHCNVELAERASIRLFDLEPTNAGNYVLLADIYAEAGMWDGVKRVKKLLEARGLQKVPGRSWIEVKRKIYSFVSVDEVNPRMEQLHALLVKLSMELKEEGYVPQTKVVLYDLKAAEKERIVLGHSEKLAVAFGLINSSKGEVIRITKSLRLCEDCHSFTKFISKFANKEILVRDVNRFHHFRDGVCSCGDYW

>Pt09G04470

MATLGNPLASVPISSNPTILTANNEQKSNPSTVPILIDKCANKKHLKQLHAHMLRTGLFFDPPSATKLFTACALSSPSSLDYACKVFDQIPRPNLYTWNTLIRAFASSPKPIQGLLVFIQMLHESQRFPNSYTFPFVIKAATEVSSLLAGQAIHGMVMKASFGSDLFISNSLIHFYSSLGDLDSAYLVFSKIVEKDIVSWNSMISGFVQGGSPEEALQLFKRMKMENARPNRVTMVGVLSACAKRIDLEFGRWACDYIERNGIDINLILSNAMLDMYVKCGSLEDARRLFDKMEEKDIVSWTTMIDGYAKVGDYDAARRVFDVMPREDITAWNALISSYQQNGKPKEALAIFRELQLNKNTKPNEVTLASTLAACAQLGAMDLGGWIHVYIKKQGIKLNFHITTSLIDMYSKCGHLEKALEVFYSVERRDVFVWSAMIAGLAMHGHGRAAIDLFSKMQETKVKPNAVTFTNLLCACSHSGLVDEGRLFFNQMRPVYGVVPGSKHYACMVDILGRAGCLEEAVELIEKMPIVPSASVWGALLGACRIYGNVELAEMACSRLLETDSNNHGAYVLLSNIYAKAGKWDCVSRLRQHMKVSGLEKEPGCSSIEVNGIIHEFLVGDNSHPLSTEIYSKLDEIVARIKSTGYVSDESHLLQFVEEEYMKEHALNLHSEKLAIAYGLIRMEPSQPIRIVKNLRVCGDCHSVAKIISKLYNRDILLRDRYRFHHFSGGNCSCMDYW

>Pt09G08900

MAEGEWSSLGGMYTSEEADFMAQLLGNCPNQVDSSSNFGVPSSFWPNHEPTTDMEGANECLFYSLDFANINLHHFSQGSSSYSGGSGILFPNTSQDSYYMSDSHPILANNNSSMSMDFCMGDSYLVEGDDCSNQEMSNSNEEPGGNQTVAALPENDFRAKREPEMPASELPLEDKSSNPPQISKKRSRNSGDAQKNKRNASSKKSQKVASTSNNDEGSNAGLNGPASSGCCSEDESNASHELNRGASSSLSSKGTATLNSSGKTRASRGAATDPQSLYARKRRERINERLRILQTLVPNGTKVDISTMLEEAVQYVKFLQLQIKLLSSEDLWMYAPIAYNGMDIGLDHLKVTAP

>Pt10G09640

MEEHLSPLAVTHLLQHTLRSLCVHENSQWVYAVFWRILPRNYPPPKWDGQGAYDRSRGNRRNWILVWEDGFCNFAASTAEINSGDCPSSSVYGNCEFQHYQGLQPELFFKMSHEIYNYGEGLIGKVAADHSHKWIYKEPNDQEINFLSAWHNTADSQPRTWEAQFQSGIKTIALIAVREGVVQLGAVHKVIEDLSYVVLLRKKFSYIESIPGVLLPHPSSSAYPYKVDGYGTVSDTWHYQGSNIAPQSPTEFYDHFNQVPFKITPSMSSLEALLSKLPSVVPPPQPAAAHHQAAGYCDSQSHYVSSIQRGMEKVAKEEIDEEYTRADQQDVGESSSSLSAADYRKQQFQQFQDLNVNVRNTSGYFE

>Pt10G16880

MATAPFPPVIQQSPSQHSSYYNTTYPSSLLSCLPKCTSLKELKQIQAFSIKTHLQNDLQILTKLINSCTQNPTTASMDYAHQLFEAIPQPDIVLFNSMFRGYSRSNAPLKAISLFIKALNYNLLPDDYTFPSLLKACVVAKAFQQGKQLHCLAIKLGLNENPYVCPTLINMYAGCNDVDGAQRVFDEILEPCVVSYNAIITGYARSSRPNEALSLFRQLQARKLKPNDVTVLSVLSSCALLGALDLGKWIHEYVKKNGLDKYVKVNTALIDMYAKCGSLDGAISVFESMSVRDTQAWSAMIVAYAMHGQGQDVMSMFEEMARAKVQPDEITFLGLLYACSHTGLVDEGFRYFYSMSEVYGIIPGIKHYGCMVDLLGRAGLLHEAYKFIDELPIKPTPILWRTLLSSCSSHGNLELAKQVMNQILELDDSHGGDYVILSNLCARAGKWEDVDTLRKLMIHKGAVKIPGCSSIEVDNVVHEFFSGDGVHYVSTALHRALDELVKELKSVGYVPDTSLVVHPDMEDEEKEITLRYHSEKLAISFGLLNTPPGTTIRVVKNLRVCGDCHSAAKLISSLIDREIILRDVQRFHHFKDGKCSCGDYW

>Pt11G05230

METKQPHKLPHSPAYNFLLQAGPRLYLLHQVHAHIIVSGYGRSRSLLTKLLNLACAAGSISYTRQIFLAVPNPDSFLFTSLIKSTSKSHNFSIYSLYFYSRMVLSNVSPSNYTFTSVIKSCADLSALKHGRVVHGHVLVHGFGLDVYVQAALVALYGKCGDLINARKVFDKIRERSIVAWNSMISGYEQNGFAKEAIGLFDRMKETGVEPDSATFVSVLSACAHLGAFSLGCWVHEYIVGNGLDLNVVLGTSLINMYIRCGNVSKAREVFDSMKERNVVAWTAMISGYGTNGYGSQAVELFHEMRRNGLFPNSITFVAVLSACAHAGLVNEGRRLFASIREEYHLVPGVEHNVCLVDMLGRAGLLDEAYNFIKEEIPENPAPAILTAMLGACKMHKNFDLGAQVAEHLLAAEPENPAHYVILSNIYALAGRMDQVEIVRNNMIRKCLKKQVGYSTVEVDQKTYLFSMGDKSHSETNAIYHYLDELMWKCSEAGYVPVSDSVMHELEEEEREYALRYHSEKLAIAFGLLKTSHGTPIRIVKNLRMCEDCHSAIKFISAISSREIIVRDKLRFHHFKVGSCSCLDYW

>Pt11G11640

MVGSGGADRSKEAVGMMALHEALRSVCLNSDWTYSVFWTIRPRPRVRSGNGCKVGDDNGSLMLMWEDGFCRGRVGDCLEEIDGEDPVRKSFGKMSIQLYNYGEGLMGKVASDKCHKWVFKEPTECEPNISNYWQSSFDALPPEWTDQFESGIQTIAVIQAGHGLLQLGSSKIIPEDLHFVLRMRHAFESLGYQSGSYLSQLFSPTRNTSSSSLPTKQSAIPTRPPPPLFNWSQRPLPSAASLLSSPNFQNHAARLGFPQAKDEPHMFTLPHSSETRVEEIMGEHENDIKWPNGLSFFNALTGRADNAKLLFSPESLGNKGDQNHHPLILEGKSPNPNSDASNMNNSDVMNPNEFLSLDSHPDSARKMENKYKRSFTFPARMTSSSSSTSIDHHQHQAVDYRNPEPGVYSDIMETFLE

>Pt12G04420

MKVLEQILAHAIQLGYEQSLFVAGKIIVFCAAAEHGDMSYAVSVFEKTENPDAFIWNTMIRGFGKTNDAPKAFEFYKRMQERGLVADDFTFSFLLKDIETSRQQIFPALVAWNTMIGSYVSCGKFKEALGMFSRMMELGVEPDEATLVETFSAGSSLGALDFGRWDHSCISNTDHGSIIEVNYSLLNMYAKCGALEEAYETFDGMSKKNTVTWNTMILGLAVMALQMMHCHGGMVDEGRRFFDVMKKEYHIQPNDKAQWVHGGYFGASWCNFVWRTLWAACGLHGNVELGKQLESVLPST

>Pt12G11080

MLISSSSHITPQNYHSIIIKLIKQCKTISQLHQLHAYTITNTPLSFHSSPSLLTKFLYTLTTKAKSKSSTSLLHYAKSIFNSIQNPSTFCYNTIIRVHTLLSFPIPALHFFTQMRRLSVPLDSHSFPFTLKACAQLGGVFLARCLHCQVLKFGFLSDLYVMNSLIHGYMVCDMSNDAYKVFDESPQRDVVSYNVLIDGFVKAGDVVKARELFDLMPVRDSVSWNTIIAGCAKGDYCEEAIELFDFMVDLEIRPDNVALVSTLSACAQLGELEKGKKIHDYIERNAMKVDTFLSTGLVDFYAKCGCVDIALKIFDSSSDKNLFTWNAMLVGLAMHGYGELLLEYFSRMIEAGVKPDGISILGVLVGCSHSGLVDEARKLFDEMESVYGVPREPKHYGCMADLLGRAGLIKKVMEMIKDMPSGGDMSVWSGLLGGCRIHGDVEIAEKAAKHLMELKPDDGGVYSILANVYANAERWEDVMNIRRSLSSNRVVTKIAGFSLIQLDGVAHEFIAGDSLHSESDKIYLVLNGIREHQSELS

>Pt13G00130

MPLSELLYRMAKGKTDSSQEKNPACSTDLSFVPENDFGELIWENGQIQYSRARKIQTCNSLPPKIRDKDIGNGTNTKTGKFGTMESTLNELLAVPAVEVRANQDDDMVPWLNYPLDEPLQHDYCSDFLPELSGVTVNEHSSQSNFPSFDKRSCNQSITDSHTVSVHNGLNLEQGDVVMNSSAGDIDAKRPRTSASQLYPSSSEQCKTSFPFFRSRDSTKKDDSTSNAVHHVIAPDSIRAPTSGGGFPSIKMQKQVPAPSPINSSLINFSHFARPAALVKANLQNVGMRASSGTSSMERMQNKDKGSIGLPKETDSHCRPNMMSSKVEVKPTEVKPAEGSVPAELPEEMSQEGDSKSDRNCHQNFGESAIKGLEDVEKTTEPLVASSSVGSGNSAERPSDDPTENLKRKHRDTEESEGPSEEIIYVQDAEEESVGAKKPASARAGNGSKRGRAAEVHNLSERRRRDRINEKMRALQELIPNCNKVDKASMLDEAIEYLKTLQLQVQIMSMGAGMYMPSMMLPPGMPHMHAAHMGQFLPMGVGMGMGFRMGMPDMNGGYSGCPMYQVPPMHGAHFPGSQMSGPSALHGMGGPSLQMFGLSGQGLPMSFPRAPLMPMSGGPPPKTNREPNACGVVGPMDNLDSATASSSKDAIQNINSQVMQNNVANRSMNQTSSQCQATNECFEQPAFAQNNGEGSEVAESGVLKSAGGTDITPSRATGCD

>Pt13G15200

MAKSLPFLTRPPNKVSSFIALCPQPFKIRPLDPPPFRFPASTLKFLETHYSSSLNLTTHFNNNKDDCNESTFKPDKFYASLIDDSIHKTHLNQIYAKLLVTGLQYGGFLIAKLVNKASNIGEVSCARKLFDKFPDPDVFLWNAIVRCYSRHGFFGHAIEMYARMQVACVSPDGFSFPCVLKACSALPALEMGRRVHGQIFRHGFESDVFVQNGLVALYAKCGEIVRANAVFGRLVDRTIVSWTSIISGYAQNGQPIEALRIFSEMRKTNVRPDWIALVSVLRAYTDVEDLEHGKSIHGCVIKMGLECEFDLLISLTSLYAKCGHVMVARLFFNQVENPSLIFWNAMISGYVKNGYAEEAIELFRLMKSKNIRPDSITVTSSIAACAQIGSLELARWMDEYISMSEFRNDVIVNTSLIDTYAKCGSVDMARFVFDRIPDKDVVVWSAMMVGYGLHGQGRESIILFHAMRQAGVSPNDVTFVGLLTACKNSGLVEEGWDLFHRMRDYGIEPRHQHYACVVDLLGRAGHLDRAYNFVMNMPIEPGVSVWGALLSACKIHRHVTLGEYAAERLFSLDPYNTGHYVQLSNLYASSRLWDCVAKVRVLMREKGLTKHLGYSVIEINGKLQAFQAGDKTHPRSKEIFEEVEDLERRLKEAGFVPHTESVLHDLNYEETEETLCNHSERLAIAYGLISTPPGTTLRITKNLRACDNCHAAIKLISKLVSREIVVRDACRFHHFKDGACSCGDYW

>Pt14G01710

MQTMGDFPDGEWDFSRMFSMEDPDFTPELLGQCSFPQENDEGLQFTIPPAAFFPTPEANVSMAGNESLFYSWNALNPNLHFDSQESGNNSNCSSGVFLPSSSHESYFFNDSNRIQAANDNSMSMDISLMDEKTIGLFMPFFPEIAMAETACMNGDMSSDKIGDLDDNLQPAANTVLAKGLQLKRKLDVPESNTLDDMKKKPRITRNVQKSKKSVQSKKNQRSTPKISNEEEESNAGPDGQSSSSCSSEDDNASKDSDSKVSEVLSSSGKTRASRGAATDPQSLYARKRRERINERLKILQHIVPNGTKVDISTMLEEAVHYVKFLQLQIKVKALVNFFLHAAK

>Pt14G02400

MEGGLPMLNCLLQHTLRSLCSCTDSSNPSKWVYAVFWRILPRNYPPPKWDYGGTALDRSKGNKRNWILVWEDGFCDVYECERAGTGYMKGRFGTDVFFKMSHEVYNYGEGLVGKVAADNSHKWVFKENPNESDPNLISSWNMSIEPQPRAWEFQFNSGIQQTIAIISVREGVIQLGSFDKIVEDLNLVISIQRKFSYLQSIPGIFAIQRPYLPIQHPYITKPNTHTIENQEIAFSVDDKRQITGVKRLFHESLDDFPIKAINMGWNSPQNGIPGPPIWSIPPLLPTMSCSLGALLSKLPSATPSYSNIEALGTSLLNNNNNNRTSISQRVRVDDLGVTREGQLVSSGHLDAAREEKPTSIKPSLNLPDKVVGFGHVREGESALSLN

>Pt14G08390

MAPVVQGQQVVPDNLRKQLAVAVRSVQWSYAVFWSQSTRQQGVLEWGDGYYNGDIKTRKVEAMELKADKIGLQRSEQLRELYESLLEGETGLQATRSSPALSPEDLSDEEWYYLVCMSFVFNPGEGLPGRALANKQPIWLCNAQYADSKVFSRSLLAKSASIQTVVCFPYLEGVIELGVTELVTEDPGLIQHIKASLLDFSKPVCSDKSFSAAHNKDDDKDPMSTRISHEIVDTLALENLYTPTEDIESEQEGINYLHGNVCEEFNRNSPDDFSNGYEHNLVTEDSFMLEDLKEGASQVQSWHSMDDEFSDDVRDSMNSSDCISEVFVKQGKVVPSSKGKDISHLQLKVLQEGNHTKLSSLDPGADDDLHYRRTAFVILKSSSQLIENPCFQSGDYKSSFVGWKKGAADGYKPRIQQKMLKKILFAAPLMHGGHSIRSDKENAGKDCLKNLEGCETCKLHFESEKQKENEKYLALESIVASINEIDKASILSDTINYPRQLESRVAELESCTGSTDYEARSRSYMGMVDRTSDNHGIKKPWINKRKARDIDEAELELDEVAPKDGMPVDLKVCMKEKEILIEMRCPYREYMLLDILDEANKRQLDVLSVHSSTLDGIFTLTLKSKRGSASFTRRDDQTSTSENCRQDLSRHLSTYQPALERFLHSVEISS

>Pt14G09970

MKIELGVGGGAWNDEDKTMVAAVLGTKAFNYLLSNSVANQNLLMAMCGDESLQNKLSDLVDRPNASNFSWNYAIFWQISCSKSGDWVLGWGDGSCREPKEGEESEVTRILNIRHEDETQQRMRKRVIQKLQTLFGESDEDNYALGLDQVTDTEMFFLASMYFSFPHGEGGPGKCYASGKHMWISDALKPGPDYCVRSFLAKSAGFQTIVLVATDVGVVELGSVRSVPESIEMVQSIRSWFSTRNSSIRAKPMAAAAAAAAAMPAVSEKKDENSPFSNFGIVERVGVPKIFGQDLNSNHGHGHGFREKLVVRKMEERPSWNAYQNGTRLALPGAQNGLHGSGWAQSFGMKQGTPSDVYGSQATANNLQELVNGVREEFRLNHYQPQKQVQMQIDFSGASSGPSVIGKPLSAESEHSDVEASCKEERPGTADDRKPRKRGRKPANGREEPLNHVEAERQRREKLNQRFYALRAVVPNISKMDKASLLGDAISYINELQTKLKVMEAEREKSGSISRDASALDANTNGESHNQAPDVDIQASHDELMVRVSCPLDSHPASRVIQAFKEAQITVVESKLSAANDTVFHTFVIKSQGSEQLTKEKLMAAFSRESSSLHSLSSTG

>Pt14G10370

MEGLIISPSSSSSLVSLSHETPPTLQQRLQFIVQSQPDRWSYSIFWQASKDDSGQIFLAWGDGHFQGSKDTSPKLSTTNNSRMSTSNSERKRVMKGIHSLLDECHDLDMSLMDDTDSTDTEWFYVMSLTRSFSPGDGILGKAYTTGSLIWLTGGHELQFYNCERVKEAQMHGIETLICIPTSCGVLELGSSCVIRENWGIVQQAKSLFVSDLNSCLVPKGPNNPCQEPIQFLDRNISLADGGIIAGLQEDDHTIEHGEKRTQERAETKKDNVNKLGQSYVDSEHSDSDFHFVAVNIERRIPKKRGRKPGLGRGAPLNHVEAERQRREKLNHRFYALRAVVPNVSRMDKASLLSDAVSYINEMKAKVDKLESKLQRESKKVKLEVADTMDNQSTTTSVDQAACRPNSNSGGAGLALEVEVKFVGNDAMIRVQSDNVNYPGSRLMSALRDLEFQVHHASMSSVNELMLQDVVVRVPDGLRTEEALKSALLGRLEQ

>Pt14G13210

MDLRISYTQKGERINHSLLTAAAFLYIDLLKNTSFGAASPAHVMRTFQELSPMINKIINLLNKCASPTHIHQIQSQIILHHLHSNTTLAYHFITASQNFSLLRSSLPLFFTHLHRPHVFICNTLIRAFSRIHDPHIPYSIYIHMHYNSILPNNYTFPFLFKSLSDCRDYLKCQCVHAHVIRLGHLNDIYVQNSLLDVYASCGYMGLCREVFDEMSDRDVVSWTVLIMGYRNAKNYADALIAFEKMQYAGVVPNHVTMVNALGACGSFRAIEMGVWIHDFISRNGWDLDVILGTSLVEMYLKCGRIDEGLNAFRSMKEKNVFTWNVVIQGLGFAKSGQEAVWWFNRMEEEGFEADEVTLANVLNACIHSGLVDMGRQIFSSLINGNYGFSPSLKHYACMIDLLTRAGCLDEAFKLIKEMPFEPSKSIWGSFLTGCRACGNLELSELAAKKLAEL

>Pt15G03490

MIINYLCRLKSNLKALQLGSFRSRISTINTVSISKEETVISFLKNCSCMKDLKQIHARVIQLGFEQNRFVVGKVIVFCAAAEHGDMNYAVSVFEKIGDPDAFIFNTMIRGFGKANDPRKAFDYYKRMQERGLVSDSFTFSFLLKVCGQLGLVLLGRLMHCSTLKRGLNSHVFVRNTLVHMYGTFKDIEASRQLFEEIPNPELVAWNIIIDCHVSCGKFNEALEMFSRMLKFGIEPDEATFVVILSACSALGALDFGRWVHSCISNIGHGCITEVNNSLLDMYAKCGALQEAFEIFNGMNKKNTVTWNTMILGLASHGYANEALALFSNMLEQKLWAPDDITFLVVLSACSHGGMVDKGWRFFDIMKKEYHIQPTIKHYGCMVDILGRAGFVEEAYRLISNMPMQCNAIVWRTLLAACRLHGNVELGKQVRKQLLELEPDHSSDYVLLANTYASAGRWNEAMRVRKTMHKRGVQKPEPGSRWVAWCYAHLLTGRNFWCLSAPWRSLLNSESDLGSNSGLSGNGENVYPVRVSYQLRFKVRIGAGQQLDLIN

>Pt16G05130

MAIQGPEIKFDGYNMLLNECVNKRAVREGQRVHAHMIKTCYLPPVYLSTRLIILYTKCECLGCARHVFDEMRERNVVSWTAMISGYSQRGFASEALHLFVQMLRSDTEPNEFTFATVLSSCTGFSGFELGRQIHSHIFKRNYENHIFVGSSLLDMYAKAGRIHEARGVFECLPERDVVSCTAIISGYAQLGLDEEALELFCRLQREGMSSNYVTYASLLTALSGLAALDHGKQVHSHVLRCELPFYVVLQNSLIDMYSKCGNLNYARKIFNNMPVRTVISWNAMLVGYSKHGKGIEVVKLFKLMREENKVKPDSVTFLAVLSGCSHGGLEDKGLEMFDEMMNGGDEIEAGIEHYGCVIDLLGRAGRVEEAFELIKKMPFEPTAAIWGSLLGACRVHSNTNIGEFVGCRLLEIEPENAGNYVILSNLYASAGRWEDVRNVRELMMEKAVIKEPGRSWIELDQTIHTFYASDRSHPRREEVFLKVRELLVKFKESGYVPDQSCVLYDVDEEQKEKILLGHSEKLALAFGLISTSEGVPLRVIKNLRICVDCHNFAKFVSKVYGRQVSIRDKNRFHHVAGGICSCGDYW

>Pt16G06550

MSGFSNLVLRKLELKNPKLSLLESCTTLSHLKIIHAHLIRAHTIFDVFAASCLISISINKNLLDYAAQVFYQIQNPNLFIYNSFIRGFSGSKDPDKSFHFYVQSKRNGLVPDNLTYPFLVKACTQKGSLDMGIQAHGQIIRHGFDSDVYVQNSLVTMYSTLGDIKSASYVFRRISCLDVVSWTSMVAGYIKSGDVTSARKLFDKMPEKNLVTWSVMISGYAKNSFFDKAIELYFLLQSEGVHANETVMVSVIASCAHLGALELGERAHDYILRNKMTVNLILGTALVDMYARCGSIDKAIWVFDQLPGRDALSWTTLIAGFAMHGYAEKALEYFSRMEKAGLTPREITFTAVLSACSHGGLVERGLELFESMKRDYRIEPRLEHYGCMVDLLGRAGKLAEAEKFVNEMPMKPNAPIWGALLGACRIHKNSEIAERAGKTLIELKPEHSGYYVLLSNIYARTNKWENVENIRQMMKERGVVKPPGYTLFEMDGKVHKFTIGDKTHPEIQQIERMWEEILGKIRLAGYSGNNDDALFDIDEEEKESNIHRHSEKLAIAYAIMRTKGHDPIRIVKNLRVCEDCHTATKLISKVYERELIVRDRNRFHHFKGGACSCMDYW

>Pt16G09640

SNYHSPQTNTDCLIEVDDLFNLLRLSVKYTDIDLARALHASILKLGEDTHLGNAVIAAYIKLGLVVDAYEVFMGMSTPDVVSYSALISSFSKLNRETEAIQLFFRMRISGIEPNEYSFVAILTACIRSLELEMGLQVHALAIKLGYSQLVFVANALIGLYGKCGCLDHAIHLFDEMPQRDIASWNTMISSLVKGLSYEKALELFRVLNQNKGFKADQFTLSTLLTACARCHARIQGREIHAYAIRIGLENNLSVSNAIIGFYTRCGSLNHVAALFERMPVRDIITWTEMITAYMEFGLVDLAVDMFNKMPEKNSVSYNALLTGFCKNNEGLKALNLFVRMVQEGAELTDFTLTGVINACGLLLKLEISRQIHGFIIKFGFRSNACIEAALIDMCSKCGRMDDADRMFQSLSTDGGNSIIQTSMICGYARNGLPEEAICLFYRCQSEGTMVLDEVAFTSILGVCGTLGFHEVGKQIHCQALKTGFHAELGVGNSIISMYSKCYNIDDAIKAFNTMPGHDVVSWNGLIAGQLLHRQGDEALAIWSSMEKAGIKPDAITFVLIVSAYKFTSSNLLDECRSLFLSMKMIHDLEPTSEHYASLVGVLGYWGLLEEAEELINKMPFDPEVSVWRALLDGCRLHANTSIGKRVAKHIIGMEPRDPSTYVLVSNLYAASGRWHCSEMVRENMRDRGLRKHPCRSWVIIKKQLHTFYARDKSHPQSNDIYSGLDILILKCLKAGYEPDMSFVLQEVEEQQKKDFLFYHSAKLAATYGLLKTRPGEPIRVVKNILLCRDCHTFLKYATVVTQREIIFRDASGFHCFSNGQCSCKGYW

>Pt16G10180

MLDQDELFTIGCSNNEAHDRCIAEQAMLCDDRGVFLEMAQVIRSLDLTILKGVIESRSNDTWAHFIVELLHRRRNSISSKDLMHPHPVKYEGRLCFSSDELCISELD

>Pt16G12080

MDYYSTTNPSSSGSSSAPKNGKEKKKGGRKSSGGVKLSTDPQSVAARERRHRISDRFKILQSLVPGGTKMDTVSMLEEAINYVKFLKTQVLLHQTIMNFVDDERSLDHYLPADYSTALPTEQPSYLDGNFAPVVHPSISVLPCPDSYLLGENYMQYDAFDIKN

>Pt16G12890

MSSLIVTKSAGIKNRLIQGFSCLKHLKHIHAALLRLGLDEDTYLLNKVLRFSFNFGNTNYSFRILDQTKEPNIFLFNTMIRGLVLNDCFQESIGIYHSMRKEGLSPDSFTFPFVLKACARVLDSELGVKMHSLVVKAGCEADAFVKISLINLYTKCGFIDNAFKVFDDIPDKNFASWTATISGYVGVGKCREAIDMFRRLLEMGLRPDSFSLVEVLSACKRTGDLRSGEWIDEYITENGMARNVFVATALVDFYGKCGNMERARSVFDGMLEKNIVSWSSMIQGYASNGLPKEALDLFFKMLNEGLKPDCYAMVGVLCSCARLGALELGDWASNLINGNEFLDNSVLGTALIDMYAKCGRMDRAWEVFRGMRKKDRVVWNAAISGLAMSGHVKDALGLFGQMEKSGIKPDRNTFVGLLCACTHAGLVEEGRRYFNSMECVFTLTPEIEHYGCMVDLLGRAGCLDEAHQLIKSMPMEANAIVWGALLGGCRLHRDTQLVEVVLKKLIALEPWHSGNYVLLSNIYAASHKWEEAAKIRSIMSERGVKKIPGYSWIEVDGVVHQFLVGDTSHPLSEKIYAKHGELAKDLKAAGYVPTTDHVLFDIEEEEKEHFIGCHSEKLAVAFGLISTAPNDKILVVKNLRVCGDCHEAIKHISRIAGREIIVRDNNRFHCFTDGLCSCKDYW

>Pt17G08610

MIRKRTNDRSTNRKQPSSLWKNCNNLLALKQIHATLIIKGFNSNRAALRELIFAGAMTISGAINYAHQVFAQITEPDIFMWNTMMRGSSQSKNPSKVVLLYTQMENRGVKPDKFTFSFLLKGCTRLEWRKTGFCVHGKVLKYGFEVNSFVRNTLIYFHSNCGDLVIARSIFYDLPERSVVSWSALTAGYARRGELGVARQIFDEMPVKDLVSWNVMITGYVKNGEMENARTLFDEAPEKDVVTWNTMIAGYVLRGEQRQALEMFEEMRNVGECPDEVTMLSLLSACADLGDLQVGRKLHCSISEMTRGDLSVLLGNALVDMYAKCGSIEIALQVFKKMREKDVTTWNSVIGGLAFHGHAEESIKLFAEMQALKNIKPNEITFVGVIVACSHAGNVEEGRRYFKLMRERYDIEPNMIHHGCMVDLLGRAGLLSEAFELIAKMEIEPNAIIWRTLLGACRVHGNVELGRLANERLLKLRRDESGDYVLLSNIYASAGEWDGAEEVRKLMDDGGVRKEAGRSLIEADDRAVMQFLFDPKPKLNSRGQVS

>Pt17G08700

MAEACNSMSRLKQIHAHSLLTGLHDHSIILAKMLRFAAVSPSGDLAYAQRLFDQLPHPNTFFYNTLIRGYAKSSIPSYSLHLVNQMRQNGVDPDEFTFNFLIKARSRVRVNINRNLPLVVECDEIHGAVLKLGFSSHLFVRNALIHLYAARGNPVVAWRVFDETVGVDVVSWSGLVLAHVRAGELERARWVFDQMPERDVVSWTTMVSAYSQAKYSREALELYVTMLDKGVRPDEVTLVSVISACTNLGDLQMGYSVHSYIDENGFRWMVSLCNALIDMYAKCGCMDRAWQVFNSMSRKSLVTWNSMISACANNRNPEDAFGLFSRMFNYGVAPDGVTFLAVLTAYAHVGLVDEGYRLFESMQRDHGIEARIEHYGCVVNMLGQAGWLEEAFELITSMPLPSNDVVWGVLLAACRKHGDVYMGERVVKKLLELKLDGGYYTS

>Pt17G12680

MALAKDRMDSVQTCALYGNVMGDLSSLGPNYGFDEEGDRNFEKNSALMIKNLAMSPSPPSLGSPSSANSGELVFQATDNQVEEAHSLINFKGTGFDSIMHANGSLISFEQSNRVSQTSSHKDDYSAWEGNLSCNYQWNQINPKCNANPRLMEDLNCYQSASNFNSITNSAEKENHGDWLYTHESTIVTDSIPDSATPDASSFHKRPNMGESMQALKKQRDSATKKPKPKSAGPAKDPQSIAAKNRRERISERLKMLQDLVPNGSKVDLVTMLEKAISYVKFLQLQVKVLATDEFWPVQGGKAPDISQVKGAIDATLSSQTKDRNSNSSSK

>Pt18G03590

MATRIGALKIRELENFFVPILQNCKNIVELKSIHAHVIKYSLSQSSFLVTKMVDVCDKTEDLGYASLLFKQVKEPNGYLYNAMIRAHTHNKVYALAILFYKEMLRLKDPESENPIFPDRFTFPFVIKSCSGLVCYNLGKQVHAHLCKFGPKSNITMENALIDMYTKCASLLDAHKVFDGMVERDAISWNSIISGHVGVGQMRKAGALFDLMPYRTIVSWTAMISGYTRLGSYADALYVFRQMQIVGVEPDEISIISVLPACAQLGALEVGKWIHMYCDRNGLLRKTSICNALMEMYSKCGCIGQAYQLFDQMSKGDVISWSTMIGGLANHGKAREAIELFKRMKKAKIEPNGITFLGLLSACAHAGFWNEGLAYFDSMSKDYHIEPEVEHYGCLVDILGRAGRLSQALDVIEKMPMKPDSKIWGSLLSSCRTHSNLDIAIIAMEHLEELEPDDTGNYVLLSNIYADLAKWDGVSRMRKLIKSKSMKKTPGSSLIDINNVVQEFVSWDDSKPFSRDIFWLLELLTSHQDTTDPHLIEIMLEDGSECLG

>Pt18G04010

MKMVVLASPVSTLQVLSFSDPSPPYKLVHDHPSLTLLSNCKTLQTLKQIHSQIIKTGLHNTHFALSKLIEFCAVSPHGDLSYALSLFKTIRNPNHVIWNHMIRGLSSSESPFLALEYYVHMISSGTEPNEYTFPSIFKSCTKIRGAHEGKQVHAHVLKLGLEHNAFVHTSLINMYAQNGELVNARLVFDKSSMRDAVSFTALITGYASKGFLDEARELFDEIPVRDVVSWNAMISGYAQSGRVEEAMAFFEEMRRAKVTPNVSTMLSVLSACAQSGSSLQLGNWVRSWIEDRGLGSNIRLVNGLIDMYVKCGDLEEASNLFEKIQDKNVVSWNVMIGGYTHMSCYKEALGLFRRMMQSNIDPNDVTFLSILPACANLGALDLGKWVHAYVDKNMKSMKNTVALWTSLIDMYAKCGDLAVAKRIFDCMNTKSLATWNAMISGFAMHGHTDTALGLFSRMTSEGFVPDDITFVGVLTACKHAGLLSLGRRYFSSMIQDYKVSPKLPHYGCMIDLFGRAGLFDEAETLVKNMEMKPDGAIWCSLLGACRIHRRIELAESVAKHLFELEPENPSAYVLLSNIYAGAGRWEDVAKIRTRLNDNRMKKVPGCSSIEVDSVVHEFLVGDKVHPQSNEIYKMLDEIDMRLEKAGFVPDTSEVLYDMDEEWKEGVLSHHSEKLAIAFGLISTKPGTTIRIMKNLRVCGNCHSATKLISKIFNREIIARDRNRFHHFKDGSCSCKDYW

>Pt18G06750

MECDFITLKMRSPSLNYRFCPRHTSSMKLPNKPRSSSTSLLLSLGSNFSVSAAALSTIEPSPTRDFNPTNLQNSPAHLPEANTKISRKFWSPNLPNLNGKSESNVTSSPVRFDAKERLKRYSVMLRECASKGDVKEGKAIHGNLITSGVELDSHLWVSLINFYAKCRSRFFARKVLAEMPQRDVVSWTALISGFVNEGCGSESVSLYCEMRKENVRANEFALATALKACSMCLNLEFGKQVHVEAIKAGLLLDLFVGSALVDLYARCGEMELAERLFFGMPEKNGVSWNALLNGYAQLGDGKKVLKLFCKMKECETKFSKFTLSTVLKGCANTGSLREGKVLHALALRSGCEIDEFLGCSLVDMYSKCGTVYDALKVFTKIRNPDVVAWSAMITGLDQQGHGQEAAELFHLMRRKGARPNQFTLSSLVSTATNMGDLRYGQSIHGCICKYGFESDNLVSNPLIMMYMKSRCVEDGNKVFEAMTNPDLVSWNALLSGFYDSQTCGRGPRIFYQMLLEGFKPNMFTFISVLRSCSSLLDPEFGKQVHAHIIKNSSDDDDFVGTALVDMYAKARCLEDAGVAFDRLVNRDIFSWTVIISGYAQTDQAEKAVKYFRQMQREGIKPNEYTLASCLSGCSHMATLENGRQLHAVAVKAGHFGDIFVGSALVDLYGKCGCMEHAEAIFKGLISRDIVSWNTIISGYSQHGQGEKALEAFRMMLSEGIMPDEATFIGVLSACSFMGLVEEGKKRFDSMSKIYGINPSIEHYACMVDILGRAGKFNEVKIFIEEMNLTPYSLIWETVLGACKLHGNVDFGEKAAKKLFEMEPMMDSSYILLSNIFASKGRWDDVRNIRALMTSRGIKKEPGCSWVEVDGQVHVFLSQDGSHPKIREIYAKLDKLGQSLMSIGYVPKTEVVLHNVSNKEKMEHLYYHSERLALSFALLSTNAVKPIRIFKNLRICEDCHDFMKLISDITNQEIVVRDIRRFHHFKRGTCSCQDRW

>Pt18G10950

MQPCSREMQGINSLLNPSSQIPLQDLQNQQNPSQIQNSHFDPNSSSNDDFLEQMLSNIPPCSWPDLKSPWDLTMPINNNDSSNSIAKPRDLSDETAPSNTDNSNLGFHNNFDESVILASKLRQHQISGGGGAAAAAKMMLQQQLLMAAARGVLPQNDVIDGSSFKGGDGSMQGLFNGFGAGSMNGTGQASNQSMQHFNHPQGGAMQAQNFGAQGAATTAVMNQPQASGSNGGAPAQPRQRVRARRGQATDPHSIAERLRRERIAERMKALQELVPNANKTDKASMLDEIIDYVKFLQLQVKVLSMSRLGGAAAVAPLVADMSSEAGGDCIQASADGGSLSRTSNGNQTARTNDSSLTVTEHQVAKLMEEDMGSAMQYLQGKGLCLMPISLATAISTATCHNRSPAINNNHHALLQSNGEGPASPSMSVLTVQSATMGNVGGDGGAVKDAASVSKP

>SI011G00300

MPDHLLIEMMCEDYEVFLEMAHVLKGLEVSILKGVLEHRSDKLWARVVVEASGGFSQTQILCPLMHLLHRRFS

>Sl01G081020

MSGTSFFSSPPLYTTSTTTNYSHSPSPDRLKFLIDKSKNIRQLLQIHAFLIRNGLESDPVLNFRLQQSYSSLGHLQHSVKVFKRTHSPTVFSYTAIIHNHVINDLYEQAFVLYIQMLTHNIEPNAFTFSSMLKTCPLESGKALHCQALKLGYESDTYVRTALVDVYARGSDIVSACKLFDTMTERSLVSLTTMITGYAKNGHIQEAGVLFEGMEDRDVVCWNAMIDGYSQHGRPNEALVLFRKMLLSKVKPNEVTVVAALSACAQMGVLESGRWIHAYVKSNRIQINKHVGTAFIDMYSKSGSLEDARMVFDQMRDKDVITWNSMIVGYAMHGFSLEALQLFNEMCKLGLQPTDITFIGILSACANAGLLSEGWTYFQLMEKYLIEPKIEHYGCMVNLLGRAGQLEKAYEFVKSMKIDSDPILWGTLLTACRIHGDVRLAEKIMEFLVEQDLATSGTYVLLSNIYAASGDWDGVAKVRALMKRSGVDKEPGCSSIEVNNKVHEFLAGDMKHPKSKEIYIMLEEVNKLLEAHGYLPQTDIVLHNLGEVEKQQALAVHSERLAIAYGLISTQAGTTIKIVKNLRVCPDCHAVTKLISKITGRKIIVRDRNRFHHFVDGSCSCGDFW

>Sl01G096050

MVTGNMLWSGEDKAMVASVLGKEAFEYLMSGSVSAECSLMAIGNDQNLQNKLSDLVERPNAANFSWNYAIFWQISRSKSGELVLGWGDGCCREPKEAEEREVKKILNLRLDDEGQQRMRKRVLQKLHMLFGGTDEDNYAFGLDRVTDTEMFFLASMYFSFPRGEGGPGKCFGSGKYLWLSDALTSNLDYCARSFLAKSAGMQTIALIPTDVGVVELGSVRSIPESLELLQNIKSCFSSFLSLVRDKQAAGIAAVPEKNEGNNPRLSNSGAVTERTDGNPKIFGHDLNSGTHFREKLAVRKAEERPWDMYQNGNRMPFVNARNGLNPASWAQFSNVKLGKPVELYAPPTPGHNLMNGGREEFRLNNFQHQKPAARMQIDFTGATSRTIVSPAHNVESEHSDVEASCKEDRAGPVDEKRPRKRGRKPANGREEPLNHVEAERQRREKLNQRFYALRAVVPNISKMDKASLLGDAIAYITELQKKLRDMESERELRLGSTSRDAITSEDSPSSEIQIRGPDINIEAANDEVIVRVSCSLETHPLSRIIQIFKEAQINVVESKLSAGNGTVYHTFVIKSSGSEQLTKEKLLAAFSSESNSLRQLSPVGQ

>Sl01G096200

MPDVIGGPFADVLSATLESILEIVLTSKNVFIEKKSFAELSDYLNRIVPFLKEINRKNITDSTPWQNVIQILNQQTVDARQLILECSKKNKVYLLMNCRHIAKRIENITREISRALSCIPLASLDISSGIKEDIVQVMDSMRTAEFKTAIAEEEILNKIDSGIHQRNVDRSYANKLLVSIAEAIGVSTESSALRREFEEFKDEIDNARLRKDQAEALQMDQIIALLERADAATSRQEKEKKYFIKRKSLGNQPLEPLLSFYCPITGEVMTDPVETPSGHTFERCAIEKWLAEGNLCPMTSTPLKNTMMRPNKTLRQSIEEWKDRNTMITIANMKLKLSSNEGDEVLNCLEQVKDICEQREIHREWVIMEDYIPILIKLLDSKSRDIRNLVLEVLCVLAKDGDDAKERIVEVDNALESIVHSLGRRIGERKSAVALLLELSKCKSVQESIGKVQGCILLLVTMSSCDDNKAAKDARDVLENISFSDDNVILMAQANYFKYLLQRLSSGSSDVKLLMAKTLGEMELTDHNKSSLFEEGVLDSLLSSLSHSEVEVKQAGVKALLNLSSLPRNGQDMIRKGVMRPLLDMLYRHTASQSLRELVAATITNLAFSASSEALSLLDADEDVYELFSLVNLNGPAVQQSILQAFCAMCKSPSGANVKIKLAQCSAVQVLMQFCEHSNSNVRSDAIKLLCCLIENGNGGVIQEYVDQNFIEILLKIIKTSQDEEEIASAMGITSNLPKSSQISDWLFAAEGLPVFSKFLDEVKHKSSCKLQLVENAVGTLCHFTVSINQQTQRIAGLVPKLIRLLDQGTSLTKNRAAICLAQLSENSQTLSRTIPKRSGLWCFSPSQVELCPIHRGICTLETSFCLVEAGAVGPLVRVLGDTDPGACEASLDALLTLIKDEKLQSGAKVLAEENAIPSMIKLLNSPSPRLQEKVLNSLERLFRLVEYKQRYGSSAHMPLVDLTQRGTSNIKSVAAKGCLEPSRLKYQHFKVNLCFFIILNCSKRHLSDRFLHQTLLMNQLKQIHANTLRNGIDFTQFLISKIIEIPNIPYAHKVFDNITKPTVFLYNKLIQAYSSHGFPSQCFSLYIKMRRQGCSPNPHSFTFLFAACSNRSTPIQGQMFHVHFIKWGFEFDIYTLTALVDMYAKMSLLPSARKLFDEMEMKDVPIWNSLIAGYAKNGNVVELVIDRAMQLFHEIGRRRNLCSWNTMIMGLAVHGKGDEALKLFNQMLGEGNTPDDVTFVGAILACTHGGMVAKGWELLKLMEQRFSIAPKLEHYGCMVDLLGRAGKLQEAYDLIQSMPMRPDCVIWGTILGACSFYGNVELAEKAAEFLSVLEPWNPGNYVILSNIYARAGRWDGVARLRKLMKSSQITKAAGYSFIEEGGDIHKFIVEDKSHPKSNEIYSLLDLVTTRLKFDVSTMEIDLDSIAE

>Sl01G097130

MAISMRNLEKVAIFKPSEHTIPLLSFKGYQLLNSAATQLSSVDDDQQEASLRGRFRRREQAFCIFGSSLPLSRLFHSFAHCSIHFSHCQVSSLLPHPTLLCFFQEKLSLSNPRRIWPLLPKGCFYVTNSHPDVKFLEFNTKKFSTVQTSLCNSQGKLLFSNSRKISPFSSKGYFYFTGINSNLRLSCRGICSCVDNDGELESDNDVDTESDERVVESKADPKEVERVCKVIDELFSLDRNMEAVLDECGINLSHDLVVDVLERFKHARKPAFRFFCWAAMRPGYAHDSRTYNVMMAILGKTRQFETMVSVLEEMGEKGLLTMEAFLISMKAFAAAKERKKAIGMFELMKKYKFKVGVETINCLLDALGRAKLGKEAQLLFEKLEHRFTPNLQTYTVLLNGWCRVKNLMDAGKVWNEMIDKGFKPDIVAHNTMLEGLLKCKKRSDAITLFEVMKAKGPSPNTRSYTILIRDLCKQGKMDEAVAGFEEMLSSGCEADAATFTCLVTGFGNKKRMDKVFALLTEMKEKGCPPDARLYNALIKLLINRRMPVDAVTLYKKMIRNGIQPTIHTYNMLMKSFFMTKNYDMAHATWEEMSLRGCCPDENSYTVFIGGLIGQGRSMEACKYLEEMIDKGMKAPQLDYNKFAADFSRGGKPDILEELAKRMKFSGKFEVSSLFARWAEMMKSRVKRRDPS

>Sl01G097670

MITLSLPSPAKFIPPSSKSRRIRNPDFEALKDTLIRQANGGNLKQAISTLDQISQMGFNPDLTSYTVLLKSCIRTRNFQIGQLLHSKLNDSPIQPDTIVLNSLISLYSKMGSWETAEKIFESMGEKRDLVSWSAMISCYAHCGMELESVFTFYDMVEFGEYPNQFCFSAVIQACCSAELGWVGLAIFGFAIKTGYFESDVCVGCALIDLFAKGFSDLRSAKKVFDRMPERNLVTWTLMITRFSQLGASKDAVRLFLEMVSEGFVPDRFTFSGVLSACAEPGLSALGRQLHGGVIKSRLSADVCVGCSLVDMYAKSTMDGSMDDSRKVFDRMADHNVMSWTAIITGYVQRGHYDMEAIKLYCRMIDGLVKPNHFTFSSLLKACGNLSNPAIGEQIYNHAVKLGLASVNCVANSLISMYAKSGRMEEARKAFELLFEKNLASYNIIVDGCSKSLDSAEAFELFSHIDSEVGVDAFTFASLLSGAASVGAVGKGEQIHSRVLKAGIQSSQSVCNALISMYSRCGNIEAAFQVFEGMEDRNVISWTSIITGFAKHGFAHRAVELFNQMLEDGIKPNEVTYIAVLSACSHVGLVDEGWKYFDSMSIDHGITPRMEHYACMVDLLGRSGSLEKAVQFIKSLPLNVDALVWRTLLGACQVHGNLQLGKYASEMILEQEPNDPAAHVLLSNLYASRGQWEEVAKIRKDMKEKRMVKEAGCSWMEAENSVHKFYVGDTKHPKAKEIYEKLNKVALKIKEIGYVPNTDLVLHEVEDEQKEQYLFQHSEKIALAFGLISTSKQKPIRIFKNLRVCGDCHNAMKFISVAEGREIIIRDSNRFHHIKDGLCSCNDYW

>Sl01G098150

MAAVFYSPNNAIPLSLLPPRPFNLHSLFTCTSSLTTTKKTKSYIKLGCLNLAERVFDSLRSPDVVSYTAIISAFAKSNREREAFELFLEMKDLGIEPNEFTYVAILTACIRSLNLELGCQVHGLVIRLGYSSYTYVVNALMGLYSKCGLLEFVVLLFNAMPQRDIVSWNTVIACMVEHSMYDRAFEMYSELCRNKCLIADHFTLSTLLAASSRCLAVREGQELHRHALKRGFHGNLSVNNALIGFYTKCGTLKNVVDVFERMPVKDVFSWTEMIVAYMEFGHVDLAMEIFNSMPERNSVSYNALLAGFSQNHEGFKALALFCRMLEGGMELTDFTLTSVVNACGSVMERKISEQIHAFILKCGLKSNDRIETSLIDMCTRCGRMDDAEKLFDDLPLDHDNSIALTSMICAYARNGQPEEAISLFLVRHSEKSLVVDEVALATILGVCGTLGILKLGEQIHCYAWKHGLMSDAGVGNAMISMYSKCGETQSAVKTFEAMPTHDLVSWNGLLTCYVLHRQGDGALDTWAKMERLGVDPDSITCVLVISAYRHTSTNLVDCCQKFFSSMQSSYNVNPTSEHYAGFVGVLGYWGLLEEAEKIINAMPFEPKASVWHALLDGCRLHVNAIIGKRAMKNILSIVPQDPSTFILKSNLYSASGRWQCSELVRAEMREKGIQKIPGRSWIIFGDKVHSFFARDKLHSQSKDIYSGLQILILECLKAGYVPDTSLVLHEVEEHQKKDFLFYHSAKLSVTFGLLMTRPGKPVRVMKNVLLCGDCHTFFKYVSVITKRDIHVRDASGFHHFVNGKCSCGDNWC

>Sl01G100340

MNRGSREVERRILRLLHGQKTRTQLTQIHAHILRHHLHHSNQLISHFISICGSLNRMRYASIIFQHFQNPSIFLFNSMIKGYSLCGPYQNSVVFFSTMKRRGIWPDEFTFAPLLKACANLVDLELGQGVHKHVLALGFGRFGSIRIGIVELYSGCGRMTDAKKVFDEMPHRDVIIWNLMVKGYSQSGNVDMGLGLFRQMGERTVVSWNLMISLLAQNGREKEALALFHEMKNGGFEPDEATVVTVLPVCAQLGELDLGRWIHSYAKSEGLYPNLVSVGNALVDFFGKSGDLETAITIFNDMPRKNVVSWNAAISNCAFNGKGELGVQLFDKMLDEGVRPNDSTYVGALACCAHAGLVHRGGDLFDSMIANHGIEPTIEHYGCMVDLLGRGGCLKEAHKLVETMTMEPNAAIWGALVSACRTHGEMELAEYALKELIKLEPWNSGNYVLLSNIYADRGKWEDVEKVRVLMSGNSIKKAPGQSIIV

>Sl01G100790

MISTGFIRDTYAASRILKFSTDSLFIHVNYSHKIFDYIDNPNGFICNTMMRAYLQRNQPQNTIFLYKSMLKNNVCIDNYTFPLLVQASTVRLSEAEGKEFHNHVIKTGFGLDVYVKNTLINMYAVCRNLVDARKMFDESPVLDSVSWNSILAGYVQVGNVDEAKVIFDKMPMKNVIASNSMIVLLGRSGRMSEACQLFNEMMQKDVVSWTALISCYEQHGMHTQALDLFMQMCSNGISIDEVVVLSVLSACAHLLVVQTGESVHGLVIRVGFESYVNLQNALIHMYSTCGDVMAAQRLFDTSSHLDQISWNSMISGYLKCGSVEKARELFDSMAEKDVVSWTTMISGYAQHDHFSETLALFQEMLHEDSKPDETTLVSVLSACTHLSALDQGKWIHAYIRKNGLKVNSILGTTLVDMYMKCGCVENALEVFNAMEEKGVSSWNALILGLAMNGQVERSLDMFQKMKECGVTPNEVTFVAVLGACRHMGLVDEGRSYFNAMTTHYNVEPNIKHYGCMVDLLARTGLLKEAETLIDSMPIAPDVATWGALLGACRKHGNSEMGERVGRKLLELQPDHDGFHVLLSNLYASKGNWDSVLDIRVAMTRKGVVKVPGCSMIEANGAVHEFLAGDKSHSQINEIEEMLAEMEKRLKIMGYAPGTDEVLLDIDEEEKESTLFRHSEKLAIAYGLIAIAPPTVIRIIKNLRICSDCHAAAKLISKAFDREIVVRDRHRFHHFKDGSCSCMEFW

>Sl01G102790

MSHLHTAALKATKLIHLKQFHAQLFQRSLCSDNYWVAQLIKLCTRLHAPSTYVSRVFDSVHQPNVFVFTNILKFYSQLGAYSDVLYLFDKMQKSNVAPDAFVYPILIKASGKWGIVFHAHCIKMGHDWDRFVRNAIMDVYGKFGPLEIARELFDEIPERAVADWNAMISGCWNWGDEVEARSLFDLMPEKNVVTWTAMVTGYSRRKDLENARKYFDQMPERSVVSWNAMLSGYAQNGCAEEVIKLFNEMMSCEVCPDETTWVTVISLCSSHGDVSLAEGLVKMINEKGVRLNCFAKTALLDMYAKCGNLAMARKIFDELGTYKNLVTWNAMISAYARVGDLASARGLFDKVPEKNVISWNSIIAGYAQNGESKVAIDLFKDMIAKDVLPDEVTMVSVISACGHLGALEFGNWAVNFLEKHQIKLSISGYNALIFMYSKCGNMKDAEKVFQSMEARDVISYNTLITGVAAYGNAIEAVELLWKMKKENIEPDRITYIGVLTACSHGGLLKEGQRIFDSIKDPDSDHYACMVDLLGRNGKLDEAKCLIGSMAMHPHAGVYGSLLHASRVHKRIDLGEFAASKLFEIEPENSGNYVLLSNIYASARRWEDVDRVRGLMTIGGVKKTTGWSWIEHKGEMHKFIVGDRSHERTADIHRVLFETEKKMKLAGYMADKSCVLKDVEEEEMEEMVGTHSEKMAVAFALLVTEPHSVIRVVKNLRICRDCHTAIKIISKMEGREIIVRDNNRFHCFSEGQCSCKDYW

>Sl01G107140

MEPVVAMSEGEWSSLSGTCSTEEANFMAQLFGACPNEQQLPSSGLPNFWTNHESNIGGSSEVSIFSSQHHTNSSIYHFPTSTNHFQPMLLTTSMTMEHLPPTNNLIEADAVEFLNKQVNNDSIESGENIMSESVLHGKSLQLGREYDQMHQPESSKKRSQSPVDHKNKRSVKPKKNMKSSVADDEETGNNNNNNTVLHRQSSFSCCSEDESNVSSYDIYGLASSDNSKGVSLPNGKSRANRGSATDPQSLYARKRRERINERLRILQSLVPNGTKVDISTMLEEAVQYVKFLQLQIKLLSSDDLWMYSPIAYNGMDIGLDLKIGIPNPKP

>Sl01G110010

MNSISCQQLEHKLSPFGRKLISSVAFAGSTSSFPEQPPIQFLTKSRNIIHSKPEISHAHLIKTQNLEGNTHAANSVLHNYGEYSRMDNAAKVLEEMPKQNSVSWNLMISNSNKALLYQDSWRLFCRMHMLGFDMNMYTYGSILSACGALTSTLWGEQVYGLVMKNGFFSDGYVRCGMIELFSRSCRFSDALRVFYDYLCDNVVCWNAIISGAVKNREYWVALDIFRLMWGEFLKPNEFTIPSVLNACVSLLELQFGKMVHGAAIKCGLESDVFVGTSIVDLYAKCGFMDEAFRELMQMPVSNVVSWTAMLNGFVQNDDPISAVQIFGEMRNKGIEINNYTVTCVLAACANPTMAKEAIQIHSWIYKTGYYQDSVVQTSFINMYSKIGDVALSELVFAEAENLEHLSLWSNMISVLAQNSDSDKSIHLFRRIFQEDLKPDKFCCSSILGVVDCLDLGRQIHSYILKLGLISNLNVSSSLFTMYSKCGSIEESYIIFELIEDKDNVSWASMIAGFVEHGFSDRAVELFREMPVEEIVPDEMTLTAVLNACSSLQTLKSGKEIHGFILRRGVGELHIVNGAIVNMYTKCGDLVSARSFFDMIPLKDKFSCSSMITGYAQRGHVEDTLQLFKQMLITDLDSSSFTISSVLGVIALSNRSRIGIQVHAHCIKMGSQSEASTGSSVVTMYSKCGSIDDCCKAFKEILTPDLVSWTAMIVSYAQNGKGGDALQVYESMRNSGIQPDSVTFVGVLSACSHAGLVEEGYFFLNSMMKDYGIEPGYRHYACMVDLLSRSGRLTEAERFICDMPIKPDALIWGTLLAACKLHDEVELGKLVAKKIIELEPSEVGAYVSLSNIWASLGQWDEVLKIRGSLRGTGISKEPGWSSL

>Sl02G065560

MSNGSISIALESSSSLFTKILHYTRCKNLPKGQSLHSHLIKTGSSSSCIYIANSIVNLYAKCHRLSDAHLAFQEIQTKDVVSWNSLINGYSQLGRRDSSLSALNLFKLMRQENTLPNPHTFAGIFTSLSTLGDSFTGKQAHCLAFKLGYLSDVFVGSSLLNVYCKAGHHLGDARNMFDEMPERNSVSCTTMISGYALQRMVKEAVGVFSVMLLKRGEDVNEFVFTSVLSAIALPEFVYVGKQIHCLSLKNGFLSAVSVANATVTMYAKCGRLDDACRAFELSSEKNSITWSALITGYAQNGDCEKALKLFSEMHYRGMIPSEYTLVGVLNACSDFDALREGKQVHGYLVKLGFEPQMYILTALVDMYAKCGNISDARRGFEYLKEPDIVLWTSMIAGYVKNGDNESAKGMYCRMLMEGVMPNELTMASVLKACSSLAALEQGKQIHAHIVKHGFSLEVPIGSALSTMYAKSGSLHDGNLVFRRMPARDLVSWNSMMSGLSQNGCGTEALELFEEMLHEGTRPDYVTFVNILSACSHMGLVKRGWSIFRMMSDEFGIEPRLEHFACMVDMLGRAGELYKAKEFIESAASHVDHGLCLWRILLSACRNYRNYELGAYAGEKLMELGSQESSAYVLLSNIYSSLGRLEDVERVRRLMNLRGVSKEPGCSWIELKSQFHVFVVGDQLHPQIIHIRDELWKLTKLMKDEGYKPDFDPCLELEGVMD

>Sl02G072010

MAATSPTPSTLSSFLEMANSISELHQAHAVMLKTGLFRDPFAASRLLTKATVLPLSSTETLSYALSVFTHIEEPNSYIYNTIIRAYSTSPFPQLALIIFLKMLNSVNKVFPDKYTLTFIVKACATMENAKQGEQVHGLVTKIGLEEDVYVYNTLVHMYAKCGCFGVSRGMIDGLIEDDVIAWNALLSVYAERGLFELARELFDEMPVKNVESWNFMVSGYVNVGLVDEARKVFDEMLVKDVVSWNVMITGYTKADKFNEXXXXXX

>Sl02G084830

MVSLARKLIASRNPIPTSRSYEFGNPEVFVPFSRLQETKLLESHLVSLLDNCSSLNQIKQVHAHVIRRGLDQCCYVLAKLIRLLTKINVPMDPYPRLVFHQVEYRNPFLWTALIRGYSIQGPLKEAVSLYNAMRRESISPVSFTFTALLKGSSDELELNLGRQIHCQSIKLGGFCEDLFVHNILIDMYVKCGWLDYGRKVFDEMSERDVISWTSLIVAYSKAGDMAAAAEMFERLPVKDLVAWTAMVSGFAQNAKPREALEFFHRMQSEGVETDELTLVGVISACAQLGAAKYANWVRDMAEGYGFGPANHVMVGSALIDMYSKCGNVEEAYKVFEKMKEKNVFSYSSMIMGFAMHGCANAALDLFEEMVKTEVKPNKVTFIGVLMACTHAGLVERGRNLFDTMEKHYSVEPSVEHYACMIDLLGRAGQLEEARELIKAMPMEPNSGVWGALLGACRIHAGRWEDVLGVRKSIKQKLLRKDPSRSWIEGKEGVIHEFYAGDMTHPNSKEIKEALEDLIGRLKSHGYEPNLSSVPYDLNEEHKRRILLTHSEKLALAYGLLITDSAGSTIKIMKNLRICEDCHSFMGGASQITGREIIVRDNKRFHHFRNGVCSCGNFW

>Sl02G087610

MNCLSNALVVVRTNNSFQIWKWCMSMAEKCNNMRQLKAIHAIYITLGLQRNTYAVSKLLDFCALSNSGDLSYASRIFAQVQTPNAFLYNALIRAYSSSPQPQVSLNYFNLMVQTSNAAAPDSFTFPFLLIACANGPLEVEGKQIHSWIIKNSFSASNAHVQTALIRFYTNCKALDDARKVFDEITDIDVIQCNVLMSGHLQSGLAKEALSIFQDMLGRGVGPDEYCVTTALGACAQLGALEQGKWIHEHVTKSEWLEYDVFIGSALVDMYAKCGSINLASEVFESMPTRNKHSWATMIRGFAVHGRPELALSCLERMQVADGLKPDGVVILAVLAACAHSGLQKEGQGLLDEMESLYGVTPEHEHFSCVVDLLCRAGRLDDALKLIRRMPMKPRASVWGALLSGCRNHNNVNLAELAVKEILLVEDGNEAEEDSAYVQLSNIYLAARQCDDARRIRRRIGDRGLRKTPGYSAIEIDGMVNEFISGDVSHICLADIHKVLDLTYLDPHFDNLI

>Sl02G088170

MIKKTIASRSSNLHRQSSLWSKCKDLQSLKQIHALMIINGFNSNRIALRELIYASAVTFSASIHYAHKLFAQITQPDLFMWNTMLRGSAQSHRPSLAVSVYTHMEKRSIRPDSYTFPFLLKACTKLSWLVSGLVVHGKIVKHGFESNKFARNTLIYFHANVGDIRIAGQLFDGSAKRDVVAWSALTAGYARRGKLDAARRLFDDMPVKDLVSWNVMITGYVKQGKMDNARELFDIVPKRDVVTWNAMISGYVLCGENEKALKMYEEMRGAGEYPDEVTMLHLLSACTDSAFLDVGELIHRSIIEMGAGELSVFLGNALVDMYARCGSIRKALEVFQGXXXXDVSSWNIIILGLAFHGHSEECISLFEDMRRMKYIPNEITFVGVLVACSHAGKVDDGREYFSMMRTDYNIETNIRHYGCMVDMLARAGLLNEAFEFINTMDIKPNAIIWRTLLGACKVHSNVKLGRYANKQLLKLGREDSGDYVLLSNIYASRDEWDGVERVRKLMDDNGVWKEPGCTLIEADDYDLKNFYFDSKCKHS

>Sl02G088470

MQMPTPIRAPTWVSRRRYFEQKLWDLDKCRDINQLKQMQALVYKSNLEQDPFIAPKLIAAFSNCRQMGSVLKIFDQVRDPNVHLYNALIRAHIYNFQPSQAFDTFFDMQSSGIFPDNFTFSFLLKGCSGKCWLSVVSMIHTHVVKWGFEDDIYVPNSLIDAYSKCGLVGVRIAGQLFWGMKERDVVSWNSMISALLKVGDLSEARKLFDEMPQRDRVSWNTMLDGYTKAEQMSVAFELFKTMPQRDVVSWSTMVSGYCKAGDLEMARMLFDKMPSKNLVSWTIMISGYAEKGLINEAILLFMQMEETGLRLDVAAFVSILAACAESGMLSLGKKVHDSVERSMYKCNTLVCNALIDMYAKCGCLHKAYKVFNGLKKRDLVSWNAMIHGLAMHGRGKKALELFIRMKQEGFVPDKVTLVGILCACNHTGLVDEGILFFYSMEKDYGVKPEVEHYGCLIDLLGRGGYVREAFELARKMPLEPNIKIWGSLLGACRMHKDVELADDVRSLLVKLEPKNAGKLSALSNIYASAGDWDNVANIRLMMKNIGRPNQSGGSLLLLNDEYREFTVMDKSHVKSDKIYQMIDRLGQHLKLLSPVPAGLCDE

>Sl02G091440

MSMALAKEHVIMSDTKMGMVDNYDQYYEGEFGINDHSSPELYGIHEEPPKSIFEECENSEKTSPKIAKNFALSSSNSSLSSPSSSNSNAQSVINFKGVYGNFMHSANGSLLSFEQSERFCPNPRMISNINQVEGSVWEDNNLHYQNCVTPKGSSNTSPRVINDNSNNNGIPFGWLNSEANASTTTHIDESRFNKRPSTEESMQTNKKQCSAGSKKGKPNNNNNNSIGTKDPQSIAAKNRRERISERLKILQELVPNGSKVDLVTMLEKAIGYVKFLQLQVKVLATDEFWPTQGGKAPDISQVKEAIDAILATQRDRN

>Sl03G115790

MSVEESVPAKQRGSRALQQRLFSLLQSCKSIKQLTQIHAHVITNGFTQKNFILVRLLSPFLTSYSLKYADHIFSQVRSPSTTLWNQIIRGHARSENPQKSIELFNQMEISTAMPDGYTYSYVLNGCAKGGLFREGQQVHGKIVKNWSLLNVFVQTNLVNLYSTTGGENCIDNAQKMFDEMTEKNVVTWNSLLFGYFRNGDADEALRLFDEMPHKNVVSWTTVISGCTQNGRCQHALALFRLMQRRHVEFDQVTLVAVLSACAESGALDLGKWIHSTVVESSQLRNEPVLVSFYNALIHMYASCGEIEEAYRVFEEMPRRNSISWTSMITGFAKQGYAHEALTIFQQMESWGGNDVRPDEITFLGVLFACSHAGYVNEGYRYFRCMKETWDIEPRIEHYGCMVDMLSRAGLFDEATTLVETMPMKPNEAVWGALLGGCKIHKNVRLASSIAQKLAVELEADRAAGYFVLLSNLYATEKRWQDVVITRQKMYGMGLKKSPGQSKIEADGTTHNFLASDLSHKHTCSVYEMLGLLTSQAKLQGYPQNISEGELIV

>Sl03G116000

MRAISPHCSSVSAFRWLSSHAAVRVDNHRQCLTGPTNHTSHLHTEKTEPLISLIKSTSSKPHLLQIHAHLIRKSLFQDPIFFSPFLFGIALPPFHDLGYASQVFSKFRKPDVFQYNIMIRAYGMSDSPGNGFMLYQEMLRSGVSPNSLTSSFVTNCCIKIGSLFGGLQIHARILRDGHQSDGRLLTTLMDFYSSNEKYTEACKVFDEMSHRDTIAWNVLISVYMRNRRTRDALGLFDMMQSSYDCQSDDVTCLMLLQACANLNALAFGERVHRYCEEHGFDKAMNICNALITMYSRCGCLEKAFEVFKGMTEKDVVSWTAMISGLASNGYGRDAIEAFREMQRVGVSPDDQTFTGVLSACSHSGLLDEGRMFFNSMSKEFGISPNIHHYGCVVDLMGRAGMVDEAYNLINSMKVKPDATIWRTLLGACRIHHQAELGEQVIERLIELKAQEAGDYVLLLNIYSSLGDWGKVVNVRKMMKDRGIQTNPACSTIEFRGKIHEFVANDFSHPRKTEIYETLDEINQQLRIAGYVAETVAELHNVGTEDKQIALSYHSEKLAIAFAVLSTPPGTSIRVAKDLRICVDCHNFAKILSAVYSREVIIRDRNRFHHFREGRCSCNDYW

>Sl03G116840

MNYARKKKQGVTMDTLTTLNEIADSEAQPNKFRRKKSIVWEHFTIERIGADCTRACCKKCKKSFAYISGSKLAGTSHLKRHIALGICPVGRTNQDKNQLTSFNSAAPTNGSAGATGKSRKRYRANPGPTSVPFDQARCYHDIAKMIIQHDYPLEMVEHSGFNKFVQNLQPLFSSVSVDTIQEHIFNIYLGEKQNLLNIIGAIPGRVSLTLNLRTSDQNLGYVFITGYFVDSDWKLRCRLLNVIMVPFPDSDVAFNHAVAACLTDWCLETKLFTLTLDQSVANVNVRKNLGHLLSIKGVNILNGQLIIGSCCARVLSDLAQYALHYMRAIVEKVRQSVKFVKTADAHEEKFLELKRQLQVPSAKELIVDDQTKWDTTYQMLMTASELKEVFSCLDTSDPDYKVTPTMDEWKQAEILCEYLKLFFDAANLLTSPTYSTADVLFHEVWKIQLDLMQAARSQDRFIRDLTRPLQEKFNEYWNDCNLVLAVAVVMDPRFKMKLVEFTFNKIYGEEAETWIKIVDEGVHEVFCDYIVQSLPPPPASFVEEANDNFVIKSEFSQEDSFLATNGDAFPDFEVYLDIINNQQMKTELDQYLEESLMPRSQDFDVLGWWRINRCKYPTLSKMASDILSIPVCTVTPDSVFDTVSRDLDRHRSSLRPITIEALSCSKDWLQYESWEPPYGTPDATVKMNMMLSSLRGLSNLTYTFHGKGVFLLASQCPNQVFQLAHLHLSPEIHLPLQSLIEKCKSMDQLRQIQSVIIQKGLISDPKLCSNMITFCSNNESGDMKYARSVFDIMPERGVFIWNTMIKGYSRENIPHDGVSIYREMLNNNVKPDNYTFPFLLKGFTREVSLKLGRSVHAHICKFGFELNEFVHHALIHVYCLCGQVDMARGVFDLSAKSDILIWNSMISGYNRSKQFGESRKLFYAMEEKQLQPTSVTLISVISALSQLKDLDTGNRVHQYVKDYKVQSSLVLDNAIVDLYASSGKMDVALGLFQSMKHKDVISWTTIVKGFVYIGQVDVARIYFDQMPKRDNISWTAMMDGYVKENRFKDVLMLFREMQAAKIRPDEFTMVSILTTCAHLGALELGEWIKTYIDKHKIYVDIHLGNAVIDMYFKCGSVEKALVMFTQMPSRDKFTWTAMIIGLASNGHEREALDMFFEMLRASETPDDVTYIGVLSACTHMGLVEEGKSFFANMASQHGIQPNVIHYGCLVDLLGRAGRLEGAYEVIMRMPVKPNSIVWGALLGACRIHKDVQMAEIAAQQLLQLEPGNGAVYVLLCNIYAACKKWDNLRETRRIMTDRGIKKTPGCSLIEMHGIVHEFVAGDQSHPQSKSIYSKLAELIGELKFSGYVPDTSEVSLDIGEEEKENSINRHSEKLAIAFALINSEPGFTIRIVKNLRICTDCHHVAKLISKRYNRKLIIRDRTRFHHFVQGSCSCKDYW

>Sl03G119950

MAAVSAKTPTISLPLDPQNPQFSKLHSSKSLNFSHIFQKSHLFSLKKPHQNSILSSSSTSTPTTDPNSHLIQLCFHNQLEQAIVFLKSIKDLHGTIEEDTFVTLARLCEFKRASNEACEVFSCIHNCMTQLSLRLGNALLSMFVRLGNLGDAWYVFGKMEERDVFSWNVLIGGYAKNGYFDEALDLYQRMLWVGIRPDVYTFPCVLRTCGGLPDWRMGREIHAHVIRFSYDSEIDVVNALITMYVKCGDVCSARVLFDGMSKRDRISWNAMISGYFENGEFLEGLVLFSSMREFGFFPDLMTMTSVISACEALGDDRLGRALHGYVARMEFYSDVSAHNSLIQLYSAIGSWEEAEKIFDRIQCKDVVSWTAMISGYESNGFPEKAVKTYKMMELEGVMPDEITIASVLSACTSLGLLEMGVKLQHVAERRGLIAYVIVSNTLIDLFSKCNCIDKALEIFHRIPDKNVISWTSIILGLRINNRSLEALNFFREMKRHQDPNSVTLMSVLSACSRIGALMCGKEIHAYVLRNGMEFHGFLPNALLDFYVRCGRRAPALNLFHMQKEDVTAWNILLTGYAQRGLGALAIELFDGMISSRVKPDEITFISLLRACSRSGLVTEGLDYLNSMESKYCIVPNLKHYACVVDLLGRAGLVEDAYDFILSLPVKPDSAIWGALLNACRIHRQIELGELAARHILETDERGVGYYVLLCNFYSDNGRWDEVVRLRKIMIEKGLTIDPGCSWIEVKGNVHAFLSGDNLHPQSKEINAVLEGFYEKMEAARRSKSERHTVNEVKDSKAEIFCGHSERLAIGFGLINTAPGTPIWVTKNLYMCKSCHDTIKFISEVVRREIAVRDTEQFHHFKDGRCTCGDENYLGET

>Sl04G009000

MTKDTSLFKKLITWDLFTLSNSTPFAKILDSYINTKSQYVIQTVHCRVLKTHFSSEVFINNKLIDTYGKSGVLKYAKNVFDKMPERNTFTWNSMMNAYTASRLVFEAEELFYMMPEPDQCSWNLMVSSFAQCELFDSSIEFLVRMHKEDFVLNEYGYGSGLSACAGLRDSRMGTQLHASVAKSRYSRSVYMGSALIDMYSKTGDVDCAAKVFNGMCERNVVSWNSLLSCYEQNGPVKEALVVFARMMEFGFKPDEKTLASVVSACASLCAIREGKEIHAQIVKSDKLRDDLIICNALVDMYAKSGRIAEARWIFDRMPVRSVVSDTCLVSGYARVASVKTARAVFSGMIERNVVSWNALIAGYTQNGNNEEALNLFLMLKRESVWPTHYTFGNLLNACANLADLKLGRQAHTHILKHGFRFQNGPEPDVFVGNALIDMYMKCGSVEDGSCVFTKMLDRDWVSWNAVIVGYAQNGHAMEALETFNAMLVYGEKPDHVTMIGVLCACSHAGLVEEGRRYFYSMDRDYGLTPFKDHYTCMVDLLGKAGCLEEAKDLIESMPMPPDSVVWGSLLAACKIHREIELGKYVAEKLLEIDPTNSGPYVLLSNMYAEQGRWQDVKMIRKLMRQRGVVKQPGCSWIEIQSQVHVFMVKDKRHTQKKEIYLILNTLTKLMKLSGYVPNAGHLDGDEEQTMLDFNSSEEFEEPVTAAIAC

>Sl04G009210

MLNVTVSSNSATLTQKFITFIEKCKSISELKKLHALLITCGISKETQFSSRILCFTALSDSSSIDYAHRVFLQIKTPTIFDYNALIRGYSSSKNPCKSLSLFVEMLQNEVFPNYFTYPFVVKCLAKLSEVRIGRSVHGGVLKNGFDVDLYVSNSLIHMYGSCGDVLCARKVFDEMPVRNLVSWNSMMDGYGKCGDVVLMREVFDSMIERDVVSWSSLIDGYVKDGEYAEALAMFEKMRVEGPKANEVTIVSVLGACAHLGALEQGRVMHEYVVENKLPMTLVLRTSLVDMYAKCGAVEEALVVFREALGRKTDVLIWNAMIGGLATHGLVTESLELYKEMHVLKVRPDEITYLCLLCACAHGGLVKEAWCFFDSLGKDGMTAKCEHYACMMDVLARAGRLTEAYRFLCEMPMEPTASMLGALLSGCINHGRLDLAEIVGKKLIDLEPFHDGRYVGLSNVYALKKRWDEAKAMREAMDTRGVKKLPGFSVVEIFGALHRFIAHDKAHPESDQIYTILDFVLWQMKLDKDCEEPEQLSCDINGGLSNGVDSSALQMNDFSL

>Sl04G064750

MRHIVSSLRQYSLLLHRRRKFSATTQAKQSRNEDPEPQSAPVTLLYDNLLKICLQECKNLQSRRVFDEMPQRVARAVKACKTIHLQSXXXGFASQGHLGNSIVDLYAKCGDMVSAEKAFFWLENKDGMAWNSIILMYSRNGLLENVVEAFGSMWNSGVWPNQFSYAIVLSACARLVEVEIGKQVHCSVVKTGFEFDSFTEGSLIDMYAKCGYLIDARRIFDGAVEPDNVSWTAMISAYIQVGLPQKAMEVFEEMQERGCVPDQVASVTIINACVGLGRLDAARQLFTQMTCPNVVAWNVMISGHAKGGKEVEAIQFFQDMIKASIRPTRSTLGSVLSATASVANLSFGLQVHAVAVKQGLESNVYVGSSLINMYAKCQKMEAASEIFNSLGEKNEVLWNALLAGYAQNGSACKVVELFRSMRLSTFETDEYTYTSILSACACLEDVEMGRQLHSIIIKNKFASNLFVGNALIDMYAKCGALGDARQQFDKMLTRDHISWNAIIVGYVQDEEEEEAFNMFHKMTLERIIPDEACLASVLSACANIHDLNKGKQVHSLLVKYGLESGLFAGSSLVDMYCKCGDITSASEVFFCLPDRSVVSTNALISGYAQKNINYAVHLFHNMLVEGLRPSEVTFASILDACSDHAYMLGMYYDSGKLEDASFLFSEFTKLNSPVLWTAMISGNIQNDCCEEALIGYQEMRKFNVMPDQATFASALKACSTLAFMQDGRKIHCLIFHTGFDMDELTSSSLIDMYAKCGDVKCSVQVFSEMVSKKDIISWNSMIVGFAKNGFAEDALEVFEEMKRASVKPDDITFLGVLTACSHAGMVSEGRQIFKDMTSLYDVRPRADHCACMVDLLGRWGNLKEAEEFIERFDFELDAMIWSAYLGACKLHGDDTRGQKAAEKLIELEPQNSSSYILLSNIYAASGNWGGVNFLRKEMKERGVRKPPGCSWIIVGQKTNMFVAGDKFHPCAGDIHALLKDLTALMKDEDGHVVVLWIGHFGFGSWRKGYTEEDLVKALSGQPKVGFRQYAGYMDVNVKAGKSLFYHFVEAKVKPDDKTLTLWLNGGPGCSSIGGGDFT

>Sl04G074160

MELDLQSCARLLNTVNSNQSLPNGKQLHLVFLKRGILNSALTIANRLLQMYTRCGQMADAQLLFDEMSQRNCFTWNTMIEGYMKWGKMNNSLDLFRLMPSKNEFSWNVVILGLVKAEELGVARRLLSEMPRKNEVVWNGLIHGYAKMGFPGVALCLFKEFIDWDFREMGGASHIDSFVLATALGACADTRSADLGKQIHARIIVNEVEVDSVLASSLVNMYGKGGDLDNANYILNRMQNPDNFSLSALISAYSKRGRMDDARKIFNLITDPSIVLWNSMISGFVSCDEVLEALLLFGEMHREGVIGDSSTLASVLNACASAYALKNCLQVHVYGFKLGLLDDLVVASALIDTYAKCGCPDEASNVFNELKTQDTILLNSMITIYFNCNRIEDARQLFESMPYKSLISWNSMIIGLNQNGCPVEALDLFYRMNRMDFRMDKFSFSSVISACASIASVELGEQIFARVVIIGLDGDQIITTSLIDFYCKCGFVSDARKLFDQMMKSDEVSWNSMLMGYATNGYGNEALNLFHEMRSVGVSPTNITFIGVLSACDHCGLLEEGKRWFYSMNYDYHIDPGIEHYSCMVDLYARAGCLEEAVNLIKKMPFEADSSMWLSILRGCVAHGNKILGQLVAQRIIELDPENSGAFVQLSNIFATSEDWERSALVRRLMIEKKIHKSSGRSWSDM

>Sl04G078520

MEGGLPMLNCLLQHTLRSVCTCSDSSSNASEWVYAVFWRIVPRNYPPPKWDHGGGLLDRAKGNKRNWILVWEDGFCDFYECERSKREHVTINFGPEIFFKMSHEVYSFGEGLVGKVAADNSHRWVSKDAPNEKDSNFTCSWNMSIEAQPRAWGVQFNSGIQTIAIISVREGIIQLGSFNKAFEDHNLVLNIQRKFSYLQSIPGIYAIQRPFLPIQHPYTYKPNNVTLVNETDNQMDDKNQIIGSKRVHEFPFKSINFGYNSPQTMASLPLWSMPIAAPSCYANAAHEMSSLHDRVTRNTTSKDVKVVDELGHLKFETDEENDQFSLNQNLGLENKVVEVGFRQLGNGGAAPNPN

>Sl05G005810

MEEQLNSLAITHLLQHSLRSLCIHENSQWVYAVFWRILPRNYPPPKWDNQGGAYDRSRGNRRNWILVWEDGFCNFAASTAEINANECPGSSSNNNNNNNYGEYQHYQGLQPELFFKMSHEIYNYGEGIIGKVAADHSHKWIYKEPNEQEINFLSAWHNSADSHPRTWEAQFRSGIKTIALIAVREGVIQLGAVHKVIEDLSYVVLLRKKFSYIESIPGVLLPHPSSSAYPFKVDGYGASPDAWHFQTNLPTPTPTPTELYEHFNQHQHMRITPSMSSLEALLSKLPSVIPADVAAGMTGGSIPTTYCHEYQQQPQYRPNVEILGLEKVAKEEYEDEEEEKENNNNEEKTRNNNNSNERLDHNGGESSSSMSSYSQHHHNYHHQHYGYHHDLNVSSSMPNNGY

>Sl05G006800

MILSTTPTRPTTTLTINTNTSEHRRHILQILTQCTSISQLKQVHAYTLRTTPLDHPDALFLYSRILHFASMNDLDYSFKLFGNLENPNSFIWNTLIRGCAHSNDRKGEAFMLFQRMVESVEPDKHTFPFVLKGCAYLFALSEGKQAHGVALKLGFDSDVYVNNSLVHFYSSCGCLKDARKVFDEMPERSLVSWNVMIDALVQSGEFENALRMFSEMQKVFEPDGYTMQSVLDACAGLGALSLGMWCHAYILRKCESFLDFELLVNNCLLNMYCKCGSVDIAVQVFERMSRHDLNSWNTMILGFAMHGEVEAAFHCFNQMVSKRVMPDSITFVGILSACNHRGFVDEGRSYFDKMVSEYKIRPVLEHYGCLVDLLARAGCIDKALDVVSNMPMKPDAAIWRSLLDGCCKKNADIEFSEEVARKIMESDGSDTSGVYVLLSRVYATANRWDEAGMIRTLMTDKGIRKDPGCSLIEINGVFHEFFAGDTSHLHTREIYEFLDVIDKRLLAAGYVPDLSHASTVDELDNGKRQSLKLHSERLAIAYGLLKLKPGTPLRIFKNLRICSDCHNVTKLISKVFDVEIIVRDRVRFHHFRNGSCTCKDYW

>Sl05G013200

MAIQTPLLIVGPFSCQEITKKNSRKDTFTPIDPLALVPKCKSLRDLKQIQAFSIKTQLQNDIFFMSKLINFCTKNPTPACMYHAHLLFDKIPQPDIVLFNFLARGYAHSDTPLNAFVLFLKILTLGVVPDFYTFPSLLKACAGAEALEEGKQLHCLLIKYGLNGDMYVCPALMNMYIEFKDNDSARRVFDRIADPCVVTYNAIIIGYVRSSEPNEALLLFRELQVKKIKPTDVTILGVVSSCALLGTLGFGKWVHEYIKKNSFDQYVKVNTALIDMYAKCGSLADAISVFESMPYRDTQAWSAMIMAYAIHGRARCAITLFQEMQNTKVNPDGITFLGLLYACNHSGLIEEGFRFFNSMTENYRIVPGVKHYGCMLDSLARAGRLTDAFKFLTELPIPPTLLLWRTLLAACSIHGNVDLGKLVLERIFELDKSHSGDYVIFSNMCARAGKWEEVNYIWNLMKERGIKKIPGCSSIEVNNVLHEFFSGEVTCIEHRELHQEVDKLIEKLKLVGYVPDTSIAFRPGLNDEDKEATLRYHSEKLAITFGLLNSPPGKTIRVVKNLRICGDCHSAAKFISLIFKRNIIIRDLHRFHHFEGGNCSCGDFW

>Sl05G050560

MTMLWSDEDKTMVAAVLGTKAFDYLMSSLVSAECSLMAMGSDENLQNMLSDLVERPNASNFSWNYAIFWQISRSKLGELVLGWGDGCCREAREGEESELTRILNIRLADEAQQRMRKRVLQKLHMFFGGTDEDNYVSGLDKVTDTEMFFLASMYFSFPRGQGGPGKCFTAGKHVWLSDVMRSSVDYCSRSFLMKSAGMQTVVLIPTDIGVMELGSVRTIPESLELVHSIKSCFSSFLAQVRAKQAAPLAAVVAEKKNGNNSVFPSSFPFDQSKENPKIFGQNLESGSTEFREKLALRKPVDGPLEMYRNGNRAPIINTQNGVRPVSWASFGNVKPGNSVDLYSPQAPPNNLREFVNGGREELRLNSLQHQKPGGMQIDFTNSRPVVSPVPTVESEHSDVEVSCKEKHAGPADERRPRKRGRKPANGREEPLNHVEAERQRREKLNQRFYALRAVVPNISKMDKASLLGDAIAHITDMQKRIRDAEYKLEKRGSTSVDAADINIEAASDEVIVRARCPLGTHPVAKVVEAFKETQVSVVESKLAVGNDTVYHTFVVKSSGPEQLTKEKLMAAFAGESNSL

>Sl06G053850

MCYWKLPTVHFSGNLSIRTYPNVQTTHKYMNQAVSRLLQTCKTLHSLKSVHAHLLVCGSIASSDLVLNKIIRLYTRFGATNYARKVFDEIPERNPFLWTSMIHGYVENSQHTQAFSLFLDMHIGDVTPLNFTISSILKALGRLKWSRHSEGMLGIIWKCGFGFDLLVQNSVIDCFMRCGEVDCARRVFDGMEEKDVVSWNSMLSGYVTNDKLEIARELFDSMDEKNVVSWTSVICGYARKGDMEEARNLFDTMPTKDMAAWNVMISGYTDVGDMQTANSLFQAMPVRDTGTWNLMISGYCKVTELERARDYFEQMPYRNVVSWTMMIDGYVKSGKFHEARCLFDEMPEKNLVTWSTMISGYAKNGKPSAALELFRNFKKQNLEVDETFILSIISACSQLGIVDAVESVMSGDVGSRYFSDTRVVNSLVDLYAKCGNIEKASQVFEMADKKDFYCYSTMIAAFANHGLVEKALHLFEDMQRENIEPDEVTFLAVLTACNHGGLIDEGRRYFKQMTEEFRIQPSEKHYACMVDILGRGGFFEEAHEMILSMHVAPTSAVWGAMLAACNVHRNVQMAEVAASELFKIEPDNSGNYILLSNIYAAAGRWHDVARVRALIREHHVKKNRGSSWIELDSAVHEFVMGDVSHVEVDRICFILSLLNEDMKLSGYTKDTDLHPISTRYPSYLSLSSDTEMDEELF

>Sl06G065040

MQRGTTGDGGGGLSRFRSAPATWLEALLESDTESEVILNPSSPILHTPNKPPPHPSTPKLKLETGGATRFTGDPGLFESGGSSNFLRQNSSPAEFLSHISSDGYFSNYGIPSSLDYLSPSVDVSQSAKRTRDDDSESSPRKLVSQLKGESSGQLHGSGGSLDAEMENLMDDLVPCKVRAKRGCATHPRSIAERVRRTRISDRIRKLQELVPNMDKQTNTADMLEEAVEYVKFLQRQIQELTEHQKKCTCSMKDQ

>Sl06G069600

MSHHTWNFSHQKQEQQVVEKEEEENRYTRGHVHNQQNQVDPMSNKCEVAELTWENGQVAMHRLGSNLSNEQTKHTWGKAGDTLESIVHQATFQKQHHSYIMGSDGQNQANINREKNVSYGAQQTRGVLKRMRSSDSDPQLYIGGISLEHLNARASAKDNDITMITWPCNEDSACHGGSENKEEERETKSSNPSKRSRRAAVHNQSERRRRDRINEKMKALQKLVPNASKTNKASMLEEVIKYLKQLQAQIQLISYAKNMEQQMMMMSLGMQPAHIQMPLLATMGMCSSTTGILNNMTSNLAPAPYQSLIGGRAPLIYPTSSMPTLFPPFMSPPFATASSIPSTPPQPINAESISPKLTKYAAPPNIAASTSFPFSHPYNAYLPHSMKMEFNNEMAAQYLQRGNQENVNIQGQKK

>Sl06G074110

MGYLLKEVLKTLCGVNQWSYAVFWKIGCQNTKILIWEESYYETSTLSNIHGTSGVENPELAFQDWSTGWAFGGVQNSQLQNQAGENLHLLINKMMMDNQFNLVGEGLIGRAAVTGKHQWVLSEGLSRNVHPPEVLRELRQQFSAGIQTISVIPVLPHGVVQFGSYLHIMENMGFVEDVKTLMSQLGCVPGVLLSDENATKEPALETSRSVYLGSSVSTEYCGRAKVMNSASIIDKGNSIQTEGFVGQTSFSLVDATFQDSNFTQTFADCHDNHLHKKISPQVKPCMYMNNQLTNSVIKTEVIPPNTDMWKKQQDSQYIPKPPFCQESSVGSLPLDSDSIMLTEQQISGENSLAKSNLTLPNFLGSSHGRSHHAVMYKSIPHPNFIADASRPPQKIISCTEHIGDGLQIGSSDLMASSKYDVNHVINNHSLDGQGAEYLLDGSKRMVENDLFQALGPILTQNENPSSSECIQDFYSEKIEHGARFPLFDSAYGDVHVQCQSGDDLFDVLGADFKKNHLNGSWNNGQCKEPNSNTKDWIKNSSTSTISQDASSTINQGNSDSCMFSMTGFDRILDTMVSSHSAKQSLDDNVSSRTTITNLSSSSAPNASCSYDRVGVSSQIQGEQFVSPKTLLKSGAISSSYKSECSKEDTGMYSQSSSIYGSTISSWVESGYDTKPSSSVSTGYSKKPDEMSKTSRKRLKPGENPRPRPKDRQMIQDRVKELREIVPNGAKCSIDALFERTIKHMLFLQSVTKHADKLKQTGESKIISKEGGLLLKDNLEGGATWAYEVGSQSMVCPIIVEDLNQPRQMLVEMLCEERGLFLEIADIIRGLGLTILKGVMETRNDKIWAQFAVEANRDVTRMEIFISLVHLLEQTAKGGTEPVNAADNNTAMVHSYHQAAAKPATGRSCSLL

>Sl06G076900

MKLSYGCMSLPSSMQQMITKVRAIGRKCTSTVQHFASGLLVIEQSRASTAVLVHLQDLGQLPASLSRSCTSDNGILPVNSWFLSKHLVSLRAFELTLKCGCPNKVKIQINVKESISTITVLDFISDEEIMATPYTQVLPLPRHQHFPKPNPISKTVINDRYFENHPLVLLIDKSQSINQLKQIHAYMLRIGLFFDPFSASKLIEASSLSHFSSLDYAHKVFDEIPQPNLFSWNALIRAYSSSQDPIQSILMFVNMLCEGREFPSKFTYPFVFKASAKMKAIRFGRGLHGMVVKGRDVGLDIFVLNSLIHFYADCGCLDEAYLIFENMQTRDVVSWNTMILGFAEGGYADEALKIFHRMGEENVRPNDVTMMAVLSACAKKLDLEFGRWVHAFIKRNGIRESLILDNAILDMYMKCGSIEDAERLFRKMGEKDIVSWTTMLVGYARAGNFNAARSILNTMPSQDIVAWNALISAYEQSGKPKEALSVFNELQLIKKAEPDEVTLVCALSACAQLGAIDLGGWIHVYIKKQGIKFNCHLTTALIDMYSKCGDVEKALEMFDSVNIRDVFVWSAMIAGLAMHGRGKEAISLFLKMQEHKVKPNSVTLINVLCACSHSGLVEEGRAIFNQMEYVYGIVPGVKHYACLVDILGRAGELEVAEKLINNMPVTPGPSVWGALLGACRLHGNLELAEQACNRLVELEPENHGAYVLLSNIYAKSGKWDEVSMLRKRMRECGLKKEPGCSSIEVHSIVHEFLVGDNTHPQSQKIYAKLDEIAARLKHVGYVSNKSQILQLVEEEDMQEQALNLHSEKLAMAFGLISVAPSQPIRIVKNLRVCADCHAVAKLLSKLYDREIILRDRYRFHHFKEGNCSCKDYW

>Sl06G082880

MEHRPGRFTTLAAAANQMHPNNYRRTFSHIYQECAKHCTQQPGRQAHARMIISGFQPTVFVTNCLIQMYVKCSNLGYADKVFDKMPLRDTVSWNAMIFGYSMVSELDKAQLMFDLTPERDAISWNSLISGYMQNRNYGKSIQTFLEMGRDGIAFDRTTFAVILKACSGIEDSWLGMQVHGLVVRLGLATDVVTGSAMVDMYSKCKRLDESICFFNEMPEKNWVSWSALIAGCVQNNKFSDGLHLFKNMQKGGVGVSQSTYASVFRSCAGLSDLKLGSQLHGHALKTDFGYDVIVATATLDMYAKCNSLSDARKVFNWLPNHNLQSYNALIVGFARGDQGYEAVILFRLLLKSYLGFDEISLSGVFSACAVFKGRLEGMQLHGVACKTPFLSNVCVANAIMDMYGKCEAPQEALRLFDEMEIRDAVSWNAIIAAYEQNGHEDETLILFFRMLKSRMEPDEFTYGSVLKACAARQDFNTGMVIHNRIIKSGMGLECFIGSAVIDMYCKCEKVEEAEKLHERMKEQTIVSWNAIISGFSLCEQSEEAQKFFSRMLEEGVKPDNFTFATVLDTCANLATVGLGKQIHAQIIKQELQSDVFITSTLVDMYSKCGNMQDSRLMFEKAPKKDFVTWNALVCGYAQHGLGEEALQIFEKMQLEDVRPNHATFLAVLRACAHIGLVEKGLQHFNSMSNNYGLDPQLEHYSCMVDILGRAGQISDALKLIQDMPIEADDVIWRTLLSMCKMHRNVEVAEKAAKCLLELDPEDSSSHILLSNIYAAAGMWKEVSEMRKVMRYGGLKKEPGCSWIEIKSVLHMFLVGDKAHPRCNEIYDNLDALICEMKRTSQILDNELLLSCESLLAVDQFSLNIGWLLMEVCKGKKLRFMLTQGKAWASLGYPREILSCMWLVTGLHLNIVSLPNYMHRTLSIFRDNSTV

>Sl07G005400

MNSLLSQQQQSQISLQDLQNGGNGGSTGGVGGLSQHSMGHSHFDPTSSHDDFLEQILSSVPSSSPWPDLSKSWDPHHHLSSPPHNPSSGEDQPPSNPFHSQFHYDDQASSLLASKLRQHQITSGGGAAAAAKALMLQQQLLLSRTLAGNGLRSPNGASGDNGLLSLPLNLSNGDQNDGVANPTNDNSVQALFNGFTGSLGQTSNQPQHFHHPQGGSMQSQSFGAPAMNQTPAASGSAGGGGGSTPAAQPKQQRVRARRGQATDPHSIAERLRRERIAERLKALQELVPNANKTDKASMLDEIIDYVKFLQLQVKVLSMSRLGGAPLVADMSSEGRGEGNVGRGGNGRASSSSNNETMTVTEHQVAKLMEEDMGSAMQYLQGKGLCLMPISLATAISTSTTRISNNPLLAPEAGGSTSPTLSALTVQSATAGKDATSLSET

>Sl07G005740

MPLTTKSYSQLNNISTQILNFLNKTSISPSQISQIQAQIIHNNLHFNTTIAHNFISVSKSLGLFNSAYTLYTKLIKKPHIFICNTLIQECSHSEIPVLKQNSISMYVHMHKESIFPNNYTYPFVLKSLSDLKELKLGKSVHTHVVKWGYVCDIYVQNSLLNLYASCGEIEFCQQVFDEMPERDVVSWTVLIMGYRDCGKFGDALVVFEKMKDSGVAPNRVTMVNALSACANCGALDMGMLIHDEIRRSGWAMDVILGTSLIDMYGKCGKIEHGFWVFQEMKHRNVYTWNAVIRGLALAKSGEEAVRWFFIMERENVKPDEITLVAVLCACAHTGMVEQGREIFSWLMNEKYGFPPGVKHYACMVDLLARSGHLEDALRMITDMPMEPTKSVWGGLLAGCRLHGNQELSEFAAWKLIGLAPRNSAYYVVLANLYGAMGRWNDAEKIRALMKERGLSKDLGSSSVELEDQKDLQELLR

>Sl07G006990

MNERQRLAELLRNCSKILSLDVGKQVHGAVLRMGYAFDLMIGNDLIDMYGKCSRVELARSVFHKMPERNVVSWTALMCGYLHHSNAQESLLLLSRMLFANVRPNEYTFSTNLKACGILGVLENGQQIHGLCAKSGFEKHPVAGNSIIDMYSRCGKLGEAEKKFHEMPEKSLITWNVMIAGYAMGGFGDKSLCLFKKMQQQGEMPDEFTFASTLKACSGFKAVREGSQIHGFLITKGFLISSQKVIAGALIDLYVKSGNLFEAHKVFSQVEQKSVISWTTLTVGYAQEGKLTEAMNLFKQLRESSITLDGFVLSSMMGIFADFTLIELGKQLHCCAVKIPSGLDISVLNSIMDMYLKCGLIEEAETLFDVMPEKNVISWTVMITGYGKYGLGGEAVELFKKMHMDRIEPDEVSYLALLTACSHSGLVQESEEFFSKLCNSNCLKPSVEHYACMVDILGRAGRLREAKVVIENMPVKPNVGIWQTLLGACRVHKNVEIGREVGEILLKLDGNNPVNYVMMSNIFADARLWEECEGLRGLVKTKGLRKEAGQSWVEIDKKMHFFYNRDETHPLTKAIHEFLYKMEKKMKYELGYTREVSFSLHDVEDETRDESLRFHSEKLAIGLALLSGSDEIEGKPIRVFKNLRVCGDCHEYIKGLSKILKKIFLVRDANRFHKFENGTCSCRDYW

>Sl07G062950

MVGSGTADRSKEAVGMMALHEALRSVCLNTDWTYSVFWTIRPRPRVRGGNGCKVGDDNGSLMLMWEDGFCRGRGTDCLEEMDGEDLVRKAFSKMSIQLYNYGEGLMGKVASDKCHKWVFKEPTECEPNISNYWQSSFDALPPEWTDQFESGIQTIAVIQAGHGLLQLGSCKIIPEDLHFVLRMRHTFESLGYQSGFYLSQLFSSTRTSSPSSAIPLKQPTMPIRAPPPLFNWGPRPMPSASSLLSSPNFQNSARLGIPQSKDESHMFLQLPHSSEPRMEDMMGAAADHESDIKWPNGLTFFSALTGRNDDSRILFNPDSLGSKPDHNQHPLSLDGKTSNPNSDASSLHNNGGANPNDFLSLDSHPDSIRKMDKFKRSYTLPARMASSSNSSTSLDQHANNPGEYRNEGGMYPDVMERFLE

>Sl07G066110

MTAASTSARAPLQNLTELQRGKRNEIPSSVLLLQMCREKKEVKQLHGQLILNGLIHRFPNGAKLVESYVGVSETDDALLVFNSSIQSPDTFAYNVMIRGLILSKRPIESLFLYERLVTDGLSPDSHTYTFVLKACSHMKAVLEGKTVHAQIMKKGIKPNTHICSSLISMYSCAGDMASARQVLDEFSEPNNVICPFNSMITGYMNEGLVEEAIEIFDTMGNKDTATWSVMLSGYVKNGMHEDALATFQKMMSYNVPLNEASLVCTLSSCGELGALDQGRWLHKYIINKRETVMSVNLGTALVDMYAKCGCIEFSYQLFEKMPRRDVVTWGVILSSFATHGQAKLCFQLFDKMIESGVEPNGVVFVAILSACSHAGLVEEGCHYFDQMVHQFGIRPSIEHFGCMVDLLGRAGRLAEAEQLILSMPEEPNSVILGALFNACRIHNDVERGRHLFKRLINLEPSPDRYKLAASLFANNGEDHEIRKLLNDKILETRCGLSNIQVDGVDHEFMVSDIANDKAQDVYETLGG

>Sl08G005390

MIQTGMISHIFPISRLISFCALDVNGDINYANALFSEISEPNVYIWNTMIRGFVKKQFLEMSFCLFRRMVREKVEMDKRSYVFVLKGCGVLKGVGVHCRIWKVGFLGDLIVRNGLVHFYGGSGKIVDAQKVFDESPVRDVVTWTSLIDGYVKMKMVDEALRLFDLMCSSGVEFNDVTLITVFSACSLKGDMNLGKLVHELVENRGVECSLNLMNAILDMYVKCGCLPMAKEMFDKMEIKDVFSWTSMIHGYARNGEVDLAKKCFSVMPERNVVSWNAMIACYSQNNRPWEALELFHEMEKRGLVPMESTLVSVLSACAQSGSLDFGRRIHDYYIKQKQVKFSVILANALIDMYGKCGNMDAAGELFHEMPERDLVSWNSVIVGCASHGLAQKAVTLFEQMKCSGLKPDSITFVGVLSACAHGGLVNQGWEYFRCMELNGLIPGVEHYACMADLLGRSGHLKEAFEFTKQMPVEPDKAVWGALLNGCRMHGNVELAKVAAEKLIELDPQDSGIYVLLASLCANERKWADVRMVRSLMRAKGVKKNPGHSLIEVDGNFYEFVAADDSHHESQAIHKILDEIILLSKLEEYVSDAQPEQT

>Sl08G008600

MENLNISTSSTPSQPNTLQKTLQYIIHNRQEWWVYAIFWQASKDVNNRLILSWGDGHFRGTKDTTGSTKTGHGQYHQFQKKFGFNDISETNNNVTDTEWFYMVSMPQCFVADDDLVIRAYTSASHVWLASYYELQIYNCERAKEANLHGIRTIVCISTTSGVVELGSSDVIQENWEFVQFIRSLFGSNNNMNTTSHLPVNQVTLGDDHKVAKCGSNIIVKQEMTIGNLLSESGISDFENDDSLTINNVMNGSIKRAKKGDSSHIRREMAMDVHVEAERKRREKLNHRFYALRSVVPYVSKMDKASLLGDAVTYINELKAKIKNLESKLIEPQKKHILMEQHDSHSASSTIVTDHGANNKSLFSSNGVRNGMEIEVKIIGSEGVIRVQSLDMNYPCTRLMNAMKEMKFQIYHASISSVKDLMLQDIVIRVPEEFSNEETLKSAIISKLSVMEN

>Sl08G014100

MNTLSPKSAHPIFKIIEQCKNIATLKQVHAQMITTGLIFHTYPLSRILISSSTIDATISYALSIFNHVTNPTIFLFNTLISSSLSRKKDDQTHFALALYNRILTQTTLIPNSYTYPSLFKACGSQPWIQHGRALHTHVLKFLEPPYDHFVQASLLNFYSKCGELGVARFLFDQITGPDLASWNSILAAYAHNYSVYYEADLDSVYDSSSLSLEVLLLFSQMQKSLTCPNEVSLVALISACADLGALSHGIWAHSYVLRNDLKLNRFVGTALIAVYSNCGRLDFARQVFDQLLERDTYCYNAMIRGLAVHGLGVEALELFKKMDLEGLVPDDVTMLVIMCACSNVGLVDQGCKFFESMKEDYGIEPKLEHYGTLVDLFGRAGRVKEAEEIVQTIPMKPNAVLWRSLLGAARVHGNLEVGESALKQLIQLEPETSGNYVLLSNMYASLNRWDDVKQLRKLMKDQGIEKAPGSSIVDIDGAMHEFLIGDKTHPELKWIYVKLDEMHRRLQEHGHKSGTREVLFDIEEEEKESALTYHSERLAIAYALIASDSGAPIRIIKNLRVCNDCHTATKLISRIYEREIIVRDRSRFHHFKNGTCSCLDYW

>Sl08G023640

MSGESTKPYIVKKCITLLLSCASSTYKFKQVHAFSIRRRIPLSNPYMGKYLIFTLVSLSGPMCYAQQIFNQIQFPNIFTWNTMIRGYAESINPYPAIEIHNDMCVNSVAPDTHTYPFLLKAIAKVIDVREGEKVHCIAIRNGFESLVFVQNSLVHFYGAISQAENAHKVFEEMSDKNLVAWNSVINGYALNSRPNETLTLFRKMVLEGVRPDGFTLVSLLTASAELGALALGRRAHVYMLKVGLDKNLHASNALLDLYAKCGNVNEAEQVFHELEEDSVVSWTSLIVGLAVNGFCEKALELFEEMERKGFVPTEITFVGVLYACSHCGLVDKGFAYFERMQKLFGVKPKIEHYGCMVDLLGRAGLVEKAYKYIKDMPLQPNAVIWRTLLGACSIHGHLALAEMTRNHLKQLEPNHSGDYVLLSNLYAAERRWSDVHQLRTTMLKEGVKKVPGHSLVELGNRVHEFVMGDRSHPKNEAIYAMLGEMTRLLRLEGYVPHTSNVLADIEEEEKETALAYHSEKIAIAFMLISTPPGTPIRIVKNLRVCADCHLAIKLISKVFEREIVVRDRSRFHHFTNGSCSCKDYW

>Sl08G075110

MMTMQQLLFDFNYDVEQNFSDENQDCYFDPDEFILPIEMNNSCCFMPEYSVLEKQPKDNHSCFIPEYSVFENIPKRQKIFQDDFFPNPNSNTITPSTHNSCFMPEYSVFEKQQKLFQDNFHEEGFLPNPPMFEDFALPEIPVPVFSAGVVAKKGGSSNNEKKMSAQSMAARQRRKKISDKTQELGKLIPGGHRMNTAEMLQATYKYIKLLQAQAGILAFIGSYQESFETPNLQKLVGSSLVQEKLYSSEHCLVPKVFVEALENNQEFQNSQILEEIKTLMKEGK

>Sl08G076150

MVRTSVLYQTPFLIPKEYHAKAQELNFSLKEQEWISMIKKCNNMRELKQVHGQILKLGFICSSFCAGNLLSTCALSEWGSMDYACLIFDEIDDPGSFEYNTVIRGYVKDMNLEEALLWYVHMIEDEVEPDNFSYPTLLKVCARIRALKEGKQIHGQILKFGHEDDVFVQNSLINMYGKCGGVRQSCIVFEQMDQRTIASWSALIAANANLGLWSECLRVFAEMNSEGCWRAEESTLVSVISACTHLNALDFGKATHGYLLRNMTGLNVIVETSLIDMYVKCGCLEKGLFLFQRMANKNQMSYSAIISGLALHGRGEEALRIYHEMLKARIEPDDVVYVGVLSACSHAGLVEEGLKCFDRMRLEHRIEPTIQHYGCMVDLLGRTGRLKEALELIKGMPMEPNDVLWRSLLSACRVHQNVELGEVAAKNLFMLKSRNASDYVMLCNIYAQAKMWEKMSYLVLA

>Sl08G081140

MAMGHQDQDGVPGNLRKQLALAVRGIQWSYAIFWSTAVTQPGVLKWIDGYYNGDIKTRKTVQAGEVNEDQLGLHRTEQLKELYSSLLTSESEEDLQPQAKRPSASLSPEDLTDTEWYFLVCMSFVFNVGQGLPGKTLATNETVWLCNAHQAESKVFSRSLLAKSASIQTVVCFPYLGGVIELGVTELVTEDPNLIQQIKNSFLEVDYSVILKRPNYVSNDAKNDTNIGSQKPDHNALENDAYPVEINSPHDSSNGFVANQEAEDSLMVVDGIGETSQAQSWRFMDDNISNGANNSLNSSDCISQNNANCEKLSPLSSGEKETKPCPLDRQENDQKKPHLLDHQGDDAQYQAVLSTLLKSSDQLTLGPHFRNMNKKSSFASWKTDIQMPRFGTAQKLLKKVLLEVPRMHAGVIHKFSRENGKKNSLWRPEVDDIDRNRVISERRRREKINERFMHLASMLPTSSKVDKISLLDETIEYMKELERRVQELEARSARRSNDTAEQTSDNCGTSKFNDIRGSLPNKRKACDMDEIEPESSNGLLKCSSADSIVINMIDKEVSIKMSCLWSESLLLKIMEALTDLHMDCHTVQSSNLDGILSIAIESKSTGSKTLAVGTIREALQRVVWKS

>Sl08G082930

MCDSMAKETLKRLCRSHGWSYGVFWGFDQTNSFFI

>Sl08G082940

MGSVIDEMLLQVHILGRGIIGQTAFSKKYKWMFTAANHERQISIRSSNNSNLFLDDNEFEQQFSAGIKTIAVLSVEPLGVLQFGSTNKLQESTCFVEQARTLFQGIGGSPTSSSCENLHFVNSTVFPTANNESLMKESHFLENLIQSVTCNAESQIMNSDVATAFLSENQFQDVNQFNNCSSQFDTQLQQAMFPSAGLFTSFHDSCLTSTWEDLPSDMSIQDFSYVLPTGINQFEYGTGATQSFHDNTTFGSLGGFGVLANEDTTGPLNGYIVQCPINQRNDGAVSTISDNILDTTGIISASAGINEQFRFNSDSDASVSIQSSITNAFETVEKANCSNMSAIEKMTNLVGVKHDSKKPCNWGDVSNPVVSTSNSEWTYSNANELRSRPANRLFSKLGLDQFLDGALSSSYSFAGSFSDGQLSETNKRRRVGSSSECNYLQKPLGFSNFDKNAKLVQPECGLDRTSNLEAKSEIITKLDASTLIGDRCSINNCRGNEKSSKPTKKKAKPGTRPIPKDRQLIYERLSELRGLIPNGEKMSIDRLLHRTVKHLLFLQGVTKHAEGLKKAESLKDSETRLNSKSNGNGVTWACEIGDQTMVCPLIVEDLSTPGQMLIEILYNEQGFFLEMVDIIRGFGLNILKGVMQSRETKMWAHFVVEAEGNRLVTRHEIFSSLVQLLHLTSASKVGLNNQLQYTSGGRNTLINDCPNSAVPISGCLPETIRCVR

>Sl08G083170

MQDFISTSSSFSSTSLLQKRLHYIIHNRQEWWVYGIFWQASKDANGRLIFSWGDGHFRDLALAKVHNANVSDMEMFYAVSAPNCFLSEDDLIVHAYNSGSYVWLNNYYELQIYNYDRAKEAHLHGIRTLLCISTPHGVVELGSSQVIQENLELVQLIKSLFGQINDHGFNFVPLGDPMDTKTITMGSDSGNSDESSAMNKDSPKKRARKSTTAKNHVEAERQRREKLNHRFYALRSVVPNVSKMDKASLLADAVTYINELKAKVEELKAKIEVSTKKLIQKRNCVSSSAVVDGTNINININSSFVDGMEVEVKIIGVEAMIRVRSPNVNYPCARLMNVLRELEFQIHHATVSSMKEMMQQDVVIRVPHNVTNEEAIKSVILTKLSFA

>Sl09G005070

MNSKEKKERVYSSAPKKVMKLSTDPQSIAARERRHRISDRFKILQSLVPGGSKMDTVTMLEEAIHYVKFLKTQIWLHQTMVNLVDINHEMVGYYPLVDDDQNIHKNNISSMDYQQMQQVQSYDNDAFQQVEFPFEETNISGDVFMYYN

>Sl09G008300

MKNLKKFHGHFITNGFSNDTLSLSAILYFTALSPTGDLAYAHLVFNQIDSPNTFMFNTMIRGYGSSSNLSEVMSFYIKMLQNGFFPNHYTYPFVIKALCRTQNYILGEALHCSVIKFGHVLDLHIANSLLHMYAKFGFFVEIMYLFDEMPEPDVVSWNVVIDNFVKNGCFDEVLDAVNQMCLNGVEPNAVTLLVLVSFSLKMGDFGLGKLIHLYVMKRGIHMSENLGNGLIDMYSKFGDMESAEKLFDMMKMKTVFSWTSLLDGFIQKGELQRAVVVFNLMPKDTTAWNVMLSGYSEAGDMSSAETMFRAMPDRDLVSWNTMILGYTQNKMYMKSLELLREMLGFGLRPDRITLVGLFSVCGYAGVLHIGEAIHSFMEKQNVKEEEVEVALLDMYSKCGDPEKALTVFYTIRRRKSVLAWTNMIVGLAMNGLANEALVLFHQMCDEGTDPNEITYLGVLCACSYAGLVKEGKWLFNAMSKVHGITPRSEHYGCMVDLLGRAGLVEEAEMFIQDMPEKADAGIWGALLGACRMHGEVQMGERIAKIVTQMDPYQSGRYILLSNIYAAENRWFDFG

>Sl09G011170

MSAASLRHFLESLCFKSPWNYAVFWKLQHQCPIILTWEDGYLDVPGAREPYRSQNGNYYSKNLSDLSPNCGSRSHNGYLSAHSIGLAVAEMSSTYHIAGKGVVGEVASLGIPRWISSDSVAPAELGFGSVAECPDKWMLQFVAGIKTILLVPCIPYGVLQLGSVETVAENMEMVTILAEEFDAHLKFVESFLPGGESCEFLLQSTLSETLNIPSATTTNKVNEDDVAADIPIVEDHKSSAVFPMTSLIDVQHPFQLSGQHMQNVLENENESKIGKFVEHMPNVLENAYKWEIPMQHVDMINLVKQLAHGYSDDNRSGITERSIVRSSCHTKDIDAFSYSSCNVGGVGVSNEVDFHFDGDMLDPRSLGMDCHNTILGNVSNSFSCSTERELHEAFGSTIHNLSGFSANPSSKSIYAADCTFNSEPSDGWHLKEDNAENLLEAVVASAYCFTDDYSLNKMAGLESLNMSSGKPVPSRKRLNQSAESDSVGDAVTRSTLTSASAGVDKYASTNRPHSASSFDYVVSTFDEGHHQTKVFSSLDCHKESKISNTNKKRRRSGDSHKPRPRDRQLIQDRLKELRQLVPSGAKCSIDGLLDKTIKHMLFLRSVTDQADKLRFQAQTEVAPDKNLQSPPIKSSNQQGTSWALELGSVDQICPIIVKDLEYPGHMLIEMMCDDHGRFLEISDVIHRLELTILKGVMEKRSESTWAHFIVEASGSFHRLDIFWPLMQLLQQVPSSVSRNI

>Sl09G065100

MEIIQPNSLQLQNMLQNSVQSVKWTYSIFWQFCPKQGVLVWRDGYYNGAIKTRKTVQPMEVTAEEASLHRSQQLRELYDSLSAGDSNPPARRPSAALSPEDLTESEWFYLMCVSFSFPPPIGLPGKAYSKKHHIWIMGANDVDSKVFCRAILAKTVVCIPLLDGVVELGTIEKVQEDIGFIHRVKSFFNEPQQAQPPKPALSEHSTSDPAAFSEPHFYFSNTPSSAGICPADQDGRITGEEENEDEDEDEAEDDEDENDEAELDSDGIAIQSGAGAANPMAAEASELMQLDMSEAIRLGSPDDGSNNMDTDLYLDGISQAGNTADSFKAETAISWANFQDLQHLPGIPSYDELSQEDTHYSQTVSAVLEHLSNTSSKFASSATIMGSISPDSAQSAFTLWPVTCSPNLSHCRRHDIGDGSGTTSQWLLKSILFTVPFLHSTKKLSEALSPKSRDAAAADSSAAASRFRKGCTINSCTQQEETSGNHVLAERRRREKLNERFIILRSLVPFVTKMDKASILGDTIEYVKQLRKKVQDLEARDRHTEITKKSDEKSGSPIVKAFPVKGKRRMKSTVEGSIVGAPAKMTGSPPMEEEVLQVEVSIIENDALVELRCPYKEGLLLDVMQVLRELKVEVVAIQSSLSTGLLLAELRAKVKENIYGRKASILEVKKSINQIIPRVN

>Sl09G066280

MASQLQQALRSLCCNTPWKYAVFWKLTHRARMMLTWEDAYYDNDGFPGKKSPDSTAGNLYDGHYSNNHLGVAVAKMSYHVYSLGEGIVGQVAITGKHLWLSANKVAAITNLAPEHCDGWQAQFSAGIKTIVVAAVAPHGVVQLGSLDSIPEDLRAIKHIRDVFSELQELMTSCLRSSMQHSMENSCLSEISTRTSGSEIFQDCVNNLGRSVCEDRRNMWSPLYTSFEKSVDHSCIFLQPGGYPNKILEVVNNQRLHRSSVQGSDDSTNLFCAGYELYEALGPVFQKGNSSKDWEAGKREEMAVDMLEGIGTSSLVMSNTGNEHLLEAVIANVNRHDNDCSSVKSFCKSVDSLLTTEITAEPCSSDIGTISSTGYSFDRETLNSFNSSGTCSIRSSRGLSSTSCSRGSGHVERPLEPVKMHKKRARPGESCRPRPRDRQLIQDRIKELRDLVPNGSKCSIDSLLERTIKHMLFMQSVTKHADKLSKCSASKLADKESGICGSSSHEVGSSWAVEVGNNQKVCPMRVENLGMNGQMLVEIFEDGSHFLDIAEAIRSLGLTILKGLAEAYGERTRMCFVVEGQNDRTLHRMDVLWSLMQLLQAKINL

>Sl09G082170

MAIMTHPFTTPPTTHPNTTHIHLSTSISKATSLPQLKQVHTQILRQNLSDSDSGSLLFDLILSSIPLPSSLQYSLSIFSTLQNPRTHLINKLFRELSRSKEPHNALLFLENGRRNGLEVDRFSFPPLLKAASRAFALREGMEIHGLGCKLGFISDPFIQTALLGMYANSGQIQDARLVFDKMSERDIVTWDIMIDGYCQNGLFDDVLVLLEEMRSSNVEPDSRVFTTILSACGQTGNLALGKVIHELISENNIIADSRLQSSLISMYAGCGCMDLAQNLYDELSQKNLVVSTAMISGYSKAGQVEAAHSIFNQITDKDLVCWSAMISGYAESDQPQEGLKLLDEMQASGVKPDQVTMLSVISACANLGALDQAKRIHMIVDKYRFREALPVNNALIDMYAKCGYLDGAREVFGRMRRKNVISWTSMTSAHAIHGEADQALMLFRQMKEPNWITFVAVLYACSHAGLVDEGQQIFSSMVNEYKITPKLEHYGCMVDLYGRANRLREALELVESMPMAPNVVIWGSLMAACRIHGEYELGEFAAKRLLELDPEHDGAYVFLSNFYAKGKRWENVGEVRQLMKHKGILKERGHSKIEMGNEIHKFLTADKSHKHADDIYAKLDEVVCKLMQVGYAPNTSIVLIDVDEDEKKDIVLLHSEKLALCYGLLKSSRGSPIHIIKNLRICEDCHNFMKLASKVFEREIVVRDRTRFHHYRDGSCSCKDYW

>Sl09G092150

MNMKFGFLTVNGLEDMFIAVLKNCKSSIFLKKIHAQIVKFSLTQSSYLVTKMVEICDKIGEIEYANLLFKQVEHPNNYLYNSIIRAYTHKHRYISCINVYKQMMTCAISPDEYTYPFVIRSCSAILRVDVGEQFHVHVCKFGLYCSNVIANSLLDMYVKCDRMRDAHMVFDEMSDRDVISWNSLICGHVRLRQVRKARALFDVMPDKSIVSWTAMISGYTKTGCYGDALDVFRRMQMVGVKPDWISLVSVLPACAQLGALELGKWIHFYADKYGYLRKTSVCNALMEMYAKCGSVNEAWQLFNQMSERDVISWSTMIGGLANHGRAQEALKLFHEMQRSAVEPNEITFVGLLCACAHAGLCDDGLRYFDSMKNDYNIEPGIEHYGCLVDILGRTGRLERALAIIKTMPVKPDSAIWGSLLSSCRTHRNLEIAVIAMEHLLELEPEDTGNYILLANIYADLGKWDGVSRMRKFIRSKSMKKTPGCSLIEINSLVQEFLSGDNSKPFSKDIHEVLELLALHQSKENDLVDTTLEYLSP

>Sl10G009270

MDEPIFSPSSSQSSLQHRLQYIVKNQTNYCSDWAYIIFWQSSNNRSCLTWGDGHLNMKITNNKDVEWFYLMSLAQSFCVGEGVVGKCFSSGSLVWLAGDQQFEFCHCERAKEAHYVHGINTFVCIPISSGVLELGSSTMIKQDLNLVQQVKSMFFGYETIDQFDDFGLFNCLELYGEEAKKGEVVVGTTPHENKAGLKNKTSKKRRREICETQGNHVEAERQRREKLNSRFYALREVVPNVTKMDKATLLSDAVTYITQLKAKVDELESKLHSNNYHYYYPEMKIKHKMENHDINVVDNQSSITTSRDHTMEIEVKMVGQDAMIRVQSENVNYPSTRLMCALQEVELHVYHANISSVNDFMLHDIVVKVPQGLETEDEVKYALLRSLDQQTCS

>Sl10G049720

MEHVVSMSEGDWSSFSGMCFTEEADFMAQLLGNCSFPNELPSNSGYWNIGHESNIGSSGGREHSSFFFPPLSHESHYSSNSRPILMRNDSSITTERGLMDTNNPIEADEYFANNMEFDANMAEPLLDGKGLQLGRIDYEDHSPTESSKKRVRLHCHVPKNKRSTKLQTEGKTVEMDMKSKAVLQRQNSMVSCCSEDESNVSLELRRKSRASRGSATDPQSMYARKRRERINERLKTLQSLIPNGTKVDISTMLEEAVQYVKFLQLQIKLLSSDDLWMYAPIAYNGMDLGLDLRIGNPK

>Sl10G049780

MEPVFEDEWSGMCSTDQEADFMAQLLWQPNNIDNMYFSSYNGCNSSQISFPSLESYYQSHVQSILTRNGSSITKENGMVEGENTSSPSQVLATYNPIEADDFLNQDVSMESGENTAKVLDSPPESSKKKRLCNLLGDVPKNKRSVNLKKASEMDGKNKAALQRQNSISCCSEDESINVPKSRASRGSATDSQSLYARKRREKINQRLRILQSLVPNGTKVDISTMLEEAVQYVKFLQLQIKLLSSDDLWMYAPIAYNGMDIGLDLKIGTPKS

>Sl10G084540

MNVVGKVSSFITQGVYSVATPFHPFGGAVDVIVVKQNDETFRSTPWHVRFGKFQGVLKGAEKVVRIEVNDVEADFHMYLDNSGEAYFIREVPADNQNEVNGDSRDSAKKEEVDTSDLDNVNNNDGVKEDNDSNKAELSSKDEGVTLGIERLDEGGSDGDRRSSEFLEEQSSLEDSAVAELSSSRCENADHVEEVLESQDSSSEVVMVSVDGHILKAPILSSEMNVEDVKLDTPRFHLGPGQGTDFSDSNSEFISGEATWADEYLSYQASPKVSPADACNVKKESSKVEHQTDVSEVDGRYSVNLDIKNQENGKVEITSCIIKKNAVANSCLELQASGTHVENEVNQPDVVSPAKIQEVVDDLENKPPGKLGDCSSLNPAKTLPQGESVMDSNGSDGSIDHQVVSDKHPEKQNNVSLATEAIQSGQKGPDECTQCVDVKHLIEAFLKVIVTGVEISLCGSLLHAGMGSSAAREAFDANYISEEEFRSSSESIIKNKNLVVRIRGNYILWDRAAPIILALAECDTELPVECADFIPVELEETSKSGEDGSEISSVPSGRRWTLWPNPFRRVKTLEKSNCNSSNEEVFVDSESGSVYQPMEQTAIAQGGKGSPPLQLVRTNIPTSSQITSLNLKEGQNVVTFIFSTRVLGEQKVESHIYLWKWNAKIVISDVDGTITKSDVLGQFMPLVGKDWTHFGTARLFSAIKENGYQLLFLSARAIVQAYLTKNFLFNLKQDGKTLPTGPVVISPDGLFPSLYREVIRRAPHEFKIACLEDIKALFPQDYNPFYAGFGNRDTDEFSYRKIGIPKGKIFIINPKGEVSINNQIDVKSYTSLHSLVDDMFPPTSMAEQEDYNIWNYWKMPISDIDYFNTLSLSSHIYINESRLHAKVLMNGLWQQTHFALSLLKSPNKLSTLNHLKSLHVYLLRTGLHRSSFAVGNFITHCASLGLMSYAALLFDQMPEPNSFVWNTLIRGFQQNRAPKYTLYYFDQMRANNVQPDRFTYPFAIRACSGLLECAKGVSLHGQVVKIGVNFDVFVGTSLVDFYTAMGDLNMTKRVFEELPEKDEITWYAMLSSYVNKFNDMRKARDLFEKIPCKDLVIWHTLILGYVKAGDLELAKKYFDEAPVKDLLMYNTILGCLAKNGEVECLLRLFREMPCRDLVSWNTVIGGLVRDGRINEAMRFFYEMERVNLSPDDVTLASLLSACAQAGALDTGKWLHSYIDRRCSELNAVIGTALVDMYSKCGDLGSAADVFNKMSERDVVAWSAMIMGSSMNGQSRTALNFFYRMKDESERPNDATILGVLCACVHGGLVDEGKKCFYGMSEEFGLTPKLEHYGCMVDLLGRAGLLDEAYSLIQSMPCEPHTGAWGALLGACKIHRNVELAEKAIEHLIQLDLDDGGYLAIMSNIYANAGRWEDVSKVRKLMKEKGIGKSRGISSIEVNGVIHEFGVQEKKHPQAREIYDMIDEIYRRLKRAGHVASTREVFFDVEEEEKEKALFFHSEKMAVAFGLIATDKTTIIRVVKNLRICPDCHAAMKLISASFEREIVIRDRHRFHHFKNGVCSCRDYW

>Sl11G006940

MSLIRKCTSTKPSSITVLNLLRESRCIKHLEQVHTHIIQKGYEQDHFIINQFISLCNVFSPDVSYATSVFEHVIQPNVYVWNTLIKGYSKKSSLVDCFVVLKQMRTSVNVIPDEFTFPSLVKSCSNVLALKEGEIIHGLLVRYGLDSDVFVGSSLIDLYGKCKEIEYARRVFDEISLKNEVIWTAMIVGYVYAGDLLEARKLFDEMPQRNVASGNAMIRAFVKFGDLSGAKKLFDSMPDKDVVSFTTMIDGYAKAGDMASARFLFDRSSNRDIISWSALMSGYAQNGQPNEAVKIFHEMLSMNVRPDEFIMVSLMCACSQLGRLDLANWVEHYMSQNSFDLNQVHIAAALVDMNAKCGNMERAKMLFEGMPKRDLVSYCSMIQGLSIHGCGSQAVDFFDRMLNEGLVPDDVSLKIILTACSRAELVKEGFRIFNLMTTKYSVKLSPDHYACVVDLLGRSGKLQDAYELIKSMPVEPHAGAWGALLGACKLHCDIEVGEEISNRLFELEPQNTGNYVLLSDIYAAANRWLDVALLRQKMSEKGLRKIPGCSSWLNANRLVTGTLCGFDQFMNLVIDNTVEVNGNDKNEIGMVLIRGNSMVTIEALEPVA

>Sl11G007230

MSFIQLIRGCTSSKSLFNGKSLHAQLLKLGSLKADIFTNNHLLTMYLKLNQFDDAQQLFDRMPERNIISWTTLISTYTQLGMYEKALGCFRSMNLEDGFGPNGYTYVAALSACSSLGAERTGKELHGRMLRTEERLNSFVSNCLVNFYGKCGLLISARIVFDGILEPNSVTWASLISCYFHCGEYGEGLNMFVLSLRGGVIVNEFFCGSVLGACAVVKSLQLGMQIHGLIVKLSLGMDQFVVTGLINFYAKCGRLELARQAFDEADGPELHAWTAIIGGCVQLGSGREAIELFCKLLSSGLKPSERTFSSVIGAFADVKEVRVGKQIHCRIVKMGFDSFSFVCNALLDFYSKSDLFEESLKLFQEMKEQDVVSWNTLIAGCVSSSRYEEALRFLREMLLEGFEPSLYTYSSILSICGDLPAIEWGKQTHCRVLKSRLDSNVVVDSALIDMYAKCGRLGYARRVFDILPAKNLVSWNTMVVGYAQHGFGKEALEIYGMMQSSGVKPNDITFLGVLSACGHVGLLDEGLHHFTSMTKVHGIIPRTDHLACVVSLFARKGKTKEAYHFIQSFSVEPDKVVWRCLLSGCKANRDFVLGKYAAEKILDIDPDDTSAYIVLSNIYAELQMWDETAKIRKLINAKALKKETGHSWIELQNKMYTFSACNIMSLQESYLQQVLTGLTAQLLDSDYNESIVVKPHYVRARNSTCGEKKDDKKTAAEFNKKEAKDQNANATRHGGAKGYTTTGEPEYSTTAYDPTNDTIGGGGGGEGPVLFDHPTIKTQDPSFPC

>Sl11G008940

MWTLLQTPPPPPPCGVQNPYSSRRISTAFGHSTGNDSASKSINLCLRCNYNDSIQSYCKKGELKQALQLLSQEPNPNQHTYELLILSCSEKNSFHDGLTVHRKLIDDGFEQDPFLATKLINMYSNLDSIDHARQVFDKINNRTIFLWNAFFRALTLAGNGVEVLKLYTRMNSIGIISDRFTYTYVLKACAVASLPKHKEIHAQLFRHGYHTHTHIMTTLIDVYARFGYVENAACVFDQMPQKNMVSWSAMIGCYAKNGKPLEAFDVFRDMMNHVLLPNSVTMVSVVQACAALGALEQGKVLHGYILRKGLDSILPVLSALVTMYARCGALELGRRVFDQMGKRDVVAWNSMISSYGIHGFGGKAIETFREMIRHGVSPSPISFVSVLGACSHAGLVEEGKELFDSMLKEHNICPTVEHYACMVDLLGRANQLEEAAIIIQDMRIEPGPKVWGSLLGSCRIHCNVELAERASRRLFELEPTNAGNYVLLADIYAEAKMWDEVKQVRKLLEAKGLRKVSGCSMIEVKRKIYSLQSVDELNIQIEQIHALLLKLSTEMKQNGMYVPDTRIVLYDLEEEEKERILLGHSEKLAVAFGLINSSKGDPIRISKNLRLCEDCHSFTKLISKYTNREILVRDINRFHHFANGVCSCGDYW

>Sl11G064750

MTDLKKIHAHLIKSGLIKDKIAASRVLAFSAKSPPIGDINYANLVFTHIENPNPFTWNTIIRGFSESSTPQYAIHLFIEMLNNSQVQPHLLTYPSVFKAYARGGIAKNGAQLHGRIMKLGLEFDTFIRNTLLYMYASCGFLVEARKLFDEDEIEDVVSWNSMIIGLAKSGEIDDSWRLFSKMPTRNDVSWNSMISGFVRNGKWNEALELFSTMQEENVKPSEFTLVSLLNACGHLGALEQGNWIYKYVKKNNVELNVIVVTAIIDMYCKCANVEMAWHVFVSSSNKGLSSWNSMILGLATNGFEDDAIKLFARLQCSILKPDSVSFIGVLTACNHSGLVEKAKDYFQLMKMEYGIEPSIKHYGCMVDILGRAGLVEEAEEVIRSMKMEPDAVIWGSLLSACRSHGNVELARWSAENLLELDPNESSGYVLMANMYAASGLFDEAMNERISMKEKHIAKEPGCSSVEINGEVHEFASGSTFMVEI

>Sl11G068960

MGYLLKEVLKTLCGVNQWSYAVYWKIGCQNTKLLIWEESYYEPSTYSGIHGVPMVENPELPFHDWGVCWGPGEVRNPQLMNQAGERVQLLVNKMMMESQINIVGEGLVGRAAVTGSHQWFHSEGLSRVVHPQEVLKELTGQFSAGIQTIAVVPVLPHGVLQFGSCLHIMENMSFVEDVRILISQLGCVPGVLLSDETAMKDPMVNTGTPVYMGSSVLVDSSVSTNVMNSAPVIASCSYQGNSSQNDGFIRQTSSSLDAQVQDRIMQSIDSSFQASNMTQRFVESHDDRQFHKKIIPEVKSHLSPNSQLINSVIKAEVISPSSNLWMSQQAPLHIPRPPFHQQSFTDSLTVDSSSLSNVSQLNGFTASDPRPNDVLISSYHGNSISPSNGENELCKRRDGHHRSIPCPNSIADANGLSNIISSCTKSTGNGLQTTSKFNVGDDSSTSHISDGQNAQFMWDESNGIVENDLFQALGIMLTQNEHPCSTSKSVQEVCVEKHEYVGQSALLENNKYEDSCVQRHSGDDLFDVLGADFKNKLFNGSRNSYQSNGPDSNTWDRVKSNSTSVLSQQASSIVNQGKSDSGSFFAAGFERLLDSISSKPSAKQNMDDDVSCRTTLTNLSNSSAPNVSSSYGRAGFSSQIQGNVFGKPKTLAKPGSTVSRSFRSEKEKTGAYSQSSSIYGSQISSWVEQGHDTKPTSSVSTGYSKKPDETSKTGRKRLKPGENPRPRPKDRQMIQDRVKELREIVPNGAKCSIDALLERTIKHMLFLQSVTKHADKLKQTGESKIISKEGGLLLKDNFEGGATWAYEVGSQSMVCPIIVEDLNQPRQMLVEMLCEERGLFLEIADIIRGLGLTILKGVMETRNDKIWAKFAVEANRDVTRMEIFISLVRLLEQTSKGAEESAKAIDNNTAMVHSYHQAASIPATGSRPCN

>Sl12G010170

MQAMNSFQSTGENGASSGEHMSHSHFDPSSSHDDFLQQILSSVPSSSPWPEISGDGHPYNFDDHQSTLLASKLRQHQINGGTSAAAAAKALMLQQQLLLSRGIAGNGGSGINGDQNDDGLNSGNDISVQALYNGFAGSLGQTSNQSQHFHHSQAQSFGAPAASLSMNQTPAASGSAGGAQPKQQKVRARRGQATDPHSIAERLRRERIAERMKSLQELVPNANKTDKASMLDEIIDYVRFLQLQVKVLSMSRLGGAAAVAPLVADRSSEGGGDCVQGNVGRGGSNGTTSSANNDSSMTMTEHQVAKLMEEDMGSAMQYLQGKGLCLMPISLATAISTSTCHSMKPNNPLLLAGGSAINGVGETGGGPSSPTLSASTVQSATMGNGGT

>Sl12G042400

MVRLPSKFKPPKPKTLISLLTSQTQKSLHHTFSPSLLQTHKLFNHFTQILSHAITSGYFTNPFISSQLLQHCLTNSDFTLTFSHTLFTQIPNHNVFSWNFLIRAYCHSSFPQEAVSLYNQMMRESLSLDNFTFPFVLKACGRLVSFHKGREVHCVSLKMGFEFDVFVQNSLIYMYFQCGVVEFAQKVFDFIPDSVRDVVTWNSMISGYLQGGCCYQAVKVFGKLLRESDVRLNDVSIVSALTACARTELLDVGKRIHGLIVVYGVVMDVFLGSSLVDMYTKCGCLEDAKTVFDQLIDKNIVCWTSMIAGYLKCDMCKEAIELFREMLIAGVMAEPATIACVVSACGHSGALDLGRWVHNYSERSGIEMNLIVKNSLIDMYSKCGLVDKALEVFHSLTNKDVYSWSMMISGLAMNARSKEALQLFSLMEGSSEVSPNEVTFLGILSACSHGGFVDEGFYFFEKMTTRYKIFPGIEHYGCLVDLLGRANLLVEAMDLIGSMSIQRDAVIWRSLLFACSRHGNVELAEVAAKKINELEPEKCGAHVLLSNAYASASRWKDVKMVRRNMGAEGIQKQPGCSFIEIDGFVHEFLVADRAHCQIDIICDTLLAMNVVLKSRAPELVFLDFFH

>Sl12G088790

MVAMREMIFRIAAMQPIQIDPSSVKAPKRRNVKISSDPQSVAARHRRERISEKIRILQRLVPGGTKMDTASMLDEAIHYMKFLKKQVQSLEKVEINRPMSMSLSGNYLPNYNYPYHQSVQP

>Sl12G096510

MQATITSVIPTPQSPLSHQTSNVNTLRGLESCSSMSKLKQFHAQIVKHGLSHDNDAMGRVIKFCAISELGDLSYALQVFDKLPEPDPFIYNTVIRGLLKSQMLRDSLLFYLCMLESSVRPNSFTFPPLIRVCCIDNAVEEGKQIHGHVVKFGFGFDRFSQNNLIHMYVSFRCLEEAKRVFDNMCERDDVSLTTLISGYAQLGYVDEAFQVFDSIREKSCVIWNAMISAYVQSNRFHEAFALFERMGCEGVVVDKFLAASMLSACTRLGALKQGEWIVEHVKKSGIELDSKLSATIVDMYCKCGCLDKAVEFFQGLSRKGISSWNCMIGGLAMHGEGEAAIEVLKEMESERVAPDYITFVNLLSACAHSGLVEEGKHYFRYMKETYGIEPGMEHYGCLVDLLGRAGLLVEARRVVNDMPMSPDVGVLGAFVGACRIHKNVELGEKIGKQVIELEPQNSGRYVLLANLYANAGRWEDVASIRKLMNDRGVKKVPGFSIVELEGVVNEFIAGGRTHPQAEEIYVKANEMLECIKSAGYSPDSDGMQYDIDEEETENPLNYHSEKLAIAFGLLKSKPGDILRITKNLRVCKDCHQASKFISKVYNLEIIVRDRNRFHHFKGGECSCKDYW

>Sl12G096680

MKTLLRFKENVISVHGLVISGPLLPLLFQHLRCCSGATIQWKNERVPIKCSDLLDEFTTYCYARDLPRAMNALNALQIQKIWADAVTYSELIKCCLARGAVEHGKRVHQHVFSNGYEPKTFLVNTLMNMYVKFNMLEEAQALFDQMSERNVVSWTTMIAAYSSAKINNKALEFLIFMMRDGVKPNMFTYSSVLRACDDLSNLRQLHCSLLKVGLESDVFVRSALIDVYSKMGQLECAMCTFNEMVTGDLVVWNSIIGGFAQNSDGDEALTLFKRMKRAGFSADQSTLTSALRACTSLALLEVGRQVHVHVLKFKRDLILDNALLDMYCKCGNLEDAHQIFSQMVEKDVISWSTMIIGYAQNGFSRKALELFKEMKVSGIRPNYITVLGVLFACSHAGLVEDGQFYFHSMKKLFGIDPGREHYGCMVDLLGRSGKLDEAVKLIHEMGCEPDAVTWRTLLGACRVHRNMDLAEYAAKQIIKLDPSDAGTYILLSNIYARTQKWEDVMDLRRSMKERGVKKEPGCSWIEVNKQIHAFIMGDNSHPQKEEINKELNQIIWRLKEVGYVPDTNFVLQDLEDEQMEDSLLYHSEKIAVAFGILSLSREKTIRIRKNLRICGDCHLFVKLLAQIEHRSIVIRDPIRYHHFQDGICSCGDYW

>Sm00001G02690

MDPFQGDVFGEFAHDDESLNQLVSSGSFCCAFSQDHHQHHHQCPSFSLDAASSTCDHLLSPSDTDSFGIPAKNHSPGSASENLSIISSSCGGASCFLEFGGAGAGAGSTDSSPASSIVEQAAAYTTAAGFDHPPPPPPPPVDLHGAASLLESWHFLQYESLPGLLHIPEPVDHFSKLSGSSSSPPSFDLLPGGTASSHDDVIILGSLLDHHHPHSRAHEVMIQQQENAPRAGDDRHVPEQELVPGARHAPCSDDDHTDIDMIDRRELCSSKEQPKLMEVEASPPMAASHQSNKRQRGCFTLQNVSESLQHHSVIFHNVGVAAKLSEAMLLQINSSVARNPSPVSPVAASARNPSSTITSSSATTTTSTGSGGGGHGSFSYRFKETVEPQSIAARQRRKKISERVRELEKLVPGGNKLDTASMLDEAIRFVKFLQIQVQLLEAVGNGGHGNYGANNHCLNVPLNHQARMFPFSSSPSSSIAISSPCSSSSSSSSSSCASSSCNLASMAIATSSVVAMSPASIADSVLLLGSRRSSSASSSSPLVLTETLQHALFKHKRCVATIQQCPVEHLLL

>Sm00002G08130

MDDCCSSLLPPASARAHHHSHSSPDQDQASSNNSLSDISSSFFDQDYDWASLQQPQRSNADSSTELLNSSTLISQSQQTLDQFGGFNLNFLIASSRPYGQQYETAVPCFQMVSPAMQVKPLKSLKRESLDQSTLVNCEDNSPGSARSGVFNPNCSELEQRQPFTAASDLISNKRQCLDQRLQGWSSPTNESKHKLSSVNFAGPALNTNGKPRAKRGSATDPQSVYARHRRERINERLKTLQHLVPNGAKVDIVTMLEEAIHYVKFLQLQVNMLSSDEYWTYAPTTYNGPETPLGLKLPLTGS

>Sm00011G04120

MGIVCEPLFFGGDLTRRSCRRRQEVACRSSKGDCDPGGGTGDSCLTSDHEAAAAAAAEDCSTELETAPIRGSCSVNDSSAGSSGGSSSSWTWQSNSPGRIVENVLEQVQSKASSPSSSPAAAEAAVQGFSEGERAVENWHDLLQATDLRDDGIETLPSLVSAAFPSDQEQHKARFPSWDNRPSKLSSHHQADYGCFMQQDGAAFGSSSNVFRPPPQFVDSHNLSLSSQAMLFERQHQHQQEFNSFSSHVSYPPGGTFNCHDGVAATTGNSRSNLELLLGSSAIDFKQPQQEEQQITSQCQGAFGGSWSANGNLGINGQAARRNHTGDGGAGQSQAQQLQSIFLPGQGSSRKPGIKTVEEMSRRCSLQDNNRDAKRHKGSTAQRTRFGASSKRGNSHKREPALNTNLKPRAKQGCANDPQSIAARQRRERISDRLKILQELIPNGSKVDLVTMLEKAINYVKFLQLQVKVLMNDEYWPPKGDGEEDYPVSQKYLFLSKIS

>Sm00018G02390

MGVVERSREVMAVMALHEALRNVCISSDWTYSVFWTIRPRPRLRGGLGCKAGDEGSLMLMWEDGYCRPAGCCSGDAAADNKNEEDPVRKAFRKMSIQLYNYGEGLMGKVASDKCHKWVFRESSETEVGPLNYWQSSFDSHPPEWMDQFSAGIQTIAVIQAGHGLLQLGSCKIVAEDLHFVLRMRYTFESLENQAGGGFLSQTFAPMR

>Sm00020G01060

MATAAASWVVEHCTQAAFEDEHKTRICKVPDPSCSCSTSREGSLDSEICFSSPPPQHQDVVGGSSNHFLDFGIDHHQQHHPHLINLLSDHEWINFLRWRSWQIESSSSNSAAAMPVEFPPSATDFSQLELFPSNLEPGLSLPLTSFSSHHSHQSSTVLEQFPQCLEIPSAATPLSSCPTLYDEAGYQDPFLAAFSSSTLHHAVDPNRRIDNVDTNHALVASNHPGIMTPCSTDQGDKGIISGKKAKPSSSSTSVSNESVNRSSVVDGPRNPRKRRALEDRQDAASSKQQKKIGSSSSSPAWAIDAAVALGSQEPIIHISHGGRTTPHDHELILQQIAAASSSSGKAPRVPALNTNFKPRARQGSANDPQSIAARHRRERISDRLKILQELVPNSTKVDLVTMLEKAINYVKFLQLQVKVLTSDDYWPSGATWQNSSKADTAL

>Sm00030G01020

VNHLLQHTLRSICTESQWVYAVLWRILPRNYPPPKILTWEDGFCDFAACVKSFKQSSSGDSEAASAEPLQPELFFKMSHEVYSYGEGLMGKVASDSSHKWIFRDPADQDSSLVSPWHGSLDPHPRTWEAQFKAGIQTIAIVAVQEGLLQLGSLDKVLEDLNLVILLQRKFTYLQTIPGVFAPHPNTLNHSIHQAAMIAGSSGRKIPERPPLSKSLNTGHSSPGVVPHLHPSMSSLHALLSKLPSVSDNALVPPP

>Sm00038G00990

MDVVRRTRSSSSPSFTSITNSSSSRISSCDPGDFSFLDYSQVCCRSLDLLSPDLPPPLVEDSSPVNSHGYKEYSSIAPKSLYHPLEFNQSFQAQLELQQTLMGDPSFGNSSLLTPLDDHSQDQRNPEDFHGQELAYYCQVEPLDEKYLSLMEHPPLYVNYLTDANRRMPPLGSSQALDQPGAPRIISNSSSSPTRKRSADDQNTTNAFSKREKIDSSPASSCCTTALNTNLKPRSRQGTANDPQSIAARQRRERISQRLKILQDLVPNGSKVDLVTMLEKAINYVKFMQLQLQASVVD

>Sm00042G01080

MEHEDFAGSQFCSIWESFSSDECGSGASSLNLPELCNSPDHNDNGSWLPPPPSSQIFHSKPATALALDNHHHHHQNLDRIIDSFPSPSTSSSHHHTLISSPDSPLVFNFLLPQHQHTSSSIQFGFPPVQLLQSPEEVIPSKPCVRTPPPKSSKASSSSSLNQNSSSSSMAVLENHCARIKKERAEVQDENCPAWKKKCLESRGLESRTKSSVTLSAPEESSTVTSALGPALNTDGKPRAKRGSATDPQSIYARQRRERINERLRALQGLVPNGAKVDIVTMLEEAINYVKFLQLQLLSSDEYWMYAPTNYNGMNISLGMHLNMQSYLKIKENVSS

>Sm00068G01040

MAKAIVPGLQERLRSLVNGKGWDYVVYWKLNDDQRFIDWVGCCCGGVTAGHGIGADFFSPQPHYLDGGAPCPDISFPHPSTKTCTILSAMTLSPSIPLDSAGVHAQVLLSGQPRWINFSLESNQHEGVQTKVYIPIQNGIVELGSSSQIAENAMVIQSVKAKCGDPWQDFQGFQENVDQQGFKSLYKNQEDGLDKHFNGHGDQVHWTSALDTHPWEHDVGFSDNQFSLNIAAPTQLNFLGQPSKTGGQPSDHFDKQVDCSRPEKQGPPFVQGLQDVPPLAPPNHSFSESPHGSGVSKENSEVKQETRADSSDCSDQVDEDDEKATGRSGRRHLSKNLVAERKRRKKLNERLYSLRALVPKITKMDRASILGDAIEYVKELQQQVKELQDELEDDSQAANNIPTMTDVCGGGHKHPGSEGITIADVDTNKCALKADDINDKKVEDLTQPMQVEVSKMDAHLLTLRIFCEKRPGVFVKLMQALDALGLDVLHANITTFRGLVLNVFNAEMRDKELMQAEQVKETLLEMTSQPEARSSSSLVSESHMGVITIDGGSGSDCSCKSVEYNNVANFTILEDGYNFKPGVSRQLLDCKDHNKRNAIARWLNMEYGRHEIEDNNPTYSYFVEDLQCFEITSFNGWGHFRTTNGWYGWSFWAFTGSFIQSSNHLSVDFL

>Sm00073G01190

MLVAASFQNQVGGGGAASTSQAGSRNHQGLAATLDHSDNTGQVSSWSHHFVPGNALSFAKLGDGGASSDPNSVKKRYRDEELQQMYGAAGQMQIDQQQKHQQDQQQRSQAPLAYDSNAPVAPVNPPPAGISARPRVRARRGQATDPHSIAERLRREKIAERMKALQELVPNANKTDKASMLDEIIDYVKFLQLQVKVLSMSRLGGAGATMAPLVADLPLEGAGQELVSSSQLCRQISVNLSPQDGIALTEHQVARLMEDDMGSAMQYLQSKGLCLMPISLATSISSCSTKVPTTMSLAATAAAAAAAGSSPTTAAIAGALQHATLSDSEASAMEGCNGRAATKDFPEQSEEQWLK

>Zm01G02730

MATPLLLPSRAAHAASATTCASQHLKATTSKEPPPRARPKHGEGGAPRPVPVPSSAPRPKSLVLSHIAAGRMDEAADAFAGVSSPGAFLHNVMIRGFADAGLPLDALAAYRAMLAASARPDRFTFPVVVKCCTRAGALGEGRAAHAAVIKLGLGADMYTANSLVALYAKLGLVGDAERVFDGMPARDIVSWNTMVDGYVSNGMGALALACFRDMNDALRVRHDGVGVIAALAACCLESALAQGREIHGYAIRHGLEQDVKVGTSLVDMYCKCGNVLYAENVFATMLLRTVVTWNCLIGGYALNERPVDAFDCFMQMRAEGFQVEAVTAINLLAACGQTESSLYGRSVHAYVVRRHFLPHVVLETALLEMYGNVGKVESSEKIFGQMTEKTVVTWNNMIAAYIYMEMYQEAIALFLELLNQPLYPDYFTMTTVVPAFVLLGSLRQCRQMHSYIIKLGYGDSTLIMNAVMHMYARCGDIVASREIFDKMPGKDVISWNTIIIGYAIHGQGKTALEMFDEMKCNGIEPNESTFVSVLTACSVSGLETEGWKEFNSMQHEYGMVPQIEHYGCMTDLLGRADPFASGKTVLPNKHSVRLAVAFGLISSEPGTPILVKKNVRVCNHCHRALKLISKYSRRKIVVGDTKIYHIFSDGSCCCGDYW

>Zm01G07720

MATVWSGCSTSCGSFGQELPPRGKRGGGEGTRFLPRRRGRIRGDARQTQALSAAMVTVGACVCRAAPCLLDSVVDGTEDVSVGFGGIDGEPLADGYDGGKRRGPRRRPTRPTVFQKEVGARTAPPAMASEPDSKSEHGVSRLHFLEERDEETLSRRLIKLSQNNKVRSATELFDSMRASGLQPSAHACNSLLACYVRRSPLADAMRMFEFMKAKGMATGHTYTMILKAVASNQGYVSALEMFDKIEEEEGSKKIIDVIVYNTMISVCGRAKEWMLVERLWRRLEQHSLNGTLLTYDLLVSIFVQCGQSELAIAAYQEMLQKGLDPSEDIMKAIIASCTKEGKWEFAMSTFSRMLRAGMKPSLILFNSIINSLGKAGQDELAFRMYHLLEKSGLKPDQYTWSALLLALYRSGRCWDCLKLFHGIKGKHPTLLNDHLYNTALMSCEKLGQWEHGLQLLWLMEKRGLKLSVVSYNHVIGACEVASKPKVALKVYQRMINQRCSPDTFTHLSVIRACIWGSLWTEVEDILEEVAPDSSIYNAVIHGLCLRGKTGLANKVYAKMRSMGLVPDGKTRAFMLQHIATD

>Zm01G18510

MRMLAIRYLHRTVAAKADTIMALKLCRRLSGFIVGANSRCCRLSDVDKFALLFQNCTDVRSLKKLHARVLTLGLGRDVILGSEILICYASLGVLCKTRLCFQGFLNNDLAEWNSVMVDIFRAGYPEEAILLYRGLKLRQIDLDEKTVTFGLKSCIELRNLLLGKGMHADSVKLGLSRDKFVGSSLVGLYSKLARMDDSQKAFEEILDKDIVSYTSMITGYSENMDSTSWNAFKIVSDMSWSNLEVNRVTLVSLLQVAGNLGAIREGKSVHCYSIRRDIGISDEVLETSLVHMYMQCGACQLASAVLKNSAQSVASWNAMLAGLVRTGQSGNAIHYLYIMLYEHKVVPDSVTYANVISACAELCNSGYAASVHAYIIRRSIPLDVVLATALIKVYLKCTRITISKRLFNQLVVKDTVSYNAMIYGYLQNGMVNEAIALLKEMVTECVAPNFVTILSLLAAIADHKDFARGRWIHGFSIRHGFCSNVDIANQIIRMYSGCGKIASARIVFASFENKNLISWTTMMMGCLFCGHGGQTVELFQLLMQQHDNKPDSIAVMTAIQAVSEFGHLKGVKQVHCFVYRALLEKDTKTMNSLITAYAKCGRLDLSVSLFLSLEHRDLDSWNSMISAYGMHGFYTKVLEMFKLMEEGNINPDGLTFSSVLSACSHAGLIKEGLHIFQSMTSMYSVRPQEEHYGCFVDLMSRAGHLEEGYKFIKLSTLNDKSSVLCALLSACRTYGNTMLGQVISNELLEVGQQNPGTYALISEVFAQKGQWNKSASIRNRAKENGLRKLPGSSLIESVEQVSKVC

>Zm01G21810

MPVPGEPSPPASPRQYSGLLAALHSCISGGNAPAAVSLLPTLARAGLRAPFPLLSSLAGLLLLRPAAPSFPSLAGRLLLYVRLAGLKHLVPCSTQLADQLLSLNFLLGRPRDARRLFTKMPRPSVHSYNAMLAGYARLSLAAPASEVFSSMPHRELVSYNAVMLALGRGGEAQGAVELYSELRNMYPSLGYNYHTFLALLVAGAQLMDRELARQFHCHLVVLGFLSDVKIASSLLDVYRKCHCVDDARKLFGEMHIKDVRMWTTVVCACAEDGQLATARQLFDQMPERNAFTWTALIEGYACHGQPMDALNMFQHMMKEGLQPDQITLGCCLSACAAMRSLNLGQQIHGMLLRCGFDRSVMIVTAIIDMYSKCSFLSGAIKVFSLTGKERKNAVLWNGMLSALYDHGHEQDVIGLFVQMIHGRHKPDANTFFLVLVACCHCGLVEEGMNFFDLMNERYRIVPGQEHYFCMADLVSHASSNNKVIEWIKSSPFSFNLRVWEVLVRNCTIHGNIELLNKVEEELARLDHPE

>Zm01G32730

MNSLFKRLPRKRVAATRRSSRRVHAQPRPHPSLTTFSRLCVEGPLPAALALLSELAAAGLRADPVSLTRLVKLCVRHGTADHGRLIHRHVEAHGPLPHDGAGGLFVSNSLASMYAKFGLLDDALRMFDGMPVRNVVTWTTVVAALASADGRKQEALRFLVAMRRDGVAPNAYTFSSVLGACTTPGMLTAVHASTVKAGLDSDVFVRSSLIDAYVKLGDLDGGRRVFDEMVTRDLVVWNSIIAGFAQSGDGVGAIELFMRMKDAGFSSNQGTLTSVLRACTGMVMLEAGRQVHAHVLKYDRDLILHNALLDMYCKCGSLEDADALFHRMPQRDVISWSTMVSGLAQNGKSVEALRVFDLMKSQGVAPNHVTMVGVLFACSHAGLVEDGWHYFRSMKRLFGIQPEREHHNCMVDLLGRAGKLDEAVEFIHGMSLEPDSVIWRTLLGACRMHKNASLAAYAAREILKLEPDDQGARVLLSNTYADLRQWTDAEKPWKAMRDRGMRKEPGRSWIELEKRVHVFIAGDLSHPCSDTIIQELNRLIGRIKSLGYVPQTEFVLQDLPTEQKEDLLKYHSEKMAIVFGTMHAVDGKPIRIMKNLRICGDCHAFAKLVSKSEGRVIVIRDPVRFHHFQDGACSCGDYW

>Zm01G50590

MESVLTSHPLTQSFPPSSDAPSHLSPSPAFSPNSASTPTFSQASEPSHSGVSSQVGRSKFQRWSDGGGGPFSGEPGSYKDATLIQLSAVPSSPSRGSQLRSAKEVVVRGHVEDLRQAVSGRRQVGLVRHSLPPSPRRQEVDGWQTVERRRGRRFEQAPRRPVDLRGKCFNCLSSNHFVAVCRRPTRCFRCRLLGHQARRCPSLLPVEKGTSGKPPRVQGGRVQVWQRLGPPVPSSVLNHRMSVWQRIGPQLPPSKFSHSPAAEASLVESNISSHSGKKKRRRAKRHKEVSWQSTVPVGAASTLEPSPGVGTQSSVKTCMLEFSTAMAREEAALRRALFVSIVGSRPEISGAEVRDEVARAFGIDVNSISIHQSLPEDFLLLLPDESSADRVFSGGKLFRGPLFNLQFRRWSRLAHAEAARLPALVEVEVHGIPAHAWDRSTVEYLLRDSCIITDIHPSTSLKNDLSSFRLRAWCLNTDLFPRPMKLLIVEPGTDVQEKRCLSYDIEVAVTAVMVPTNFDPPPAHSPAADGRDRHEDGGGRHSDDSNSPRPDGGRLQRPIFQRLGPRGQSTSGGRGAAVQRCALSAAGPAAGIEVVASPTRLLDGSPLSLTIGAGAEEDLAAGIAEVRRLHCPLPLLLVAAGVEEVPPPVRRLAVNSVAGVEEVASSMGRAVHSPSWLAAGTGTEAHSGRMLAADIGTEAHSGGMLAVSSGAVAHTGSMVAVPAREVLAVQMELTAADMGTGPGTVCAKVGPVGCAVDAPVVQQPQMDGHGGPMWTRLAGSLKSLGSGSPRQRFVGLQRLTKTVSPRNSFNVGIELVPRLTSTQSLVGTSEADRNSPAGMSLDAQVLESPRLKVYSRLKKGRLQIPAPVSSQMEVSAPVLDLVPAGVSSQLEEATPVMKLVANITRDIDCLLPQPPIQKRRTKQLPPDFVPRRSSRLSKKREGLNSTVRQVQAELMLKLNVTNSQVAVTDEMLEEFGQLFDKPLSNSHIKALAALFGWSVPDNVQECLDVPLLRELSIETEIVRMFHPPIRKSEEAIATIAPRYTHSVRVLDERFIRILKIFKWGPDAEKALEVLMLRVDHWLVREVMKTDVGVNVKMQFFRWAAKRRNYEHDTSTYMALIHCLEVVEQYGEMWKMIQEMVRNPICVVTPTELSDVVRMLGNAKMVRQAITIFYQIKTRKCQPIAQAYNSMIIMLMHEGQYEKVHQLYNEMSTEGHCFPDTVTYSALISAFCKLGRRDSAIQLLNEMKEIGMQPTTKIYTMLIALFFKFNDAHGALSLFEEMRHQYCRPDVFTYTELIRGLGKAGRIDEAYHFFCEMQREGCRPDTVFMNNMINFLGKAGRLDDAMKLFQEMETLRCIPSVVTYNTIIKALFESKSRASEVPSWFERMKESGISPSSFTYSILIDGFCKTNRMEKAMMLLEEMDEKGFPPCPAAYCSLIDALGKAKRYDLACELFQELKENCGSSSARVYAVMIKHLGKAGRLDDAINMFDEMNKLGCAPDVYAYNALMSGLARTGMLDEALSTMRRMQEHGCIPDINSYNIILNGLAKTGGPHRAMEMLSNMKQSTVRPDVVSYNTVLGALSHAGMFEEASKLMKEMNTLGFEYDLITYSSILEAIGKVDHEYTDQG

>Zm02G09570

MAARSLWQLEDAIIARLRTCATFRDLLRAHGLAVRLCLSQSSYVATQIVHICNAHGRAAHAARVFAQIPAPNLHLHNAMIKAYAQNHLHRDAVEVYVRMLRCLPHPSTAGFSVGDRFTYPFLLKACGALAASRLGRQVHAHEEDARVQHD

>Zm02G24090

MAVGDALRRLCEEIGWSYAVFWKAIGAADPVHLVREDGYCGHTSCPVGSEPSESLPLPSDAGCSVPAADTICWLVNNDMASQVHVVGQGIVGRAAFSGNHQWIVHGTANGHGLSSEVAAEVNNQFRVGIKTIAIIPVLPRGVLQLGSTGLVMENTNFVMFAKKLCSQLNNRSSMAASASVKNTSSQHGQSRPDTATVSTSTPPNALLSASLLKVAQQNGHPVREHSVYAERDLRFIQQASICESQFGSNLQSGVMNSCLTSPSFASVKKQSLLMNSIGQLQFSNNVHSSADLAREIILKSLGRQDSSVCENTNMNMHRGRYVVSNDINGSRNFDFLPIGAKESRANLSIPASSQILDHMPGTLQQKQSLVPFRVTQSSEFSKKVENPESGSFQIPSVTTSDSGGQVSNSFNMVQENRLSMSNNVRQVQKIDRFSDPSANVSTQRMKNRDECEPGMPIERASSFLVEPGADNDLFDMFGSEFHQSSHNVGADLVTWSGTESQNSDRDVPESSICFNSSPLFSSLDNELHYSGILSLTDTDQLLDAVISNVNPSGKQCPDNSASCKTALMDVPSTSHLCSIDLKRCESPSVPSLLIKHELAQFAKQPCLVDKSEDGCLSQNNGMHKSQIRLWIESGQNMKCESASASNSKGLDVPSKANRKRSRPGESPKPRPKDRQLIQDRIKELRELVPNGAKCSIDGLLEKTVKHMLFLQSVTKNADKLKDSTESKILGSENGPLWKDYFEGGATWAFDVGSQSMTCPIIVEDLDRPRQMLVEMLCEDRGIFLEIADFIKGLGLTILRGVMETRKSKIWARFTVEVIQLNLKISILSVQNLKQ

>Zm02G30550

MQPTTREMQAMAAAGQQISLDDLRAAAGGVHDDFLDQMLGGLPPSAWPELSSAAGGKAPDGGAQAEQMQHQPQHLGGGGGLYDDSALLASRLRQHQISGGGGGEAVKQMVLQQLADLRQEHHVLLQGMGRSTSTGGSRDGGLLLPLSLGSGGSGGDVQALLKAVADSAGGEAAGVFGGSFAGSLHHHQQQQQHFQSHPQQTAPLPGQGFGGGAGASGGVSQPQAGAACGGAAAPPRQRVRARRGQATDPHSIAERLRRERIAERMKALQELVPNANKTDKASMLDEIVDYVKFLQLQVKVLSMSRLGGAAAVAPLVADMSSEGRGGVAVAAGSDDGLAVTEQQVAKLMEEDMGTAMQYLQGKGLCLMPVSLASAISSATCHMRPPVGGPGLGVAAAAAHHMAAMRLPPHAMNGGAGAGADAVPASPSMSVLTAQSAMAPNGAGGGTDGEGSHSQQQQRHHPKDAASVSKP

>Zm02G38040

MEIDMMAQFLGAADDHCFTYEYEHVDESMEAIAALFLPTLDTDSANFSSSCFNYAVPPQCWPQPDHSSSVTSLLDPAENFEFPVRDPLPPSGFDPHCAVAYLTEDSSPLHGKRSSVIEEEAANAAPAAKKRKAGASATTKGSKKSRKASKKDNIGDADDDGGYACVDTQSSSSCTSEDGNFEGNTNSSSKKTCARASRGAATEPQSLYARKRRERINERLRILQNLVPNGTKVDISTMLEEAAQYVKFLQLQIKLLSCDDTWMYAPIAYNGINIGNVDLNIYSLQK

>Zm03G14980

MEDGGLVSEAGAWAELGTGGDESEELVAQLLGAFFRSHGEEGRHQLLWSDDQASSDDVHGDGSLAVPLAYDGCCGYLSYSGSNSDELPLGSSSRAAPAGGPPEELLGAAETEYLNNVAAADHPFFKWCGNGEGLDGPTSVVGTLGLGSGRKRARKKSGDEDEDPSTAIASGSGPTSCCTTSDSDSNASPLESADAGARRPKGNENARAAGRGAAAATTTTAEPQSIYARKRRERINERLKVLQSLVPNGTKVDMSTMLEEAVHYVKFLQLQIRLLSSDDTWMYAPIAYNGMGIGIDLRMHGQDR

>Zm03G15560

MEDGGLVSEAGAWAELGTGGDESEELVAQLLGAFFRSHGEEGRHQLLWSDDQASSDDVHGDGSLAVPLAYDGCCGYLSYSGSNSDELPLGSSSRAAPAGGPPEELLGAAETEYLNNVAAADHPFFKWCGNGEGLDGPTSVVGTLGLGSGRKRARKKSGDEDEDPSTAIASGSGPTSCCTTSDSDSNASPLESADAGARRPKGNENARAAGRGAAAATTTTAEPQSIYARKRRERINERLKVLQSLVPNGTKVDMSTMLEEAVHYVKFLQLQIRLILCEASPQLTDSRNGGDTSNEAHKRNSTDGNNKQTCELDAFIYKSLEGIQDQEFCNIIELLSVRPPIDAKLKLMKEIAEEHEIGPTYFNGSTLPLPKEKHDETVAASATKLPDEDYESNTGLDSLDLPEVPKAAICPPSLNHIFNLSFSFSVVIFLAVDVLLDNIRSIDRAEEFAFRAEEDAVWSQVAKAQLREGLVSEAIESFIRADDAAHFLDVIRAAEEANVYNDLVKLTDSMPNVADLQNVGDRLYDEELYEAAKIIYAFISNWAKLAVTLVKLKQYQGAVDAARKANSAKTWKEVDDLEEVSEYYQNRGCFSELIALMESGLGLERAHMGIFTELGVLYARYRSEKLMEHIKLFSTRLNIPKLIRACDEQQHWKELTYLYIQYDEFDNAATTIMNHSSDAWDHMQFKDVCVKVSNVELYYKAVQFYLQEHPDLINDMLNVLALRLDHTRVVDIMRKEDYERLRESVDTHDNFDQIGLAQKLQKHELLEMRRIAAYIYKKADRWKQSIALSKKDNMYKDCMETCSQSGDRELSEDFLVYFIEQGKKECFVSCLFICYDLIRPDVALELAWMNNMVDFAFPYLLQFIREYTSKVDDLVKDKIESQKEERAKEKEEKDLVAQ

>Zm03G19680

MAVGDALRRLCEEVGWSYAVFWKAIGAADPVHLVWEDGSCGHASCYTGSDAPEAGCEPGTSVCTLVGKVMASQIHVIGEGTIGRAAFTGNHQWIVHDPATASDHNLRPEVVAEMNHQFAAGIQTIAIIPVLPRGVLQLGATSVVVENTNLVLQYKKLCSQLNNRSSMASSSSVKNDLNPKVQSHPLNGPTSIYPADTRSKFFSGSPMSYQQCYGPDGTTVSSSILTNMGSNASMLMVAQRNAQVGKEHILYAPDMRFRLQNPYCDRRVESNTQSSVVSSGFISSISASMERHPWMTNSQQLEQGNTGEPSDLRNVLLKSLAYRHPLFNDNTNMVLVHGKGQVPGFVNGHGGFDFLPEGTMVFKGNLYGSTANQILDQRCSSTSEMSGHRPTISYKMPQSTQCVRKTESPKRETCETSAALSSGSDIQVSGGLKTVISQENNMSSSDLISPKKPNEVHAPTGAIVKAVKNMDSRRLPDISNERSPPLRMDAAAESDLFDMFDSEVHHLCRNVDNDLTCKAAKPENSNRDVPESSVRLGSSSSFNSVDDKFPYSGIFSLTDTDQLLDAVVSNVNPGGKQISGDSTSCKTSAADIPTTSYCRLKEPKQSESSSPLLIKNELAVSNFVKQSSFQEKAEDGCLSQNNAIQKSQIRLWIENGQNMKCESASASNSKGVDMSSKASRKRSRPGENPKPRPKDRQLIQDRIKELRELVPNGAKCSIDALLEKTIKHMLFLQSVTKHADNLKDSNKSDILGGENGPLKDYFEGGATWALDVGSRSMTCPIIVEDLERPRQMLVEMLCEDRGIFLEIADFIKGLGLTILRGVMEARKNKIWARFTVEANRNVTRMEIFLSLMRMLEPSCGAGENSNNNIKMPLGIVQYPIIPATGHLR

>Zm03G27910

MDPYHYQTMYDPRGFPIIHPQPYLQHPVAGALGDSRVRGGGSGARRRPGAKLSTDPQSVAARERRHRISDRFRVLRSLVPGGSKMDTVSMLEQAIHYVKFLKTQISLHQAALMQHEEGCHAELAAYSAVAVVGDNEVTLASHGRTGACDEMMQLQVAAEEALSYGVDAHQPYGLDPRQLSGGHELPPLPASCIFLEEPADACYSVCDLDDGDTGLPGSY

>Zm03G29530

MVGSGADGGGGGDDARSKEAAGMMALHEALRNVCLDSDWTYSVFWTIRPRPRCRGGNGCMVGDDNGSLMLMWEDGFCRPRVAECLADMSGGEDPVRKAFIKMSIQLYNYGEGLMGKVASDKCHKWVFKEPSECEPNISNYWQSSFDALPPEWTDQFASGIQTIAVIQAGHGLLQLGSCKIIPEDLHFVLRMRHTFESLGYQSGFFLSQLFSSARGASPTPPPPAARPPPAQLFNWPGAHQPQLFPPGPGAFHHPSARAMPPLFPGGGKDEGHMFHIPPAHHGSKPLHMDELQHRHQQTMGAGGEAPDGELRWPNGLSFFTALTGRDDDAKLLFGGPGAADDEKAAPDAQTGHGGAGNVEEYLSLESHSDKAPSKVENGAQHSTKSKRSFTLPARVSSSTTTSPSVSASTAPQGMEYRGTHEGGVYSDLMETFLE

>Zm03G37780

MDHNLSDMTILSEHAWGDEAGAVLLSPPEVGSVIEGGGGMTVLERLVLDEALAAAILELQAIQQAAPGPASCAGKQQLAAGGGGGVVGAEAAVAFPSMMAMAMAAPTPAYADADVDAGVLQRQQHHHRPHAAMGMPSPDYDLAPAAAATPAVAVTAAAPPAFTTNTNAAVDSGCSGLVVLGRSPPPPVFSASNETAATASLDCEHGGGGRGGGRRQRRPNRKRKAAAEAERSRPVRVAAAAAQEESTLCSLLASTAPTTAGGIQIAFSTAAVHQQQHAAGGAGGAKTSAKPSSLSLSSSGSSSNISFDGRNPARNVRISSDPQTVAARQRRERISDRLRVLQKLVPGGAKMDTASMLDEAASYLRFLKSQVRDLQTLDRRNYGAASSNDAAATMAAVVGSSLSYNRGTGAMPAFTFPAAGETLRGAG

>Zm04G01700

MEANCASIWSEESETIAHLQSMFWSGSNADACLSSPNSSASSCVEPSTSPTALFLPLAENQRCDKEQCQNTGAVRCFGHKSQVLAPITNEVTANKRVCLMDENRKSTDAKRPCTVPPWVSRKNSIAPADEINTTEPVNKSCSWCCSSEDDSAGACEEPIVLKQSTSSRGPSRSSKDSQSLYAKRRRERINEKLRTLQQLIPNGTKVDMSTMLEEAVQYVKFLQLQIKLLLACKCEQEVHLE

>Zm04G24210

MAACLLHPYPPGLHSSRANPYSASRQPTHLKWELGRWRRHLLLCVAASAGILQRELIMLPSARTTQNPGPVNPRNQFDIMPERSTENTKSAANPLDESLGTGGQIASGRDPHEVGEKSESAVFVALAHAGRHAEVVELFRSMRREGVPVNRFVLPSVFRACADLWDSRMLRAVHGLVINCSLCQHVVVGTALAAAYIDLGLVDDASKVFDDISQPNVVSWSVIIGGYARSSQWDKAWDAFSAMQCSGLPPNGSVLVMAIQACGALGCLVRGKQAHTVAVVLGFERNATVWNCLIDMYGKCGSVESCRRVFDTTVSRDQVSWNTIISCYARLGLCEEALETIVRMQESGFTIDRFVLGSGVAACANLADIDSGRAFHGYLIRRALDTDVIQGSALVDMYGKCGYMDLARLVFDRMDERNDVSWDALLSGFVENGLVDSALDTFRQMESAKIKSNQHTFVNLLRLCGDRRYREYGRQIHGHAIRVINQMNVVLETELIDMYAKCGCIEVSQLLFLRMNERNLVSWNTLLSGYVGDGQPVATVNIYRQMELARVGPDHYTLAGLLNMCRFQGLLRYGRQIHAHLIKTGSEMNVVLQTLLVHMYFKCRRWRDAHNVCTLIRERNSHVYEAFFKVYGDDYLI

>Zm04G33360

MVRSRGHLALSTRLMNVGLASLCRGGWLDRAESVLVDAIRLGLPPDVVTYNTLLAAYCRVDGLDAGLAVVHRMREAGVRPDAVTYNSLITGADRSGLTVRALDLFDEMLRSGVAPDSWSYNALMHCLFRSGHPEHAYRVFADMADRGVAPCATTYNTLLDGLLKAGHVTNAFRMFRYLQRAGLPVGIVTYNTMINGLCKSGKVGYARMVLKELGRTEHAPNAVTYTTVMKCCFRYGRFEQGLETFLSLLEGGYISDAFPYTTVISALVKKGRMQEANTYCELMIQSGSTLDNACYNTLIHLRCQEGKINDAFELLNMMEEGGLESDEYTFSILVNGLCKMGQIEAAEKQIWSMEMMGMQSNVVAYNCLIDALCKCHEVDAAIKILHSMKLKDDFSYTSLVHGLCKVGRYHMASKFLRICLHEGNNVLASAKRAVISGLRSAGFKNDLRKVRSALYMARLLRS

>Zm04G34530

MAVGDALRRLCEEIGWSYAVFWKAIGAADPVHLVQEDGYCGHTSCPVGSEPSESLPSDAGCSVPAANTICSLVHSVMASQVHVVGQGTVGRAAFSGNHQWIVRGTANGHGLSSEVAAEMNNQFRVGIQTIAIIPVLPRGVLQLGSAGLVMENINFVMFAKKLCSQLNNRSSMAASASAKNALSQHGQSRPDTATVSTRTPPNALLNASLLKVAQQSGHPVREHVVYAEPDLRFRQQASFCENQFGSNLQSVVMNSSLTSPSLASVKKQPLLMNSIGQLQFSNYVHSSADLARDVILKSLVRQDSSVCETTNMNMHRGRYMVSNDINGSGDFDFLPVGAKASRANLSTSVSSQILDHTTRTLQQKQSLVPFKVTQSSEFNKKMENPENGSFQTPSVPTSESGGQVSNSFNMDQENQLIMSNNVRQVQKINRFSDPSANVSTQRMKNRDVCESGMPIERGSSLLVEPGADNDLFDMLGSEFHQFSHNVGSDLVTWSGTESQNSDRDGPESSIYLNSSPLFSSLDNELHYSGILSLTDTDQLLDAVISNVNPSSKQCPDDSASCKTALTDVPSTSHLGSIDLKRCESSGVPSVLIKHELAQLVKQPCIFDKSENCLSQNNGMHKSQIRLWIESGQNMKCESTSASNSKGLDTPSKANRKRSRPGESAKPRPKDRQLIQDRIKELRELVPNGAKCSIDGLLEKTVKHMLFLQSVTKNADKLKDSTESKILGSENGPLWKDYFEGGATWAFDVGSQSMTCPIIVEDLDRPRQMLVEMLCEDRGIFLEIADFIKGLGLTILRGVMEMRKSKIWARFTVEANRDVTRMEIFLSLVRLLEPNCDGSAAAENASNMNMPLGLVHQPVIPATGSIQ

>Zm05G31220

MDGFVDPFVPEQAWPQDAMFIGSSWPCAGAGATSLADPAGTYAYLGAQAAPGQGAGFHLQDGSSTALVPPLELHQQFLSAHLPGDDVTVTQEEGLGFEVNSALVAGMLGPVLSAPCAVSLADSAPVVCSSSNDSSGGSEQSGLPPPPRFLVLGEQPASWPSAFPRISSLAGEETSRSFGFGAVSDNDLVRGSSCVADVNKYPQIGNARLHVEDDVEFNTGKMLSFAPGLDFGDLQLSQKELSGLRHLNSGQQLSSFDATRNPEQSSNEASGGGTGLNAPPFMVPANGAAGNGAPKPRVRARRGQATDPHSIAERLRREKISDRMKNLQELVPNSNRTDKASMLDEIIEYVKFLQLQVKVLSMSRLGATEAVVPLLTQSQTENSGGGLLLSPRSGSGRQQQARGGSLPPPSSEVRDGAAFEQEVAQLMESDMTTAMQYLQSKGLCLMPVALASAISGQKGASSAAVQPENGGAKEMLRAVKPLASPIHGR

>Zm05G36270

MLLTVGECHCEDEARKVVDKMVNQVHVVGEGLIGRTLISGEHQWISDDIPFSFSQISDEDNLGLCQGYTWWQHQFLSGIKTIAVVPMPAFGVAQFGSMQKVSESLQFLDQVKGATFLMESLSWSPSTKDAQKDAFMYNPQFQLDSSTTREGLPRTKAEPENSRLENAISIDSLKNFAITSNGLNPHVVAMPVNSKSISTVKLIQSDSNLRYNNISENAQQFKSAKQPDSSWASAATCFSNLTNLQRIERGLSCTPNKLRHCLQSEKSSGFLDSYSSIVSTDAELKCTLFDNATPFVQSDVIQEVGTSGSTRACGLHELPNEIWAEIASGEMKPVIKEVNENNGFLESTAFDPVMNDWWGDTALLAGNTSHLSATTMNSVSGQASSEQLSIEDRGLFSESVFEELLGFDGNVGPVMDSTEPLAGFVSGCHLPRYNLQGSLSVCKAQVPPLILPSGSCTSENVVMGSSKEAPVSLQNLSMDDCGSLNTASSKISQVKKPEGEKVVKKRARPGESTRPRPKDRQQIQERVKELREIVPNSAKCSIDALLDRTIKHMLFLQSVTKYAEKIKQADEPNMISKDNGSVLNDNSHGVVLKDNPSGGSNGGATWAYEVAGQTMVCPIIVEDLAPPGQMLVEMLCEERGFFLEIADTIRGFGLTILKGLMELRDGKIMARFLVEANKNVTRMDIFLSLVQLLQQNSRNRSSDQLVKVMNNGIPSFVDHQQSPASIPVWPC

>Zm05G37360

MVGGLVLSQAQTHQAATPRTAAPAPAPAQALEKRPRDQAQPCGTVAAAAAVSVSAPSLDEVRKVHARHIKLGLDRCPRYARPLLAACAVSGWPGGMELAASIFESLVEPEAFDYNTLMRGHVSGGGGDRDPAAALRLYIDGLEAGVSPDSYTFPFVLKACAQLAALQEGRQLQAHIVKLGFQHDEHVQNSLISFYGKCGAPAMARLAFDRVEAEERTTVSWSALLAAYTKAGLWGECLASFGSMMLDGWRPDESSMVSALSACAHLGSFDAGRSVHCALLRNTARLNTIMLTSLLDMYAKCGSIEKAAAAFDAMDDKSAWAYSAMLSGLALHGDGRKALQVFDAMVREGHAPDAAAYVGVLNACSRAGLLEDGLRCFDRMRLEHKVAPNALHYGCMVDLMARAGRLDDARALIGTMPTGPTDTAWRSLLNACRIHGDLDLAWHALQELRRLGAANAGDYVIVADMHARAKNWTAAAALRTEAVDSGLAQSLGFSAVEVRGELHRFVSQDMAHPRTRDIYEMLHQMEWQLRFDGYKPDTSEVALDAGEEEKGRVVAAHSQKLAMAFGLLSTPEGAPVRVVTNLRMSKECHAYSALISEIFGRDIVIRDRNRLHRFRCGTCSCRDYW

>Zm05G38190

MALVREHGGYYGGFDSVEAAAFDTLGYGHGASLGFDASSALFGEGGYAAGGGDAWAGAGASTVLAFNRTTAAAAVGVEEEEEECDAWIDAMDEDDQSSGPAAAAPEARHALTASVGFDASTGCFTLTERASSSSGGAGRPFGLLFPSTSSSGGTPERTAPVRVPQKRTYLSAEPQAVSPNKKHCGAGRKASKAKLASTAPTKDPQSLAAKNRRERISERLRALQELVPNGTKVDLVTMLEKAISYVKFLQLQVKVLATDEFWPAQGGKAPEISQVREALDAILSSASQSQRGQLN

>Zm06G14150

MMATGFFPDEVTLSSVMSACAGLAAEREGRQVHAHMVKRDRLRDDMVLNNALVDMYAKCGRTWEARCIFDSMPSRSVVSETSILAGYAKSANVEDAQVVFSQMVEKNVIAWNVLIAAYAQNGEEEEAIRLFVQLKRDSIWPTHYTYGNVLNACGNIAVLQLGQQAHVHVLKEGFRFDFGPESDVFVGNSLVDMYLKTGSIDDGAKVFERMAARDNVSWNAMIVGYAQNGRAKDALHLFERMLCSNENPDSVTMIGVLSACGHSGLVDEGRRHFHFMTEDHGITPSRDHYTCMVDLLGRAGHLKEAEELIKDMPTEPDSVLWASLLGACRLHKNVELGERTAGRLFELDPENSGPYVLLSNMYAEMGKWADVFRVRRSMKDRGVSKQPGCSWIEIGSKMNVFLARDNRHPCRNEIHSTLRIIQMEMCRTSIDGIDDDLMNYCPEACG

>Zm06G29890

MEDFGGWNMQYAAELCLPSRRDDDDDGLLGDFLGGGGLDIRSDHGGLNLPSSHPAVQNLVPCHGGVSLSVSDAFMGLDTADMLASVTGALDCSLLDTFACGSDAVVVAEEPAAQPTASSNAFSGYSSTTGGNYGNISSGESNTYGGSGGGHDTEVASPTCASKRKLAGKYPAIAAASATATTKTAEGQVAAPRRGTKRDSATSSSSTSATVAGHGGGNHAAGGGGSSSVGDGNGYEPDSEAIAQVKEMIYRAAAMRPVHQLVCGAAGGEPPSSSQNKPRRRNVRISSDPQTVAARLRRERVSERLRVLQRLVPGGSRMDTASMLDEAAGYLKFLKSQVKALERANPSSNGGYNDSSFLPQSYTGCLGVDGGTSAAFGSDGAIGGYGYVKPGRNMHR

>Zm06G31830

MPLLPTKPAPALNPFLLPLASLALPRALPPIRLHAKKRLRGLSSAAVSAAASTSSSADPSTELRALCSHGQLAQALWLLESSAEPPDEDAYVALFRLCEWRRAVEPGLRACAHADDRHAWFGLRLGNAMLSMLVRFGETWHAWRVFAKMPERDVFSWNVMVGGYGKSGLLDEALDLYHRMMWAGVRPDVYTFPCVLRSCGGVPDWRMGREVHAHVLRFGFGEEVDVLNALMTMYAKCGDVMAARKVFDSMTVMDCISWNAMIAGHFENGECNAGLELFLTMLHDEVQPNLMTITSVTVASGLLSDVTFAKEMHGLAVKRGFAGDVAFCNSLIQMYASLGMMRQARTVFSRMDTRDAMTWTAMISGYEKNGFPDKALEVYALMEVNNVSPDDITIASALAACACLGSLDVGVKLHELAESKGFISYIVVTNAILEMYAKSKRIDKAIEVFKCMHEKDVVSWSSMIAGFCFNHRNFEALYYFRHMLADVKPNSVTFIAALAACAATGALRSGKEIHAHVLRCGIEYEGYLPNALIDLYVKCGQTGYAWAQFCAHGAKDVVSWNIMIAGFVAHGHGDTALSFFNQMVKIGECPDEVTFVALLCACSRGGMVSEGWELFHSMTEKYSIVPNLKHYACMVDLLSRAGQLTEAYNFINEMPITPDAAVWGALLNGCRIHRHVELGELAAKYVLALEPNDAGYHVLLCDLYADACLWDKLARVRKTMREKGLDHDSGCSWVEVKGVVHAFLTDDESHPQIREINTVLEGIYERMKASGYAPVESHCPEDEVLKDDIFCGHSERLAVAFGLINTTPGTSISVTKNQYTCQSCHRILKMISNIVRRDIIVRDSKQLHHFKDGSCSCGDEGFG

>Zm07G00060

MAMWLLMKEDVLCTCENGCHLQFNRKVKGLKAAHAQAVHHILVSAGMSTHIIQLLQGLCSDGLWRYGVFWSFKSETNGWILTWGDGYVNKMIEDWQMGDLSASPTVRKNQTIPSSWCKKSYPFCPIEAALMRMPSHLYPLGEGIIGKVALTGQHCWISANELCSTAMHKYQEDWQLQFAAGIKTVLLVPVVPHGVLQLGSLDTVFESAVLVALIKDMFHKICDASVSHASLSTGSAYSNNLSLPTATLSINHPDVNLIYSSAQIVNADHHSLTHPFSTSEIPILEDITIGSYRTIPTGWPNGLLGDNGTIGHEYFEGFSLTDMAHWNQENTHGSTIVVNDGAMLSNTSIHSEFHGDVMVMSREEHELFTWRCRLKQEPTSPALPQLNDNSADLYVQLETNNYAELLLDTIIYQTGHTSNIESAPSAESPISCKTQVKKDHSLRVDESPISDIPGRQDLSPISINEGFISCAMTDGRVEINKTIPEECLVENTHRINSAEIKRRCRKVELQRPRPRDRQLIQDRMKGLRELIPNASKCSIDALLDKTIAYMLFLQSVSEKAEKRRCPMPYFCFAVTLRQIQNTLEDKESHDETKKQLERCPLRVEELNQPGHLLIEMLCEDYEVFLEMAHVLKGLKVSILKGVLEYRSDKFCARFVIEGSDGFNQMQILCPLMHLLRRR

>Zm07G14220

MAGTGIRGGREGAGWRRAGGGDGTPAAALWWGVVRRLVAGRGEIAGSVFGGSFVGSLQQQQHFQSHHPQQQTAPPLPGQGFGGGGRGGGRAGAPGGMSQPQAGAAVGGAAAPPRQRVRARRGQATDPHSIAERLRRERIAERMKALQELVSNANKTDKASMLDEIIDYVKFLQLQVLSMSRLGGAARSRQGRDGAAAAAGSDGLAVTEQQVAKLMEEDMGTAMQYLQGKGLCLMPVSLAAAISSATCHMRPPVGGPGVAAGHHMAAMRLPHVTNGGAGADDVSASPSMSMLTTPSAMPNGAGAGVDGKDAASVSKP

>Zm07G25890

MEIDMMAQFLGADDDHCFTYDYEHDVDESMEAIAALLLPTLDTDSNSSSSCFNDDVPPQCWPQPGHSSSVTTLLGPAESFEFPVMDLSFPTSDFDPQSHCATPYLTEDLGSLQHGKHSPVMEEEAADVEPAAKKRKASATATATAKGSKKSRKASKKDCIVDDDDVYVDPQSSGSCTSEEGNFEGNTYSSAKKTCTRASRGGATDPQSLYARKRRERINERLRILQNLVPNGTKVDISTMLEEAAQYVKFLQLQIKLLSSDDMWMYAPIAYNGINISNVDLNIPALQKAVGIHRRPSMRASSCVDGDLSMPLRYATRMSRRVLVMVEAVTVHVTQKYFVSASKDNSWCFYDISTGSCLTHVGEASGQEGYTSASFHPDGLILGTGTTDAVVKIWDVKAQSNVAKFEGHVGPVTAMSFSENGYFLATAAHDGVKLWDLRKLRNFRTFSSYDLDTPTNTVEFDFSGNYLAIGLI

>Zm07G29800

MPLRRLPLPLPCPARATAKGKPHPLSTAHHPFAETPHPPPAGAATPLAALRGRLQAGLPVSPTAFSAAVARSGPDALPALHALAVISGLAAFAPVTNSLAARYAKGNSFAAAARVFAAAPSRDTSSYNTILSATPDPDDALAFAARMLRAGDVRPDAITFTVTLSLAAGRGEGRLVRQLHALVSRAGIAADVFVGNALVTAYARGASLDAARKVFEEMPARDLVSWNALVCGLAQDGECPAEVIRVFLRMLKHGGVRPDRISVCSVISACGGEGKLELGRQIHGFAVKLGIEGHVSIANVLVAMYYKCGTPGCARRLFEFMGERDVVSWTTVMSMDREDAVSLFNGMMRDGVAPNEVTFVAILSAMPGHCPAREGQMVHAVCIKTGLSDKPAAANSFITMYAKLRRMDDAKMIFGLMPHPEVIAWNALISGYAQNEMCQDALEAFLSMVKITKPSETTFASILSAVTAVETVSMAYGQMYHCQTLKLGLGASEYVSGALIDLYAKRGSLEESWKAFGETVHRSLIAWTAIISANSKHGNYDGVVSLFNDMARSGVTPDGVVLLSVLTACRYSGFASLGREIFESMATKHGAELWPEHYACVVDMLGRAGRLEEAEELMLQMPSGPSVSAMQSLLGACRIHGNTDVGERVAGVLLETEPTESGAYVLLSNIYAEKGDWGAVARVRRKMRGMGVKKEVGFSWVDAGGANDSLHLHKFSSDDTTHPQREEIYRVAEGLGWEMKFFKNCLQVEMEFLD

>Zm08G01090

MPPSPPLSPALHRLLSFLPRLASPRHLLQAHAFLLPRGGQRDPRLLSALLLASLCLVPHSQQTSDHAAALLRRVYPSVYLRDVARLPPHLLRRSLLGQLHSLLLRTGHVASDAHVSSSLVQLYCTCGHVADARSVFDEMAVRDVVAWNVMIAGYVKAGDQAHARELFDAMPERNVVSWTTVIGGYAQMKRPEKAVEVFRRMQVEGIEADGVALLSVLAACGDLGAVDLGEWVHRFVVRRGLCQEIPLMNSIIDMYMKCGCIEKAVEVFEGMEEKSVVTWTTLIAGFALHGLGLQAVEMFCRMERENMAPNAVTFLAILSACSHVGLTDLGRWYFNIMVSQYRIKPRVEHYGCMVDLLGRAGCLKEAQDLVKNMPLKANAAIWGALLAAARTHGDAGLGEQALLHLIELEPNNSGNYILLSNIFAEQERWGDVSKLRKAMRERGLRNVPGASSIEVDGMVHEFTSRDWSHPFIHTICEVLCEIDTNMKSVGFASVFPEVLHVIEEG

>Zm08G02120

MAQASKRSAFLRPVMYDDDEPSSISMSLELFGYHNGGVGVVVDDGDEAADGGSASALSLQLAFDDDSFVKGGCECDDAGGGGASAADYYYYGSWAGAYGGSGATSSGSNSSVLSFEQAGSGGGQRHHLVYGEDGCALWMDDAGAGMVVEHPPAQQQQQHGSGCDFGLIGSSPDDAGLPIQLEPGTVQLPAKAPPHKRARRDGDQVQAAAAAKKQCGGVGARMKSKQAKLAAPAPTKDPQSVAAKVRREKIAEKLKVLQDLVPNGTKVDLVTMLEKAITYVKFLQLQVKVLAADEFWPAQGGKAPDLSQVKDALDAILSSSQQCPK

>Zm08G22550

MDHNFSDMISSESAWSGGVGAGDGAGSGMTVLERLVLDEALAAAILELQGIQQAPACAGKEVVMVAEPTAGGGVVGEAAVAFPSMGMATPTPAYADVDADVLQRQQHHHRHHGAMGMPPDYDLTPATPAVTVTAVPPAFTSAAAYSGGNFVVGQSPAVFSTSNETTATAAAVTSTTSLNCEQGGGGGGGRRQRRANRKRKAAAERSPVAAAQESTLCSLLASTTTAGEGGIQIAFSNAAAAAKRSAKPSSLSSSGSSSISFDGRNPETNCGGVDDPMYEPDTEALAQVKEMIYRAAAMRPVSLGSEEDAGERPRRRNVRISSDPQTVAARQRRERISERLRVLQKLVPGGAKMDTASMLDEAASYLRFLQSQVRELQTLDRRNYGLTTADNNASRATATMAASGPLMSYNNGHGALPAFTFPAAAGETLY

>Zm08G33530

MGKVASDKCHKWVFKEPSECEPNISNYWQSSFDALPPEWTDQFASGIQTIAVIQAGHGLLQLGSCKIIPEDLHFVLRMRHMFESLGYQSGFFLSQLFSSSRGASPTPTPPFPLKQPPAAPAARAPPQLFSWPGHHQPQQLPSPAGASPLFPPGPGAFHPSSARPMPPFPGGKDEGHMFHLPPAAHHGTNKQPPQHMDEHHHHQQQPMVGAGGDAPDGDLRWPNGLSFFTALTGRADDAKLLFGGPGAGAGAADDDDKAAPDAHGGAENVEEYLSLESHSNKARKVESPAAQSTNKFKRSFTLPARMSSSSSTSAFPSVSASTAPPQQQQAGMEYRGPHEGGVYSDLMETFLE

>Zm09G02380

MDFSAGSYFTSWPVNSASESYSLADGSVESYGGEGIMPPSSYFMTARSDHNLKFSVHEQDSTMLPNDQLTYAGARQTDLLPGETPSRDKLCENLLELQRLQNNSSLPSNLVPPGVLQHNSTPGAFHPQLNTPGLSELPHALSSSIDSNGSEVSAFLADLNAVSSASALCSTFQNASSFMEPVNLEAFSFQGAQSDSVLNKTTHPNGNISVFDSAALASLHDSKEFISGRLPSFASVQETNVAASGFKTQKQEQNAVCNVPIPTFTARNQIAVAAMPGSLIPQKIPSWINENKSEGPVSHPSDVQIQPNSVGNGVGVKPRVRARRGQATDPHSIAERLRREKISDRMKNLQDLVPNSNKADKASMLDEIIDHVKFLQLQVKVLSMSRLGAPGAVLPLLAESQTEGYRGQLLSAPTNAQGLLDTEESEDTFAFEEEVVKLMETSITSAMQYLQNKGLCLMPVALASAISTQKGVSAASIPPEQ

>Zm09G02760

MQPSGRAMAGGEGGGGAQQQTDDFFDQMLSTLPSAWADLGAGAAEDLAAQSHFGDDSSALLASRLRQHQIGGGDVKSSASHQSQVMLQLSDLHRHGGLGGEESGLFTDRSAPAPEEMEGGFKSPNSAGGDHSLFNGFGVHGAATVQPPFGQSGSMSPESLGGTAASGGGAPPAGGAASSAGGGAAPPRQLRQRQRAKRGQATDPHSIAERLRRERIAERMKSLQELVPNANKTDKASMLDEIIDYVRFLQLQVKVLSMSRLGGAAGGMAPLVASMASSEGKSNGSGGGGNTNATTKSGNGGGLRVAEHQVAKMMEEDMGTAMQYLQGKGLCLMPISLASAISSATSSASLLSRPPPAAAAGGGGGQMHGANGGAAAAATTISSPASANSSG

>Zm09G17300

MNLWTDDNASMMEAFMASADLPAYPWGAPAGGGNPPPPQMPPAMAMAPGFNQDTLQQRLQAMIEGSRETWTYAIFWQSSLDAATGASLLGWGDGYYKGCDDDKRRHRPPLTPAAQAEQEHRKRVLRELNSLISGGASAAPAPAPDEAVEEEVTDTEWFFLVSMTQSFLNGSGLPGQALFAGHHTWIAAGLSSAPCDRARQAYNFGLRTMVCFPVGTGVLELGSTDVVFQTAETMAKIRSLFGGGPGGGSWPPVQPQAAPQQQHAAEADQAAETDPSVLWLADAPVVDIKDSYSHPSAAEISVSKPPPPPPPPQIHFENGSTSTLTENPSPSVHAPPAPPAPPQRQQQNQGPFRRELNFSDFASNPSLAAAPPFFKPESGEILSFGVDSNAQRNPSPAPPASLTTAPGSLFSQSQHTATAAANDAKNNNNNNKRSMEATSLASNTNHHPAAAANEGMLSFSSAPTARPSAGTGAPAKSESDHSDLDASVREVESSRVVAPPPEAEKRPRKRGRKPANGREEPLNHVEAERQRREKLNQRFYALRAVVPNVSKMDKASLLGDAISYINELRGKLTSLESDRETLQAQVEALKKERDARPHPHPAAGLGGHDAGGPRCHAVEIDAKILGLEAMIRVQCHKRNHPSARLMTALRELDLDVYHASVSVVKDLMIQQVAVKMASRMYSQDQLSAALYSRLAEPGSVMGR

>Zm10G02310

MAVGDALRRLCEEVGWSYAVFWKAIGAADPVHLVWEDGCCGHVSCSAGSEAPEAGCELGTSVCTLVRKVMTSQIHVVGEGTIGRAAFTGNHQWIVHDPANDHNLRPEVSAEMNHQFAAGIQTIAIIPVLPRGVLQLGATSVVMENTNLVLQYKNLCSQLNNRSSMASSSSVKNDLNQKIQSRPLNGLTSIHPADTRSKFFSGPPLTYQQCYGPDGRTVSSSTLANMGSNASMLMVAQRNGQVGKEHIVYAPDMRFRPQNPYCDRRVGSNTQNSVVSSGFISSISASMERHPLMTNNRQLEQGNTGEPSDSRSVLLKSLAYHNPLVHENTNMTLLHGRGQVPGFVNGHGGFDFLPEGTRVVKSNLYGSTTNQILDQICSSTSGMKGHGPTISYKIPQPTQFVGKTESSKRETCQASVALLSDSDIQVSSVLKTAISQENQTSRSDLIGSKKANEVHDPTDAIVQAVKNMDRRRLPDISNERSPPFLMDPAAESDLFDMFGSEFHHLCHNVDNDLTWKAAKPENSNRDVPEFSDHLGSSPAFNSVDDEFPYSGIFSLTDTDQLLDAVVSNANPGGKQISGDSASCKTSVTDIPSTSYCRLKEPKQSESSAPPLLIKNELAVSNFVKHPSFPEKAADGCLSQNNAIQKSQIRLWIESGQNLKCESASASNSKGVDTSSKASRKRSRPGENPKPRPKDRQLIQDRIKELRELVPNGAKCSIDALLEKTIKHMVFLQSVTKHADNLKDSNESKILGGENGPLKDYFEGGATWAFDVGSQSMTCPIIVEDLDRPRQMLVEMLCEDRGIFLEIADFIKGLGLTILRGVMEARKNKIWARFTVEANRDVTRMEIFLSLMRLLEPSYDGGGAGENPNNNIKMPLGIVQYPVIPVSGHLR

>Zm10G16990

MARHLHHHRGQELVAHLRASSPLADLLRSAPSLPGARAAHGCVLKSPVAGETFLLNTLVSTYARLGRLREARRVFDGIPLRNTFSYNALLSAYARLGRPDEARALFEAIPDPDQCSYNAVVAALARHGRGHAGDALRFLAAMHADDFVLNAYSFASALSACAAEKDLRTGEQVHGLVARSPHADDVHIGTALVDMYAKCERPVDARRVFDAMPERNVVSWNSLITCYEQNGPVGEALVLFVEMMATGFFPDEVTLSSVMSACAGLAAEREGRQVHAHMVKRDRLRDDMVLNNALVDMYAKCGRTWEARCIFDSMPSRSVVSETSILAGYAKSANVEDAQVVFSQMVEKNVIAWNVLIAAYAQNGEEEEAIRLFVQLKRDSIWPTHYTYGNVLNACGNIAVLQLGQQAHVHVLKEGFRFDFGPESDVFVGNSLVDMYLKTGSIDDGAKVFERMAARDNVSWNAMIVGYAQNGRAKDALHLFERMLCSNENPDSVTMIGVLSACGHSGLVDEGRRHFHFMTEDHGITPSRDHYTCMVDLLGRAGHLKEAEELIKDMPTEPDSVLWASLLGACRLHKNVELGERTAGRLFELDPENSGPYVLLSNMYAEMGKWADVFRVRRSMKDRGVSKQPGCSWIEIGSKMNVFLARDNRHPCRNEIHSTLRIIQMEMCRTSIDGIDDDLMNYCPEACG

>Zm10G22170

MEAALGALCRAGGWSYAAVWRFHPQDPRLLTLGDCYCEDEAKAMVQKMLNLVEVVGEGILGQALVSGECQWIYDDTCHALNQISHADNGDLHQGYTWWQHQFLSGMKTIAVLPLQLHGLVQFGSTRKVPRSSVFLNQVRDIFDRMKNASSDGPMEYTQNNSLACGQQPILTSSRPVNDMLVHNKVNPLQTEKLLEDIERTESIRSSICSPSNSQRSLNDLTSHGTGNSSMDTNILAMPVNSKSIYDLKRFDNVTELFHRSSNDRIGFQVSSSKEPDSTIAGVMSEYKSLNNIHMTENDSCGQNIGYTPYLYTTTNSSNSGLDELSYSGVALSSSFTAVSGKVKDCLQTESDKFLCKSVSFGSNPCASKFQENRLSPYHALMHEQSLIPDPGESVRLLSQEESFIVQGDSSQFKDTADFTGQGNSTFPELPNRSPEEATAETLENNMKDCSGNNSLLETMMLDLSTNSFVQDWWDNSVLLAENLPNSSEKDPEPAMELANRHSLPTGERGSPFIPVIEQLLGDVAHPPTPTGHSSLEAEATALAGRVSDYQLPQFAFRDCLTVDNAQIPALACSNYTCRTRSAQNGSSEVPSANVSVDDTCSLNTANSNGSQSSNPEGTKVAKRRARAGESTRPRPKDRQLIQDRVKELREIVPNGAKCSIDALLERTIKHMLFLQSVNKYAEKIKQADEPKIIDKGSGVILKDNPVAGTNGGATWAYEVAGQTMVCPIIIEDLSPPGQMLVEMLCEEQGLFLEIADNIRGIGLTVLKGRMELCDGKIWSRFLVEANREVTRMDIFLSLVQLLEQNSLVRSTDQMAKVMNNGVPSFANHQRSPLLVPVGIPERLQ

>Zm10G22550

MALSASRVQQAEELLQRPAERQLMRSQLAAAARSINWSYALFWSISDTQPGVLTWTDGFYNGEVKTRKISNSVELTSDHLVMQRSDQLRELYEALLSGEGDRRAAPARPAGSLSPEDLGDTEWYYVVSMTYAFRPGQGLPGRSFASDEHVWLCNAHLAGSKAFPRALLAKSILCIPVMGGVLELGTTDTVPEAPDLVSRATAAFWEPQCPTYSEEPSSSPSGRANETGEAAADDGTFAFEELDHNNGMDIEAMTAAGGHGQEEELRLREAEALSDDASLEHITKEIEEFYSLCDEMDLQALPLPLEDGWTVDASNFEVPCSSPQPAPPPVDRATANVAADASRAPVYGSRATSFMAWTRSSQQSSCSDDAAPAAVVPAIEEPQRLLKKVVAGGGAWESCGGATGAAQEMSGTGTKNHVMSERKRREKLNEMFLVLKSLLPSIHRVNKASILAETIAYLKELQRRVQELESSREPASRPSETTTRLITRPSRGNNESVRKEVCAGSKRKSPELGRDDVERPPVLIMDAGTSNVTVTVSDKDVLLEVQCRWEELLMTRVFDAIKSLHLDVLSVQASAPDGFMGLKIRAQFAGSGAVVPWMISEALRKAIGKR

>Sm157919

MTGRKRLRPGEAPRPRPKDRQQIQDRVRELRDIVPNATKCSIDALLEKTIRHMKFLQSVTQHGDKWKAGGDVKGERSAENSSGLENGASWAMELDAKGSGVPILVENLKQPRQMLVEMLCEERGLFWEIADNIRGLGLTILKGVMESRNDKIWARFNVEVHAAKEVNRLKVVWNLTQLLRPSGKNTSAPSFVGSEMSDVCSESFGSRFQPQVVASHGITVSPGSAW

>Sm46989

RKRLRPGEAPRPRPKDRQQIQDRVRELRDIVPNATKCSIDALLEKTIRHMKFLQSVTQHGDKWKAGGDVKVNEGRNGASWAMELDAKGSGVPILVENLKQPRQMLVEMLCEERGLFWEIA
